# QTL and genetic analysis controlling fiber quality traits using paternal backcross population in Upland Cotton

**DOI:** 10.1101/554147

**Authors:** Lingling Ma, Ying Su, Yumei Wang, Hushai Nie, Yupeng Cui, Cheng Cheng, Meiyan Wang, Jinping Hua

## Abstract

In present study, F_14_ recombinant inbred line (RIL) population was backcrossed to paternal parent for a paternal backcross (BC/P) population, deriving from one Upland cotton hybrid. Three repetitive BC/P field trials and one BC/M field trial were performed including both two BC populations and the original RIL population. Totally, for fiber quality traits, 24 novel QTLs were detected and 13 QTLs validated previous results. And 19 quantitative trait loci (QTL) in BC/P populations explained 5.01% - 22.09% of phenotype variation (PV). Among the 19 QTLs, three QTLs were detected simultaneously in BC/M population. The present study provided novel alleles of male parent for fiber quality traits with positive genetic effects. Particularly, *qFS-Chr3-1* controlling fiber strength explained 22.09% of PV in BC/P population, which increased 0.48 cN/tex for fiber strength. A total of seven, two, eight, two and six QTLs explained over 10.00% of PV for fiber length, fiber uniformity, fiber strength, fiber elongation and fiber micronaire, respectively. In the RIL population, six common QTLs detected in more than one environment such as *qFL-Chr1-2*, *qFS-Chr5-1*, *qFS-Chr9-1*, *qFS-Chr21-1*, *qFM-Chr9-1* and *qFM-Chr9-2*. Two common QTLs of *qFE-Chr2-2* (TMB2386-SWU12343) and *qFM-Chr9-1* (NAU2873-CGR6771) explained 22.42% and 21.91% of PV. In addition, a total of 142 and 46 epistatic QTLs and QTL × environments (E-QTLs and QQEs) were identified in RIL-P and BC/P populations, respectively.

## INTRODUCTION

Upland cotton (*Gossypium hirsutum* L.) is one of the most important sources of natural textile fiber. Among four cultivated species, Upland cotton shows higher yield potential and stronger adaptation to diverse environments than Sea island cotton (*G. barbadence*), *G. arboreum* and *G. herbaceum*, and accounts for more than 92% of cultivated cotton worldwide (Zhang *et al.* 2015a). But fiber quality of Upland cotton is not as good as that of Sea Island cotton. To meet the diverse demands of textile industry, it is a key target to improve fiber quality in breeding project of Upland cotton.

Generally, fiber quality traits consisted of fiber length (FL), fiber uniformity (FU), fiber strength (FS), fiber elongation (FE), and fiber micronaire (FM). Different traits have different genetic mechanisms. Among 4892 QTLs in Cotton QTLdb database (Yu *et al.* 2014), 494, 289, 470, 287 and 395 were detected for FL, FU, FS, FE and FM, respectively (http://www2.cottonqtldb.org:8081/, CottonQTLdb, newly released V2.3 on January 24, 2018).

A total of 151, 132, 91, 118 and 234 QTLs were meta-analyzed for QTL-rich regions for FL, FU, FS, FE and FM, respectively (Said *et al.* 2013). A number of QTLs were located on Chr 5, Chr 19 and Chr 21 (Said *et al.* 2013, 2015). For fiber quality, the most QTLs were detected based on mapping in recombinant inbred lines (RIL) populations of Upland cotton (Wu *et al.* 2009; Sun *et al.* 2012; Ning *et al.* 2014; Tan *et al.* 2014; Shang *et al.* 2015; Tang *et al.* 2015; Zhang *et al.* 2015b; Jamshed *et al.* 2016; Li *et al.* 2016). However, RIL population can be only used to dissect additive and additive × additive effects and not to dissect dominance and dominance-related genetic effects because lacking of heterozygous genotypes. Recently, RIL populations as permanent mapping populations, were used to develop backcross populations in rice (Mei *et al.* 2005) and cotton (Shang *et al.* 2016d), which allows performing repetitive trials as doing in ‘immortalized’ F_2_ population (Hua *et al.* 2002, 2003). Seven QTLs controlled fiber length and fiber strength by using backcross population deriving from Guazuncho 2 × VH8-4602 (Lacape *et al.* 2005). And 44 QTLs for fiber quality traits were detected on Chr 1, Chr 9 and Chr 21 using (CCRI 8 × Pima 90-53) × CCRI 8 BC_1_F_1_ interspecific population (Yang *et al.* 2015). Shang *et al.* (2016d) detected 17, 6, 15, 11 and 21 QTLs for FL, FU, FS, FE and FM, respectively, in F_9_ RIL and F_9_BC_1_ progenies of a hybrid ‘Xinza 1’, While Wang *et al.* (2016) detected 22, 14, 17, 3 and 20 QTLs for the five traits in another two parental F_8_BC_1_ populations. In Wang’s work (2016), two markers of NAU5530 and CIR099 flanking *qFL-c19-2*, *qFU-LG3-1* and *qFS-LG3* were same in Shang’s work (2016d). Using a map of single nucleotide polymorphism (SNP) markers, one fiber length hotspot on Chr 5 carrying three QTLs was observed (Li *et al.* 2016). Additionally, four potential candidate genes for fiber length on Chr Dt7 were found using genotyping by sequencing by genome-wide association studies (GWAS) (Su *et al.* 2016). Previous studies indicated that the RIL population and its two BCF_1_ populations which derived from the same parents increased the power of QTL detection in cotton (Shang *et al.* 2016d; Wang *et al.* 2016). Therefore, it is also available to dissect QTLs for fiber quality using multiple populations at the same time on Upland cotton. Cotton genomes for diploid species (Paterson *et al.* 2012; Wang *et al.* 2012; Li *et al.* 2014; Du *et al.* 2018) and tetraploid genomes (Zhang *et al.* 2015; Li *et al.* 2015c; Yuan *et al.* 2015; Liu *et al.* 2015) had been released recently, so as recent genomics researches in cotton (Fang *et al.* 2017a; Wang *et al.* 2018). These genomics analysis in cotton facilitate applications of SNP markers (Ali *et al.* 2018) and GWAS for fiber quality (Fang *et al.* 2017b; Ma *et al.* 2018b). It is very important to detect novel QTLs and to validate the reported QTLs using diverse populations. In our previous study, serial genetic analyses were performed in multiple segregating populations including F_2_, F_2: 3_, RIL and BC/M population derived from the hybrid ‘Xinza 1’ across multiple years and various locations (Liang *et al.* 2013; Liang *et al.* 2015; Shang *et al.* 2015, 2016a,b,c,d; Ma *et al.* 2017, 2018a, 2019). In previous study, 111 quantitative trait loci (QTLs) were detected for fiber quality using four populations derived from RIL (XZ) and backcross (XZV) hybrids (Shang *et al.* 2016d). Recently, a total of 55 QTLs were detected which distributed in 21 chromosomes using BC/M population in three locations (Ma *et al.* 2017). In addition, 32 QTLs at five stages and 24 conditional QTLs at four intervals were detected for plant height in different years or populations or stages (Ma *et al.* 2018a). And 26 and 27 QTLs including heterotic loci were identified in TC/P and TC/M populations, respectively (Ma *et al.* 2019). In addition, 10 and 16 clusters improved more than one trait for fiber quality and yield and yield-components, respectively (Ma *et al.* 2017; Ma *et al.* 2019). In order to identify and validate genetic components related fiber quality traits, we further developed BCF_1_ progenies population based on RIL population by backcrossing with the paternal parent of ‘Xinza 1’. Here we termed as paternal backcross population (BC/P population for short). We generated additional 177 BCF_1_ crosses for BC/P populations by backcrossing the 177 RI lines as current female parents to GX100-2 (the original male parent), respectively. Detection of new and novel QTLs and comparison analysis were performed for fiber quality traits using BC/P, BC/M and RIL populations together.

## MATERIALS AND METHODS

### Plant materials and populations development

The intraspecific F_14_ recombinant inbred lines (RIL) were inbred for 177 individuals, which were derived from an Upland cotton hybrid “Xinza 1” (GX 1135 × GX 100-2) by single seed descent method (Shang *et al.* 2016a). The parental backcross (BC/P) population was obtained by backcrossing the original male parent (GX100-2) to 177 RI lines, respectively. The maternal backcross (BC/M) population referred to the previous study (Ma *et al.* 2017; Ma *et al.* 2019). The control set was performed in each experimental trial, including GX100-2, “Xinza 1”, GX1135 and a competition hybrid “Ruiza 816” in Yellow River Region.

In present study, we named the maternal and paternal backcross populations as BC/M and BC/P populations for short, respectively, and similarly referred RIL-M population and RIL-P population as RIL population used in BC/M field trials and in BC/P field trials, respectively.

### Field arrangement, sampling and trait evaluation

Three BC/P field trials and one maternal BC trial were conducted in 2015 and 2016 in two locations in China, E1: Quzhou Experimental Station in Handan City, and E2: Guoxin Seed Company Ltd in Cangzhou City (Ma *et al.* 2017, 2018a, 2019). The field trials were designed and planted same to the previous study for BC field trials (Shang *et al.* 2016a; Ma *et al.* 2017, 2018a, 2019). Field management followed the local conventional standard field practice.

Twenty-five naturally opening bolls in the middle of plants were hand-harvested for each plot at mature stage in three environments, respectively. Fiber samples were ginned and sampled for measurements of fiber quality traits with HVI 900 instrument (USTER_ HVISPECTRUM, SPINLAB, USA) at Cotton Fiber Quality Inspection and Test Center of Ministry of Agriculture (Anyang, China) (Shang *et al.* 2016d). The fiber quality traits included 2.5% fiber span length (FL, mm), fiber uniformity (FU, %), fiber strength (FS, cN/tex), fiber elongation (FE), and fiber micronaire (FM) as usual.

### Genetic Map and Data Analysis

The genetic map based on RIL population published before (Shang *et al.* 2016c), in which a total of 653 loci based on SSR markers distributed on 31 linkage groups and anchored on 26 chromosomes, covering 3889.9 cM (88.20%) of cotton genome with average interval of 6.2 cM (Ma *et al.* 2017, 2018a, 2019). The genotype for each maternal F_14_BC_1_ was deduced on the basis of the RIL genotype used as the parent for backcross (Shang *et al.* 2016a, b, c).

Basic statistical analysis was implemented by the software SPSS (Version 19.0, SPSS, Chicago). Using the variance analysis, heritability was calculated in the equation as *h^2^* = *δ^2^_G_ /* [*δ^2^_G_ +*(*δ^2^_G_* × E) */env*], where *δ^2^* and *δ^2^* × refers to the genotypic variance and genotype-by-environment interaction variance, respectively, and *env* to the number of the environments.

Composite interval mapping (CIM) method was used for QTL mapping in the confidence interval of 95%. The software QTL Cartographer (Version 2.5) (Zeng *et al.* 1994; Wang *et al.* 2007) was used to map single-locus QTL and to estimate the genetic effect. The threshold of LOD was estimated to declare a suggestive QTL after 1000 permutation times, whereas QTL in another environment or population with LOD of at least 2.0 was considered as common QTL (Liang *et al.* 2013; Shang *et al.* 2015, 2016d). According to the position linked and shared common markers, QTLs detected in different populations were regarded as common QTL (Shao *et al.* 2014; Shang *et al.* 2016d; Ma *et al.* 2017).

The QTL IciMapping 4.1 (www.isbreeding.net) was conducted by the two-locus analysis using inclusive composite interval mapping (ICIM) method (Shang *et al.* 2016d; Ma *et al.* 2017).. The main-effect QTL (M-QTL) and its environmental interaction (QTL × environment, QE), epistatic QTLs (E-QTLs) and its environmental interactions (QTLs × environment, QQE) were conducted using RIL-P and BC/P datasets under multiple environments in three parental TC trials. A threshold LOD 2.5 and 5 scores were used to declare significant M-QTL and E-QTLs, respectively.

### Data availability

All of our raw data are available as Supporting Information Table S6 for phenotype values of fiber quality traits of BC/M population in three field trials in this study, Table S7 for phenotype values of fiber quality traits in field trial of BC/P population in this study, and Supporting Information Table S8A for genotypes of RIL population (Shang *et al.* 2016a), Table S8B for genotypes of BC/M population (Ma *et al.* 2017), and Table S8C for genotypes of BC/P population.

## RESULTS

### Trait Performance in Two Populations

The ‘original’ maternal parent ‘GX1135’ of Xinza 1 performed differently with the ‘original’ male parent ‘GX100-2’ for five fiber quality traits (**Table 1**). The hybrid *‘*Xinza 1*’*showed no significant hybrid vigor of F_1_ for fiber quality traits ranging from −3.17% to 1.53% of mid-parent heterosis (MPH). Among different traits, larger phenotypic variation was observed for fiber length (FL), fiber strength (FS) and fiber micronaire (FM), ranging from 2.63% to 8.53% in both BC and RIL populations; And lighter phenotypic variation presented with 0.87% - 1.23% for fiber uniformity (FU) and fiber elongation (FE) in BC and RIL populations.

**Table 1.** Descriptive statistical analysis of fiber quality traits in multiple populations and the control set

| Trait | Env. | BC/P |  |  | RIL-P |  |  | GX1135 | GX100-2 | Xinza 1 | Ruiza 816 |
| --- | --- | --- | --- | --- | --- | --- | --- | --- | --- | --- | --- |
|  |  | Mean | Range | CV% | Mean | Range | CV% |  |  |  |  |
| FL/mm | 2016E1 | 30.38±0.80 | 27.70-32.95 | 2.63 | 30.36±1.07 | 26.35-33.75 | 3.54 | 31.15 | 30.40 | 30.73 | 31.15 |
|  | 2015E2 | 31.08±0.69 | 29.20-33.00 | 2.21 | 30.95±0.95 | 28.60-33.20 | 3.07 | 30.44 | 30.10 | 31.05 | 31.15 |
|  | 2016E2 | 29.42±0.78 | 26.65-31.45 | 2.64 | 29.15±1.08 | 26.00-31.60 | 3.70 | 28.92 | 28.80 | 30.35 | 30.35 |
|  | <u>2016E2</u> | <u>29.26±0.95</u> | <u>26.50-31.70</u> | <u>3.25</u> | <u>29.14±1.2</u> | <u>26.05-32.75</u> | <u>4.12</u> | 29.28 | 29.38 | 28.90 | 29.80 |
| FU | 2016E1 | 86.28±0.76 | 83.75-88.30 | 0.88 | 85.99±0.85 | 82.85-88.05 | 0.98 | 86.43 | 85.43 | 85.78 | 86.43 |
|  | 2015E2 | 86.06±0.70 | 83.70-87.90 | 0.81 | 85.79±0.82 | 82.70-88.35 | 0.96 | 85.28 | 86.50 | 86.13 | 86.08 |
|  | 2016E2 | 85.29±1.02 | 81.75-87.40 | 1.20 | 84.95±1.05 | 80.85-87.05 | 1.23 | 84.60 | 85.35 | 85.45 | 86.00 |
|  | <u>2016E2</u> | <u>85.14±0.98</u> | <u>82.25-87.40</u> | <u>1.15</u> | <u>84.90±1.21</u> | <u>80.2-87.45</u> | <u>1.42</u> | 85.43 | 86.05 | 85.05 | 86.20 |
| FS/(cN/tex) | 2016E1 | 28.81±0.94 | 26.05-30.85 | 3.25 | 28.99±1.25 | 24.65-32.30 | 4.32 | 31.85 | 30.13 | 30.23 | 31.85 |
|  | 2015E2 | 30.63±0.99 | 28.15-33.60 | 3.22 | 30.80±1.42 | 27.50-34.30 | 4.62 | 29.20 | 28.57 | 28.15 | 30.85 |
|  | 2016E2 | 29.11±1.02 | 26.45-32.45 | 3.50 | 29.03±1.38 | 25.6-33.90 | 4.75 | 28.50 | 28.70 | 30.18 | 31.08 |
|  | <u>2016E2</u> | <u>29.39±1.08</u> | <u>26.75-31.85</u> | <u>3.68</u> | <u>29.12±1.41</u> | <u>25.5-32.75</u> | <u>4.83</u> | 29.20 | 28.35 | 29.28 | 31.10 |
| FE | 2016E1 | 6.81±0.06 | 6.55-7.00 | 0.87 | 6.80±0.07 | 6.55-6.95 | 1.09 | 6.98 | 6.90 | 6.90 | 6.98 |
|  | 2015E2 | 6.99±0.06 | 6.85-7.20 | 0.84 | 6.99±0.08 | 6.80-7.25 | 1.16 | 6.78 | 6.77 | 6.83 | 6.78 |
|  | 2016E2 | 6.74±0.06 | 6.55-6.90 | 0.88 | 6.72±0.07 | 6.50-7.00 | 1.08 | 6.67 | 6.70 | 6.75 | 6.80 |
|  | <u>2016E2</u> | <u>6.72±0.07</u> | <u>6.55-6.90</u> | <u>0.99</u> | <u>6.71±0.08</u> | <u>6.55-6.90</u> | <u>1.17</u> | 6.73 | 6.70 | 6.70 | 6.75 |
| FM | 2016E1 | 4.81±0.34 | 3.95-5.70 | 7.12 | 4.86±0.41 | 3.80-5.75 | 8.53 | 5.33 | 5.10 | 4.50 | 5.33 |
|  | 2015E2 | 4.76±0.27 | 4.15-5.40 | 5.59 | 4.71±0.36 | 3.65-5.65 | 7.65 | 5.06 | 5.10 | 5.35 | 5.18 |
|  | 2016E2 | 4.93±0.24 | 4.30-5.70 | 4.92 | 4.97±0.37 | 4.10-5.90 | 7.45 | 5.33 | 4.55 | 4.98 | 5.55 |
|  | <u>2016E2</u> | <u>5.18±0.25</u> | <u>4.60-5.80</u> | <u>4.80</u> | <u>5.02±0.36</u> | <u>3.90-5.90</u> | <u>7.04</u> | 5.40 | 5.13 | 5.00 | 5.38 |

Env., two testing locations (E1, Handan, E2, Cangzhou) in 2015 and 2016.
Mean referred to the mean value ± standard deviation. CV%, coefficient of variation.
Data without underline was from the recombinant inbred line (RIL-P) and backcross (BC/P) populations in paternal BC trials; Data with underlines was from the RIL-M and BC/M populations in maternal BC trials.

Genotype variance and environment variance for five traits showed significant variation at level of 0.05 in RIL and BC populations (**Table 2**). Fiber length (FL) and fiber uniformity (FU) increased in BC/P population in comparison with that in BC/M population, whereas fiber micronaire (FM) reduced. Fiber length (FL) and fiber strength (FS) showed larger heritability of 91.82% and 91.10% in RIL-P population, respectively. The heritability decreased to 86.63% and 81.93% for FL and FS in BC/P population, respectively (**Table 2**). The result indicated that wider range of phenotypic variation and bigger heritability in RIL-P population were showed in BC/P population for five fiber quality traits.

**Table 2.**
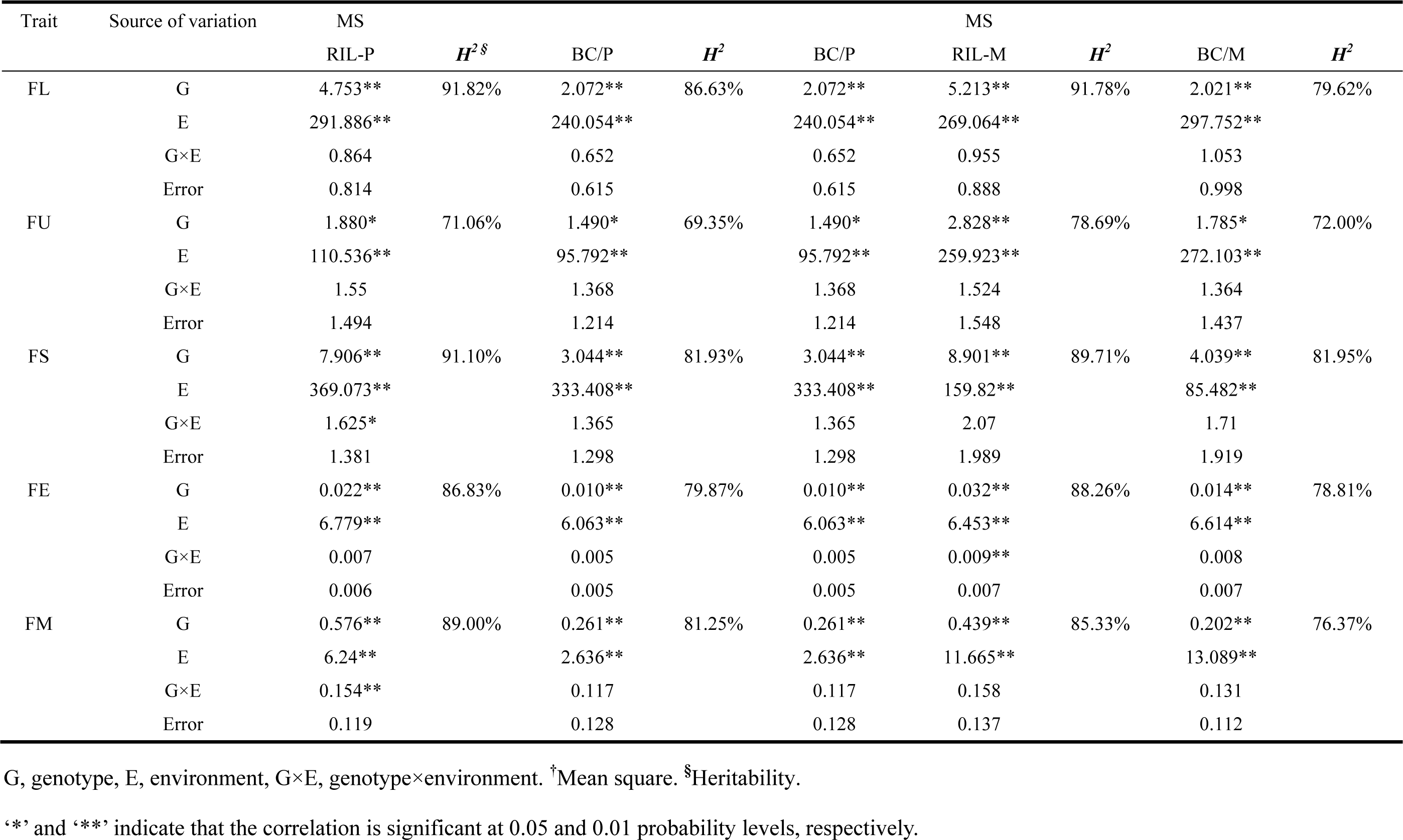
Results of ANOVA and heritability for yield and its components in different populations from two backcross trials

### Correlation Analysis among Fiber Quality Traits in Multiple Populations

The significant correlation coefficients were calculated among five fiber quality traits in BC/P, BC/M, RIL-P and RIL-M populations in 2015E2, 2016E1 and 2016E2 (Table 3). Fiber length (FL) correlated significantly and positively with fiber uniformity (FU), fiber strength (FS) and fiber elongation (FE) in these populations except FU in 2016E1. However, fiber micronaire (FM) showed significant negative correlation with FL and FS in the populations. This result was similar to the previous research (Liang *et al*, 2013; Shang *et al*, 2016d; Ma *et al*, 2019).

**Table 3.** Correlation analyses between five fiber quality traits in RIL population and its TC progenies

| Trait | Env. | FL |  | FU |  | FS |  | FE |  |
| --- | --- | --- | --- | --- | --- | --- | --- | --- | --- |
|  |  | RIL | BC | RIL | BC | RIL | BC | RIL | BC |
| FU | 2016E1 | 0.013 | -0.096 |  |  |  |  |  |  |
|  | 2015E2 | 0.226** | 0.222** |  |  |  |  |  |  |
|  | 2016E2 | 0.497** | 0.381** |  |  |  |  |  |  |
|  | <u>2016E2</u> | <u>0.438**</u> | <u>0.178*</u> |  |  |  |  |  |  |
| FS | 2016E1 | 0.656** | 0.616** | -0.030 | -0.083 |  |  |  |  |
|  | 2015E2 | 0.718** | 0.699** | 0.217** | 0.075 |  |  |  |  |
|  | 2016E2 | 0.754** | 0.581** | 0.403** | 0.060 |  |  |  |  |
|  | <u>2016E2</u> | <u>0.784**</u> | <u>0.525**</u> | <u>0.305**</u> | <u>0.043</u> |  |  |  |  |
| FE | 2016E1 | 0.659** | 0.617** | 0.128 | 0.105 | 0.530** | 0.511** |  |  |
|  | 2015E2 | 0.627** | 0.476** | 0.369** | 0.163* | 0.507** | 0.533** |  |  |
|  | 2016E2 | 0.754** | 0.737** | 0.454** | 0.252** | 0.710** | 0.696** |  |  |
|  | <u>2016E2</u> | <u>0.813**</u> | <u>0.535**</u> | <u>0.414**</u> | <u>0.249**</u> | <u>0.751**</u> | <u>0.486**</u> |  |  |
| FM | 2016E1 | -0.237** | -0.202** | 0.295** | 0.314** | -0.320** | -0.404** | -0.015 | 0.016 |
|  | 2015E2 | -0.439** | -0.334** | 0.088 | 0.049 | -0.428** | -0.390** | -0.102 | 0.047 |
|  | 2016E2 | -0.443** | -0.124 | -0.053 | 0.093 | -0.497** | -0.261** | -0.206** | 0.004 |
|  | <u>2016E2</u> | <u>-0.426**</u> | <u>-0.277**</u> | <u>0.063</u> | <u>0.142</u> | <u>-0.434**</u> | <u>-0.482**</u> | <u>-0.219**</u> | <u>-0.020</u> |

‘\*’, ‘\*\*’ indicate that the correlation is significant at 0.05 and 0.01 probability levels, respectively.
RIL, recombinant inbred line population in maternal and paternal testcross trials, respectively; BC, backcross populations including maternal and paternal BC (BC/M and BC/P) populations.
Figures underlined referred to QTLs identified in one BC/M trial, the remaining QTLs identified in three BC/P trials.

The correlations were similar tendency among BC/P and BC/M populations. In both BC/P and BC/M populations, no significant correlation was showed between FU and FS. However, the majority of correlation values decreased in both BC/P and BC/M populations in comparison with correlation values in RIL population after backcrossing to either of parents.

### Single Locus QTL Analysis

In four field trials, a total of 37 QTLs controlling fiber quality were detected in three corresponding populations of BC/P, RIL-P, BC/M and RIL-M, explaining 5.01% - 22.42% of PV (phenotypic variance) (**Table 4**, **Figure 1**). These QTLs anchored on 17 chromosomes, respectively.

**Table 4.**
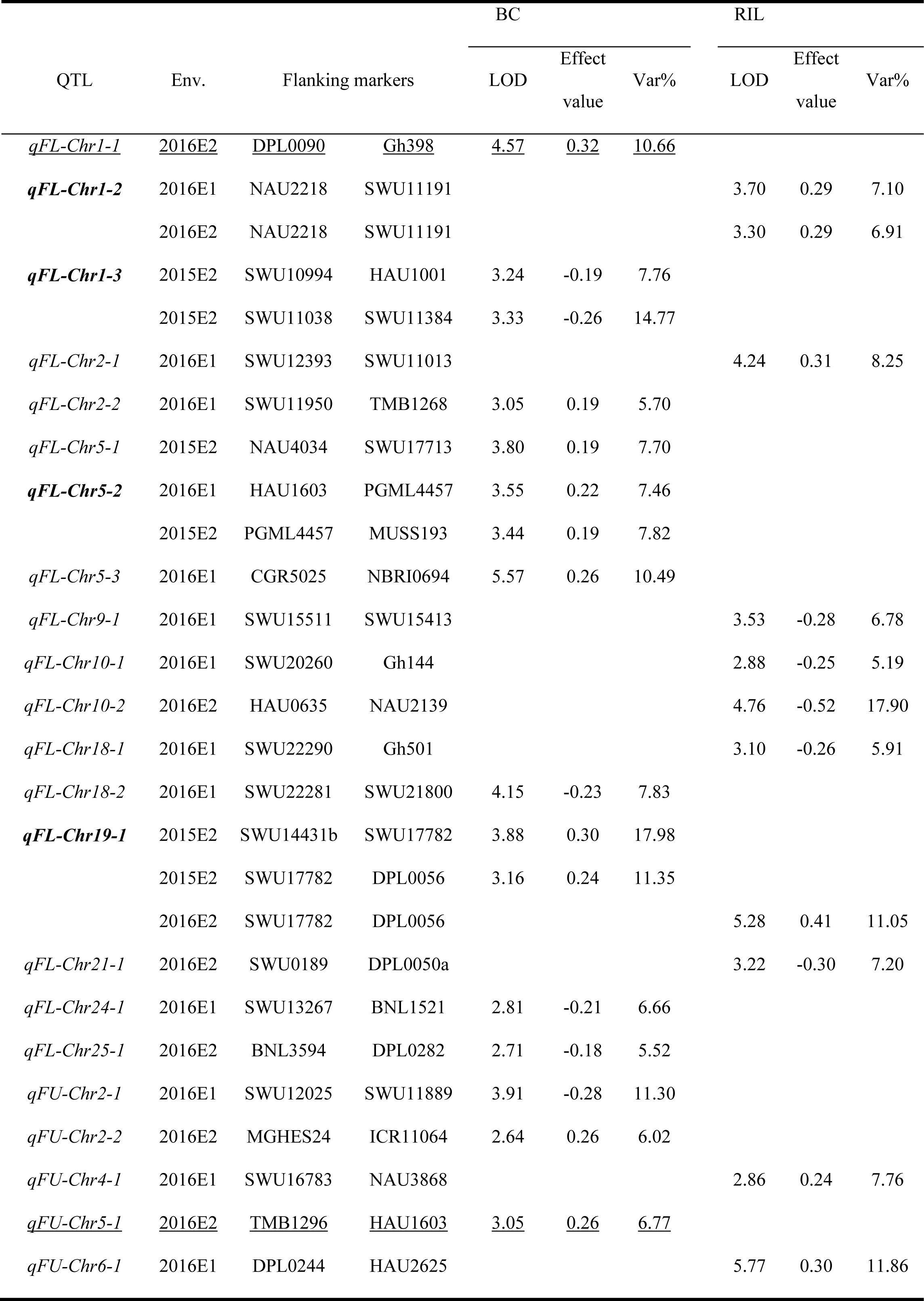
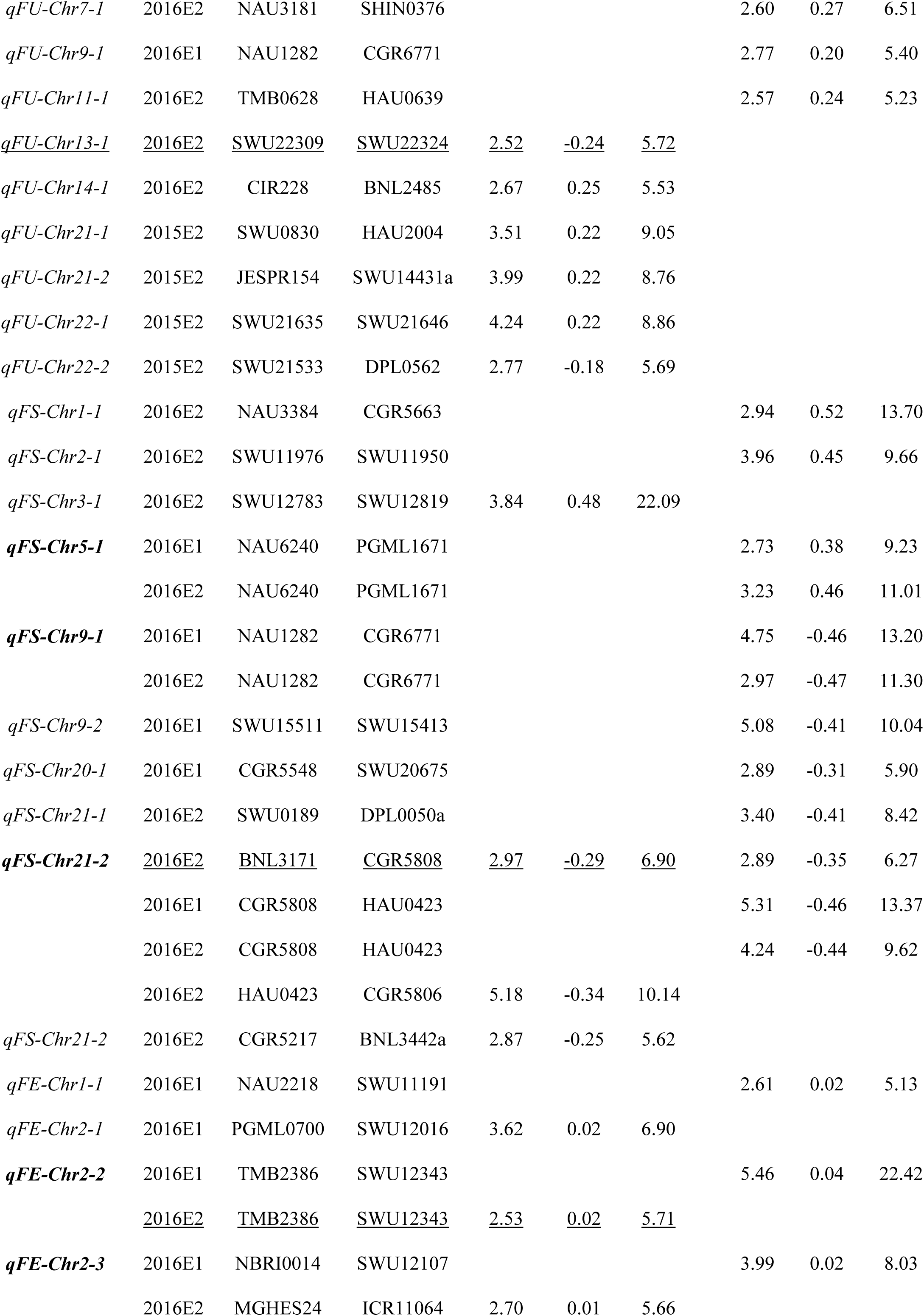
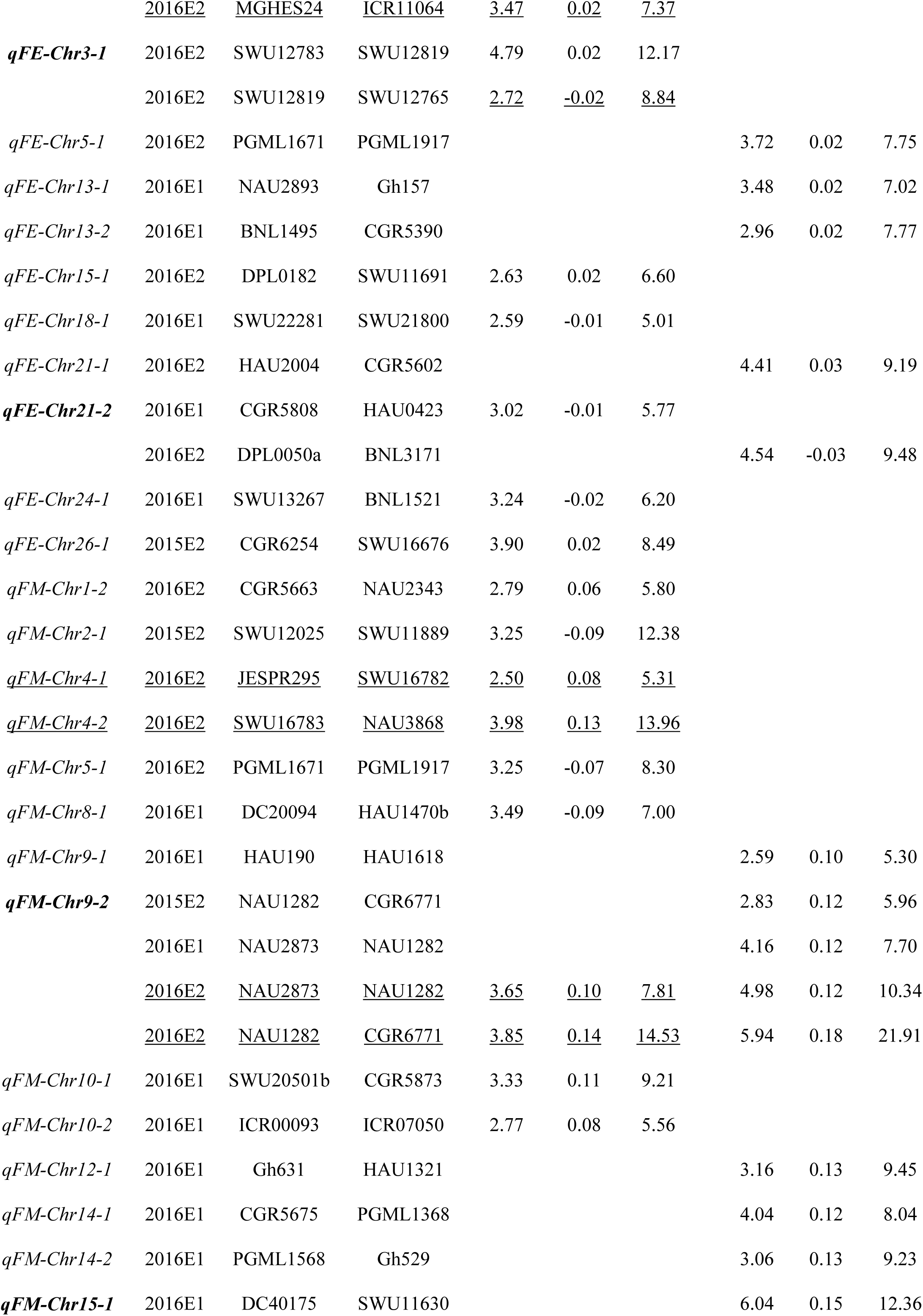
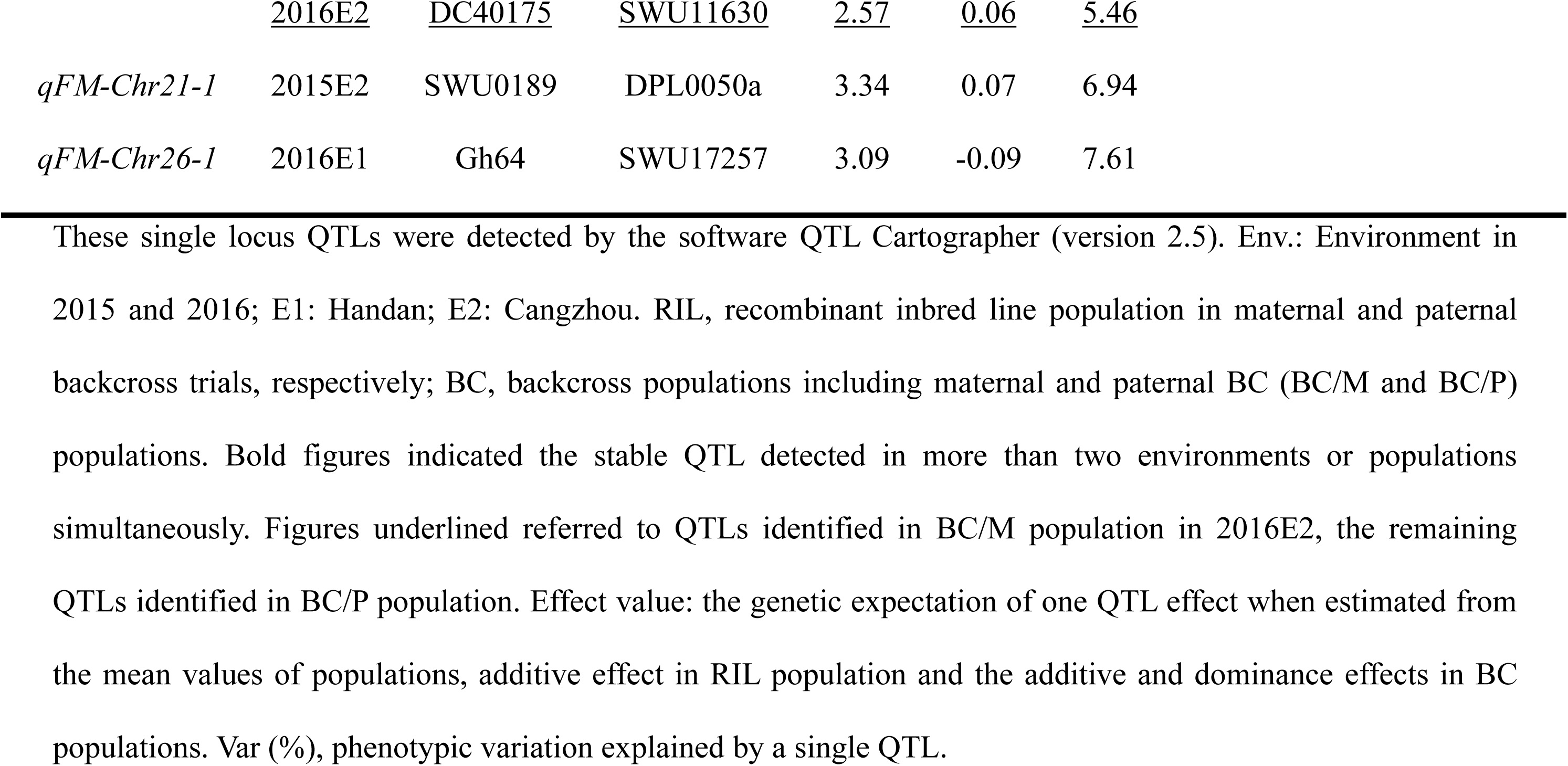
Single-locus QTLs in paternal and maternal backcross experiments by composite interval mapping method

**Table 5.** Summary on M-QTL and E-QTLs controlling fiber quality traits in BC/P and RIL-P datasets in BC/P trials

| Trait |  | BC/P |  | RIL-P |  |  |
| --- | --- | --- | --- | --- | --- | --- |
| M-QTL<sup>a</sup> | n | V(A)% | V(AE)% | n<sup>b</sup> | V(A)%<sup>c</sup> | V(AE)% |
| FL | 11 | 3.13 | 0.70 | 13 | 2.63 | 0.27 |
| FU | 3 | 3.02 | 3.00 | 7 | 3.15 | 2.40 |
| FS | 11 | 2.53 | 0.67 | 14 | 2.92 | 0.25 |
| FE | 0 | - | - | 12 | 2.77 | 0.59 |
| FM | 13 | 2.61 | 0.99 | 10 | 3.41 | 1.01 |
| Mean | - | 2.82 | 1.34 | - | 2.98 | 0.90 |
| E-QTL<sup>a</sup> | n | V(AA)% | V(AAE)% | n | V(AA)%<sup>c</sup> | V(AAE)% |
| FL | 16 | 3.44 | 0.30 | 52 | 4.33 | 0.22 |
| FU | 11 | 1.93 | 2.12 | 0 | - | - |
| FS | 5 | 3.32 | 0.52 | 44 | 4.15 | 0.24 |
| FE | 1 | 0.76 | 0.01 | 18 | 4.42 | 0.24 |
| FM | 13 | 3.44 | 0.73 | 28 | 3.77 | 0.23 |
| Mean | - | 2.58 | 0.74 | - | 4.17 | 0.23 |

<sup>a</sup> M-QTLs and E-QTLs refer to the main effect QTL and epistasis QTLs by environments.
<sup>b</sup> The number of QTLs.
<sup>c</sup> V(A)%, V(AAE)%, V(AA)% and V(AAE)%, the total proportion of phenotypic variation on average explained by single QTL, epistatic QTLs (AA) and them by environments (AE or AAE) at the current scanning position, respectively.

**Figure 1.**
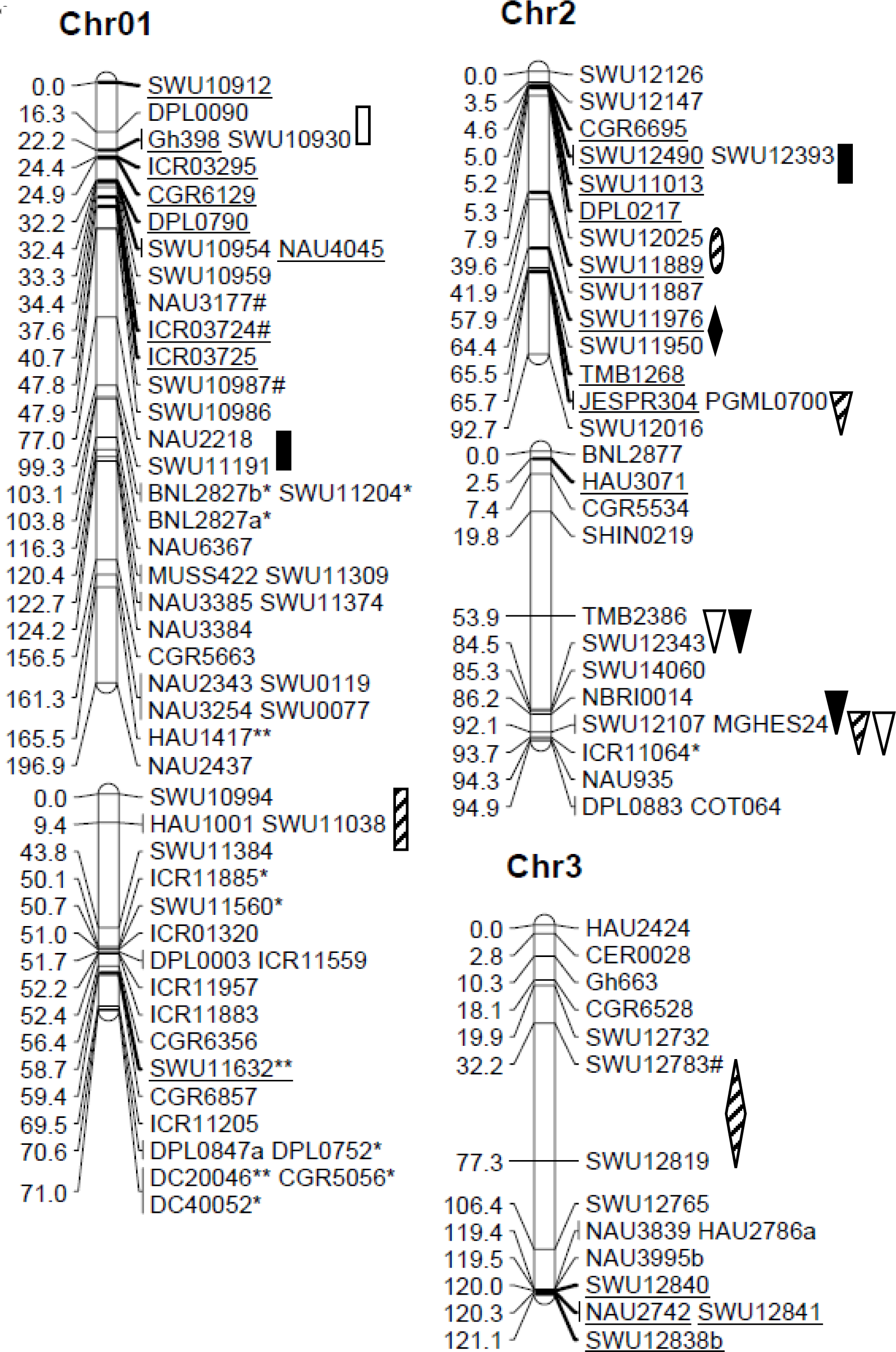
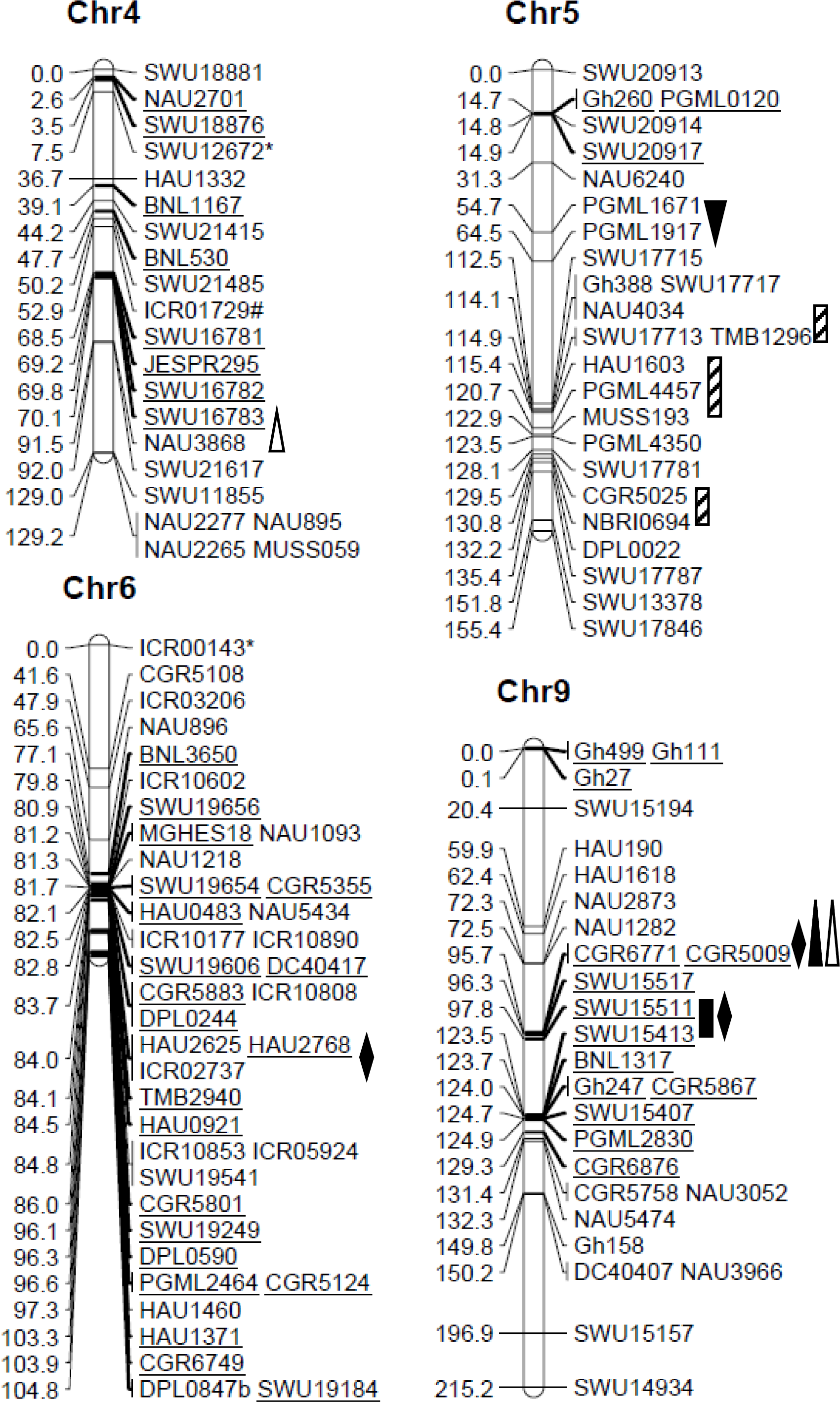
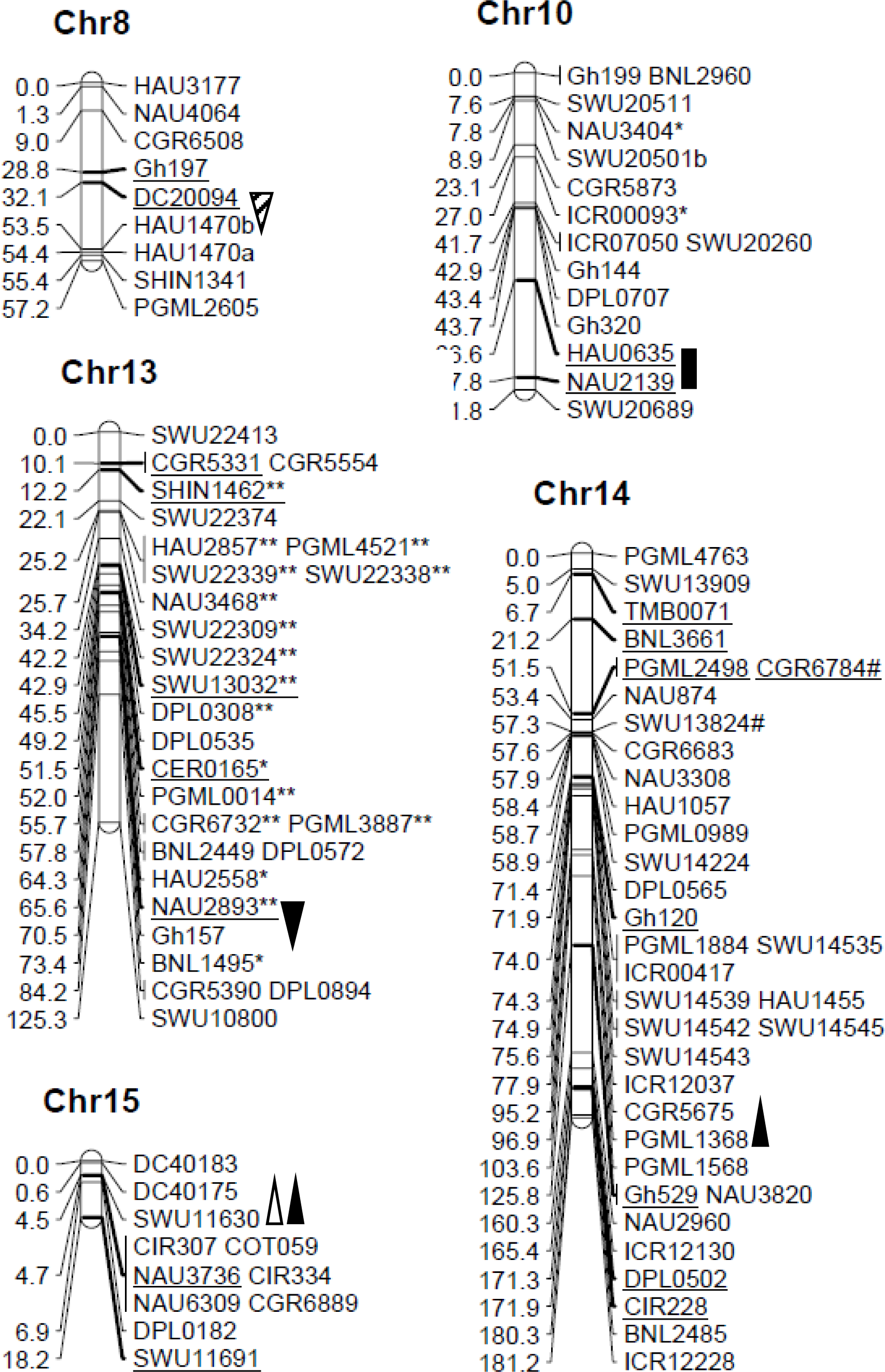
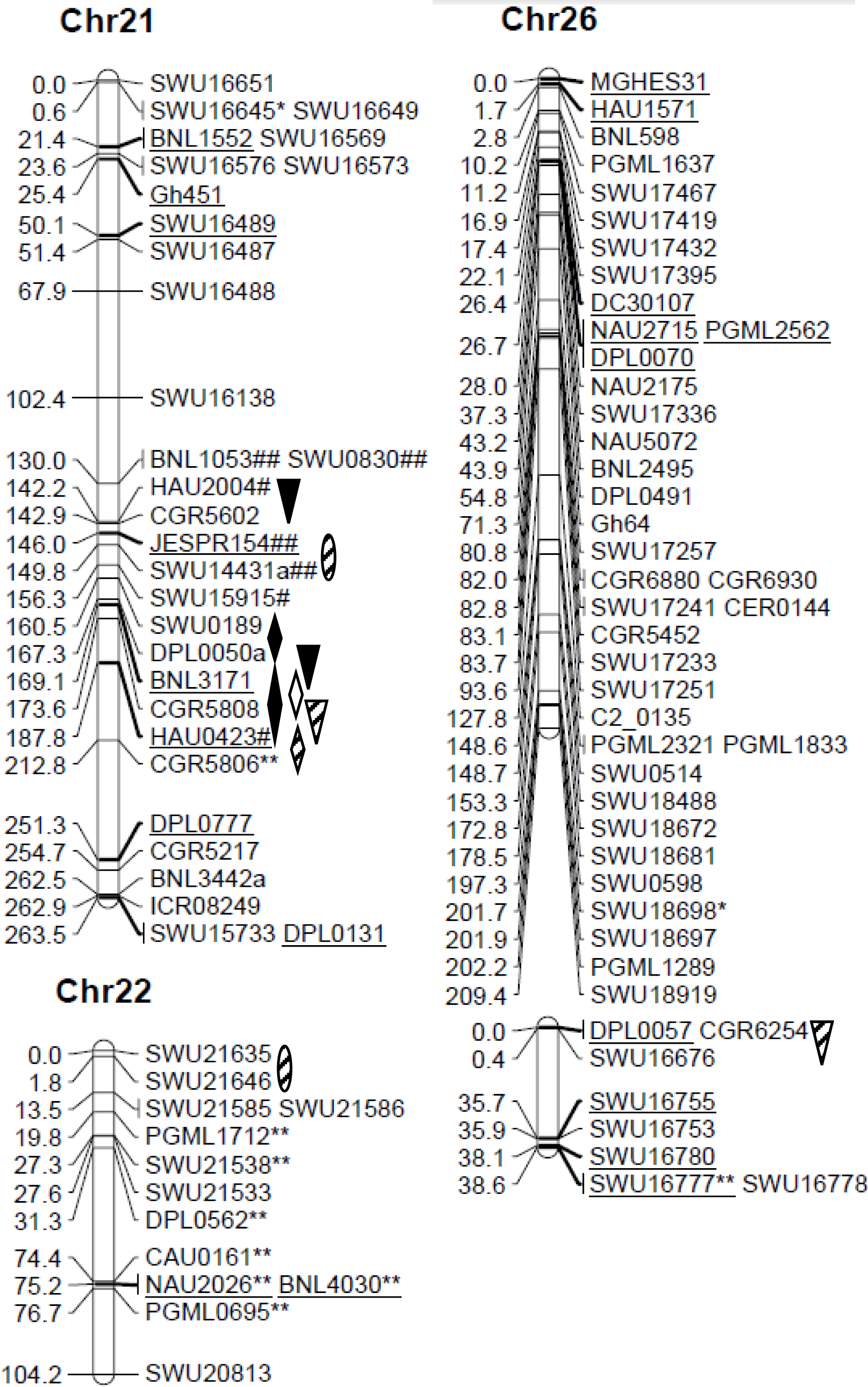
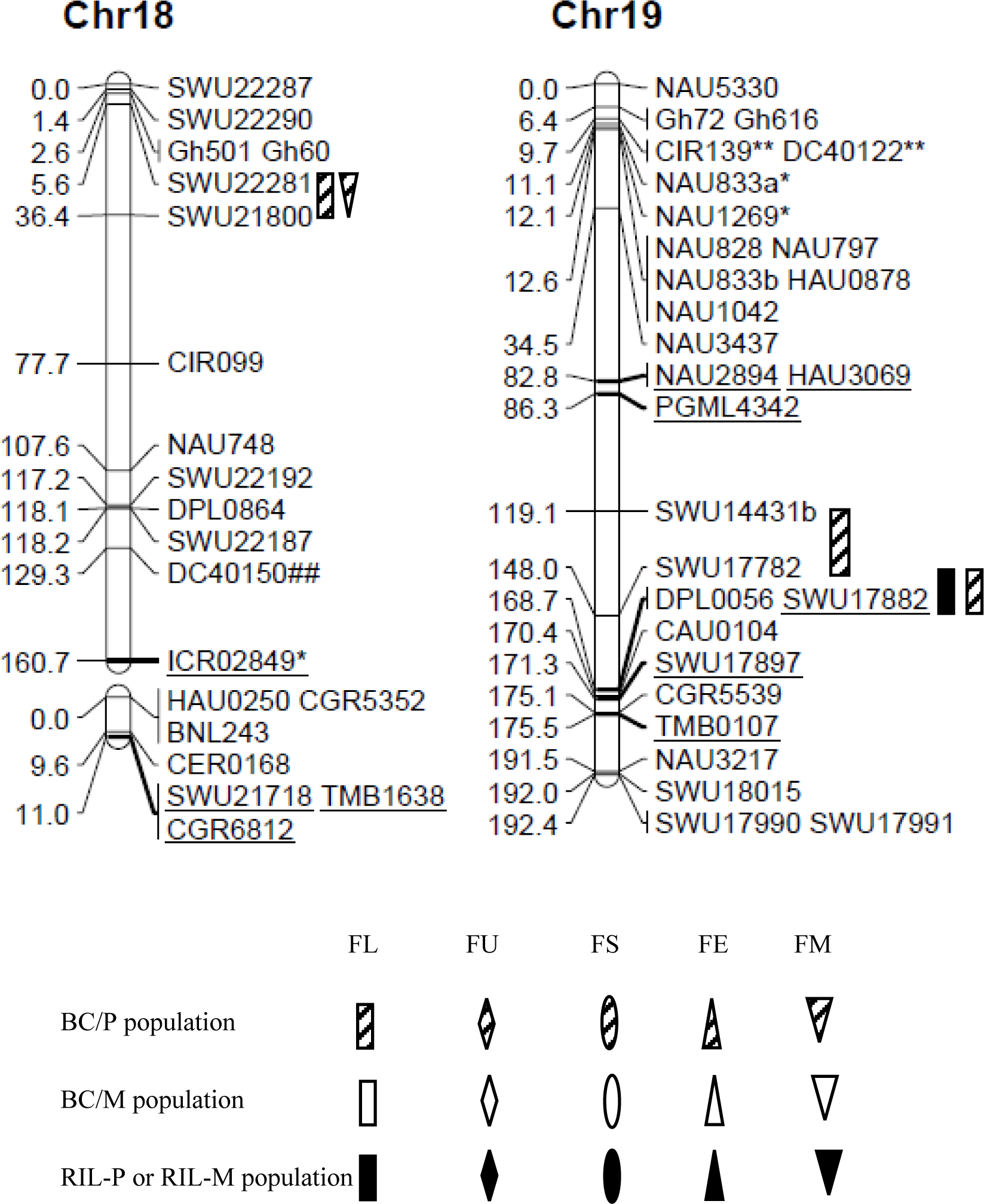
Locations of QTLs controlling fiber quality traits in BC/P and BC/M trials * and ** (# and ##), marker showed respectively segregation distortion significant at P = 0.05 and 0.01 levels; markers with * and ** skewed toward the GX1135 alleles, and markers with # and ## skewed toward the GX100-2 alleles.

For fiber length, six, one and five QTLs were identified in BC/P, BC/M and RIL populations, respectively. The *qFL-Chr5-2* was simultaneously identified in 2015E2 and 2016E1, explaining 7.46% and 7.82% of PV, respectively. The *qFL-Chr1-2* was simultaneously detected in 2016E1 and 2016E2, explaining 7.01% and 6.91% of PV, respectively. The *qFL-Chr19-1* explaining 17.98% of PV in BC/P population was verified in RIL population with 11.05% of PV. Among five QTLs detected in RIL population, three QTLs showed additive effects originated from GX1135 alleles whereas three QTLs showed additive effects offered by GX100-2 alleles. Three QTLs *qFL-Chr5-1*, *qFL-Chr5-2* and *qFL-Chr5-3* were distributed on chromosome 5 (Chr 5), and three QTLs *qFL-Chr1-1*, *qFL-Chr1-2* and *qFL-Chr1-3* were distributed on Chr 1.

A total of five QTLs were detected for fiber uniformity (FU) explaining 8.76% - 11.86% of PV, which distributed on four chromosomes. Four QTLs and one were identified in BC/P and RIL populations, respectively. No common QTL was identified in multiple populations or multiple environments. The *qFU-Chr6-1* increased FU providing alleles by GX1135 in RIL population.

For fiber strength (FS), a total of six QTLs were detected on four chromosomes, explaining 6.27% - 22.09% of PV. Two common QTLs were identified. The *qFS-Chr3-1* was detected in BC/P population alone explaining high to 22.09% of PV in 2016E2. The *qFS-Chr3-1* increased 0.48 cN/tex fiber strength in 2016E2. The *qFS-Chr9-1* was simultaneously detected in RIL population in 2016E1 and 2016E2, providing alleles by female parent GX100-2 in both environments. The QTL explained 9.23% of PV in 2016E2 and 11.01% of PV in 2016E1. All of four QTLs provided increasing effect alleles donated by female parent GX100-2 in both environments in RIL population. However, *qFS-Chr9-1* increased 0.46 cN/tex and 0.47 cN/tex fiber strength in 2016E1 and 2016E2. The *qFS-Chr21-2* was detected in BC/P, BC/M and RIL populations at the same time across 2016E1 and 2016E2, explaining 6.27% - 13.37% of PV. Four QTLs distributed on A-subgenome of Chr 2, Chr 3, Chr 9 whereas the two remaining QTLs distributed on D-subgenome of Chr 21. Six, two and six QTLs for fiber elongation (FE) were identified in the BC/P, BC/M and RIL populations, respectively. Four common QTLs were detected in at least two populations, including *qFE-chr2-2*, *qFE-chr2-3*, *qFE-chr3-1*, and *qFE-chr21-2*. The *qFE-chr2-3* detected in BC/P population was verified in BC/M and RIL populations, which explained 7.02% of PV on average. The *qFE-chr3-1* explaining 12.17% of PV in BC/P population were also detected in BC/M population in the same environment of 2016E2. The *qFE-Chr2-2* explained 22.42% of PV in RIL population in 2016E1 and was observed in BC/M population in 2016E2. The *qFE-Chr21-2* detected in BC/P population was also identified in RIL population.

A total of 5 QTLs underlying fiber micronaire were located on 5 chromosomes. The *qFM-Chr9-2* was identified in BC/M and RIL populations in 2016E2 explaining 14.53% and 21.91% of PV, respectively. At the same time, the QTL were detected in three environments of 2015E2, 2016E1 and 2016E2. Another common QTL *qFM-Chr15-1* was simultaneously detected in RIL population in 2016E1 and in BC/M population in 2016E2, explaining 12.36% and 5.46% of PV, respectively. All of six QTLs detected in RIL population increased over 0.10 fiber micronaire value, which donating increasing additive effect alleles by GX1135, containing *qFM-Chr9-1*, *qFM-Chr9-2*, *qFM-Chr12-1*, *qFM-Chr14-1, qFM-Chr14-2*, and *qFM-chr15-1*. On summary, 19 QTL explained 10.14-22.09% of PV on average in BC/P population in 2015E2, 2016E1 and 2016E2. Among them, a total of eight QTLs explained larger than 10% of PV. Then, we identified 8 QTLs in BC/M population, explained 9.03% of PV on average. At last, 20 QTLs existed in RIL-P and RIL-M populations explaining on average 10.39% of PV in four environments above. Totally, 12 common QTLs were detected in multiple environments or in multiple populations of BC/P, BC/M, RIL-P and RIL-M populations (Table 4, S5).

### Pleiotropic Effects

We also observed 5 pleiotropic regions controlling at least two fiber quality traits on 2 chromosomes of Chr 9, Chr18 and Chr 21 (Figure 1). Of these, a pleiotropic region flanking with SWU15511- SWU15413 on Chr 9 increased the values for FL and FS, showing increasing additive effects originated from alleles of GX100-2. The NAU2873-CGR6771 on Chr 9 contributed alleles to FS but also increased the FM. The region of SWU0830-HAU2004-CGR5602 contained *qFU-Chr21-1* in BC/P population and *qFE-Chr21-1* in RIL-P population. And SWU0189-CGR5808 on Chr 21 flanked along *qFS-Chr21-1*, *qFS-Chr21-2 and qFE-Chr21-2*, all of which showing increasing additive effects originated from alleles of GX100-2. The region of SWU15511- SWU15413 on Chr 18 controlled fiber length and fiber elongation at the same time.

### Digenic and Environmental Interactions in Three BC/P Trials

In the three repetitive BC/P trials, a total of 38 and 56 M-QTLs and environmental interactions (QTL × environment, QE) were identified in BC/P and RIL-P populations, respectively (**Table 5**, **Table S1, S2**). Of these, they explained 2.53% - 3.13% and 2.63% - 3.41% of PV on average in the populations, respectively. Environmental effect prevailed in both BC/P and RIL-P populations. However, environment and M-QTL interacted with 1.24% of PV in the BC/P population while with 0.90% of PV in RIL-P population.

Then, 46 and 142 E-QTLs and their environmental interactions (digenic interactions × environment, QQE) were respectively identified in BC/P and RIL-P populations, respectively (**Table S3, S4**). Eighteen pairs of E-QTL and QQE explained 4.17% of PV on average in RIL-P population, while nine pairs of E-QTL and QQE explained 2.58% of PV on average in BC/P populations (**Table 5**). On average, the number of both types of M-QTL and E-QTL was larger in RIL populations than that in BC populations. To our surprise, epistatic interactions contributed more to fiber quality than M-QTLs did in RIL-P population.

Totally, 20 (42.5%) and 101 (71.13%) pairs of E-QTLs and QQE contained M-QTLs and QEs in BC/P and RIL-P populations, respectively (**Table 6**). We detected about 3-fold epistatic QTLs in RIL populations than QTLs in BC/P population, and 19.01% M-QTLs participated epistasis between M-QTL and M-QTL. Three types of epistasis were checked: I) both loci were M-QTLs; II) either locus between two loci was M-QTL; III) both loci were no M-QTLs (Shang *et al.* 2016d; Ma *et al.* 2019). Apparently, 27 (57.45%) epistatic QTLs of type III was the most popular type in epistatic styles in BC/P population whereas it was 31 (52.11%) epistatic QTLs of type II in RIL-P population (Table 6). The results indicated that epistasis played more vital role in improving fiber quality both in RIL populations of Upland cotton. The result was consistent to the previous result that epistatic QTLs with significant additive × additive effects were identified for fiber quality traits (Wang *et al.* 2017; Ma *et al*, 2019).

**Table 6.** Types of epistasis detected for fiber quality traits in the RIL-P and BC/P populations

| Trait | Type of epistasis<sup>a</sup> |  |  |  |  |  |  |  |
| --- | --- | --- | --- | --- | --- | --- | --- | --- |
|  | RIL-P |  |  |  | BC/P |  |  |  |
|  | I | II | III | Total | I | II | III | Total |
| Fiber length | 8 | 29 | 15 | 52 | 2 | 5 | 9 | 16 |
| Fiber uniformity | 0 | 0 | 0 | 0 | 2 | 5 | 4 | 11 |
| Fiber strength | 10 | 22 | 12 | 44 | 0 | 1 | 5 | 6 |
| Fiber elongation | 4 | 6 | 8 | 18 | 0 | 0 | 1 | 1 |
| Micronaire | 5 | 17 | 6 | 28 | 1 | 4 | 8 | 13 |
| Total | 27 | 74 | 31 | 142 | 5 | 15 | 27 | 47 |

<sup>a</sup> type I, both loci were M-QTLs, type II, either locus among two loci was M-QTL, and type III, both loci were no
M-QTLs.

## DISCUSSION

In present study, the paternal BC (BC/P) population was constructed to explore the genetic mechanism for fiber quality, following the maternal BC (BC/M) population (Shang *et al.* 2016d; Ma *et al.* 2017, 2019). The backcross design has the obvious advantages: (I) dissecting the genetic components between paternal and maternal backcross populations; (II) identifying more new and novel QTLs for important traits using multiple corresponding populations (BC/P, BC/M and RIL) originated from the same hybrid; and (III) allowing to generate enough hybrid seeds when needed, similar to IF_2_ population.

Here, we detected 19 and 8 QTLs alone in BC/P and BC/M populations, respectively. Three QTLs for fiber strength and fiber elongation shared in both BC populations, including *qFS-Chr21-2, qFE-Chr2-3 and qFE-Chr3-1.* The result indicated that remaining elite alleles (84.21%) showed increasing additive effects originated from male parent for fiber quality in BC/P population. Therefore, the present study was significant for separating the novel elite alleles of male parent for fiber quality.

The identification of stable QTLs (including common QTLs) across multiple environments and multiple populations plays an essential role in marker-assisted selection (MAS) (Jamshed *et al.* 2016). In present study, a total of 12 common QTLs were simultaneously identified in more than one environment(s) or population(s) (**Table 4**). They distributed on Chr 1, Chr 2, Chr 3, Chr 5, Chr 9, Chr 15, Chr 19 and Chr 21. In present study, a total of 13 single locus QTLs (35.14%) for fiber quality were common in comparison with the previous studies in multiple years and multiple locations see Table S5 (Shang *et al.* 2016d; Ma *et al.* 2017). The QTLs verified each other in the RIL population and its BC progenies, suggesting that it is reasonable and effective to map QTLs using different populations across multiple environments and multiple years. The experiment design and the continuous study in our lab contributed to these results in the present study. Among these 13 QTLs, 10 QTLs explained larger than 10% of PV. Five QTLs were identified in Shang *et al.*’ results (2016d) and Ma *et al.*’ results (2017), including *qFL-Chr5-1*, *qFL-Chr5-2*, *qFL-Chr5-3*, *qFS-Chr21-1 and qFE-Chr2-1* (Table S5). For example, region of NAU4034- SWU17713 flanking with *qFL-chr5-1* explained 12.31% of PV on average across 2012 and 2015 in multiple locations. The QTL increased 0.31 mm fiber length (FL) on average, suggesting the major roles and important regions for FL. These verified QTLs were valuable for follow-up breeding program, so as to facilitate fine mapping and favorable gene pyramiding project (Shao *et al.* 2014).

The epistatic effects and environmental interactions existed commonly for fiber quality traits as well as other traits. At two-locus level, we detected a number of interactions under environments for fiber quality traits in both populations (**Table 5**). Three types of epistasis combinations were observed (**Table 6**). However, epistasis QTLs influenced fiber quality by Type III (57.45%) in BC/P population. Differently, epistasis influenced fiber quality by Type I and Type II (71.13%) in RIL population. In addition, 3-fold epistasis QTLs was detected in RIL-P population than that in BC/P population. The results indicating that epistasis played roles in different genetic mode to control fiber quality. Especially, no E-QTL was identified for fiber uniformity (FU). Another interesting result detected that epistasis is another vital genetic effect affecting fiber quality traits (Shang *et al.* 2016d). Similarly to the previous study, Wang *et al.* (2006) indicated that both epistasis effect and single-locus effect of QTLs played important genetic roles in cotton fiber quality.

In the present study, the five QTLs increased fiber micronaire (FM) values from 0.06-0.18 (**Table 4**). The mean values ranged from 4.71-5.18 on average in the RIL-P and BC/P populations of Xinza 1. In fact, fiber quality ranks from B grade (3.5-3.6, 4.3-4.9) to C grade (<3.4, >5.0) for fiber micronaire, in which the fiber change larger thickness to be worse. At the same time, FM displayed negative correlation with FL, FU, FS and FE. Therefore, we should avoid exploiting these QTL regions for FM when breeding in cotton. Therefore, we will not focus on the region of NAU2873-CGR6771 on Chr 9, contributing alleles to FS and increased the FM values. At the same time, larger lint yield and well fiber quality are the key aims in cotton breeding program. However, negative correlation between yield and fiber quality hinders the cotton breeding (Yang *et al.* 2015). Many important heterotic loci were detected to increase yield or yield-components in previous studies in our lab (Shang *et al.* 2016a; Ma *et al.* 2019). Some heterotic loci were also identified for improving fiber quality in present and previous studies (Shang *et al.* 2016d; Ma *et al.* 2017). The important pleiotropic regions should be paying more attention in further research so as to improve fiber quality and to increase yield in breeding program.

## AUTHOR CONTRIBUTIONS

LM performed the experiments, analyzed the data and prepared the manuscript. YW maintained the experimental platform and attended bench work. YS, HN, YC, CC and MW attended field experiments and data collection. JH conceived the experiments, provided experimental platform and revised the manuscript.

## ACKNOWLEDGMENTS

We thank Shihu Cai (China Agricultural University), Dongyong Xu and Huaiyu Lu (Guoxin Seed Company Ltd, Cangzhou, Hebei Province) for their contributions on field experiments and data acquisition. Thanks to Qingzhi Liang, Liangguang Shang, Abdugheni Abduweli, Xiaocui Wang (China Agricultural University) for the contributions in SSR marker evaluation and map construction, Kunbo Wang and Fang Liu (Institute of Cotton Research, Chinese Academy of Agricultural Sciences) for providing part of SSR markers, Dr. Zhengsheng Zhang (Southwest University) for providing SWU and ICR SSR primers. This research was supported by the National Key R & D Program for Crop Breeding (2016YFD0101407) to J.H.

---

**Table S2.**
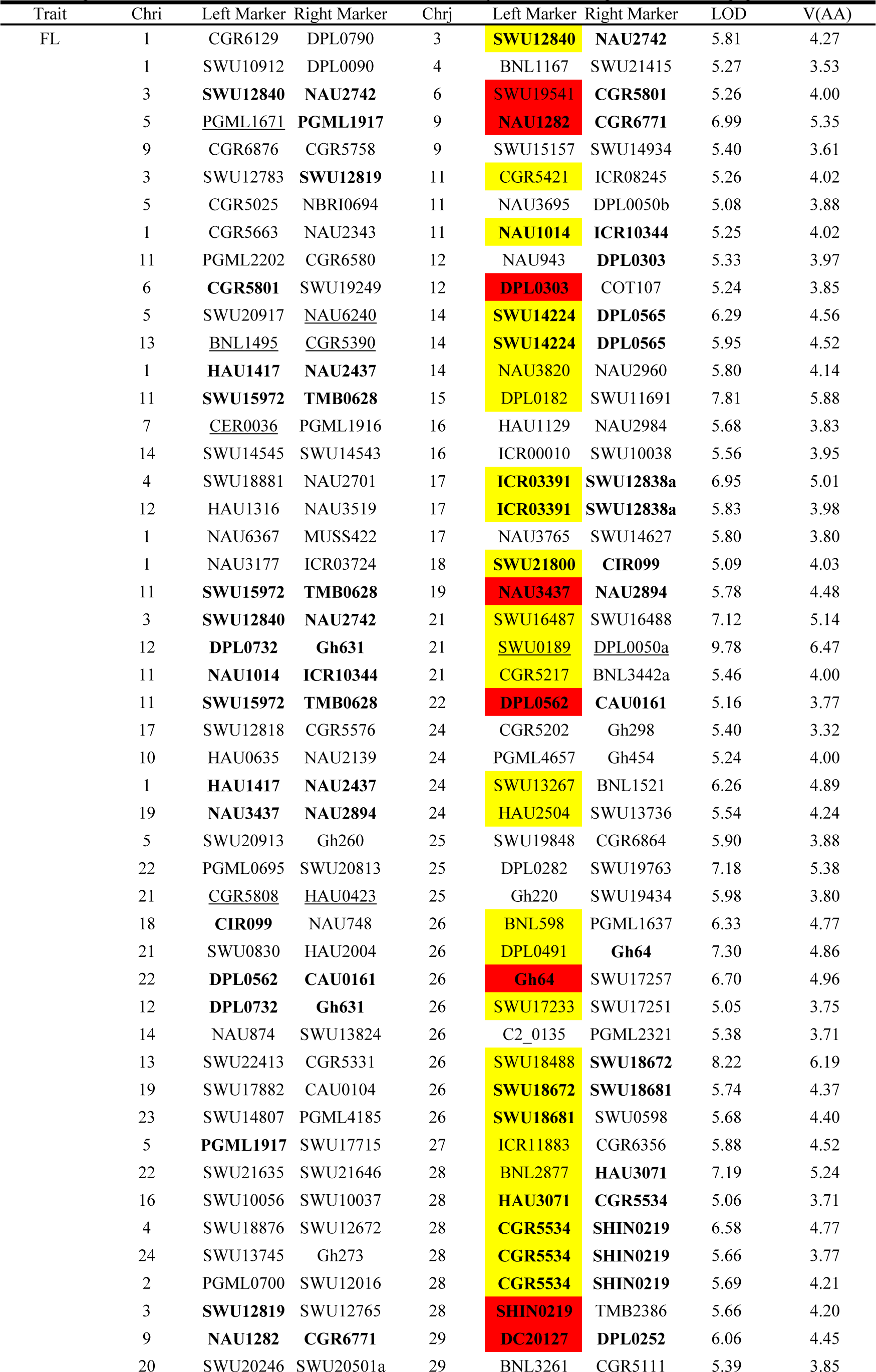
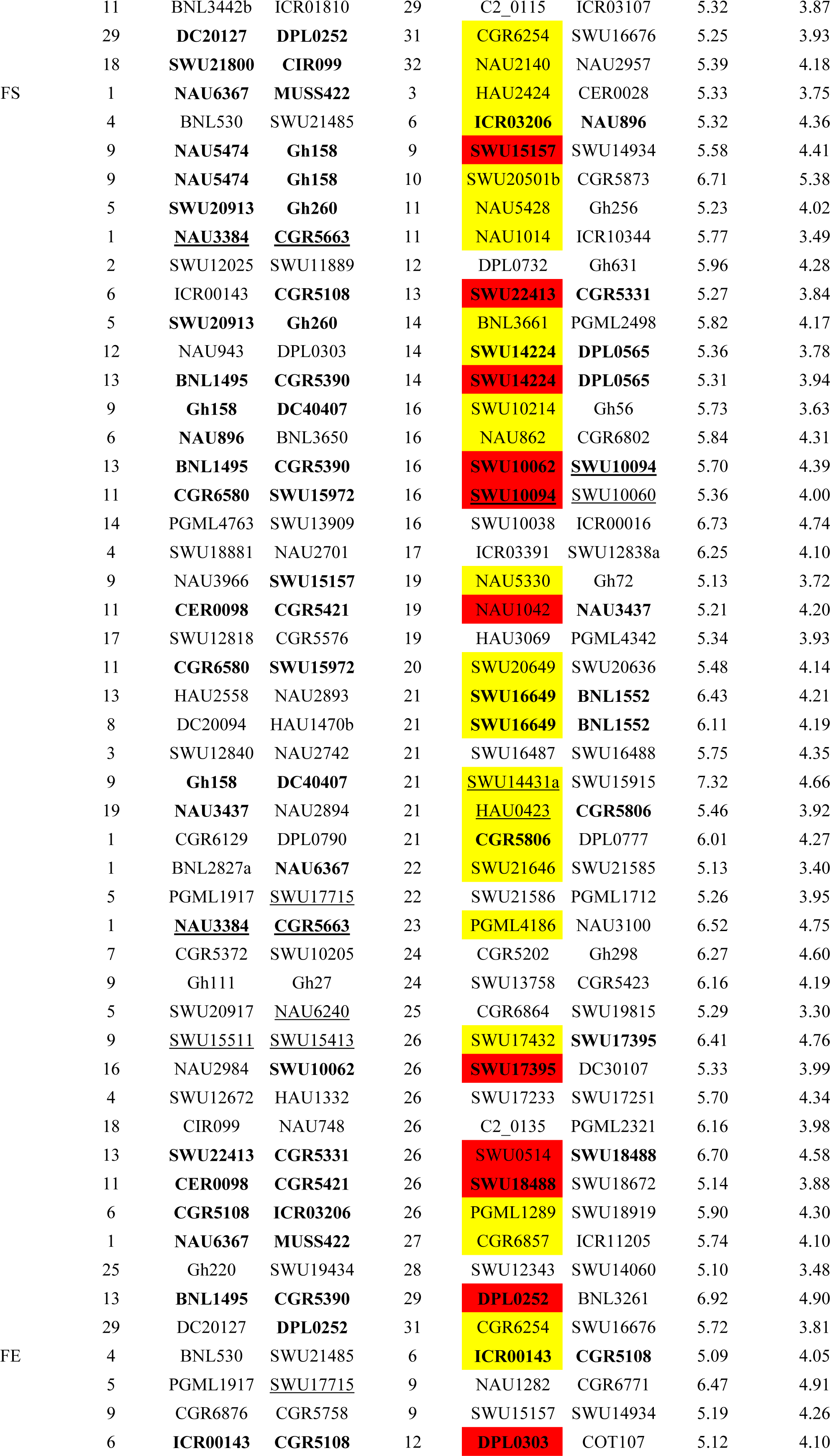
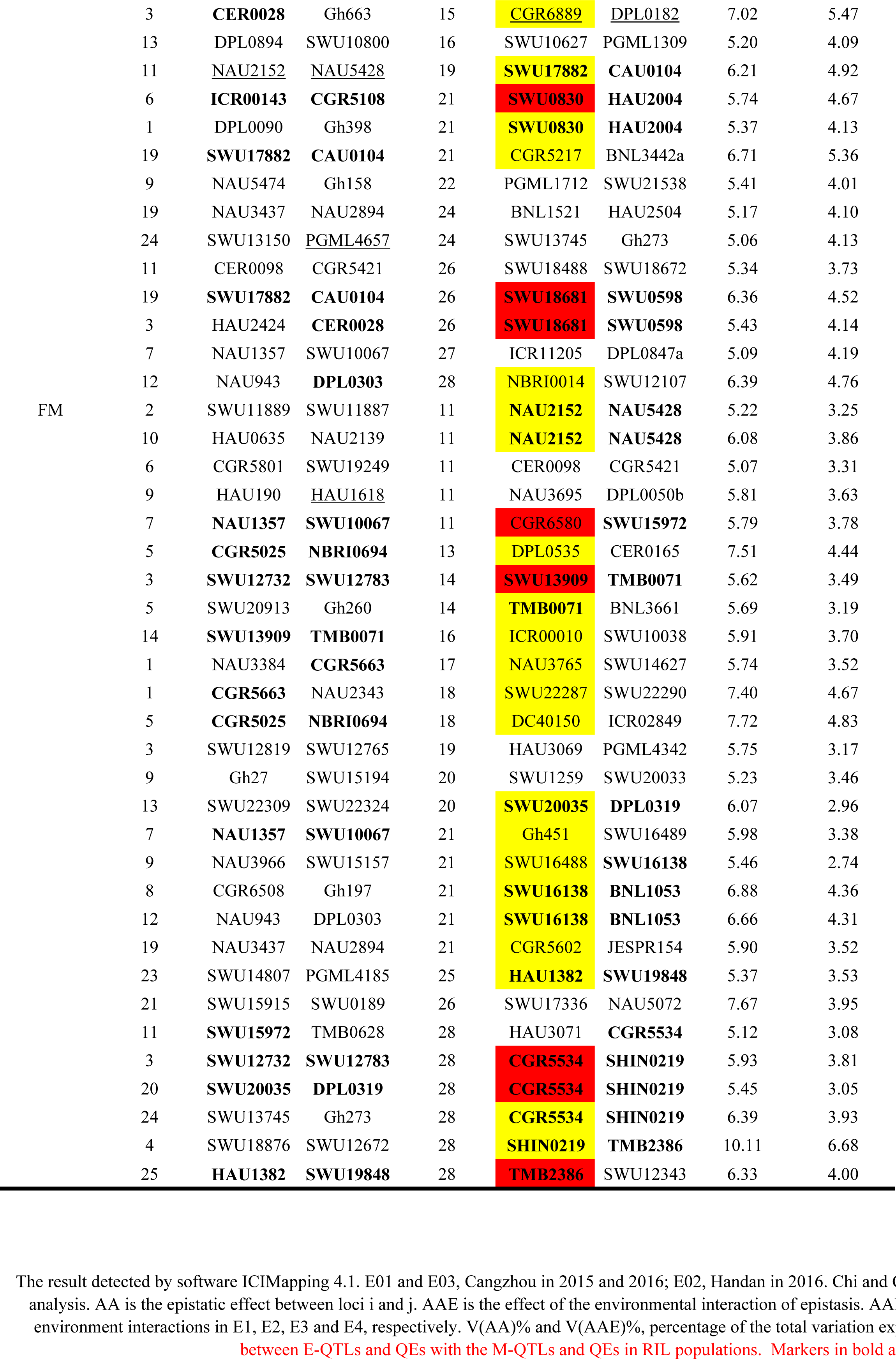
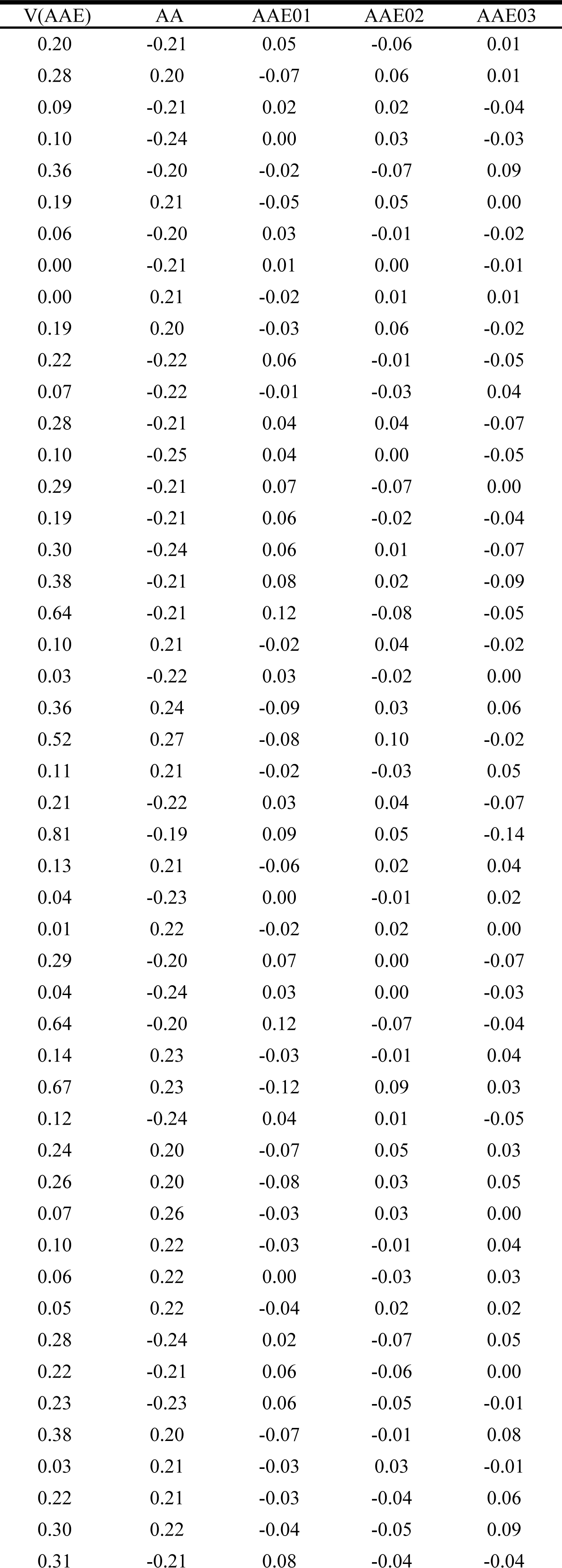
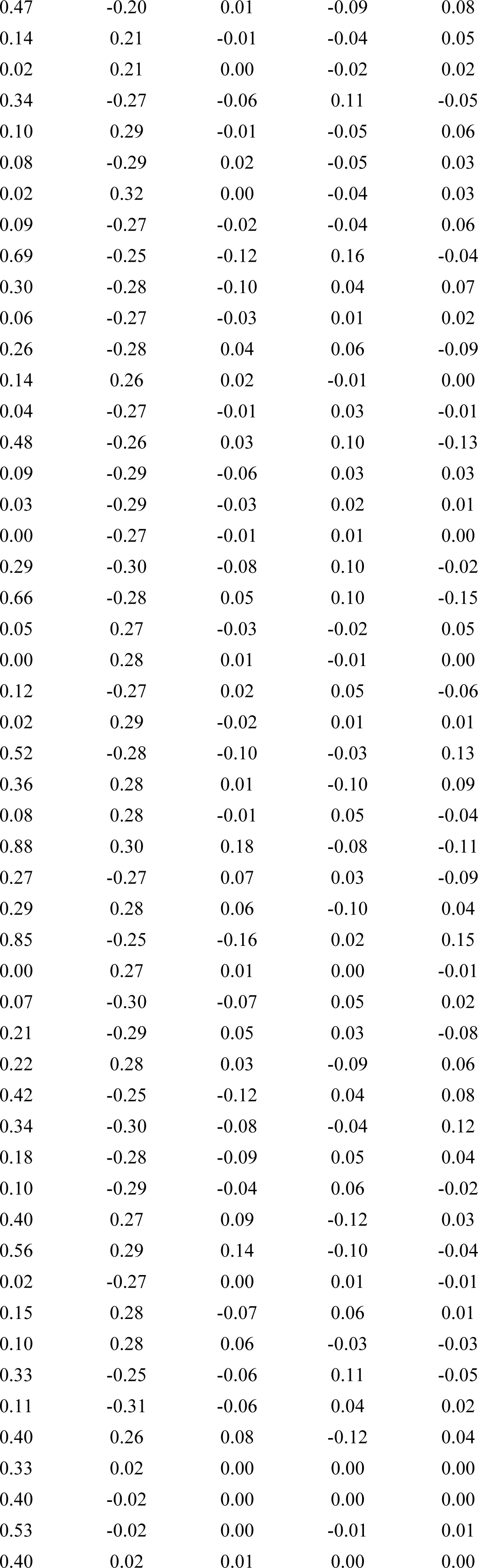
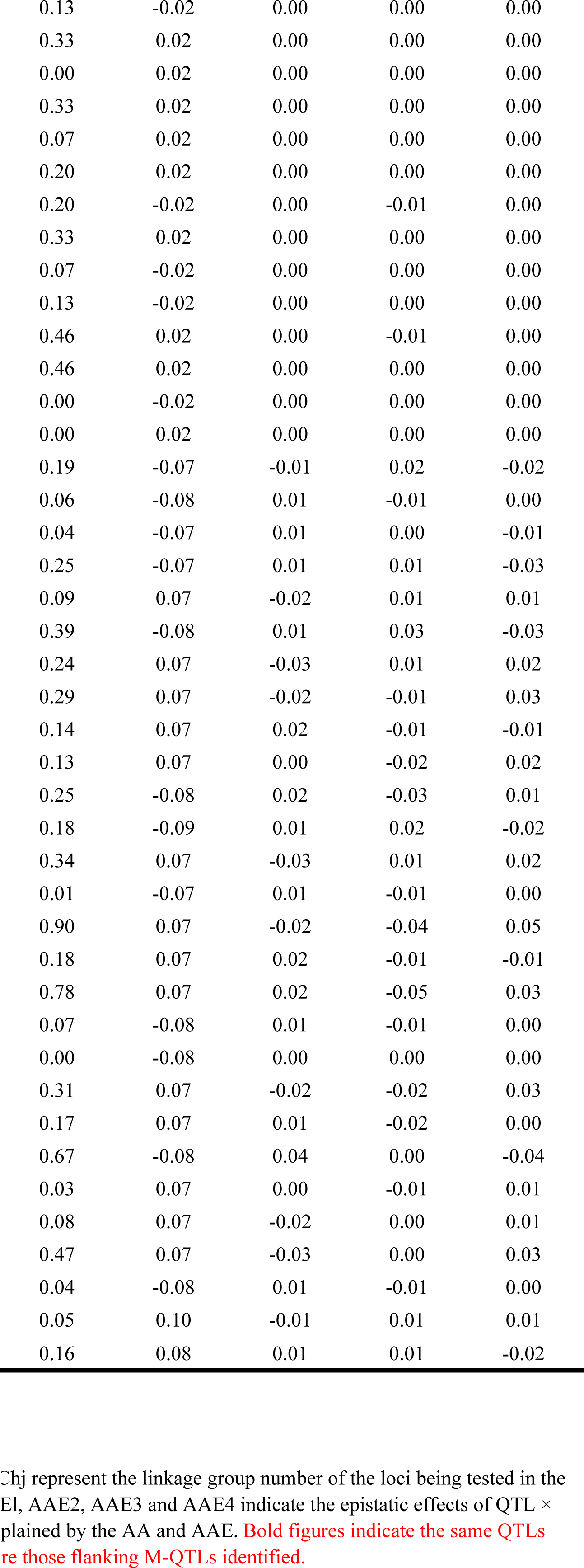
Epitasis effects and environmental interactions detected for yield and its components in RIL-P population

**Table S5.**
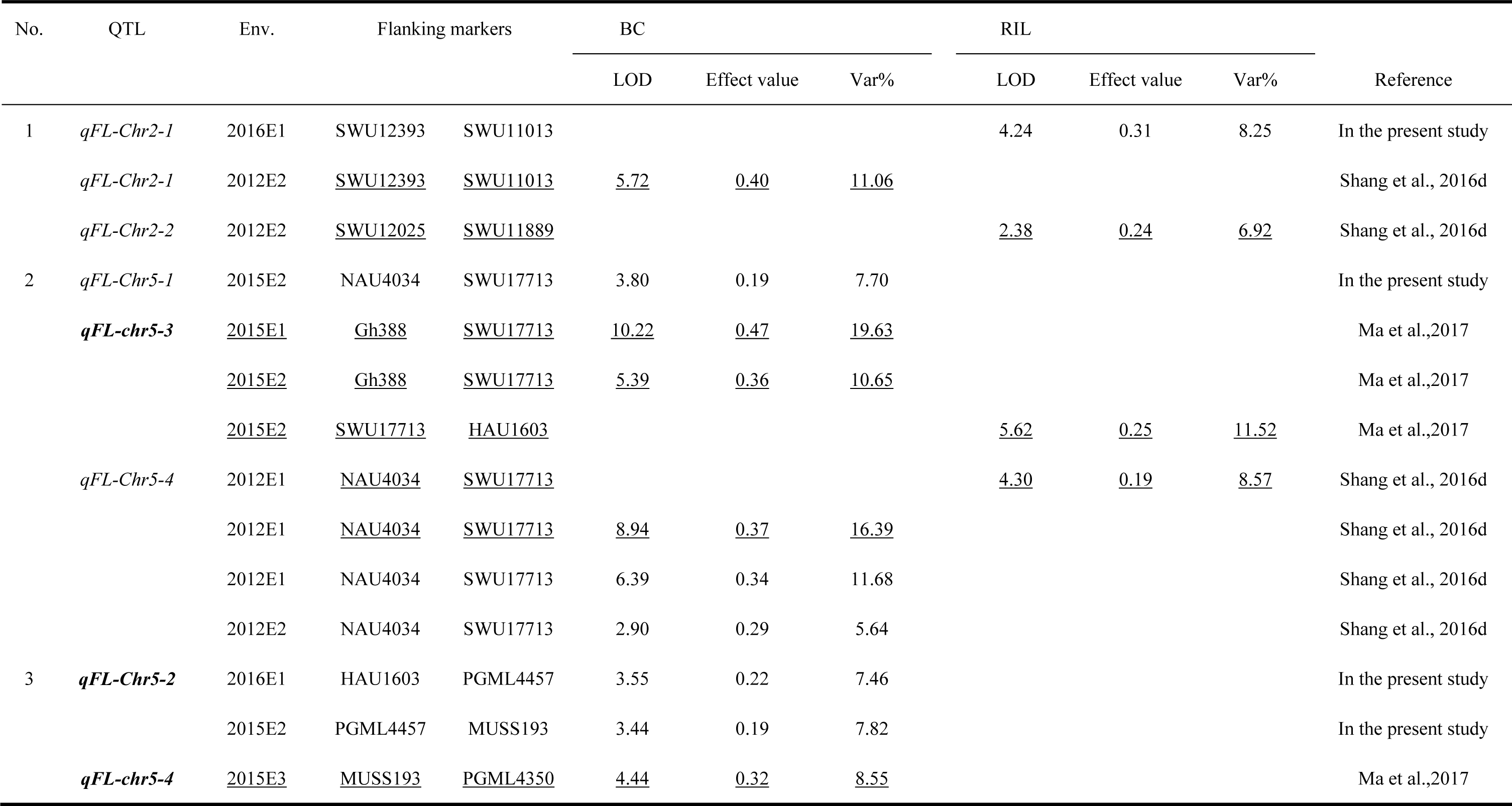
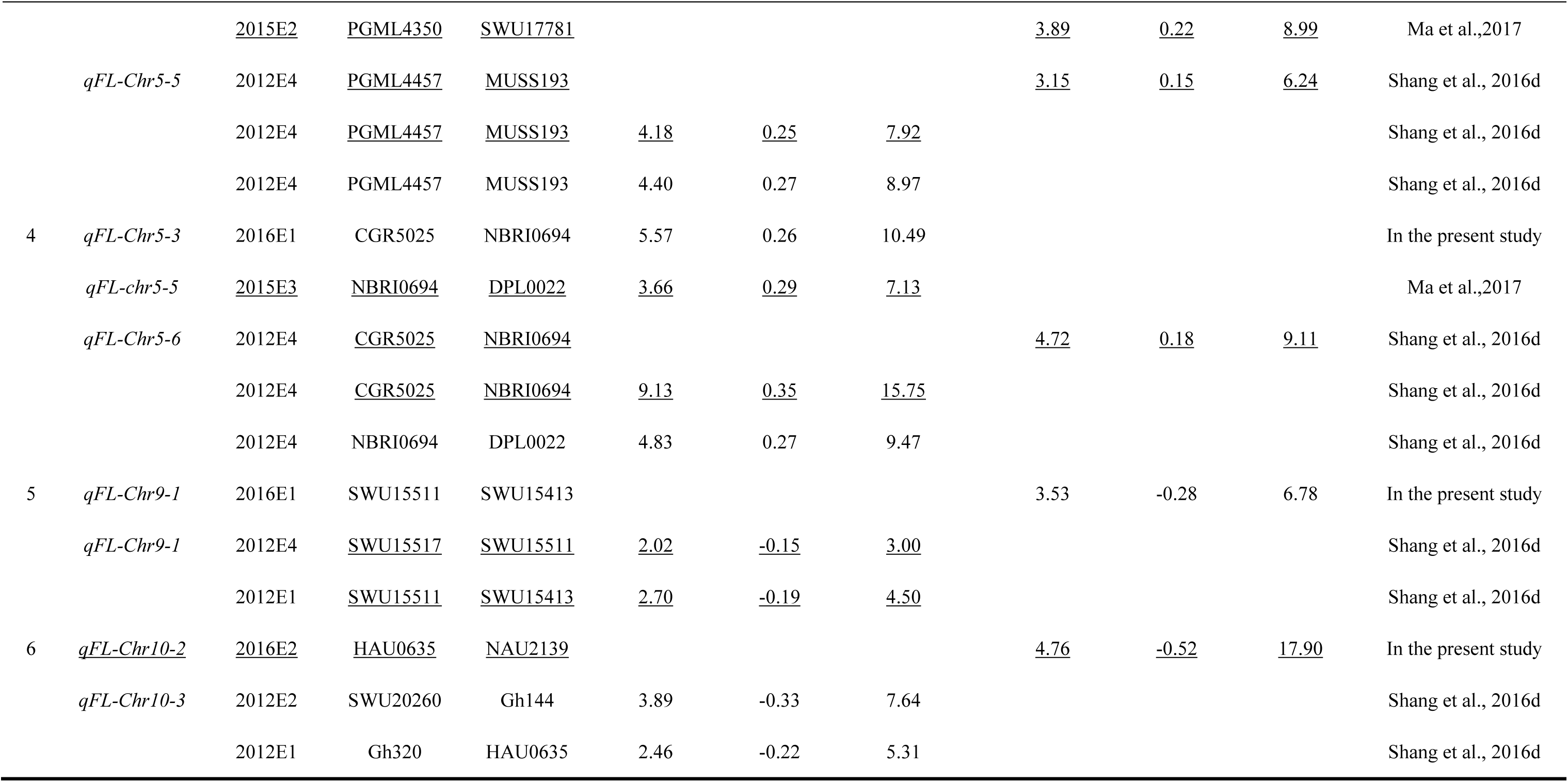
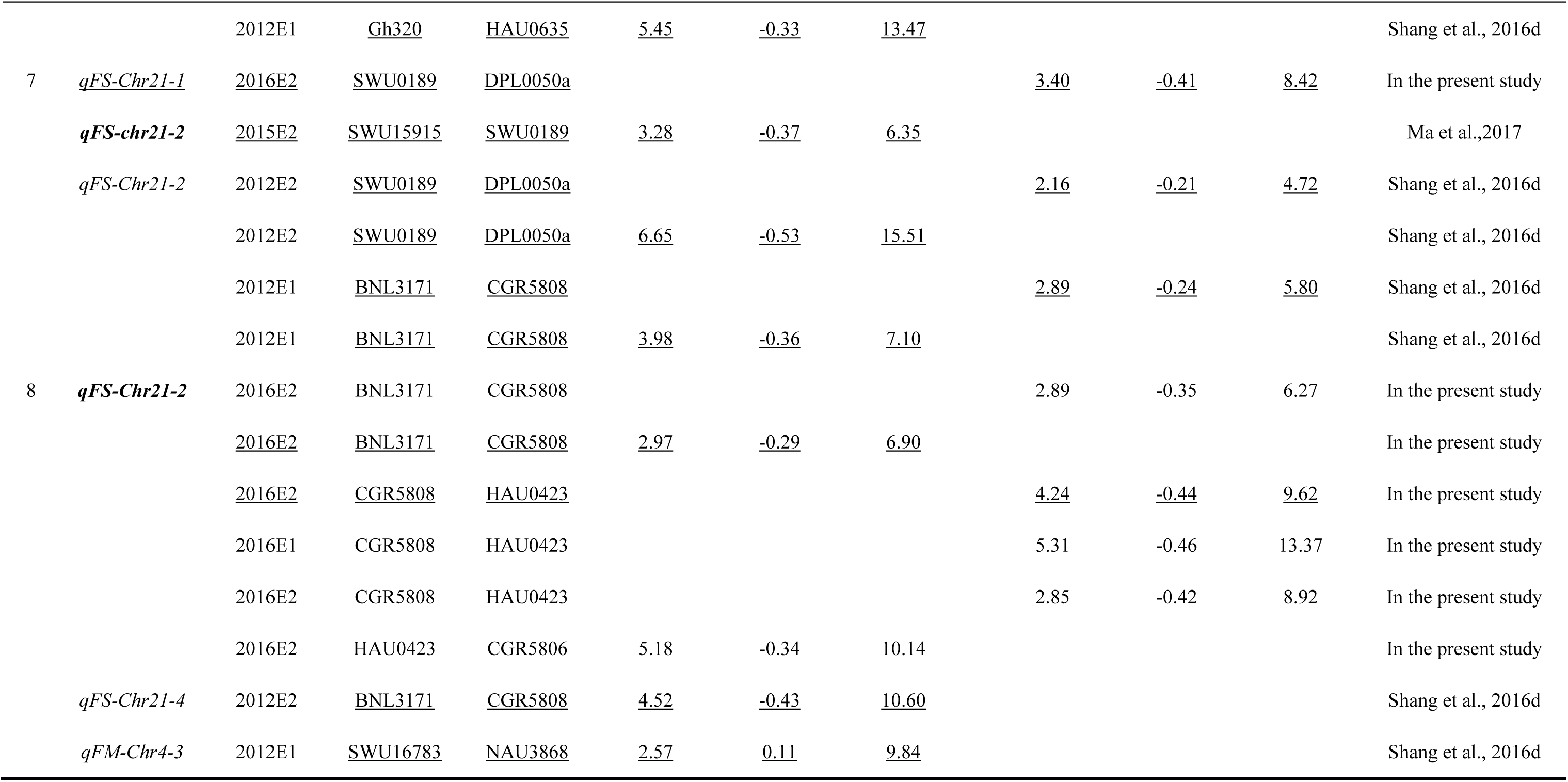
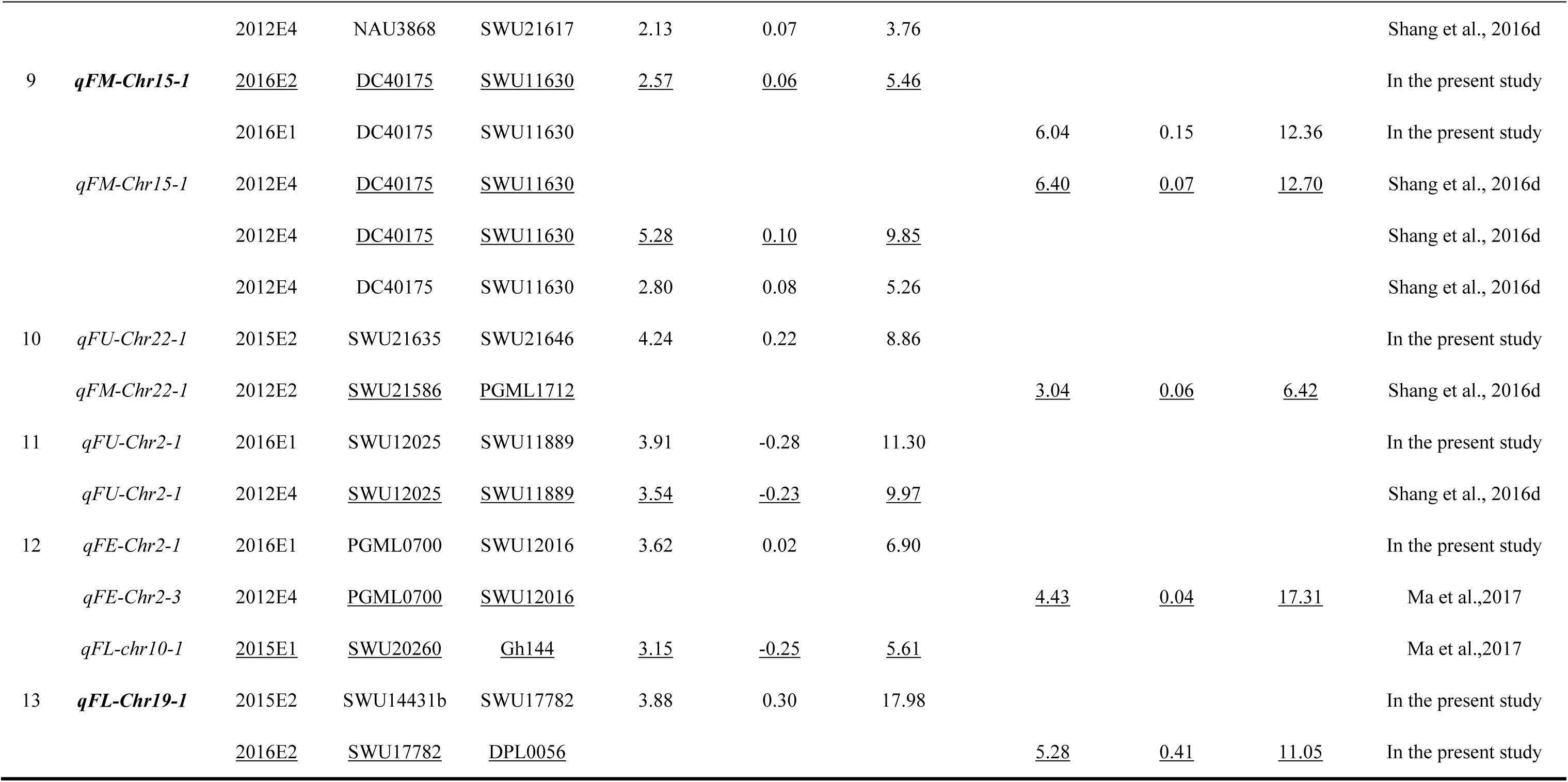
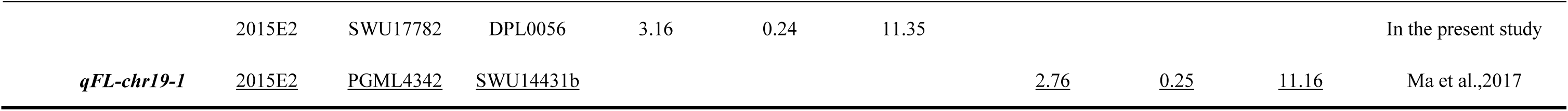
Common single locus QTLs in comparison with the previous studies in multiple years and locations

**Table S6.** Phenotype values of fiber quality traits of BC/M population in three field trials in this study

| Number of field trial<br>plot | Material | FL/mm | FU/% | FS(cN/tex<br>) | FE/% | FM | FL/mm | FU/% |
| --- | --- | --- | --- | --- | --- | --- | --- | --- |
|  |  | 2015E2 | 2015E2 | 2015E2 | 2015E2 | 2015E2 | 2016E1 | 2016E1 |
| 001 | 14QM003 | 28.9 | 85.5 | 30.2 | 6.9 | 5.0 | 29.6 | 87.3 |
| 002 | 14QM003H | 29.6 | 87.3 | 28.9 | 6.9 | 5.0 | 30.3 | 88.2 |
| 003 | P♂ | 28.1 | 86.5 | 29.3 | 6.8 | 4.1 | 29.5 | 88.0 |
| 004 | 14QM004H | 29.6 | 85.5 | 30.5 | 7.0 | 4.6 | 29.8 | 87.9 |
| 005 | 14QM004 | 29.0 | 84.1 | 30.1 | 7.0 | 5.0 | 30.4 | 87.6 |
| 006 | 14QM005 | 29.3 | 86.3 | 29.8 | 6.9 | 5.2 | 32.3 | 86.0 |
| 007 | 14QM005H | 28.6 | 84.3 | 29.8 | 6.8 | 4.6 | 30.3 | 86.1 |
| 008 | P♂ | 29.7 | 86.0 | 29.9 | 6.9 | 4.9 | 29.5 | 88.0 |
| 009 | 14QM006H | 31.1 | 86.3 | 31.1 | 6.9 | 4.6 | 31.1 | 86.8 |
| 010 | 14QM006 | 30.6 | 86.1 | 31.4 | 7.0 | 5.3 | 30.7 | 86.9 |
| 011 | 14QM007 | 29.5 | 86.2 | 29.2 | 6.9 | 5.0 | 29.8 | 86.3 |
| 012 | 14QM007H | 30.7 | 85.2 | 31.1 | 7.0 | 5.2 | 31.6 | 88.5 |
| 013 | P♂ | 29.6 | 85.1 | 30.0 | 6.9 | 4.4 | 29.5 | 88.0 |
| 014 | 14QM008H | 30.9 | 84.7 | 30.7 | 7.0 | 4.5 | 30.8 | 87.6 |
| 015 | 14QM008 | 29.9 | 85.2 | 28.9 | 6.9 | 4.6 | 31.3 | 86.7 |
| 016 | 14QM009 | 28.1 | 85.4 | 28.4 | 6.8 | 4.1 | 29.9 | 86.0 |
| 017 | 14QM009H | 28.7 | 85.4 | 29.0 | 6.8 | 4.8 | 29.0 | 86.7 |
| 018 | P♂ | 27.8 | 84.7 | 27.6 | 6.9 | 5.0 | 29.5 | 88.0 |
| 019 | 14QM010H | 30.4 | 86.6 | 30.7 | 7.0 | 4.7 | 30.8 | 87.8 |
| 020 | 14QM010 | 31.4 | 85.6 | 32.3 | 7.0 | 4.3 | 32.1 | 86.1 |
| 021 | 14QM011 | 31.5 | 86.0 | 31.9 | 7.0 | 4.9 | 28.9 | 86.4 |
| 022 | 14QM011H | 31.2 | 87.5 | 31.2 | 7.0 | 5.1 | 32.1 | 87.5 |
| 023 | P♂ | 30.4 | 87.0 | 30.2 | 6.9 | 4.3 | 29.5 | 88.0 |
| 024 | 14QM012H | 30.7 | 87.3 | 31.8 | 7.0 | 5.1 | 30.9 | 85.4 |
| 025 | 14QM012 | 31.6 | 84.9 | 35.6 | 7.0 | 4.3 | 28.8 | 84.3 |
| 026 | 14QM013 | 29.6 | 85.2 | 30.3 | 7.1 | 4.9 | 28.7 | 85.9 |
| 027 | 14QM013H | 30.3 | 84.9 | 30.4 | 7.0 | 4.5 | 29.0 | 86.1 |
| 028 | P♂ | 29.9 | 86.0 | 29.2 | 6.9 | 4.4 | 29.5 | 88.0 |
| 029 | 14QM014H | 32.5 | 86.6 | 31.3 | 7.0 | 4.9 | 30.6 | 87.0 |
| 030 | 14QM014 | 29.6 | 83.4 | 29.4 | 6.8 | 3.9 | 29.9 | 85.1 |
| 031 | 14QM015 | 30.3 | 85.5 | 29.6 | 7.0 | 4.8 | 30.0 | 85.6 |
| 032 | 14QM015H | 29.2 | 86.1 | 28.3 | 6.8 | 4.6 | 30.5 | 86.9 |
| 033 | P♂ | 28.6 | 85.7 | 31.0 | 6.8 | 3.9 | 29.5 | 88.0 |
| 034 | 14QM016H | 30.4 | 85.7 | 30.0 | 6.9 | 5.1 | 29.9 | 87.8 |
| 035 | 14QM016 | 31.1 | 86.0 | 32.8 | 7.0 | 4.7 | 31.8 | 87.4 |
| 036 | 14QM017 | 30.5 | 86.1 | 32.6 | 7.0 | 4.2 | 30.7 | 86.5 |
| 037 | 14QM017H | 30.3 | 84.8 | 30.8 | 7.2 | 5.0 | 31.1 | 87.5 |
| 038 | P♂ | 29.7 | 85.6 | 29.7 | 6.9 | 4.4 | 29.5 | 88.0 |
| 039 | 14QM018H | 29.9 | 84.3 | 30.8 | 6.9 | 4.6 | 30.5 | 86.0 |
| 040 | 14QM018 | 32.0 | 86.7 | 34.1 | 7.0 | 4.0 | 29.9 | 84.9 |
| 041 | 14QM019 | 30.7 | 86.2 | 32.4 | 7.0 | 4.7 | 28.8 | 84.1 |
| 042 | 14QM019H | 29.7 | 85.3 | 29.1 | 6.8 | 4.8 | 30.2 | 86.6 |
| 043 | P♂ | 29.6 | 84.5 | 27.9 | 7.0 | 4.9 | 29.5 | 88.0 |
| 044 | 14QM020H | 30.4 | 86.0 | 30.7 | 7.0 | 4.5 | 30.5 | 86.9 |
| 045 | 14QM020 | 30.9 | 82.9 | 29.6 | 6.9 | 4.3 | 32.3 | 86.2 |
| 046 | 14QM021 | 30.3 | 85.2 | 31.2 | 6.9 | 4.7 | 30.9 | 85.1 |
| 047 | 14QM021H | 31.0 | 85.8 | 30.8 | 7.0 | 4.7 | 29.9 | 87.3 |
| 048 | P♂ | 29.6 | 84.8 | 28.1 | 6.9 | 4.7 | 29.5 | 88.0 |
| 049 | 14QM022H | 30.7 | 86.4 | 32.1 | 7.1 | 4.7 | 30.7 | 87.2 |
| 050 | 14QM022 | 30.1 | 84.4 | 30.4 | 7.1 | 4.7 | 30.3 | 86.3 |
| 051 | 14QM023 | 31.9 | 85.2 | 30.5 | 7.0 | 4.5 | 29.9 | 86.0 |
| 052 | 14QM023H | 30.8 | 84.8 | 29.1 | 7.0 | 4.8 | 30.0 | 86.7 |
| 053 | 14QM380 | 30.9 | 86.5 | 29.7 | 7.0 | 4.7 | 28.9 | 86.4 |
| 054 | 14QM024H | 32.2 | 86.5 | 32.5 | 7.3 | 4.5 | 31.4 | 86.1 |
| 055 | 14QM024 | 33.1 | 86.7 | 33.0 | 7.3 | 4.4 | 30.4 | 86.7 |
| 056 | 14QM025 | - | - | - | - | - | 30.7 | 85.9 |
| 057 | 14QM025H | 31.6 | 87.2 | 31.9 | 7.1 | 4.9 | 29.5 | 87.8 |
| 058 | P♂ | 29.8 | 85.5 | 29.0 | 6.9 | 4.5 | 28.9 | 86.4 |
| 059 | 14QM026H | 30.2 | 87.0 | 30.0 | 7.0 | 5.0 | 30.1 | 86.9 |
| 060 | 14QM026 | 29.6 | 86.2 | 28.1 | 7.1 | 4.9 | 30.3 | 87.3 |
| 061 | 14QM027 | 30.6 | 85.8 | 28.5 | 7.0 | 4.9 | 29.1 | 87.0 |
| 062 | 14QM027H | 30.8 | 87.4 | 29.3 | 7.0 | 5.1 | 29.9 | 88.2 |
| 063 | P♂ | 32.1 | 85.9 | 31.7 | 7.1 | 4.8 | 28.9 | 86.4 |
| 064 | 14QM028H | 30.3 | 85.9 | 29.7 | 6.9 | 4.6 | 31.0 | 86.8 |
| 065 | 14QM028 | 29.8 | 84.9 | 29.6 | 7.0 | 5.0 | 29.7 | 86.1 |
| 066 | 14QM029 | 32.1 | 86.7 | 29.9 | 7.1 | 4.9 | 31.6 | 87.3 |
| 067 | 14QM029H | 31.1 | 87.2 | 30.6 | 7.0 | 4.8 | 30.4 | 84.1 |
| 068 | P♂ | 30.0 | 87.3 | 29.9 | 6.9 | 4.7 | 28.9 | 86.4 |
| 069 | 14QM030H | 32.1 | 84.8 | 32.7 | 7.0 | 4.2 | 31.3 | 86.8 |
| 070 | 14QM030 | 32.6 | 86.2 | 33.8 | 7.2 | 4.1 | 31.7 | 86.8 |
| 071 | 14QM031 | 32.0 | 87.0 | 31.9 | 7.1 | 4.0 | 31.4 | 84.4 |
| 072 | 14QM031H | 32.0 | 85.8 | 31.4 | 7.0 | 4.5 | 32.0 | 84.8 |
| 073 | P♂ | 31.5 | 86.4 | 31.2 | 7.1 | 4.4 | 28.9 | 86.4 |
| 074 | 14QM032H | 32.1 | 86.4 | 31.7 | 7.0 | 5.2 | 30.6 | 86.3 |
| 075 | 14QM032 | 30.6 | 84.0 | 31.9 | 7.1 | 5.1 | 30.2 | 87.5 |
| 076 | 14QM033 | 29.9 | 86.0 | 31.1 | 6.9 | 4.9 | 30.3 | 86.5 |
| 077 | 14QM033H | 31.3 | 85.7 | 31.3 | 7.1 | 5.3 | 30.7 | 87.0 |
| 078 | P♂ | 30.3 | 87.2 | 29.8 | 6.9 | 4.8 | 28.9 | 86.4 |
| 079 | 14QM034H | 30.4 | 85.6 | 31.0 | 7.0 | 5.0 | 30.6 | 87.8 |
| 080 | 14QM034 | 31.4 | 86.0 | 30.7 | 7.1 | 4.6 | 30.8 | 87.1 |
| 081 | 14QM035 | 32.5 | 86.6 | 33.8 | 7.0 | 4.4 | 30.2 | 86.1 |
| 082 | 14QM035H | 30.5 | 85.1 | 30.7 | 7.0 | 4.8 | 30.6 | 87.0 |
| 083 | P♂ | 30.6 | 86.6 | 29.1 | 6.9 | 4.8 | 28.9 | 86.4 |
| 084 | 14QM036H | 31.4 | 85.4 | 30.4 | 7.0 | 4.6 | 31.0 | 86.6 |
| 085 | 14QM036 | 31.4 | 84.9 | 30.8 | 6.9 | 4.1 | 31.1 | 85.8 |
| 086 | 14QM037 | 29.8 | 84.8 | 30.2 | 6.9 | 5.3 | 30.7 | 87.5 |
| 087 | 14QM037H | 31.2 | 86.7 | 32.1 | 7.0 | 4.5 | 30.7 | 85.7 |
| 088 | P♂ | 29.5 | 85.2 | 30.6 | 6.9 | 4.8 | 28.9 | 86.4 |
| 089 | 14QM038H | 30.1 | 86.9 | 27.3 | 6.9 | 4.8 | 29.6 | 86.7 |
| 090 | 14QM038 | 31.4 | 86.1 | 30.5 | 7.0 | 4.7 | 29.2 | 86.5 |
| 091 | 14QM039 | 30.3 | 86.7 | 30.0 | 7.0 | 5.5 | 29.7 | 85.8 |
| 092 | 14QM039H | 31.0 | 86.5 | 30.8 | 7.0 | 5.3 | 30.4 | 88.1 |
| 093 | P♂ | 29.9 | 87.1 | 28.8 | 6.9 | 5.1 | 28.9 | 86.4 |
| 094 | 14QM040H | 30.9 | 86.9 | 30.0 | 7.0 | 4.8 | 30.1 | 86.0 |
| 095 | 14QM040 | 30.5 | 87.2 | 31.0 | 7.0 | 4.8 | 30.0 | 86.9 |
| 096 | 14QM041 | 32.5 | 87.3 | 32.0 | 7.2 | 4.8 | 31.8 | 86.7 |
| 097 | 14QM041H | 31.6 | 86.1 | 32.0 | 7.1 | 4.3 | 30.5 | 86.1 |
| 098 | P♂ | 30.6 | 84.3 | 29.1 | 6.9 | 4.5 | 28.9 | 86.4 |
| 099 | 14QM042H | 30.9 | 85.8 | 30.6 | 7.1 | 5.3 | 29.9 | 88.2 |
| 100 | 14QM042 | 31.2 | 87.0 | 30.2 | 7.1 | 5.1 | 30.2 | 87.3 |
| 101 | 14QM043 | 30.2 | 84.7 | 29.8 | 7.0 | 5.2 | 29.8 | 87.2 |
| 102 | 14QM043H | 32.2 | 87.2 | 30.8 | 7.1 | 5.2 | 30.6 | 87.6 |
| 103 | 14QM380 | 29.5 | 86.4 | 29.1 | 6.9 | 5.0 | 28.1 | 83.1 |
| 104 | 14QM044H | 30.7 | 86.6 | 29.9 | 7.1 | 5.2 | 29.6 | 86.4 |
| 105 | 14QM044 | 30.8 | 84.8 | 29.4 | 7.0 | 5.0 | 31.1 | 87.7 |
| 106 | 14QM045 | 32.4 | 87.1 | 31.6 | 7.1 | 4.7 | 30.7 | 87.3 |
| 107 | 14QM045H | 31.9 | 85.9 | 29.0 | 7.0 | 5.0 | 31.5 | 87.1 |
| 108 | P♂ | 30.9 | 86.1 | 30.2 | 7.0 | 4.4 | 28.1 | 83.1 |
| 109 | 14QM046H | 29.7 | 86.2 | 29.2 | 7.0 | 4.7 | 29.9 | 88.0 |
| 110 | 14QM046 | 29.3 | 84.4 | 28.6 | 6.9 | 4.9 | 29.0 | 87.1 |
| 111 | 14QM047 | 31.5 | 85.4 | 30.7 | 7.0 | 5.1 | 30.9 | 86.1 |
| 112 | 14QM047H | 32.2 | 86.3 | 31.6 | 7.0 | 5.1 | 30.3 | 87.1 |
| 113 | P♂ | 30.5 | 86.6 | 30.4 | 7.0 | 5.0 | 28.1 | 83.1 |
| 114 | 14QM048H | 31.2 | 84.3 | 31.1 | 7.0 | 4.4 | 30.0 | 87.9 |
| 115 | 14QM048 | 30.3 | 85.4 | 30.6 | 7.0 | 4.7 | 29.3 | 84.5 |
| 116 | 14QM049 | 31.1 | 85.1 | 30.6 | 7.1 | 4.1 | 29.5 | 87.5 |
| 117 | 14QM049H | 32.0 | 85.4 | 30.5 | 7.1 | 4.6 | 30.7 | 86.8 |
| 118 | P♂ | 31.1 | 85.8 | 31.6 | 7.0 | 5.1 | 28.1 | 83.1 |
| 119 | 14QM050H | 29.8 | 86.1 | 28.9 | 6.9 | 4.7 | 31.9 | 86.9 |
| 120 | 14QM050 | 32.0 | 86.1 | 31.2 | 7.0 | 4.6 | 31.2 | 86.3 |
| 121 | 14QM051 | 31.7 | 86.5 | 33.2 | 7.0 | 4.6 | 31.2 | 88.3 |
| 122 | 14QM051H | 31.6 | 86.2 | 30.9 | 6.9 | 4.4 | 30.2 | 87.5 |
| 123 | P♂ | 28.7 | 85.8 | 28.6 | 6.9 | 4.7 | 28.1 | 83.1 |
| 124 | 14QM052H | 30.7 | 85.1 | 30.1 | 7.0 | 4.7 | 30.3 | 87.5 |
| 125 | 14QM052 | - | - | - | - | - | 29.9 | 86.6 |
| 126 | 14QM053 | 33.0 | 85.7 | 34.3 | 7.1 | 3.6 | 30.9 | 86.4 |
| 127 | 14QM053H | 31.1 | 85.3 | 31.9 | 7.2 | 4.4 | 31.9 | 87.0 |
| 128 | P♂ | 29.4 | 86.5 | 28.9 | 6.9 | 5.0 | 28.1 | 83.1 |
| 129 | 14QM054H | 29.9 | 85.7 | 29.9 | 6.9 | 4.8 | 28.6 | 84.8 |
| 130 | 14QM054 | 30.3 | 86.0 | 28.8 | 7.0 | 4.8 | 29.4 | 86.9 |
| 131 | 14QM055 | 29.9 | 85.4 | 31.4 | 6.9 | 5.4 | 30.0 | 88.1 |
| 132 | 14QM055H | 31.2 | 85.5 | 30.9 | 6.9 | 4.7 | 28.9 | 85.8 |
| 133 | P♂ | 30.4 | 83.8 | 29.7 | 6.9 | 4.3 | 28.1 | 83.1 |
| 134 | 14QM056H | 31.4 | 85.4 | 33.1 | 7.0 | 4.4 | 32.0 | 86.4 |
| 135 | 14QM056 | 29.9 | 82.7 | 33.6 | 6.9 | 3.4 | 32.1 | 85.7 |
| 136 | 14QM057 | 29.6 | 84.1 | 30.2 | 6.9 | 3.9 | 29.3 | 85.8 |
| 137 | 14QM057H | 30.8 | 83.7 | 31.2 | 7.0 | 4.5 | 28.0 | 85.4 |
| 138 | P♂ | 29.4 | 82.9 | 30.6 | 7.0 | 5.0 | 28.1 | 83.1 |
| 139 | 14QM058H | 28.9 | 84.6 | 30.7 | 6.9 | 4.8 | 30.2 | 85.5 |
| 140 | 14QM058 | 31.3 | 83.5 | 32.6 | 7.0 | 4.6 | 31.2 | 85.6 |
| 141 | 14QM059 | 28.2 | 85.0 | 28.3 | 6.8 | 4.6 | 31.0 | 86.3 |
| 142 | 14QM059H | 30.4 | 85.3 | 29.8 | 7.0 | 4.9 | 30.0 | 88.4 |
| 143 | P♂ | 30.8 | 86.5 | 29.4 | 6.9 | 4.7 | 28.1 | 83.1 |
| 144 | 14QM060H | 31.1 | 86.4 | 30.4 | 7.0 | 4.6 | 30.3 | 87.7 |
| 145 | 14QM060 | 33.3 | 85.5 | 33.9 | 7.1 | 4.3 | 31.6 | 86.4 |
| 146 | 14QM061 | 29.5 | 86.4 | 30.0 | 6.9 | 5.3 | 29.1 | 85.3 |
| 147 | 14QM061H | 30.1 | 84.7 | 29.2 | 6.8 | 5.1 | 28.8 | 86.3 |
| 148 | P♂ | 31.1 | 85.5 | 29.9 | 7.0 | 4.5 | 28.1 | 83.1 |
| 149 | 14QM062H | 31.7 | 86.7 | 29.9 | 7.0 | 4.6 | 30.6 | 88.5 |
| 150 | 14QM062 | 32.3 | 85.7 | 33.4 | 7.0 | 4.2 | 32.5 | 86.4 |
| 151 | 14QM063 | 31.2 | 86.4 | 31.4 | 7.0 | 5.0 | 29.1 | 86.1 |
| 152 | 14QM063H | 31.1 | 86.1 | 32.8 | 7.0 | 4.6 | 30.9 | 87.5 |
| 153 | 14QM380 | 30.7 | 85.7 | 30.2 | 7.0 | 4.6 | 30.1 | 87.5 |
| 154 | 14QM064H | 30.8 | 85.2 | 30.7 | 7.0 | 5.3 | 29.5 | 87.8 |
| 155 | 14QM064 | 32.3 | 85.6 | 32.2 | 7.1 | 5.1 | 29.9 | 86.4 |
| 156 | 14QM065 | 30.5 | 85.9 | 29.9 | 6.9 | 4.9 | 30.9 | 87.6 |
| 157 | 14QM065H | 30.9 | 85.7 | 32.0 | 7.0 | 4.7 | 30.6 | 87.6 |
| 158 | P♂ | 30.8 | 86.6 | 30.1 | 7.0 | 4.6 | 30.1 | 87.5 |
| 159 | 14QM066H | 31.4 | 87.4 | 29.2 | 7.0 | 5.1 | 28.8 | 87.4 |
| 160 | 14QM066 | 30.9 | 85.6 | 31.5 | 7.0 | 5.0 | 30.5 | 88.1 |
| 161 | 14QM067 | 31.1 | 86.5 | 31.0 | 7.0 | 4.7 | 30.2 | 87.0 |
| 162 | 14QM067H | 30.7 | 84.2 | 29.6 | 7.0 | 4.9 | 28.5 | 85.3 |
| 163 | P♂ | 30.8 | 85.4 | 29.9 | 6.9 | 4.8 | 30.1 | 87.5 |
| 164 | 14QM068H | 32.2 | 86.4 | 30.4 | 7.1 | 4.8 | 28.2 | 86.7 |
| 165 | 14QM068 | 30.7 | 85.2 | 29.8 | 7.0 | 5.5 | 31.1 | 86.8 |
| 166 | 14QM069 | 28.6 | 84.8 | 28.1 | 6.8 | 4.9 | 28.3 | 86.5 |
| 167 | 14QM069H | 29.9 | 85.0 | 29.6 | 6.8 | 5.2 | 28.6 | 87.9 |
| 168 | P♂ | 29.7 | 85.6 | 29.6 | 6.9 | 5.1 | 30.1 | 87.5 |
| 169 | 14QM070H | 31.2 | 84.6 | 30.8 | 7.1 | 4.9 | 30.7 | 88.1 |
| 170 | 14QM070 | 30.8 | 85.0 | 30.4 | 7.0 | 4.6 | 31.7 | 85.7 |
| 171 | 14QM071 | 30.5 | 82.3 | 29.9 | 7.0 | 4.8 | 30.1 | 86.4 |
| 172 | 14QM071H | 31.0 | 84.7 | 30.6 | 6.9 | 4.8 | 29.9 | 87.7 |
| 173 | P♂ | 30.3 | 84.9 | 31.0 | 6.9 | 4.6 | 30.1 | 87.5 |
| 174 | 14QM072H | 31.3 | 85.3 | 31.2 | 7.0 | 4.8 | 30.3 | 89.4 |
| 175 | 14QM072 | 30.6 | 84.8 | 30.0 | 6.9 | 4.9 | 29.0 | 85.3 |
| 176 | 14QM073 | 31.5 | 86.4 | 32.7 | 7.0 | 4.8 | 30.1 | 88.4 |
| 177 | 14QM073H | 31.6 | 86.4 | 30.2 | 7.0 | 5.2 | 30.1 | 88.1 |
| 178 | P♂ | 30.6 | 86.9 | 29.7 | 6.9 | 4.6 | 30.1 | 87.5 |
| 179 | 14QM074H | 32.2 | 85.0 | 31.3 | 7.0 | 4.7 | 30.3 | 87.5 |
| 180 | 14QM074 | 30.9 | 84.2 | 31.4 | 6.9 | 4.5 | 30.2 | 86.7 |
| 181 | 14QM075 | 31.4 | 86.4 | 30.0 | 7.0 | 4.5 | 31.1 | 86.1 |
| 182 | 14QM075H | 31.7 | 86.5 | 31.1 | 7.1 | 4.5 | 29.5 | 86.6 |
| 183 | P♂ | 30.8 | 86.7 | 30.3 | 6.9 | 4.2 | 30.1 | 87.5 |
| 184 | 14QM076H | 30.2 | 86.2 | 29.3 | 7.0 | 5.2 | 29.4 | 86.3 |
| 185 | 14QM076 | 29.1 | 84.9 | 28.1 | 6.9 | 5.4 | 28.2 | 84.2 |
| 186 | 14QM077 | 29.7 | 86.1 | 30.9 | 7.1 | 5.2 | 28.8 | 88.0 |
| 187 | 14QM077H | 30.4 | 86.8 | 31.9 | 7.1 | 4.7 | 30.3 | 86.1 |
| 188 | P♂ | 30.2 | 85.1 | 31.7 | 7.1 | 4.2 | 30.1 | 87.5 |
| 189 | 14QM078H | 30.5 | 85.0 | 30.7 | 7.0 | 5.2 | 30.9 | 86.5 |
| 190 | 14QM078 | 30.1 | 84.9 | 29.6 | 6.9 | 5.1 | 29.0 | 86.3 |
| 191 | 14QM079 | 30.8 | 85.7 | 29.4 | 6.9 | 4.9 | 32.1 | 85.7 |
| 192 | 14QM079H | 31.4 | 85.6 | 29.9 | 6.9 | 4.5 | 29.8 | 87.3 |
| 193 | P♂ | 29.5 | 83.5 | 28.6 | 6.9 | 4.7 | 30.1 | 87.5 |
| 194 | 14QM080H | 30.8 | 86.5 | 31.5 | 7.0 | 4.5 | 29.9 | 87.3 |
| 195 | 14QM080 | 30.2 | 86.1 | 29.4 | 6.9 | 4.5 | 28.8 | 84.1 |
| 196 | 14QM081 | 29.8 | 85.9 | 30.9 | 7.0 | 4.4 | 30.3 | 85.6 |
| 197 | 14QM081H | 31.0 | 86.4 | 31.9 | 7.1 | 4.9 | 30.1 | 87.2 |
| 198 | P♂ | 29.8 | 85.6 | 29.3 | 6.9 | 4.2 | 30.1 | 87.5 |
| 199 | 14QM082H | 30.7 | 85.6 | 29.1 | 6.9 | 4.9 | 30.0 | 85.7 |
| 200 | 14QM082 | 32.8 | 85.2 | 34.5 | 7.0 | 4.0 | 31.5 | 86.0 |
| 201 | 14QM083 | 31.9 | 85.7 | 31.0 | 7.1 | 4.7 | 30.8 | 87.1 |
| 202 | 14QM083H | 30.4 | 86.2 | 30.7 | 7.1 | 5.1 | 28.3 | 86.4 |
| 203 | 14QM380 | 29.4 | 86.4 | 28.6 | 6.9 | 4.7 | 30.8 | 87.5 |
| 204 | 14QM084H | 30.5 | 87.3 | 28.6 | 7.0 | 5.2 | 31.2 | 87.6 |
| 205 | 14QM084 | 31.1 | 84.2 | 30.8 | 7.1 | 5.2 | 29.8 | 85.2 |
| 206 | 14QM085 | 31.3 | 85.7 | 33.0 | 7.1 | 4.5 | 29.9 | 79.3 |
| 207 | 14QM085H | 31.7 | 86.7 | 31.1 | 7.1 | 5.0 | 31.1 | 88.0 |
| 208 | P♂ | 30.3 | 86.1 | 30.3 | 7.0 | 4.6 | 30.8 | 87.5 |
| 209 | 14QM086H | 30.1 | 86.8 | 30.8 | 7.0 | 4.4 | 29.0 | 86.2 |
| 210 | 14QM086 | 31.4 | 87.6 | 31.8 | 7.2 | 4.4 | 30.8 | 87.7 |
| 211 | 14QM087 | 30.8 | 85.0 | 30.9 | 6.9 | 4.4 | 30.5 | 87.1 |
| 212 | 14QM087H | 30.2 | 85.2 | 29.9 | 6.9 | 4.8 | 29.0 | 85.7 |
| 213 | P♂ | 31.4 | 85.6 | 30.7 | 7.0 | 4.3 | 30.8 | 87.5 |
| 214 | 14QM088H | 32.5 | 86.9 | 33.9 | 7.1 | 4.4 | 32.0 | 85.8 |
| 215 | 14QM088 | 32.4 | 86.1 | 32.3 | 7.1 | 4.7 | 30.3 | 86.5 |
| 216 | 14QM089 | 29.5 | 84.6 | 29.4 | 6.9 | 5.5 | 28.9 | 85.9 |
| 217 | 14QM089H | 30.4 | 85.3 | 28.2 | 6.9 | 5.1 | 29.8 | 86.2 |
| 218 | P♂ | 30.5 | 84.8 | 30.9 | 7.0 | 4.4 | 30.8 | 87.5 |
| 219 | 14QM090H | 32.1 | 84.7 | 30.8 | 7.0 | 3.8 | 33.3 | 86.5 |
| 220 | 14QM090 | 33.3 | 84.8 | 34.8 | 7.1 | 3.9 | 34.3 | 85.5 |
| 221 | 14QM091 | 31.5 | 85.3 | 29.9 | 7.0 | 4.8 | 30.0 | 85.4 |
| 222 | 14QM091H | 30.7 | 85.4 | 30.5 | 7.0 | 4.5 | 30.2 | 87.0 |
| 223 | P♂ | 30.4 | 87.6 | 31.3 | 7.0 | 4.5 | 30.8 | 87.5 |
| 224 | P ♂ | 30.0 | 87.3 | 29.4 | 6.9 | 4.9 | 29.5 | 86.0 |
| 225 | F1 | 31.3 | 85.0 | 31.3 | 7.0 | 4.7 | 30.4 | 86.1 |
| 226 | P ♀ | 30.5 | 84.9 | 29.7 | 6.8 | 4.9 | 30.6 | 85.2 |
| 227 | P♂ | 29.6 | 85.9 | 30.3 | 6.9 | 4.5 | 30.8 | 87.5 |
| 228 | 14QM092H | 32.7 | 84.7 | 32.5 | 7.1 | 4.4 | 30.9 | 85.4 |
| 229 | 14QM092 | 32.3 | 84.8 | 32.5 | 7.0 | 3.8 | 31.4 | 87.2 |
| 230 | 14QM093 | 30.6 | 85.2 | 29.1 | 6.9 | 4.8 | 31.7 | 87.9 |
| 231 | 14QM093H | 31.5 | 83.6 | 32.9 | 7.1 | 4.1 | 30.1 | 85.5 |
| 232 | P♂ | 30.4 | 84.6 | 29.8 | 6.9 | 4.7 | 30.8 | 87.5 |
| 233 | 14QM094H | 30.2 | 85.8 | 29.7 | 6.9 | 4.8 | 28.9 | 85.9 |
| 234 | 14QM094 | 30.5 | 85.1 | 28.5 | 6.9 | 4.7 | 28.5 | 86.1 |
| 235 | 14QM095 | 29.2 | 86.0 | 28.7 | 6.9 | 5.2 | 28.4 | 89.2 |
| 236 | 14QM095H | 30.1 | 86.2 | 29.7 | 6.9 | 4.8 | 30.2 | 86.2 |
| 237 | P♂ | 29.7 | 85.0 | 29.6 | 6.9 | 4.2 | 30.8 | 87.5 |
| 238 | 14QM096H | 31.1 | 84.8 | 31.1 | 7.0 | 4.7 | 30.7 | 86.5 |
| 239 | 14QM096 | 31.7 | 85.8 | 31.4 | 7.1 | 4.3 | 30.7 | 86.8 |
| 240 | 14QM097 | 30.2 | 84.7 | 29.4 | 6.9 | 4.9 | 29.6 | 86.1 |
| 241 | 14QM097H | 29.8 | 84.8 | 29.9 | 6.9 | 4.4 | 30.7 | 87.1 |
| 242 | P♂ | 30.2 | 85.5 | 29.5 | 6.9 | 4.6 | 30.8 | 87.5 |
| 243 | 14QM098H | 30.1 | 86.7 | 29.3 | 7.0 | 5.0 | 30.5 | 87.2 |
| 244 | 14QM098 | 30.7 | 86.0 | 31.3 | 7.0 | 4.5 | 30.1 | 86.4 |
| 245 | 14QM099 | 30.2 | 84.9 | 29.4 | 6.9 | 5.3 | 29.6 | 87.5 |
| 246 | 14QM099H | 30.7 | 86.3 | 31.9 | 6.9 | 4.1 | 28.6 | 86.8 |
| 247 | P♂ | 31.2 | 86.4 | 31.2 | 7.0 | 4.5 | 30.8 | 87.5 |
| 248 | 14QM100H | 31.0 | 85.5 | 28.9 | 6.8 | 4.0 | 29.7 | 88.8 |
| 249 | 14QM100 | 30.0 | 86.6 | 28.9 | 6.9 | 5.1 | 30.6 | 85.4 |
| 250 | 14QM101 | 30.1 | 87.6 | 29.7 | 6.9 | 4.8 | 29.4 | 87.2 |
| 251 | 14QM101H | 30.8 | 85.6 | 32.0 | 7.1 | 4.8 | 31.2 | 87.1 |
| 252 | 14QM380 | 30.0 | 84.6 | 27.8 | 6.8 | 4.8 | 30.5 | 86.2 |
| 253 | 14QM102H | 31.0 | 84.6 | 29.8 | 7.0 | 4.7 | 30.3 | 85.8 |
| 254 | 14QM102 | 31.1 | 86.7 | 29.9 | 7.0 | 4.6 | 29.7 | 85.7 |
| 255 | 14QM103 | 32.8 | 85.8 | 31.2 | 7.0 | 5.0 | 32.4 | 87.4 |
| 256 | 14QM103H | 32.3 | 85.8 | 30.3 | 7.0 | 4.7 | 31.4 | 87.3 |
| 257 | P♂ | 29.1 | 84.1 | 28.6 | 6.9 | 4.7 | 30.5 | 86.2 |
| 258 | 14QM104H | 32.7 | 86.8 | 31.3 | 7.0 | 5.1 | 30.2 | 86.8 |
| 259 | 14QM104 | 32.7 | 85.7 | 32.4 | 7.0 | 4.3 | 31.1 | 87.4 |
| 260 | 14QM105 | 31.2 | 83.8 | 33.4 | 7.1 | 4.5 | 30.2 | 85.7 |
| 261 | 14QM105H | 31.5 | 84.9 | 30.0 | 7.0 | 5.0 | 29.9 | 86.0 |
| 262 | P♂ | 29.6 | 84.6 | 28.8 | 6.9 | 4.9 | 30.5 | 86.2 |
| 263 | 14QM106H | 32.0 | 85.9 | 29.7 | 7.0 | 5.0 | 30.4 | 86.9 |
| 264 | 14QM106 | 32.6 | 85.7 | 31.7 | 7.2 | 5.0 | 31.6 | 88.2 |
| 265 | 14QM107 | 30.0 | 85.1 | 31.0 | 6.9 | 5.1 | 30.4 | 86.3 |
| 266 | 14QM107H | 31.8 | 85.9 | 30.3 | 7.0 | 5.4 | 29.6 | 86.8 |
| 267 | P♂ | 32.3 | 85.3 | 30.0 | 7.0 | 4.4 | 30.5 | 86.2 |
| 268 | 14QM108H | 31.1 | 86.4 | 29.4 | 7.0 | 5.3 | 30.2 | 87.3 |
| 269 | 14QM108 | 31.4 | 85.1 | 28.1 | 6.9 | 4.7 | 30.6 | 85.4 |
| 270 | 14QM109 | 29.5 | 85.3 | 29.2 | 6.9 | 4.4 | 29.1 | 84.1 |
| 271 | 14QM109H | 30.8 | 85.0 | 29.1 | 6.9 | 4.8 | 30.7 | 87.4 |
| 272 | P♂ | 31.8 | 86.1 | 30.7 | 7.0 | 4.5 | 30.5 | 86.2 |
| 273 | 14QM110H | 31.1 | 86.2 | 31.4 | 7.0 | 5.3 | 30.3 | 86.5 |
| 274 | 14QM110 | 30.5 | 85.5 | 31.4 | 6.9 | 4.9 | 29.8 | 87.6 |
| 275 | 14QM111 | 31.4 | 87.1 | 29.3 | 7.1 | 4.6 | 29.7 | 87.8 |
| 276 | 14QM111H | 31.0 | 85.7 | 30.1 | 6.9 | 4.6 | 31.3 | 87.6 |
| 277 | P♂ | 31.0 | 85.8 | 30.8 | 6.9 | 4.4 | 30.5 | 86.2 |
| 278 | 14QM112H | 31.4 | 85.6 | 29.6 | 7.0 | 5.4 | 31.2 | 86.1 |
| 279 | 14QM112 | 30.2 | 85.4 | 27.5 | 6.9 | 5.3 | 31.8 | 87.7 |
| 280 | 14QM113 | 29.8 | 83.5 | 28.8 | 6.8 | 4.9 | 31.8 | 85.9 |
| 281 | 14QM113H | 31.4 | 85.8 | 31.5 | 7.0 | 4.0 | 30.6 | 87.0 |
| 282 | P♂ | 30.2 | 86.4 | 29.8 | 7.0 | 4.3 | 30.5 | 86.2 |
| 283 | 14QM114H | 31.4 | 86.0 | 28.9 | 6.9 | 4.8 | 30.1 | 85.9 |
| 284 | 14QM114 | 32.2 | 85.8 | 31.5 | 7.1 | 5.2 | 31.6 | 86.8 |
| 285 | 14QM115 | 32.0 | 86.1 | 29.8 | 7.0 | 5.1 | 30.4 | 87.1 |
| 286 | 14QM115H | 31.9 | 86.2 | 31.9 | 7.0 | 4.7 | 30.6 | 87.0 |
| 287 | P♂ | 29.9 | 84.8 | 28.2 | 6.8 | 4.5 | 30.5 | 86.2 |
| 288 | 14QM116H | 31.6 | 87.6 | 29.7 | 7.1 | 5.1 | 30.6 | 86.7 |
| 289 | 14QM116 | 31.8 | 85.4 | 29.3 | 7.0 | 4.7 | 29.7 | 87.3 |
| 290 | 14QM117 | 29.4 | 82.9 | 27.3 | 6.8 | 5.3 | 29.5 | 85.7 |
| 291 | 14QM117H | 30.8 | 83.6 | 27.0 | 6.9 | 5.3 | 30.1 | 85.2 |
| 292 | P♂ | 29.4 | 85.0 | 27.8 | 6.8 | 4.4 | 30.5 | 86.2 |
| 293 | 14QM118H | 30.7 | 86.2 | 29.9 | 6.9 | 4.5 | 31.7 | 86.1 |
| 294 | 14QM118 | 31.4 | 86.5 | 31.9 | 7.1 | 5.1 | 28.8 | 86.7 |
| 295 | 14QM119 | 31.1 | 85.5 | 30.4 | 7.0 | 5.0 | 32.1 | 84.6 |
| 296 | 14QM119H | 30.7 | 85.9 | 30.3 | 7.0 | 4.6 | 31.1 | 85.9 |
| 297 | P♂ | 29.8 | 85.9 | 28.6 | 6.9 | 4.7 | 30.5 | 86.2 |
| 298 | 14QM120H | 31.4 | 87.5 | 29.5 | 7.0 | 5.1 | 30.5 | 86.1 |
| 299 | 14QM120 | 30.4 | 86.2 | 31.3 | 6.9 | 4.5 | 29.0 | 84.6 |
| 300 | 14QM121 | 30.5 | 84.9 | 30.5 | 6.9 | 4.4 | 30.5 | 87.4 |
| 301 | 14QM121H | 31.0 | 85.7 | 29.8 | 7.0 | 4.7 | 30.8 | 87.0 |
| 302 | 14QM380 | 29.1 | 83.9 | 28.9 | 6.9 | 4.6 | 30.2 | 87.2 |
| 303 | 14QM122H | 31.1 | 85.6 | 30.3 | 6.9 | 4.0 | 30.5 | 87.2 |
| 304 | 14QM122 | 31.8 | 87.0 | 33.1 | 7.1 | 4.2 | 31.1 | 86.8 |
| 305 | 14QM123 | 31.2 | 86.6 | 31.8 | 7.0 | 4.5 | 30.9 | 87.6 |
| 306 | 14QM123H | 30.6 | 87.1 | 31.0 | 6.9 | 4.3 | 31.3 | 88.3 |
| 307 | P♂ | 29.5 | 85.6 | 28.6 | 6.9 | 5.1 | 30.2 | 87.2 |
| 308 | 14QM124H | 30.0 | 85.3 | 31.5 | 7.0 | 5.0 | 30.4 | 87.0 |
| 309 | 14QM124 | 31.2 | 86.0 | 29.6 | 7.0 | 5.1 | 30.5 | 87.0 |
| 310 | 14QM125 | 29.5 | 84.6 | 28.2 | 6.8 | 4.8 | 28.4 | 86.8 |
| 311 | 14QM125H | 30.4 | 84.9 | 28.7 | 6.9 | 5.1 | 28.0 | 86.1 |
| 312 | P♂ | - | - | - | - | - | 30.2 | 87.2 |
| 313 | 14QM126H | 30.8 | 86.4 | 31.4 | 7.0 | 4.7 | 28.7 | 83.7 |
| 314 | 14QM126 | 30.7 | 84.5 | 30.9 | 7.0 | 5.4 | 29.8 | 86.9 |
| 315 | 14QM127 | 30.9 | 86.6 | 31.0 | 6.9 | 4.7 | 30.3 | 86.1 |
| 316 | 14QM127H | 31.3 | 85.7 | 31.0 | 6.9 | 4.3 | 30.4 | 86.3 |
| 317 | P♂ | 31.5 | 85.1 | 30.4 | 6.9 | 4.6 | 30.2 | 87.2 |
| 318 | 14QM128H | 32.1 | 85.3 | 29.5 | 7.0 | 4.5 | 31.3 | 86.3 |
| 319 | 14QM128 | 31.3 | 83.5 | 30.6 | 6.9 | 3.7 | 31.3 | 84.6 |
| 320 | 14QM129 | 31.0 | 86.0 | 30.4 | 7.0 | 4.6 | 31.1 | 86.4 |
| 321 | 14QM129H | 31.3 | 85.6 | 31.3 | 6.9 | 4.2 | 30.7 | 85.9 |
| 322 | P♂ | 30.6 | 85.4 | 29.7 | 6.9 | 4.6 | 30.2 | 87.2 |
| 323 | 14QM130H | 32.3 | 86.7 | 31.6 | 7.0 | 4.3 | 30.8 | 87.2 |
| 324 | 14QM130 | 30.9 | 84.3 | 31.6 | 7.0 | 4.2 | 30.8 | 84.9 |
| 325 | 14QM131 | 32.4 | 86.4 | 32.2 | 7.0 | 4.1 | 30.1 | 85.3 |
| 326 | 14QM131H | 32.3 | 85.6 | 31.5 | 7.0 | 5.0 | 32.1 | 85.5 |
| 327 | P♂ | 31.5 | 86.4 | 29.2 | 6.9 | 4.6 | 30.2 | 87.2 |
| 328 | 14QM132H | 29.6 | 84.3 | 27.8 | 6.9 | 5.5 | 30.2 | 84.5 |
| 329 | 14QM132 | 30.2 | 85.4 | 30.1 | 7.0 | 4.4 | 29.7 | 87.2 |
| 330 | 14QM133 | 31.1 | 86.7 | 30.3 | 6.9 | 4.6 | 30.3 | 83.8 |
| 331 | 14QM133H | 31.9 | 85.8 | 31.9 | 7.1 | 5.1 | 29.6 | 86.8 |
| 332 | P♂ | 30.0 | 85.2 | 28.1 | 6.8 | 4.5 | 30.2 | 87.2 |
| 333 | 14QM134H | 30.4 | 85.6 | 29.6 | 6.9 | 4.6 | 29.6 | 86.9 |
| 334 | 14QM134 | 30.9 | 85.2 | 29.9 | 7.0 | 4.9 | 28.9 | 86.0 |
| 335 | 14QM135 | 31.3 | 82.7 | 28.8 | 6.9 | 3.7 | 31.0 | 86.2 |
| 336 | 14QM135H | 32.5 | 85.5 | 30.6 | 7.0 | 5.0 | 30.6 | 85.8 |
| 337 | P♂ | 30.6 | 85.2 | 28.7 | 6.9 | 4.3 | 30.2 | 87.2 |
| 338 | 14QM136H | 32.2 | 86.2 | 30.7 | 7.0 | 4.7 | 31.6 | 85.8 |
| 339 | 14QM136 | 31.5 | 85.8 | 31.2 | 6.9 | 4.9 | 31.3 | 85.3 |
| 340 | 14QM137 | 31.6 | 81.7 | 34.9 | 7.0 | 3.8 | 31.2 | 84.9 |
| 341 | 14QM137H | 30.3 | 85.0 | 32.1 | 6.9 | 4.5 | 31.1 | 86.2 |
| 342 | P♂ | 32.2 | 84.4 | 35.3 | 7.1 | 4.1 | 30.2 | 87.2 |
| 343 | 14QM138H | 31.1 | 86.1 | 31.2 | 6.9 | 4.6 | 29.6 | 83.7 |
| 344 | 14QM138 | 31.4 | 86.0 | 31.0 | 6.9 | 4.3 | 29.7 | 86.0 |
| 345 | 14QM139 | 30.9 | 86.1 | 29.9 | 7.0 | 4.7 | 28.5 | 85.4 |
| 346 | 14QM139H | 31.5 | 87.4 | 31.3 | 7.0 | 4.3 | 31.4 | 85.7 |
| 347 | P♂ | 31.2 | 85.8 | 29.7 | 7.0 | 4.8 | 30.2 | 87.2 |
| 348 | 14QM140H | 31.8 | 85.8 | 31.8 | 7.0 | 4.9 | 33.1 | 86.9 |
| 349 | 14QM140 | 31.4 | 84.7 | 31.6 | 6.9 | 4.3 | 30.8 | 86.4 |
| 350 | 14QM141 | 30.3 | 86.7 | 30.2 | 7.0 | 5.2 | 30.9 | 85.8 |
| 351 | 14QM141H | 31.3 | 86.3 | 31.4 | 7.0 | 4.7 | 30.3 | 85.9 |
| 352 | 14QM380 | 31.4 | 86.9 | 30.0 | 7.0 | 4.3 | 29.3 | 87.2 |
| 353 | 14QM142H | 32.3 | 86.8 | 30.2 | 7.0 | 4.6 | 30.3 | 86.9 |
| 354 | 14QM142 | 31.5 | 86.1 | 32.1 | 7.0 | 4.1 | 31.3 | 84.7 |
| 355 | 14QM143 | 31.8 | 86.7 | 34.5 | 7.1 | 4.6 | 29.5 | 86.6 |
| 356 | 14QM143H | 31.2 | 86.4 | 33.2 | 6.9 | 3.9 | 29.6 | 86.3 |
| 357 | P♂ | 31.0 | 86.3 | 29.2 | 6.9 | 4.7 | 29.3 | 87.2 |
| 358 | 14QM144H | 31.7 | 86.5 | 31.3 | 7.0 | 4.6 | 29.7 | 85.4 |
| 359 | 14QM144 | - | - | - | - | - | 27.3 | 82.7 |
| 360 | 14QM145 | 31.0 | 86.2 | 28.4 | 6.9 | 4.4 | 29.7 | 87.4 |
| 361 | 14QM145H | 31.6 | 86.1 | 31.2 | 7.0 | 4.7 | 29.9 | 86.5 |
| 362 | P♂ | 29.9 | 85.8 | 29.1 | 6.8 | 4.4 | 29.3 | 87.2 |
| 363 | 14QM146H | 30.3 | 87.8 | 30.6 | 6.9 | 4.8 | 31.1 | 87.6 |
| 364 | 14QM146 | 29.8 | 86.7 | 29.7 | 6.9 | 4.2 | 31.5 | 86.7 |
| 365 | 14QM147 | 32.7 | 88.3 | 30.4 | 7.2 | 4.4 | 31.6 | 84.8 |
| 366 | 14QM147H | 32.6 | 86.6 | 30.4 | 7.0 | 4.4 | 32.1 | 85.3 |
| 367 | P♂ | 31.1 | 86.2 | 31.0 | 6.9 | 4.7 | 29.3 | 87.2 |
| 368 | 14QM148H | 32.2 | 87.5 | 32.2 | 7.0 | 4.1 | 31.5 | 87.5 |
| 369 | 14QM148 | 31.9 | 86.6 | 32.6 | 7.0 | 4.2 | 31.4 | 86.4 |
| 370 | 14QM149 | 31.9 | 86.7 | 32.0 | 7.0 | 4.2 | 31.9 | 86.0 |
| 371 | 14QM149H | 31.3 | 85.6 | 30.9 | 6.9 | 3.9 | 32.2 | 84.6 |
| 372 | P♂ | 30.3 | 85.4 | 30.2 | 6.9 | 4.5 | 29.3 | 87.2 |
| 373 | 14QM150H | 31.9 | 86.9 | 30.3 | 7.0 | 4.7 | 30.6 | 85.4 |
| 374 | 14QM150 | 31.8 | 87.8 | 33.5 | 7.1 | 4.6 | 30.9 | 86.3 |
| 375 | 14QM151 | 30.7 | 87.8 | 31.3 | 6.9 | 4.5 | 30.7 | 86.3 |
| 376 | 14QM151H | 31.3 | 87.4 | 33.5 | 7.0 | 4.2 | 30.3 | 86.6 |
| 377 | P♂ | 28.2 | 85.7 | 26.7 | 6.8 | 4.8 | 29.3 | 87.2 |
| 378 | 14QM152H | 31.4 | 87.0 | 31.7 | 7.0 | 4.2 | 30.4 | 85.7 |
| 379 | 14QM152 | 31.2 | 86.6 | 31.9 | 6.9 | 4.3 | 30.2 | 87.5 |
| 380 | 14QM153 | 29.9 | 87.2 | 31.0 | 6.9 | 4.7 | 30.0 | 86.8 |
| 381 | 14QM153H | 31.2 | 87.4 | 31.4 | 7.0 | 5.1 | 30.5 | 85.6 |
| 382 | P♂ | 31.3 | 86.8 | 29.7 | 6.9 | 4.5 | 29.3 | 87.2 |
| 383 | 14QM154H | 31.3 | 85.9 | 30.3 | 7.0 | 4.5 | 29.7 | 85.3 |
| 384 | 14QM154 | 31.4 | 86.6 | 29.8 | 6.9 | 4.7 | 31.9 | 85.3 |
| 385 | 14QM155 | 31.4 | 87.7 | 34.2 | 7.1 | 4.8 | 30.6 | 85.4 |
| 386 | 14QM155H | 31.0 | 86.6 | 29.9 | 6.9 | 4.8 | 29.3 | 87.6 |
| 387 | P♂ | 30.4 | 88.2 | 31.2 | 6.9 | 4.1 | 29.3 | 87.2 |
| 388 | 14QM156H | 31.6 | 86.0 | 30.3 | 7.0 | 4.7 | 31.3 | 83.5 |
| 389 | 14QM156 | 30.8 | 85.0 | 31.1 | 7.0 | 4.4 | 30.4 | 86.7 |
| 390 | 14QM157 | 32.5 | 85.0 | 30.4 | 7.0 | 4.6 | 31.0 | 86.4 |
| 391 | 14QM157H | 31.9 | 86.3 | 30.0 | 7.0 | 4.4 | 31.6 | 85.0 |
| 392 | P♂ | 31.7 | 86.7 | 31.4 | 7.0 | 4.2 | 29.3 | 87.2 |
| 393 | 14QM158H | 30.2 | 86.0 | 29.2 | 6.9 | 4.8 | 28.2 | 84.7 |
| 394 | 14QM158 | 29.2 | 86.3 | 28.4 | 6.9 | 5.1 | 24.4 | 82.7 |
| 395 | 14QM159 | 32.9 | 87.8 | 29.7 | 7.1 | 4.7 | 32.5 | 86.3 |
| 396 | 14QM159H | 32.8 | 86.9 | 31.4 | 7.2 | 4.6 | 31.0 | 86.3 |
| 397 | P♂ | 31.6 | 86.4 | 30.9 | 6.9 | 4.6 | 29.3 | 87.2 |
| 398 | 14QM160H | 31.3 | 87.3 | 31.3 | 7.0 | 4.7 | 31.2 | 85.6 |
| 399 | 14QM160 | 31.9 | 86.2 | 31.3 | 6.9 | 4.6 | 30.9 | 86.2 |
| 400 | 14QM161 | 31.3 | 85.4 | 29.4 | 6.9 | 4.6 | 29.6 | 85.9 |
| 401 | 14QM161H | 30.8 | 86.1 | 29.2 | 7.0 | 5.1 | 30.5 | 87.1 |
| 402 | 14QM380 | 29.9 | 85.7 | 29.4 | 6.9 | 4.8 | 29.7 | 87.5 |
| 403 | 14QM162H | 31.0 | 86.7 | 31.0 | 7.1 | 4.8 | 29.4 | 85.5 |
| 404 | 14QM162 | 30.3 | 87.5 | 30.2 | 7.0 | 5.5 | 28.7 | 86.4 |
| 405 | 14QM163 | 32.4 | 87.0 | 32.4 | 7.1 | 4.5 | 31.3 | 86.1 |
| 406 | 14QM163H | 31.3 | 87.9 | 32.0 | 7.0 | 4.7 | 31.3 | 85.7 |
| 407 | P♂ | 30.1 | 86.4 | 30.5 | 7.0 | 5.1 | 29.7 | 87.5 |
| 408 | 14QM164H | 31.3 | 87.5 | 33.7 | 7.1 | 4.1 | 29.4 | 85.1 |
| 409 | 14QM164 | 30.3 | 87.9 | 30.2 | 7.0 | 4.6 | 28.4 | 87.0 |
| 410 | 14QM165 | 31.4 | 84.0 | 33.5 | 6.9 | 4.2 | 29.2 | 85.0 |
| 411 | 14QM165H | 31.9 | 87.1 | 32.5 | 6.9 | 4.2 | 31.5 | 86.5 |
| 412 | P♂ | 30.7 | 87.6 | 28.7 | 7.0 | 4.8 | 29.7 | 87.5 |
| 413 | 14QM166H | 33.4 | 86.9 | 32.2 | 7.0 | 4.5 | 30.4 | 85.6 |
| 414 | 14QM166 | 33.4 | 86.8 | 32.8 | 7.0 | 4.2 | 31.4 | 87.6 |
| 415 | 14QM167 | 32.2 | 85.8 | 32.8 | 7.0 | 4.0 | 31.4 | 85.6 |
| 416 | 14QM167H | 32.4 | 86.8 | 33.5 | 7.0 | 4.3 | 31.1 | 83.5 |
| 417 | P♂ | 30.4 | 86.5 | 30.2 | 6.9 | 4.7 | 29.7 | 87.5 |
| 418 | 14QM168H | 31.3 | 86.8 | 30.7 | 7.0 | 4.6 | 29.9 | 85.0 |
| 419 | 14QM168 | 30.0 | 85.9 | 27.7 | 6.8 | 5.1 | 29.2 | 85.0 |
| 420 | 14QM169 | 30.0 | 85.3 | 29.1 | 6.9 | 5.2 | 28.7 | 86.9 |
| 421 | 14QM169H | 29.8 | 87.5 | 28.3 | 6.9 | 5.3 | 28.7 | 86.0 |
| 422 | P♂ | 30.7 | 88.0 | 30.3 | 7.0 | 4.6 | 29.7 | 87.5 |
| 423 | 14QM170H | 29.8 | 85.6 | 28.7 | 6.9 | 4.8 | 29.6 | 86.0 |
| 424 | 14QM170 | 30.1 | 86.7 | 29.2 | 7.1 | 4.9 | 29.7 | 86.2 |
| 425 | 14QM171 | 30.7 | 86.7 | 31.8 | 7.0 | 5.1 | 29.3 | 86.7 |
| 426 | 14QM171H | 30.2 | 86.4 | 29.2 | 6.9 | 4.8 | 30.2 | 85.5 |
| 427 | P♂ | 31.4 | 86.2 | 29.9 | 7.0 | 5.1 | 29.7 | 87.5 |
| 428 | 14QM172H | 31.5 | 84.4 | 31.7 | 7.0 | 4.3 | 30.7 | 84.9 |
| 429 | 14QM172 | 32.0 | 86.6 | 30.7 | 7.1 | 5.0 | 29.5 | 85.5 |
| 430 | 14QM173 | 31.3 | 85.2 | 30.8 | 7.1 | 4.8 | 31.0 | 85.3 |
| 431 | 14QM173H | 30.8 | 85.8 | 30.7 | 7.0 | 4.8 | 30.4 | 84.2 |
| 432 | P♂ | 29.9 | 86.6 | 30.2 | 7.0 | 4.4 | 29.7 | 87.5 |
| 433 | 14QM174H | 30.6 | 85.7 | 30.8 | 6.9 | 4.4 | 26.8 | 84.2 |
| 434 | 14QM174 | 30.5 | 84.9 | 31.3 | 6.9 | 4.3 | 25.2 | 84.2 |
| 435 | 14QM175 | 30.8 | 86.3 | 30.5 | 7.0 | 5.3 | 27.5 | 83.0 |
| 436 | 14QM175H | 31.6 | 86.9 | 31.3 | 7.1 | 5.0 | 31.1 | 86.3 |
| 437 | P♂ | 30.2 | 85.5 | 29.3 | 6.9 | 5.0 | 29.7 | 87.5 |
| 438 | 14QM176H | 32.4 | 85.4 | 32.9 | 7.0 | 4.5 | 31.4 | 85.0 |
| 439 | 14QM176 | 32.5 | 85.0 | 33.3 | 7.0 | 4.5 | 31.4 | 84.2 |
| 440 | 14QM177 | 29.7 | 85.7 | 29.5 | 7.1 | 5.2 | 30.1 | 85.5 |
| 441 | 14QM177H | 30.3 | 87.0 | 29.3 | 6.9 | 4.9 | 29.3 | 87.3 |
| 442 | P♂ | 31.0 | 87.1 | 29.3 | 6.9 | 5.0 | 29.7 | 87.5 |
| 443 | 14QM178H | 31.9 | 87.4 | 32.6 | 7.1 | 5.1 | 29.2 | 84.2 |
| 444 | 14QM178 | 30.7 | 85.6 | 31.2 | 7.0 | 4.5 | 31.6 | 85.6 |
| 445 | 14QM179 | 31.7 | 85.9 | 31.5 | 7.1 | 5.0 | 31.0 | 85.6 |
| 446 | 14QM179H | 31.0 | 86.1 | 29.2 | 7.0 | 5.2 | 31.5 | 86.2 |
| 447 | P♂ | 31.6 | 85.5 | 32.8 | 7.1 | 4.2 | 29.7 | 87.5 |
| 448 | P♂ | 31.9 | 85.9 | 30.9 | 6.9 | 4.4 | 29.2 | 86.7 |
| 449 | F1 | 31.4 | 85.7 | 31.8 | 7.0 | 4.3 | 31.4 | 86.2 |
| 450 | P♀ | 30.6 | 85.8 | 31.2 | 7.0 | 5.4 | 31.8 | 85.4 |
| 451 | 14QM003 | 29.7 | 85.6 | 30.7 | 7.0 | 5.2 | 29.4 | 85.2 |
| 452 | 14QM003H | 30.1 | 85.8 | 29.7 | 7.0 | 5.4 | 29.2 | 84.0 |
| 453 | 14QM380 | 30.1 | 86.2 | 28.8 | 7.0 | 5.0 | 29.7 | 86.0 |
| 454 | 14QM004H | 30.4 | 85.3 | 31.2 | 6.9 | 5.1 | 31.0 | 86.1 |
| 455 | 14QM004 | 30.2 | 85.6 | 29.3 | 6.9 | 5.0 | 30.9 | 86.6 |
| 456 | 14QM005 | 32.1 | 83.8 | 32.3 | 7.0 | 5.0 | 32.0 | 85.0 |
| 457 | 14QM005H | 30.9 | 85.9 | 31.9 | 7.0 | 5.3 | 30.6 | 85.7 |
| 458 | P♂ | 31.2 | 86.6 | 30.6 | 7.0 | 5.0 | 29.7 | 86.0 |
| 459 | 14QM006H | 31.3 | 87.2 | 31.7 | 7.1 | 5.3 | 30.3 | 85.0 |
| 460 | 14QM006 | 32.1 | 86.2 | 33.5 | 7.0 | 5.2 | 31.5 | 86.0 |
| 461 | 14QM007 | 30.9 | 86.9 | 30.1 | 7.1 | 5.1 | 29.5 | 84.3 |
| 462 | 14QM007H | 30.8 | 86.8 | 29.7 | 7.0 | 5.2 | 29.6 | 87.2 |
| 463 | P♂ | 30.0 | 84.9 | 29.3 | 6.9 | 4.5 | 29.7 | 86.0 |
| 464 | 14QM008H | 32.1 | 85.4 | 31.5 | 7.0 | 4.1 | 31.4 | 85.3 |
| 465 | 14QM008 | 30.9 | 84.9 | 31.5 | 7.0 | 4.0 | 30.6 | 85.8 |
| 466 | 14QM009 | 29.1 | 85.8 | 28.6 | 6.9 | 4.9 | 29.1 | 87.2 |
| 467 | 14QM009H | 31.7 | 85.9 | 31.4 | 7.0 | 5.0 | 29.7 | 87.8 |
| 468 | P♂ | 29.7 | 86.8 | 29.8 | 6.9 | 4.1 | 29.7 | 86.0 |
| 469 | 14QM010H | 31.2 | 84.9 | 32.9 | 7.0 | 4.3 | 31.4 | 84.9 |
| 470 | 14QM010 | 30.8 | 82.1 | 33.1 | 7.0 | 4.3 | 31.6 | 87.8 |
| 471 | 14QM011 | 31.4 | 86.4 | 32.1 | 7.0 | 5.3 | 30.0 | 86.0 |
| 472 | 14QM011H | 32.0 | 87.4 | 29.9 | 7.0 | 4.9 | 31.1 | 85.6 |
| 473 | P♂ | 31.1 | 84.0 | 30.9 | 6.9 | 4.7 | 29.7 | 86.0 |
| 474 | 14QM012H | 32.3 | 86.4 | 34.1 | 7.1 | 4.6 | 30.3 | 85.4 |
| 475 | 14QM012 | 31.3 | 84.5 | 32.9 | 6.9 | 4.3 | 30.0 | 86.6 |
| 476 | 14QM013 | 30.4 | 85.7 | 29.1 | 7.0 | 4.9 | 29.5 | 84.1 |
| 477 | 14QM013H | 30.9 | 86.1 | 30.4 | 7.0 | 4.5 | 28.8 | 87.3 |
| 478 | P♂ | 31.0 | 86.2 | 31.3 | 7.0 | 4.6 | 29.7 | 86.0 |
| 479 | 14QM014H | 30.1 | 85.4 | 29.5 | 6.9 | 4.7 | 29.7 | 87.0 |
| 480 | 14QM014 | 30.5 | 85.8 | 28.5 | 6.9 | 4.4 | 30.2 | 85.3 |
| 481 | 14QM015 | 30.7 | 86.9 | 29.3 | 7.1 | 4.7 | 29.3 | 85.1 |
| 482 | 14QM015H | 30.2 | 85.4 | 28.0 | 6.9 | 5.2 | 28.7 | 87.6 |
| 483 | P♂ | 31.3 | 86.3 | 30.8 | 7.0 | 4.8 | 29.7 | 86.0 |
| 484 | 14QM016H | 31.7 | 86.3 | 31.0 | 7.0 | 5.0 | 30.1 | 85.8 |
| 485 | 14QM016 | 31.7 | 85.2 | 32.2 | 7.0 | 4.5 | 30.1 | 86.3 |
| 486 | 14QM017 | 32.1 | 85.3 | 31.4 | 7.2 | 4.3 | 31.0 | 86.6 |
| 487 | 14QM017H | 31.1 | 85.6 | 33.0 | 7.1 | 4.0 | 30.4 | 85.6 |
| 488 | P♂ | 30.1 | 86.9 | 31.7 | 6.9 | 4.3 | 29.7 | 86.0 |
| 489 | 14QM018H | 32.1 | 85.6 | 32.5 | 7.0 | 4.0 | 31.5 | 84.9 |
| 490 | 14QM018 | 32.2 | 85.8 | 34.4 | 7.0 | 4.1 | 30.8 | 84.8 |
| 491 | 14QM019 | 31.2 | 85.5 | 33.1 | 7.1 | 4.3 | 30.0 | 85.9 |
| 492 | 14QM019H | 29.9 | 85.9 | 31.7 | 7.1 | 4.8 | 31.0 | 85.3 |
| 493 | P♂ | 30.7 | 85.9 | 30.4 | 7.0 | 5.0 | 29.7 | 86.0 |
| 494 | 14QM020H | 31.9 | 86.0 | 31.9 | 7.1 | 4.9 | 30.2 | 86.2 |
| 495 | 14QM020 | 31.1 | 84.9 | 31.4 | 6.9 | 4.2 | 30.6 | 83.2 |
| 496 | 14QM021 | 30.0 | 85.0 | 30.9 | 7.0 | 4.9 | 31.2 | 85.0 |
| 497 | 14QM021H | 29.9 | 85.4 | 29.9 | 7.0 | 4.9 | 30.2 | 86.2 |
| 498 | P♂ | - | - | - | - | - | 29.7 | 86.0 |
| 499 | 14QM022H | 31.5 | 85.9 | 31.0 | 7.0 | 4.5 | 31.3 | 86.4 |
| 500 | 14QM022 | 31.2 | 85.4 | 30.6 | 6.9 | 3.9 | 32.1 | 86.7 |
| 501 | 14QM023 | 31.8 | 85.6 | 31.2 | 7.1 | 4.2 | 31.8 | 84.6 |
| 502 | 14QM023H | 30.8 | 85.9 | 30.8 | 7.0 | 4.5 | 31.0 | 85.5 |
| 503 | 14QM380 | 30.6 | 86.4 | 30.0 | 6.9 | 4.9 | 29.4 | 85.9 |
| 504 | 14QM024H | 31.9 | 86.2 | 31.9 | 7.1 | 4.6 | 31.7 | 87.6 |
| 505 | 14QM024 | 32.2 | 86.1 | 32.8 | 7.2 | 4.4 | 32.1 | 86.3 |
| 506 | 14QM025 | - | - | - | - | - | 30.5 | 87.2 |
| 507 | 14QM025H | 31.5 | 87.2 | 33.1 | 7.0 | 4.3 | 30.4 | 86.7 |
| 508 | P♂ | 30.5 | 86.2 | 30.2 | 7.0 | 4.7 | 29.4 | 85.9 |
| 509 | 14QM026H | 30.5 | 88.2 | 28.6 | 7.0 | 5.1 | 30.8 | 87.7 |
| 510 | 14QM026 | 31.5 | 87.1 | 30.1 | 7.1 | 4.9 | 31.2 | 86.5 |
| 511 | 14QM027 | 30.4 | 86.3 | 30.1 | 7.1 | 4.7 | 30.2 | 87.0 |
| 512 | 14QM027H | 30.8 | 88.4 | 29.7 | 7.2 | 5.2 | 29.5 | 88.4 |
| 513 | P♂ | 29.9 | 87.0 | 29.6 | 6.9 | 4.4 | 29.4 | 85.9 |
| 514 | 14QM028H | 31.4 | 86.8 | 30.3 | 7.0 | 5.0 | 29.4 | 86.6 |
| 515 | 14QM028 | 30.6 | 86.6 | 31.9 | 7.1 | 4.6 | 29.7 | 88.9 |
| 516 | 14QM029 | 31.3 | 86.7 | 30.4 | 7.2 | 4.6 | 32.3 | 86.5 |
| 517 | 14QM029H | 31.5 | 86.5 | 30.3 | 7.0 | 4.7 | 31.0 | 86.5 |
| 518 | P♂ | 30.2 | 86.4 | 29.1 | 7.0 | 4.6 | 29.4 | 85.9 |
| 519 | 14QM030H | 31.8 | 85.6 | 31.9 | 7.1 | 4.1 | 31.3 | 85.4 |
| 520 | 14QM030 | 31.8 | 86.4 | 31.0 | 7.1 | 4.1 | 30.8 | 85.8 |
| 521 | 14QM031 | 32.2 | 85.6 | 31.2 | 7.3 | 4.5 | 31.7 | 86.3 |
| 522 | 14QM031H | 32.0 | 86.4 | 31.7 | 7.1 | 4.1 | 31.2 | 85.6 |
| 523 | P♂ | 31.7 | 86.7 | 30.8 | 6.9 | 4.7 | 29.4 | 85.9 |
| 524 | 14QM032H | 31.3 | 86.5 | 33.0 | 7.0 | 4.3 | 30.2 | 86.4 |
| 525 | 14QM032 | 31.4 | 86.0 | 31.6 | 7.1 | 5.0 | 30.6 | 85.7 |
| 526 | 14QM033 | 30.4 | 86.7 | 31.0 | 7.0 | 5.6 | 28.1 | 88.0 |
| 527 | 14QM033H | 30.6 | 87.2 | 30.3 | 7.1 | 5.4 | 29.4 | 85.6 |
| 528 | P♂ | 29.8 | 81.8 | 29.1 | 7.0 | 5.2 | 29.4 | 85.9 |
| 529 | 14QM034H | 31.4 | 86.0 | 30.5 | 7.1 | 5.3 | 30.0 | 87.6 |
| 530 | 14QM034 | 31.1 | 86.7 | 33.0 | 7.1 | 5.1 | 31.1 | 87.2 |
| 531 | 14QM035 | 31.6 | 85.9 | 31.9 | 6.9 | 4.5 | 29.6 | 85.8 |
| 532 | 14QM035H | 31.7 | 85.2 | 31.5 | 7.0 | 4.5 | 31.4 | 86.5 |
| 533 | P♂ | 30.6 | 86.6 | 30.1 | 7.0 | 4.7 | 29.4 | 85.9 |
| 534 | 14QM036H | 31.3 | 85.8 | 30.4 | 7.0 | 5.0 | 29.8 | 87.3 |
| 535 | 14QM036 | 31.8 | 85.1 | 30.6 | 7.1 | 4.4 | 29.4 | 86.4 |
| 536 | 14QM037 | 30.3 | 86.1 | 29.4 | 6.9 | 5.2 | 31.0 | 87.2 |
| 537 | 14QM037H | 31.3 | 86.7 | 30.2 | 6.9 | 5.2 | 30.2 | 86.9 |
| 538 | P♂ | 30.6 | 85.7 | 29.8 | 6.9 | 5.0 | 29.4 | 85.9 |
| 539 | 14QM038H | 31.1 | 85.8 | 29.6 | 7.0 | 5.1 | 30.1 | 87.7 |
| 540 | 14QM038 | 29.4 | 85.8 | 27.7 | 6.9 | 4.6 | 28.2 | 85.8 |
| 541 | 14QM039 | 30.1 | 87.3 | 30.6 | 7.0 | 5.1 | 29.4 | 86.5 |
| 542 | 14QM039H | 31.0 | 84.9 | 31.3 | 7.0 | 4.9 | 30.2 | 85.9 |
| 543 | P♂ | 30.7 | 84.7 | 29.9 | 6.9 | 4.9 | 29.4 | 85.9 |
| 544 | 14QM040H | 30.1 | 84.7 | 30.5 | 7.0 | 5.1 | 29.5 | 85.8 |
| 545 | 14QM040 | 29.8 | 85.6 | 28.7 | 6.9 | 5.1 | 29.0 | 86.0 |
| 546 | 14QM041 | 30.6 | 84.6 | 29.3 | 6.9 | 4.8 | 31.4 | 86.6 |
| 547 | 14QM041H | 30.6 | 85.3 | 31.4 | 7.0 | 4.4 | 30.1 | 86.0 |
| 548 | P♂ | 28.8 | 87.2 | 28.2 | 6.9 | 5.0 | 29.4 | 85.9 |
| 549 | 14QM042H | 30.0 | 86.8 | 28.4 | 7.0 | 5.2 | 28.3 | 85.6 |
| 550 | 14QM042 | 30.0 | 85.8 | 28.7 | 7.0 | 5.1 | 27.1 | 84.2 |
| 551 | 14QM043 | 29.6 | 85.4 | 28.0 | 6.9 | 5.5 | 28.0 | 85.2 |
| 552 | 14QM043H | 30.1 | 85.5 | 28.2 | 6.9 | 5.3 | 30.3 | 85.8 |
| 553 | 14QM380 | 30.3 | 85.8 | 29.1 | 6.8 | 5.0 | 28.4 | 83.5 |
| 554 | 14QM044H | 31.3 | 85.8 | 30.5 | 7.0 | 4.8 | 29.2 | 86.4 |
| 555 | 14QM044 | 30.4 | 85.8 | 29.6 | 7.0 | 5.0 | 30.3 | 86.5 |
| 556 | 14QM045 | 31.3 | 87.3 | 31.6 | 6.9 | 4.6 | 30.8 | 85.8 |
| 557 | 14QM045H | 31.6 | 85.9 | 29.2 | 7.0 | 5.1 | 31.0 | 86.4 |
| 558 | P♂ | 30.0 | 84.3 | 30.6 | 6.9 | 5.0 | 28.4 | 83.5 |
| 559 | 14QM046H | 30.1 | 87.2 | 30.4 | 7.0 | 4.9 | 30.4 | 86.7 |
| 560 | 14QM046 | 29.8 | 86.3 | 28.3 | 6.9 | 5.0 | 29.5 | 87.0 |
| 561 | 14QM047 | 31.1 | 85.5 | 31.1 | 7.0 | 5.6 | 31.1 | 86.0 |
| 562 | 14QM047H | 31.9 | 85.5 | 32.3 | 7.2 | 5.1 | 30.9 | 86.8 |
| 563 | P♂ | 29.0 | 85.5 | 28.0 | 6.8 | 4.6 | 28.4 | 83.5 |
| 564 | 14QM048H | 29.3 | 85.8 | 29.2 | 7.0 | 5.1 | 30.4 | 85.7 |
| 565 | 14QM048 | 30.6 | 86.4 | 30.8 | 7.0 | 4.8 | 28.9 | 86.5 |
| 566 | 14QM049 | 30.6 | 86.4 | 30.7 | 7.0 | 4.7 | 29.1 | 87.3 |
| 567 | 14QM049H | 30.4 | 85.6 | 30.1 | 6.9 | 4.9 | 29.2 | 87.6 |
| 568 | P♂ | - | - | - | - | - | 28.4 | 83.5 |
| 569 | 14QM050H | 31.2 | 86.0 | 30.8 | 7.0 | 5.1 | 30.0 | 86.7 |
| 570 | 14QM050 | 33.5 | 85.1 | 32.4 | 7.1 | 5.1 | 31.5 | 85.4 |
| 571 | 14QM051 | 31.8 | 85.3 | 30.0 | 6.9 | 4.8 | 31.5 | 87.8 |
| 572 | 14QM051H | 31.8 | 85.9 | 30.9 | 7.1 | 5.0 | 32.0 | 85.6 |
| 573 | P♂ | 31.3 | 86.5 | 31.1 | 7.0 | 4.9 | 28.4 | 83.5 |
| 574 | 14QM052H | 31.5 | 86.4 | 31.1 | 7.0 | 4.7 | 31.0 | 86.4 |
| 575 | 14QM052 | 30.7 | 85.1 | 29.6 | 6.9 | 4.5 | 31.1 | 85.8 |
| 576 | 14QM053 | 33.1 | 85.3 | 32.0 | 7.1 | 3.7 | 33.1 | 85.8 |
| 577 | 14QM053H | 30.3 | 84.5 | 30.0 | 6.9 | 4.2 | 31.6 | 85.6 |
| 578 | P♂ | 30.4 | 85.4 | 29.1 | 6.9 | 4.6 | 28.4 | 83.5 |
| 579 | 14QM054H | 30.1 | 86.7 | 28.8 | 6.9 | 4.7 | 30.0 | 86.2 |
| 580 | 14QM054 | 29.7 | 84.7 | 29.0 | 7.0 | 4.9 | 28.2 | 85.4 |
| 581 | 14QM055 | 29.8 | 86.4 | 30.6 | 6.9 | 5.4 | 28.5 | 86.8 |
| 582 | 14QM055H | 30.9 | 86.4 | 29.8 | 7.0 | 5.1 | 29.1 | 86.9 |
| 583 | P♂ | 31.0 | 85.2 | 29.8 | 7.0 | 4.5 | 28.4 | 83.5 |
| 584 | 14QM056H | 32.7 | 85.4 | 34.1 | 7.1 | 4.4 | 30.9 | 85.6 |
| 585 | 14QM056 | 32.5 | 84.9 | 31.6 | 7.0 | 4.6 | 31.1 | 85.0 |
| 586 | 14QM057 | 30.0 | 86.8 | 30.8 | 7.0 | 3.5 | 27.8 | 84.8 |
| 587 | 14QM057H | 29.3 | 83.7 | 27.4 | 6.8 | 4.4 | 29.5 | 86.5 |
| 588 | P♂ | 30.7 | 86.3 | 29.5 | 6.9 | 4.8 | 28.4 | 83.5 |
| 589 | 14QM058H | 29.8 | 85.1 | 29.8 | 6.9 | 5.1 | 30.2 | 86.1 |
| 590 | 14QM058 | 31.3 | 86.3 | 30.8 | 7.0 | 4.9 | 31.5 | 83.2 |
| 591 | 14QM059 | 30.4 | 80.4 | 30.1 | 7.0 | 4.9 | 31.3 | 85.6 |
| 592 | 14QM059H | 29.7 | 86.4 | 29.5 | 7.0 | 4.8 | 28.4 | 85.8 |
| 593 | P♂ | 30.1 | 85.9 | 28.5 | 6.9 | 4.6 | 28.4 | 83.5 |
| 594 | 14QM060H | 32.1 | 85.7 | 31.9 | 7.1 | 4.7 | 30.3 | 86.6 |
| 595 | 14QM060 | 32.1 | 85.7 | 30.5 | 7.0 | 4.9 | 30.8 | 84.5 |
| 596 | 14QM061 | 29.6 | 84.7 | 29.5 | 6.9 | 4.4 | 30.4 | 84.7 |
| 597 | 14QM061H | 30.4 | 85.7 | 30.6 | 7.1 | 4.9 | 29.9 | 84.9 |
| 598 | P♂ | 30.0 | 85.5 | 29.3 | 6.9 | 4.2 | 28.4 | 83.5 |
| 599 | 14QM062H | 31.1 | 84.5 | 31.2 | 7.0 | 4.2 | 31.2 | 87.3 |
| 600 | 14QM062 | 33.2 | 86.8 | 33.0 | 7.2 | 4.2 | 31.0 | 84.9 |
| 601 | 14QM063 | 28.7 | 85.0 | 30.1 | 6.9 | 4.5 | 29.2 | 86.2 |
| 602 | 14QM063H | 30.9 | 85.6 | 31.5 | 7.0 | 4.5 | 30.2 | 85.6 |
| 603 | 14QM380 | 30.1 | 84.9 | 31.8 | 7.0 | 4.5 | 31.0 | 86.2 |
| 604 | 14QM064H | 31.3 | 86.3 | 29.3 | 6.9 | 5.0 | 30.1 | 86.9 |
| 605 | 14QM064 | 31.3 | 85.4 | 31.0 | 7.0 | 5.1 | 30.6 | 87.2 |
| 606 | 14QM065 | 31.8 | 85.7 | 31.1 | 7.1 | 5.0 | 30.0 | 85.8 |
| 607 | 14QM065H | 30.7 | 85.7 | 30.2 | 6.9 | 4.3 | 30.1 | 86.2 |
| 608 | P♂ | 30.6 | 86.8 | 29.6 | 7.0 | 4.6 | 31.0 | 86.2 |
| 609 | 14QM066H | 31.7 | 87.1 | 31.0 | 7.1 | 4.8 | 31.6 | 85.3 |
| 610 | 14QM066 | 31.7 | 85.3 | 31.2 | 7.0 | 4.5 | 31.3 | 85.5 |
| 611 | 14QM067 | 30.5 | 86.0 | 29.8 | 6.9 | 4.4 | 30.7 | 86.2 |
| 612 | 14QM067H | 30.8 | 86.2 | 30.2 | 7.0 | 4.4 | 29.7 | 86.8 |
| 613 | P♂ | 30.7 | 86.0 | 28.9 | 6.9 | 5.2 | 31.0 | 86.2 |
| 614 | 14QM068H | 31.2 | 85.9 | 32.3 | 7.1 | 4.8 | 30.1 | 86.3 |
| 615 | 14QM068 | 30.1 | 87.6 | 29.0 | 7.0 | 5.2 | 30.2 | 86.6 |
| 616 | 14QM069 | 29.4 | 85.4 | 29.1 | 6.9 | 5.1 | 27.3 | 86.7 |
| 617 | 14QM069H | 30.5 | 87.8 | 29.9 | 7.0 | 5.0 | 29.5 | 86.0 |
| 618 | P♂ | 29.9 | 87.0 | 29.9 | 7.0 | 4.6 | 31.0 | 86.2 |
| 619 | 14QM070H | 31.6 | 86.4 | 31.7 | 6.9 | 4.6 | 31.6 | 85.9 |
| 620 | 14QM070 | 31.1 | 86.9 | 30.8 | 7.0 | 4.7 | 32.0 | 86.6 |
| 621 | 14QM071 | 31.1 | 85.8 | 29.8 | 6.9 | 4.5 | 30.0 | 84.8 |
| 622 | 14QM071H | 31.0 | 87.7 | 32.0 | 7.0 | 4.3 | 30.0 | 85.1 |
| 623 | P♂ | 29.4 | 85.3 | 29.4 | 6.9 | 4.5 | 31.0 | 86.2 |
| 624 | 14QM072H | 30.5 | 86.1 | 29.4 | 6.9 | 4.8 | 30.2 | 86.8 |
| 625 | 14QM072 | 29.6 | 85.9 | 28.6 | 6.8 | 5.0 | 29.8 | 85.7 |
| 626 | 14QM073 | 31.1 | 87.1 | 31.6 | 6.9 | 4.5 | 30.6 | 86.0 |
| 627 | 14QM073H | 31.6 | 86.7 | 30.2 | 6.9 | 4.6 | 29.7 | 86.5 |
| 628 | P♂ | 31.3 | 85.9 | 28.7 | 7.0 | 4.5 | 31.0 | 86.2 |
| 629 | 14QM074H | 31.3 | 84.5 | 32.2 | 7.0 | 5.0 | 30.5 | 86.2 |
| 630 | 14QM074 | 30.9 | 86.8 | 31.2 | 6.9 | 4.5 | 30.6 | 85.0 |
| 631 | 14QM075 | 30.4 | 85.3 | 29.9 | 6.9 | 4.0 | 31.8 | 85.5 |
| 632 | 14QM075H | 31.5 | 87.2 | 30.0 | 7.0 | 4.3 | 29.9 | 86.0 |
| 633 | P♂ | 31.6 | 85.7 | 30.2 | 7.0 | 5.2 | 31.0 | 86.2 |
| 634 | 14QM076H | 31.6 | 87.1 | 31.7 | 7.2 | 5.1 | 29.0 | 84.2 |
| 635 | 14QM076 | 30.5 | 85.0 | 27.6 | 7.0 | 5.5 | 30.8 | 86.1 |
| 636 | 14QM077 | 29.1 | 87.0 | 29.2 | 7.0 | 5.3 | 28.9 | 87.1 |
| 637 | 14QM077H | 29.6 | 87.1 | 28.1 | 6.8 | 4.8 | 30.5 | 86.1 |
| 638 | P♂ | 31.3 | 88.2 | 31.4 | 7.0 | 4.8 | 31.0 | 86.2 |
| 639 | 14QM078H | 30.5 | 85.9 | 28.1 | 7.0 | 5.3 | 31.0 | 85.2 |
| 640 | 14QM078 | 29.2 | 84.5 | 26.7 | 6.9 | 5.0 | 29.1 | 85.5 |
| 641 | 14QM079 | 30.3 | 86.3 | 31.5 | 6.9 | 5.1 | 30.9 | 86.3 |
| 642 | 14QM079H | 31.8 | 85.7 | 30.8 | 6.9 | 5.0 | 31.2 | 85.2 |
| 643 | P♂ | 30.1 | 87.0 | 28.5 | 6.9 | 5.3 | 31.0 | 86.2 |
| 644 | 14QM080H | 30.3 | 86.3 | 30.5 | 7.0 | 4.6 | 30.0 | 87.5 |
| 645 | 14QM080 | 30.4 | 86.7 | 28.6 | 6.9 | 4.9 | 29.2 | 85.8 |
| 646 | 14QM081 | 30.0 | 84.7 | 29.7 | 6.9 | 4.6 | 30.1 | 86.5 |
| 647 | 14QM081H | 31.1 | 86.2 | 31.6 | 7.0 | 4.2 | 30.6 | 86.2 |
| 648 | P♂ | 29.7 | 83.6 | 29.1 | 6.9 | 4.7 | 31.0 | 86.2 |
| 649 | 14QM082H | 30.8 | 86.7 | 31.0 | 6.9 | 4.3 | 31.3 | 86.0 |
| 650 | 14QM082 | 32.6 | 86.1 | 34.1 | 7.1 | 4.5 | 31.3 | 84.6 |
| 651 | 14QM083 | 32.1 | 87.8 | 29.6 | 7.0 | 4.8 | 30.4 | 86.1 |
| 652 | 14QM083H | 31.5 | 86.3 | 31.0 | 7.0 | 4.7 | 30.2 | 87.2 |
| 653 | 14QM380 | 30.9 | 86.3 | 29.5 | 7.0 | 4.9 | 30.5 | 86.8 |
| 654 | 14QM084H | 30.6 | 87.5 | 29.2 | 6.9 | 5.3 | 31.2 | 86.5 |
| 655 | 14QM084 | 31.4 | 87.4 | 29.1 | 7.0 | 5.0 | 29.9 | 87.7 |
| 656 | 14QM085 | 31.7 | 87.6 | 31.8 | 7.1 | 4.9 | 31.1 | 86.4 |
| 657 | 14QM085H | 31.5 | 86.8 | 29.8 | 7.0 | 4.7 | 29.5 | 85.1 |
| 658 | P♂ | 30.3 | 87.4 | 27.4 | 6.9 | 5.3 | 30.5 | 86.8 |
| 659 | 14QM086H | 30.8 | 86.7 | 29.1 | 7.0 | 5.1 | 31.0 | 86.0 |
| 660 | 14QM086 | 29.6 | 89.1 | 30.6 | 7.1 | 5.5 | 29.0 | 86.9 |
| 661 | 14QM087 | 31.7 | 86.5 | 31.8 | 7.0 | 5.0 | 31.0 | 86.5 |
| 662 | 14QM087H | 31.8 | 86.5 | 30.7 | 7.1 | 5.4 | 29.7 | 87.3 |
| 663 | P♂ | 31.2 | 87.9 | 29.7 | 6.9 | 4.9 | 30.5 | 86.8 |
| 664 | 14QM088H | 30.9 | 86.6 | 28.8 | 6.9 | 5.0 | 29.9 | 86.5 |
| 665 | 14QM088 | 31.3 | 86.9 | 33.1 | 6.9 | 3.8 | 29.0 | 84.2 |
| 666 | 14QM089 | 31.0 | 86.6 | 30.0 | 7.0 | 5.2 | 29.6 | 86.4 |
| 667 | 14QM089H | 31.4 | 86.0 | 29.4 | 7.0 | 4.8 | 30.5 | 85.8 |
| 668 | P♂ | 30.0 | 86.2 | 28.6 | 6.9 | 4.9 | 30.5 | 86.8 |
| 669 | 14QM090H | 33.0 | 85.8 | 30.6 | 7.0 | 4.5 | 32.6 | 84.9 |
| 670 | 14QM090 | 33.1 | 87.3 | 33.4 | 7.0 | 4.7 | 33.2 | 86.5 |
| 671 | 14QM091 | 30.3 | 86.4 | 29.8 | 7.0 | 4.8 | 28.0 | 84.0 |
| 672 | 14QM091H | 31.6 | 88.0 | 31.7 | 7.1 | 4.4 | 30.3 | 84.2 |
| 673 | P♂ | 29.7 | 87.0 | 29.4 | 7.0 | 4.4 | 30.5 | 86.8 |
| 674 | P ♂ | 31.0 | 84.6 | 30.5 | 6.9 | 4.2 | 30.3 | 86.5 |
| 675 | F1 | 29.9 | 87.6 | 29.1 | 6.9 | 4.7 | 31.0 | 85.8 |
| 676 | P ♀ | 30.2 | 86.1 | 29.6 | 6.9 | 5.1 | 29.0 | 84.0 |
| 677 | P♂ | 30.2 | 88.4 | 29.0 | 6.9 | 4.8 | 30.5 | 86.8 |
| 678 | 14QM092H | 31.5 | 85.6 | 30.7 | 7.0 | 4.9 | 31.6 | 85.7 |
| 679 | 14QM092 | 31.9 | 85.9 | 31.3 | 7.0 | 4.0 | 32.2 | 86.7 |
| 680 | 14QM093 | 31.2 | 86.2 | 29.5 | 7.0 | 4.8 | 30.5 | 85.7 |
| 681 | 14QM093H | 31.9 | 85.3 | 30.8 | 7.0 | 4.9 | 30.0 | 85.2 |
| 682 | P♂ | 29.7 | 85.4 | 29.0 | 6.9 | 4.7 | 30.5 | 86.8 |
| 683 | 14QM094H | 31.1 | 85.9 | 29.4 | 6.9 | 4.8 | 30.0 | 87.0 |
| 684 | 14QM094 | 29.3 | 85.1 | 29.2 | 6.9 | 4.9 | 29.4 | 86.3 |
| 685 | 14QM095 | 30.8 | 87.2 | 33.8 | 7.1 | 4.5 | 29.7 | 86.4 |
| 686 | 14QM095H | 29.7 | 87.3 | 28.8 | 6.9 | 4.6 | 28.9 | 86.4 |
| 687 | P♂ | 30.4 | 86.5 | 29.1 | 6.9 | 4.5 | 30.5 | 86.8 |
| 688 | 14QM096H | 32.1 | 87.1 | 30.9 | 7.1 | 4.7 | 30.9 | 85.2 |
| 689 | 14QM096 | 32.7 | 86.2 | 32.1 | 7.1 | 4.7 | 31.1 | 86.5 |
| 690 | 14QM097 | 30.0 | 84.5 | 29.5 | 7.0 | 4.5 | 28.9 | 86.0 |
| 691 | 14QM097H | 31.1 | 86.1 | 31.1 | 7.0 | 4.4 | 29.5 | 85.9 |
| 692 | P♂ | 30.4 | 86.2 | 30.9 | 6.9 | 4.7 | 30.5 | 86.8 |
| 693 | 14QM098H | 31.6 | 86.3 | 29.3 | 7.0 | 4.1 | 29.7 | 86.5 |
| 694 | 14QM098 | 30.6 | 86.5 | 28.6 | 6.9 | 4.8 | 28.6 | 85.5 |
| 695 | 14QM099 | 30.9 | 86.1 | 31.8 | 6.9 | 4.2 | 29.5 | 84.9 |
| 696 | 14QM099H | 30.8 | 84.0 | 30.9 | 7.0 | 4.5 | 28.3 | 85.5 |
| 697 | P♂ | 31.7 | 85.3 | 29.8 | 7.0 | 4.4 | 30.5 | 86.8 |
| 698 | 14QM100H | 31.6 | 88.1 | 31.0 | 7.1 | 4.8 | 29.7 | 85.9 |
| 699 | 14QM100 | 29.4 | 84.8 | 27.4 | 6.8 | 4.7 | 29.2 | 85.3 |
| 700 | 14QM101 | 29.6 | 86.6 | 29.6 | 6.9 | 4.6 | 29.9 | 86.7 |
| 701 | 14QM101H | 30.6 | 87.1 | 29.0 | 6.9 | 4.8 | 29.7 | 87.9 |
| 702 | 14QM380 | 31.4 | 85.3 | 31.9 | 7.0 | 4.5 | 29.2 | 86.9 |
| 703 | 14QM102H | 31.3 | 85.8 | 30.8 | 7.1 | 5.0 | 31.4 | 86.0 |
| 704 | 14QM102 | 32.5 | 84.9 | 31.3 | 7.0 | 4.6 | 30.6 | 84.6 |
| 705 | 14QM103 | 32.9 | 85.2 | 33.2 | 7.2 | 4.4 | 32.6 | 86.6 |
| 706 | 14QM103H | 31.5 | 84.4 | 30.9 | 7.0 | 4.8 | 31.7 | 85.1 |
| 707 | P♂ | 29.7 | 85.4 | 28.0 | 6.9 | 4.4 | 29.2 | 86.9 |
| 708 | 14QM104H | 31.4 | 86.9 | 29.9 | 7.1 | 4.7 | 30.6 | 86.0 |
| 709 | 14QM104 | 31.2 | 86.7 | 31.0 | 7.1 | 4.8 | 31.1 | 84.9 |
| 710 | 14QM105 | 31.6 | 84.4 | 31.2 | 7.1 | 4.7 | 31.4 | 84.7 |
| 711 | 14QM105H | 31.6 | 86.6 | 29.8 | 7.0 | 4.7 | 29.5 | 86.7 |
| 712 | P♂ | 31.1 | 84.7 | 30.9 | 7.0 | 4.3 | 29.2 | 86.9 |
| 713 | 14QM106H | - | - | - | - | - | 31.1 | 86.6 |
| 714 | 14QM106 | 32.5 | 85.5 | 30.4 | 7.1 | 4.6 | 30.7 | 86.0 |
| 715 | 14QM107 | 30.9 | 85.7 | 31.6 | 7.0 | 5.0 | 28.9 | 82.7 |
| 716 | 14QM107H | 31.2 | 86.2 | 30.8 | 7.0 | 5.2 | 30.8 | 86.2 |
| 717 | P♂ | 29.7 | 85.9 | 28.5 | 7.0 | 5.1 | 29.2 | 86.9 |
| 718 | 14QM108H | 30.5 | 86.4 | 28.9 | 7.0 | 5.3 | 30.0 | 84.3 |
| 719 | 14QM108 | 32.0 | 85.8 | 28.5 | 7.0 | 5.0 | 30.6 | 84.9 |
| 720 | 14QM109 | 29.5 | 85.9 | 28.2 | 6.9 | 4.7 | 30.7 | 85.5 |
| 721 | 14QM109H | 30.1 | 84.3 | 28.8 | 6.8 | 4.1 | 29.4 | 86.9 |
| 722 | P♂ | 29.0 | 85.6 | 27.7 | 6.9 | 5.1 | 29.2 | 86.9 |
| 723 | 14QM110H | 30.9 | 86.6 | 31.4 | 6.9 | 4.7 | 30.0 | 85.9 |
| 724 | 14QM110 | 30.2 | 86.8 | 30.9 | 7.0 | 5.2 | 30.1 | 85.8 |
| 725 | 14QM111 | 31.5 | 86.4 | 32.2 | 7.0 | 4.0 | 30.6 | 86.4 |
| 726 | 14QM111H | 32.0 | 86.3 | 30.4 | 7.0 | 4.9 | 31.1 | 85.8 |
| 727 | P♂ | 31.4 | 85.9 | 32.2 | 7.0 | 4.0 | 29.2 | 86.9 |
| 728 | 14QM112H | 30.6 | 87.3 | 31.2 | 7.0 | 4.8 | 31.4 | 85.9 |
| 729 | 14QM112 | 29.6 | 86.2 | 27.9 | 6.9 | 5.0 | 30.8 | 85.7 |
| 730 | 14QM113 | - | - | - | - | - | 31.1 | 85.5 |
| 731 | 14QM113H | 32.1 | 86.7 | 30.0 | 7.0 | 5.3 | 31.6 | 85.5 |
| 732 | P♂ | 30.5 | 87.0 | 28.6 | 6.9 | 4.6 | 29.2 | 86.9 |
| 733 | 14QM114H | 32.6 | 86.4 | 29.7 | 7.0 | 5.2 | 29.7 | 85.6 |
| 734 | 14QM114 | 31.5 | 86.9 | 30.5 | 7.0 | 5.0 | 31.2 | 86.6 |
| 735 | 14QM115 | 31.9 | 87.0 | 33.0 | 7.0 | 4.2 | 31.4 | 84.4 |
| 736 | 14QM115H | 30.6 | 84.7 | 30.1 | 7.0 | 5.3 | 31.9 | 86.2 |
| 737 | P♂ | 29.9 | 86.6 | 27.6 | 6.9 | 4.6 | 29.2 | 86.9 |
| 738 | 14QM116H | 30.2 | 85.4 | 29.4 | 7.0 | 4.9 | 30.3 | 85.3 |
| 739 | 14QM116 | 30.7 | 86.1 | 29.5 | 6.9 | 4.9 | 28.6 | 85.7 |
| 740 | 14QM117 | 31.8 | 85.9 | 30.4 | 7.0 | 4.9 | 31.3 | 85.3 |
| 741 | 14QM117H | 31.9 | 84.6 | 31.6 | 7.0 | 4.4 | 31.1 | 82.3 |
| 742 | P♂ | 29.8 | 86.1 | 27.9 | 6.8 | 4.6 | 29.2 | 86.9 |
| 743 | 14QM118H | 31.3 | 87.1 | 31.7 | 7.1 | 4.6 | 32.4 | 85.3 |
| 744 | 14QM118 | 31.7 | 84.9 | 30.8 | 7.0 | 4.9 | 30.5 | 86.4 |
| 745 | 14QM119 | 32.4 | 85.0 | 32.3 | 7.0 | 4.5 | 31.3 | 84.7 |
| 746 | 14QM119H | 31.5 | 86.2 | 30.6 | 7.0 | 4.8 | 31.3 | 86.5 |
| 747 | P♂ | 30.4 | 86.1 | 28.6 | 6.9 | 5.0 | 29.2 | 86.9 |
| 748 | 14QM120H | 31.0 | 85.5 | 32.3 | 7.0 | 4.5 | 30.4 | 86.6 |
| 749 | 14QM120 | 30.3 | 85.8 | 30.7 | 6.9 | 4.1 | 29.9 | 87.0 |
| 750 | 14QM121 | 31.3 | 85.3 | 30.9 | 7.0 | 5.0 | 30.6 | 86.2 |
| 751 | 14QM121H | 32.1 | 86.8 | 30.6 | 7.0 | 5.2 | 30.5 | 86.5 |
| 752 | 14QM380 | 30.2 | 86.2 | 28.9 | 6.9 | 4.9 | 30.1 | 85.2 |
| 753 | 14QM122H | 32.1 | 86.0 | 30.8 | 7.0 | 5.2 | 30.5 | 86.0 |
| 754 | 14QM122 | 32.2 | 86.7 | 31.2 | 7.2 | 5.3 | 31.3 | 85.5 |
| 755 | 14QM123 | 31.8 | 86.8 | 31.7 | 6.9 | 4.5 | 29.3 | 86.3 |
| 756 | 14QM123H | 31.2 | 85.8 | 28.8 | 7.0 | 5.4 | 30.2 | 86.5 |
| 757 | P♂ | 30.1 | 85.2 | 28.7 | 6.9 | 5.7 | 30.1 | 85.2 |
| 758 | 14QM124H | 30.8 | 86.1 | 29.3 | 6.9 | 5.3 | 29.9 | 85.7 |
| 759 | 14QM124 | 29.8 | 83.7 | 29.5 | 7.0 | 5.4 | 30.3 | 84.1 |
| 760 | 14QM125 | 29.8 | 86.4 | 28.2 | 6.9 | 5.3 | 28.6 | 82.9 |
| 761 | 14QM125H | 29.4 | 85.9 | 29.1 | 6.9 | 5.1 | 29.9 | 87.0 |
| 762 | P♂ | 29.3 | 86.4 | 28.1 | 6.9 | 5.1 | 30.1 | 85.2 |
| 763 | 14QM126H | 29.2 | 84.3 | 28.9 | 7.0 | 5.3 | 29.3 | 86.3 |
| 764 | 14QM126 | 30.4 | 87.0 | 30.7 | 7.2 | 5.9 | 29.7 | 86.5 |
| 765 | 14QM127 | 30.4 | 85.4 | 31.9 | 6.9 | 4.2 | 29.9 | 84.3 |
| 766 | 14QM127H | 31.0 | 86.5 | 30.6 | 6.9 | 4.7 | 30.2 | 85.7 |
| 767 | P♂ | 30.6 | 86.8 | 29.0 | 7.0 | 5.0 | 30.1 | 85.2 |
| 768 | 14QM128H | 31.7 | 87.1 | 30.2 | 7.0 | 4.9 | 31.1 | 86.3 |
| 769 | 14QM128 | 31.4 | 85.9 | 29.8 | 7.0 | 4.5 | 31.9 | 85.3 |
| 770 | 14QM129 | 31.7 | 86.7 | 29.7 | 7.0 | 5.3 | 31.1 | 84.9 |
| 771 | 14QM129H | 32.2 | 86.2 | 31.5 | 7.0 | 4.8 | 29.7 | 87.6 |
| 772 | P♂ | 30.8 | 86.4 | 30.2 | 7.0 | 4.7 | 30.1 | 85.2 |
| 773 | 14QM130H | 31.6 | 86.7 | 30.2 | 7.0 | 5.1 | 31.8 | 85.2 |
| 774 | 14QM130 | 33.2 | 86.1 | 32.8 | 7.2 | 4.0 | 31.4 | 84.5 |
| 775 | 14QM131 | 32.7 | 86.9 | 32.5 | 7.2 | 4.6 | 31.2 | 86.0 |
| 776 | 14QM131H | 32.0 | 86.6 | 29.9 | 7.0 | 4.9 | 32.5 | 85.8 |
| 777 | P♂ | 30.5 | 86.6 | 29.0 | 6.9 | 5.1 | 30.1 | 85.2 |
| 778 | 14QM132H | 30.3 | 86.0 | 29.4 | 7.0 | 5.3 | 30.0 | 86.6 |
| 779 | 14QM132 | 29.4 | 85.0 | 28.8 | 6.8 | 4.5 | 29.6 | 85.1 |
| 780 | 14QM133 | 32.0 | 87.2 | 31.9 | 7.1 | 5.0 | 32.1 | 86.4 |
| 781 | 14QM133H | 32.5 | 87.0 | 31.3 | 7.1 | 5.2 | 31.2 | 83.2 |
| 782 | P♂ | 30.7 | 85.5 | 29.3 | 6.9 | 4.9 | 30.1 | 85.2 |
| 783 | 14QM134H | 29.9 | 85.9 | 28.6 | 6.9 | 4.7 | 29.9 | 87.1 |
| 784 | 14QM134 | - | - | - | - | - | 28.9 | 86.8 |
| 785 | 14QM135 | 32.1 | 85.2 | 33.1 | 7.1 | 4.6 | 30.3 | 84.7 |
| 786 | 14QM135H | 30.6 | 85.8 | 29.5 | 7.0 | 4.6 | 31.1 | 84.7 |
| 787 | P♂ | 30.7 | 85.6 | 29.9 | 7.1 | 4.7 | 30.1 | 85.2 |
| 788 | 14QM136H | 31.3 | 84.7 | 31.0 | 7.0 | 4.8 | 31.3 | 85.0 |
| 789 | 14QM136 | - | - | - | - | - | 32.8 | 85.1 |
| 790 | 14QM137 | 30.5 | 84.5 | 29.5 | 6.8 | 4.6 | 32.2 | 84.1 |
| 791 | 14QM137H | 31.7 | 85.3 | 33.0 | 7.1 | 4.6 | 31.4 | 85.8 |
| 792 | P♂ | 31.4 | 85.4 | 30.1 | 7.0 | 4.5 | 30.1 | 85.2 |
| 793 | 14QM138H | 31.9 | 85.9 | 31.0 | 7.0 | 4.7 | 30.0 | 85.6 |
| 794 | 14QM138 | 30.6 | 85.2 | 29.8 | 6.9 | 4.1 | 29.3 | 84.6 |
| 795 | 14QM139 | 29.6 | 85.6 | 28.4 | 6.9 | 4.7 | 30.4 | 86.0 |
| 796 | 14QM139H | 30.7 | 85.7 | 30.4 | 7.0 | 4.5 | 30.6 | 85.7 |
| 797 | P♂ | 29.5 | 85.2 | 28.4 | 6.8 | 4.7 | 30.1 | 85.2 |
| 798 | 14QM140H | - | - | - | - | - | 31.9 | 86.2 |
| 799 | 14QM140 | 32.4 | 86.3 | 31.8 | 7.0 | 4.5 | 31.0 | 86.2 |
| 800 | 14QM141 | 31.3 | 87.5 | 32.1 | 7.1 | 4.9 | 30.5 | 86.5 |
| 801 | 14QM141H | 31.5 | 86.2 | 31.1 | 7.0 | 4.6 | 30.9 | 86.1 |
| 802 | 14QM380 | 30.4 | 87.3 | 30.3 | 7.0 | 4.8 | 29.9 | 85.0 |
| 803 | 14QM142H | 31.0 | 87.7 | 31.3 | 7.0 | 4.5 | 28.0 | 84.2 |
| 804 | 14QM142 | 30.8 | 86.7 | 31.9 | 7.1 | 4.5 | 31.1 | 85.2 |
| 805 | 14QM143 | 31.1 | 86.6 | 32.8 | 7.0 | 5.0 | 30.4 | 87.1 |
| 806 | 14QM143H | 30.8 | 85.1 | 30.9 | 6.9 | 4.5 | 29.8 | 87.0 |
| 807 | P♂ | 30.7 | 86.5 | 31.6 | 7.0 | 4.5 | 29.9 | 85.0 |
| 808 | 14QM144H | 31.0 | 87.1 | 32.1 | 7.0 | 4.9 | 31.0 | 86.0 |
| 809 | 14QM144 | 31.4 | 86.4 | 32.1 | 7.0 | 5.1 | 30.0 | 86.3 |
| 810 | 14QM145 | 29.8 | 86.3 | 29.1 | 6.8 | 4.4 | 31.7 | 86.4 |
| 811 | 14QM145H | 30.6 | 86.6 | 30.4 | 7.0 | 4.4 | 30.2 | 86.0 |
| 812 | P♂ | 30.5 | 85.6 | 32.5 | 7.0 | 4.7 | 29.9 | 85.0 |
| 813 | 14QM146H | 29.3 | 86.2 | 29.6 | 6.9 | 4.8 | 29.1 | 87.4 |
| 814 | 14QM146 | 30.6 | 86.9 | 32.3 | 7.0 | 4.1 | 31.5 | 86.2 |
| 815 | 14QM147 | 32.4 | 86.3 | 31.3 | 7.2 | 4.3 | 32.7 | 84.2 |
| 816 | 14QM147H | 32.4 | 87.4 | 29.5 | 7.1 | 4.6 | 29.2 | 84.2 |
| 817 | P♂ | 29.6 | 86.6 | 28.2 | 6.9 | 4.7 | 29.9 | 85.0 |
| 818 | 14QM148H | 31.5 | 84.6 | 32.0 | 7.1 | 4.9 | 31.0 | 85.8 |
| 819 | 14QM148 | 31.9 | 86.7 | 33.1 | 7.1 | 4.3 | 32.0 | 86.5 |
| 820 | 14QM149 | 30.5 | 85.4 | 31.6 | 6.9 | 4.2 | 32.2 | 85.4 |
| 821 | 14QM149H | 31.7 | 85.6 | 30.0 | 7.0 | 4.9 | 32.3 | 85.8 |
| 822 | P♂ | 30.3 | 86.4 | 30.6 | 7.0 | 4.1 | 29.9 | 85.0 |
| 823 | 14QM150H | 30.2 | 87.6 | 30.3 | 7.0 | 4.3 | 31.3 | 85.5 |
| 824 | 14QM150 | 31.7 | 87.6 | 31.5 | 7.0 | 4.4 | 31.5 | 86.1 |
| 825 | 14QM151 | 30.3 | 85.3 | 29.3 | 6.9 | 4.6 | 29.9 | 87.1 |
| 826 | 14QM151H | 31.2 | 85.8 | 31.0 | 7.0 | 4.1 | 31.3 | 85.7 |
| 827 | P♂ | 29.6 | 85.8 | 28.3 | 7.0 | 4.6 | 29.9 | 85.0 |
| 828 | 14QM152H | 29.8 | 84.9 | 29.3 | 6.9 | 4.9 | 29.8 | 85.8 |
| 829 | 14QM152 | 30.1 | 85.4 | 30.8 | 6.9 | 4.9 | 29.6 | 86.8 |
| 830 | 14QM153 | 29.5 | 86.0 | 30.4 | 6.9 | 5.3 | 29.2 | 87.5 |
| 831 | 14QM153H | 30.8 | 87.2 | 30.3 | 7.0 | 4.8 | 31.5 | 86.4 |
| 832 | P♂ | 29.6 | 87.5 | 29.3 | 6.9 | 5.3 | 29.9 | 85.0 |
| 833 | 14QM154H | 30.4 | 84.7 | 31.5 | 6.9 | 4.7 | 30.3 | 86.1 |
| 834 | 14QM154 | 32.1 | 87.2 | 32.6 | 7.0 | 4.7 | 31.7 | 86.1 |
| 835 | 14QM155 | 31.0 | 84.8 | 32.7 | 7.0 | 4.9 | 30.4 | 86.0 |
| 836 | 14QM155H | 31.5 | 86.0 | 30.5 | 7.1 | 5.0 | 31.2 | 86.8 |
| 837 | P♂ | 30.7 | 86.5 | 31.2 | 6.9 | 4.4 | 29.9 | 85.0 |
| 838 | 14QM156H | 31.4 | 86.8 | 30.6 | 7.1 | 4.9 | 30.7 | 86.1 |
| 839 | 14QM156 | 30.1 | 87.1 | 30.8 | 7.1 | 5.1 | 30.0 | 85.2 |
| 840 | 14QM157 | 30.7 | 85.6 | 31.7 | 7.2 | 4.9 | 31.6 | 86.2 |
| 841 | 14QM157H | 31.3 | 87.8 | 32.3 | 7.1 | 4.4 | 30.0 | 86.6 |
| 842 | P♂ | 30.7 | 86.3 | 31.9 | 7.0 | 4.6 | 29.9 | 85.0 |
| 843 | 14QM158H | 28.7 | 86.5 | 27.1 | 6.8 | 5.4 | 27.8 | 85.7 |
| 844 | 14QM158 | 28.2 | 84.2 | 26.6 | 6.8 | 4.8 | 28.3 | 85.9 |
| 845 | 14QM159 | 32.2 | 86.1 | 30.5 | 7.0 | 4.2 | 31.9 | 86.3 |
| 846 | 14QM159H | 33.2 | 87.3 | 31.2 | 7.2 | 4.7 | 30.9 | 86.8 |
| 847 | P♂ | 30.4 | 86.7 | 30.0 | 6.9 | 4.4 | 29.9 | 85.0 |
| 848 | 14QM160H | 29.9 | 84.3 | 32.2 | 7.0 | 5.0 | 29.1 | 86.1 |
| 849 | 14QM160 | 31.6 | 86.7 | 31.7 | 7.1 | 4.7 | 30.5 | 85.2 |
| 850 | 14QM161 | 30.8 | 86.0 | 31.3 | 7.0 | 4.8 | 30.5 | 85.9 |
| 851 | 14QM161H | 30.7 | 85.8 | 30.7 | 6.9 | 4.2 | 31.0 | 85.4 |
| 852 | 14QM380 | 29.5 | 84.7 | 30.5 | 6.9 | 3.9 | 29.3 | 85.5 |
| 853 | 14QM162H | 30.5 | 87.0 | 30.3 | 7.0 | 5.2 | 31.1 | 85.8 |
| 854 | 14QM162 | 29.2 | 87.2 | 31.8 | 7.1 | 5.4 | 30.1 | 84.6 |
| 855 | 14QM163 | 32.2 | 87.2 | 32.6 | 7.2 | 4.7 | 31.4 | 85.7 |
| 856 | 14QM163H | 31.6 | 87.0 | 32.3 | 7.0 | 4.5 | 31.1 | 85.2 |
| 857 | P♂ | 30.4 | 87.5 | 30.8 | 7.0 | 4.4 | 29.3 | 85.5 |
| 858 | 14QM164H | 31.2 | 86.7 | 32.9 | 7.2 | 5.1 | 29.7 | 87.9 |
| 859 | 14QM164 | 29.4 | 84.9 | 29.5 | 6.9 | 5.2 | 28.7 | 86.3 |
| 860 | 14QM165 | 29.8 | 87.3 | 32.8 | 6.9 | 4.3 | 31.4 | 84.0 |
| 861 | 14QM165H | 32.1 | 87.0 | 32.5 | 7.0 | 4.5 | 32.2 | 85.3 |
| 862 | P♂ | 30.2 | 85.9 | 30.3 | 6.9 | 4.6 | 29.3 | 85.5 |
| 863 | 14QM166H | 32.5 | 88.1 | 31.9 | 7.1 | 4.6 | 31.3 | 85.2 |
| 864 | 14QM166 | 32.5 | 86.9 | 34.0 | 7.0 | 4.2 | 31.1 | 85.0 |
| 865 | 14QM167 | 31.8 | 86.8 | 32.0 | 7.0 | 4.6 | 31.2 | 84.3 |
| 866 | 14QM167H | 31.5 | 87.0 | 31.4 | 7.1 | 4.6 | 29.9 | 85.3 |
| 867 | P♂ | 30.1 | 87.3 | 30.2 | 6.9 | 5.1 | 29.3 | 85.5 |
| 868 | 14QM168H | 31.3 | 86.3 | 32.7 | 7.0 | 4.5 | 30.8 | 85.7 |
| 869 | 14QM168 | 29.2 | 85.2 | 29.3 | 6.8 | 5.0 | 28.7 | 86.1 |
| 870 | 14QM169 | 28.9 | 85.2 | 26.9 | 6.8 | 4.9 | 28.0 | 85.0 |
| 871 | 14QM169H | 28.6 | 85.5 | 28.3 | 6.9 | 4.7 | 29.4 | 87.4 |
| 872 | P♂ | 29.4 | 84.6 | 31.6 | 7.1 | 4.5 | 29.3 | 85.5 |
| 873 | 14QM170H | 30.6 | 87.5 | 31.5 | 7.0 | 4.9 | 30.5 | 86.6 |
| 874 | 14QM170 | 29.9 | 84.5 | 29.2 | 7.0 | 5.5 | 30.1 | 86.4 |
| 875 | 14QM171 | 30.2 | 86.5 | 30.1 | 6.9 | 5.1 | 31.2 | 84.9 |
| 876 | 14QM171H | 30.7 | 85.1 | 31.2 | 6.9 | 4.1 | 30.4 | 85.4 |
| 877 | P♂ | 31.0 | 85.8 | 28.7 | 7.0 | 5.2 | 29.3 | 85.5 |
| 878 | 14QM172H | 31.2 | 85.3 | 31.2 | 7.0 | 4.7 | 29.8 | 83.5 |
| 879 | 14QM172 | 31.1 | 85.2 | 32.5 | 7.0 | 4.4 | 30.4 | 85.3 |
| 880 | 14QM173 | 32.1 | 85.4 | 32.3 | 7.1 | 4.7 | 30.4 | 85.0 |
| 881 | 14QM173H | 31.0 | 85.6 | 31.4 | 7.0 | 5.0 | 30.7 | 86.1 |
| 882 | P♂ | 30.3 | 86.6 | 29.8 | 6.9 | 4.6 | 29.3 | 85.5 |
| 883 | 14QM174H | 28.7 | 85.3 | 30.1 | 7.0 | 4.8 | 28.6 | 86.3 |
| 884 | 14QM174 | 30.2 | 86.7 | 30.6 | 7.0 | 4.9 | 30.8 | 87.2 |
| 885 | 14QM175 | 30.4 | 85.3 | 32.0 | 7.1 | 4.8 | 31.0 | 84.4 |
| 886 | 14QM175H | 31.4 | 86.3 | 31.0 | 6.9 | 4.5 | 30.0 | 86.1 |
| 887 | P♂ | 30.5 | 84.5 | 31.3 | 6.9 | 4.4 | 29.3 | 85.5 |
| 888 | 14QM176H | 31.6 | 87.0 | 30.4 | 6.9 | 4.6 | 31.9 | 86.2 |
| 889 | 14QM176 | 33.1 | 86.5 | 32.1 | 7.0 | 4.4 | 32.8 | 85.1 |
| 890 | 14QM177 | 29.9 | 85.9 | 30.4 | 6.9 | 4.2 | 29.0 | 84.5 |
| 891 | 14QM177H | 31.6 | 84.9 | 32.0 | 7.0 | 4.1 | 31.2 | 88.1 |
| 892 | P♂ | 30.4 | 85.2 | 30.0 | 7.0 | 4.4 | 29.3 | 85.5 |
| 893 | 14QM178H | 31.0 | 83.8 | 32.3 | 7.0 | 4.7 | 30.1 | 84.8 |
| 894 | 14QM178 | 32.4 | 85.2 | 31.7 | 7.0 | 4.7 | 32.6 | 84.7 |
| 895 | 14QM179 | 31.2 | 85.3 | 30.9 | 7.0 | 4.7 | 30.6 | 85.6 |
| 896 | 14QM179H | 31.0 | 85.8 | 30.5 | 7.1 | 5.4 | 30.6 | 85.6 |
| 897 | P♂ | 29.7 | 83.6 | 29.3 | 6.9 | 4.6 | 29.3 | 85.5 |
| 898 | P ♂ | 30.0 | 85.3 | 30.1 | 6.9 | 4.5 | 30.8 | 86.3 |
| 899 | F1 | 31.3 | 86.0 | 30.9 | 7.1 | 5.1 | 31.4 | 86.4 |
| 900 | P♀ | 30.3 | 84.9 | 30.0 | 6.9 | 5.0 | 31.3 | 85.8 |

4QM003H- 14QM179H refer to 177 F<sub>14</sub>BC progenies in BC/P population.
| FS(cN/tex<br>) | FE/% | FM | FL/mm | FU/% | FS(cN/tex<br>) | FE/% | FM |
| --- | --- | --- | --- | --- | --- | --- | --- |
| 2016E1 | 2016E1 | 2016E1 | 2016E2 | 2016E2 | 2016E2 | 2016E2 | 2016E2 |
| 28.1 | 6.7 | 5.5 | 29.3 | 86.2 | 28.9 | 6.8 | 5.0 |
| 29.0 | 6.8 | 5.1 | 30.7 | 86.3 | 31.9 | 6.9 | 4.8 |
| 26.3 | 6.8 | 5.5 | 28.4 | 84.9 | 29.2 | 6.6 | 4.9 |
| 27.5 | 6.8 | 5.0 | 30.1 | 87.2 | 30.2 | 6.8 | 4.9 |
| 27.0 | 6.7 | 5.1 | 30.5 | 86.0 | 29.6 | 6.8 | 4.9 |
| 31.8 | 6.8 | 4.9 | 29.3 | 82.9 | 31.1 | 6.7 | 4.8 |
| 28.0 | 6.7 | 4.3 | 30.8 | 85.3 | 30.6 | 6.7 | 4.3 |
| 26.3 | 6.8 | 5.5 | 28.4 | 84.9 | 29.2 | 6.6 | 4.9 |
| 27.6 | 6.7 | 5.2 | 29.9 | 85.7 | 28.6 | 6.7 | 5.1 |
| 31.1 | 6.9 | 4.8 | 30.3 | 85.6 | 30.5 | 6.7 | 4.8 |
| 27.2 | 6.8 | 5.5 | 29.5 | 85.0 | 28.2 | 6.8 | 5.6 |
| 27.2 | 6.9 | 5.5 | 29.8 | 86.3 | 29.0 | 6.8 | 5.3 |
| 26.3 | 6.8 | 5.5 | 28.4 | 84.9 | 29.2 | 6.6 | 4.9 |
| 28.7 | 6.8 | 4.5 | 29.3 | 84.1 | 29.0 | 6.7 | 4.7 |
| 29.6 | 6.8 | 4.5 | 27.8 | 84.6 | 28.8 | 6.6 | 4.8 |
| 28.6 | 6.7 | 4.2 | 28.6 | 85.3 | 26.7 | 6.7 | 5.7 |
| 25.9 | 6.6 | 5.3 | 28.9 | 85.0 | 28.4 | 6.7 | 5.4 |
| 26.3 | 6.8 | 5.5 | 28.4 | 84.9 | 29.2 | 6.6 | 4.9 |
| 28.1 | 6.9 | 4.4 | 29.9 | 86.8 | 28.5 | 6.8 | 5.1 |
| 30.8 | 6.8 | 4.7 | 29.0 | 86.7 | 28.9 | 6.7 | 4.9 |
| 27.0 | 6.6 | 5.6 | 30.6 | 87.5 | 29.4 | 6.8 | 5.6 |
| 29.3 | 6.9 | 5.3 | 31.4 | 87.2 | 29.3 | 6.8 | 4.8 |
| 26.3 | 6.8 | 5.5 | 28.4 | 84.9 | 29.2 | 6.6 | 4.9 |
| 31.0 | 6.9 | 4.5 | 30.3 | 85.6 | 30.9 | 6.8 | 4.9 |
| 29.5 | 6.8 | 5.2 | 29.8 | 85.2 | 29.8 | 6.7 | 5.1 |
| 27.0 | 6.7 | 5.5 | 29.1 | 86.3 | 27.7 | 6.8 | 5.7 |
| 28.2 | 6.7 | 3.8 | 29.5 | 85.2 | 27.3 | 6.7 | 5.3 |
| 26.3 | 6.8 | 5.5 | 28.4 | 84.9 | 29.2 | 6.6 | 4.9 |
| 30.5 | 6.8 | 4.4 | 30.3 | 87.1 | 27.7 | 6.7 | 5.3 |
| 26.8 | 6.7 | 4.9 | 29.4 | 86.0 | 27.9 | 6.7 | 5.0 |
| 27.7 | 6.8 | 4.5 | 30.6 | 86.7 | 28.4 | 6.9 | 4.8 |
| 27.4 | 6.8 | 5.1 | 29.9 | 84.1 | 29.0 | 6.8 | 5.0 |
| 26.3 | 6.8 | 5.5 | 28.4 | 84.9 | 29.2 | 6.6 | 4.9 |
| 27.8 | 6.8 | 5.1 | 29.9 | 87.0 | 28.7 | 6.8 | 5.3 |
| 28.9 | 6.8 | 5.2 | 29.0 | 85.2 | 27.8 | 6.7 | 5.4 |
| 29.8 | 6.9 | 4.4 | 29.7 | 86.8 | 30.1 | 6.7 | 4.6 |
| 28.6 | 6.9 | 4.7 | 30.3 | 85.5 | 29.5 | 6.8 | 5.1 |
| 26.3 | 6.8 | 5.5 | 28.4 | 84.9 | 29.2 | 6.6 | 4.9 |
| 28.9 | 6.8 | 4.7 | 28.9 | 85.2 | 29.4 | 6.7 | 5.0 |
| 31.9 | 6.9 | 4.3 | 30.7 | 86.4 | 30.2 | 6.8 | 4.5 |
| 28.0 | 6.7 | 4.6 | 28.5 | 85.7 | 29.1 | 6.7 | 5.6 |
| 31.4 | 6.8 | 4.3 | 28.9 | 85.3 | 29.1 | 6.8 | 5.6 |
| 26.3 | 6.8 | 5.5 | 28.4 | 84.9 | 29.2 | 6.6 | 4.9 |
| 28.4 | 6.8 | 4.5 | 30.4 | 85.6 | 30.1 | 6.8 | 5.1 |
| 29.3 | 6.9 | 4.6 | 30.6 | 83.4 | 30.5 | 6.8 | 4.8 |
| 29.2 | 6.8 | 4.6 | 28.4 | 84.3 | 28.6 | 6.7 | 5.7 |
| 27.6 | 6.8 | 4.8 | 29.5 | 85.4 | 28.4 | 6.8 | 5.4 |
| 26.3 | 6.8 | 5.5 | 28.4 | 84.9 | 29.2 | 6.6 | 4.9 |
| 28.5 | 6.8 | 4.6 | 29.9 | 87.2 | 28.9 | 6.8 | 5.4 |
| 28.5 | 6.8 | 4.6 | 30.7 | 87.4 | 32.4 | 7.0 | 4.9 |
| 28.9 | 6.7 | 4.4 | 28.0 | 85.1 | 27.4 | 6.6 | 5.0 |
| 29.5 | 6.8 | 4.5 | 29.0 | 85.2 | 27.8 | 6.7 | 5.3 |
| 26.4 | 6.7 | 4.4 | 28.6 | 84.8 | 27.1 | 6.8 | 5.9 |
| 30.1 | 6.9 | 4.5 | 29.5 | 84.1 | 29.7 | 6.8 | 4.5 |
| 28.9 | 6.8 | 4.5 | 27.6 | 85.2 | 29.8 | 6.7 | 4.3 |
| 30.7 | 6.7 | 3.8 | 29.3 | 85.6 | 28.7 | 6.8 | 4.9 |
| 28.7 | 6.9 | 5.7 | 28.2 | 86.6 | 26.3 | 6.6 | 5.0 |
| 26.4 | 6.7 | 4.4 | 28.6 | 84.8 | 27.1 | 6.8 | 5.9 |
| 28.0 | 6.8 | 4.9 | 30.2 | 86.6 | 28.0 | 6.8 | 4.9 |
| 28.6 | 6.9 | 5.3 | 29.4 | 86.6 | 27.8 | 6.8 | 5.4 |
| 27.4 | 6.8 | 4.7 | 29.3 | 85.1 | 28.1 | 6.7 | 5.0 |
| 27.8 | 6.8 | 4.7 | 28.7 | 84.9 | 28.0 | 6.7 | 5.4 |
| 26.4 | 6.7 | 4.4 | 28.6 | 84.8 | 27.1 | 6.8 | 5.9 |
| 27.9 | 6.8 | 5.1 | 28.7 | 86.2 | 28.1 | 6.7 | 5.4 |
| 27.9 | 6.8 | 5.3 | 26.5 | 84.0 | 26.8 | 6.6 | 5.6 |
| 28.8 | 6.9 | 4.5 | 30.4 | 86.8 | 29.5 | 6.8 | 4.7 |
| 28.2 | 6.8 | 4.1 | 30.5 | 85.6 | 29.5 | 6.8 | 5.1 |
| 26.4 | 6.7 | 4.4 | 28.6 | 84.8 | 27.1 | 6.8 | 5.9 |
| 29.1 | 6.9 | 3.9 | 29.9 | 87.8 | 27.7 | 6.7 | 4.7 |
| 29.1 | 6.8 | 4.1 | 30.2 | 84.5 | 28.7 | 6.8 | 4.7 |
| 27.8 | 6.7 | 3.5 | 29.9 | 86.9 | 28.9 | 6.7 | 4.8 |
| 30.2 | 6.9 | 4.0 | 29.5 | 86.0 | 28.4 | 6.8 | 4.7 |
| 26.4 | 6.7 | 4.4 | 28.6 | 84.8 | 27.1 | 6.8 | 5.9 |
| 28.3 | 6.8 | 5.3 | 29.7 | 83.9 | 28.2 | 6.7 | 5.3 |
| 29.7 | 6.9 | 5.4 | 25.9 | 83.0 | 26.2 | 6.6 | 5.8 |
| 28.0 | 6.8 | 5.6 | 30.0 | 87.0 | 29.8 | 6.8 | 4.9 |
| 27.8 | 6.8 | 5.7 | 29.4 | 85.3 | 28.5 | 6.7 | 5.1 |
| 26.4 | 6.7 | 4.4 | 28.6 | 84.8 | 27.1 | 6.8 | 5.9 |
| 28.1 | 6.8 | 5.5 | 26.6 | 83.5 | 26.2 | 6.6 | 5.4 |
| 29.2 | 6.8 | 5.4 | 29.1 | 84.5 | 29.8 | 6.7 | 4.6 |
| 29.1 | 6.7 | 4.6 | 30.1 | 85.4 | 28.2 | 6.7 | 4.8 |
| 29.1 | 6.7 | 4.6 | 29.8 | 84.8 | 29.3 | 6.7 | 4.6 |
| 26.4 | 6.7 | 4.4 | 28.6 | 84.8 | 27.1 | 6.8 | 5.9 |
| 27.8 | 6.8 | 5.1 | 29.3 | 87.0 | 29.1 | 6.7 | 4.9 |
| 27.4 | 6.9 | 4.8 | 29.7 | 87.4 | 28.9 | 6.8 | 5.3 |
| 29.5 | 6.7 | 4.7 | 30.0 | 87.1 | 31.2 | 6.7 | 4.2 |
| 27.7 | 6.7 | 4.9 | 29.8 | 85.2 | 29.4 | 6.7 | 4.6 |
| 26.4 | 6.7 | 4.4 | 28.6 | 84.8 | 27.1 | 6.8 | 5.9 |
| 27.3 | 6.7 | 4.9 | 28.3 | 84.4 | 28.8 | 6.7 | 5.0 |
| 26.4 | 6.6 | 4.9 | 30.1 | 85.5 | 29.2 | 6.8 | 5.1 |
| 27.8 | 6.8 | 5.6 | 28.7 | 85.4 | 27.7 | 6.7 | 5.5 |
| 28.0 | 6.8 | 5.4 | 29.8 | 86.4 | 28.8 | 6.8 | 5.4 |
| 26.4 | 6.7 | 4.4 | 28.6 | 84.8 | 27.1 | 6.8 | 5.9 |
| 28.5 | 6.7 | 4.4 | 30.0 | 87.8 | 28.1 | 6.8 | 5.4 |
| 29.3 | 6.8 | 5.0 | 27.5 | 82.2 | 27.5 | 6.6 | 5.1 |
| 30.3 | 6.9 | 4.9 | 29.1 | 83.6 | 28.4 | 6.7 | 4.6 |
| 29.6 | 6.7 | 3.9 | 27.9 | 84.6 | 28.8 | 6.7 | 4.5 |
| 26.4 | 6.7 | 4.4 | 28.6 | 84.8 | 27.1 | 6.8 | 5.9 |
| 27.9 | 6.9 | 5.4 | 29.9 | 86.0 | 28.1 | 6.7 | 5.1 |
| 28.8 | 6.9 | 5.3 | 30.9 | 85.6 | 28.9 | 6.9 | 5.8 |
| 26.7 | 6.8 | 5.3 | 28.2 | 86.2 | 28.3 | 6.7 | 4.9 |
| 29.0 | 6.9 | 4.7 | 28.7 | 85.1 | 27.4 | 6.7 | 5.0 |
| 28.0 | 6.8 | 3.9 | 27.6 | 84.1 | 27.0 | 6.6 | 5.0 |
| 27.7 | 6.8 | 5.1 | 29.6 | 85.3 | 31.1 | 6.8 | 5.1 |
| 27.2 | 6.8 | 5.0 | 27.6 | 82.1 | 26.8 | 6.6 | 5.7 |
| 29.7 | 6.8 | 5.1 | 28.1 | 83.9 | 28.8 | 6.7 | 4.8 |
| 31.0 | 7.0 | 4.8 | 30.2 | 87.4 | 29.2 | 6.8 | 4.7 |
| 28.0 | 6.8 | 3.9 | 27.6 | 84.1 | 27.0 | 6.6 | 5.0 |
| 28.8 | 6.8 | 4.6 | 29.8 | 86.0 | 26.8 | 6.7 | 5.4 |
| 27.1 | 6.7 | 5.2 | 29.9 | 86.6 | 28.3 | 6.7 | 4.9 |
| 29.4 | 6.8 | 5.3 | 29.2 | 86.2 | 29.3 | 6.7 | 5.1 |
| 27.6 | 6.8 | 5.7 | 29.7 | 87.2 | 28.3 | 6.7 | 4.9 |
| 28.0 | 6.8 | 3.9 | 27.6 | 84.1 | 27.0 | 6.6 | 5.0 |
| 28.0 | 6.7 | 4.4 | 28.5 | 85.6 | 28.6 | 6.7 | 4.5 |
| 27.7 | 6.6 | 4.3 | 28.9 | 86.6 | 28.2 | 6.7 | 4.6 |
| 29.6 | 6.8 | 4.5 | 28.4 | 86.9 | 29.5 | 6.7 | 4.2 |
| 28.3 | 6.8 | 4.8 | 30.4 | 88.2 | 28.6 | 6.8 | 4.8 |
| 28.0 | 6.8 | 3.9 | 27.6 | 84.1 | 27.0 | 6.6 | 5.0 |
| 28.4 | 6.8 | 4.7 | 30.4 | 86.5 | 29.6 | 6.8 | 5.1 |
| 31.4 | 6.8 | 4.8 | 28.9 | 84.9 | 29.2 | 6.7 | 5.3 |
| 28.9 | 6.8 | 5.1 | 28.0 | 85.5 | 28.3 | 6.6 | 5.3 |
| 29.0 | 6.7 | 4.7 | 29.1 | 85.3 | 28.9 | 6.7 | 5.7 |
| 28.0 | 6.8 | 3.9 | 27.6 | 84.1 | 27.0 | 6.6 | 5.0 |
| 29.1 | 6.8 | 4.9 | 28.5 | 84.7 | 28.0 | 6.7 | 5.1 |
| 28.1 | 6.7 | 5.1 | 28.8 | 86.9 | 29.6 | 6.7 | 5.0 |
| 29.8 | 6.7 | 4.8 | 30.5 | 84.7 | 30.9 | 6.7 | 4.5 |
| 30.6 | 6.9 | 4.8 | 31.0 | 86.8 | 31.3 | 6.8 | 4.6 |
| 28.0 | 6.8 | 3.9 | 27.6 | 84.1 | 27.0 | 6.6 | 5.0 |
| 26.0 | 6.6 | 5.0 | 29.5 | 85.3 | 28.8 | 6.8 | 4.7 |
| 27.1 | 6.7 | 4.6 | 29.2 | 86.0 | 29.7 | 6.8 | 4.9 |
| 27.0 | 6.6 | 4.9 | 26.2 | 83.4 | 28.9 | 6.5 | 5.0 |
| 28.9 | 6.7 | 4.6 | 29.5 | 86.5 | 28.9 | 6.7 | 5.4 |
| 28.0 | 6.8 | 3.9 | 27.6 | 84.1 | 27.0 | 6.6 | 5.0 |
| 29.9 | 6.8 | 4.7 | 30.2 | 86.7 | 29.1 | 6.8 | 4.8 |
| 30.9 | 6.8 | 4.7 | 28.9 | 84.6 | 29.6 | 6.7 | 4.6 |
| 27.1 | 6.8 | 4.8 | 26.6 | 81.6 | 26.5 | 6.6 | 5.0 |
| 26.3 | 6.7 | 4.9 | 27.4 | 83.9 | 26.7 | 6.6 | 4.9 |
| 28.0 | 6.8 | 3.9 | 27.6 | 84.1 | 27.0 | 6.6 | 5.0 |
| 27.6 | 6.8 | 4.7 | 29.2 | 86.5 | 30.6 | 6.8 | 4.6 |
| 31.2 | 6.9 | 4.6 | 29.4 | 85.8 | 31.8 | 6.7 | 4.2 |
| 28.9 | 6.7 | 4.5 | 25.5 | 82.8 | 25.5 | 6.5 | 5.2 |
| 30.1 | 6.9 | 5.0 | 28.1 | 86.2 | 27.2 | 6.6 | 5.1 |
| 28.0 | 6.8 | 3.9 | 27.6 | 84.1 | 27.0 | 6.6 | 5.0 |
| 27.5 | 6.8 | 5.3 | 29.2 | 86.4 | 29.0 | 6.7 | 4.9 |
| 29.2 | 6.9 | 5.5 | 29.8 | 86.2 | 31.9 | 6.8 | 4.6 |
| 28.0 | 6.8 | 5.6 | 27.0 | 84.7 | 28.9 | 6.6 | 5.0 |
| 26.5 | 6.7 | 5.4 | 29.2 | 86.5 | 29.7 | 6.7 | 4.9 |
| 28.0 | 6.8 | 3.9 | 27.6 | 84.1 | 27.0 | 6.6 | 5.0 |
| 28.1 | 6.9 | 5.3 | 28.7 | 84.1 | 31.9 | 6.7 | 4.2 |
| 30.2 | 6.9 | 5.0 | 26.9 | 81.0 | 29.6 | 6.6 | 4.5 |
| 27.7 | 6.7 | 4.9 | 29.0 | 86.5 | 31.0 | 6.7 | 4.8 |
| 29.4 | 6.9 | 5.0 | 27.7 | 84.5 | 29.0 | 6.6 | 4.6 |
| 27.8 | 6.8 | 5.2 | 30.0 | 87.3 | 30.4 | 6.8 | 4.6 |
| 29.2 | 6.8 | 4.8 | 27.9 | 84.2 | 29.6 | 6.7 | 5.2 |
| 27.3 | 6.7 | 5.2 | 30.2 | 86.4 | 31.1 | 6.8 | 5.4 |
| 30.1 | 6.8 | 4.4 | 30.4 | 87.2 | 29.8 | 6.7 | 5.0 |
| 28.6 | 6.8 | 5.1 | 30.3 | 86.3 | 30.0 | 6.7 | 4.5 |
| 27.8 | 6.8 | 5.2 | 30.0 | 87.3 | 30.4 | 6.8 | 4.6 |
| 27.1 | 6.6 | 5.0 | 28.9 | 83.2 | 29.2 | 6.7 | 5.3 |
| 27.0 | 6.7 | 5.2 | 28.4 | 83.2 | 27.5 | 6.7 | 5.1 |
| 29.1 | 6.7 | 4.6 | 29.0 | 85.5 | 27.7 | 6.7 | 5.4 |
| 28.5 | 6.7 | 3.5 | 28.8 | 86.3 | 28.4 | 6.7 | 5.0 |
| 27.8 | 6.8 | 5.2 | 30.0 | 87.3 | 30.4 | 6.8 | 4.6 |
| 27.0 | 6.7 | 5.0 | 30.5 | 86.7 | 29.3 | 6.8 | 5.0 |
| 28.2 | 6.8 | 5.3 | 30.2 | 84.8 | 30.5 | 6.8 | 4.7 |
| 26.6 | 6.7 | 5.8 | 27.5 | 84.7 | 26.9 | 6.6 | 5.6 |
| 27.4 | 6.8 | 5.6 | 28.7 | 85.6 | 28.1 | 6.7 | 5.2 |
| 27.8 | 6.8 | 5.2 | 30.0 | 87.3 | 30.4 | 6.8 | 4.6 |
| 26.9 | 6.8 | 5.0 | 28.7 | 85.2 | 28.8 | 6.6 | 4.4 |
| 29.2 | 6.8 | 5.4 | 30.1 | 86.1 | 28.9 | 6.7 | 4.8 |
| 28.6 | 6.8 | 4.5 | 29.7 | 87.1 | 30.8 | 6.8 | 4.5 |
| 26.4 | 6.8 | 5.6 | 29.2 | 85.5 | 29.5 | 6.7 | 4.6 |
| 27.8 | 6.8 | 5.2 | 30.0 | 87.3 | 30.4 | 6.8 | 4.6 |
| 29.2 | 6.8 | 4.6 | 29.3 | 85.1 | 30.2 | 6.7 | 4.3 |
| 28.5 | 6.7 | 5.0 | 28.9 | 83.5 | 28.7 | 6.6 | 4.8 |
| 31.9 | 6.9 | 5.0 | 30.7 | 85.2 | 31.7 | 6.8 | 4.7 |
| 28.2 | 6.8 | 5.0 | 29.8 | 87.2 | 30.5 | 6.8 | 4.8 |
| 27.8 | 6.8 | 5.2 | 30.0 | 87.3 | 30.4 | 6.8 | 4.6 |
| 31.0 | 6.9 | 4.8 | 29.5 | 86.0 | 30.2 | 6.8 | 5.1 |
| 30.9 | 6.8 | 4.7 | 27.2 | 83.2 | 26.2 | 6.5 | 5.1 |
| 27.6 | 6.9 | 4.2 | 30.0 | 84.5 | 27.4 | 6.7 | 4.3 |
| 27.9 | 6.7 | 3.9 | 30.3 | 85.6 | 28.7 | 6.8 | 4.7 |
| 27.8 | 6.8 | 5.2 | 30.0 | 87.3 | 30.4 | 6.8 | 4.6 |
| 26.3 | 6.7 | 4.9 | 29.4 | 85.0 | 28.1 | 6.7 | 4.6 |
| 25.8 | 6.7 | 5.3 | 28.5 | 84.9 | 27.1 | 6.7 | 4.9 |
| 27.6 | 6.7 | 4.9 | 28.6 | 85.5 | 27.4 | 6.7 | 5.2 |
| 28.1 | 6.8 | 4.3 | 28.7 | 85.1 | 29.2 | 6.8 | 5.1 |
| 27.8 | 6.8 | 5.2 | 30.0 | 87.3 | 30.4 | 6.8 | 4.6 |
| 29.2 | 6.8 | 4.4 | 31.1 | 86.8 | 28.6 | 6.8 | 4.6 |
| 27.0 | 6.7 | 5.1 | 29.1 | 85.4 | 28.8 | 6.7 | 4.7 |
| 30.3 | 6.8 | 4.7 | 30.8 | 85.9 | 29.5 | 6.7 | 5.1 |
| 29.0 | 6.7 | 4.7 | 30.0 | 85.9 | 29.8 | 6.7 | 5.1 |
| 27.8 | 6.8 | 5.2 | 30.0 | 87.3 | 30.4 | 6.8 | 4.6 |
| 29.0 | 6.8 | 4.7 | 27.5 | 84.6 | 27.9 | 6.6 | 4.6 |
| 28.7 | 6.7 | 4.3 | 27.0 | 82.4 | 26.5 | 6.6 | 4.9 |
| 29.5 | 6.8 | 4.8 | 30.2 | 87.1 | 29.6 | 6.7 | 4.8 |
| 28.6 | 6.7 | 4.6 | 29.5 | 85.6 | 28.1 | 6.7 | 4.9 |
| 27.8 | 6.8 | 5.2 | 30.0 | 87.3 | 30.4 | 6.8 | 4.6 |
| 28.0 | 6.7 | 4.4 | 27.4 | 84.0 | 27.7 | 6.6 | 4.9 |
| 31.4 | 6.8 | 4.8 | 29.9 | 86.2 | 30.2 | 6.8 | 4.9 |
| 28.8 | 6.9 | 5.1 | 29.4 | 85.0 | 29.0 | 6.8 | 5.5 |
| 28.6 | 6.7 | 4.6 | 27.1 | 84.3 | 26.4 | 6.6 | 5.3 |
| 29.9 | 6.9 | 5.3 | 28.9 | 86.2 | 27.0 | 6.7 | 5.1 |
| 31.1 | 6.9 | 4.5 | 30.1 | 87.1 | 28.3 | 6.7 | 4.9 |
| 27.1 | 6.8 | 5.6 | 29.1 | 85.9 | 30.5 | 6.8 | 5.4 |
| 31.4 | 6.8 | 4.3 | 30.4 | 85.6 | 31.2 | 6.8 | 4.9 |
| 29.4 | 6.9 | 4.8 | 30.1 | 86.0 | 28.5 | 6.7 | 5.1 |
| 29.9 | 6.9 | 5.3 | 28.9 | 86.2 | 27.0 | 6.7 | 5.1 |
| 27.4 | 6.8 | 5.2 | 30.9 | 87.1 | 29.7 | 6.9 | 5.7 |
| 27.6 | 6.8 | 5.4 | 29.4 | 86.0 | 27.4 | 6.8 | 6.0 |
| 29.1 | 6.9 | 5.2 | 28.6 | 83.8 | 30.2 | 6.8 | 5.6 |
| 27.8 | 6.7 | 4.9 | 28.0 | 84.5 | 28.1 | 6.7 | 5.0 |
| 29.9 | 6.9 | 5.3 | 28.9 | 86.2 | 27.0 | 6.7 | 5.1 |
| 28.3 | 6.9 | 5.2 | 28.7 | 84.6 | 27.9 | 6.6 | 4.3 |
| 28.6 | 6.9 | 5.8 | 28.3 | 85.3 | 28.6 | 6.7 | 5.7 |
| 28.2 | 6.7 | 5.7 | 27.2 | 84.4 | 27.6 | 6.7 | 5.9 |
| 26.9 | 6.8 | 5.4 | 28.5 | 84.0 | 28.7 | 6.7 | 5.1 |
| 29.9 | 6.9 | 5.3 | 28.9 | 86.2 | 27.0 | 6.7 | 5.1 |
| 28.4 | 6.9 | 5.2 | 30.2 | 85.7 | 27.9 | 6.7 | 5.3 |
| 31.0 | 6.9 | 4.5 | 30.9 | 85.9 | 31.8 | 6.8 | 4.2 |
| 29.4 | 6.8 | 4.9 | 28.1 | 84.0 | 27.0 | 6.6 | 5.0 |
| 28.0 | 6.8 | 5.1 | 27.9 | 82.7 | 27.4 | 6.6 | 4.8 |
| 29.9 | 6.9 | 5.3 | 28.9 | 86.2 | 27.0 | 6.7 | 5.1 |
| 29.4 | 6.8 | 5.1 | 27.8 | 85.4 | 26.7 | 6.6 | 5.5 |
| 28.2 | 6.8 | 5.7 | 29.8 | 86.7 | 30.0 | 6.7 | 5.1 |
| 29.8 | 6.8 | 5.3 | 29.8 | 86.4 | 29.1 | 6.7 | 5.7 |
| 29.9 | 6.9 | 5.3 | 28.9 | 86.2 | 27.0 | 6.7 | 5.1 |
| 28.1 | 6.8 | 5.0 | 29.4 | 86.2 | 29.9 | 6.8 | 5.5 |
| 27.4 | 6.8 | 4.8 | 31.3 | 87.2 | 30.1 | 6.8 | 5.1 |
| 27.9 | 6.8 | 4.8 | 29.0 | 84.4 | 28.4 | 6.7 | 4.9 |
| 29.4 | 6.8 | 4.6 | 28.7 | 84.3 | 27.5 | 6.7 | 5.1 |
| 29.9 | 6.9 | 5.3 | 28.9 | 86.2 | 27.0 | 6.7 | 5.1 |
| 28.7 | 6.7 | 5.1 | 28.6 | 85.6 | 28.0 | 6.7 | 4.8 |
| 26.1 | 6.6 | 5.6 | 28.8 | 84.3 | 27.9 | 6.7 | 5.4 |
| 29.1 | 6.8 | 5.4 | 27.6 | 83.7 | 27.4 | 6.7 | 5.5 |
| 29.9 | 6.8 | 4.8 | 30.1 | 86.0 | 27.6 | 6.7 | 4.9 |
| 29.9 | 6.9 | 5.3 | 28.9 | 86.2 | 27.0 | 6.7 | 5.1 |
| 27.6 | 6.8 | 5.1 | 29.0 | 84.1 | 27.6 | 6.7 | 4.6 |
| 28.4 | 6.8 | 4.6 | 31.4 | 84.4 | 29.6 | 6.8 | 4.5 |
| 28.5 | 6.8 | 5.3 | 27.7 | 82.7 | 27.0 | 6.6 | 5.0 |
| 30.0 | 6.8 | 4.5 | 29.0 | 85.7 | 27.0 | 6.7 | 4.9 |
| 29.9 | 6.9 | 5.3 | 28.9 | 86.2 | 27.0 | 6.7 | 5.1 |
| 28.5 | 6.9 | 5.3 | 29.9 | 84.2 | 29.8 | 6.8 | 4.2 |
| 28.4 | 6.8 | 4.6 | 28.0 | 83.2 | 27.5 | 6.7 | 5.0 |
| 27.4 | 6.7 | 4.7 | 29.6 | 87.0 | 29.8 | 6.8 | 5.2 |
| 25.9 | 6.6 | 5.3 | 28.3 | 82.7 | 28.9 | 6.7 | 5.1 |
| 29.9 | 6.9 | 5.3 | 28.9 | 86.2 | 27.0 | 6.7 | 5.1 |
| 27.3 | 6.8 | 5.6 | 29.3 | 83.2 | 31.4 | 6.8 | 4.1 |
| 27.6 | 6.8 | 5.5 | 29.3 | 86.2 | 28.9 | 6.8 | 5.4 |
| 29.3 | 6.8 | 5.2 | 28.7 | 86.6 | 29.3 | 6.7 | 5.2 |
| 29.0 | 6.8 | 5.5 | 29.4 | 86.0 | 29.8 | 6.8 | 5.0 |
| 29.6 | 6.9 | 5.4 | 27.7 | 84.9 | 27.2 | 6.6 | 4.6 |
| 28.9 | 6.8 | 4.4 | 29.1 | 86.2 | 28.6 | 6.7 | 4.6 |
| 26.5 | 6.8 | 5.1 | 28.9 | 85.5 | 28.0 | 6.7 | 4.8 |
| 30.8 | 6.9 | 4.5 | 30.6 | 84.5 | 31.7 | 6.9 | 4.6 |
| 27.2 | 6.8 | 4.9 | 31.6 | 87.2 | 31.9 | 6.9 | 4.5 |
| 29.6 | 6.9 | 5.4 | 27.7 | 84.9 | 27.2 | 6.6 | 4.6 |
| 27.5 | 6.8 | 4.9 | 29.3 | 84.8 | 28.5 | 6.7 | 4.5 |
| 29.3 | 6.9 | 4.6 | 29.5 | 87.8 | 30.1 | 6.8 | 4.6 |
| 28.8 | 6.8 | 4.9 | 29.6 | 87.8 | 29.5 | 6.8 | 4.7 |
| 30.5 | 6.8 | 4.3 | 29.5 | 86.9 | 30.8 | 6.7 | 4.3 |
| 29.6 | 6.9 | 5.4 | 27.7 | 84.9 | 27.2 | 6.6 | 4.6 |
| 28.4 | 6.8 | 4.3 | 29.4 | 86.4 | 29.8 | 6.8 | 4.7 |
| 29.9 | 6.9 | 4.9 | 31.5 | 86.8 | 31.4 | 6.8 | 4.6 |
| 29.1 | 6.8 | 5.6 | 29.2 | 85.7 | 30.7 | 6.7 | 4.9 |
| 28.8 | 6.9 | 5.6 | 29.2 | 86.7 | 27.8 | 6.7 | 5.1 |
| 29.6 | 6.9 | 5.4 | 27.7 | 84.9 | 27.2 | 6.6 | 4.6 |
| 28.2 | 6.8 | 5.3 | 28.8 | 84.6 | 28.0 | 6.7 | 5.1 |
| 27.3 | 6.8 | 4.8 | 27.5 | 82.8 | 26.6 | 6.6 | 5.2 |
| 26.9 | 6.7 | 4.2 | 28.6 | 86.3 | 28.9 | 6.7 | 4.7 |
| 28.8 | 6.8 | 4.4 | 29.1 | 84.4 | 30.2 | 6.8 | 4.6 |
| 29.6 | 6.9 | 5.4 | 27.7 | 84.9 | 27.2 | 6.6 | 4.6 |
| 29.1 | 6.7 | 4.8 | 29.7 | 85.3 | 31.9 | 6.8 | 4.2 |
| 28.9 | 6.7 | 5.4 | 28.5 | 84.0 | 30.3 | 6.7 | 5.1 |
| 27.1 | 6.8 | 5.1 | 29.4 | 84.6 | 31.7 | 6.8 | 4.6 |
| 31.4 | 6.9 | 3.9 | 29.7 | 85.6 | 30.9 | 6.8 | 4.3 |
| 29.6 | 6.9 | 5.4 | 27.7 | 84.9 | 27.2 | 6.6 | 4.6 |
| 29.8 | 6.8 | 4.4 | 30.2 | 87.4 | 31.9 | 6.9 | 4.5 |
| 29.4 | 7.0 | 5.4 | 29.1 | 85.9 | 29.8 | 6.8 | 4.8 |
| 29.1 | 6.9 | 4.9 | 28.1 | 85.3 | 28.8 | 6.7 | 5.4 |
| 29.4 | 6.9 | 4.6 | 29.3 | 84.1 | 32.0 | 6.9 | 5.2 |
| 29.6 | 6.9 | 5.4 | 27.7 | 84.9 | 27.2 | 6.6 | 4.6 |
| 28.0 | 6.8 | 4.9 | 29.4 | 84.0 | 30.3 | 6.8 | 4.7 |
| 29.5 | 6.9 | 5.0 | 27.9 | 84.0 | 30.3 | 6.7 | 5.0 |
| 29.0 | 7.0 | 5.6 | 29.9 | 84.5 | 30.5 | 6.9 | 5.5 |
| 27.2 | 7.0 | 5.5 | 29.9 | 86.5 | 29.4 | 6.7 | 5.1 |
| 29.6 | 6.9 | 5.4 | 27.7 | 84.9 | 27.2 | 6.6 | 4.6 |
| 28.2 | 6.9 | 5.1 | 27.8 | 82.3 | 29.2 | 6.7 | 4.8 |
| 26.9 | 6.8 | 4.8 | 30.2 | 86.9 | 31.3 | 6.9 | 4.6 |
| 27.4 | 6.9 | 5.5 | 26.6 | 84.3 | 26.5 | 6.6 | 5.2 |
| 28.1 | 6.8 | 4.5 | 29.6 | 85.2 | 29.1 | 6.7 | 4.7 |
| 29.6 | 6.9 | 5.4 | 27.7 | 84.9 | 27.2 | 6.6 | 4.6 |
| 30.2 | 6.9 | 4.9 | 29.6 | 85.0 | 31.2 | 6.8 | 4.7 |
| 27.0 | 6.6 | 5.0 | 27.2 | 82.7 | 26.9 | 6.6 | 5.5 |
| 30.6 | 6.8 | 4.4 | 29.1 | 84.6 | 31.3 | 6.7 | 4.3 |
| 29.5 | 6.8 | 5.1 | 29.4 | 86.4 | 30.5 | 6.8 | 4.8 |
| 29.6 | 6.9 | 5.4 | 27.7 | 84.9 | 27.2 | 6.6 | 4.6 |
| 28.3 | 6.7 | 4.9 | 28.6 | 85.3 | 27.5 | 6.7 | 5.2 |
| 28.3 | 6.7 | 4.8 | 28.0 | 82.8 | 27.0 | 6.6 | 5.4 |
| 28.0 | 6.8 | 5.3 | 28.7 | 84.3 | 28.6 | 6.6 | 5.0 |
| 30.5 | 6.9 | 4.7 | 29.4 | 86.4 | 30.1 | 6.8 | 4.5 |
| 29.6 | 6.9 | 4.7 | 29.6 | 85.7 | 29.4 | 6.8 | 4.8 |
| 29.4 | 6.8 | 4.9 | 28.8 | 84.8 | 28.6 | 6.7 | 4.6 |
| 28.8 | 6.8 | 4.7 | 30.4 | 86.2 | 31.1 | 6.9 | 5.0 |
| 30.1 | 6.9 | 5.4 | 28.5 | 84.4 | 28.7 | 6.6 | 4.5 |
| 28.5 | 6.8 | 5.0 | 29.4 | 86.4 | 31.9 | 6.8 | 5.3 |
| 29.6 | 6.9 | 4.7 | 29.6 | 85.7 | 29.4 | 6.8 | 4.8 |
| 27.6 | 6.8 | 5.6 | 29.2 | 84.6 | 30.3 | 6.7 | 4.7 |
| 29.5 | 6.9 | 5.6 | 27.6 | 82.9 | 27.2 | 6.6 | 5.6 |
| 25.1 | 6.6 | 5.8 | 28.0 | 84.8 | 28.3 | 6.7 | 5.1 |
| 27.3 | 6.7 | 4.9 | 29.4 | 87.3 | 29.0 | 6.8 | 5.1 |
| 29.6 | 6.9 | 4.7 | 29.6 | 85.7 | 29.4 | 6.8 | 4.8 |
| 27.0 | 6.7 | 5.6 | 28.5 | 85.2 | 29.6 | 6.8 | 5.1 |
| 29.9 | 6.9 | 5.5 | 29.1 | 84.7 | 30.6 | 6.8 | 4.9 |
| 28.7 | 6.7 | 4.8 | 30.0 | 85.2 | 31.5 | 6.7 | 4.1 |
| 30.6 | 6.8 | 4.9 | 29.9 | 85.0 | 30.6 | 6.7 | 4.8 |
| 29.6 | 6.9 | 4.7 | 29.6 | 85.7 | 29.4 | 6.8 | 4.8 |
| 29.9 | 6.9 | 4.6 | 30.1 | 83.0 | 29.5 | 6.8 | 4.5 |
| 28.0 | 6.8 | 4.5 | 30.2 | 86.6 | 30.0 | 6.8 | 4.8 |
| 28.7 | 6.9 | 4.9 | 30.5 | 83.9 | 30.0 | 6.8 | 4.7 |
| 26.9 | 6.8 | 4.5 | 30.2 | 83.0 | 32.5 | 6.9 | 4.7 |
| 29.6 | 6.9 | 4.7 | 29.6 | 85.7 | 29.4 | 6.8 | 4.8 |
| 30.1 | 6.9 | 4.5 | 30.9 | 86.6 | 30.7 | 6.8 | 4.8 |
| 30.6 | 6.9 | 4.6 | 29.4 | 83.6 | 30.7 | 6.7 | 3.9 |
| 27.5 | 6.7 | 3.9 | 30.4 | 83.3 | 30.4 | 6.8 | 4.2 |
| 30.1 | 6.8 | 4.8 | 30.5 | 86.2 | 30.3 | 6.8 | 4.3 |
| 29.6 | 6.9 | 4.7 | 29.6 | 85.7 | 29.4 | 6.8 | 4.8 |
| 28.5 | 6.8 | 4.1 | 28.5 | 84.3 | 28.7 | 6.8 | 5.3 |
| 28.9 | 6.8 | 4.9 | 28.8 | 85.3 | 27.2 | 6.7 | 5.1 |
| 29.7 | 6.8 | 4.7 | 30.6 | 86.7 | 31.7 | 6.9 | 5.4 |
| 30.2 | 6.8 | 4.6 | 30.4 | 84.6 | 31.2 | 6.8 | 5.0 |
| 29.6 | 6.9 | 4.7 | 29.6 | 85.7 | 29.4 | 6.8 | 4.8 |
| 27.5 | 6.8 | 4.7 | 28.0 | 82.3 | 28.6 | 6.7 | 4.9 |
| 29.4 | 6.8 | 5.1 | 27.9 | 84.8 | 27.4 | 6.6 | 5.2 |
| 29.7 | 6.8 | 5.0 | 30.8 | 85.4 | 30.4 | 6.8 | 4.7 |
| 30.1 | 6.8 | 3.5 | 29.6 | 83.6 | 27.9 | 6.7 | 5.0 |
| 29.6 | 6.9 | 4.7 | 29.6 | 85.7 | 29.4 | 6.8 | 4.8 |
| 29.3 | 6.9 | 5.4 | 30.5 | 84.9 | 29.2 | 6.8 | 5.0 |
| 29.4 | 6.8 | 5.3 | 30.8 | 86.2 | 30.2 | 6.8 | 4.8 |
| 28.5 | 6.7 | 5.2 | 31.5 | 83.8 | 31.3 | 6.8 | 4.5 |
| 30.0 | 6.8 | 4.7 | 28.0 | 77.9 | 29.4 | 6.7 | 4.8 |
| 29.6 | 6.9 | 4.7 | 29.6 | 85.7 | 29.4 | 6.8 | 4.8 |
| 27.1 | 6.8 | 5.0 | 29.5 | 85.8 | 28.5 | 6.7 | 5.0 |
| 28.0 | 6.8 | 5.4 | 29.5 | 84.2 | 28.9 | 6.7 | 4.6 |
| 28.6 | 6.6 | 4.3 | 29.1 | 84.8 | 28.6 | 6.7 | 5.0 |
| 28.3 | 6.9 | 4.8 | 30.5 | 87.0 | 30.9 | 6.8 | 4.5 |
| 29.6 | 6.9 | 4.7 | 29.6 | 85.7 | 29.4 | 6.8 | 4.8 |
| 31.9 | 6.9 | 4.9 | 30.5 | 86.3 | 29.2 | 6.8 | 5.2 |
| 28.4 | 6.9 | 5.5 | 30.0 | 84.9 | 29.4 | 6.7 | 5.3 |
| 30.9 | 6.8 | 5.0 | 30.3 | 87.9 | 30.4 | 6.8 | 4.5 |
| 28.9 | 6.7 | 5.2 | 31.6 | 84.6 | 30.3 | 6.9 | 4.4 |
| 27.5 | 6.7 | 4.8 | 29.9 | 83.8 | 27.7 | 6.7 | 5.0 |
| 26.9 | 6.8 | 4.9 | 29.4 | 84.9 | 28.4 | 6.7 | 4.8 |
| 30.3 | 6.9 | 5.4 | 28.2 | 82.3 | 28.2 | 6.7 | 4.7 |
| 28.3 | 6.7 | 4.6 | 28.8 | 82.7 | 30.0 | 6.7 | 4.4 |
| 29.4 | 6.7 | 4.9 | 29.8 | 83.5 | 28.9 | 6.8 | 5.1 |
| 27.5 | 6.7 | 4.8 | 29.9 | 83.8 | 27.7 | 6.7 | 5.0 |
| 29.1 | 6.8 | 5.2 | 30.0 | 85.0 | 28.9 | 6.8 | 4.8 |
| 25.2 | 6.6 | 2.9 | 30.7 | 85.9 | 28.8 | 6.8 | 5.4 |
| 28.5 | 6.8 | 4.7 | 29.5 | 84.9 | 27.3 | 6.7 | 4.5 |
| 30.5 | 6.8 | 4.0 | 29.7 | 85.7 | 30.1 | 6.8 | 4.4 |
| 27.5 | 6.7 | 4.8 | 29.9 | 83.8 | 27.7 | 6.7 | 5.0 |
| 29.5 | 6.8 | 5.0 | 28.2 | 84.6 | 28.5 | 6.7 | 4.7 |
| 27.2 | 6.8 | 4.7 | 28.7 | 82.6 | 27.0 | 6.7 | 4.8 |
| 28.0 | 6.8 | 5.2 | 28.6 | 84.2 | 28.5 | 6.7 | 4.5 |
| 28.1 | 7.0 | 5.3 | 29.8 | 82.0 | 29.7 | 6.8 | 5.1 |
| 27.5 | 6.7 | 4.8 | 29.9 | 83.8 | 27.7 | 6.7 | 5.0 |
| 28.1 | 6.9 | 5.4 | 28.2 | 85.2 | 28.1 | 6.7 | 4.9 |
| 31.0 | 6.9 | 4.4 | 29.8 | 86.3 | 27.8 | 6.7 | 5.2 |
| 29.8 | 6.9 | 4.3 | 29.5 | 87.1 | 28.3 | 6.7 | 5.2 |
| 28.7 | 6.8 | 4.9 | 27.8 | 82.3 | 28.0 | 6.7 | 5.4 |
| 27.5 | 6.7 | 4.8 | 29.9 | 83.8 | 27.7 | 6.7 | 5.0 |
| 27.7 | 6.7 | 4.0 | 28.6 | 85.2 | 28.3 | 6.7 | 5.4 |
| 30.0 | 6.8 | 4.3 | 27.9 | 81.5 | 27.6 | 6.7 | 5.0 |
| 29.2 | 6.9 | 5.4 | 29.3 | 85.0 | 28.5 | 6.7 | 5.7 |
| 28.5 | 6.8 | 4.5 | 29.7 | 83.6 | 29.7 | 6.8 | 5.1 |
| 27.5 | 6.7 | 4.8 | 29.9 | 83.8 | 27.7 | 6.7 | 5.0 |
| 29.3 | 6.8 | 5.0 | 28.7 | 85.2 | 29.9 | 6.7 | 5.1 |
| 27.7 | 6.8 | 5.1 | 27.6 | 83.9 | 27.4 | 6.6 | 5.2 |
| 28.8 | 6.8 | 5.3 | 27.6 | 85.1 | 27.6 | 6.6 | 5.7 |
| 31.5 | 6.9 | 4.2 | 27.9 | 83.1 | 27.0 | 6.6 | 5.9 |
| 27.5 | 6.7 | 4.8 | 29.9 | 83.8 | 27.7 | 6.7 | 5.0 |
| 30.4 | 6.7 | 4.2 | 28.8 | 83.8 | 28.2 | 6.7 | 5.3 |
| 30.8 | 6.8 | 4.5 | 29.5 | 87.6 | 28.7 | 6.7 | 5.2 |
| 30.5 | 6.8 | 5.0 | 27.9 | 83.5 | 28.2 | 6.6 | 5.1 |
| 28.4 | 6.8 | 5.4 | 29.2 | 80.8 | 28.9 | 6.7 | 5.4 |
| 27.5 | 6.7 | 4.8 | 29.9 | 83.8 | 27.7 | 6.7 | 5.0 |
| 28.4 | 6.7 | 4.8 | 29.9 | 87.8 | 30.0 | 6.8 | 5.0 |
| 28.4 | 6.8 | 4.7 | 29.0 | 82.7 | 28.9 | 6.7 | 4.8 |
| 28.3 | 6.8 | 4.5 | 30.9 | 84.8 | 29.2 | 6.8 | 4.5 |
| 30.7 | 6.8 | 4.1 | 31.3 | 87.2 | 30.8 | 6.9 | 4.6 |
| 27.5 | 6.7 | 4.8 | 29.9 | 83.8 | 27.7 | 6.7 | 5.0 |
| 27.0 | 6.7 | 4.9 | 29.9 | 83.9 | 28.3 | 6.7 | 4.8 |
| 22.9 | 6.4 | 4.4 | 28.1 | 84.7 | 28.2 | 6.7 | 4.7 |
| 30.1 | 6.9 | 4.6 | 29.8 | 82.5 | 30.1 | 6.8 | 4.9 |
| 29.9 | 6.8 | 4.1 | 30.1 | 83.5 | 30.2 | 6.8 | 4.8 |
| 27.5 | 6.7 | 4.8 | 29.9 | 83.8 | 27.7 | 6.7 | 5.0 |
| 29.7 | 6.8 | 4.5 | 30.4 | 86.1 | 30.2 | 6.8 | 4.4 |
| 32.1 | 6.8 | 4.2 | 30.1 | 84.0 | 30.2 | 6.8 | 5.1 |
| 28.5 | 6.8 | 4.9 | 29.6 | 82.1 | 30.2 | 6.8 | 4.9 |
| 28.2 | 6.7 | 5.1 | 31.1 | 83.7 | 32.6 | 6.9 | 4.7 |
| 28.4 | 6.8 | 5.1 | 29.1 | 83.9 | 28.9 | 6.7 | 4.7 |
| 27.3 | 6.7 | 5.8 | 28.5 | 85.4 | 29.7 | 6.7 | 4.8 |
| 28.3 | 6.7 | 5.8 | 27.7 | 84.0 | 27.4 | 6.7 | 5.6 |
| 29.7 | 6.8 | 4.3 | 30.5 | 87.8 | 29.4 | 6.8 | 4.8 |
| 29.9 | 6.8 | 4.4 | 31.1 | 86.4 | 30.2 | 6.9 | 5.4 |
| 28.4 | 6.8 | 5.1 | 29.1 | 83.9 | 28.9 | 6.7 | 4.7 |
| 29.2 | 6.7 | 5.3 | 29.1 | 84.8 | 32.8 | 6.8 | 4.9 |
| 29.3 | 6.7 | 5.2 | 27.6 | 83.5 | 29.4 | 6.7 | 4.9 |
| 31.0 | 6.7 | 4.5 | 32.1 | 85.5 | 32.8 | 6.8 | 4.5 |
| 29.7 | 6.7 | 4.4 | 28.5 | 79.5 | 30.7 | 6.7 | 4.8 |
| 28.4 | 6.8 | 5.1 | 29.1 | 83.9 | 28.9 | 6.7 | 4.7 |
| 27.7 | 6.7 | 5.1 | 29.5 | 81.9 | 30.5 | 6.8 | 5.0 |
| 29.7 | 6.8 | 4.5 | 29.9 | 85.7 | 33.3 | 6.8 | 3.8 |
| 29.1 | 6.8 | 4.7 | 30.1 | 85.2 | 30.1 | 6.7 | 4.9 |
| 30.7 | 6.8 | 4.3 | 30.9 | 83.8 | 29.9 | 6.8 | 5.1 |
| 28.4 | 6.8 | 5.1 | 29.1 | 83.9 | 28.9 | 6.7 | 4.7 |
| 27.8 | 6.8 | 4.4 | 29.2 | 86.1 | 28.7 | 6.7 | 4.6 |
| 26.8 | 6.7 | 5.9 | 26.6 | 82.3 | 26.0 | 6.5 | 5.7 |
| 27.9 | 6.7 | 5.2 | 28.1 | 85.5 | 28.3 | 6.6 | 4.9 |
| 27.9 | 6.7 | 5.1 | 26.9 | 86.0 | 26.8 | 6.5 | 4.9 |
| 28.4 | 6.8 | 5.1 | 29.1 | 83.9 | 28.9 | 6.7 | 4.7 |
| 29.3 | 6.8 | 4.1 | 30.1 | 86.7 | 28.8 | 6.8 | 4.6 |
| 29.9 | 6.8 | 4.0 | 29.1 | 84.9 | 28.3 | 6.8 | 5.2 |
| 28.2 | 6.6 | 4.8 | 28.3 | 83.3 | 30.3 | 6.7 | 4.3 |
| 28.9 | 6.8 | 4.9 | 29.0 | 86.8 | 31.1 | 6.7 | 4.7 |
| 28.4 | 6.8 | 5.1 | 29.1 | 83.9 | 28.9 | 6.7 | 4.7 |
| 29.7 | 6.7 | 4.6 | 30.1 | 85.9 | 30.9 | 6.8 | 4.6 |
| 26.6 | 6.7 | 4.8 | 29.6 | 83.1 | 28.4 | 6.7 | 4.4 |
| 31.3 | 6.8 | 4.2 | 28.1 | 81.6 | 29.3 | 6.7 | 4.5 |
| 29.3 | 6.7 | 3.9 | 29.2 | 86.5 | 27.6 | 6.7 | 4.7 |
| 28.4 | 6.8 | 5.1 | 29.1 | 83.9 | 28.9 | 6.7 | 4.7 |
| 26.5 | 6.7 | 4.5 | 30.2 | 85.0 | 28.8 | 6.7 | 4.5 |
| 24.4 | 6.4 | 4.3 | 28.0 | 84.4 | 27.7 | 6.7 | 5.3 |
| 27.3 | 6.7 | 4.9 | 28.2 | 85.5 | 29.4 | 6.6 | 4.3 |
| 30.0 | 6.9 | 4.5 | 30.1 | 84.4 | 29.4 | 6.8 | 4.8 |
| 28.4 | 6.8 | 5.1 | 29.1 | 83.9 | 28.9 | 6.7 | 4.7 |
| 29.4 | 6.9 | 5.1 | 30.3 | 86.3 | 30.8 | 6.8 | 4.9 |
| 29.8 | 6.8 | 4.8 | 31.8 | 85.6 | 31.0 | 6.7 | 4.5 |
| 28.1 | 6.8 | 4.8 | 27.9 | 83.4 | 28.6 | 6.7 | 4.3 |
| 28.4 | 6.7 | 4.6 | 30.0 | 85.9 | 31.1 | 6.8 | 4.2 |
| 28.4 | 6.8 | 5.1 | 29.1 | 83.9 | 28.9 | 6.7 | 4.7 |
| 29.4 | 6.7 | 4.0 | 30.9 | 85.9 | 28.9 | 6.7 | 4.8 |
| 31.0 | 6.8 | 4.3 | 29.3 | 84.6 | 30.4 | 6.7 | 4.6 |
| 29.4 | 6.9 | 5.2 | 30.5 | 86.7 | 28.9 | 6.8 | 5.3 |
| 28.9 | 6.8 | 4.9 | 30.9 | 87.3 | 30.6 | 6.8 | 5.2 |
| 28.4 | 6.8 | 5.1 | 29.1 | 83.9 | 28.9 | 6.7 | 4.7 |
| 27.3 | 6.7 | 4.8 | 28.8 | 83.2 | 27.7 | 6.6 | 4.6 |
| 27.9 | 6.9 | 5.3 | 30.7 | 85.4 | 31.8 | 6.8 | 4.5 |
| 29.6 | 6.8 | 4.6 | 29.8 | 85.8 | 30.0 | 6.7 | 5.1 |
| 28.7 | 6.7 | 5.2 | 28.4 | 85.6 | 28.2 | 6.6 | 5.3 |
| 28.1 | 6.7 | 3.9 | 29.3 | 86.1 | 28.0 | 6.7 | 5.5 |
| 28.1 | 6.7 | 4.7 | 28.6 | 86.1 | 26.8 | 6.7 | 5.5 |
| 29.8 | 6.8 | 4.8 | 28.2 | 83.9 | 27.6 | 6.7 | 4.8 |
| 29.4 | 6.8 | 5.0 | 28.7 | 85.7 | 27.0 | 6.7 | 5.8 |
| 29.7 | 6.8 | 5.1 | 29.6 | 85.3 | 29.5 | 6.7 | 5.6 |
| 30.5 | 6.9 | 5.0 | 28.4 | 85.4 | 27.8 | 6.6 | 5.6 |
| 28.1 | 6.7 | 4.7 | 28.6 | 86.1 | 26.8 | 6.7 | 5.5 |
| 28.4 | 6.8 | 4.7 | 29.0 | 85.9 | 28.3 | 6.7 | 5.3 |
| 28.7 | 6.9 | 5.1 | 29.1 | 84.5 | 28.1 | 6.6 | 4.8 |
| 27.0 | 6.8 | 5.2 | 27.5 | 82.2 | 27.9 | 6.7 | 5.7 |
| 27.6 | 6.8 | 4.9 | 27.2 | 84.3 | 25.9 | 6.6 | 5.1 |
| 28.1 | 6.7 | 4.7 | 28.6 | 86.1 | 26.8 | 6.7 | 5.5 |
| 29.8 | 6.9 | 4.5 | 29.2 | 88.0 | 27.0 | 6.7 | 5.4 |
| 27.8 | 6.9 | 5.1 | 27.4 | 86.0 | 26.2 | 6.6 | 5.1 |
| 28.5 | 6.9 | 5.5 | 29.8 | 85.4 | 29.7 | 6.8 | 4.8 |
| 28.7 | 6.8 | 5.1 | 28.1 | 85.5 | 27.4 | 6.6 | 5.4 |
| 28.1 | 6.7 | 4.7 | 28.6 | 86.1 | 26.8 | 6.7 | 5.5 |
| 27.6 | 6.8 | 4.8 | 30.2 | 86.3 | 29.7 | 6.8 | 5.0 |
| 29.5 | 7.0 | 5.2 | 30.1 | 85.5 | 29.7 | 6.8 | 5.7 |
| 29.4 | 6.8 | 4.6 | 28.7 | 85.1 | 27.0 | 6.6 | 5.6 |
| 30.6 | 6.9 | 5.0 | 29.1 | 84.5 | 28.8 | 6.7 | 5.5 |
| 28.1 | 6.7 | 4.7 | 28.6 | 86.1 | 26.8 | 6.7 | 5.5 |
| 29.8 | 6.8 | 4.4 | 30.2 | 85.7 | 30.6 | 6.8 | 4.5 |
| 29.7 | 6.8 | 4.6 | 29.0 | 85.5 | 31.5 | 6.8 | 4.8 |
| 29.4 | 6.8 | 4.3 | 28.9 | 85.2 | 27.0 | 6.7 | 5.2 |
| 28.0 | 6.9 | 5.2 | 27.8 | 83.8 | 28.1 | 6.7 | 4.9 |
| 28.1 | 6.7 | 4.7 | 28.6 | 86.1 | 26.8 | 6.7 | 5.5 |
| 28.3 | 6.8 | 4.7 | 29.5 | 85.6 | 29.2 | 6.7 | 4.8 |
| 30.1 | 6.8 | 4.3 | 29.1 | 83.7 | 27.9 | 6.7 | 4.5 |
| 27.7 | 6.8 | 4.6 | 27.0 | 84.3 | 25.3 | 6.6 | 5.0 |
| 26.7 | 6.7 | 5.3 | 26.5 | 84.4 | 25.7 | 6.6 | 5.1 |
| 28.1 | 6.7 | 4.7 | 28.6 | 86.1 | 26.8 | 6.7 | 5.5 |
| 27.4 | 6.8 | 5.1 | 28.9 | 85.0 | 28.5 | 6.7 | 4.9 |
| 30.3 | 6.8 | 5.4 | 29.4 | 85.2 | 29.3 | 6.8 | 5.1 |
| 31.4 | 7.0 | 4.5 | 31.5 | 85.9 | 32.0 | 6.9 | 4.7 |
| 28.8 | 6.8 | 4.7 | 30.3 | 86.9 | 28.4 | 6.8 | 4.8 |
| 28.1 | 6.7 | 4.7 | 28.6 | 86.1 | 26.8 | 6.7 | 5.5 |
| 30.7 | 6.8 | 4.5 | 30.8 | 85.4 | 29.9 | 6.8 | 5.0 |
| 32.6 | 6.9 | 4.0 | 29.4 | 84.0 | 32.3 | 6.7 | 3.9 |
| 30.1 | 6.8 | 4.8 | 27.3 | 83.0 | 29.2 | 6.7 | 4.9 |
| 30.3 | 6.9 | 4.7 | 28.0 | 82.0 | 28.9 | 6.7 | 4.6 |
| 28.1 | 6.7 | 4.7 | 28.6 | 86.1 | 26.8 | 6.7 | 5.5 |
| 30.7 | 6.9 | 4.5 | 28.1 | 82.9 | 29.9 | 6.6 | 3.7 |
| 29.6 | 6.8 | 3.2 | 29.4 | 83.4 | 29.1 | 6.7 | 4.7 |
| 30.5 | 6.8 | 4.5 | 29.1 | 84.6 | 32.1 | 6.8 | 5.0 |
| 29.7 | 6.8 | 3.8 | 29.5 | 85.8 | 31.0 | 6.8 | 4.8 |
| 28.1 | 6.7 | 4.7 | 28.6 | 86.1 | 26.8 | 6.7 | 5.5 |
| 31.4 | 7.0 | 4.4 | 28.5 | 81.3 | 28.9 | 6.7 | 4.6 |
| 31.0 | 6.9 | 4.2 | 30.7 | 86.4 | 35.4 | 7.0 | 4.1 |
| 29.6 | 6.7 | 3.5 | 29.4 | 86.0 | 29.3 | 6.8 | 4.6 |
| 30.3 | 6.9 | 4.3 | 28.4 | 84.9 | 27.7 | 6.7 | 5.0 |
| 29.3 | 6.8 | 5.0 | 28.4 | 85.7 | 27.8 | 6.7 | 4.9 |
| 29.6 | 6.9 | 4.8 | 28.4 | 85.0 | 28.4 | 6.7 | 5.0 |
| 31.4 | 7.0 | 4.9 | 29.8 | 87.6 | 30.7 | 6.8 | 4.8 |
| 30.2 | 6.9 | 5.0 | 30.3 | 87.0 | 29.8 | 6.8 | 5.0 |
| 29.7 | 6.8 | 4.6 | 29.5 | 85.8 | 31.0 | 6.8 | 4.6 |
| 29.3 | 6.8 | 5.0 | 28.4 | 85.7 | 27.8 | 6.7 | 4.9 |
| 28.0 | 6.9 | 5.1 | 28.2 | 84.1 | 28.2 | 6.7 | 4.7 |
| 29.1 | 6.8 | 5.0 | 28.8 | 84.6 | 28.8 | 6.8 | 5.1 |
| 27.4 | 6.9 | 5.0 | 28.4 | 84.4 | 28.0 | 6.7 | 4.7 |
| 29.1 | 6.8 | 5.3 | 29.2 | 84.4 | 28.0 | 6.7 | 5.0 |
| 29.3 | 6.8 | 5.0 | 28.4 | 85.7 | 27.8 | 6.7 | 4.9 |
| 28.9 | 6.8 | 4.9 | 26.7 | 81.6 | 26.9 | 6.6 | 4.9 |
| 29.1 | 6.9 | 5.3 | 29.0 | 85.1 | 30.0 | 6.8 | 4.8 |
| 28.7 | 6.9 | 4.6 | 30.5 | 85.9 | 29.6 | 6.8 | 4.6 |
| 28.7 | 6.9 | 4.6 | 27.2 | 83.6 | 28.1 | 6.6 | 4.4 |
| 29.3 | 6.8 | 5.0 | 28.4 | 85.7 | 27.8 | 6.7 | 4.9 |
| 28.8 | 6.9 | 4.5 | 29.6 | 84.0 | 30.3 | 6.8 | 4.5 |
| 29.8 | 6.9 | 4.3 | 29.2 | 83.7 | 28.8 | 6.7 | 4.2 |
| 29.0 | 6.9 | 4.6 | 29.0 | 85.5 | 30.0 | 6.7 | 4.8 |
| 30.4 | 6.8 | 4.0 | 28.7 | 86.2 | 28.3 | 6.7 | 4.8 |
| 29.3 | 6.8 | 5.0 | 28.4 | 85.7 | 27.8 | 6.7 | 4.9 |
| 28.3 | 6.9 | 5.3 | 28.4 | 85.7 | 27.8 | 6.7 | 5.1 |
| 30.1 | 6.8 | 4.8 | 26.1 | 85.1 | 27.1 | 6.6 | 5.6 |
| 27.0 | 6.6 | 5.9 | 24.2 | 82.8 | 23.9 | 6.3 | 5.9 |
| 27.5 | 6.7 | 5.7 | 29.2 | 86.9 | 28.4 | 6.7 | 5.0 |
| 29.3 | 6.8 | 5.0 | 28.4 | 85.7 | 27.8 | 6.7 | 4.9 |
| 28.8 | 6.8 | 4.9 | 30.3 | 87.1 | 28.8 | 6.8 | 5.4 |
| 29.9 | 7.0 | 5.3 | 28.3 | 83.5 | 29.0 | 6.7 | 5.5 |
| 30.0 | 6.6 | 4.5 | 28.8 | 84.0 | 29.7 | 6.6 | 4.5 |
| 30.5 | 6.8 | 4.9 | 30.0 | 86.4 | 29.3 | 6.7 | 4.9 |
| 29.3 | 6.8 | 5.0 | 28.4 | 85.7 | 27.8 | 6.7 | 4.9 |
| 29.4 | 6.7 | 4.6 | 28.8 | 86.1 | 27.7 | 6.7 | 4.8 |
| 27.0 | 6.7 | 4.6 | 25.0 | 80.1 | 24.7 | 6.5 | 5.4 |
| 29.9 | 6.8 | 4.6 | 28.8 | 83.7 | 28.7 | 6.6 | 5.1 |
| 29.7 | 6.8 | 4.8 | 28.0 | 85.4 | 28.3 | 6.7 | 5.2 |
| 29.3 | 6.8 | 5.0 | 28.4 | 85.7 | 27.8 | 6.7 | 4.9 |
| 28.3 | 6.9 | 5.0 | 29.2 | 87.3 | 27.0 | 6.7 | 5.6 |
| 26.9 | 6.7 | 4.9 | 29.2 | 84.5 | 30.0 | 6.7 | 5.2 |
| 27.4 | 6.8 | 5.3 | 25.7 | 82.9 | 25.5 | 6.5 | 5.7 |
| 26.5 | 6.8 | 5.2 | 26.9 | 84.9 | 28.0 | 6.6 | 5.6 |
| 29.3 | 6.8 | 5.0 | 28.4 | 85.7 | 27.8 | 6.7 | 4.9 |
| 28.0 | 6.7 | 4.9 | 28.1 | 85.1 | 28.2 | 6.7 | 4.8 |
| 29.1 | 6.8 | 5.1 | 28.7 | 85.9 | 29.3 | 6.7 | 4.6 |
| 29.7 | 6.9 | 4.7 | 30.3 | 85.3 | 29.9 | 6.8 | 4.9 |
| 29.7 | 6.8 | 4.1 | 29.2 | 85.1 | 28.4 | 6.7 | 5.0 |
| 29.3 | 6.8 | 5.0 | 28.4 | 85.7 | 27.8 | 6.7 | 4.9 |
| 28.0 | 6.8 | 4.9 | 28.9 | 86.5 | 27.7 | 6.8 | 5.5 |
| 27.5 | 6.7 | 5.3 | 28.1 | 84.8 | 26.6 | 6.7 | 5.5 |
| 28.4 | 6.7 | 4.6 | 27.3 | 84.7 | 26.2 | 6.6 | 5.7 |
| 30.4 | 6.7 | 4.5 | 28.4 | 85.8 | 27.5 | 6.7 | 5.3 |
| 25.8 | 6.6 | 4.1 | 28.5 | 86.2 | 27.1 | 6.7 | 5.0 |
| 28.6 | 6.8 | 5.0 | 29.1 | 87.6 | 28.4 | 6.7 | 4.9 |
| 27.1 | 6.9 | 5.9 | 28.9 | 87.6 | 27.2 | 6.8 | 5.7 |
| 30.2 | 6.9 | 5.0 | 30.6 | 87.5 | 29.3 | 6.8 | 5.0 |
| 30.5 | 6.9 | 4.6 | 30.2 | 86.3 | 28.5 | 6.7 | 5.2 |
| 25.8 | 6.6 | 4.1 | 28.5 | 86.2 | 27.1 | 6.7 | 5.0 |
| 28.7 | 6.7 | 4.9 | 29.6 | 85.5 | 28.4 | 6.7 | 4.5 |
| 27.8 | 6.7 | 5.0 | 26.8 | 84.2 | 25.7 | 6.6 | 5.1 |
| 33.6 | 6.9 | 5.1 | 29.3 | 85.3 | 29.2 | 6.7 | 5.7 |
| 31.2 | 6.8 | 4.8 | 29.7 | 86.1 | 29.3 | 6.7 | 5.1 |
| 25.8 | 6.6 | 4.1 | 28.5 | 86.2 | 27.1 | 6.7 | 5.0 |
| 31.5 | 6.8 | 4.5 | 27.8 | 82.9 | 27.3 | 6.6 | 5.0 |
| 30.1 | 6.8 | 3.3 | 25.8 | 82.3 | 24.1 | 6.5 | 4.9 |
| 30.2 | 6.8 | 3.6 | 27.1 | 84.4 | 27.5 | 6.6 | 4.7 |
| 29.3 | 6.8 | 4.3 | 29.5 | 86.6 | 28.1 | 6.7 | 4.5 |
| 25.8 | 6.6 | 4.1 | 28.5 | 86.2 | 27.1 | 6.7 | 5.0 |
| 29.4 | 6.8 | 5.3 | 28.5 | 85.6 | 29.0 | 6.7 | 5.0 |
| 31.3 | 6.8 | 5.2 | 30.3 | 86.9 | 29.4 | 6.8 | 4.8 |
| 31.7 | 6.8 | 5.1 | 29.2 | 85.3 | 28.6 | 6.7 | 5.3 |
| 31.6 | 6.9 | 4.5 | 28.5 | 85.3 | 28.2 | 6.6 | 4.7 |
| 25.8 | 6.6 | 4.1 | 28.5 | 86.2 | 27.1 | 6.7 | 5.0 |
| 30.3 | 6.9 | 5.0 | 29.4 | 85.7 | 28.1 | 6.7 | 4.8 |
| 30.1 | 6.9 | 4.8 | 29.2 | 84.4 | 28.4 | 6.7 | 4.6 |
| 30.6 | 7.0 | 4.8 | 30.4 | 85.2 | 29.7 | 6.7 | 4.4 |
| 31.0 | 7.0 | 5.1 | 28.3 | 83.8 | 28.5 | 6.6 | 4.9 |
| 25.8 | 6.6 | 4.1 | 28.5 | 86.2 | 27.1 | 6.7 | 5.0 |
| 29.1 | 6.9 | 4.9 | 28.9 | 84.7 | 27.0 | 6.7 | 5.0 |
| 27.5 | 6.8 | 4.7 | 28.7 | 84.8 | 28.3 | 6.7 | 4.8 |
| 28.7 | 6.7 | 5.4 | 28.4 | 85.4 | 28.0 | 6.6 | 5.3 |
| 29.5 | 6.8 | 5.1 | 29.6 | 87.9 | 28.0 | 6.7 | 5.4 |
| 25.8 | 6.6 | 4.1 | 28.5 | 86.2 | 27.1 | 6.7 | 5.0 |
| 30.4 | 6.9 | 4.9 | 30.0 | 86.3 | 30.3 | 6.8 | 4.9 |
| 30.1 | 6.8 | 5.0 | 29.9 | 81.4 | 29.8 | 6.8 | 5.3 |
| 27.0 | 6.7 | 4.3 | 26.3 | 84.3 | 25.8 | 6.6 | 4.8 |
| 27.9 | 6.9 | 5.1 | 28.1 | 86.7 | 27.1 | 6.7 | 5.0 |
| 25.8 | 6.6 | 4.1 | 28.5 | 86.2 | 27.1 | 6.7 | 5.0 |
| 28.6 | 6.8 | 5.4 | 28.9 | 85.3 | 29.7 | 6.8 | 5.2 |
| 31.4 | 6.8 | 3.3 | 29.4 | 85.4 | 28.2 | 6.7 | 5.1 |
| 31.7 | 6.9 | 4.5 | 30.4 | 87.3 | 28.9 | 6.7 | 5.1 |
| 26.9 | 6.7 | 5.1 | 29.1 | 86.6 | 28.6 | 6.7 | 4.7 |
| 25.8 | 6.6 | 4.1 | 28.5 | 86.2 | 27.1 | 6.7 | 5.0 |
| 28.9 | 6.8 | 4.0 | 30.1 | 86.2 | 28.0 | 6.7 | 5.0 |
| 31.5 | 6.8 | 3.8 | 31.5 | 86.8 | 29.0 | 6.8 | 5.5 |
| 30.9 | 6.8 | 4.5 | 26.7 | 83.2 | 25.6 | 6.5 | 4.9 |
| 29.4 | 6.7 | 4.6 | 28.5 | 85.7 | 28.3 | 6.7 | 4.9 |
| 25.8 | 6.6 | 4.1 | 28.5 | 86.2 | 27.1 | 6.7 | 5.0 |
| 28.1 | 6.9 | 4.7 | 28.1 | 84.3 | 27.7 | 6.7 | 5.0 |
| 29.9 | 6.8 | 4.5 | 29.6 | 81.7 | 30.7 | 6.7 | 4.5 |
| 29.2 | 6.8 | 4.9 | 29.1 | 86.4 | 29.6 | 6.7 | 4.4 |
| 30.3 | 6.9 | 4.2 | 28.2 | 82.6 | 31.2 | 6.8 | 4.4 |
| 29.9 | 6.9 | 5.3 | 28.0 | 85.0 | 27.0 | 6.6 | 4.3 |
| 29.3 | 6.8 | 5.0 | 28.4 | 84.7 | 28.9 | 6.7 | 4.9 |
| 28.3 | 6.7 | 4.7 | 28.1 | 83.3 | 28.1 | 6.7 | 5.5 |
| 29.7 | 6.8 | 5.2 | 28.7 | 83.8 | 29.4 | 6.7 | 4.7 |
| 30.5 | 6.9 | 4.8 | 29.6 | 86.0 | 27.5 | 6.7 | 4.9 |
| 29.9 | 6.9 | 5.3 | 28.0 | 85.0 | 27.0 | 6.6 | 4.3 |
| 30.2 | 6.9 | 5.1 | 27.2 | 85.3 | 27.0 | 6.6 | 5.5 |
| 29.3 | 6.8 | 5.4 | 26.2 | 85.4 | 26.2 | 6.5 | 5.5 |
| 28.7 | 6.8 | 5.3 | 29.4 | 85.9 | 29.3 | 6.7 | 5.8 |
| 28.2 | 6.9 | 5.5 | 28.8 | 86.4 | 28.7 | 6.7 | 5.1 |
| 29.9 | 6.9 | 5.3 | 28.0 | 85.0 | 27.0 | 6.6 | 4.3 |
| 28.4 | 6.8 | 5.0 | 28.8 | 86.6 | 30.0 | 6.8 | 5.2 |
| 28.0 | 6.8 | 5.5 | 29.1 | 86.5 | 28.0 | 6.7 | 5.2 |
| 26.2 | 6.6 | 5.2 | 28.8 | 87.0 | 28.2 | 6.7 | 5.8 |
| 29.1 | 6.8 | 5.0 | 28.2 | 86.3 | 27.9 | 6.7 | 5.4 |
| 29.9 | 6.9 | 5.3 | 28.0 | 85.0 | 27.0 | 6.6 | 4.3 |
| 28.3 | 6.8 | 5.0 | 29.7 | 86.0 | 29.8 | 6.7 | 4.5 |
| 29.7 | 6.9 | 5.4 | 29.2 | 83.7 | 31.2 | 6.7 | 4.4 |
| 27.9 | 6.7 | 4.7 | 29.1 | 83.0 | 31.1 | 6.7 | 4.2 |
| 27.8 | 6.7 | 5.0 | 29.0 | 85.4 | 29.8 | 6.7 | 4.1 |
| 29.9 | 6.9 | 5.3 | 28.0 | 85.0 | 27.0 | 6.6 | 4.3 |
| 27.7 | 6.7 | 4.9 | 29.8 | 86.0 | 29.3 | 6.7 | 4.8 |
| 30.3 | 6.7 | 5.0 | 29.2 | 84.2 | 30.2 | 6.7 | 4.5 |
| 28.4 | 6.7 | 5.0 | 29.3 | 86.1 | 29.7 | 6.8 | 5.2 |
| 28.3 | 6.8 | 5.1 | 29.5 | 85.7 | 29.8 | 6.7 | 5.5 |
| 29.9 | 6.9 | 5.3 | 28.0 | 85.0 | 27.0 | 6.6 | 4.3 |
| 29.4 | 6.9 | 5.7 | 29.5 | 86.9 | 31.4 | 6.7 | 4.3 |
| 30.7 | 6.8 | 5.3 | 29.9 | 85.9 | 28.8 | 6.7 | 5.4 |
| 29.3 | 6.8 | 4.1 | 30.8 | 84.5 | 28.7 | 6.7 | 4.4 |
| 29.2 | 6.8 | 4.0 | 29.6 | 85.8 | 30.6 | 6.8 | 4.1 |
| 29.9 | 6.9 | 5.3 | 28.0 | 85.0 | 27.0 | 6.6 | 4.3 |
| 28.5 | 6.8 | 4.9 | 29.9 | 85.6 | 30.1 | 6.8 | 4.9 |
| 28.8 | 6.9 | 5.3 | 28.7 | 84.3 | 26.6 | 6.7 | 5.4 |
| 28.5 | 6.8 | 4.9 | 28.8 | 84.3 | 29.7 | 6.7 | 3.4 |
| 29.6 | 6.8 | 3.9 | 29.3 | 86.2 | 29.0 | 6.8 | 5.0 |
| 29.9 | 6.9 | 5.3 | 28.0 | 85.0 | 27.0 | 6.6 | 4.3 |
| 30.2 | 6.8 | 3.7 | 29.3 | 82.5 | 29.1 | 6.7 | 4.6 |
| 27.3 | 6.7 | 5.0 | 29.7 | 84.9 | 29.2 | 6.7 | 4.9 |
| 28.8 | 6.6 | 4.6 | 29.8 | 84.6 | 31.0 | 6.7 | 4.5 |
| 29.1 | 6.8 | 4.7 | 29.8 | 85.3 | 30.3 | 6.7 | 4.8 |
| 29.9 | 6.9 | 5.3 | 28.0 | 85.0 | 27.0 | 6.6 | 4.3 |
| 28.3 | 6.8 | 4.9 | 30.0 | 85.4 | 29.5 | 6.8 | 5.3 |
| 27.5 | 6.7 | 4.1 | 28.8 | 85.8 | 28.7 | 6.7 | 4.7 |
| 28.3 | 6.8 | 4.7 | 29.5 | 85.7 | 29.7 | 6.7 | 4.6 |
| 28.8 | 6.8 | 4.6 | 29.1 | 84.3 | 28.3 | 6.7 | 5.1 |
| 29.9 | 6.9 | 5.3 | 28.0 | 85.0 | 27.0 | 6.6 | 4.3 |
| 29.1 | 6.7 | 4.8 | 29.8 | 85.4 | 31.9 | 6.8 | 4.1 |
| 30.9 | 6.9 | 4.8 | 29.1 | 84.4 | 29.7 | 6.7 | 5.1 |
| 29.6 | 6.8 | 3.2 | 26.5 | 83.7 | 26.5 | 6.6 | 5.9 |
| 27.9 | 6.8 | 5.5 | 28.7 | 85.3 | 28.2 | 6.7 | 4.9 |
| 29.4 | 6.9 | 4.9 | 28.4 | 85.3 | 28.6 | 6.7 | 4.7 |
| 29.3 | 6.9 | 5.3 | 29.5 | 87.0 | 29.3 | 6.7 | 5.0 |
| 28.5 | 6.9 | 5.8 | 29.6 | 85.6 | 29.3 | 6.8 | 5.0 |
| 30.2 | 6.9 | 5.2 | 31.3 | 85.9 | 31.1 | 6.8 | 5.1 |
| 28.8 | 6.8 | 5.1 | 30.0 | 85.5 | 30.1 | 6.8 | 5.1 |
| 29.4 | 6.9 | 4.9 | 28.4 | 85.3 | 28.6 | 6.7 | 4.7 |
| 29.3 | 6.9 | 5.0 | 29.4 | 85.6 | 28.6 | 6.8 | 5.7 |
| 27.9 | 6.9 | 5.6 | 27.1 | 82.4 | 27.6 | 6.6 | 5.3 |
| 30.5 | 6.9 | 5.4 | 30.0 | 84.7 | 29.1 | 6.8 | 5.5 |
| 28.4 | 6.9 | 5.4 | 30.3 | 85.9 | 31.9 | 6.8 | 4.2 |
| 29.4 | 6.9 | 4.9 | 28.4 | 85.3 | 28.6 | 6.7 | 4.7 |
| 30.6 | 6.8 | 4.9 | 29.1 | 82.9 | 28.6 | 6.8 | 5.8 |
| 28.5 | 6.9 | 5.7 | 27.0 | 83.0 | 27.0 | 6.6 | 5.5 |
| 29.6 | 6.8 | 5.4 | 29.3 | 85.3 | 29.1 | 6.8 | 5.9 |
| 30.2 | 6.8 | 5.0 | 29.3 | 85.8 | 27.4 | 6.6 | 4.5 |
| 29.4 | 6.9 | 4.9 | 28.4 | 85.3 | 28.6 | 6.7 | 4.7 |
| 31.5 | 6.9 | 4.6 | 31.1 | 86.5 | 32.0 | 6.8 | 4.8 |
| 33.6 | 6.9 | 4.1 | 32.3 | 85.1 | 33.2 | 6.8 | 4.1 |
| 27.4 | 6.7 | 4.9 | 28.3 | 82.6 | 27.7 | 6.7 | 5.3 |
| 31.1 | 6.9 | 4.6 | 28.1 | 82.7 | 25.5 | 6.6 | 4.7 |
| 29.4 | 6.9 | 4.9 | 28.4 | 85.3 | 28.6 | 6.7 | 4.7 |
| 29.0 | 6.8 | 5.3 | 29.1 | 85.9 | 28.6 | 6.7 | 4.3 |
| 29.1 | 6.8 | 5.1 | 31.2 | 85.5 | 29.7 | 6.8 | 5.0 |
| 28.2 | 6.7 | 4.8 | 28.4 | 82.8 | 27.7 | 6.7 | 5.5 |
| 29.4 | 6.9 | 4.9 | 28.4 | 85.3 | 28.6 | 6.7 | 4.7 |
| 29.0 | 6.8 | 4.5 | 30.1 | 86.0 | 31.1 | 6.7 | 4.3 |
| 31.7 | 6.8 | 4.6 | 31.4 | 85.4 | 30.2 | 6.8 | 5.3 |
| 27.6 | 6.9 | 5.3 | 30.1 | 84.3 | 29.0 | 6.7 | 4.1 |
| 29.4 | 6.7 | 3.8 | 28.3 | 82.9 | 26.4 | 6.6 | 4.5 |
| 29.4 | 6.9 | 4.9 | 28.4 | 85.3 | 28.6 | 6.7 | 4.7 |
| 29.6 | 6.8 | 4.8 | 30.1 | 87.0 | 29.7 | 6.8 | 4.8 |
| 27.8 | 6.8 | 5.5 | 28.6 | 86.2 | 27.7 | 6.7 | 4.8 |
| 30.9 | 6.9 | 4.7 | 28.4 | 85.0 | 31.3 | 6.8 | 5.1 |
| 27.0 | 6.7 | 5.4 | 30.0 | 87.2 | 29.6 | 6.8 | 5.5 |
| 29.4 | 6.9 | 4.9 | 28.4 | 85.3 | 28.6 | 6.7 | 4.7 |
| 28.5 | 6.9 | 5.2 | 29.2 | 85.3 | 28.1 | 6.7 | 4.9 |
| 31.9 | 7.0 | 4.7 | 28.6 | 82.0 | 29.1 | 6.7 | 4.5 |
| 29.6 | 6.7 | 4.6 | 27.7 | 82.1 | 26.8 | 6.6 | 5.1 |
| 27.3 | 6.7 | 4.7 | 30.2 | 86.0 | 28.6 | 6.7 | 5.0 |
| 29.4 | 6.9 | 4.9 | 28.4 | 85.3 | 28.6 | 6.7 | 4.7 |
| 28.2 | 6.8 | 4.7 | 29.2 | 85.1 | 26.9 | 6.6 | 5.1 |
| 28.0 | 6.7 | 4.2 | 29.9 | 86.4 | 27.5 | 6.7 | 5.6 |
| 30.0 | 6.8 | 4.7 | 27.1 | 82.7 | 28.8 | 6.6 | 5.1 |
| 27.0 | 6.5 | 4.3 | 28.2 | 84.1 | 27.6 | 6.6 | 4.8 |
| 29.4 | 6.9 | 4.9 | 28.4 | 85.3 | 28.6 | 6.7 | 4.7 |
| 27.7 | 6.8 | 5.4 | 27.8 | 83.6 | 27.6 | 6.6 | 4.9 |
| 28.2 | 6.8 | 4.3 | 28.8 | 84.7 | 27.3 | 6.6 | 5.5 |
| 28.2 | 6.8 | 4.7 | 28.9 | 84.3 | 28.6 | 6.7 | 5.4 |
| 27.4 | 6.8 | 5.3 | 30.5 | 86.9 | 30.9 | 6.9 | 5.4 |
| 29.8 | 6.7 | 3.8 | 27.6 | 84.8 | 28.0 | 6.6 | 5.3 |
| 30.9 | 6.8 | 4.2 | 31.0 | 85.9 | 31.2 | 6.8 | 4.4 |
| 30.2 | 6.9 | 5.4 | 30.6 | 86.9 | 29.7 | 6.8 | 5.2 |
| 31.2 | 6.9 | 5.4 | 32.3 | 86.8 | 31.1 | 6.9 | 5.0 |
| 31.7 | 6.9 | 4.2 | 31.3 | 86.5 | 29.8 | 6.9 | 4.8 |
| 29.8 | 6.7 | 3.8 | 27.6 | 84.8 | 28.0 | 6.6 | 5.3 |
| 30.2 | 6.8 | 4.8 | 31.1 | 85.6 | 29.3 | 6.8 | 4.9 |
| 32.5 | 6.9 | 3.5 | 31.0 | 86.0 | 32.3 | 7.0 | 4.7 |
| 30.8 | 6.8 | 4.2 | 30.4 | 86.3 | 32.2 | 6.8 | 4.6 |
| 29.3 | 6.8 | 5.4 | 29.7 | 85.3 | 29.2 | 6.8 | 4.7 |
| 29.8 | 6.7 | 3.8 | 27.6 | 84.8 | 28.0 | 6.6 | 5.3 |
| 28.0 | 6.8 | 4.8 | 31.0 | 85.2 | 28.9 | 6.8 | 5.1 |
| 29.4 | 6.8 | 4.7 | 30.5 | 87.3 | 30.4 | 6.8 | 5.2 |
| 31.0 | 6.7 | 4.8 | 27.7 | 84.9 | 30.0 | 6.7 | 5.7 |
| 28.8 | 6.8 | 4.7 | 29.6 | 85.2 | 29.5 | 6.8 | 5.2 |
| 29.8 | 6.7 | 3.8 | 27.6 | 84.8 | 28.0 | 6.6 | 5.3 |
| 30.5 | 6.7 | 3.7 | 31.1 | 86.4 | 30.2 | 6.9 | 4.9 |
| 27.6 | 6.8 | 5.2 | 30.2 | 84.6 | 30.0 | 6.8 | 4.6 |
| 30.2 | 6.9 | 4.6 | 30.1 | 86.5 | 29.2 | 6.7 | 4.4 |
| 30.2 | 6.8 | 4.2 | 29.3 | 86.3 | 29.5 | 6.8 | 5.0 |
| 29.8 | 6.7 | 3.8 | 27.6 | 84.8 | 28.0 | 6.6 | 5.3 |
| 27.6 | 6.8 | 5.3 | 29.9 | 86.7 | 29.2 | 6.8 | 5.2 |
| 31.1 | 6.8 | 5.1 | 27.6 | 84.3 | 29.0 | 6.7 | 5.6 |
| 28.4 | 6.9 | 5.0 | 30.0 | 86.5 | 31.4 | 6.8 | 4.5 |
| 29.5 | 6.8 | 4.0 | 30.0 | 85.9 | 31.0 | 6.9 | 5.0 |
| 29.8 | 6.7 | 3.8 | 27.6 | 84.8 | 28.0 | 6.6 | 5.3 |
| 27.9 | 6.8 | 5.0 | 29.0 | 84.0 | 28.5 | 6.7 | 5.0 |
| 29.9 | 6.9 | 4.7 | 28.7 | 85.4 | 31.0 | 6.9 | 5.0 |
| 28.3 | 6.8 | 5.3 | 28.7 | 83.7 | 29.5 | 6.7 | 4.9 |
| 29.4 | 6.9 | 4.8 | 28.4 | 83.9 | 28.3 | 6.7 | 5.4 |
| 29.8 | 6.7 | 3.8 | 27.6 | 84.8 | 28.0 | 6.6 | 5.3 |
| 28.5 | 6.8 | 5.2 | 28.6 | 84.3 | 29.5 | 6.8 | 5.2 |
| 30.3 | 6.8 | 4.6 | 30.5 | 85.4 | 28.9 | 6.8 | 5.1 |
| 30.1 | 6.9 | 4.6 | 29.8 | 83.7 | 29.1 | 6.8 | 4.9 |
| 30.4 | 7.0 | 4.7 | 29.9 | 85.0 | 30.4 | 6.8 | 4.6 |
| 29.8 | 6.7 | 3.8 | 27.6 | 84.8 | 28.0 | 6.6 | 5.3 |
| 29.4 | 6.8 | 3.9 | 30.0 | 85.5 | 28.6 | 6.8 | 5.1 |
| 28.1 | 6.6 | 4.2 | 29.8 | 85.2 | 28.9 | 6.8 | 4.9 |
| 27.7 | 6.9 | 5.3 | 28.9 | 85.3 | 27.4 | 6.7 | 5.3 |
| 27.8 | 6.8 | 4.8 | 29.9 | 84.5 | 28.5 | 6.8 | 5.1 |
| 29.8 | 6.7 | 3.8 | 27.6 | 84.8 | 28.0 | 6.6 | 5.3 |
| 30.6 | 6.9 | 4.7 | 29.8 | 85.0 | 29.9 | 6.8 | 5.1 |
| 30.9 | 6.8 | 4.6 | 28.3 | 84.9 | 27.0 | 6.6 | 5.4 |
| 30.9 | 6.8 | 4.4 | 30.9 | 85.7 | 31.3 | 6.8 | 4.2 |
| 28.4 | 6.8 | 4.9 | 30.5 | 86.2 | 29.7 | 6.8 | 4.9 |
| 29.8 | 6.7 | 3.8 | 27.6 | 84.8 | 28.0 | 6.6 | 5.3 |
| 30.4 | 6.8 | 4.6 | 29.7 | 84.1 | 31.9 | 6.8 | 4.6 |
| 26.1 | 6.6 | 5.2 | 29.0 | 85.2 | 28.8 | 6.7 | 4.8 |
| 28.5 | 6.8 | 4.8 | 30.2 | 87.3 | 30.7 | 6.8 | 4.7 |
| 27.9 | 6.8 | 5.2 | 30.3 | 88.0 | 29.2 | 6.8 | 4.7 |
| 29.4 | 6.7 | 3.5 | 29.4 | 85.8 | 28.8 | 6.7 | 5.3 |
| 29.6 | 6.8 | 4.8 | 30.9 | 87.4 | 29.9 | 6.8 | 5.1 |
| 31.4 | 6.9 | 4.7 | 29.0 | 83.7 | 29.0 | 6.7 | 5.5 |
| 26.3 | 6.6 | 5.1 | 29.4 | 85.6 | 28.0 | 6.7 | 5.5 |
| 29.2 | 6.9 | 5.4 | 30.3 | 85.3 | 29.8 | 6.8 | 5.1 |
| 29.4 | 6.7 | 3.5 | 29.4 | 85.8 | 28.8 | 6.7 | 5.3 |
| 29.1 | 6.9 | 5.2 | 29.6 | 83.7 | 29.4 | 6.8 | 5.4 |
| 29.1 | 6.7 | 4.0 | 27.1 | 80.9 | 26.4 | 6.6 | 5.6 |
| 27.6 | 6.7 | 5.1 | 28.7 | 84.7 | 27.3 | 6.7 | 5.3 |
| 27.9 | 6.8 | 5.4 | 29.4 | 84.9 | 28.4 | 6.7 | 4.8 |
| 29.4 | 6.7 | 3.5 | 29.4 | 85.8 | 28.8 | 6.7 | 5.3 |
| 28.5 | 6.9 | 5.3 | 29.9 | 85.6 | 29.0 | 6.8 | 5.4 |
| 29.7 | 6.9 | 5.3 | 29.4 | 85.2 | 30.6 | 6.8 | 5.1 |
| 28.4 | 6.7 | 4.0 | 30.3 | 86.3 | 29.6 | 6.8 | 4.5 |
| 28.9 | 6.8 | 4.7 | 29.5 | 85.4 | 29.8 | 6.8 | 4.7 |
| 29.4 | 6.7 | 3.5 | 29.4 | 85.8 | 28.8 | 6.7 | 5.3 |
| 28.9 | 6.9 | 4.7 | 30.5 | 85.8 | 31.2 | 6.8 | 4.4 |
| 29.2 | 6.9 | 4.9 | 30.9 | 85.7 | 29.1 | 6.7 | 4.5 |
| 29.5 | 6.8 | 4.5 | 29.4 | 85.9 | 29.3 | 6.8 | 5.2 |
| 27.2 | 6.8 | 4.8 | 30.5 | 85.1 | 30.6 | 6.8 | 5.4 |
| 29.4 | 6.7 | 3.5 | 29.4 | 85.8 | 28.8 | 6.7 | 5.3 |
| 30.0 | 6.9 | 4.1 | 29.1 | 82.7 | 28.2 | 6.8 | 5.4 |
| 28.2 | 6.9 | 4.6 | 31.6 | 88.3 | 30.0 | 6.9 | 4.7 |
| 29.1 | 6.9 | 4.5 | 29.4 | 83.1 | 29.9 | 6.7 | 4.0 |
| 29.4 | 6.8 | 3.8 | 30.2 | 84.9 | 29.8 | 6.7 | 4.3 |
| 29.4 | 6.7 | 3.5 | 29.4 | 85.8 | 28.8 | 6.7 | 5.3 |
| 28.4 | 6.9 | 5.4 | 29.8 | 85.4 | 29.9 | 6.8 | 5.0 |
| 27.5 | 6.9 | 4.7 | 27.9 | 84.3 | 28.8 | 6.7 | 4.8 |
| 32.5 | 7.0 | 4.9 | 31.0 | 86.2 | 30.8 | 6.9 | 5.6 |
| 30.7 | 6.9 | 4.5 | 32.4 | 84.5 | 33.7 | 6.9 | 5.0 |
| 29.4 | 6.7 | 3.5 | 29.4 | 85.8 | 28.8 | 6.7 | 5.3 |
| 29.2 | 6.8 | 5.1 | 29.6 | 86.5 | 28.2 | 6.7 | 5.0 |
| 27.8 | 6.8 | 5.6 | 29.2 | 85.3 | 29.6 | 6.8 | 5.3 |
| 29.1 | 6.9 | 4.9 | 31.0 | 85.9 | 31.5 | 6.8 | 4.9 |
| 29.0 | 6.8 | 4.5 | 30.6 | 85.8 | 29.6 | 6.7 | 4.8 |
| 29.4 | 6.7 | 3.5 | 29.4 | 85.8 | 28.8 | 6.7 | 5.3 |
| 28.6 | 6.8 | 4.5 | 29.3 | 84.1 | 28.7 | 6.7 | 5.5 |
| 29.7 | 6.9 | 5.2 | 29.9 | 86.1 | 28.7 | 6.6 | 5.0 |
| 29.0 | 6.7 | 4.3 | 30.8 | 86.4 | 29.3 | 6.7 | 4.9 |
| 29.1 | 6.9 | 5.0 | 31.3 | 86.4 | 28.9 | 6.7 | 4.4 |
| 29.4 | 6.7 | 3.5 | 29.4 | 85.8 | 28.8 | 6.7 | 5.3 |
| 27.9 | 6.8 | 4.8 | 29.8 | 86.4 | 27.0 | 6.7 | 5.2 |
| 28.0 | 6.7 | 5.1 | 30.1 | 85.9 | 29.3 | 6.8 | 5.0 |
| 30.3 | 6.9 | 4.7 | 30.9 | 86.6 | 28.5 | 6.8 | 4.6 |
| 29.3 | 6.8 | 5.0 | 29.3 | 86.5 | 27.3 | 6.7 | 5.3 |
| 29.4 | 6.7 | 3.5 | 29.4 | 85.8 | 28.8 | 6.7 | 5.3 |
| 29.1 | 6.9 | 5.2 | 30.1 | 86.1 | 29.5 | 6.8 | 5.3 |
| 29.0 | 6.8 | 5.4 | 30.9 | 86.0 | 30.1 | 6.8 | 5.3 |
| 29.2 | 6.9 | 5.6 | 29.5 | 82.5 | 28.9 | 6.8 | 5.1 |
| 29.0 | 6.9 | 4.9 | 30.2 | 84.6 | 27.0 | 6.7 | 5.1 |
| 29.9 | 6.8 | 3.9 | 30.1 | 86.6 | 30.1 | 6.8 | 4.6 |
| 26.3 | 6.6 | 3.8 | 28.0 | 82.7 | 28.4 | 6.6 | 4.6 |
| 28.0 | 6.8 | 5.2 | 31.9 | 86.4 | 31.2 | 6.9 | 4.6 |
| 28.9 | 6.8 | 5.2 | 29.3 | 84.4 | 29.7 | 6.7 | 5.3 |
| 28.3 | 6.8 | 5.4 | 30.6 | 83.7 | 31.4 | 6.9 | 4.7 |
| 29.9 | 6.8 | 3.9 | 30.1 | 86.6 | 30.1 | 6.8 | 4.6 |
| 28.6 | 6.8 | 4.4 | 30.7 | 86.2 | 30.3 | 6.9 | 4.9 |
| 27.6 | 6.8 | 5.3 | 29.7 | 84.9 | 28.2 | 6.7 | 5.4 |
| 28.9 | 6.8 | 4.4 | 28.7 | 85.0 | 27.1 | 6.6 | 4.8 |
| 28.1 | 6.8 | 4.8 | 28.0 | 84.3 | 28.0 | 6.6 | 4.8 |
| 29.9 | 6.8 | 3.9 | 30.1 | 86.6 | 30.1 | 6.8 | 4.6 |
| 25.9 | 6.7 | 5.4 | 25.1 | 81.5 | 24.9 | 6.5 | 5.0 |
| 28.0 | 6.9 | 5.1 | 26.5 | 79.1 | 26.8 | 6.5 | 4.4 |
| 29.0 | 6.9 | 4.6 | 30.5 | 83.7 | 28.8 | 6.8 | 5.1 |
| 28.2 | 6.8 | 4.3 | 32.0 | 85.8 | 29.0 | 6.9 | 4.8 |
| 29.9 | 6.8 | 3.9 | 30.1 | 86.6 | 30.1 | 6.8 | 4.6 |
| 28.5 | 6.8 | 4.7 | 30.0 | 86.2 | 28.0 | 6.7 | 4.6 |
| 28.4 | 6.8 | 4.7 | 31.0 | 82.9 | 30.4 | 6.8 | 4.1 |
| 28.9 | 6.9 | 5.1 | 31.3 | 84.4 | 28.6 | 6.7 | 5.0 |
| 30.1 | 6.9 | 4.9 | 30.6 | 85.1 | 30.2 | 6.8 | 4.4 |
| 29.9 | 6.8 | 3.9 | 30.1 | 86.6 | 30.1 | 6.8 | 4.6 |
| 28.4 | 6.9 | 4.6 | 30.8 | 85.7 | 30.2 | 6.8 | 4.7 |
| 28.3 | 6.7 | 3.9 | 31.3 | 87.1 | 29.8 | 6.8 | 5.0 |
| 28.2 | 6.7 | 5.3 | 30.0 | 86.2 | 29.2 | 6.8 | 5.3 |
| 28.4 | 6.8 | 4.8 | 29.4 | 86.2 | 28.5 | 6.7 | 4.6 |
| 29.9 | 6.8 | 3.9 | 30.1 | 86.6 | 30.1 | 6.8 | 4.6 |
| 27.0 | 6.7 | 4.6 | 28.7 | 83.1 | 31.7 | 6.7 | 5.0 |
| 28.7 | 6.8 | 5.5 | 28.7 | 86.3 | 29.7 | 6.7 | 5.5 |
| 27.8 | 6.8 | 5.6 | 30.8 | 86.0 | 31.0 | 6.9 | 5.2 |
| 29.8 | 6.9 | 5.0 | 31.7 | 86.0 | 31.2 | 7.0 | 5.1 |
| 29.9 | 6.8 | 3.9 | 30.1 | 86.6 | 30.1 | 6.8 | 4.6 |
| 29.0 | 6.8 | 4.6 | 31.2 | 82.5 | 30.6 | 6.8 | 5.1 |
| 30.4 | 6.8 | 4.8 | 30.5 | 84.3 | 30.8 | 6.8 | 4.9 |
| 31.3 | 6.8 | 4.7 | 30.9 | 85.8 | 31.7 | 6.8 | 4.6 |
| 27.2 | 6.9 | 5.5 | 30.8 | 86.6 | 30.9 | 6.8 | 4.8 |
| 29.9 | 6.8 | 3.9 | 30.1 | 86.6 | 30.1 | 6.8 | 4.6 |
| 28.1 | 6.9 | 5.3 | 30.1 | 84.8 | 29.4 | 6.8 | 5.5 |
| 31.2 | 6.7 | 4.0 | 28.9 | 84.9 | 26.5 | 6.7 | 5.2 |
| 29.8 | 6.9 | 4.8 | 30.6 | 85.8 | 30.3 | 6.9 | 5.1 |
| 29.5 | 6.8 | 4.6 | 30.4 | 86.3 | 29.4 | 6.8 | 5.1 |
| 29.9 | 6.8 | 3.9 | 30.1 | 86.6 | 30.1 | 6.8 | 4.6 |
| 25.1 | 6.6 | 5.5 | 29.6 | 86.0 | 27.8 | 6.6 | 5.0 |
| 26.4 | 6.7 | 5.3 | 28.0 | 84.6 | 26.9 | 6.7 | 5.3 |
| 29.0 | 7.0 | 5.2 | 28.9 | 81.7 | 27.8 | 6.7 | 4.7 |
| 30.2 | 6.9 | 4.8 | 30.7 | 86.6 | 30.8 | 6.9 | 4.7 |
| 29.9 | 6.8 | 3.9 | 30.1 | 86.6 | 30.1 | 6.8 | 4.6 |
| 28.7 | 6.8 | 5.2 | 31.3 | 87.2 | 30.6 | 6.8 | 4.9 |
| 30.7 | 6.8 | 4.7 | 30.6 | 86.5 | 30.9 | 6.7 | 4.4 |
| 28.7 | 6.8 | 5.4 | 30.5 | 85.2 | 31.8 | 6.8 | 4.3 |
| 28.9 | 6.9 | 5.3 | 30.5 | 85.4 | 28.8 | 6.8 | 4.6 |
| 27.0 | 6.7 | 4.8 | 29.1 | 87.2 | 28.8 | 6.7 | 4.8 |
| 28.7 | 6.9 | 5.3 | 29.7 | 86.4 | 28.3 | 6.7 | 5.4 |
| 28.6 | 6.8 | 5.4 | 27.7 | 84.0 | 27.7 | 6.6 | 5.2 |
| 31.5 | 6.9 | 4.0 | 30.0 | 85.4 | 30.1 | 6.8 | 4.8 |
| 29.0 | 6.9 | 4.8 | 29.8 | 84.9 | 30.5 | 6.8 | 4.4 |
| 27.0 | 6.7 | 4.8 | 29.1 | 87.2 | 28.8 | 6.7 | 4.8 |
| 29.4 | 6.8 | 5.2 | 27.8 | 81.6 | 29.3 | 6.7 | 5.3 |
| 29.2 | 6.7 | 4.5 | 28.2 | 85.9 | 29.9 | 6.7 | 5.1 |
| 29.7 | 6.7 | 4.8 | 28.1 | 85.1 | 31.9 | 6.7 | 4.8 |
| 30.2 | 6.8 | 4.9 | 29.2 | 84.0 | 29.3 | 6.6 | 4.7 |
| 27.0 | 6.7 | 4.8 | 29.1 | 87.2 | 28.8 | 6.7 | 4.8 |
| 29.9 | 6.8 | 4.0 | 29.6 | 86.4 | 31.4 | 6.8 | 4.4 |
| 30.1 | 6.9 | 4.5 | 30.7 | 86.3 | 29.5 | 6.7 | 4.7 |
| 29.2 | 6.7 | 4.4 | 31.1 | 85.4 | 30.8 | 6.8 | 4.5 |
| 29.4 | 6.8 | 4.6 | 30.2 | 83.7 | 29.3 | 6.8 | 5.4 |
| 27.0 | 6.7 | 4.8 | 29.1 | 87.2 | 28.8 | 6.7 | 4.8 |
| 28.3 | 6.8 | 4.4 | 27.8 | 85.4 | 27.0 | 6.6 | 5.7 |
| 25.8 | 6.6 | 5.6 | 25.5 | 83.4 | 25.2 | 6.5 | 5.7 |
| 27.5 | 6.6 | 5.2 | 26.3 | 83.2 | 26.8 | 6.5 | 4.7 |
| 27.4 | 6.8 | 5.1 | 27.4 | 83.2 | 27.5 | 6.6 | 5.2 |
| 27.0 | 6.7 | 4.8 | 29.1 | 87.2 | 28.8 | 6.7 | 4.8 |
| 29.1 | 6.9 | 5.2 | 27.7 | 84.5 | 27.0 | 6.6 | 5.4 |
| 28.4 | 6.9 | 5.1 | 29.0 | 85.9 | 27.8 | 6.8 | 5.2 |
| 29.1 | 6.7 | 5.0 | 28.6 | 86.4 | 28.7 | 6.7 | 5.2 |
| 29.7 | 6.8 | 4.4 | 29.5 | 84.6 | 30.5 | 6.8 | 5.3 |
| 27.0 | 6.7 | 4.8 | 29.1 | 87.2 | 28.8 | 6.7 | 4.8 |
| 28.2 | 6.8 | 4.9 | 30.6 | 85.0 | 29.4 | 6.8 | 5.2 |
| 29.0 | 6.8 | 4.5 | 29.8 | 82.3 | 30.2 | 6.8 | 4.8 |
| 28.7 | 6.7 | 3.7 | 31.0 | 85.0 | 30.7 | 6.8 | 4.3 |
| 27.9 | 6.8 | 4.6 | 31.0 | 85.4 | 30.8 | 6.8 | 4.5 |
| 27.0 | 6.7 | 4.8 | 29.1 | 87.2 | 28.8 | 6.7 | 4.8 |
| 26.4 | 6.6 | 4.3 | 29.0 | 84.9 | 29.1 | 6.7 | 4.8 |
| 28.8 | 6.9 | 5.1 | 29.3 | 86.5 | 28.5 | 6.7 | 5.2 |
| 29.5 | 6.8 | 3.4 | 28.8 | 85.1 | 27.6 | 6.6 | 5.1 |
| 28.0 | 6.8 | 4.9 | 31.2 | 86.1 | 30.4 | 6.8 | 5.0 |
| 27.0 | 6.7 | 4.8 | 29.1 | 87.2 | 28.8 | 6.7 | 4.8 |
| 29.6 | 6.8 | 5.2 | 30.2 | 86.8 | 28.3 | 6.7 | 5.1 |
| 31.7 | 7.0 | 4.8 | 30.5 | 85.2 | 30.1 | 6.7 | 5.0 |
| 27.9 | 6.7 | 4.7 | 26.5 | 82.7 | 26.7 | 6.6 | 4.9 |
| 28.3 | 6.8 | 5.2 | 28.0 | 83.0 | 28.2 | 6.7 | 5.2 |
| 27.0 | 6.7 | 4.8 | 29.1 | 87.2 | 28.8 | 6.7 | 4.8 |
| 29.1 | 6.7 | 5.0 | 29.1 | 85.2 | 28.4 | 6.7 | 5.0 |
| 30.6 | 6.9 | 4.6 | 31.6 | 86.1 | 30.0 | 6.7 | 4.5 |
| 27.7 | 6.9 | 5.7 | 27.3 | 84.0 | 27.1 | 6.6 | 4.9 |
| 28.3 | 6.8 | 5.4 | 31.2 | 85.7 | 30.2 | 6.9 | 4.9 |
| 27.0 | 6.7 | 4.8 | 29.1 | 87.2 | 28.8 | 6.7 | 4.8 |
| 29.4 | 6.8 | 5.2 | 28.5 | 84.8 | 28.8 | 6.7 | 4.8 |
| 27.4 | 6.8 | 5.3 | 29.7 | 84.2 | 29.2 | 6.7 | 5.3 |
| 29.0 | 6.8 | 5.5 | 28.9 | 84.0 | 29.8 | 6.7 | 5.6 |

**Table S7.**
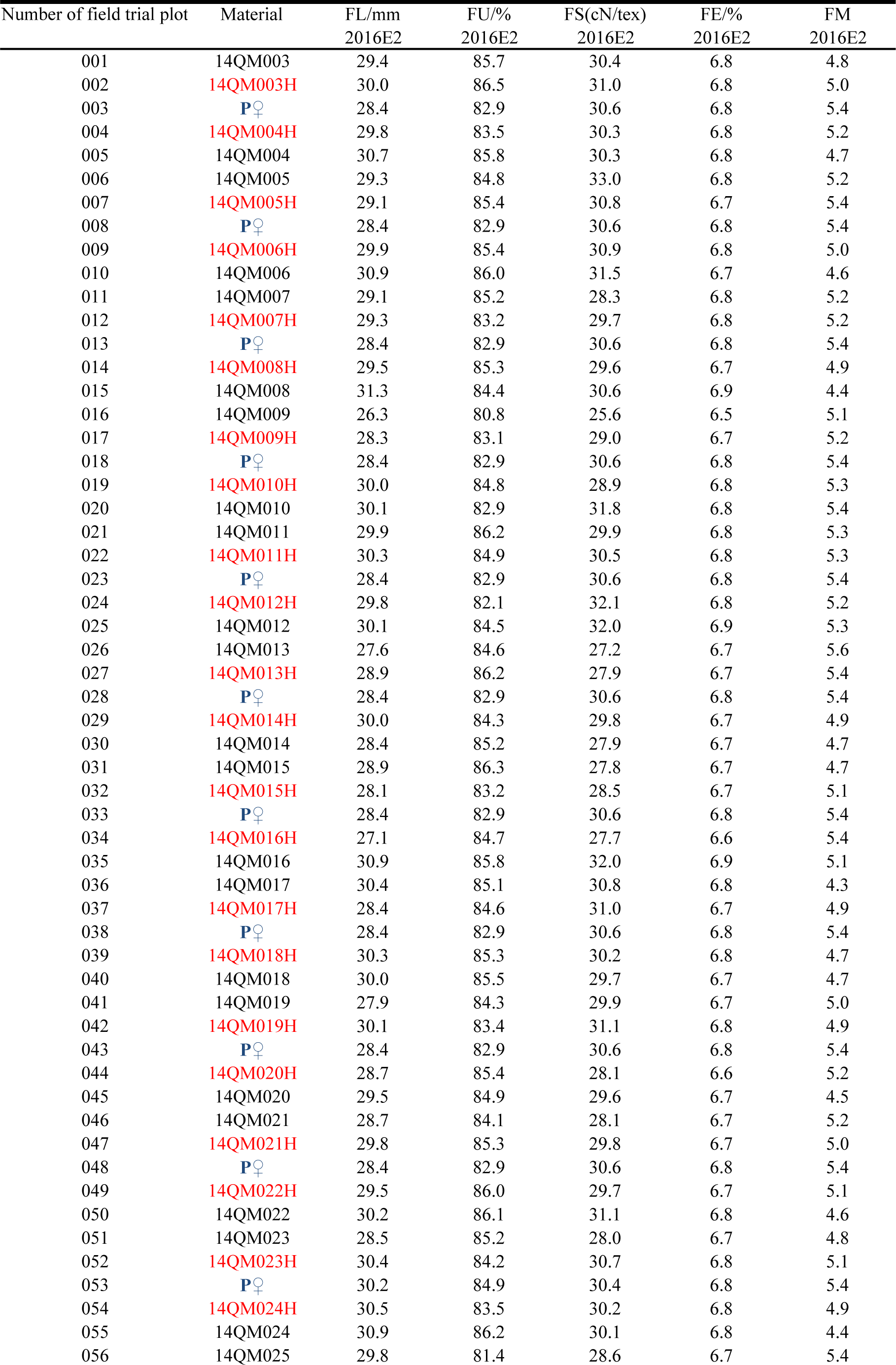
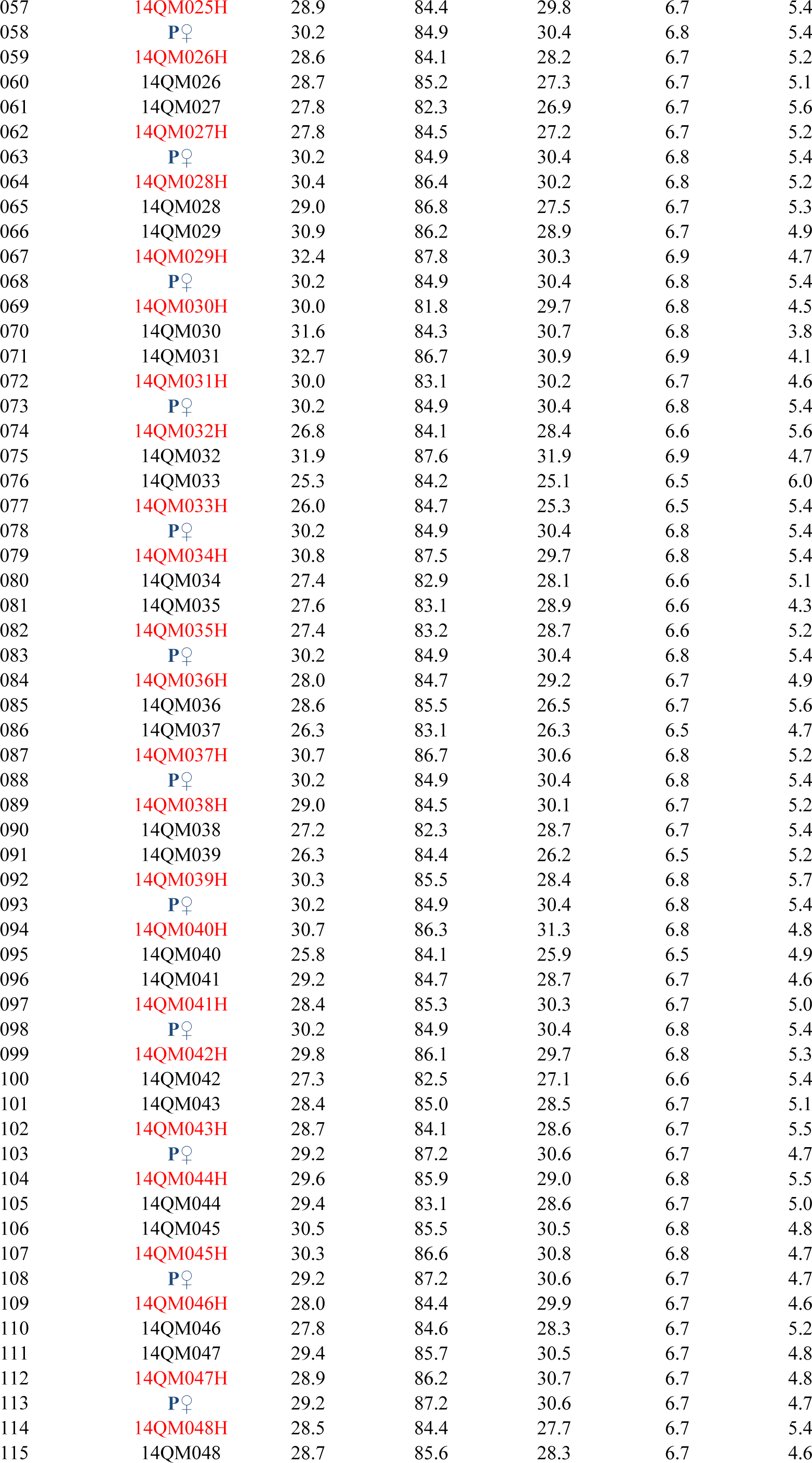
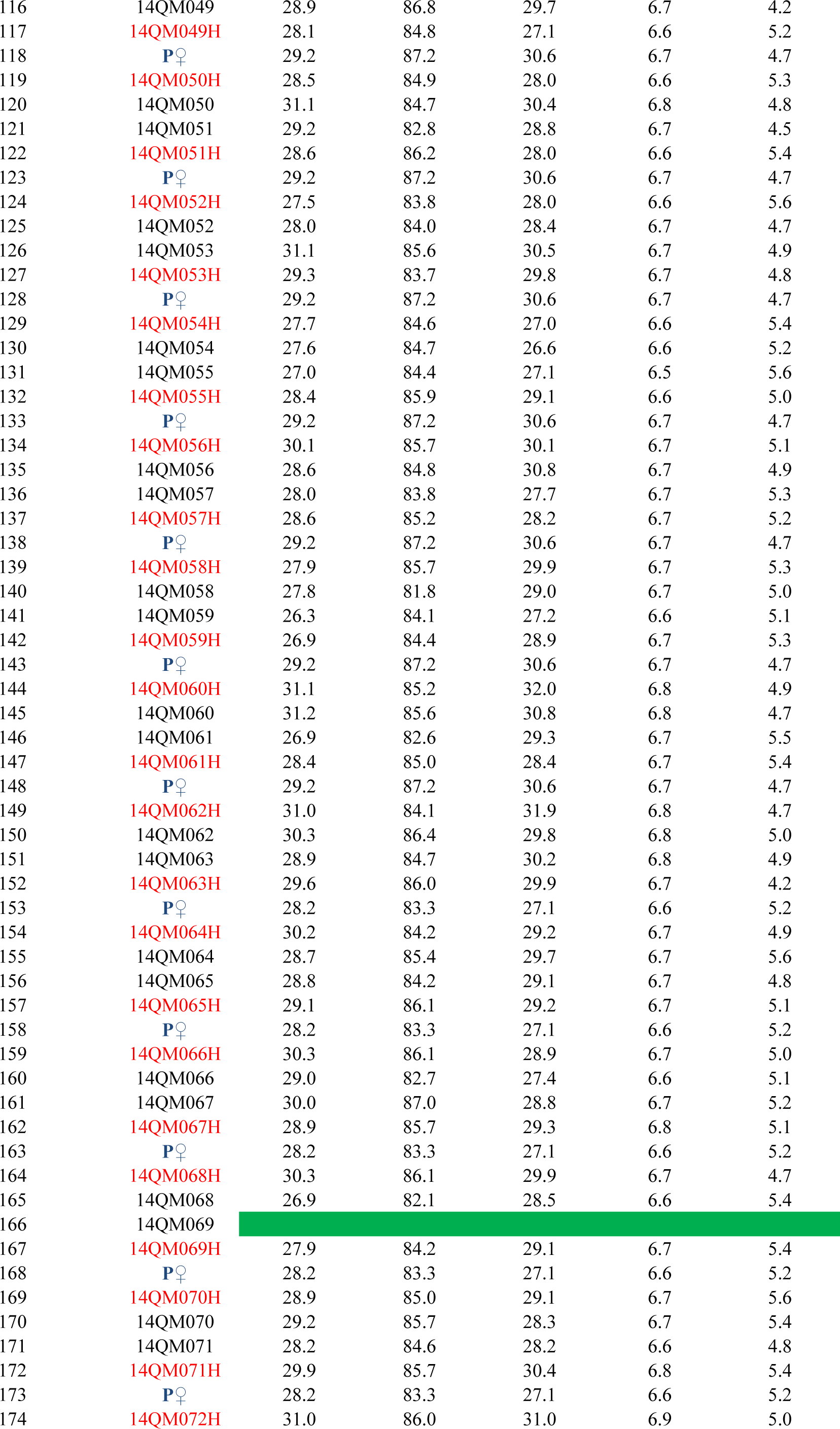
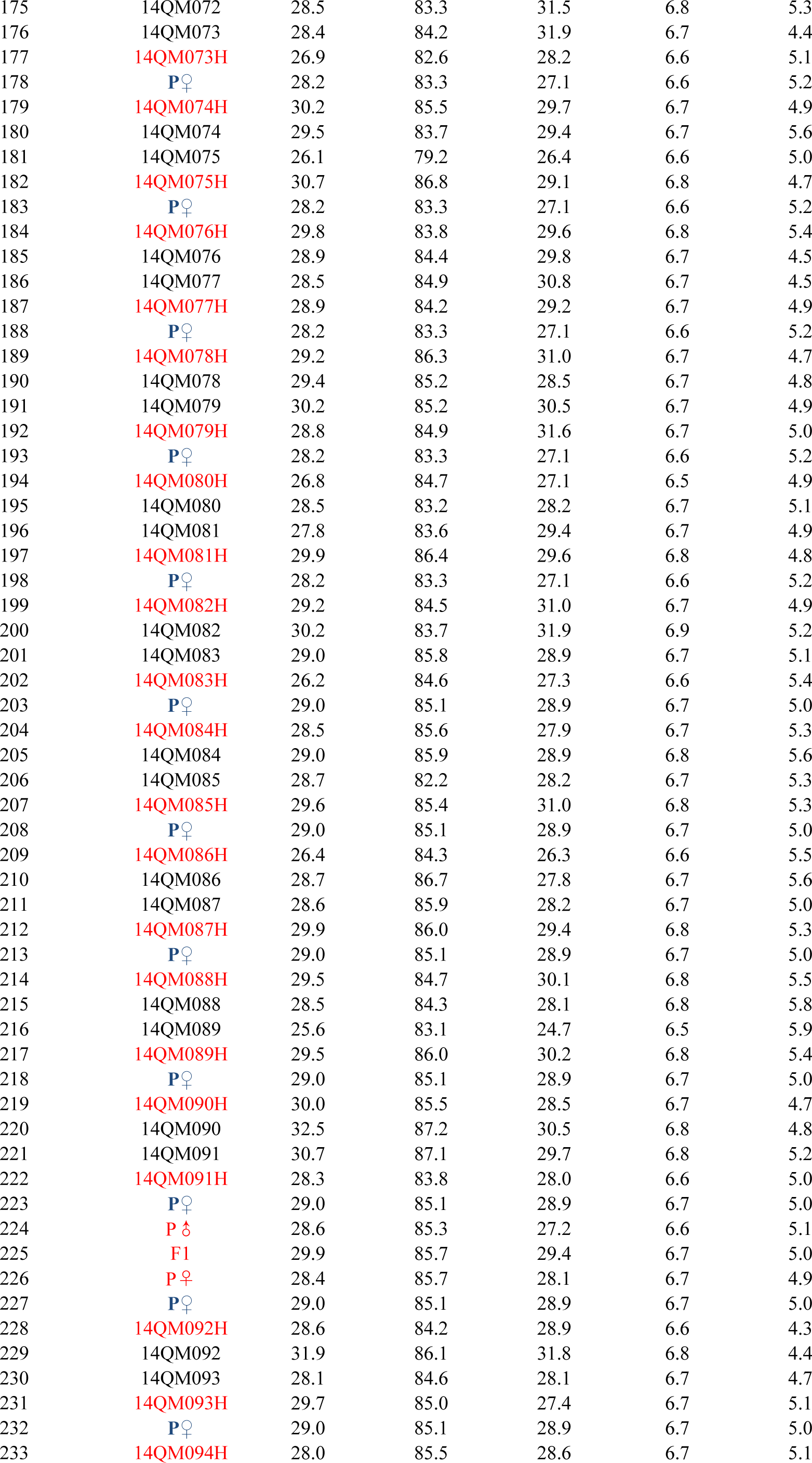
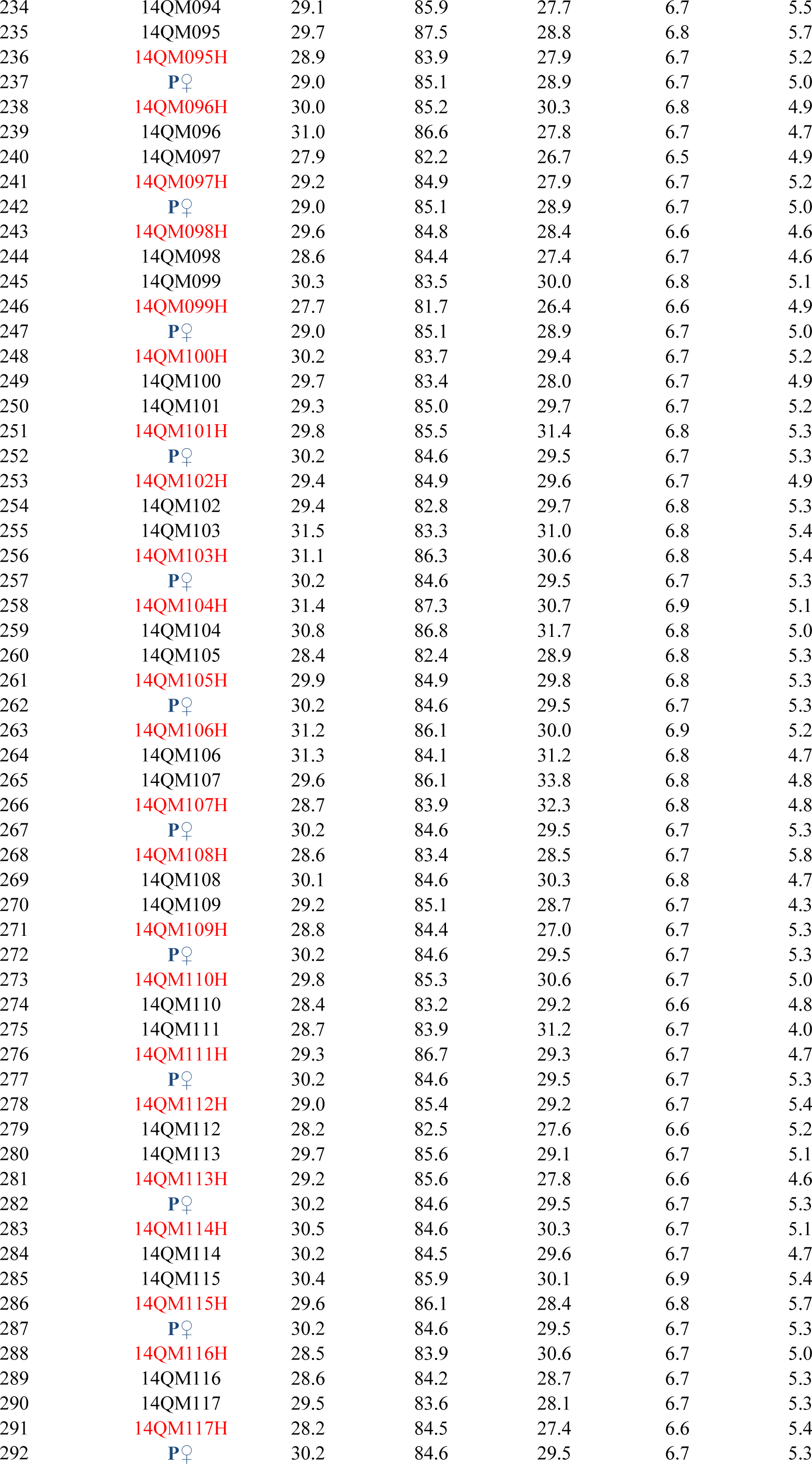
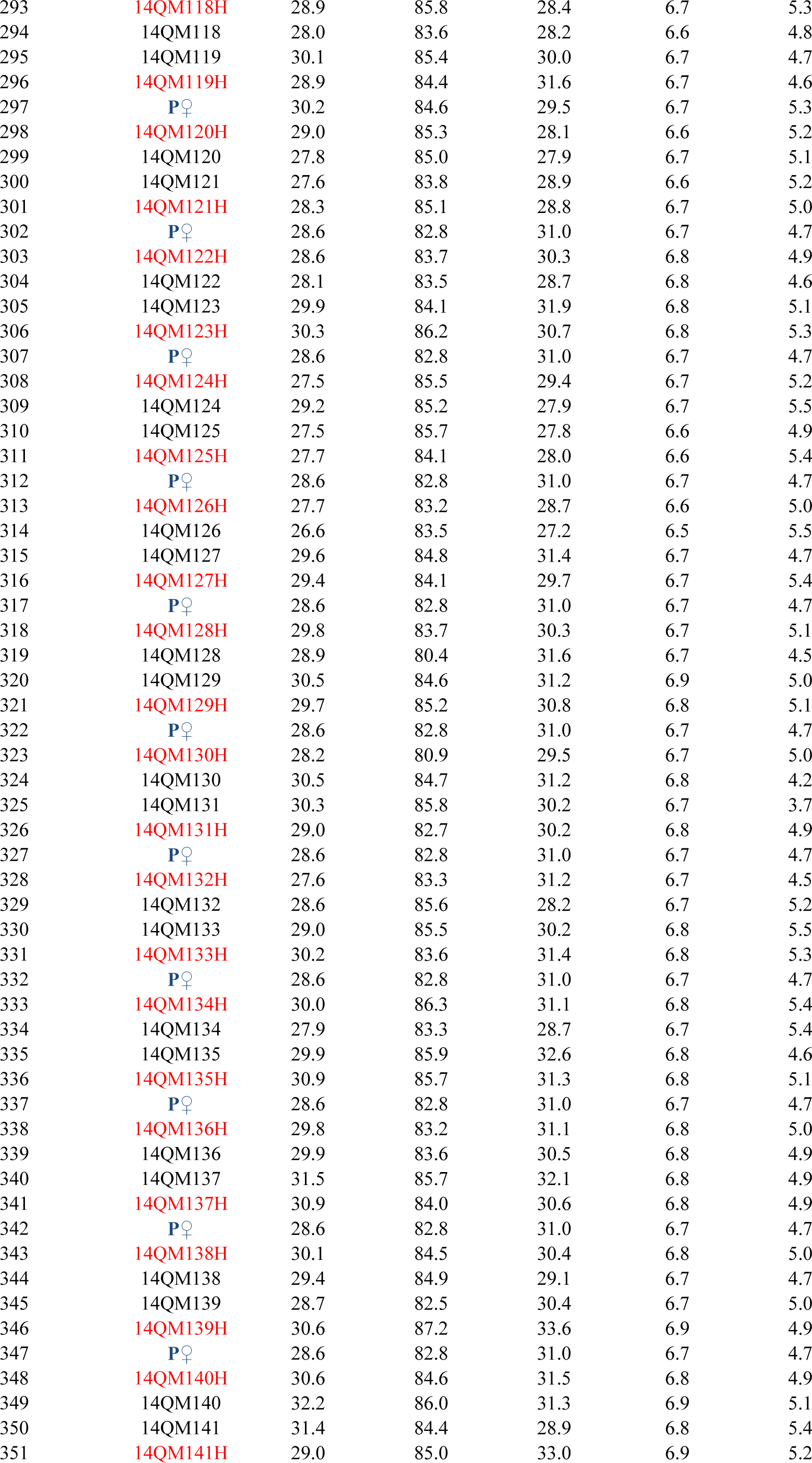
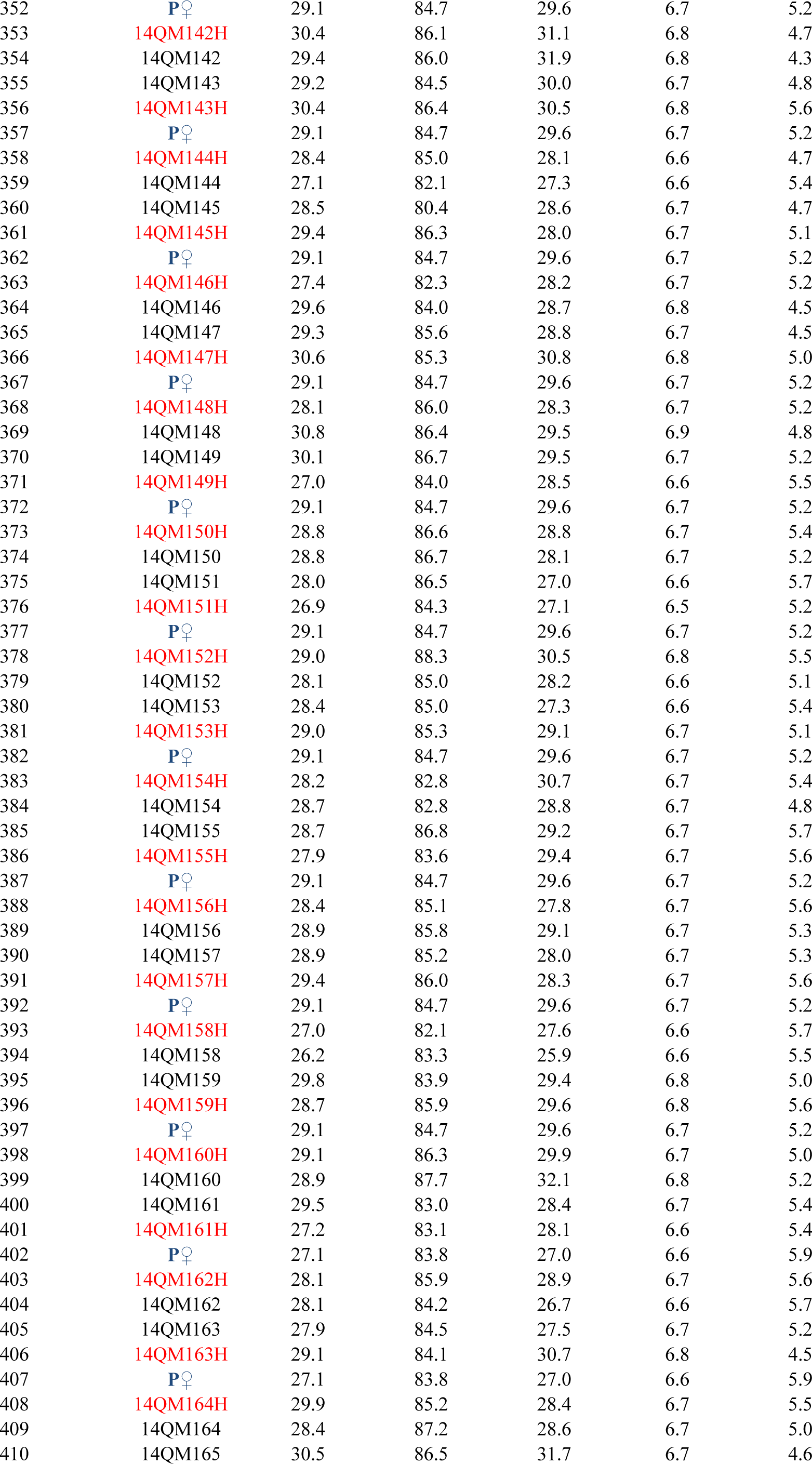
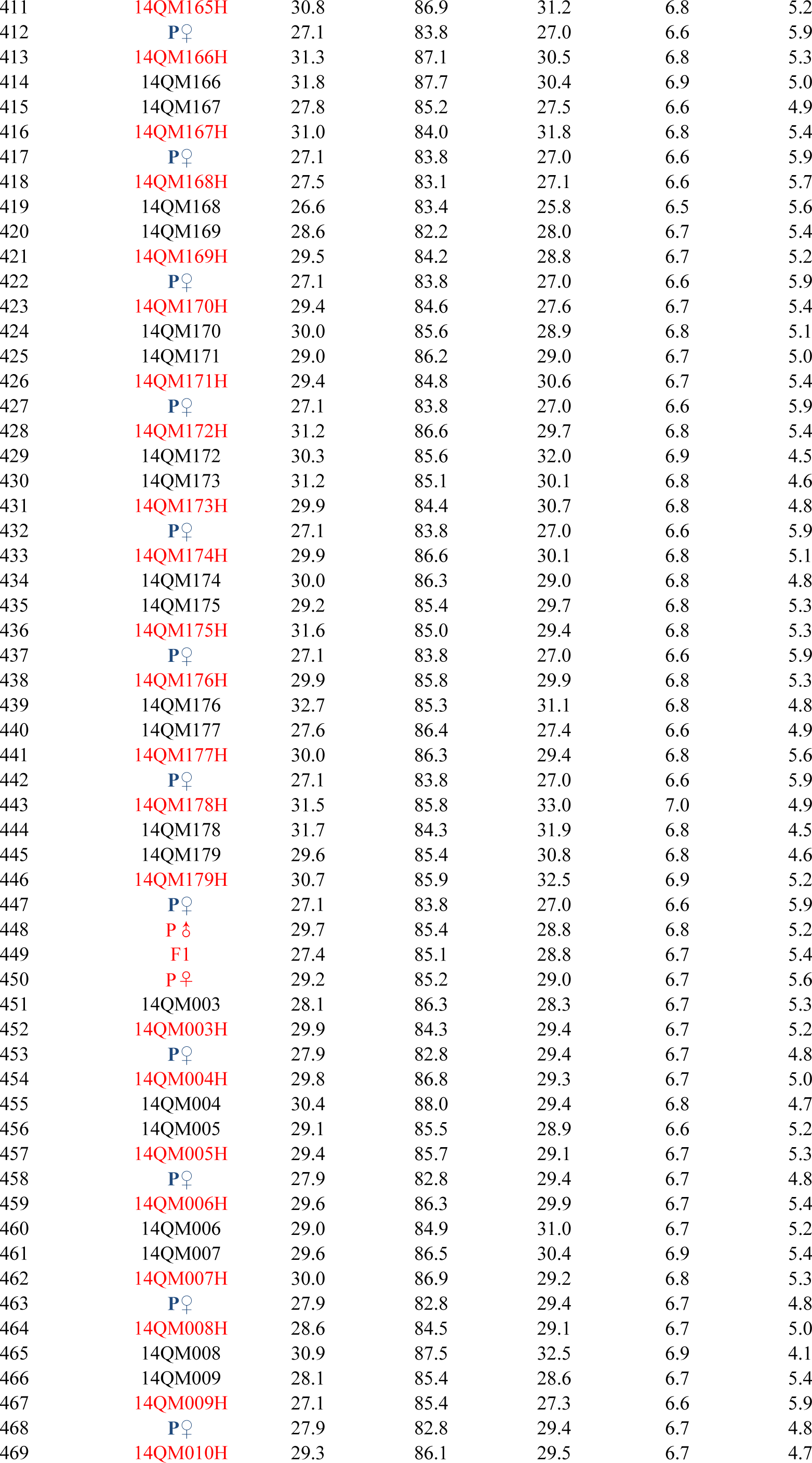
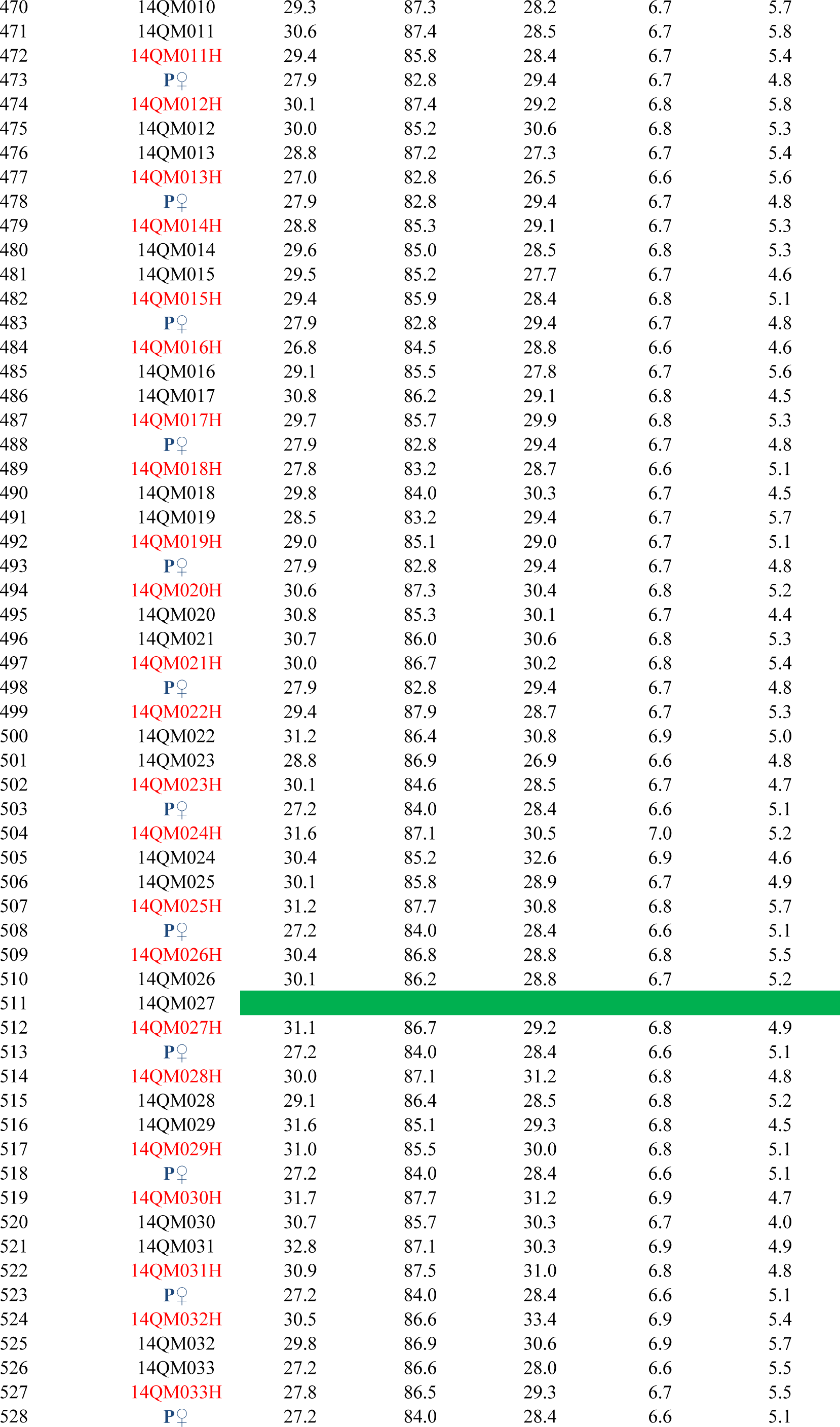
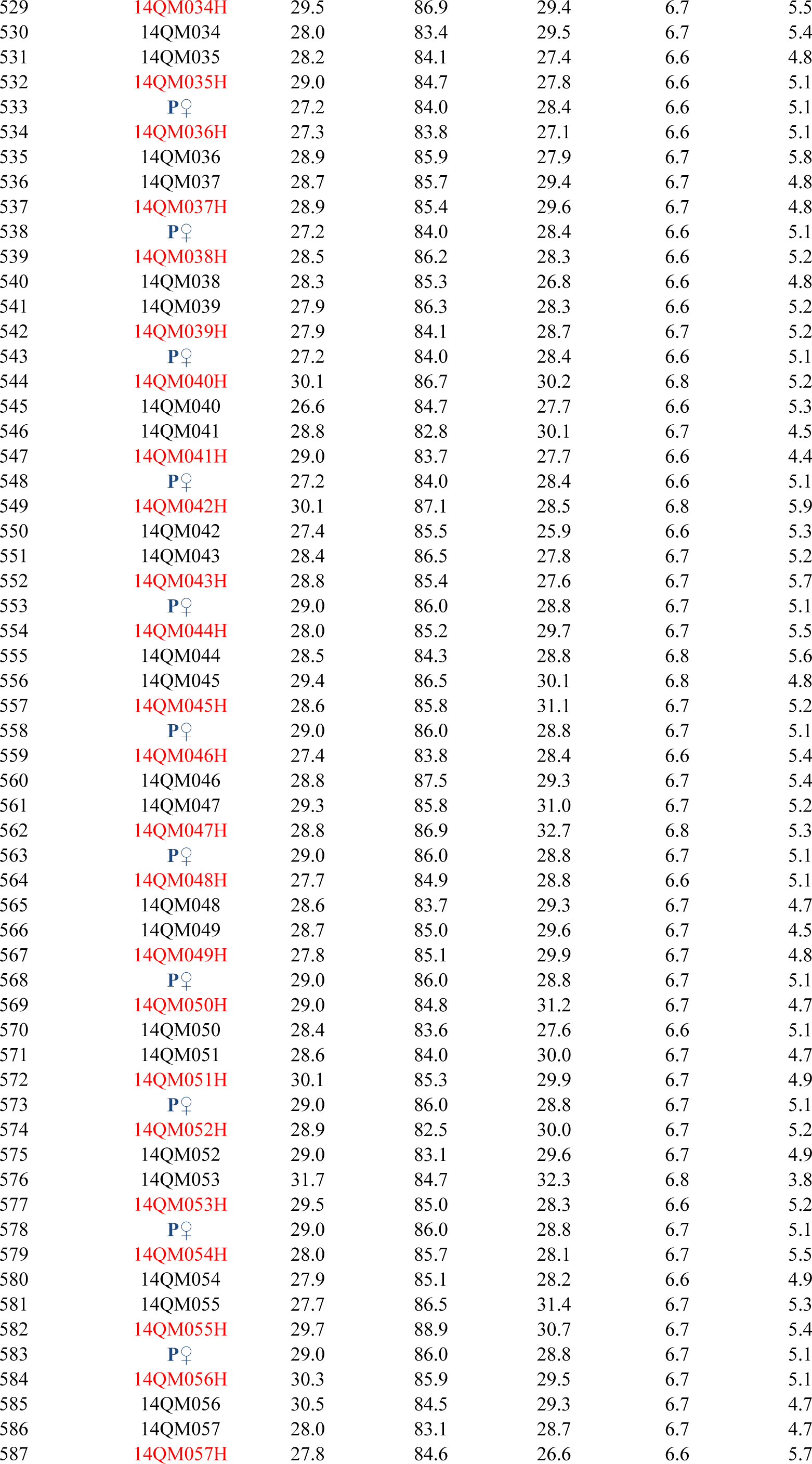
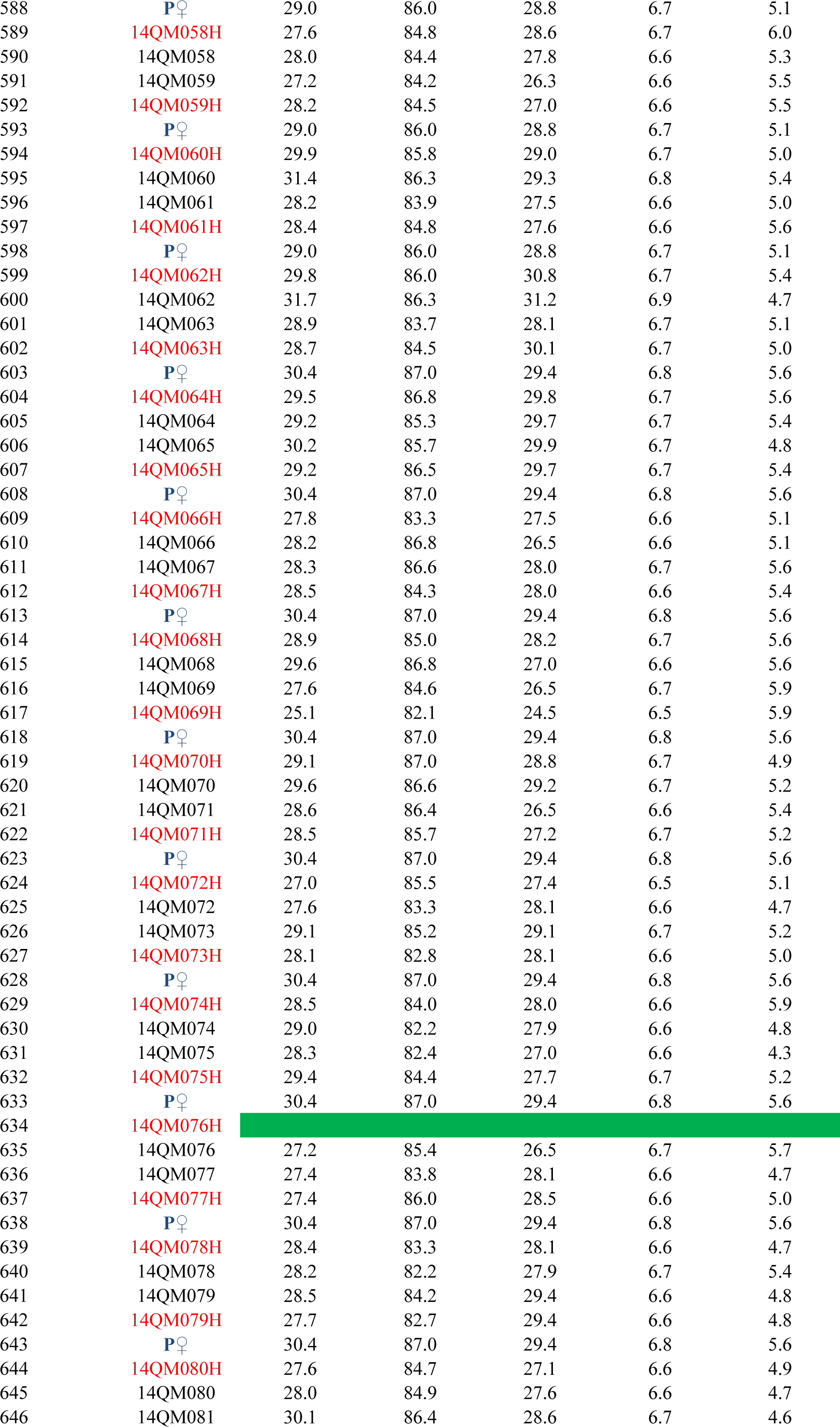
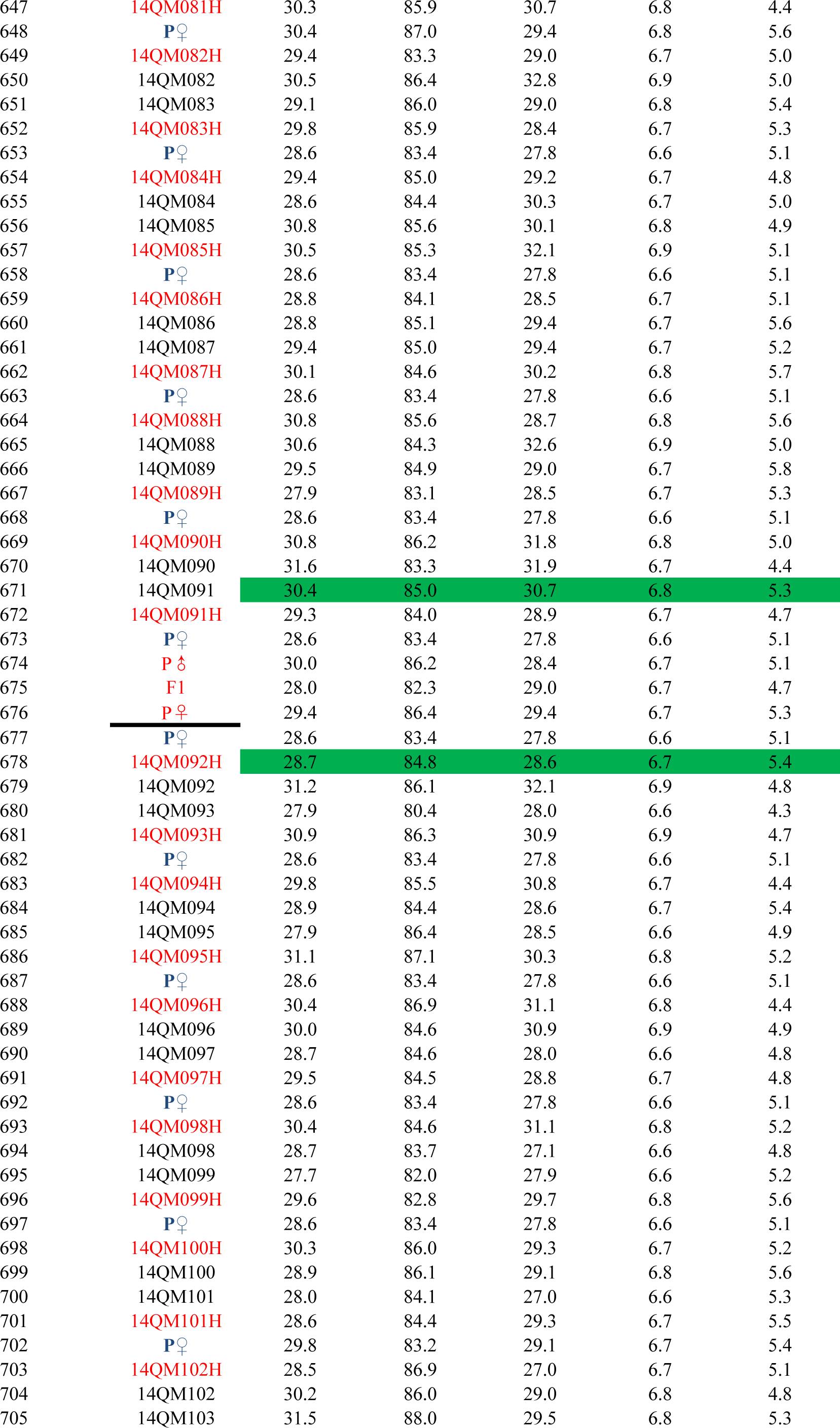
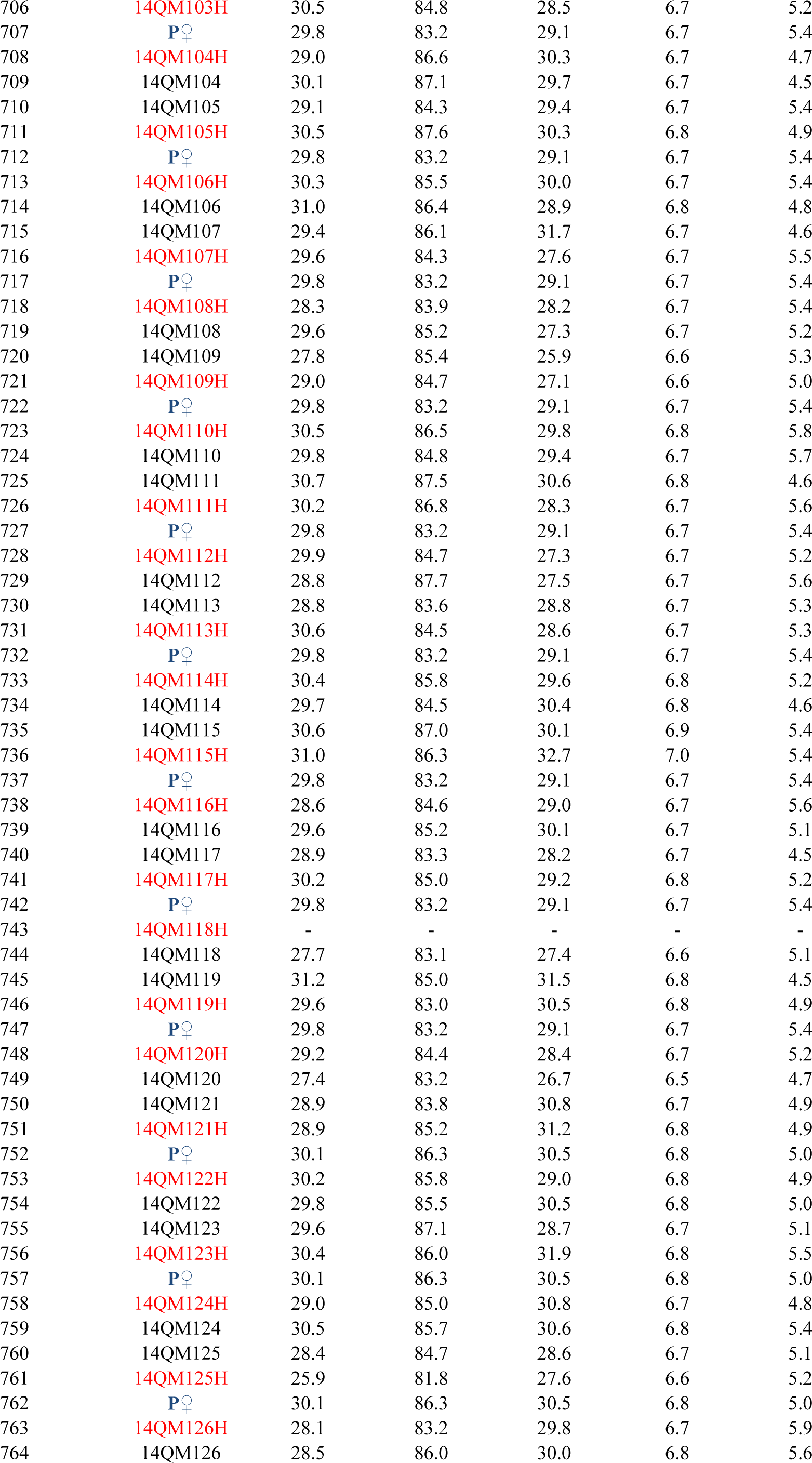
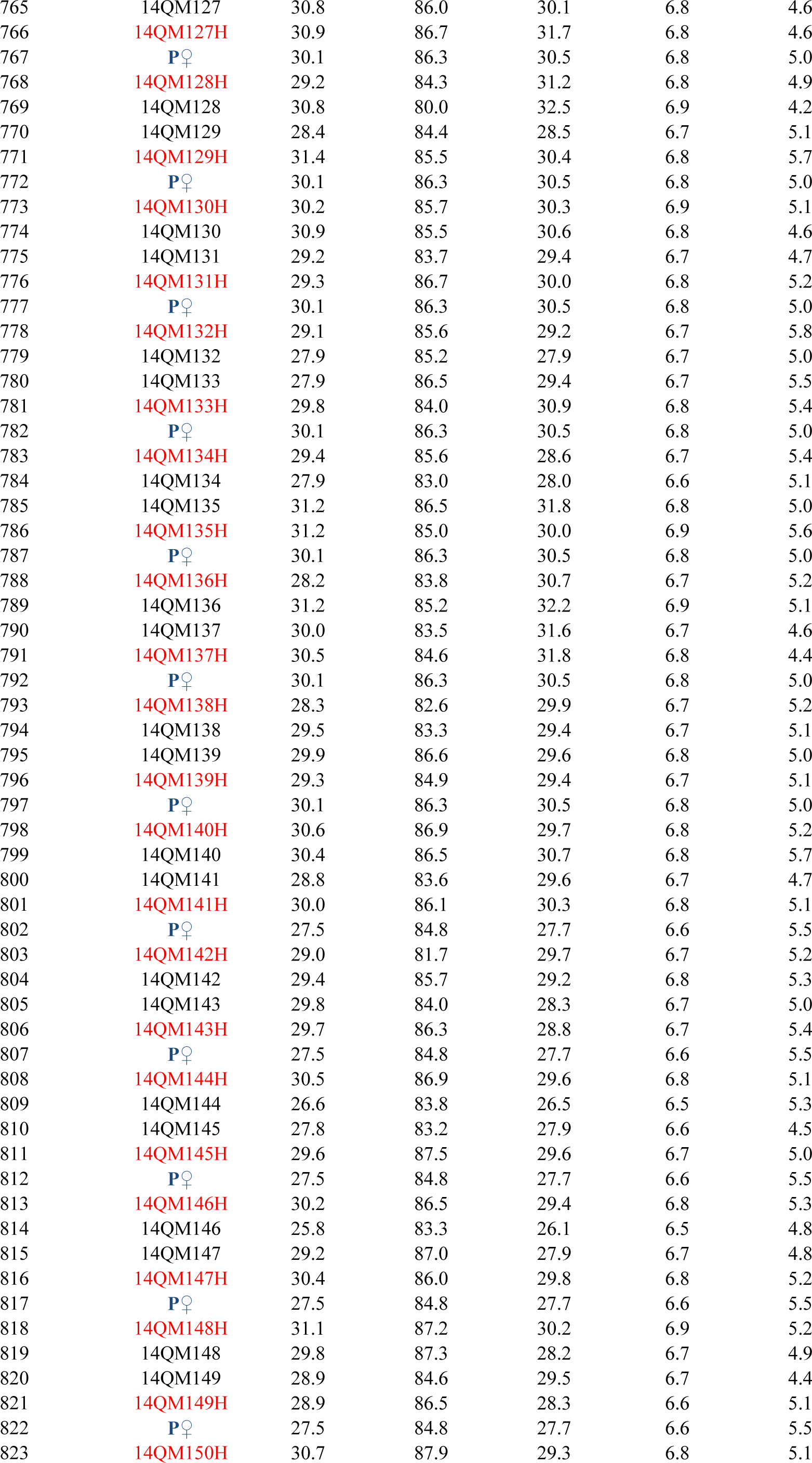
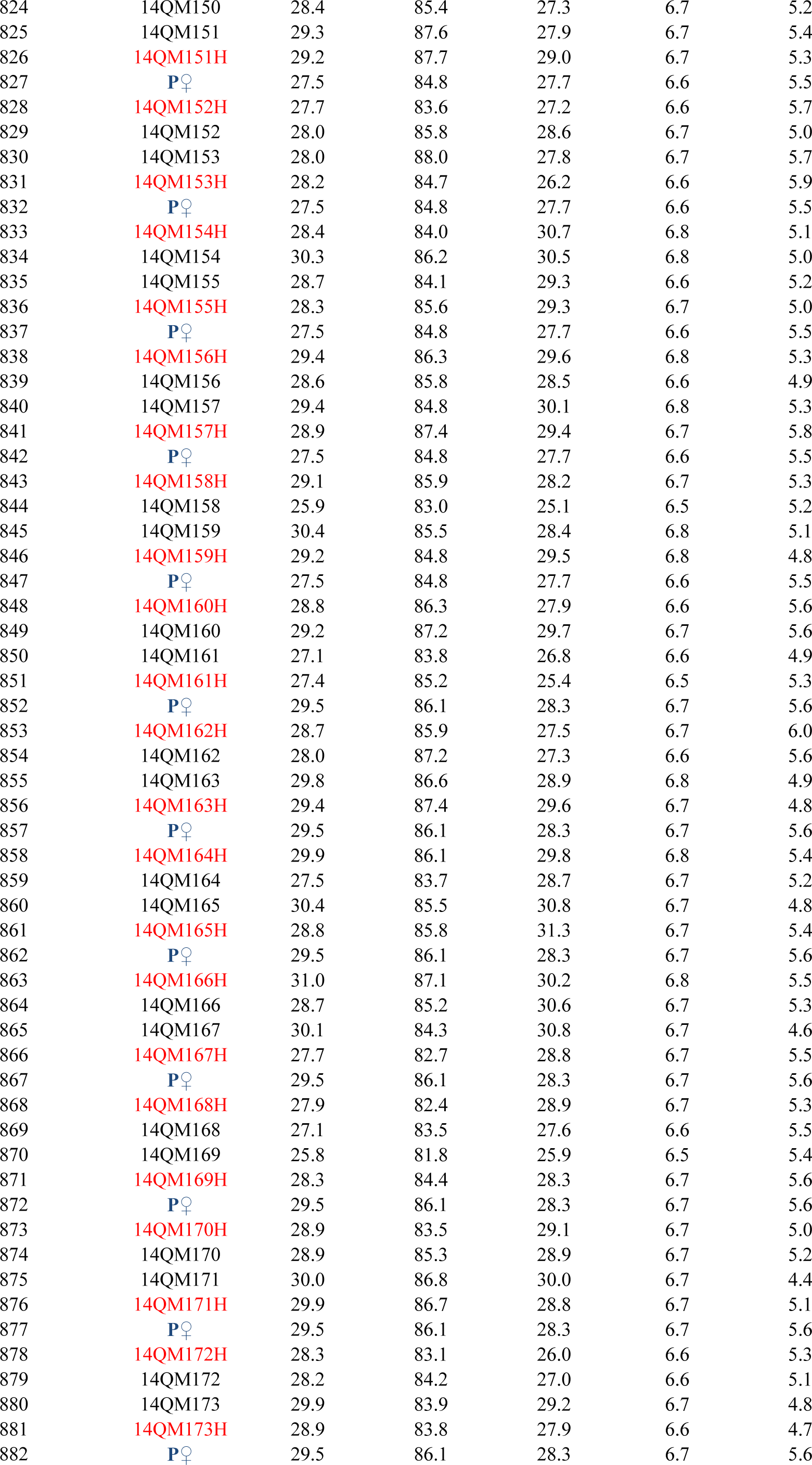
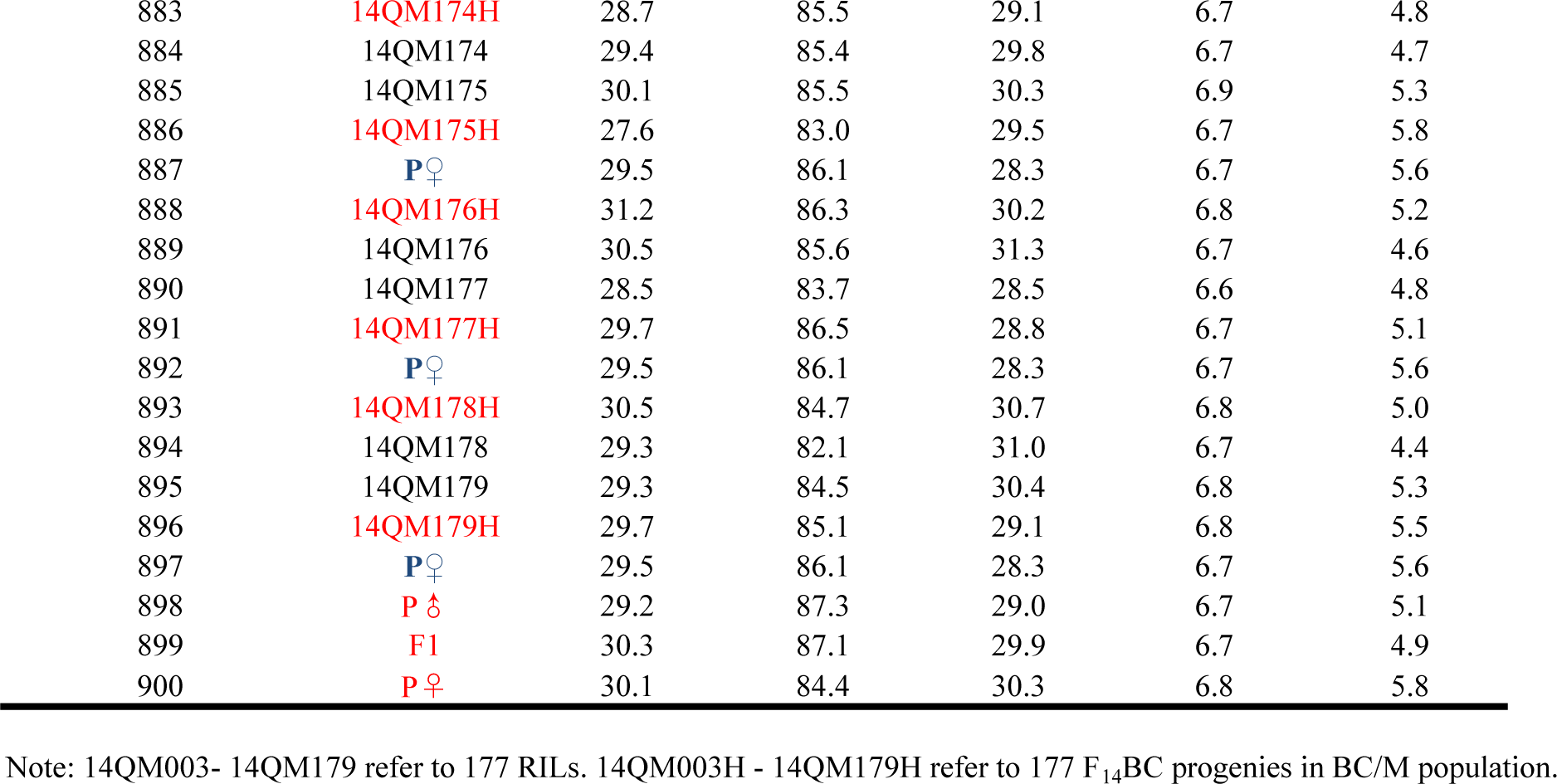
Phenotype values of fiber quality traits in field trial of BC/P population in this study

**Table S8.**
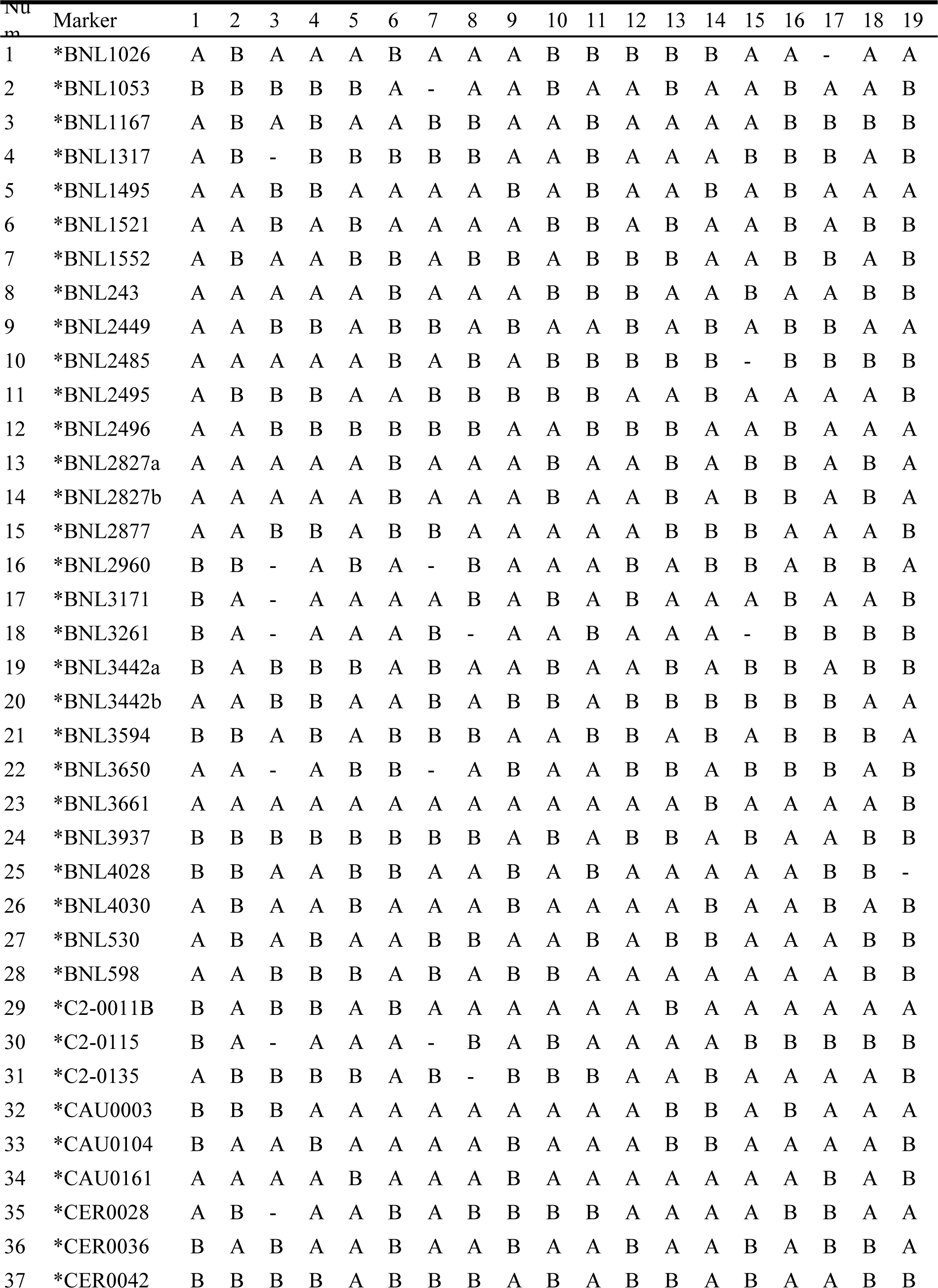
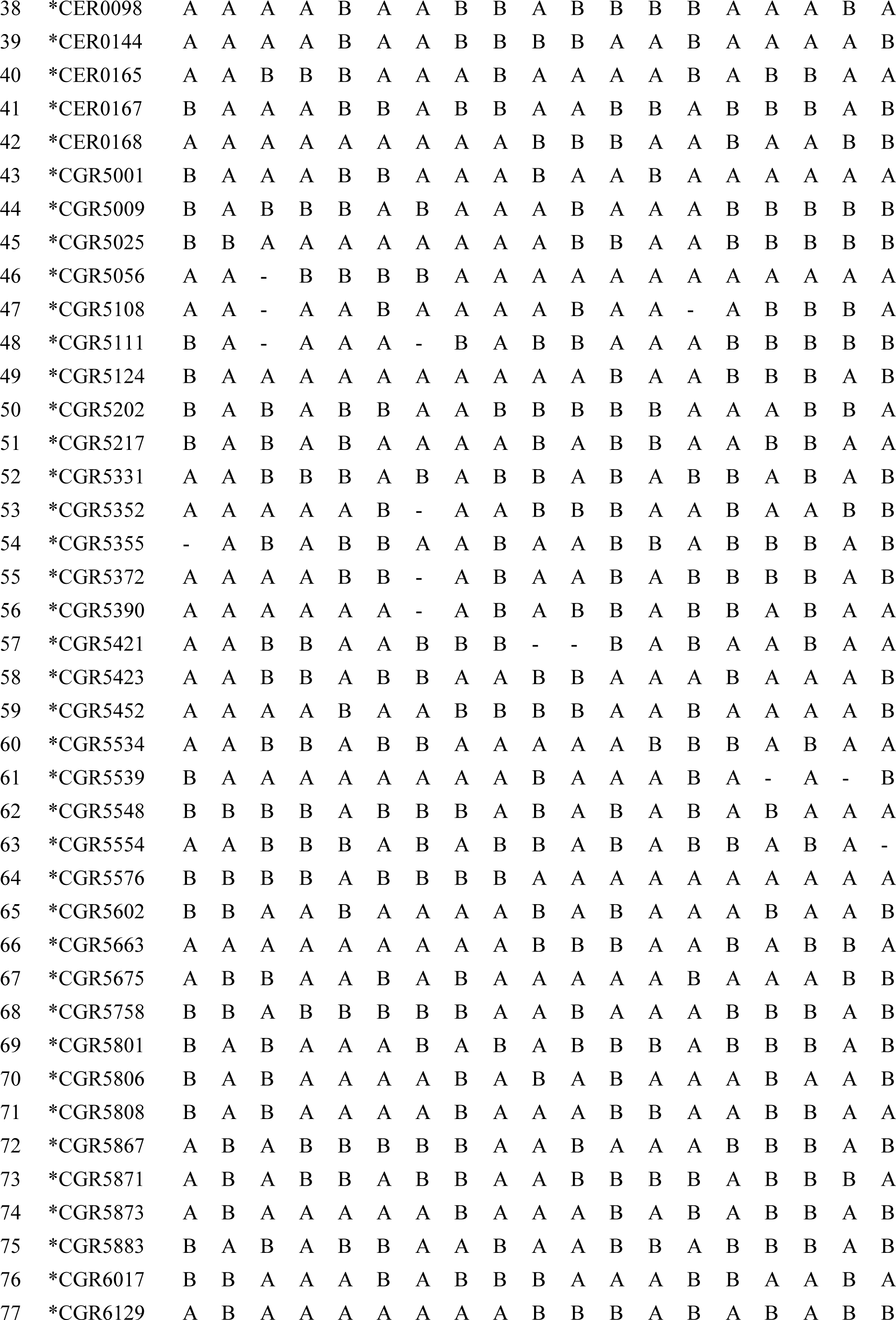
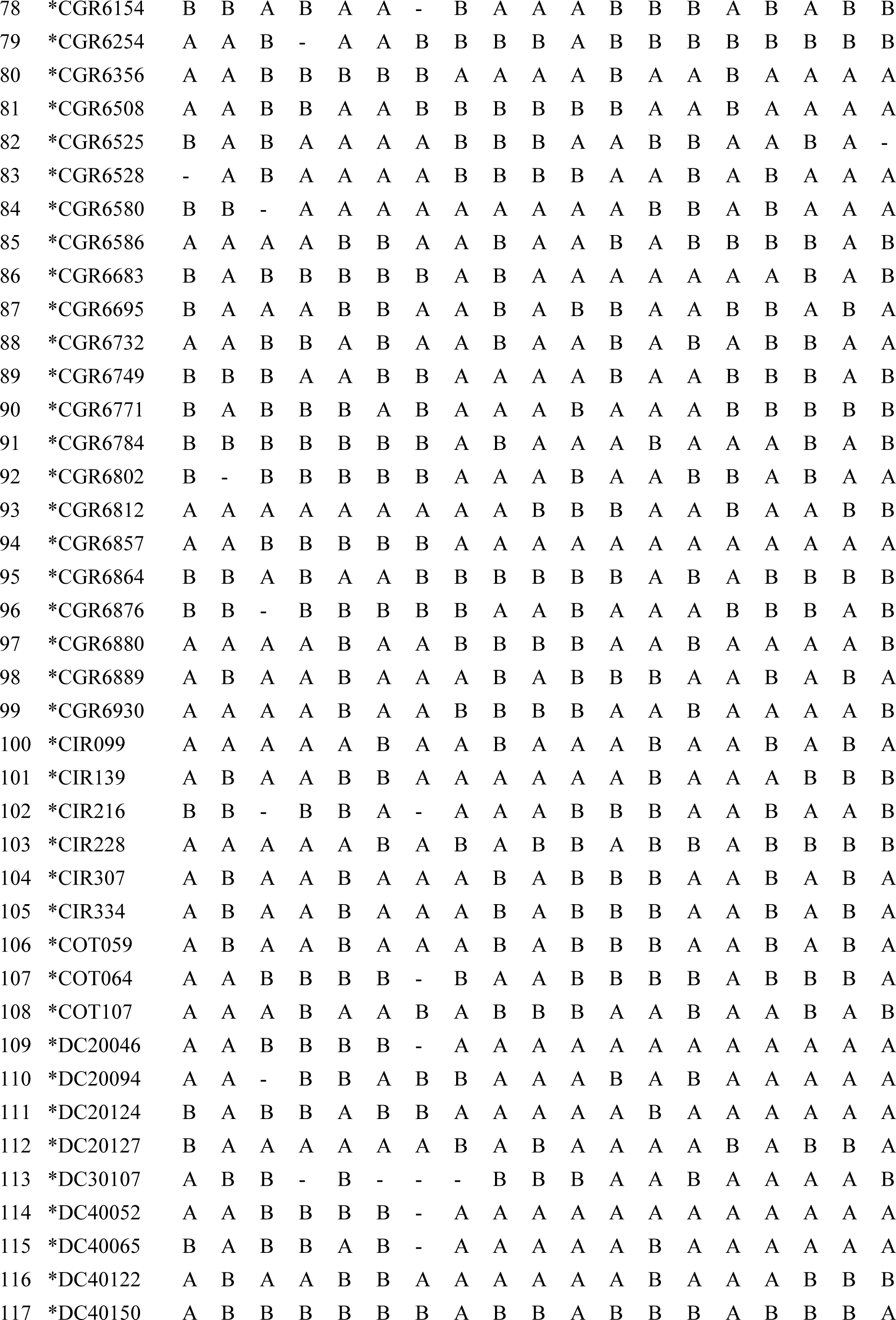
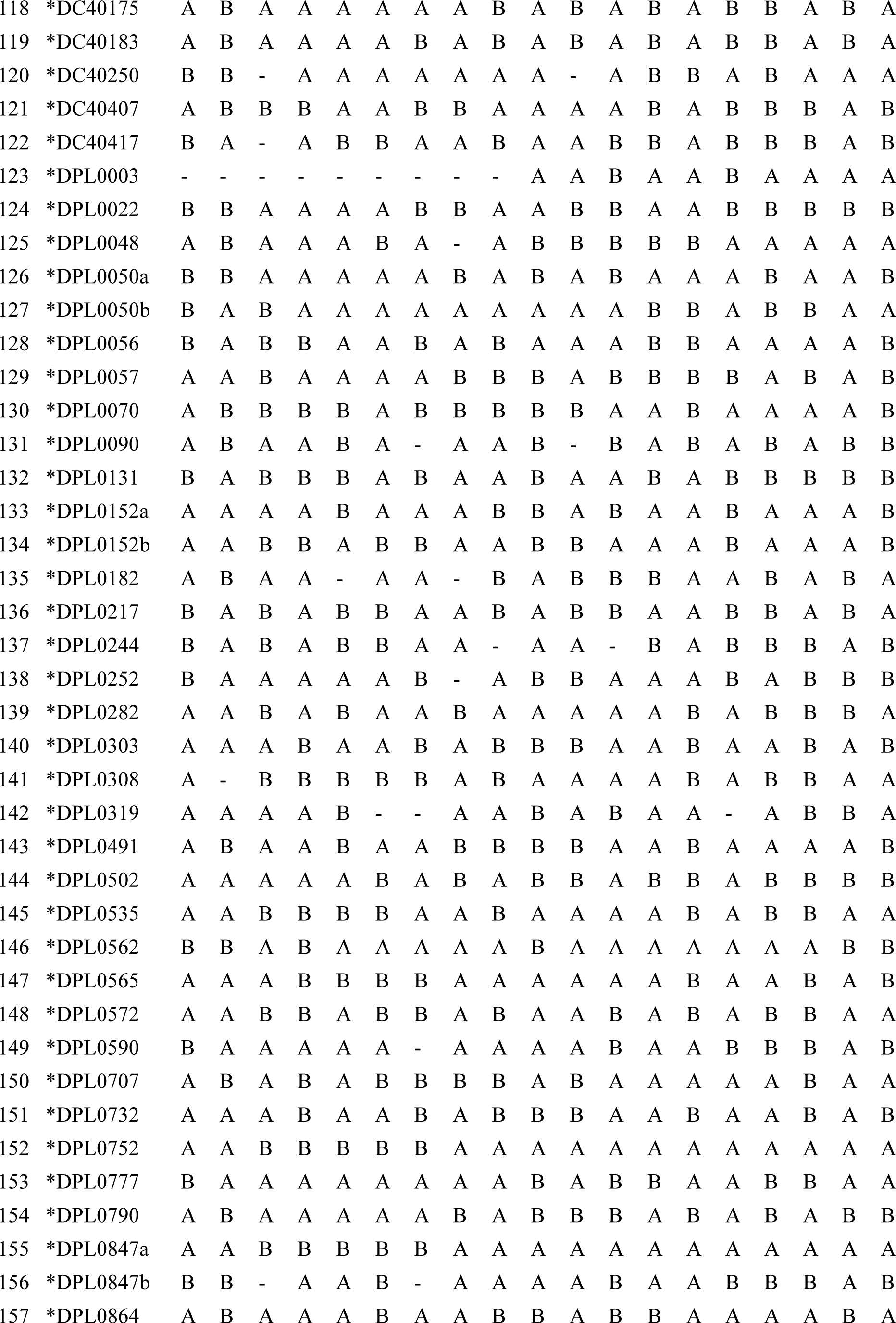
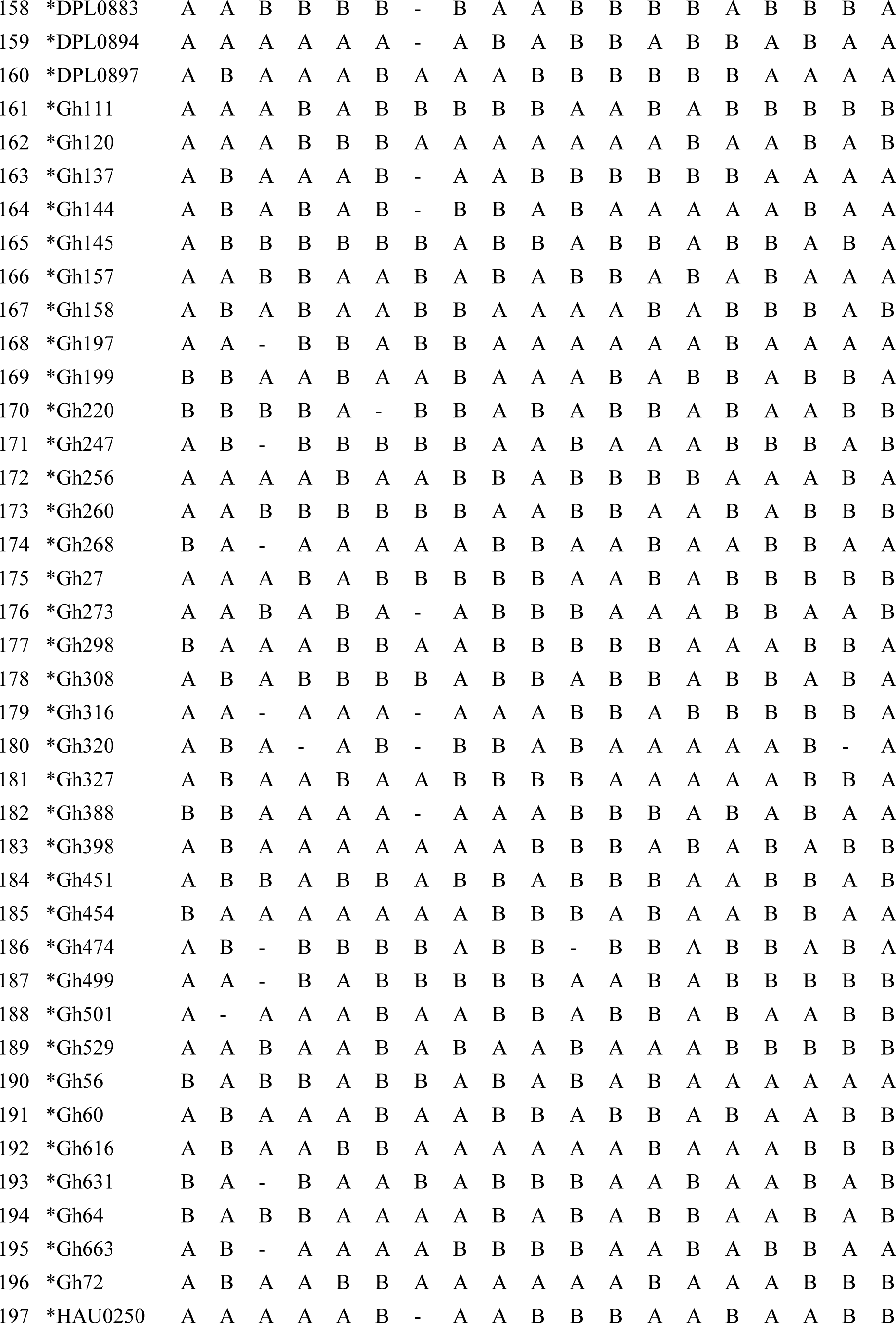
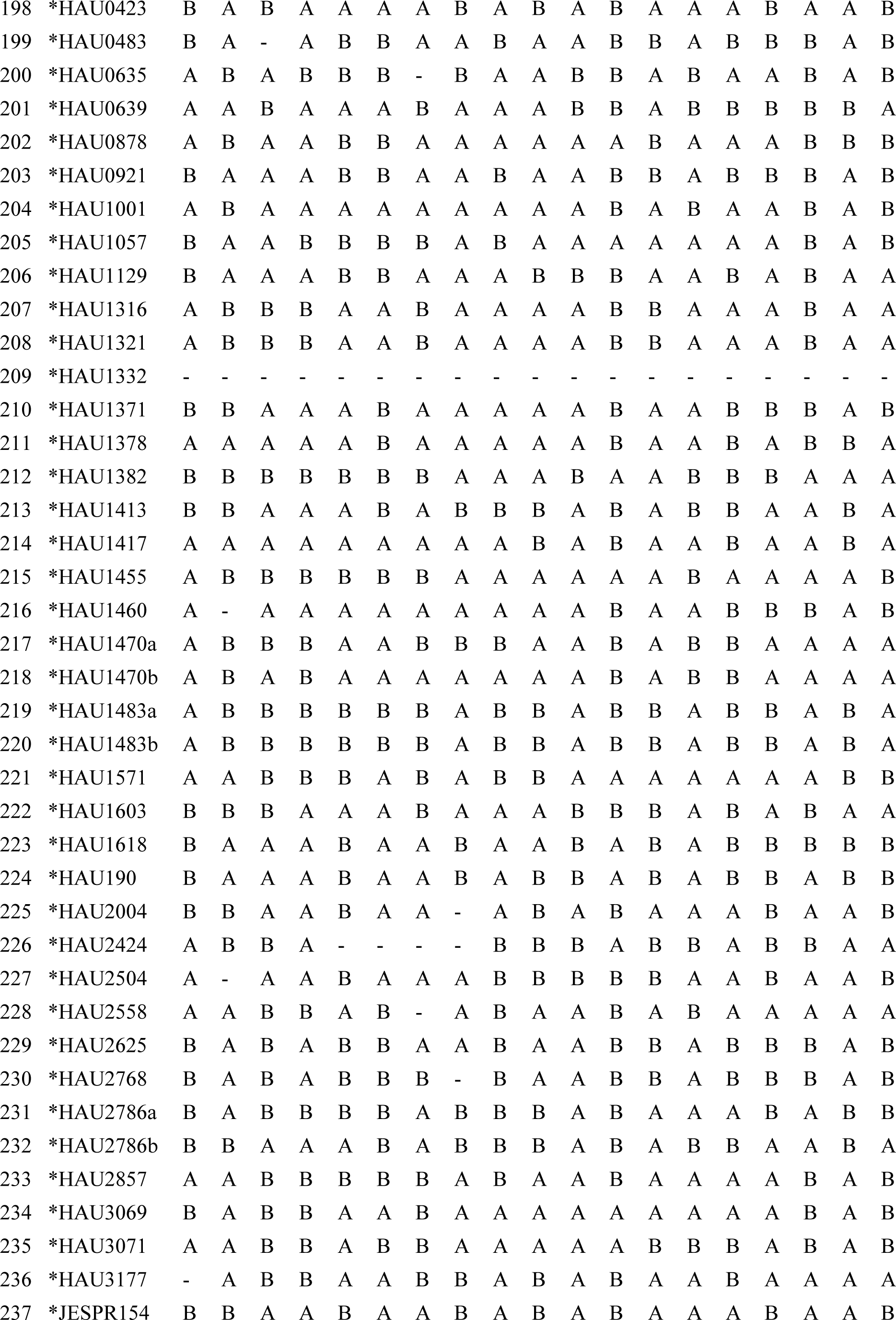
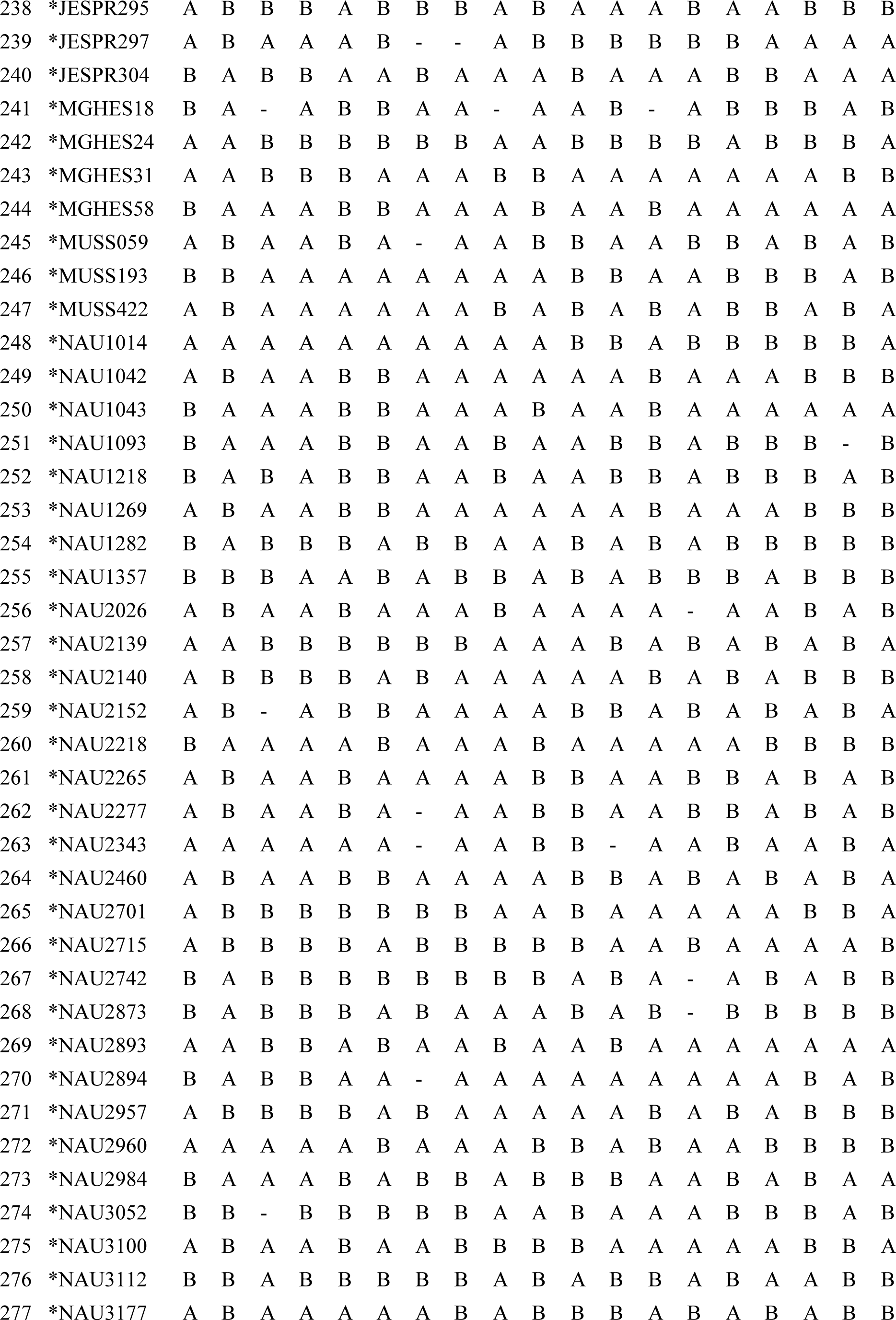
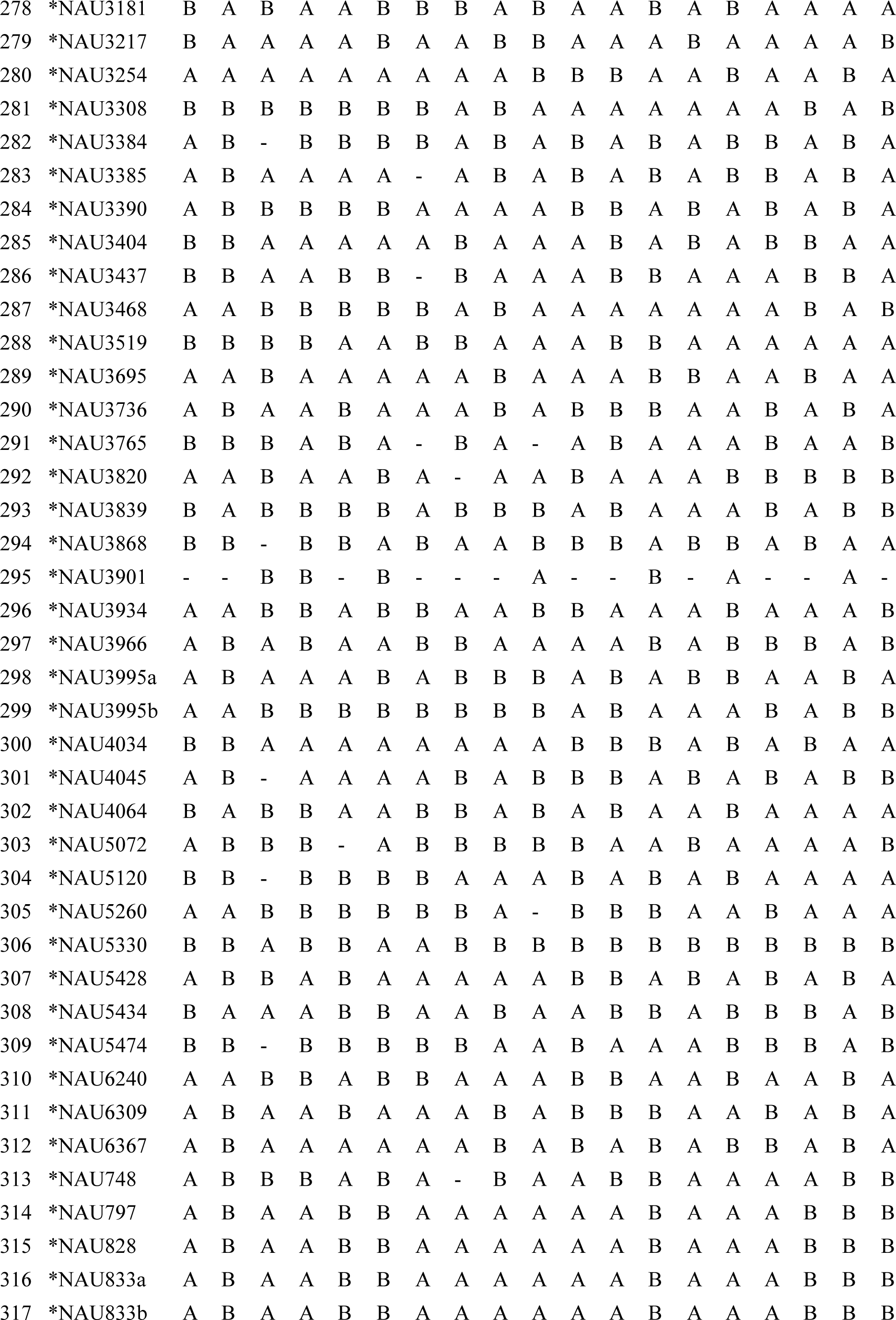
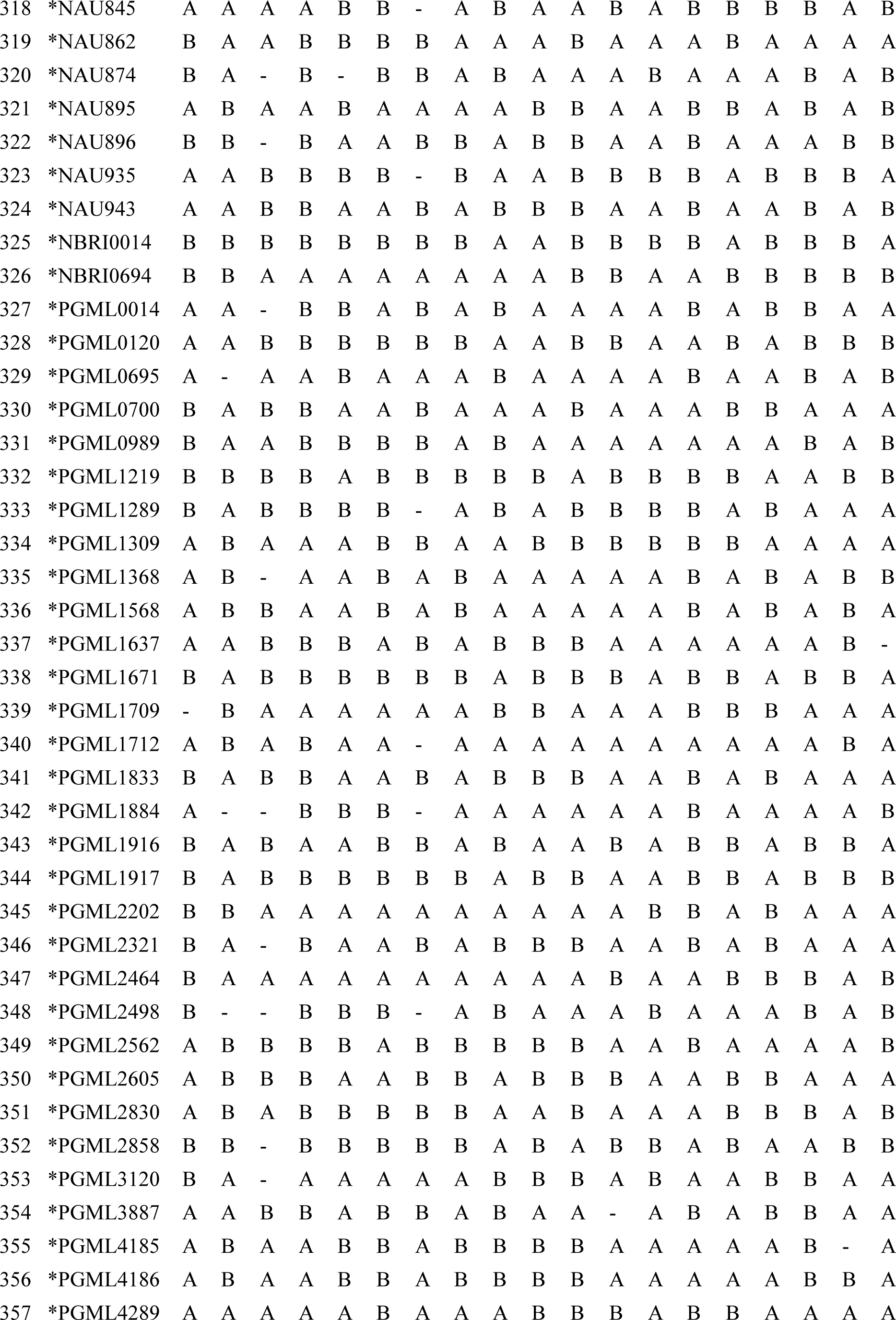
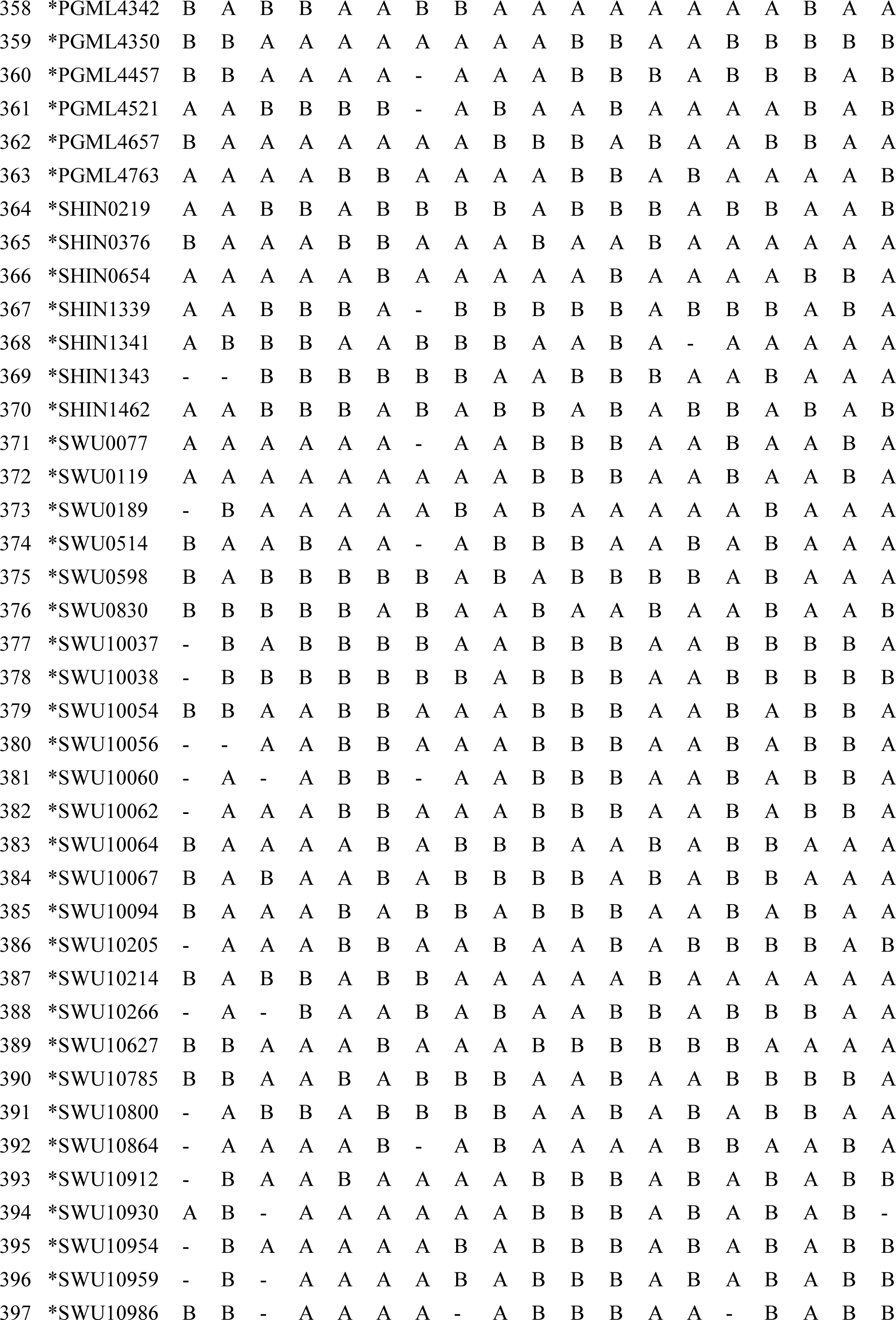
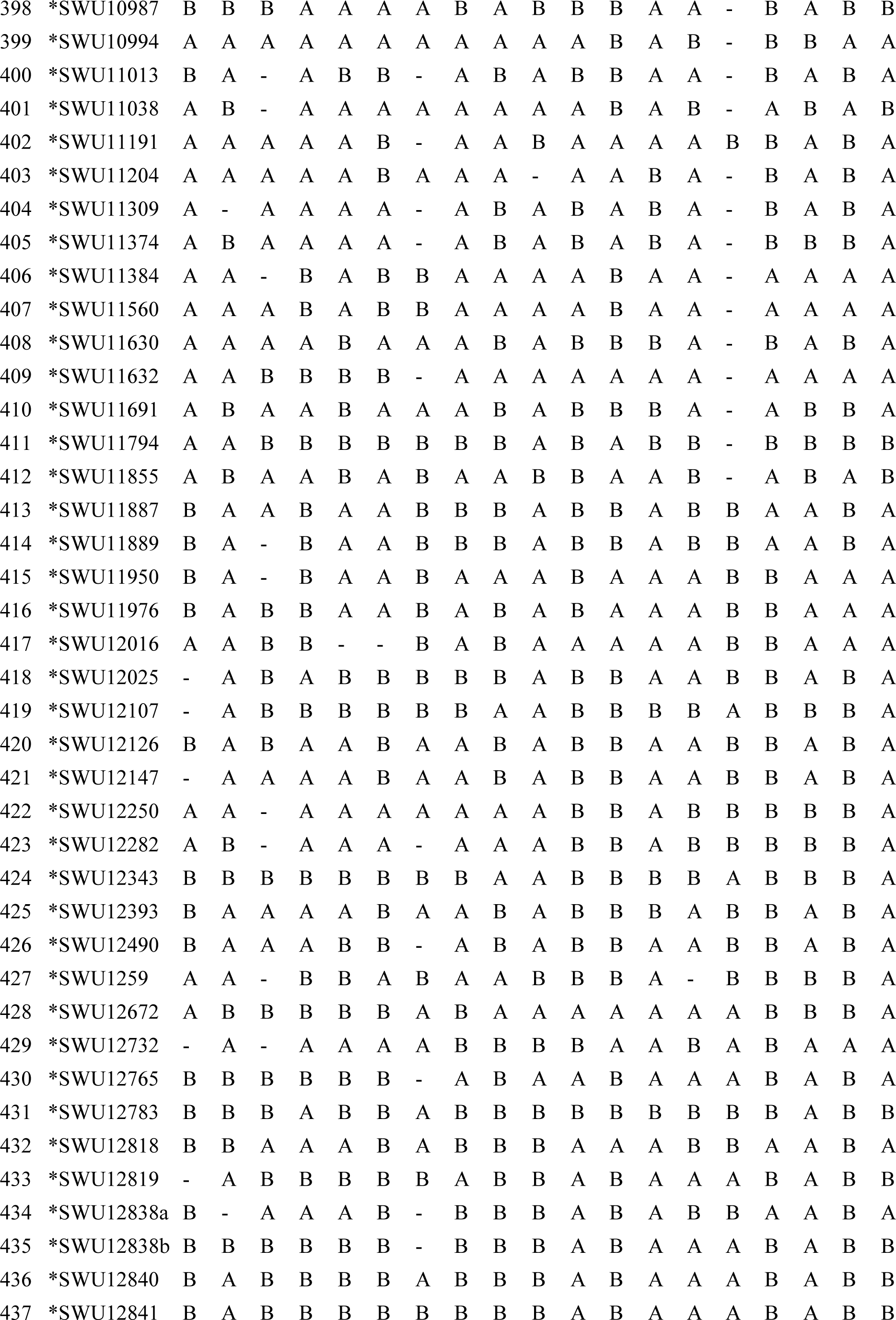
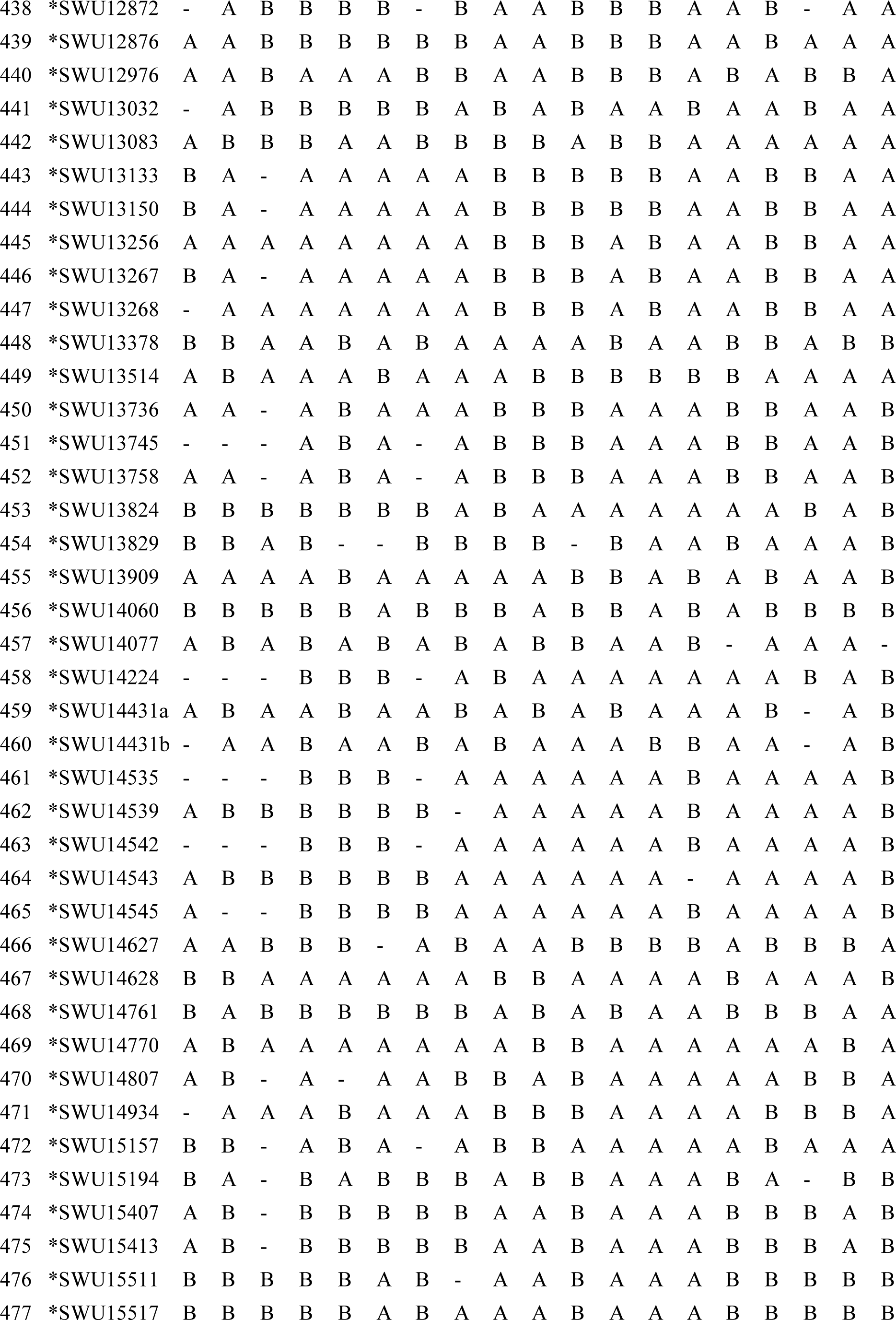
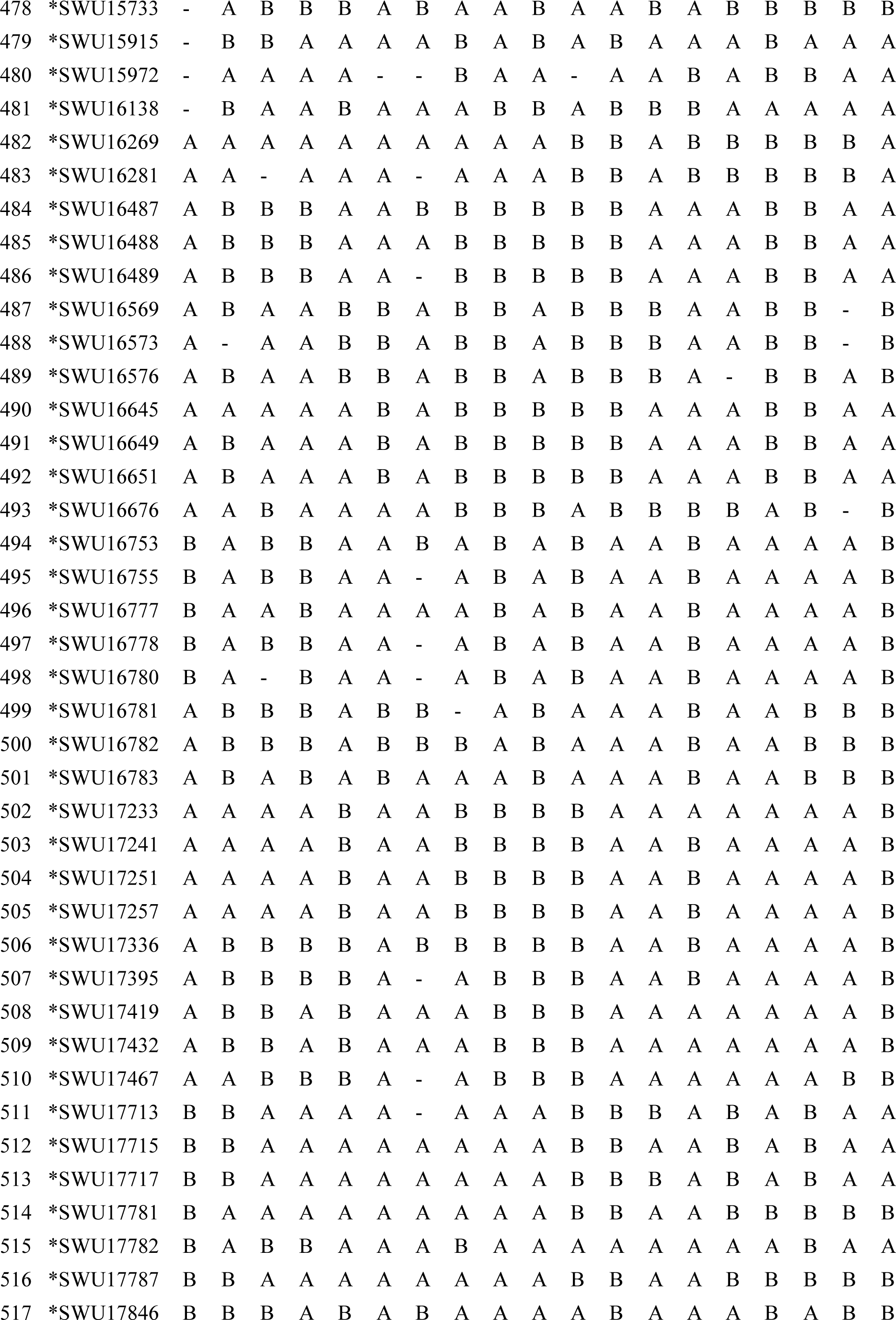
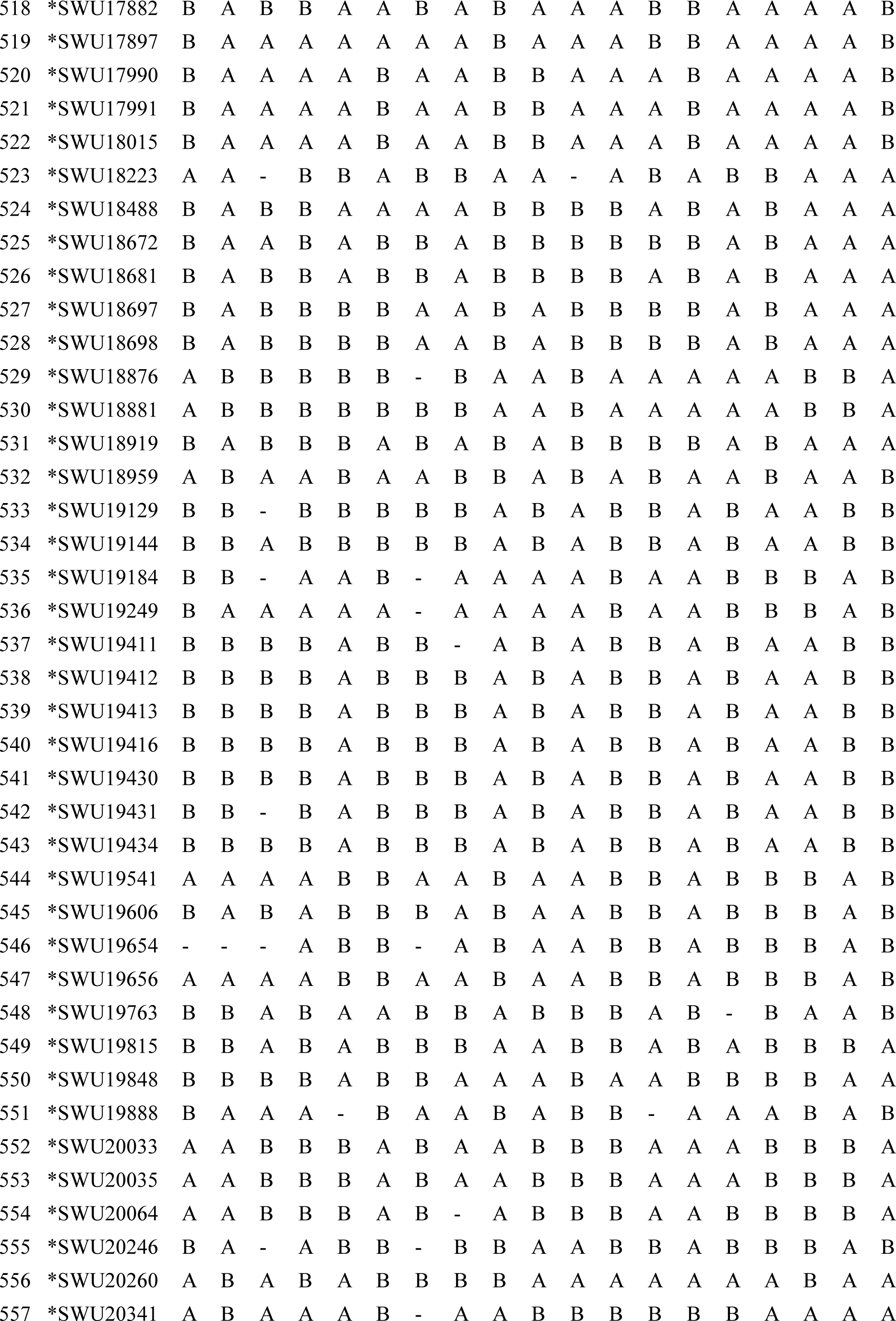
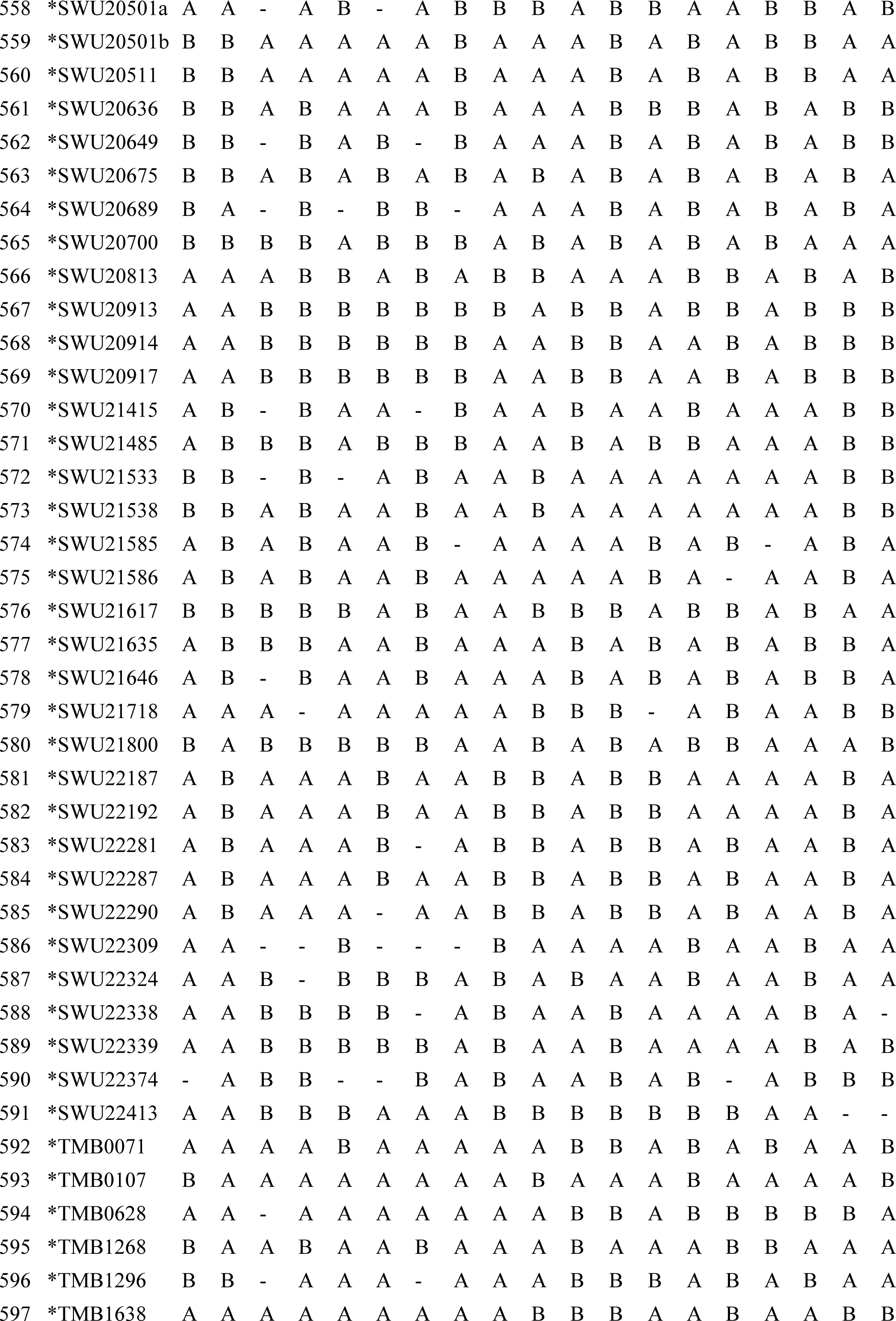
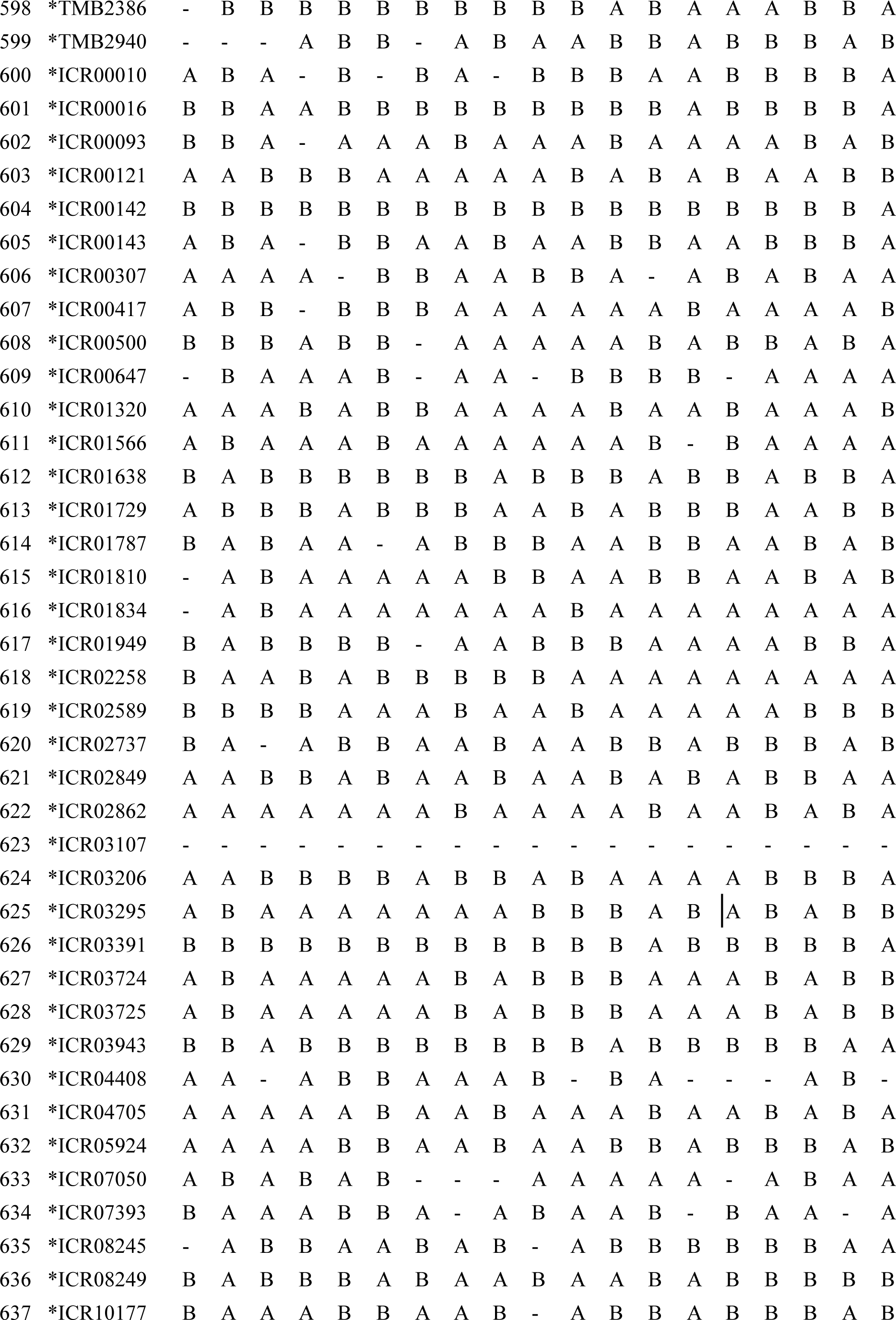
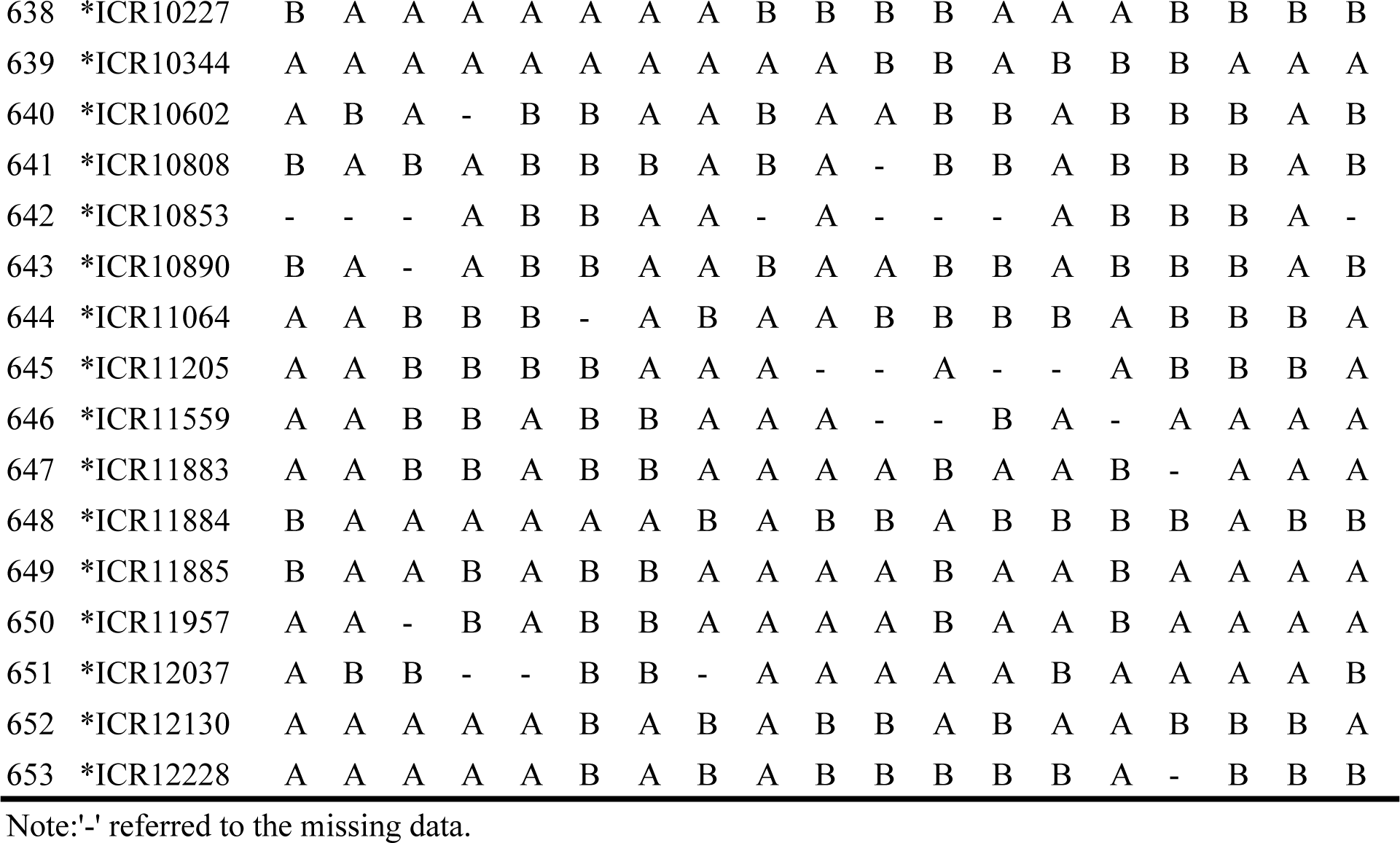
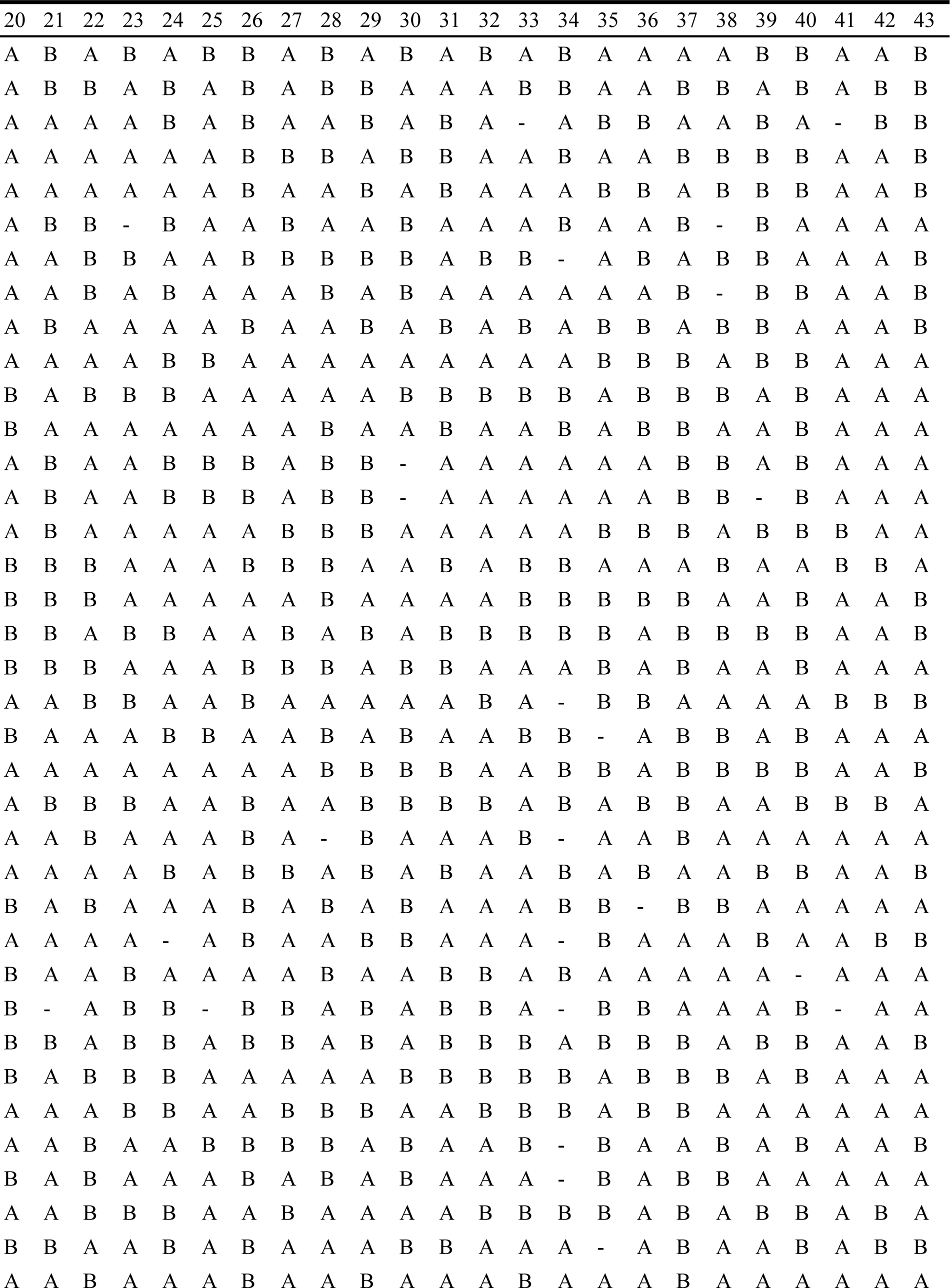
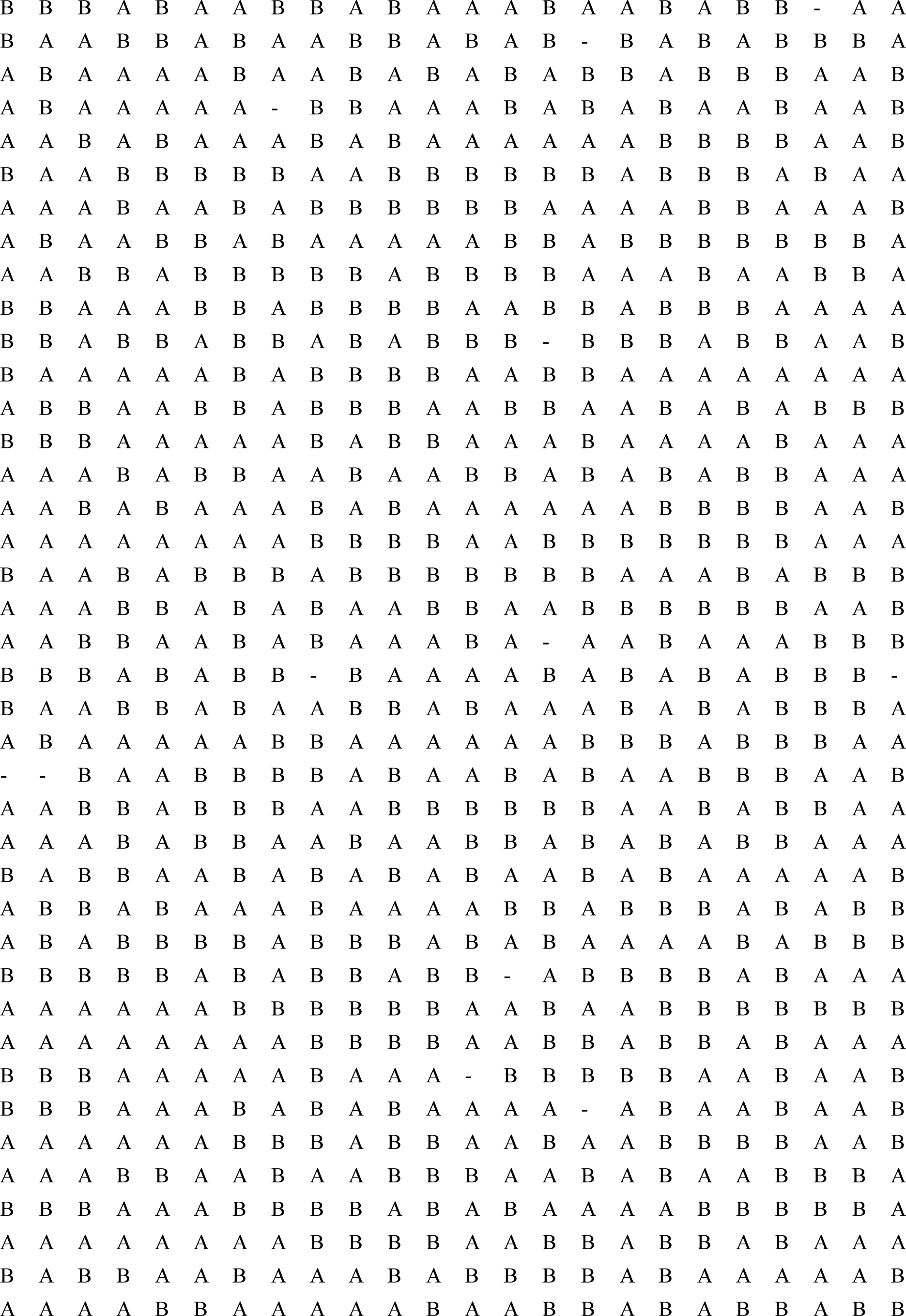
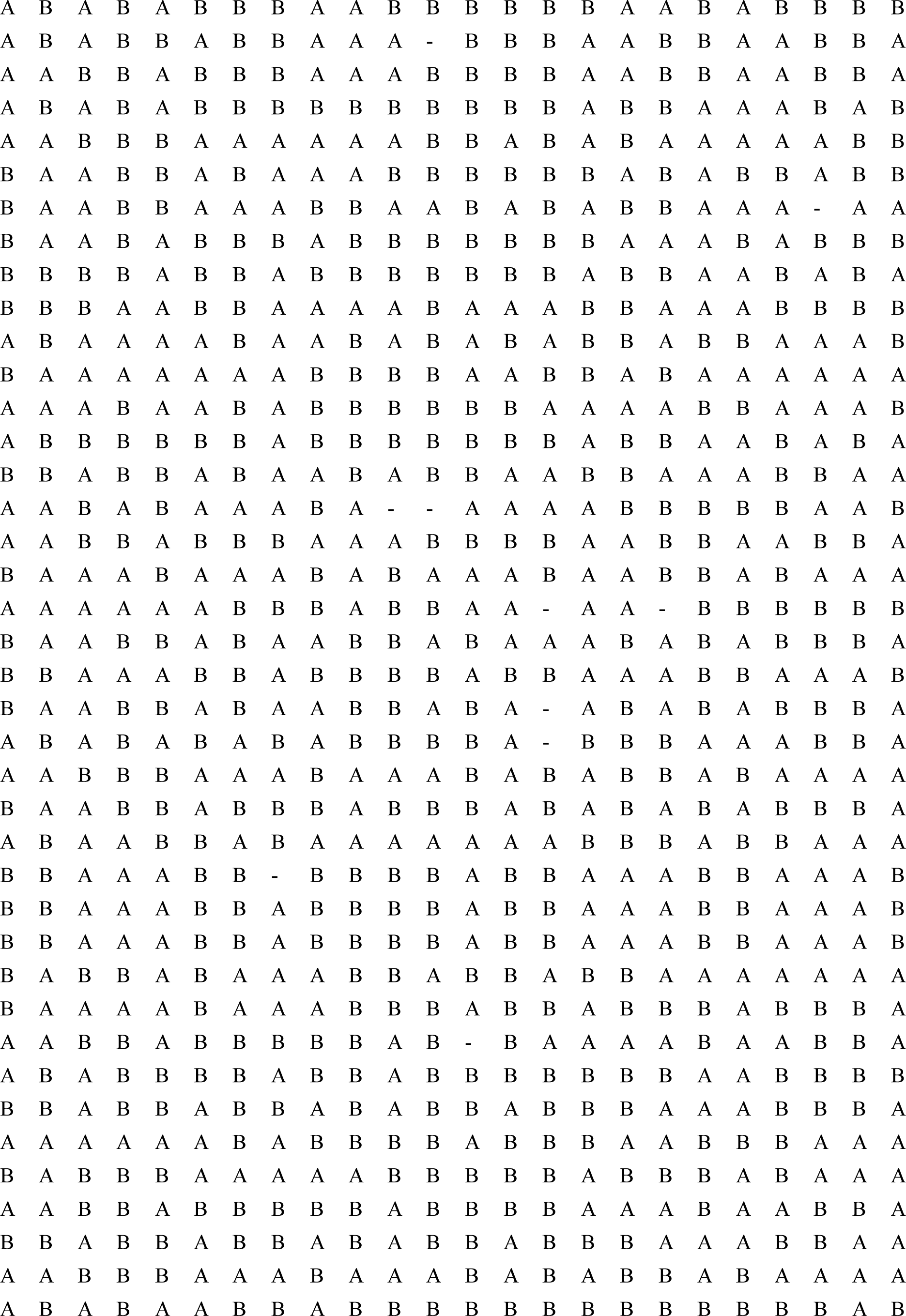
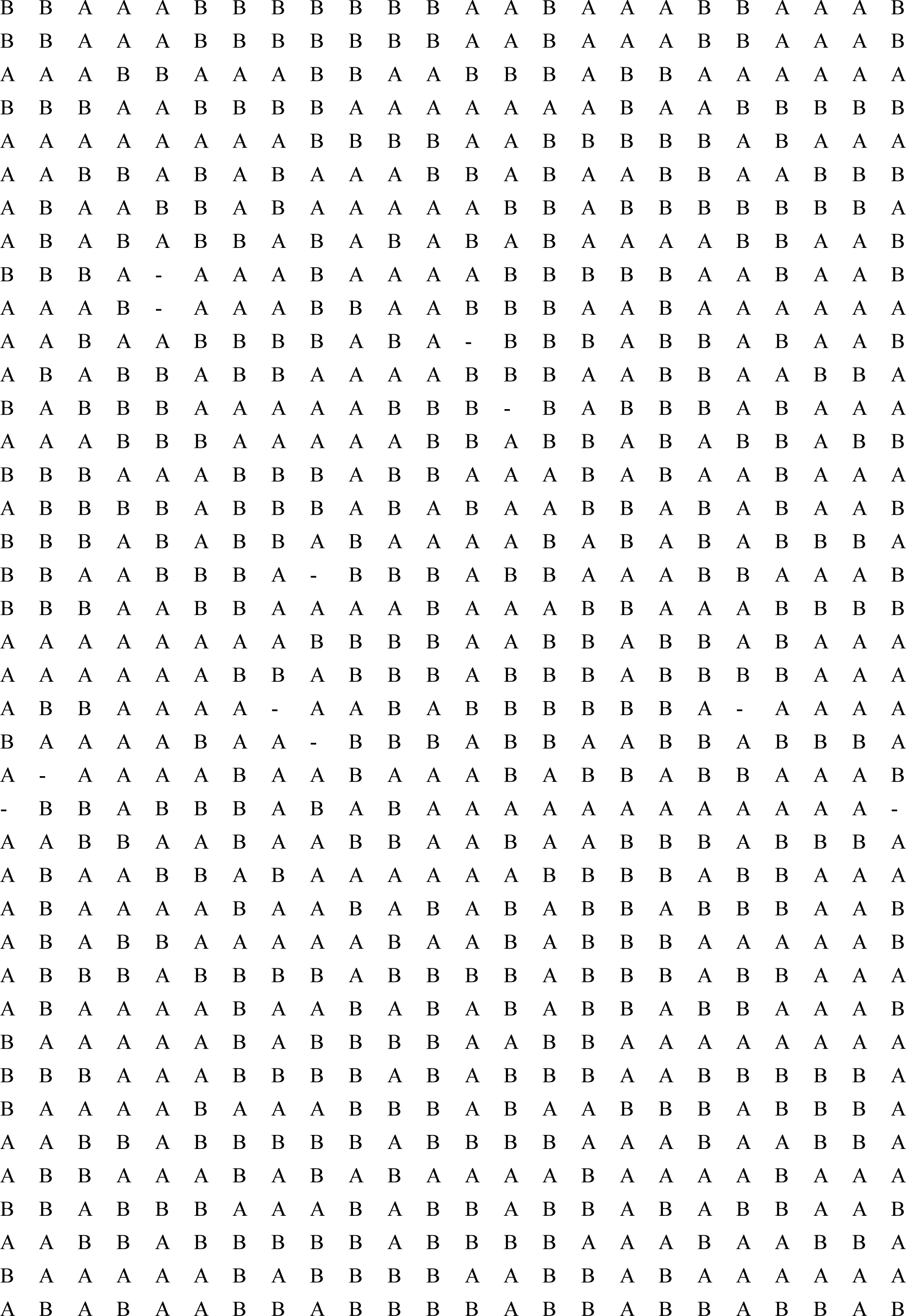
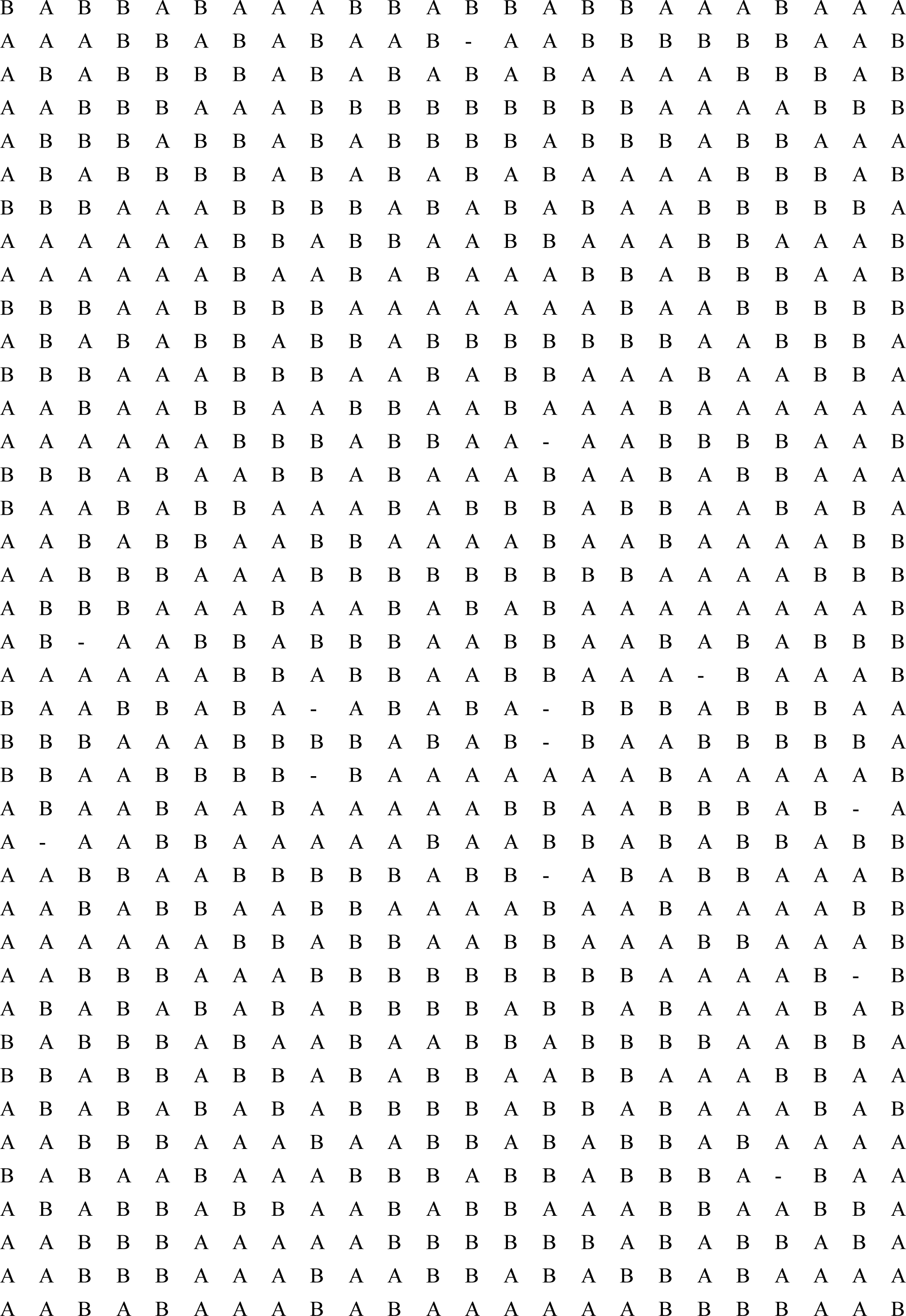
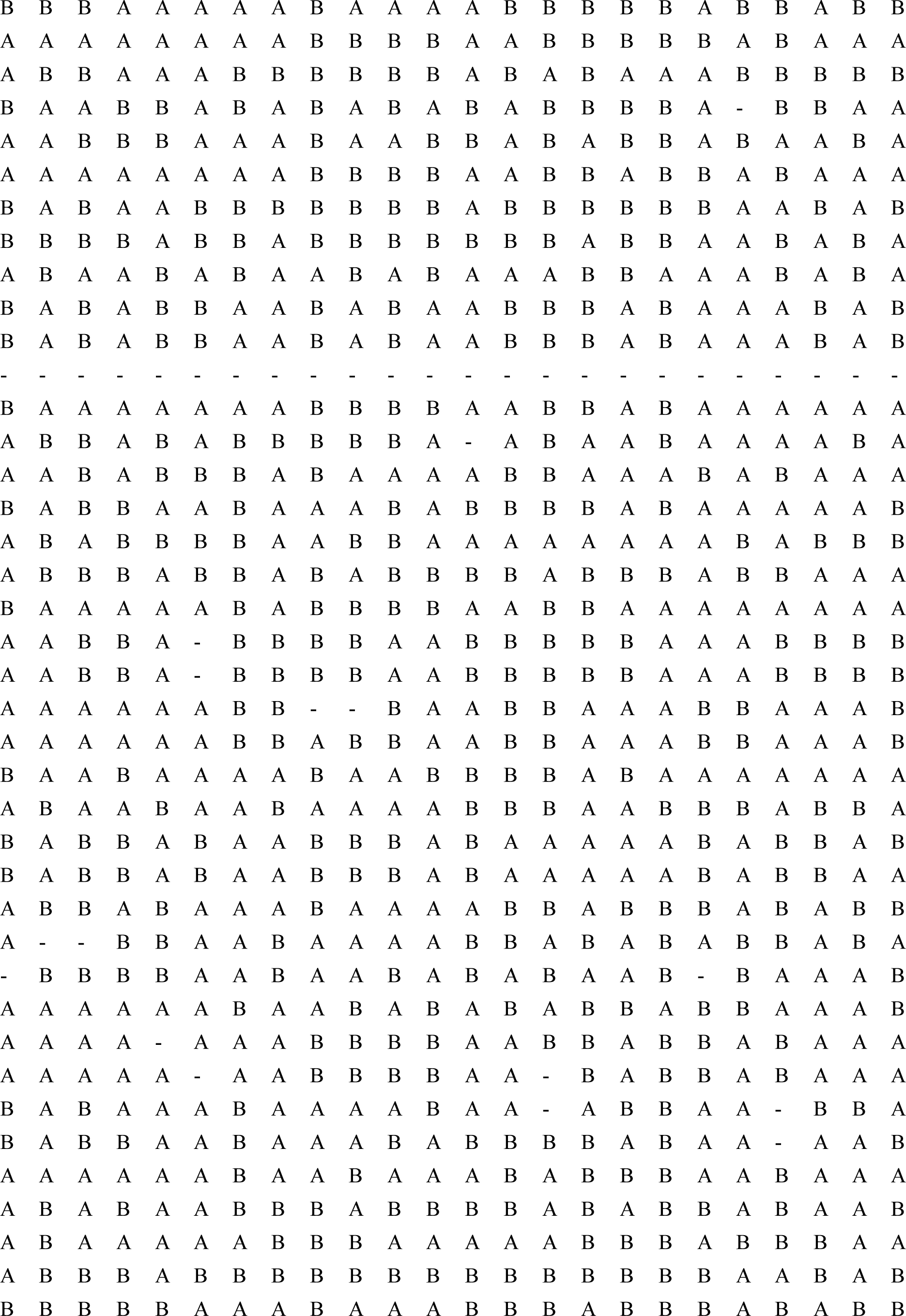
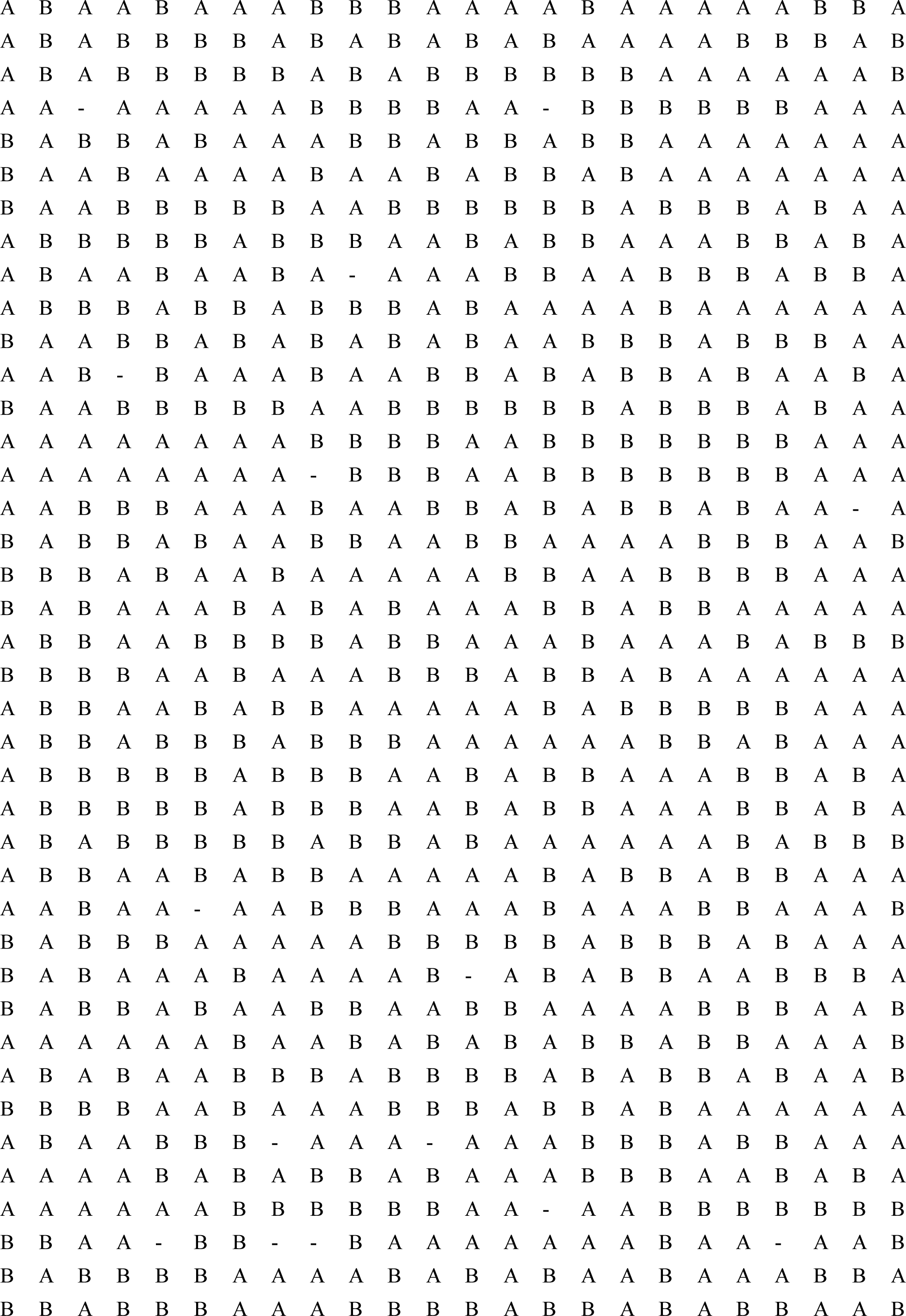
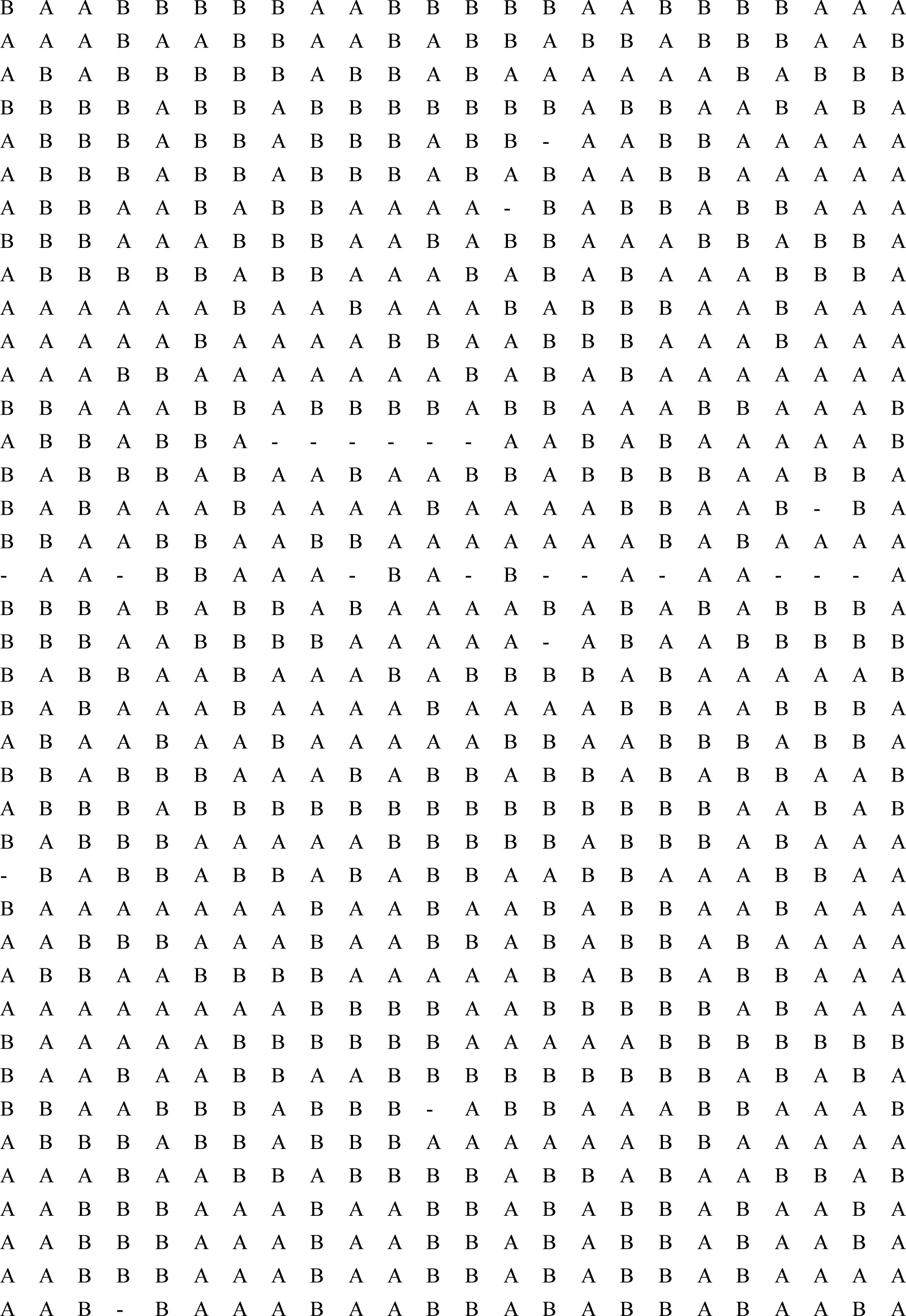
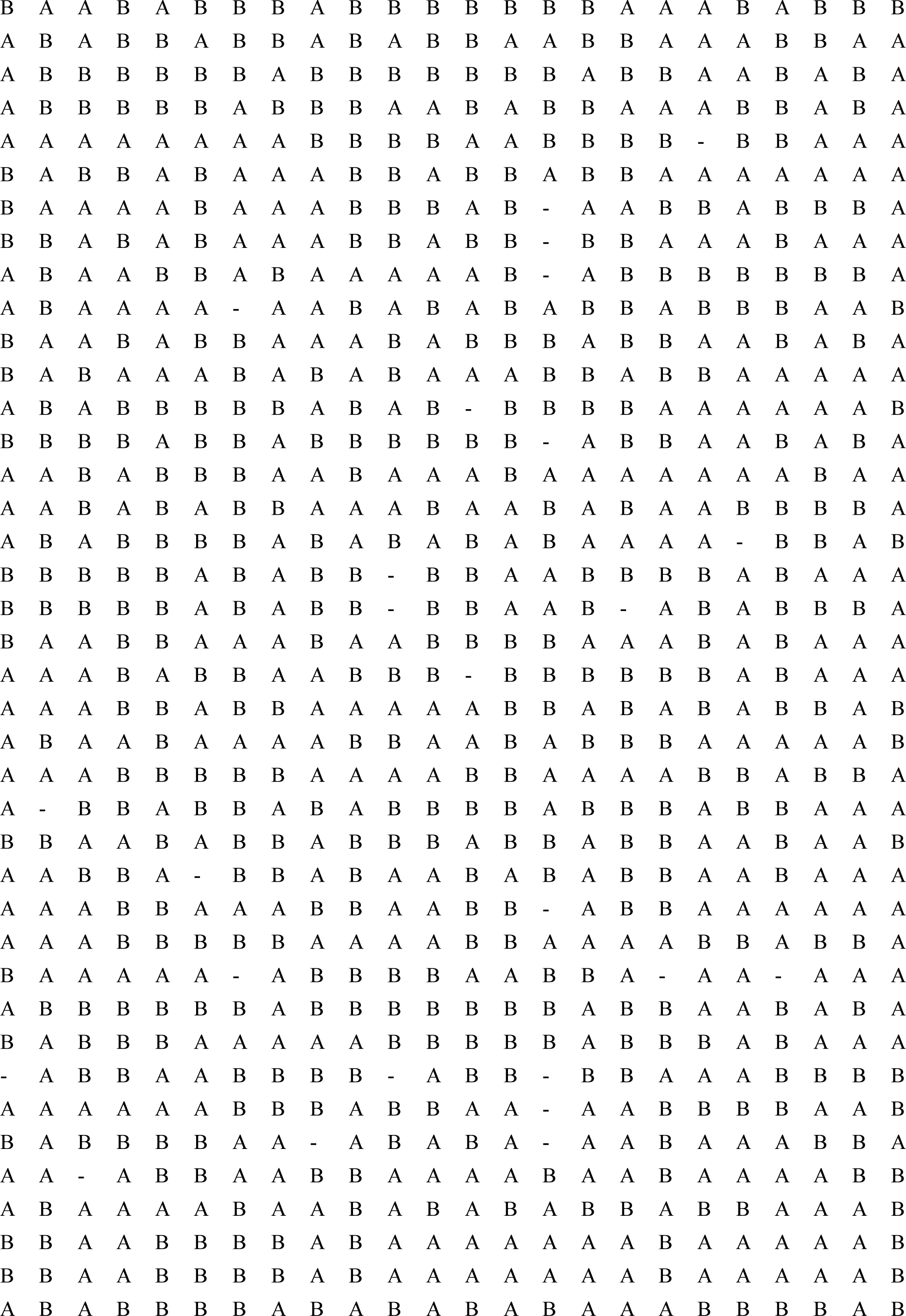
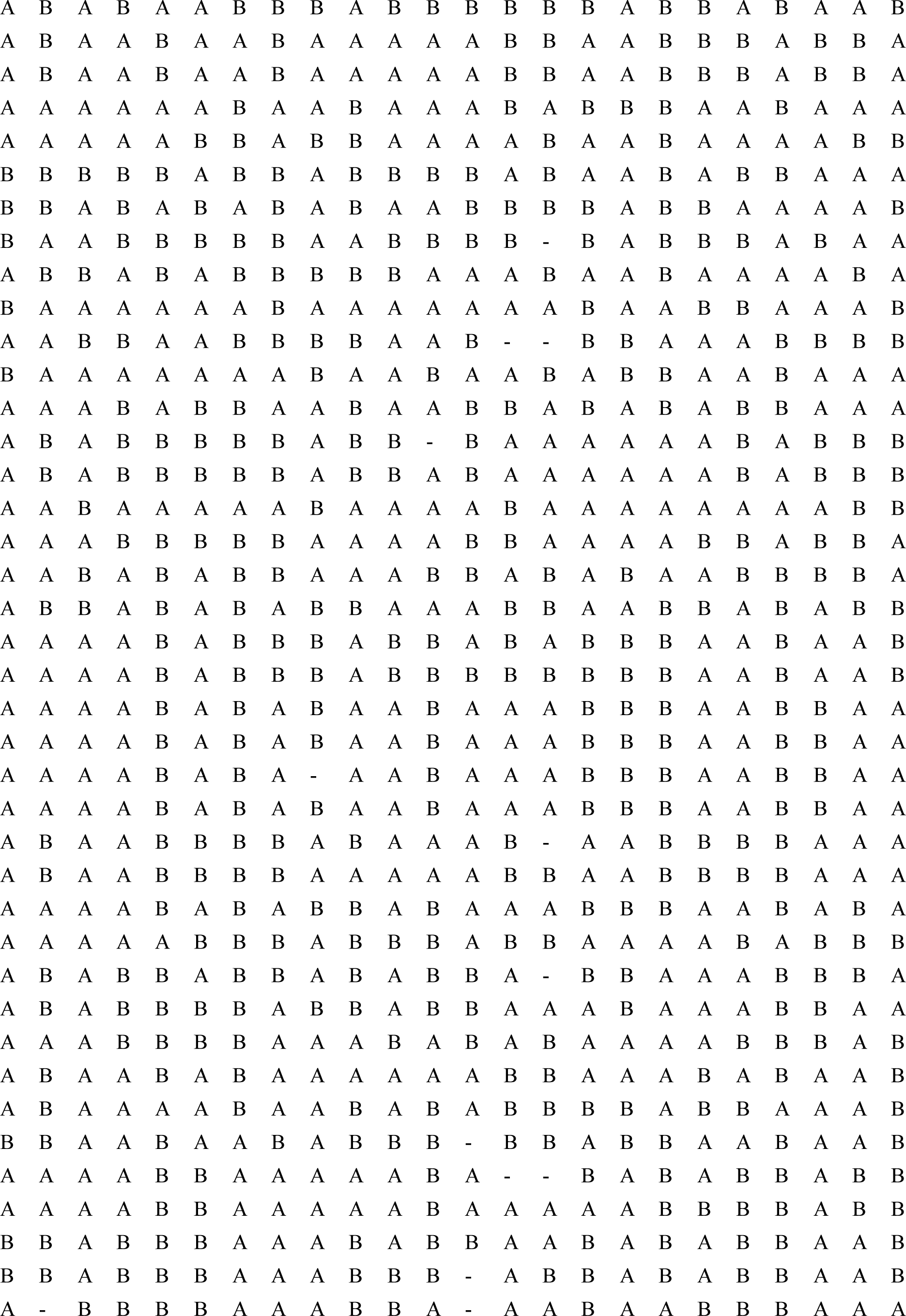
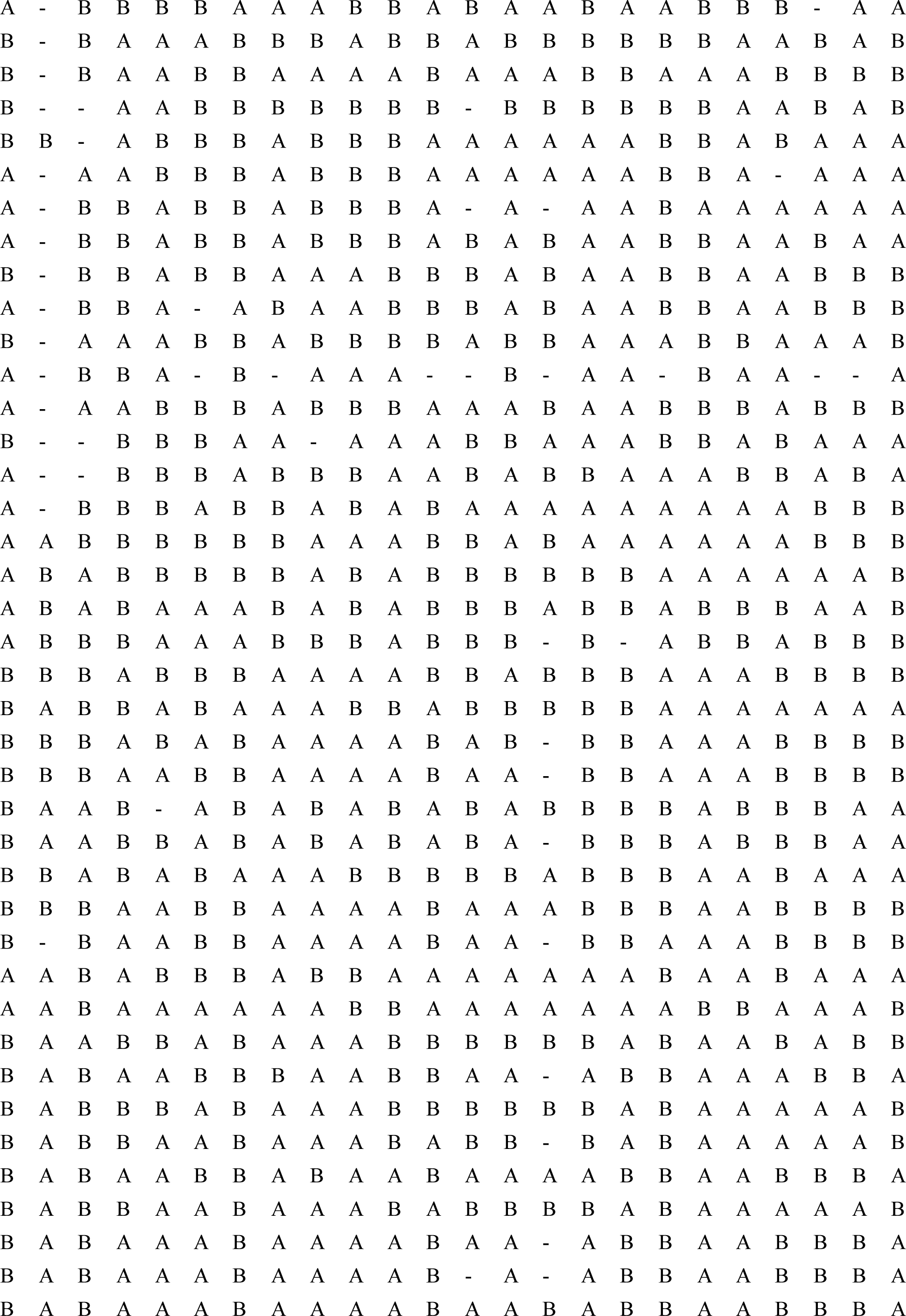
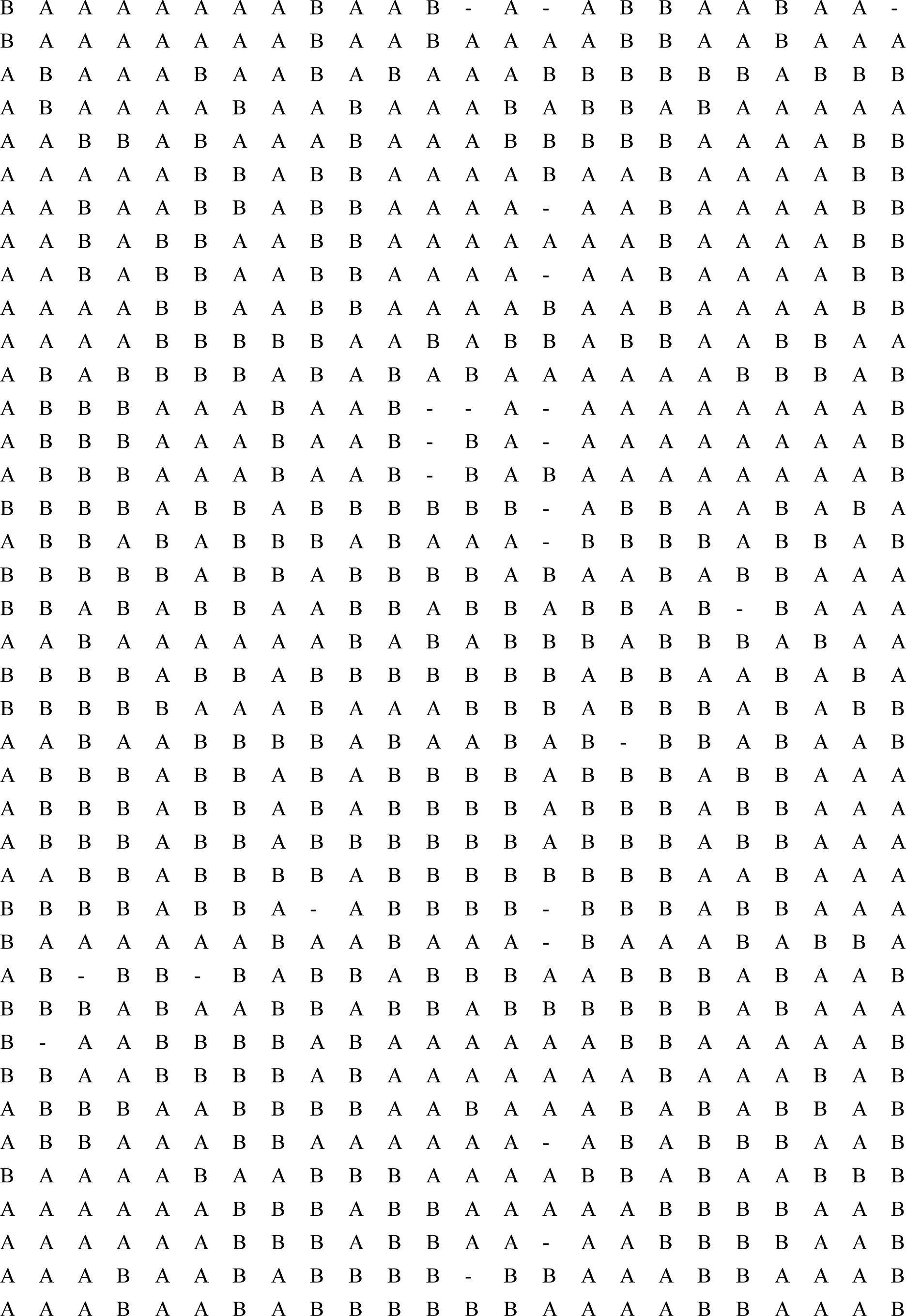
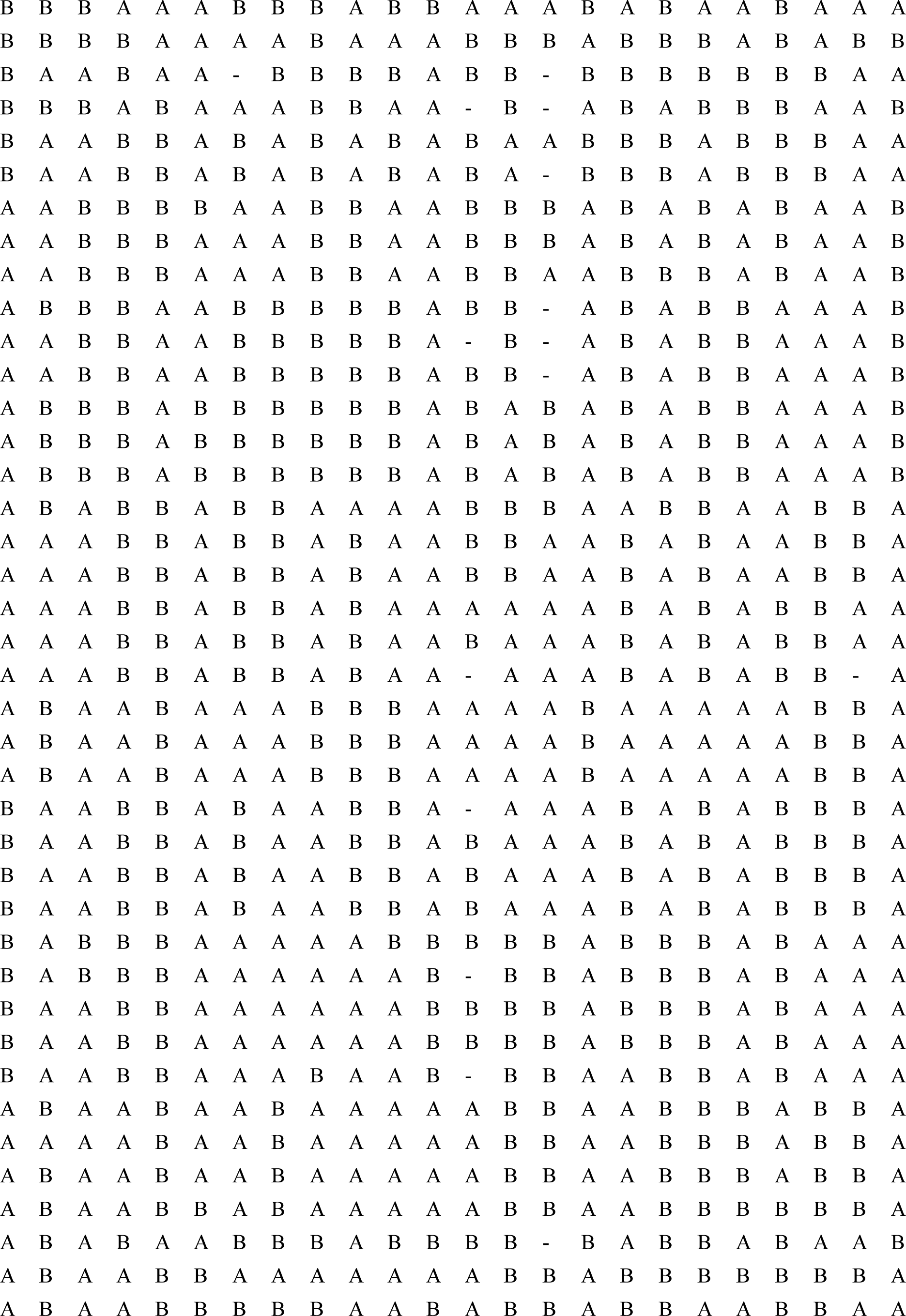
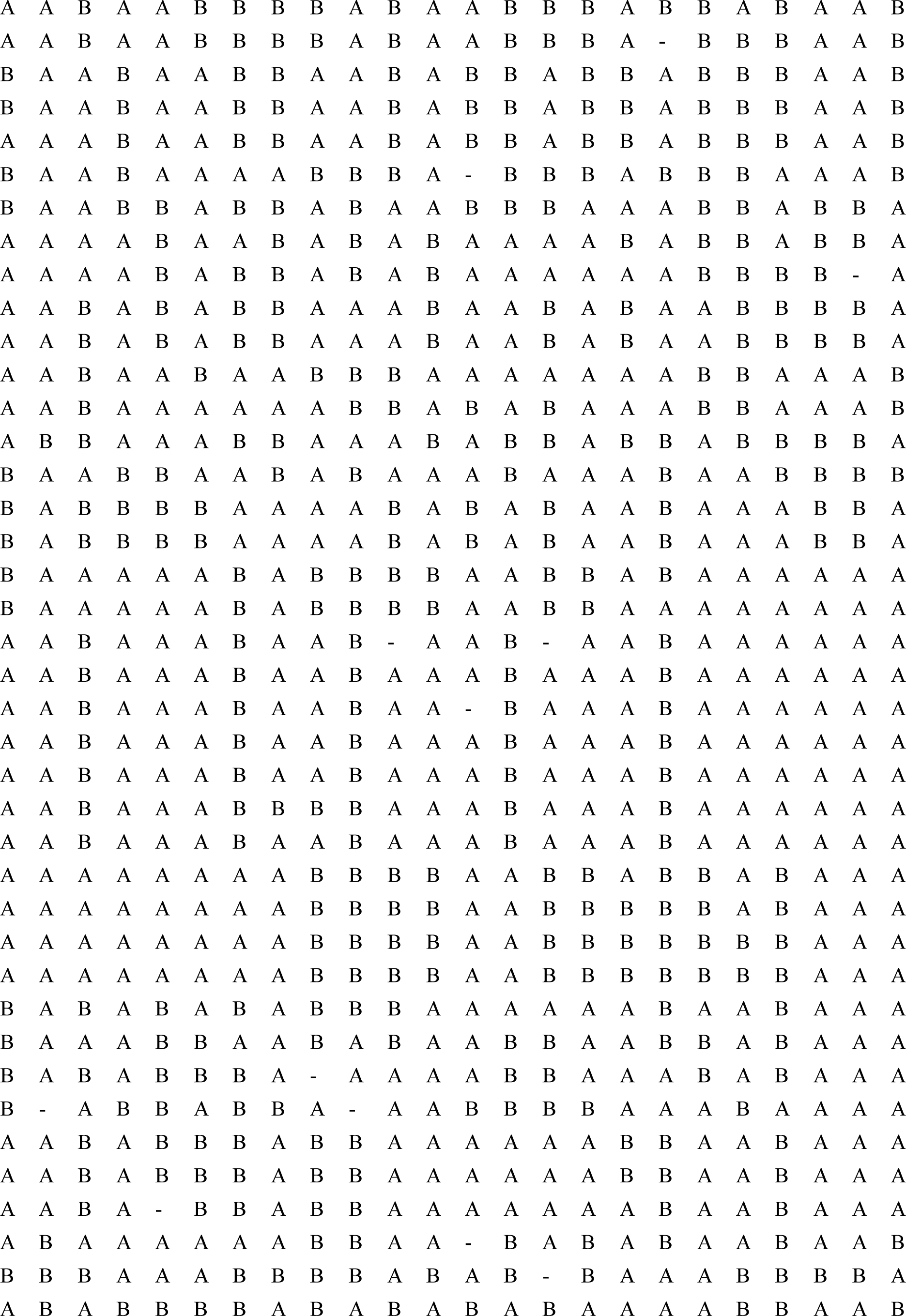
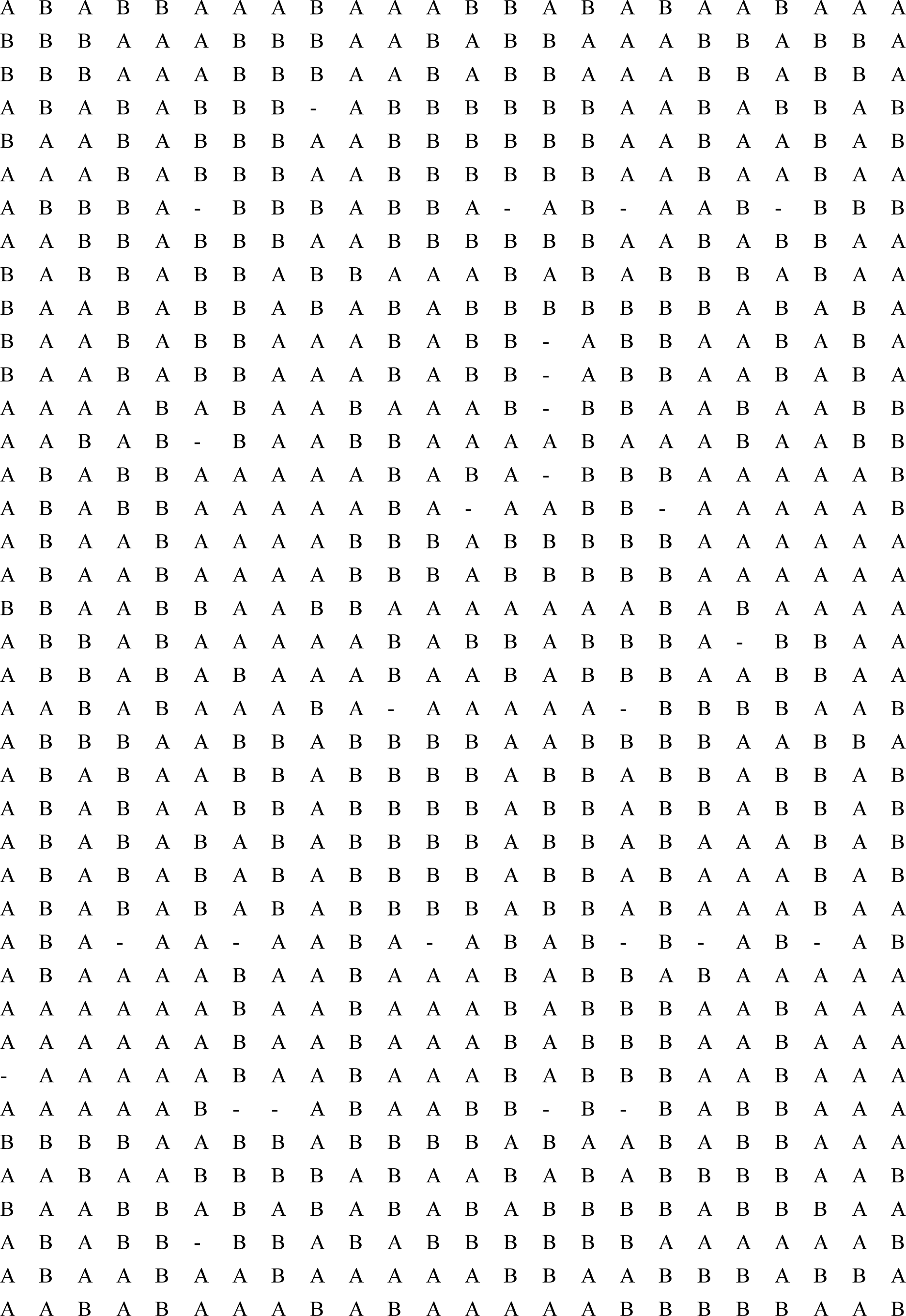
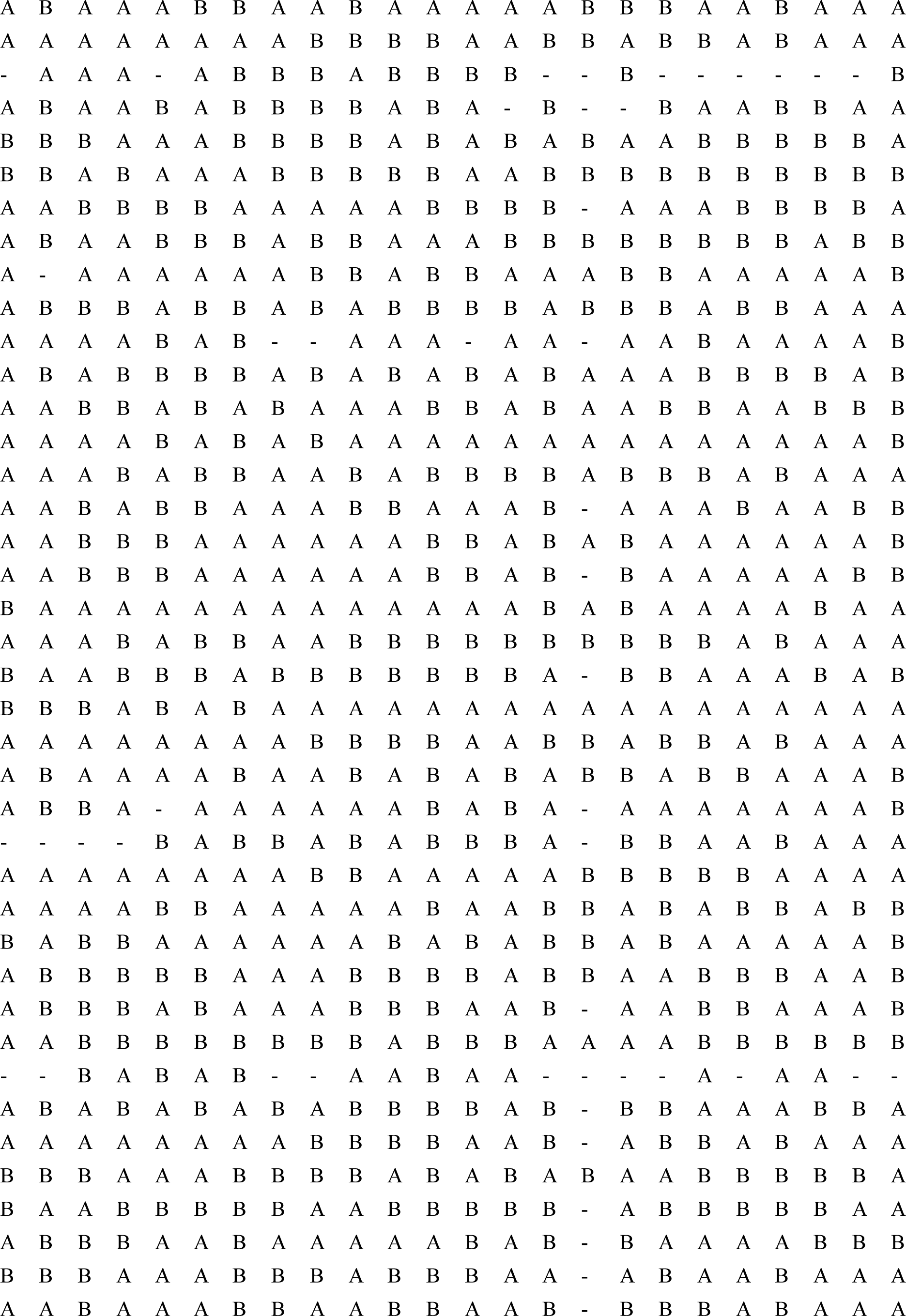
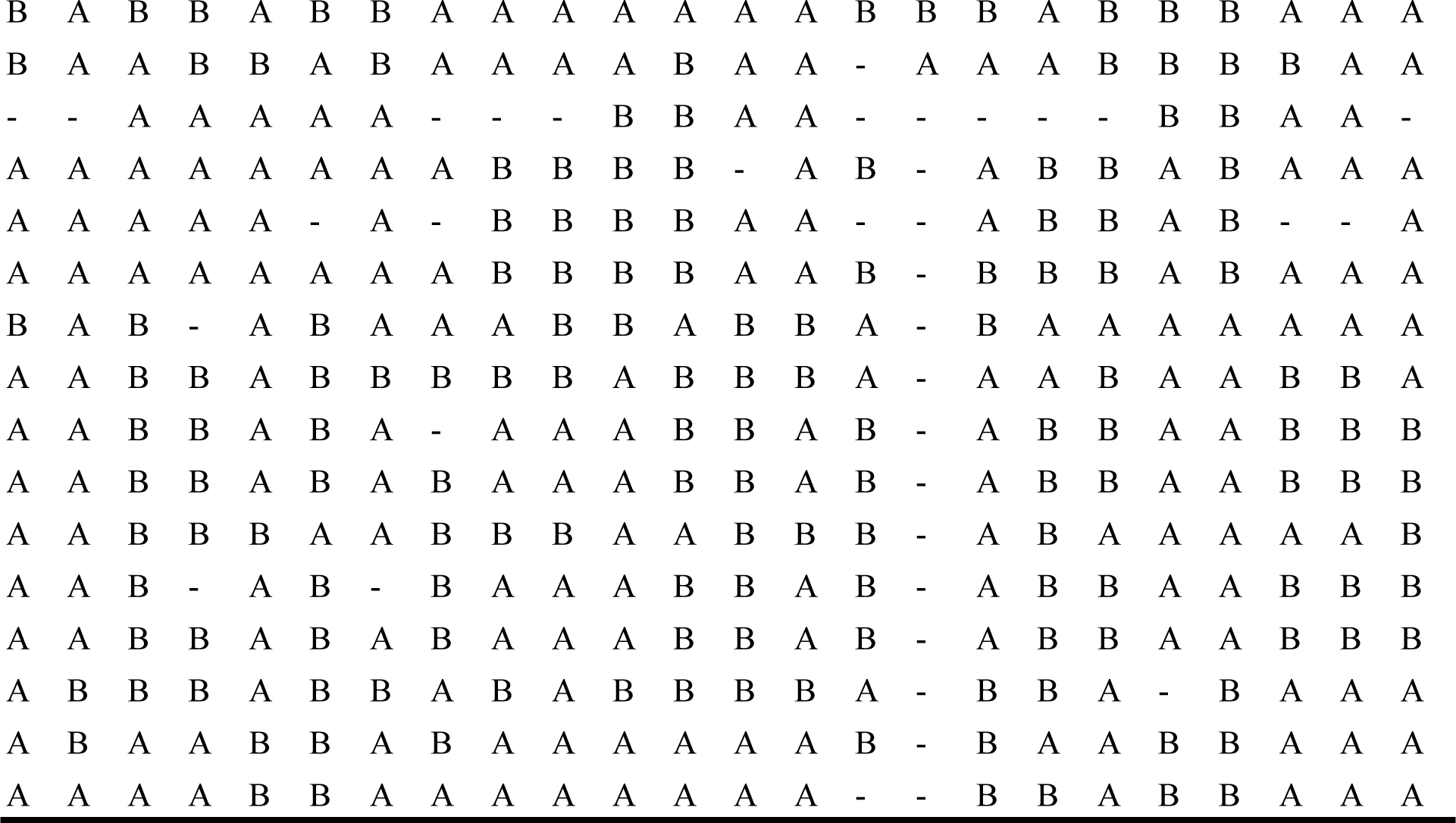
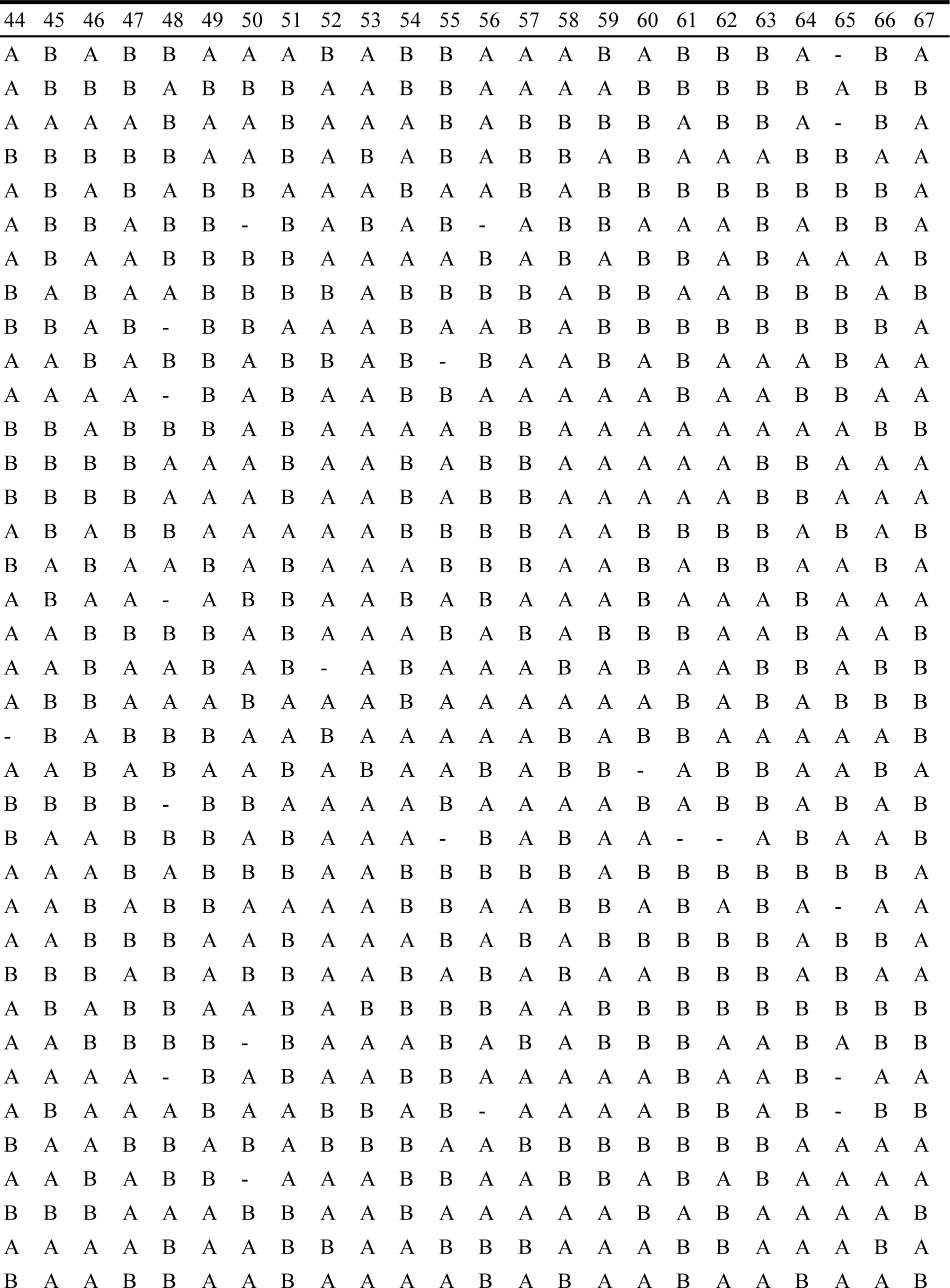
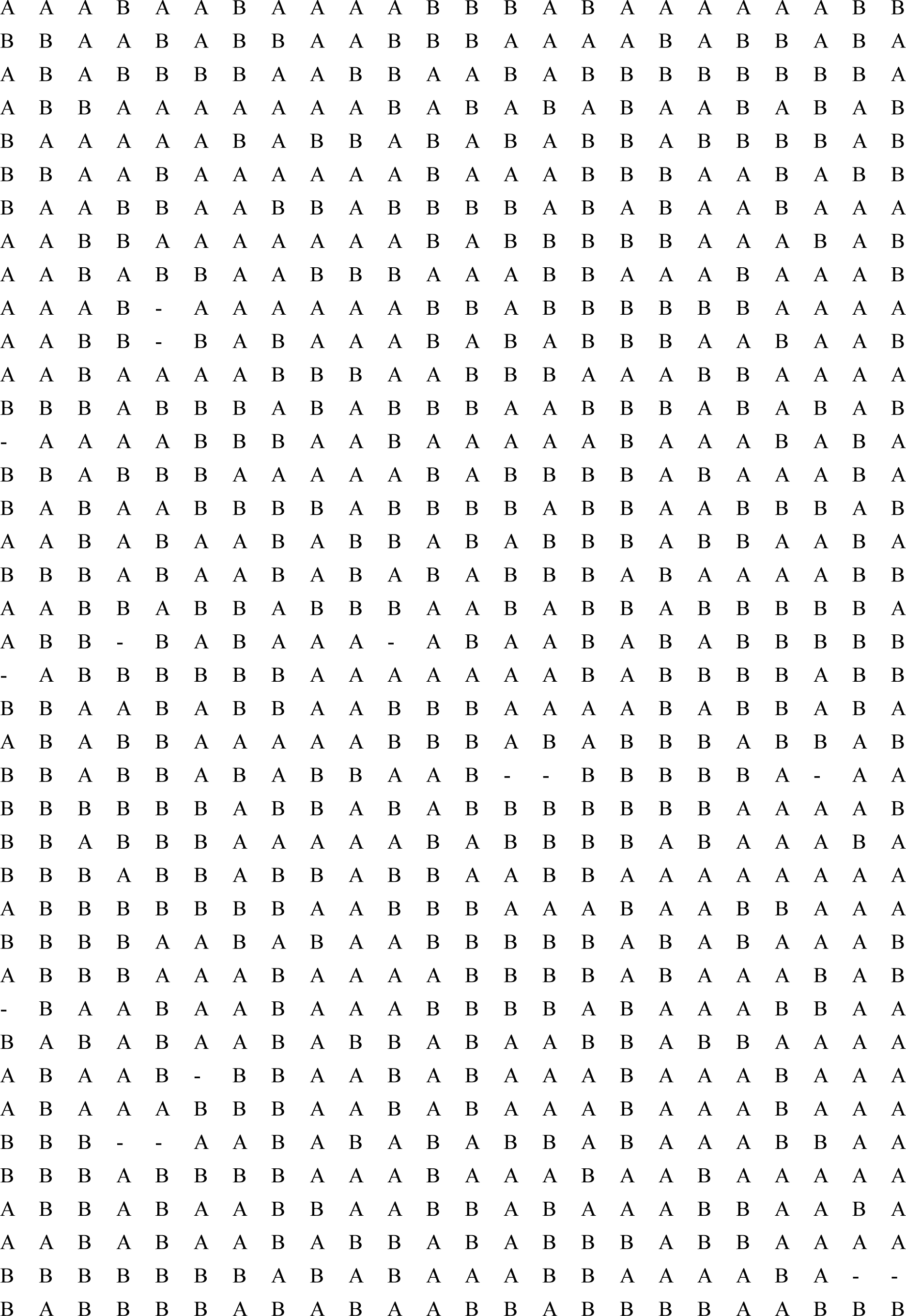
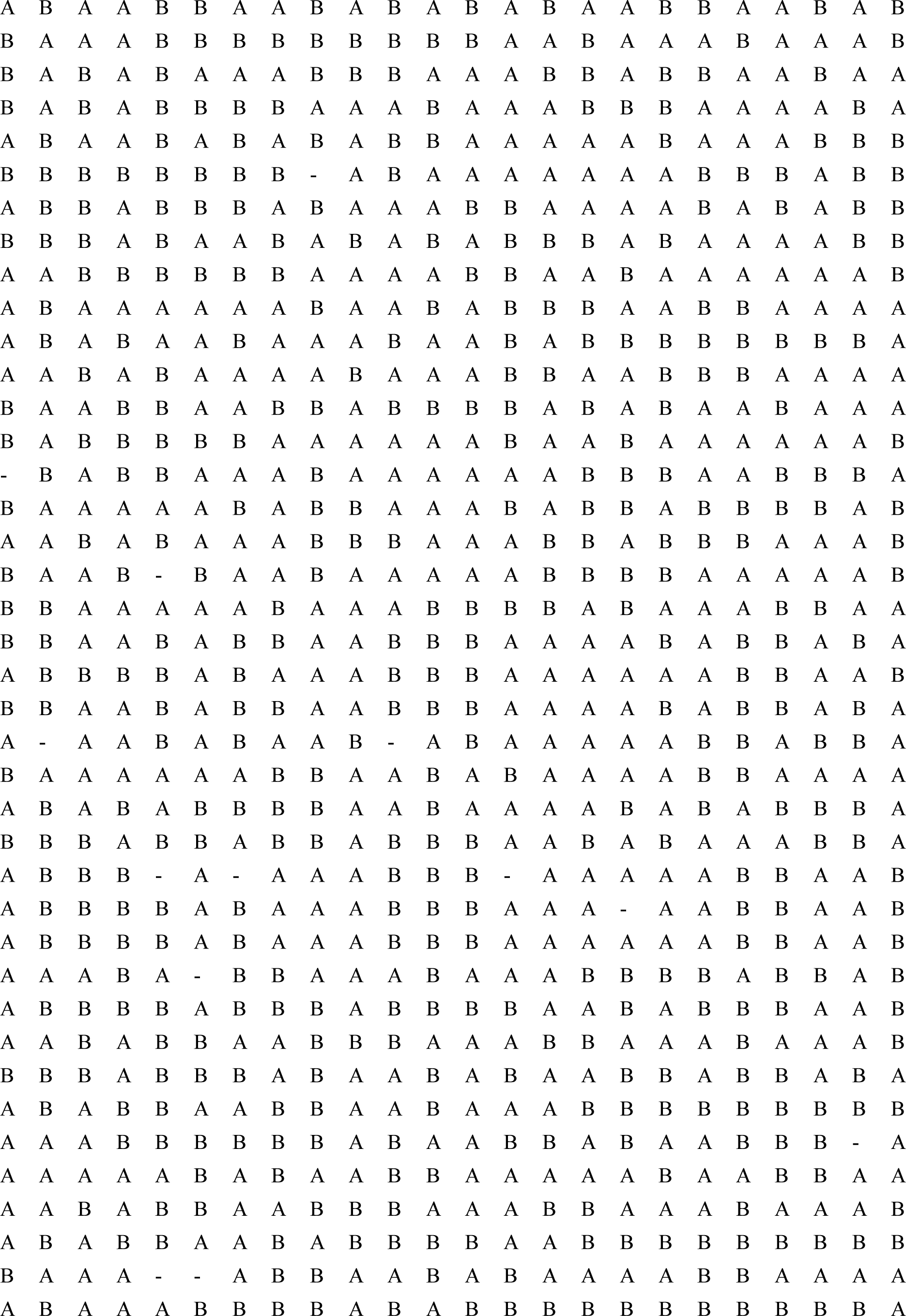
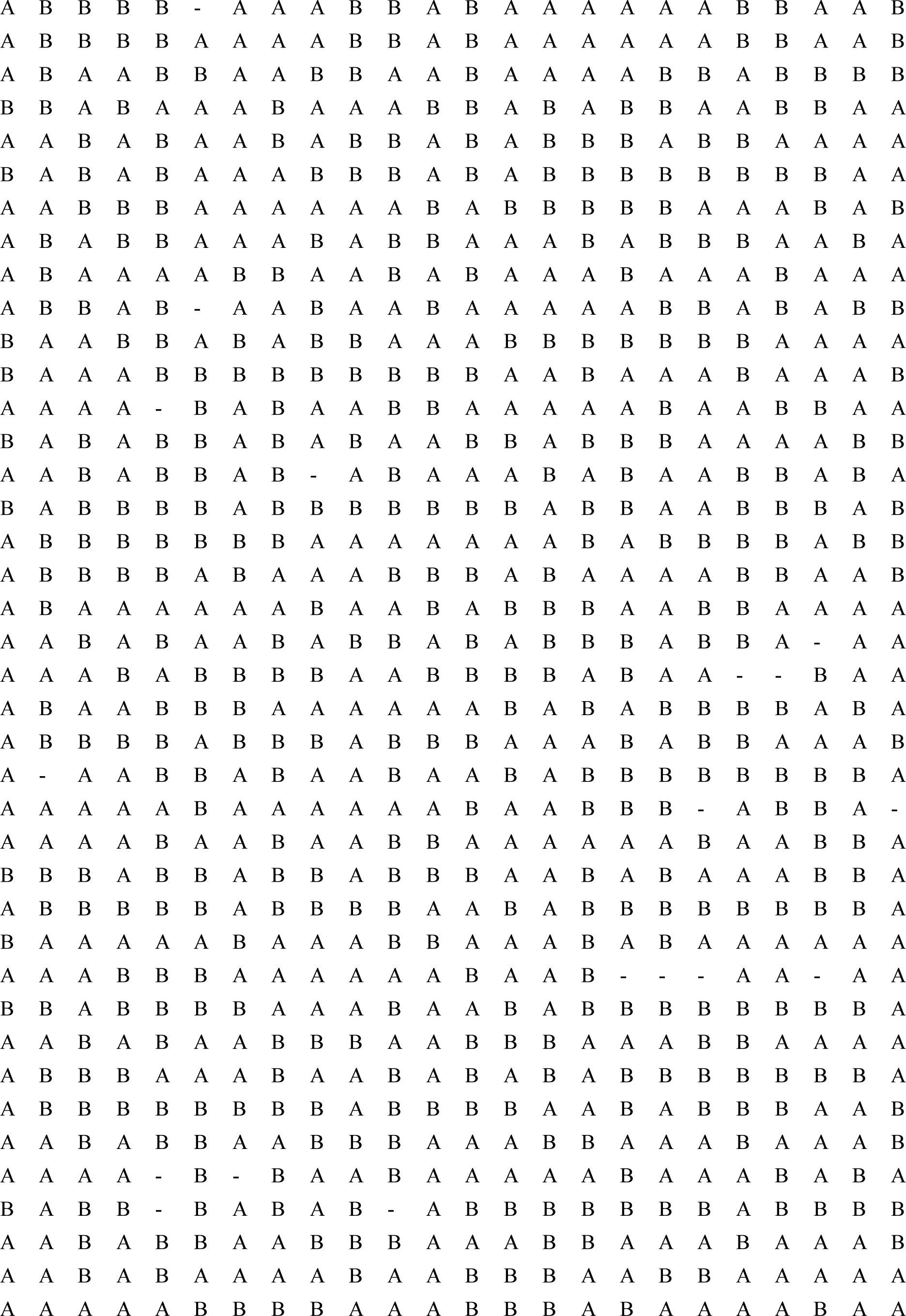
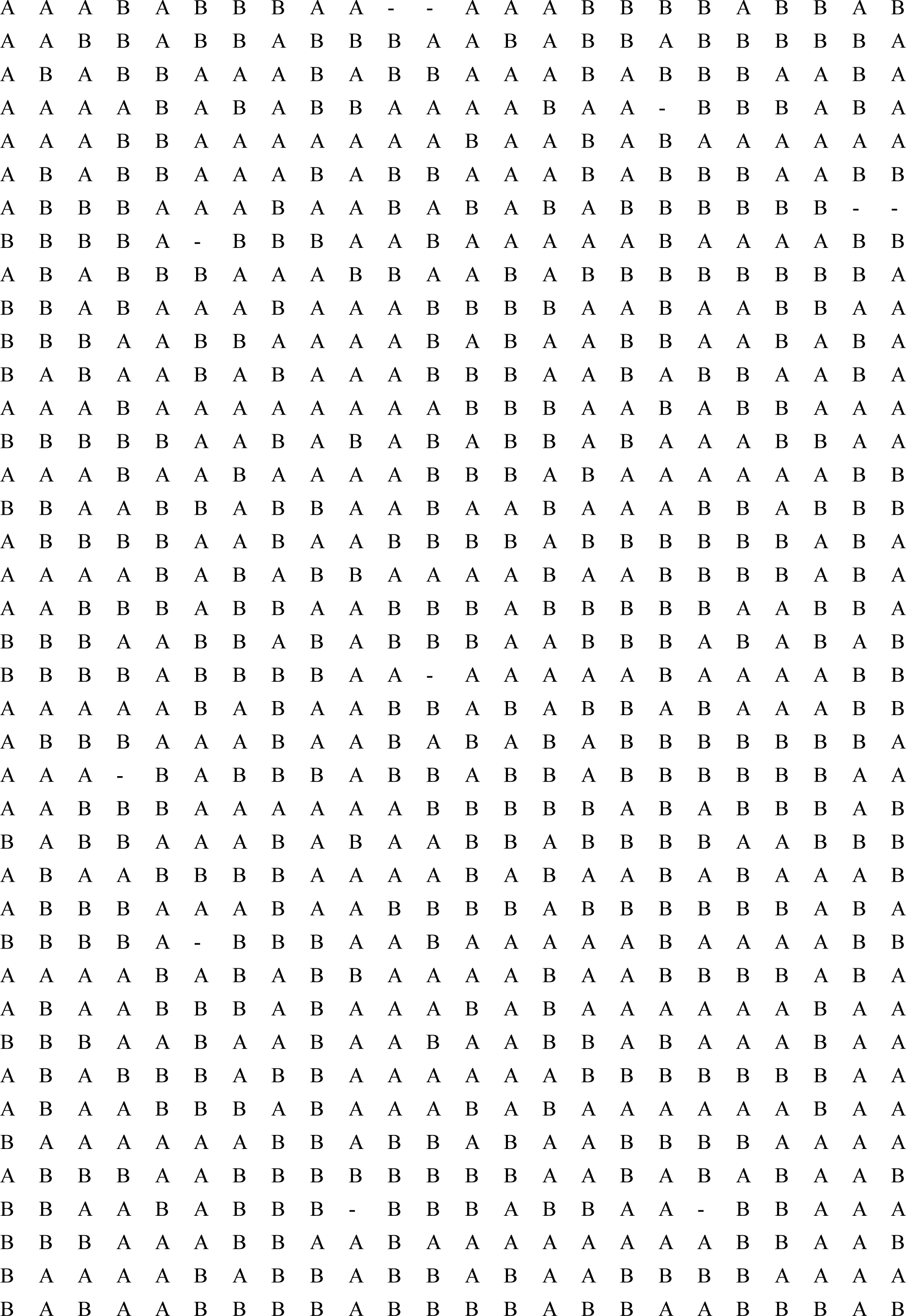
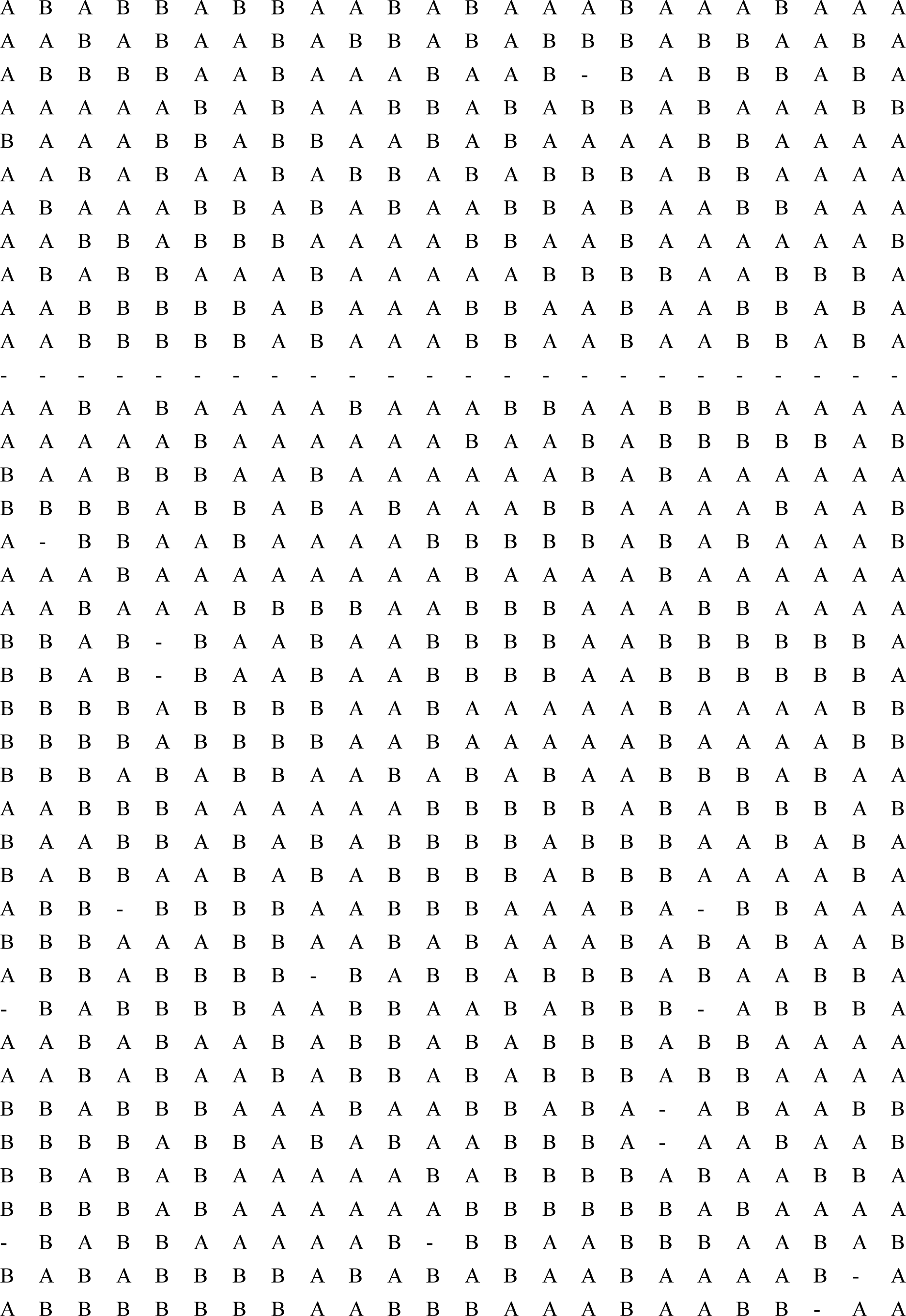
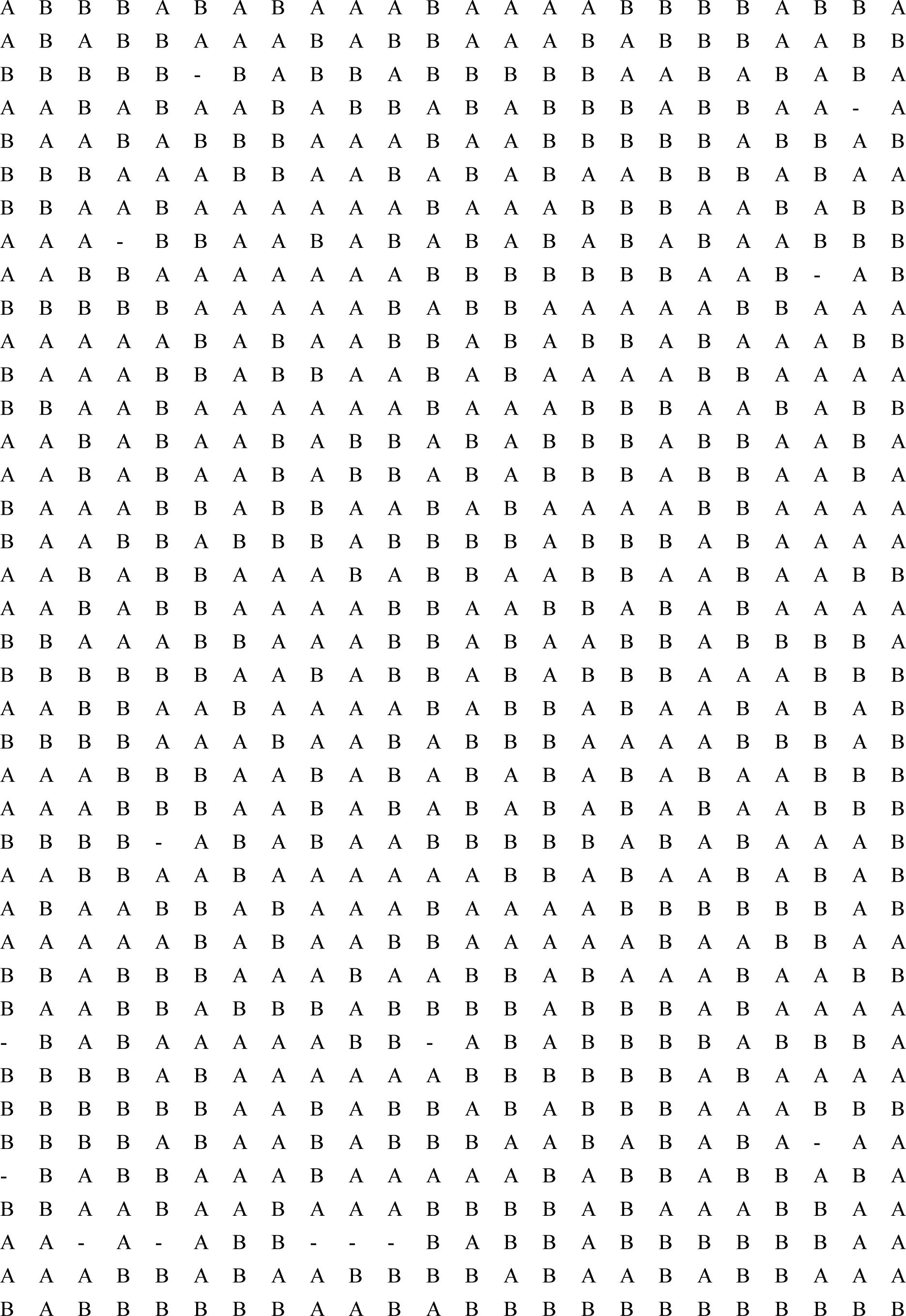
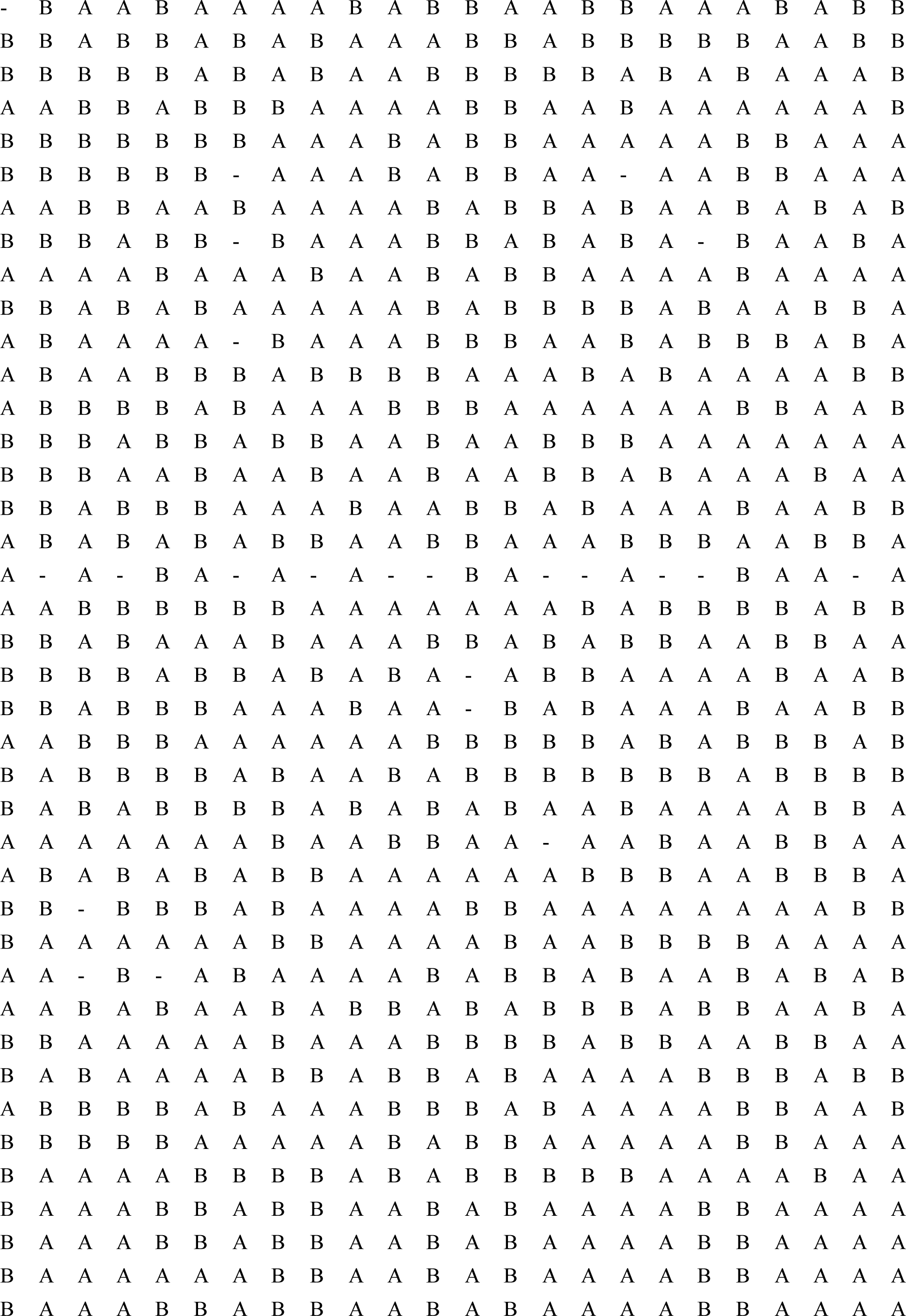
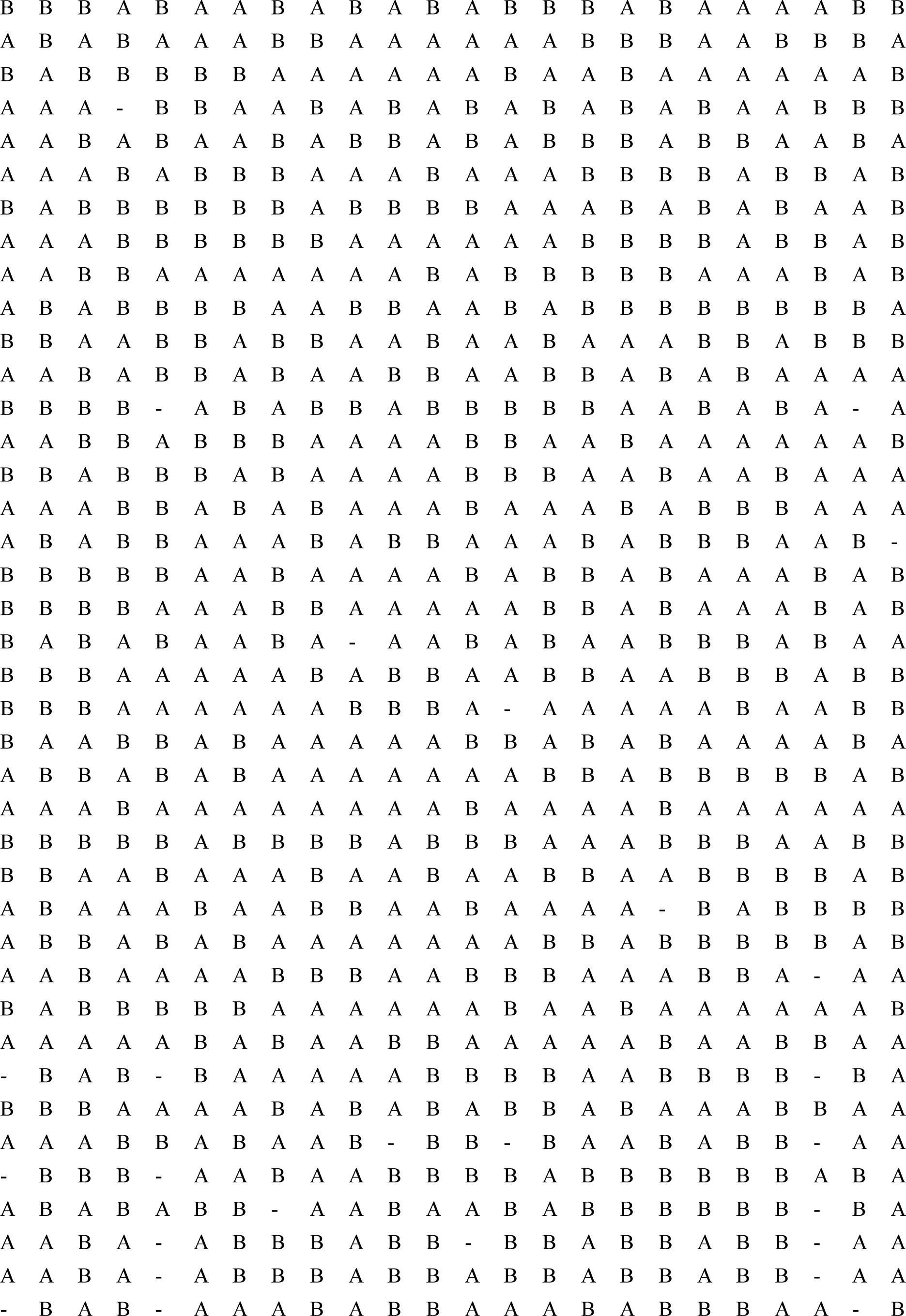
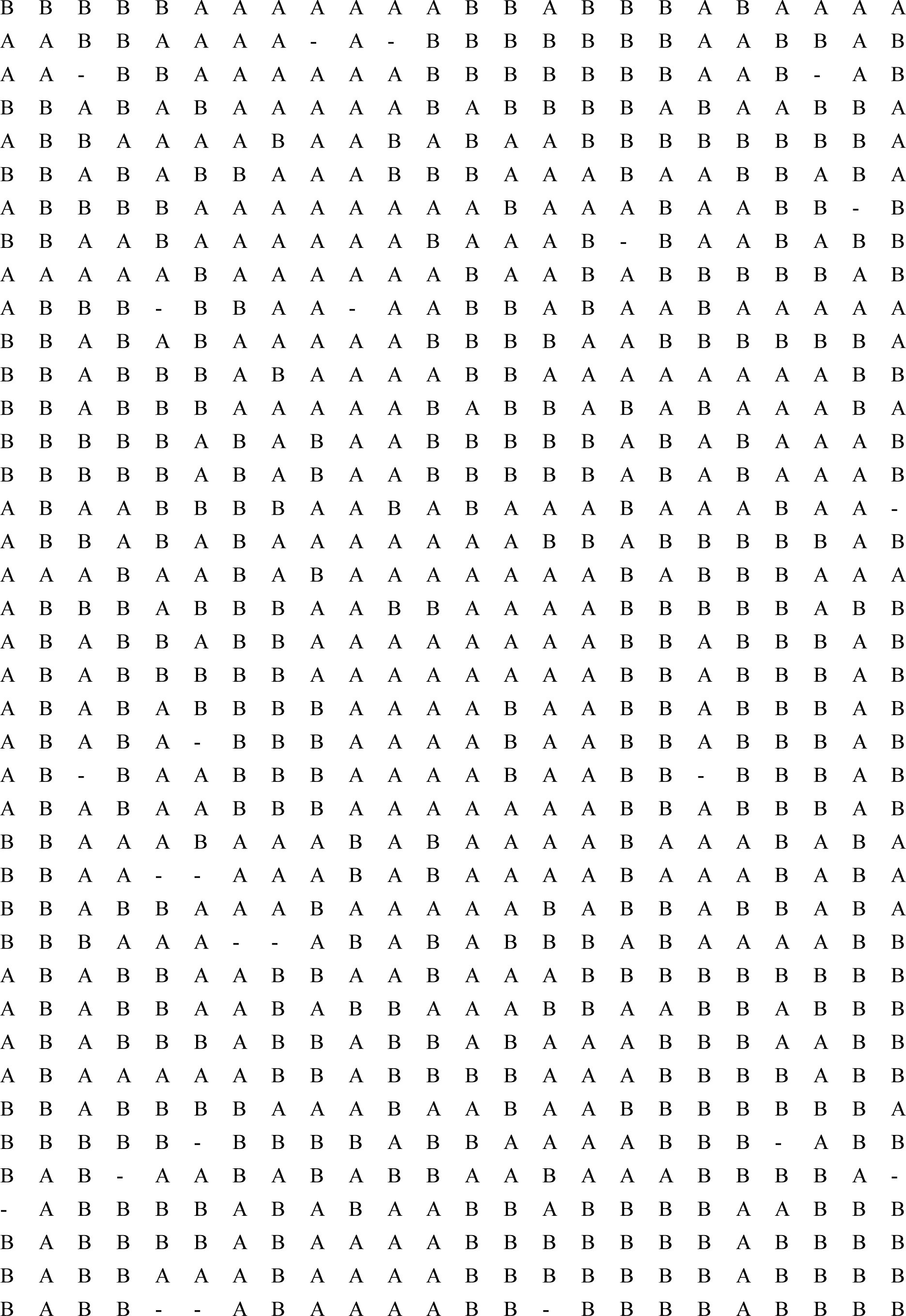
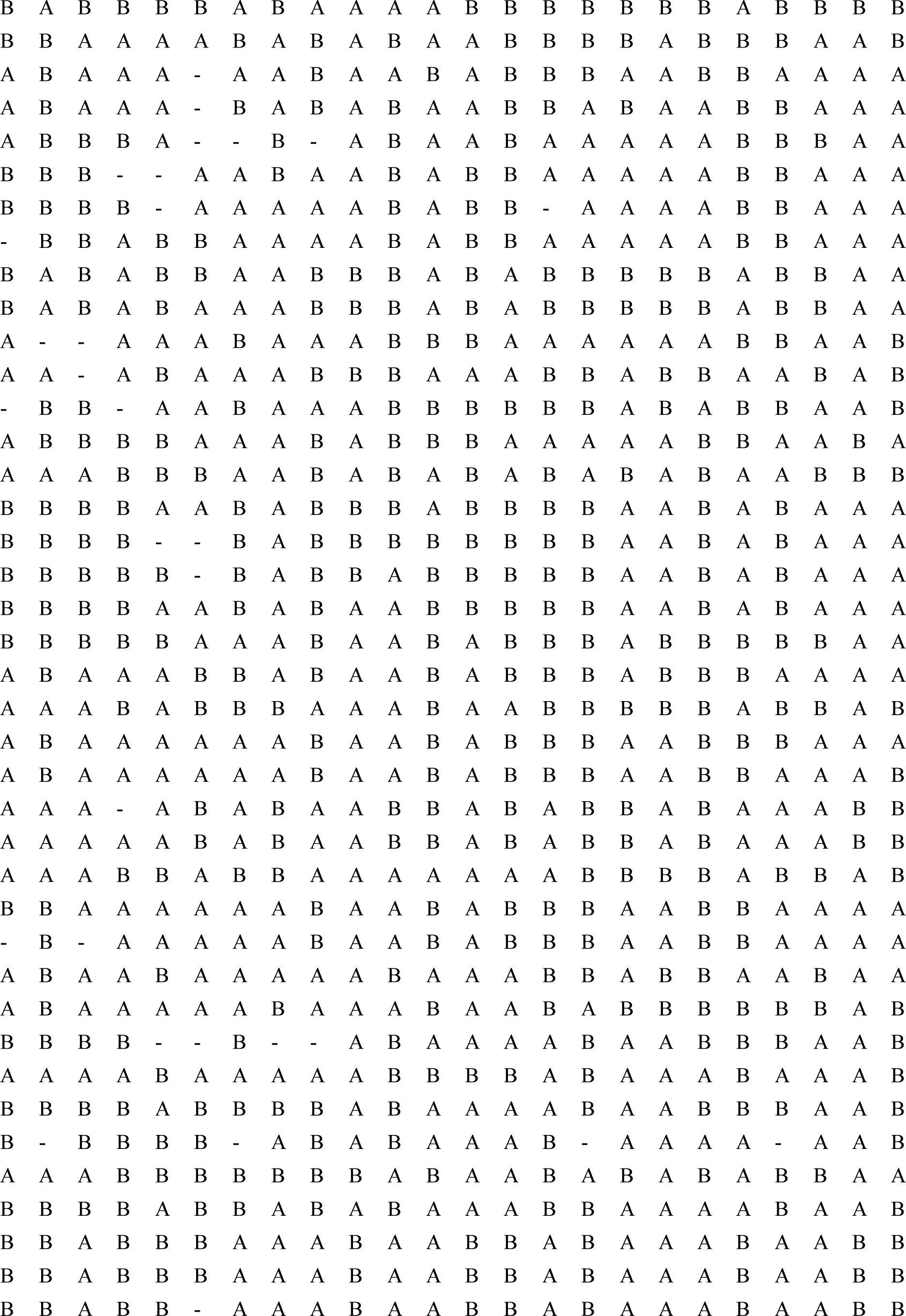
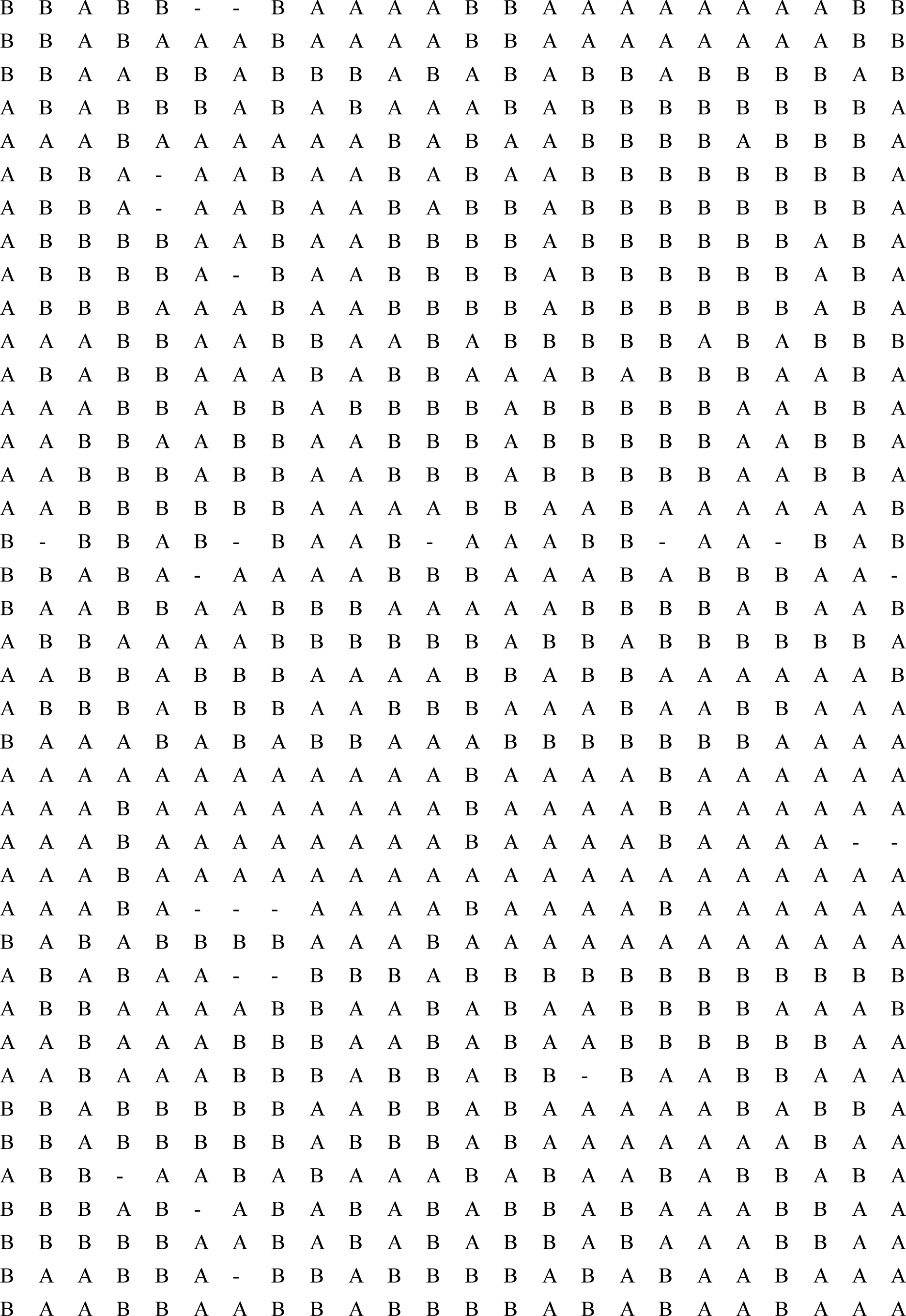
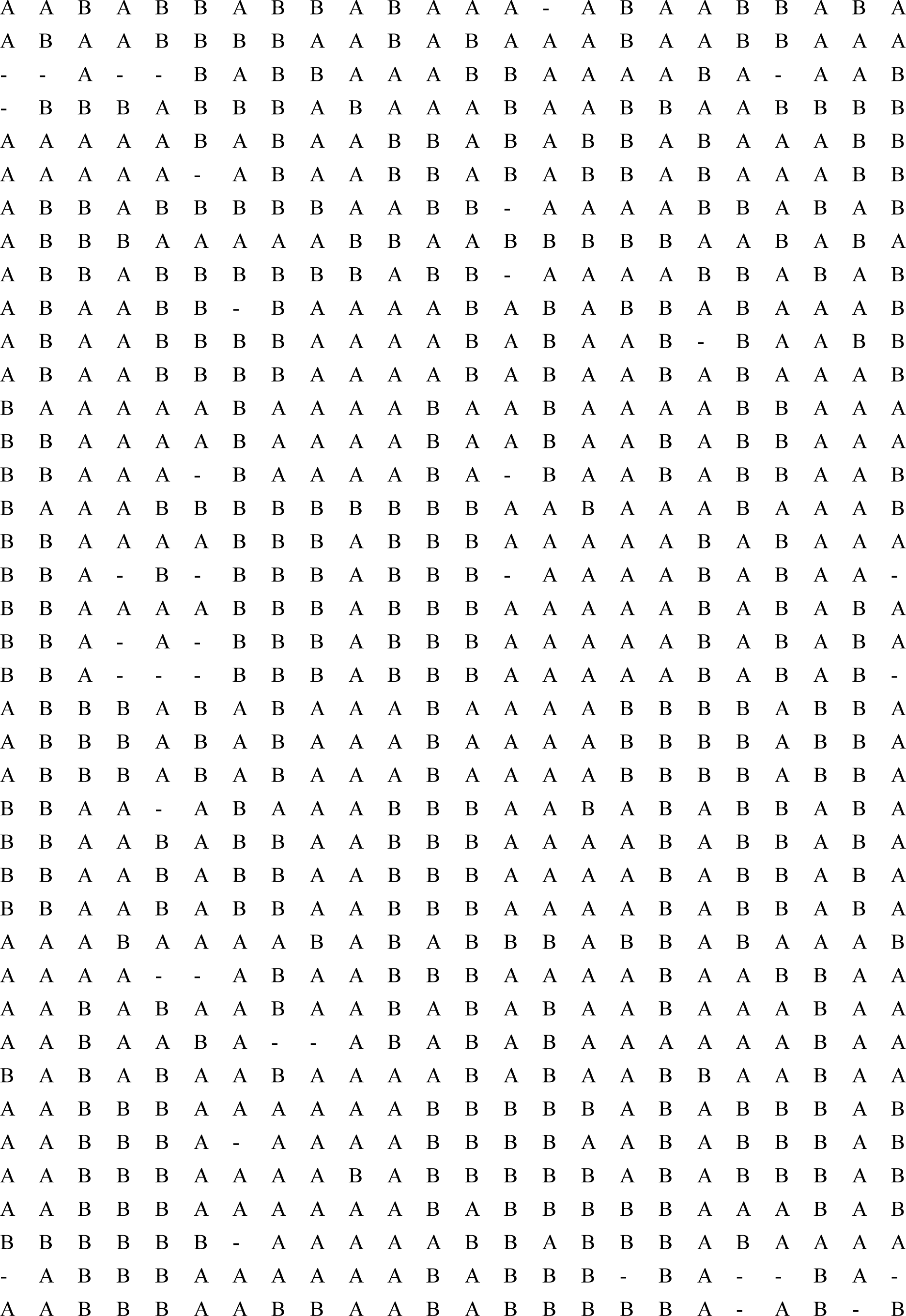
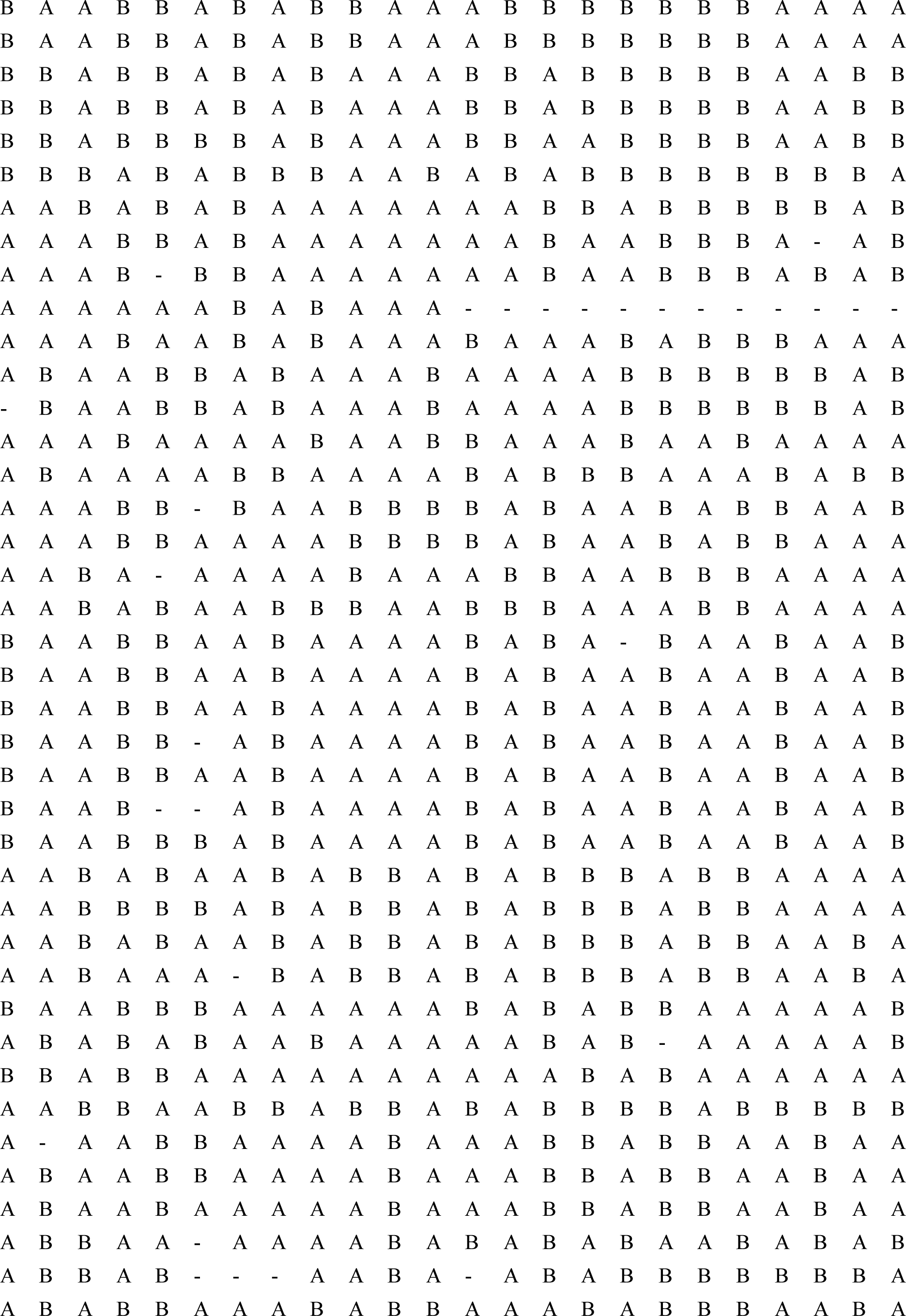
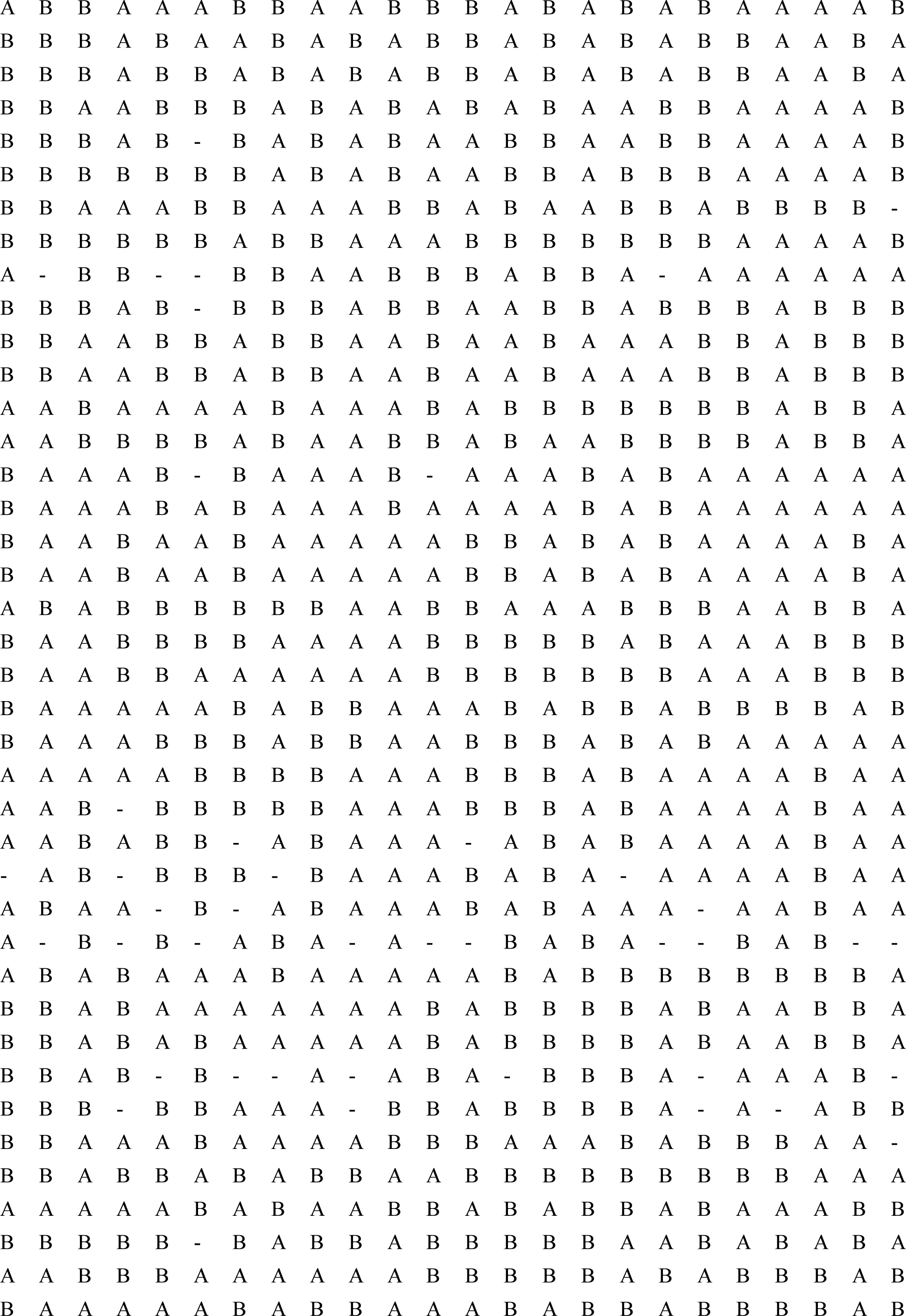
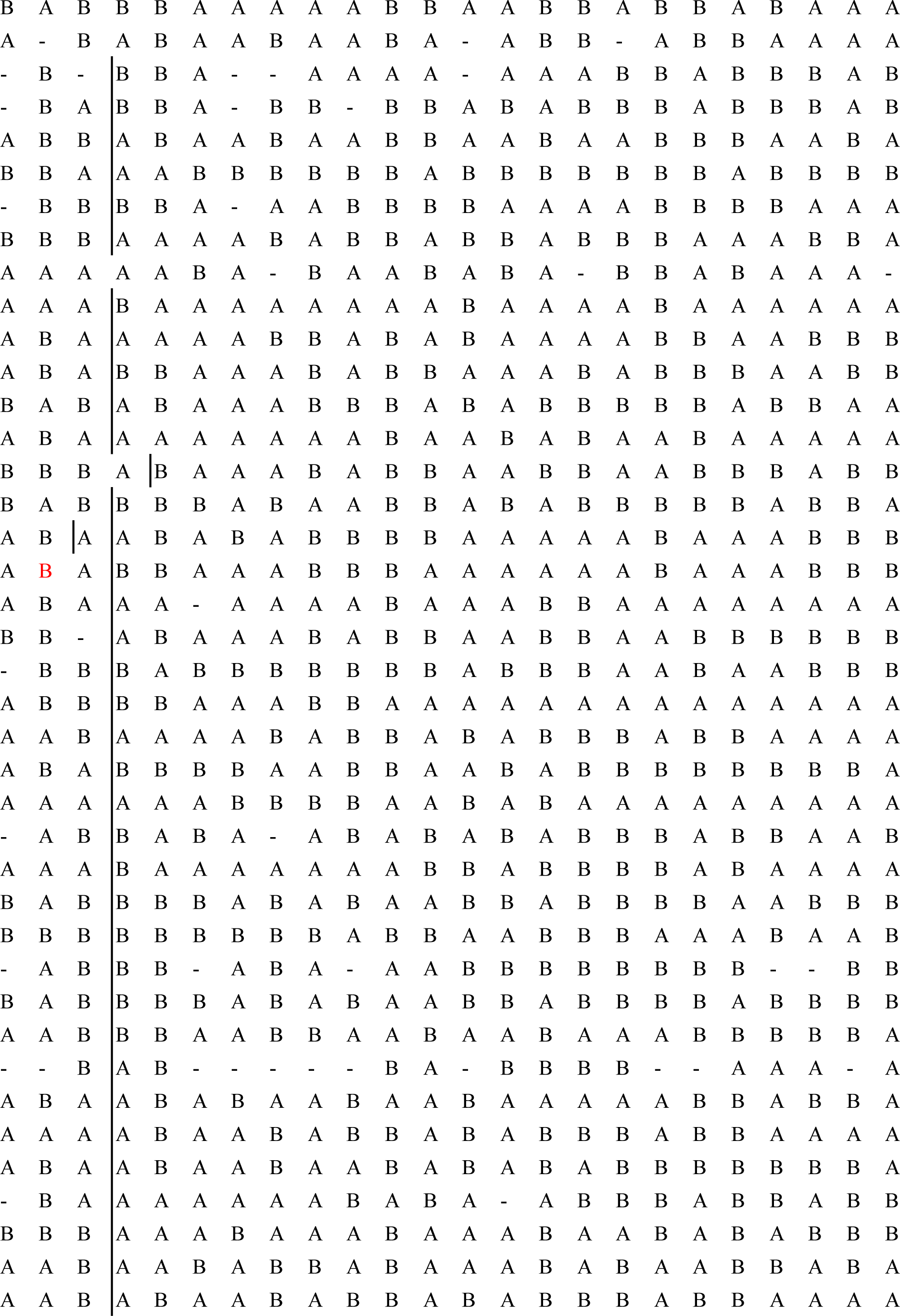
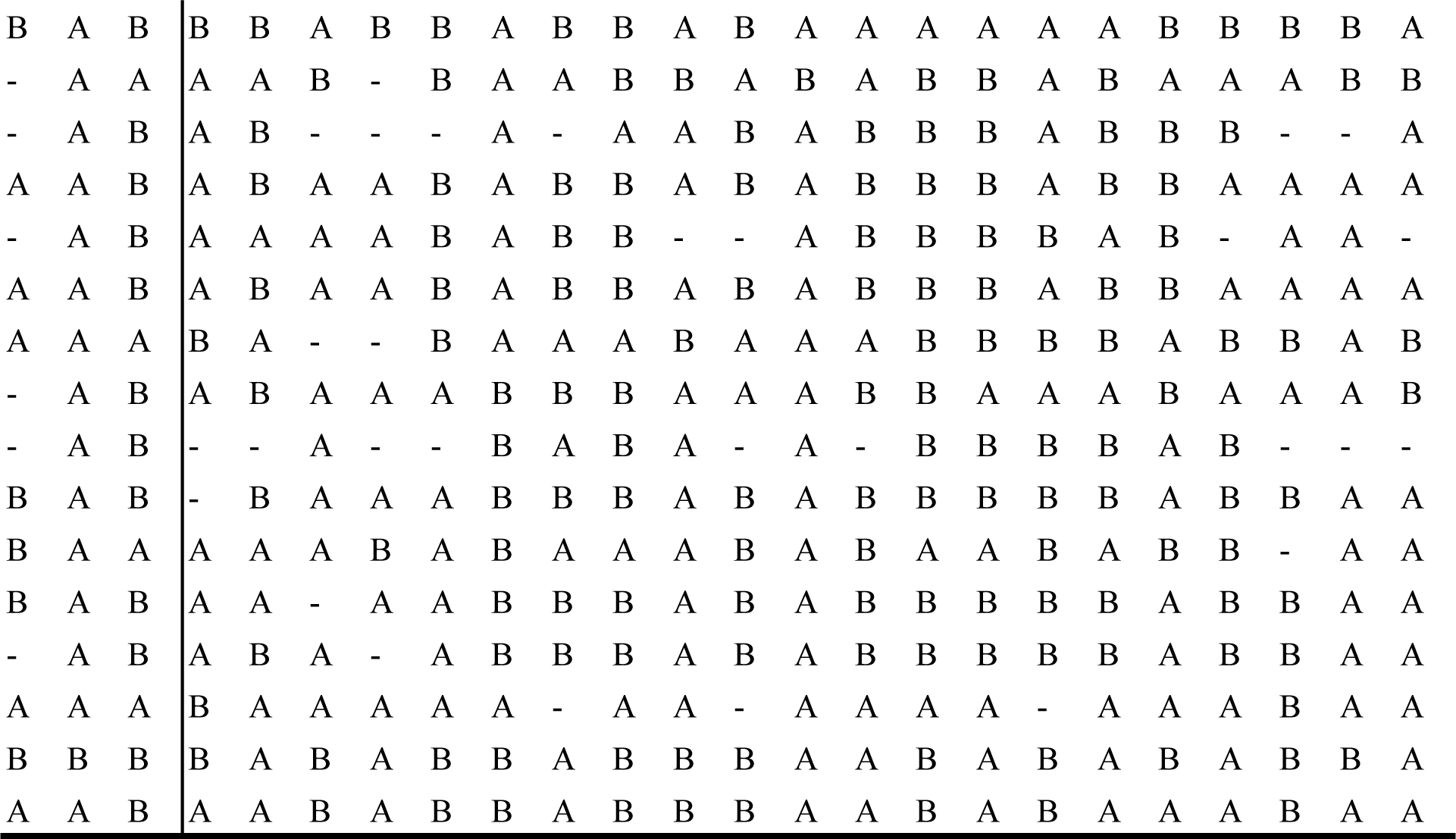
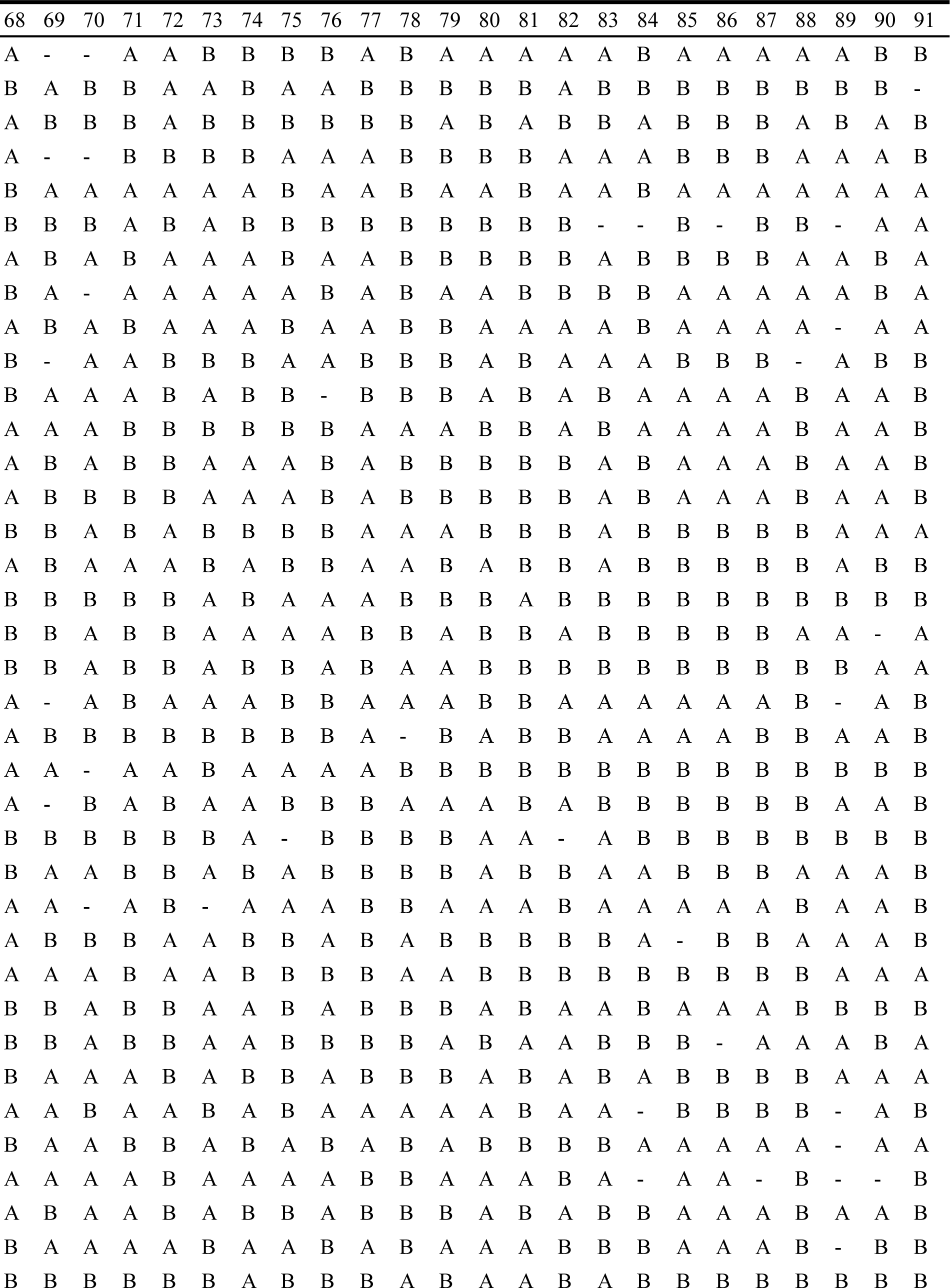
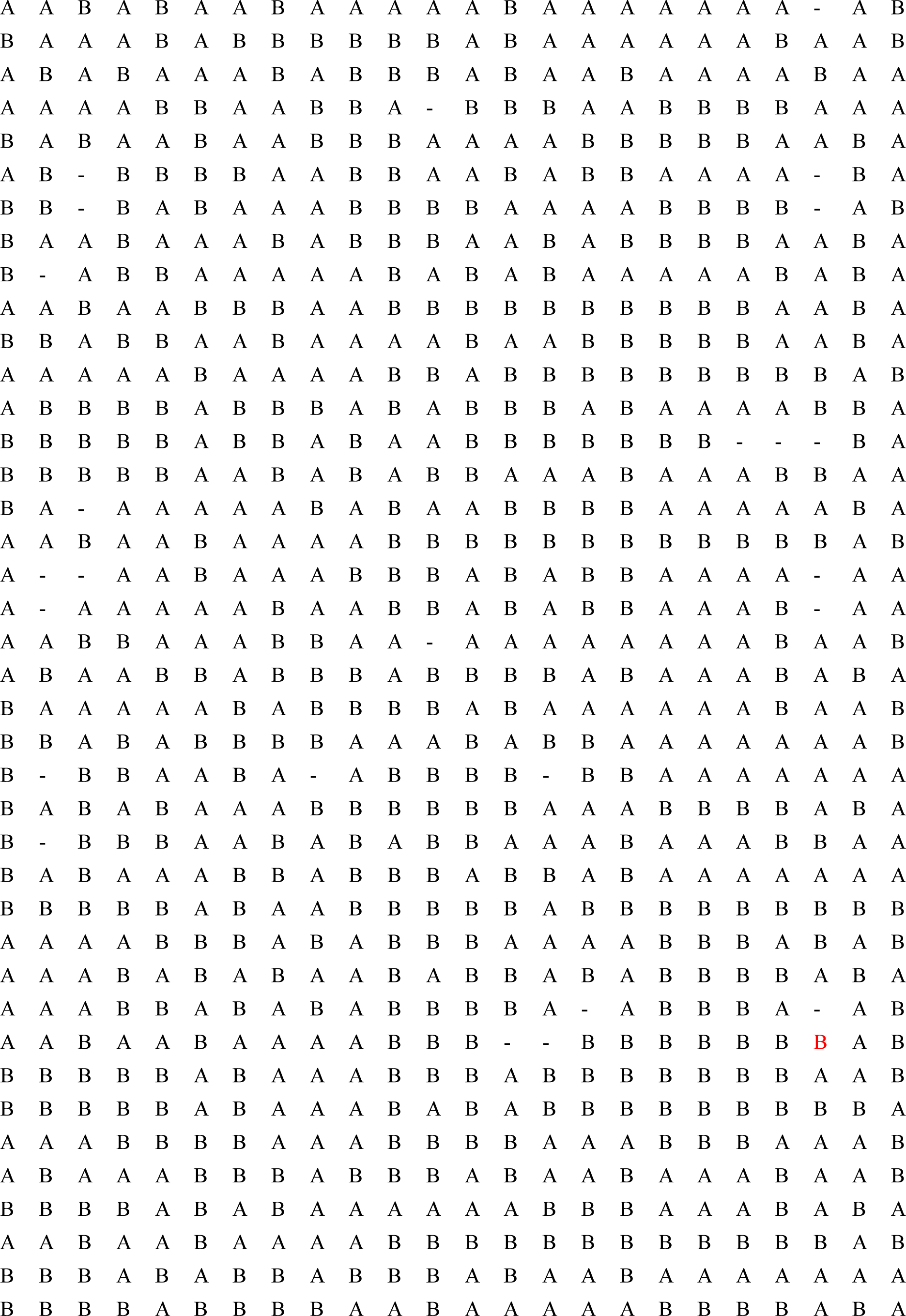
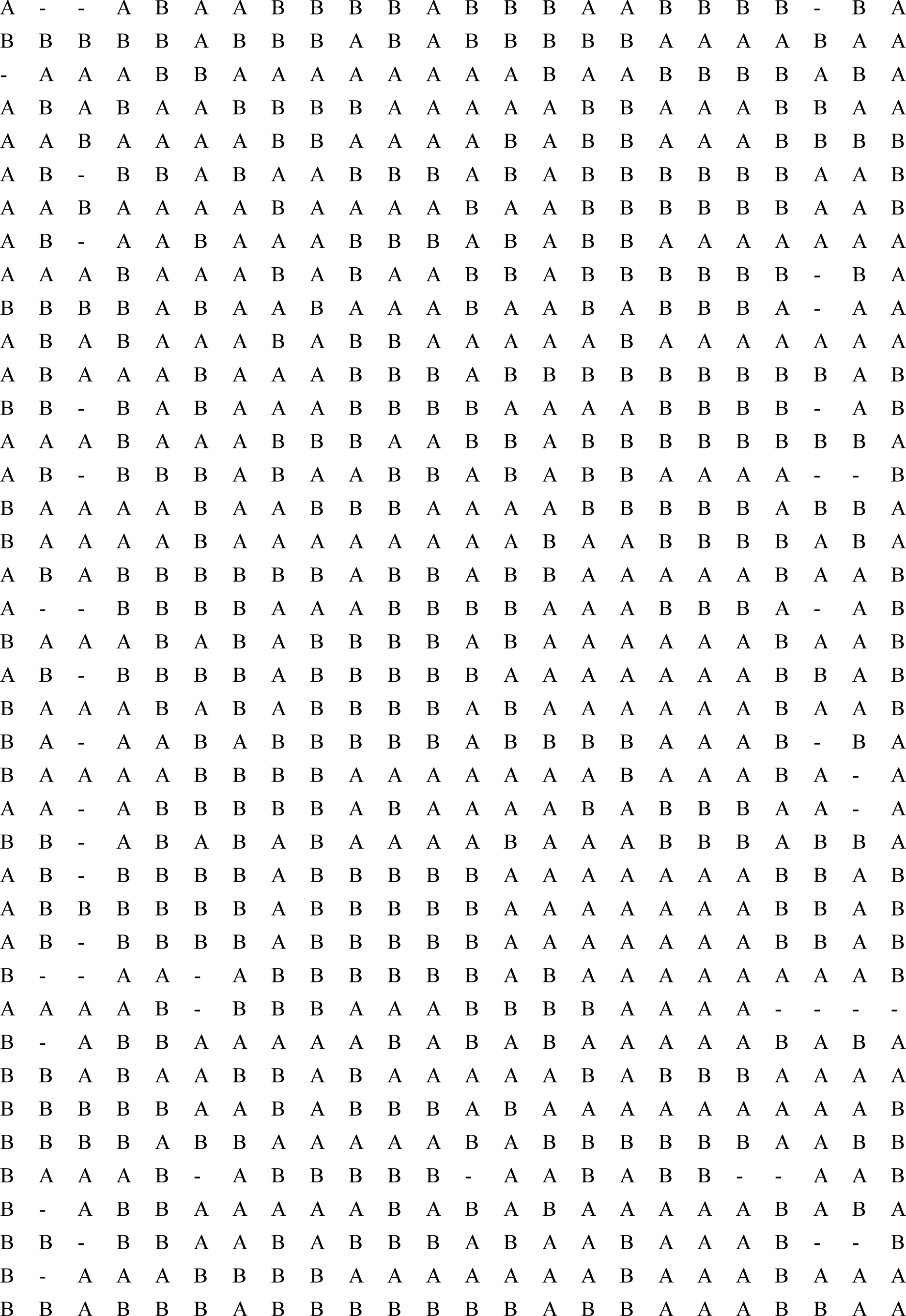
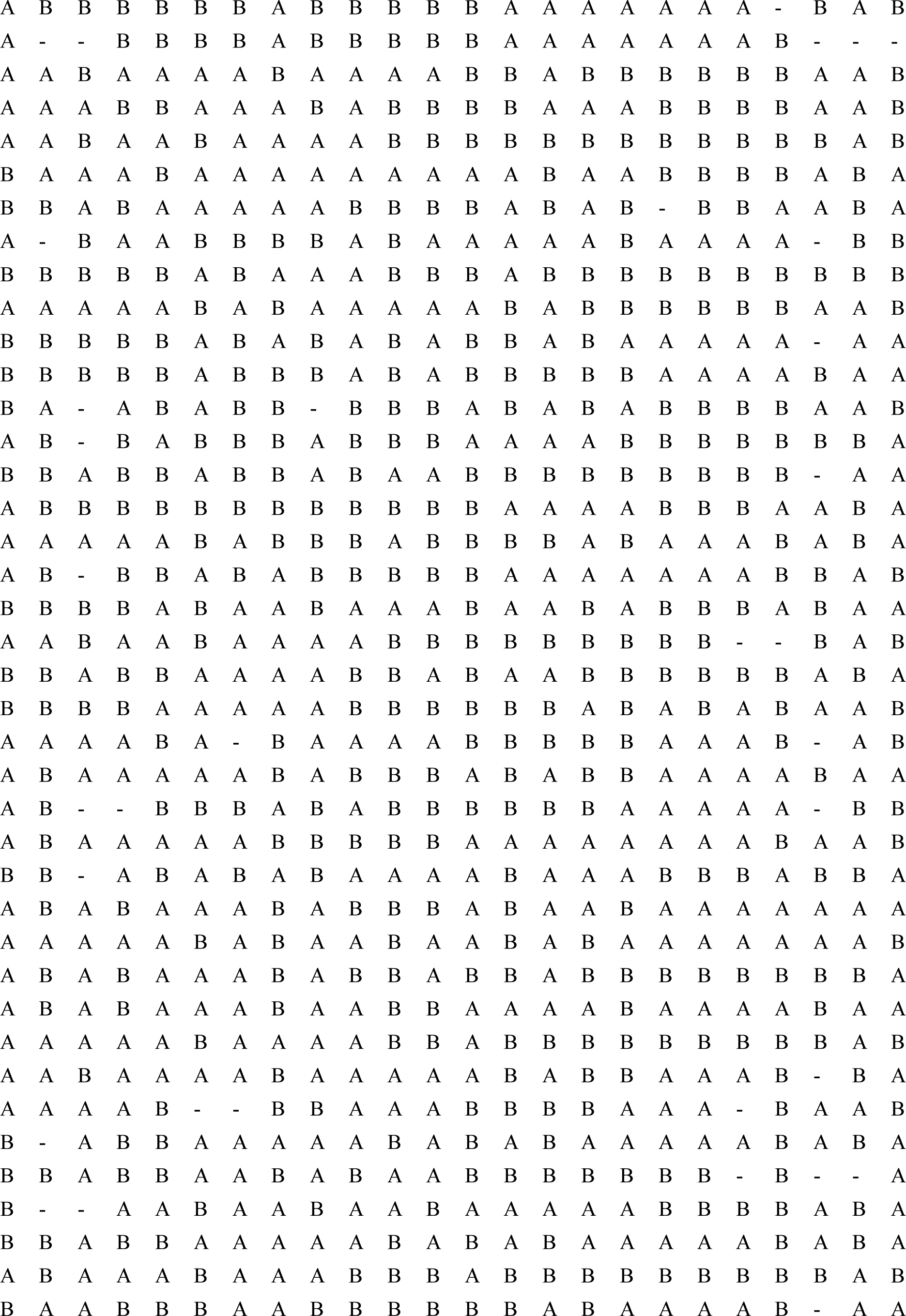
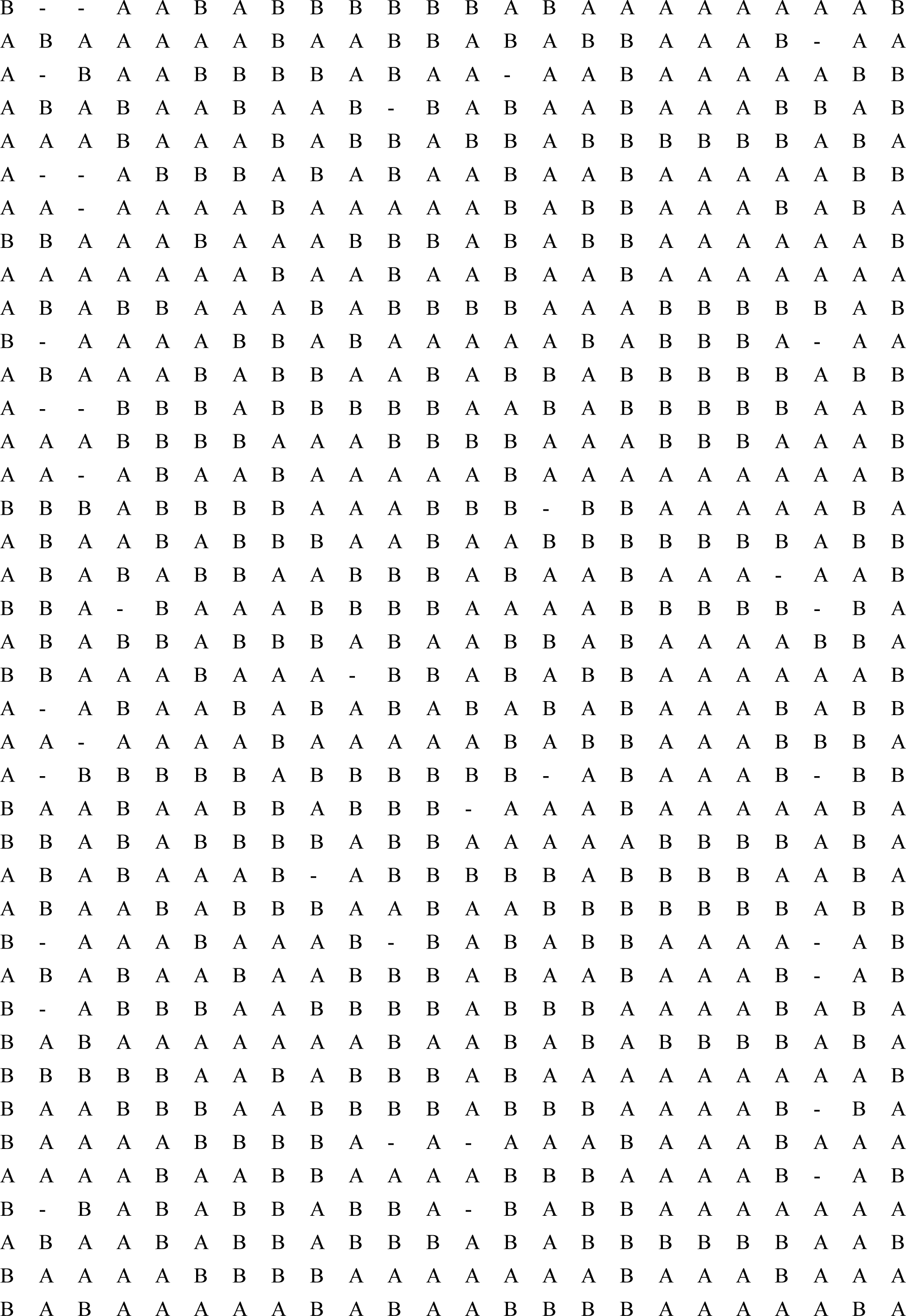
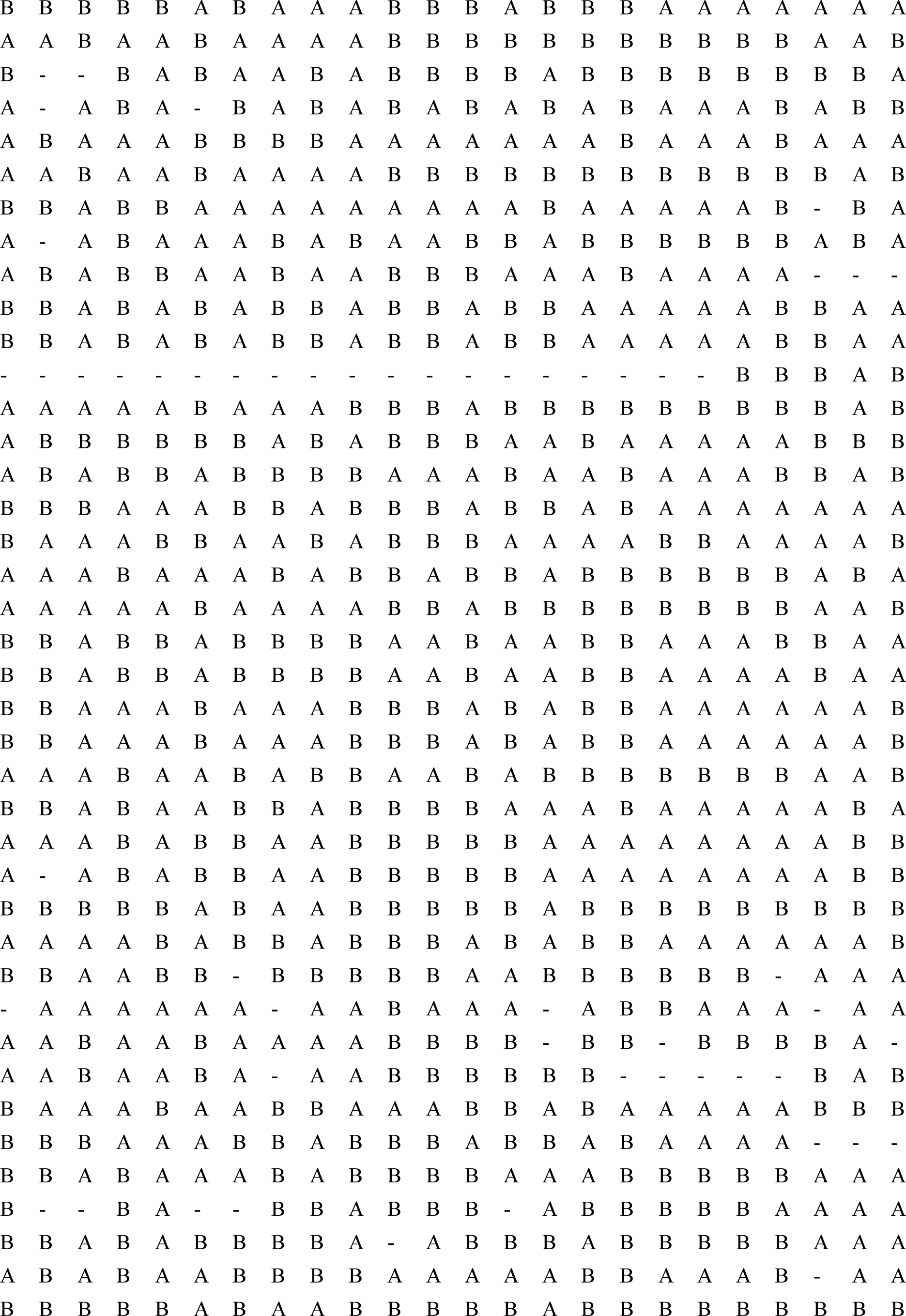
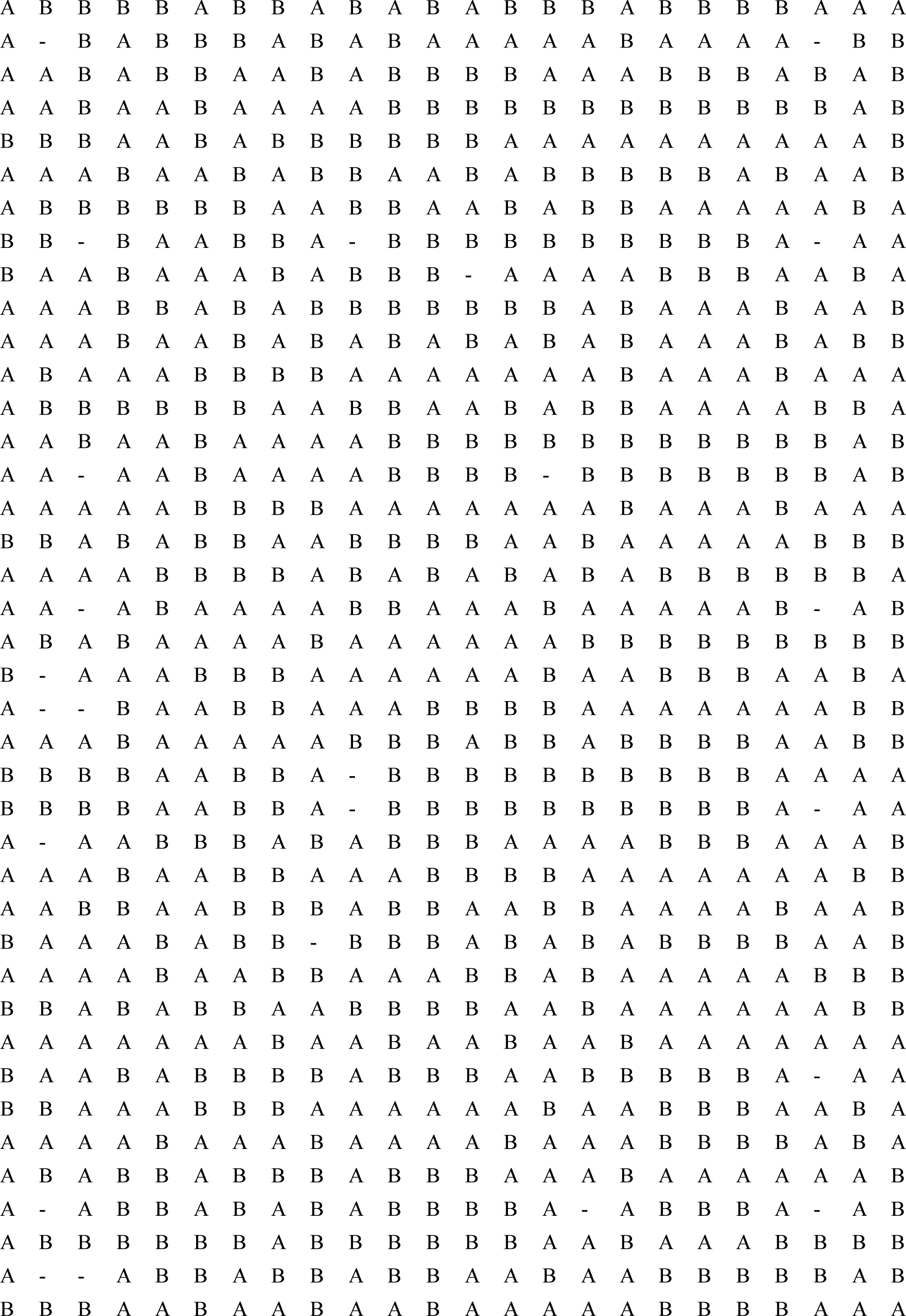
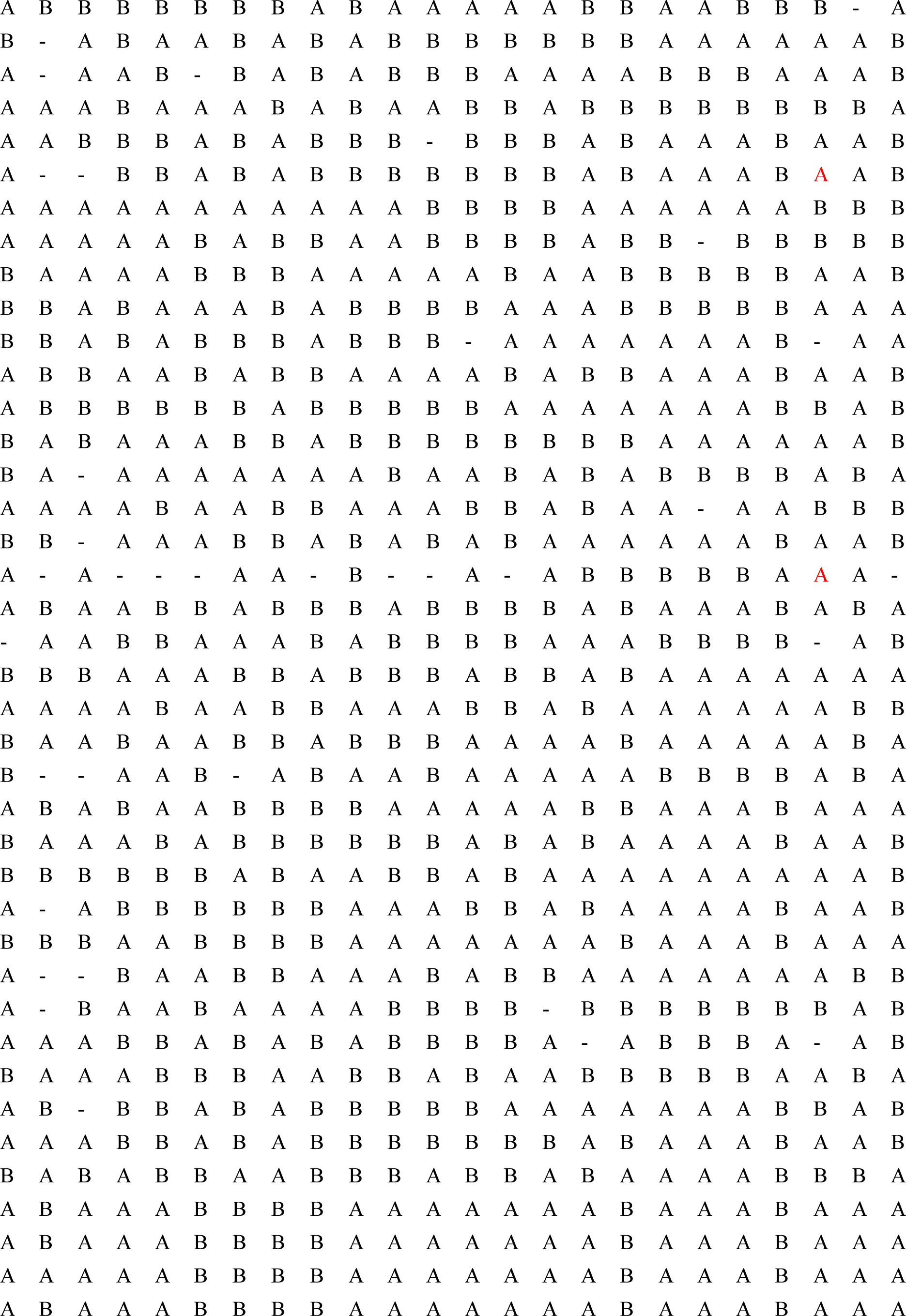
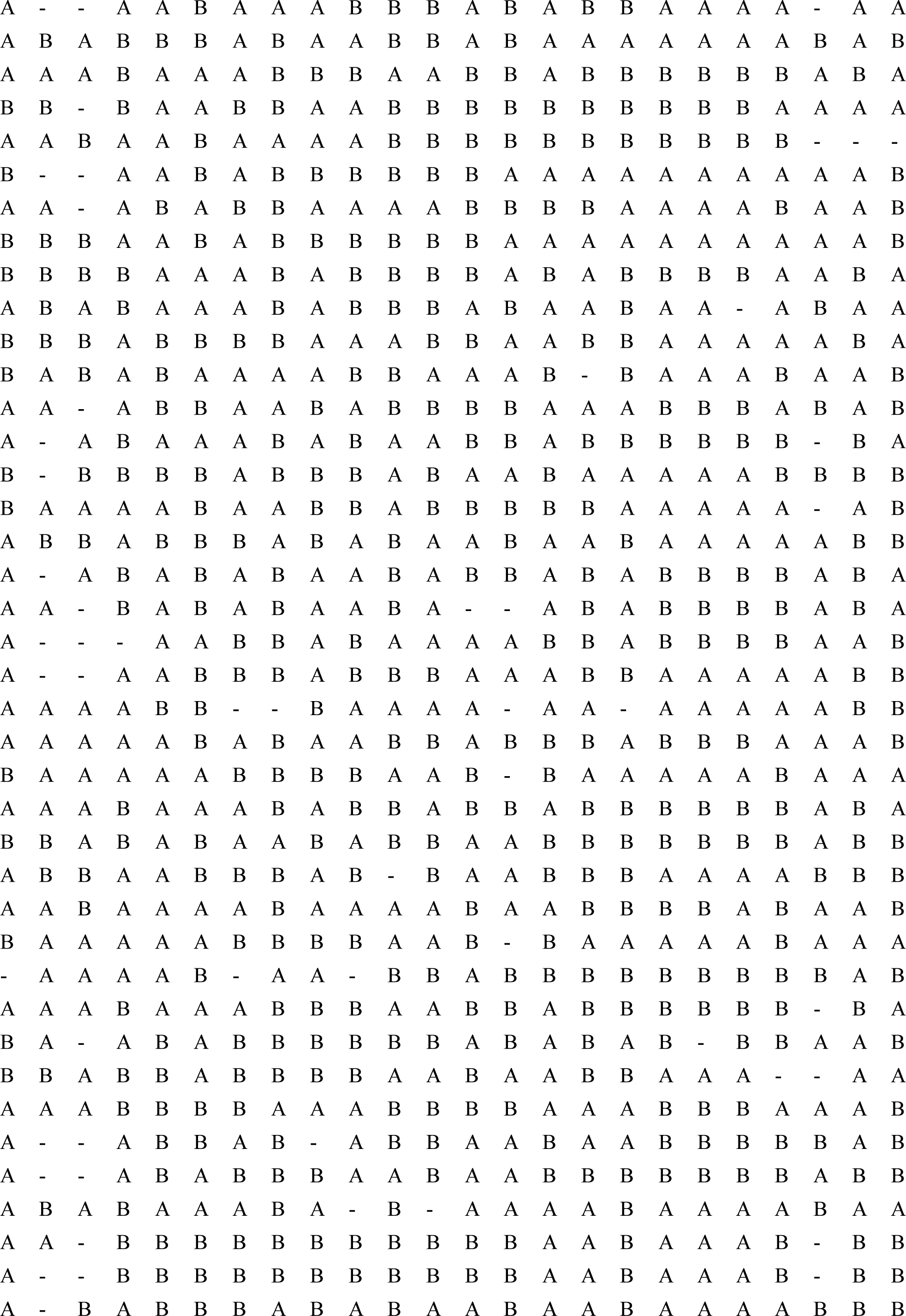
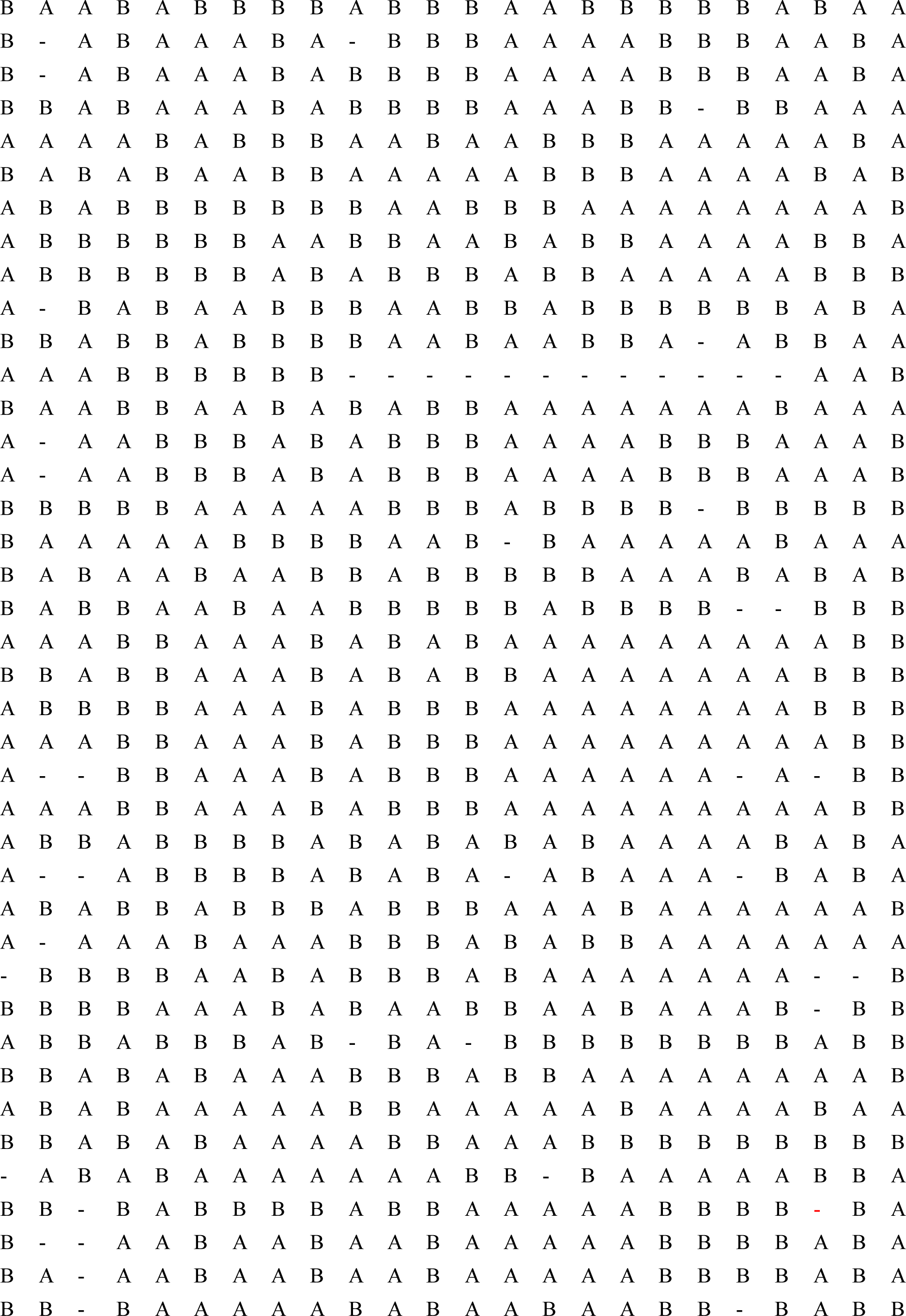
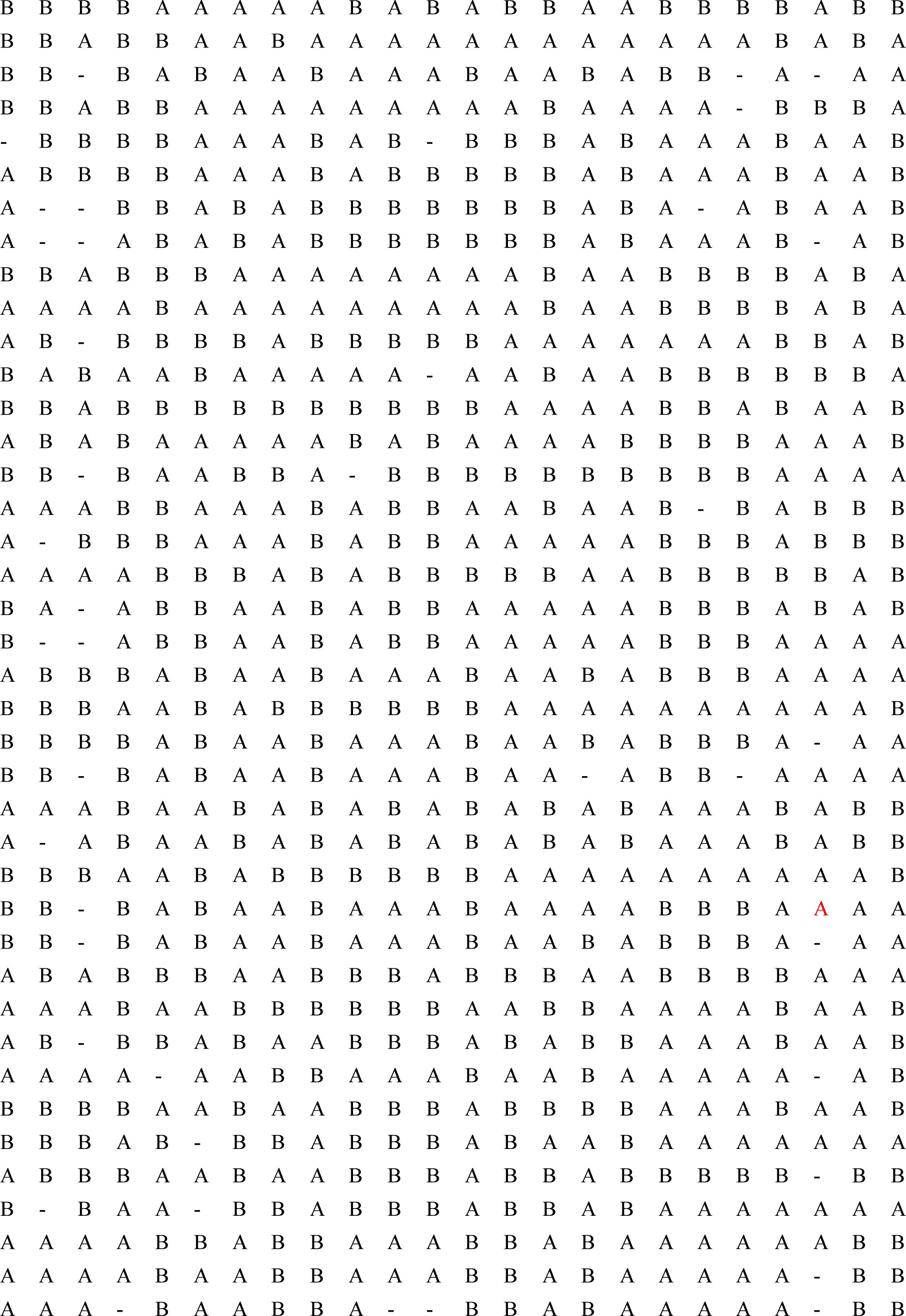
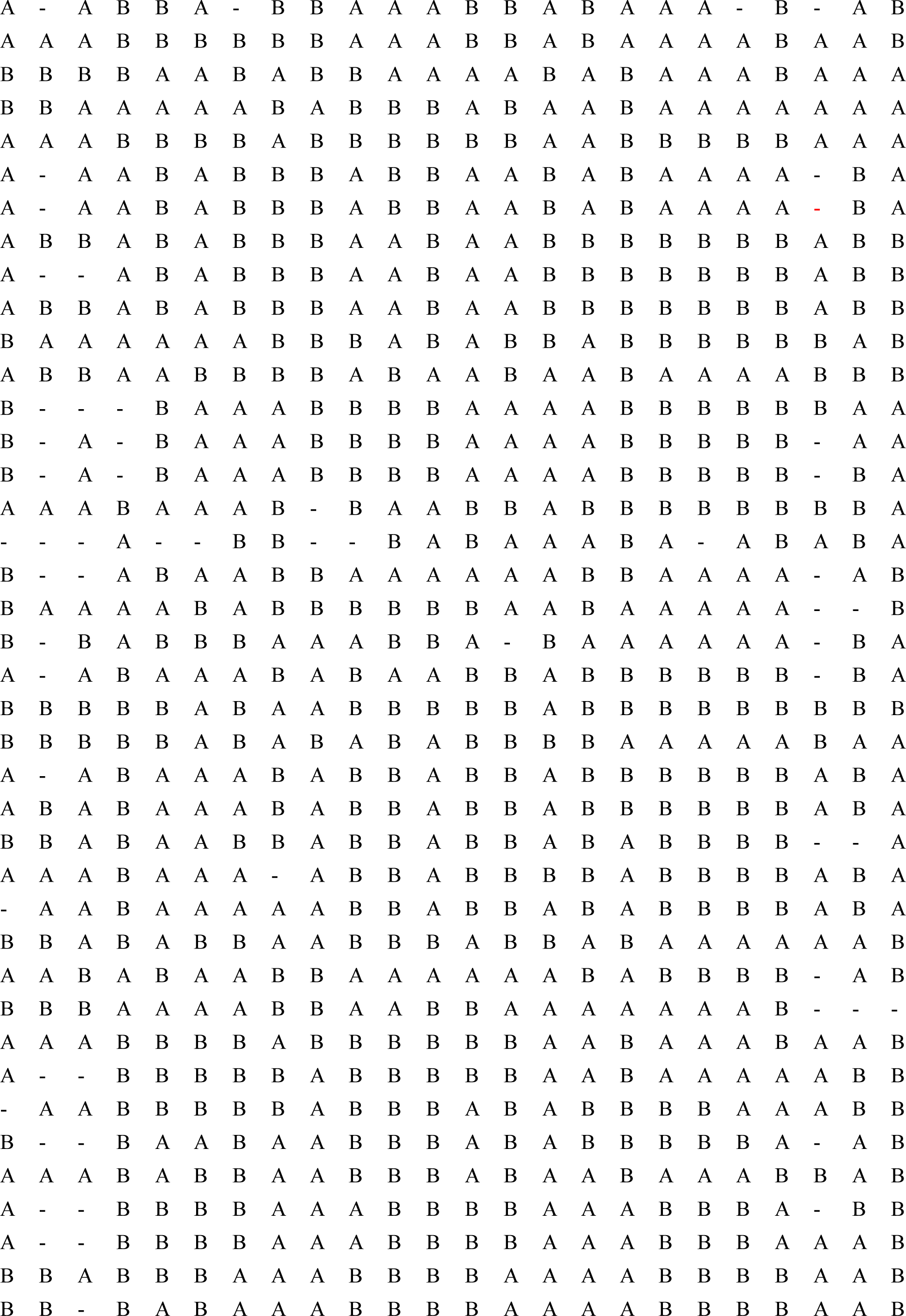
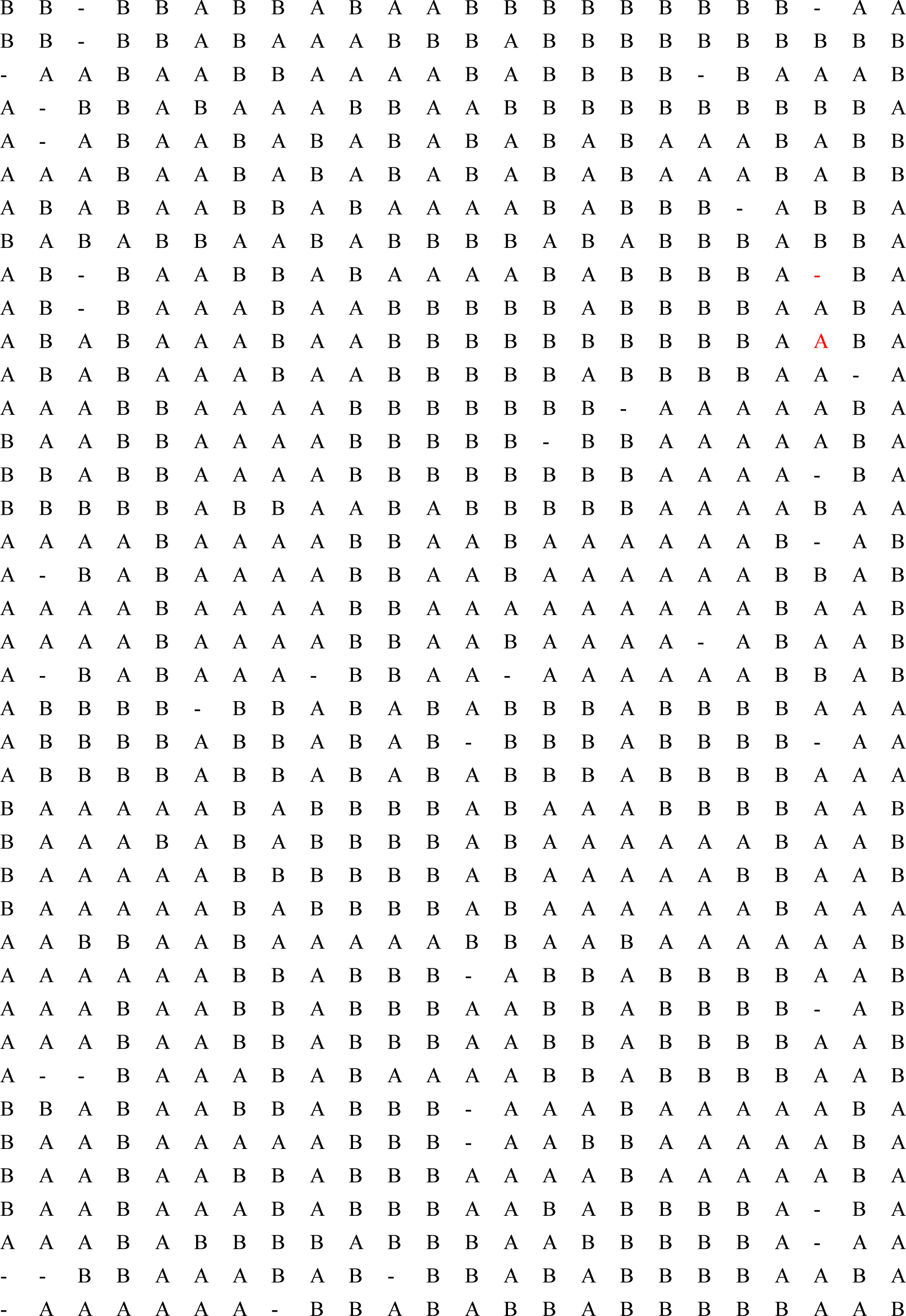
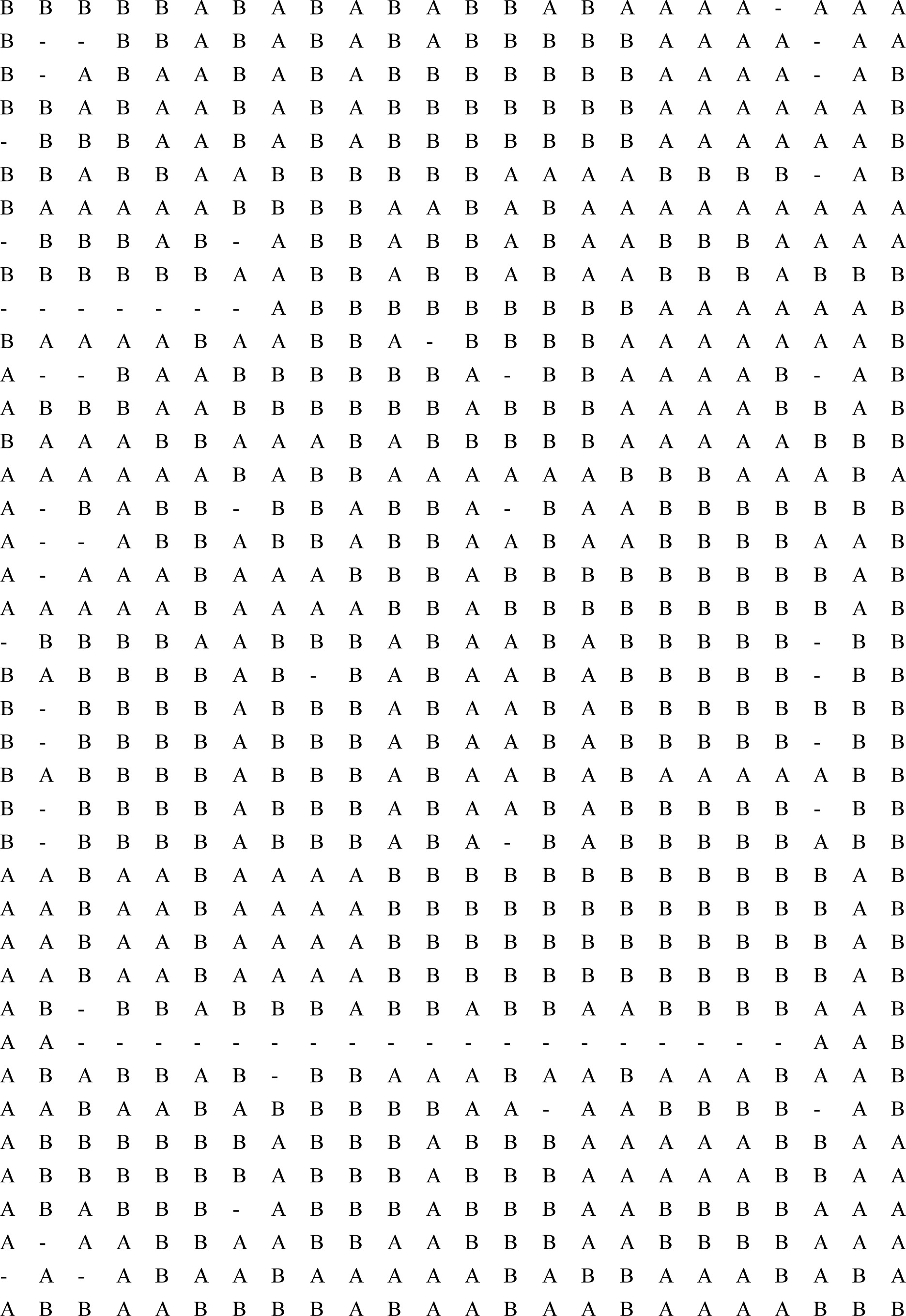
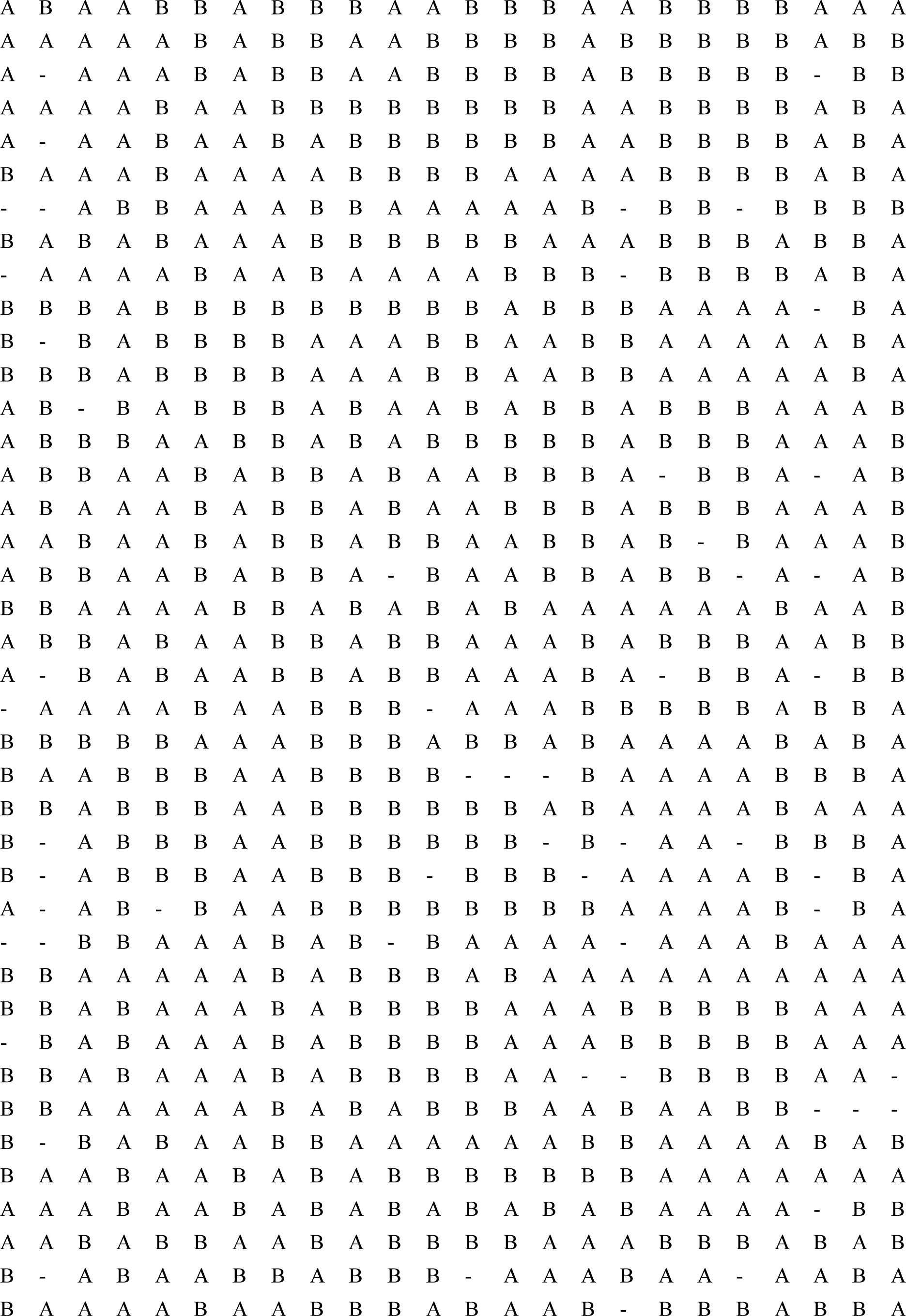
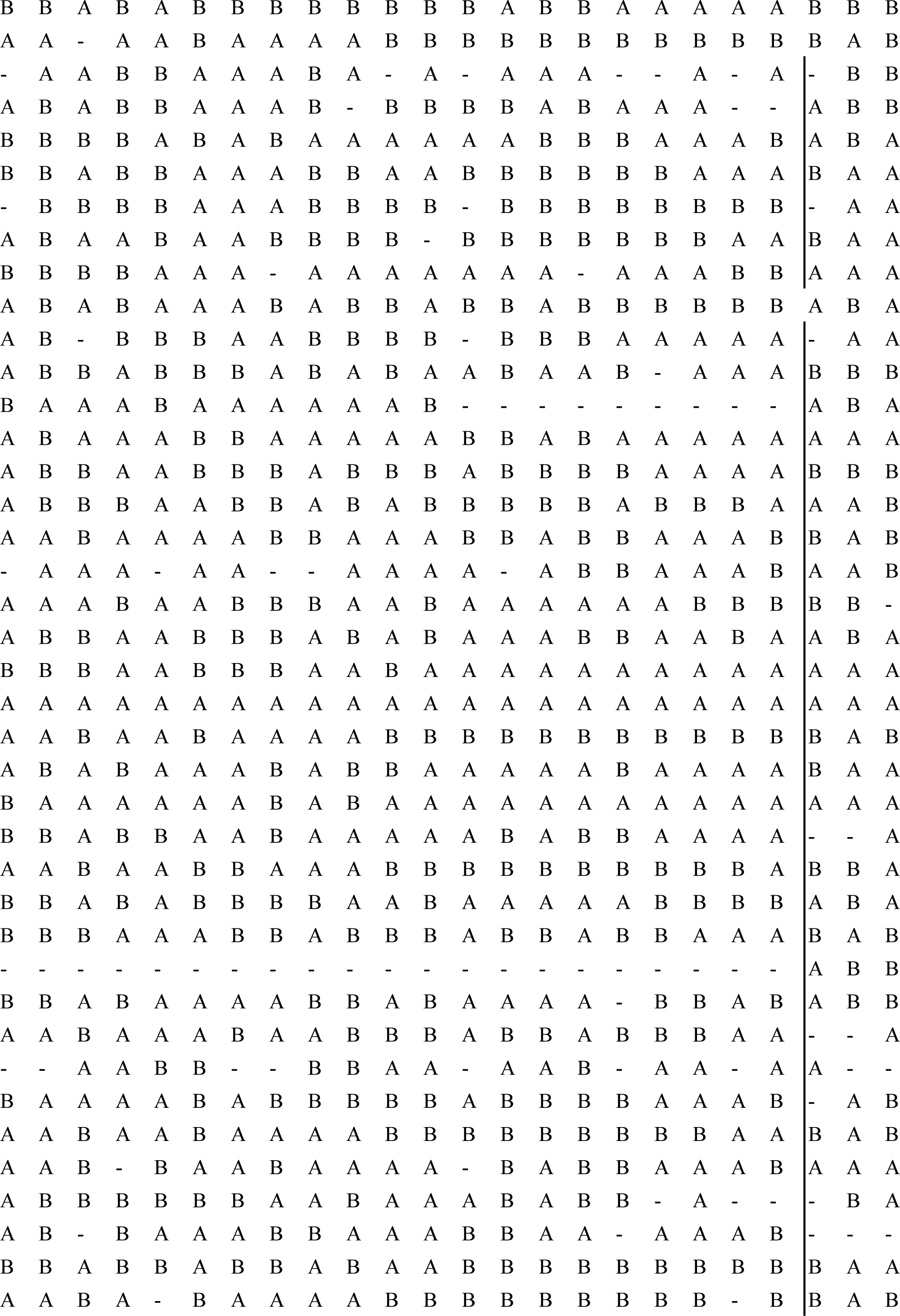
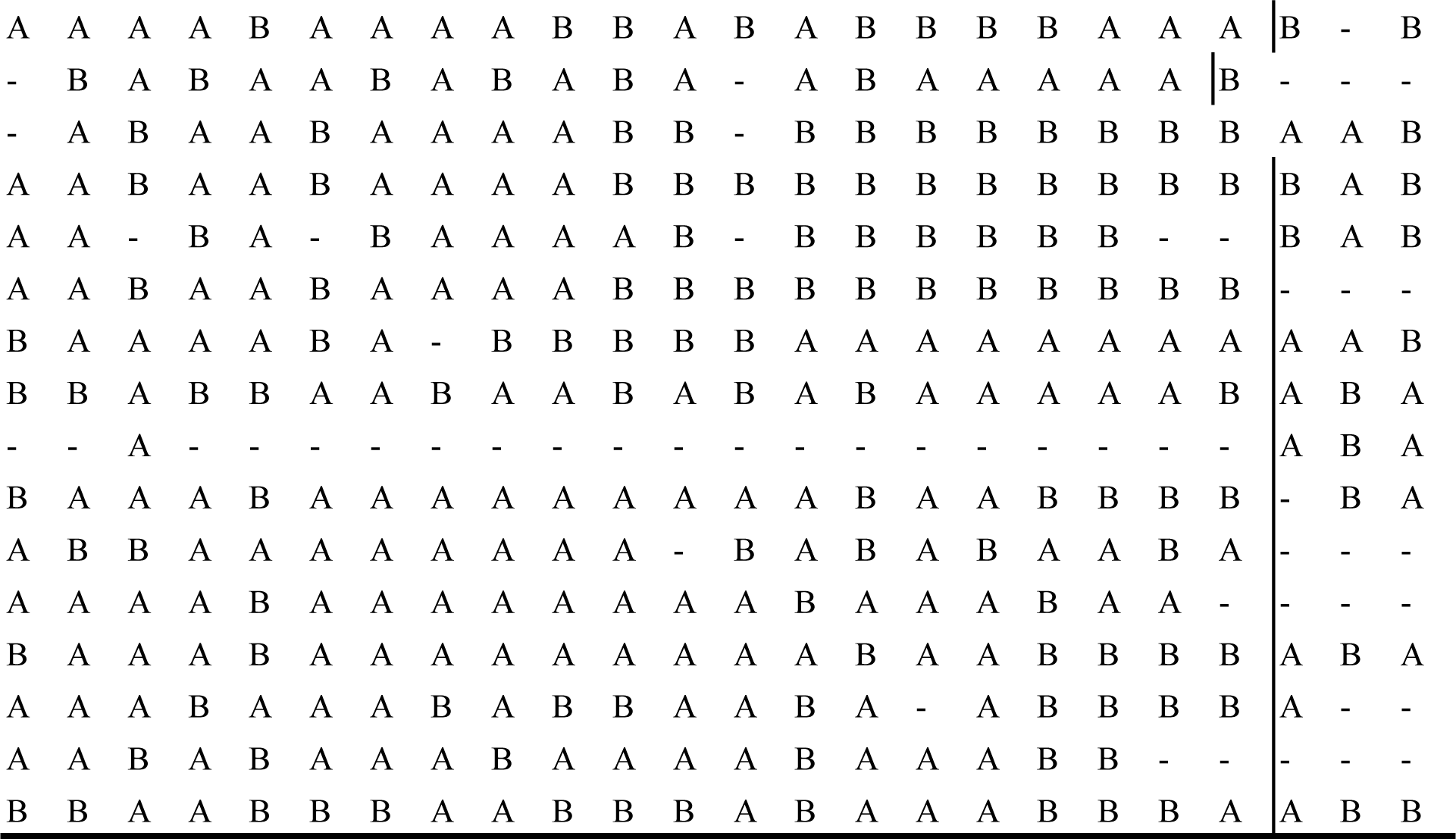
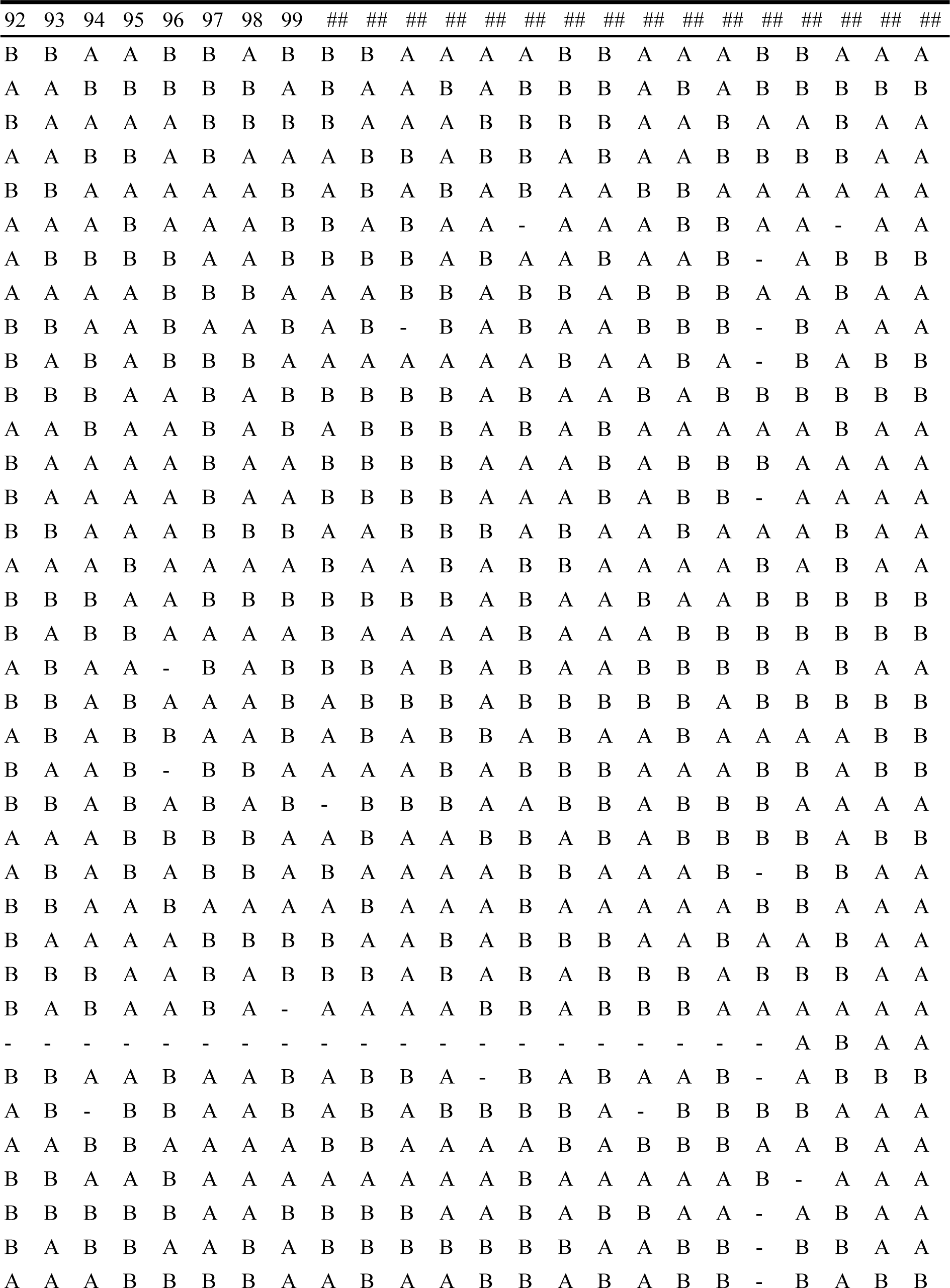
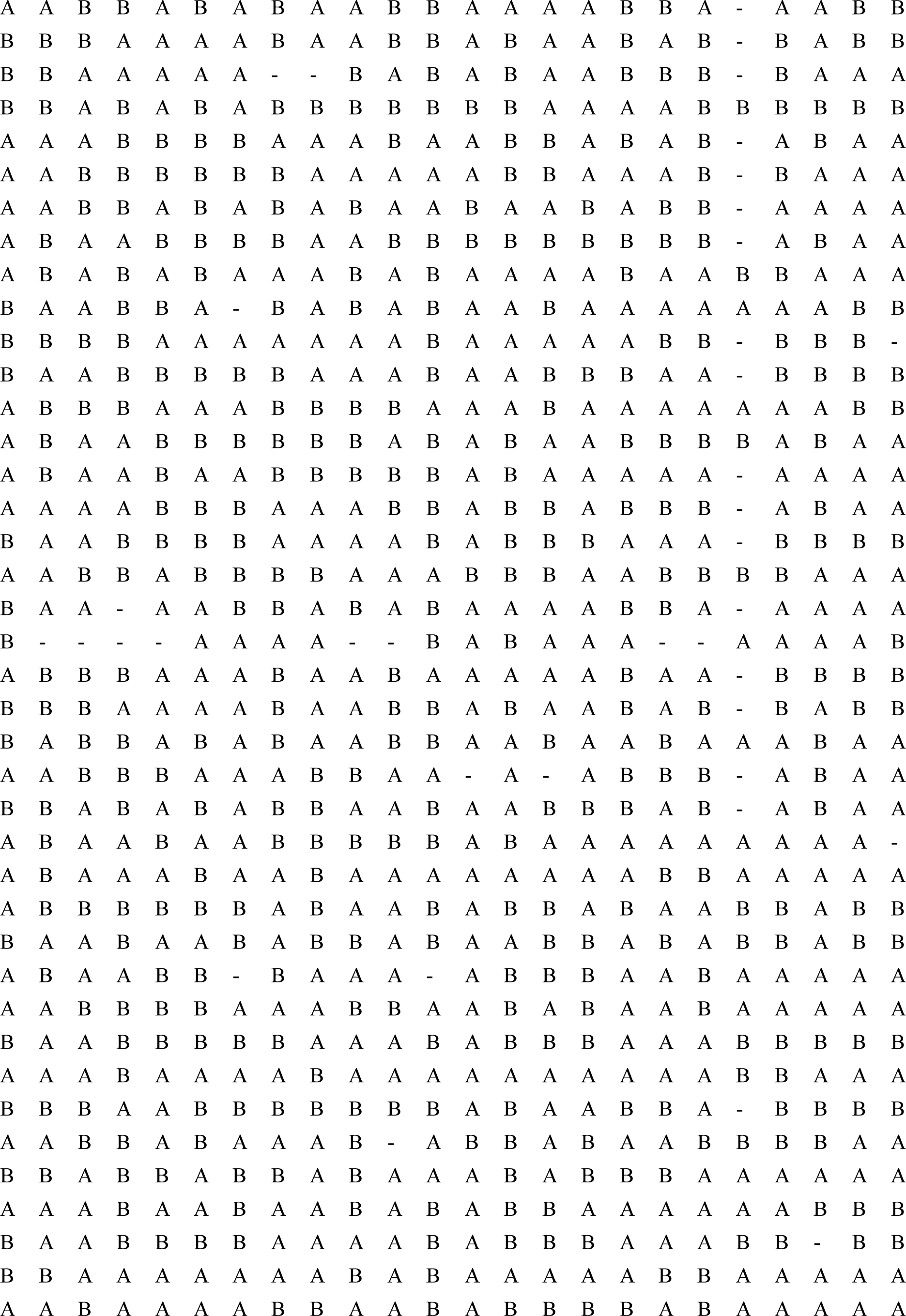
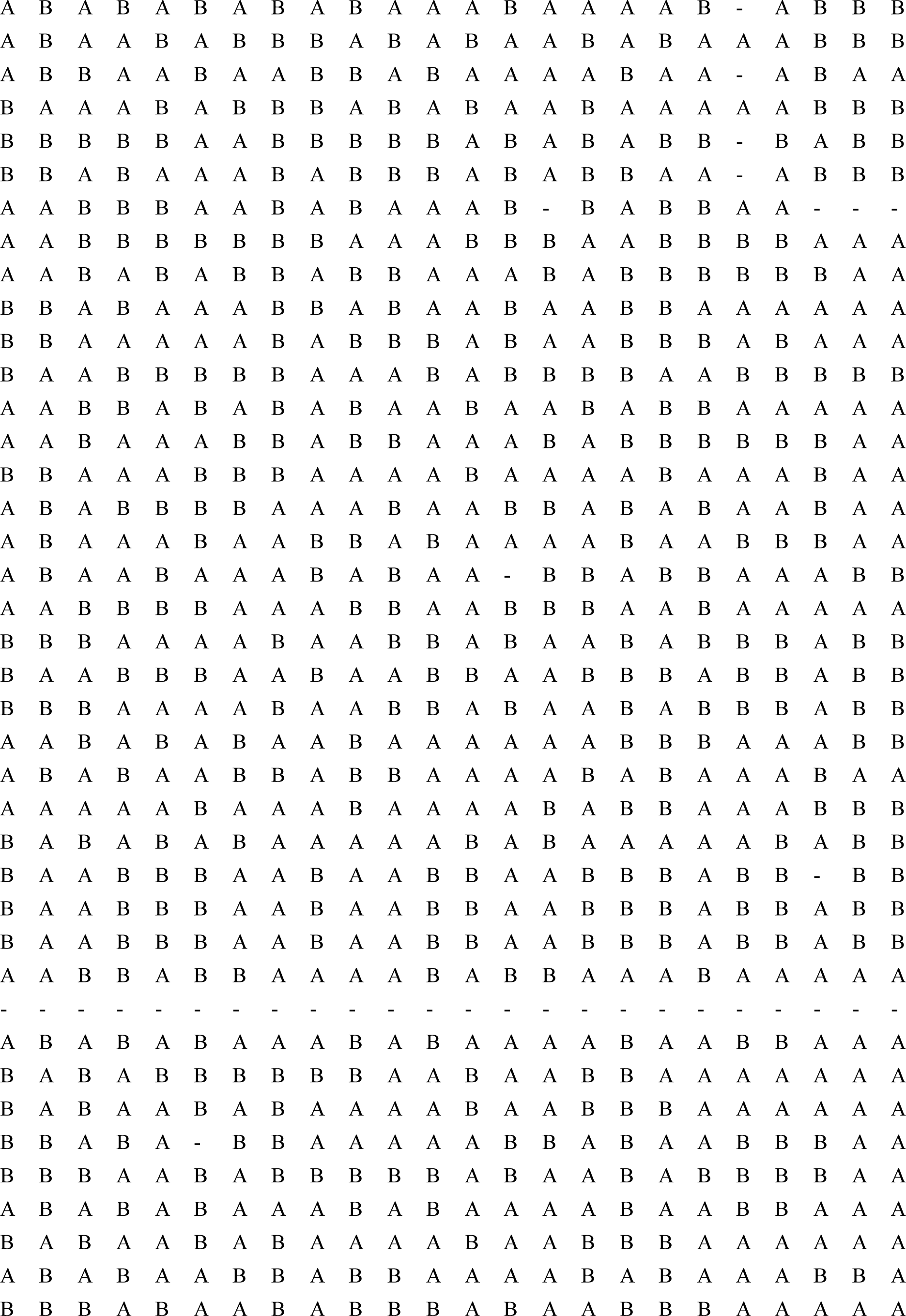
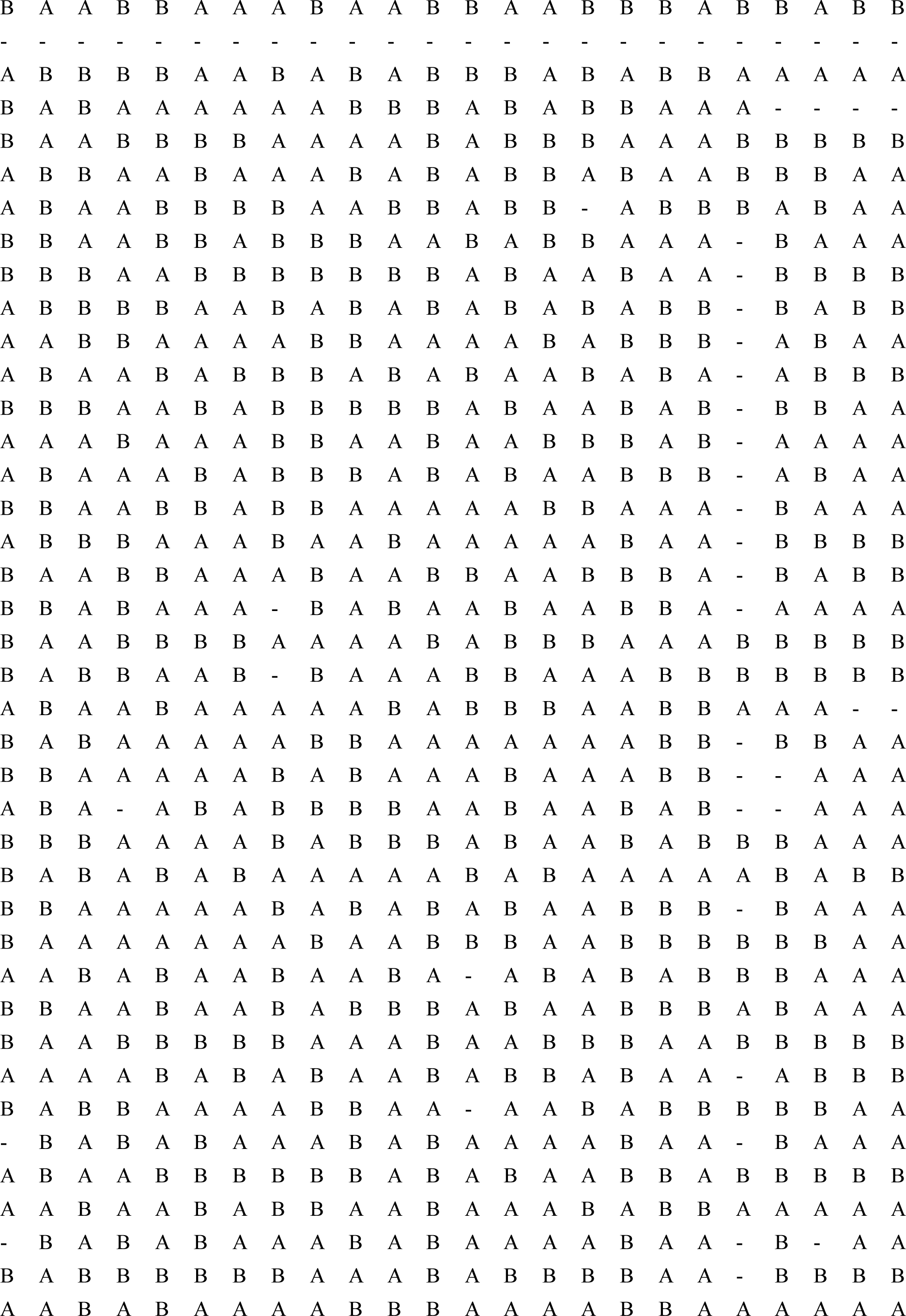
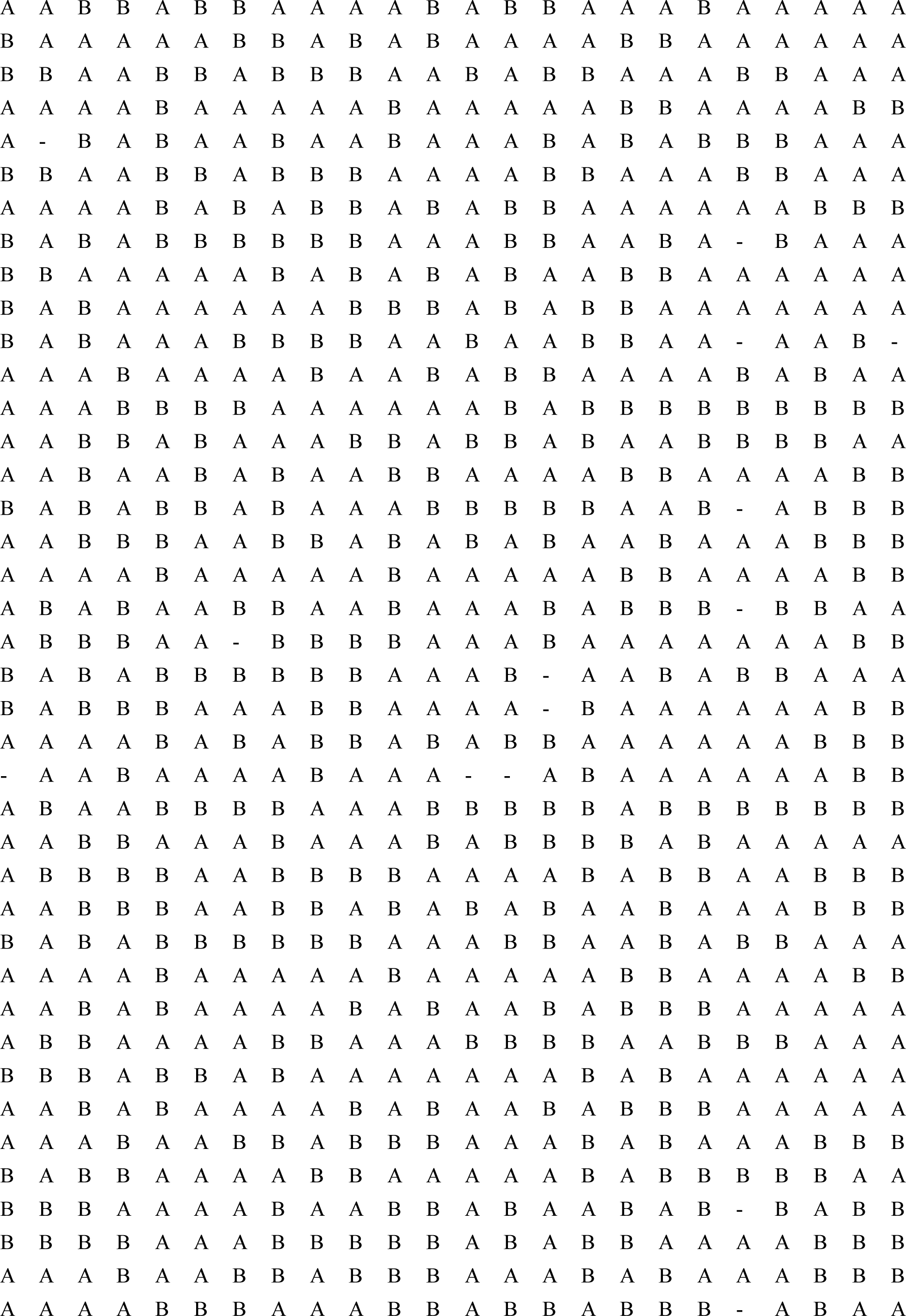
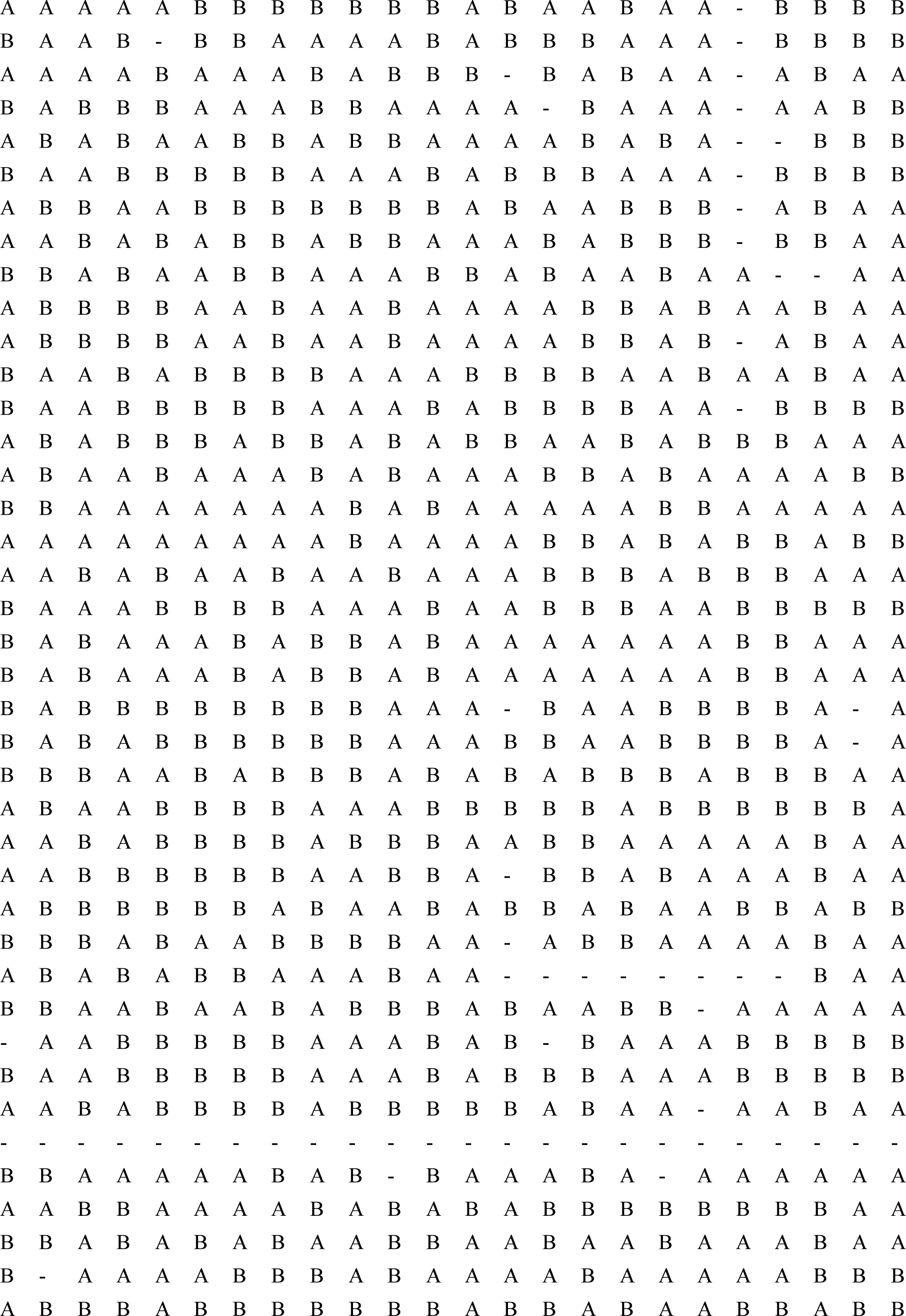
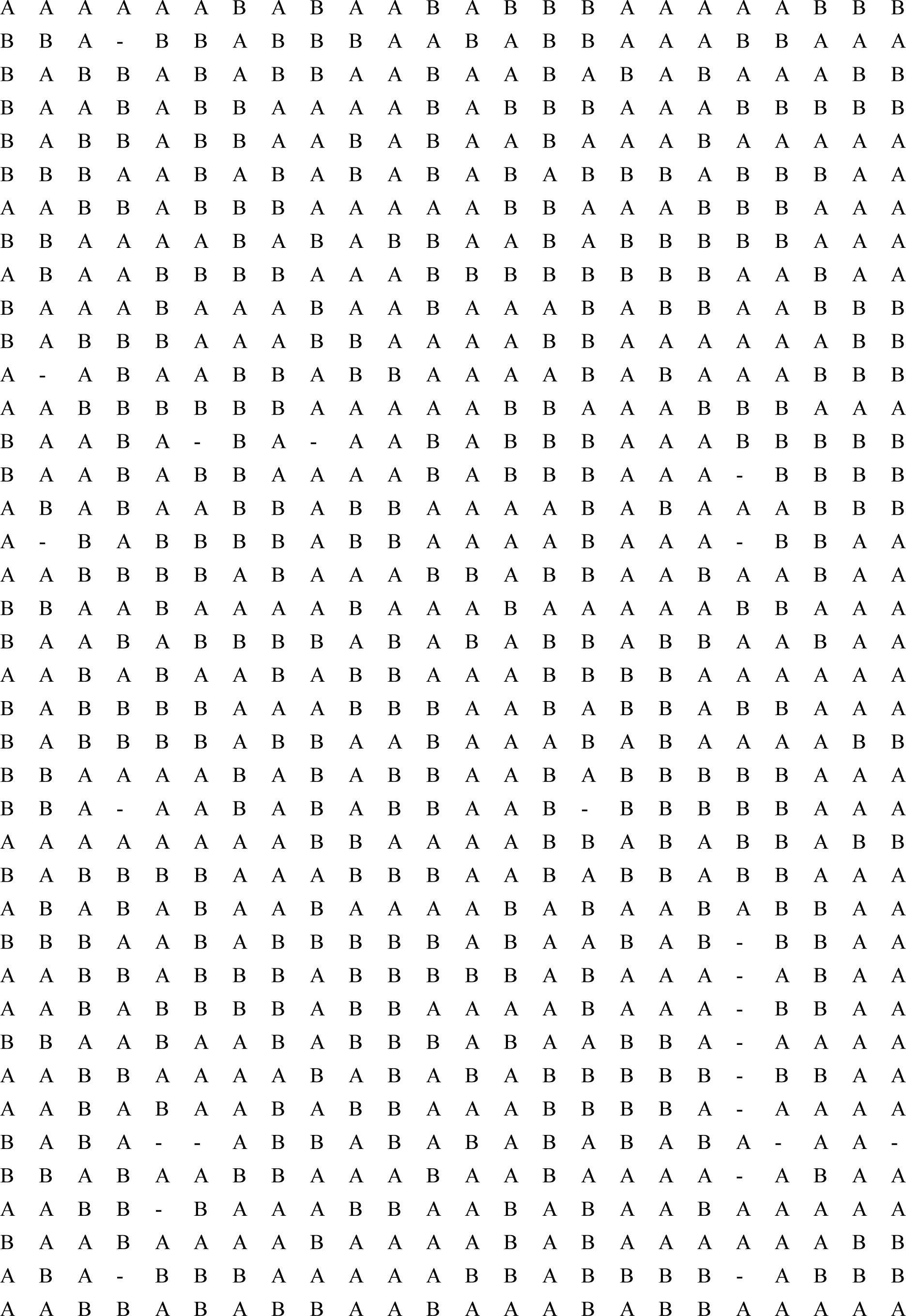
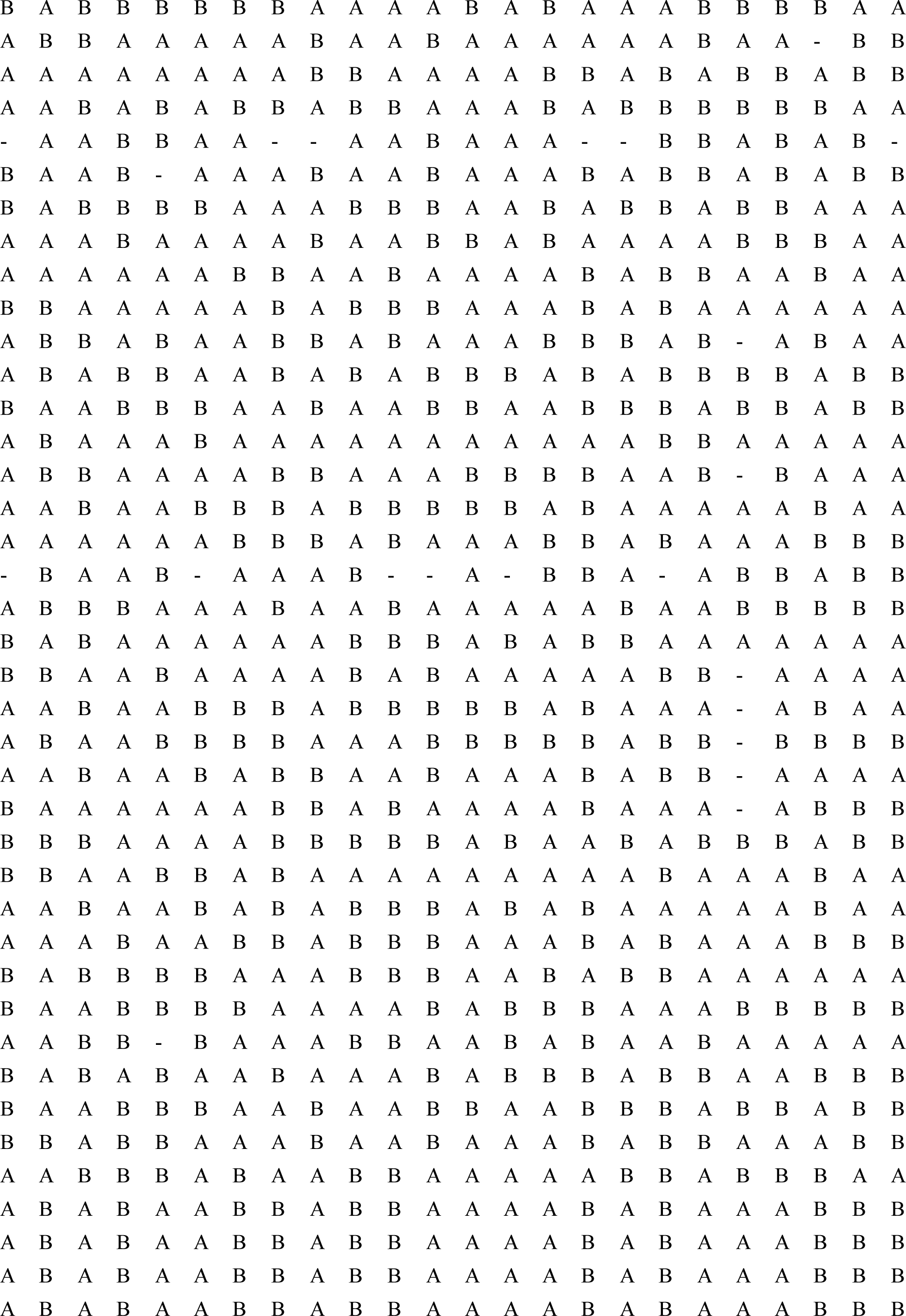
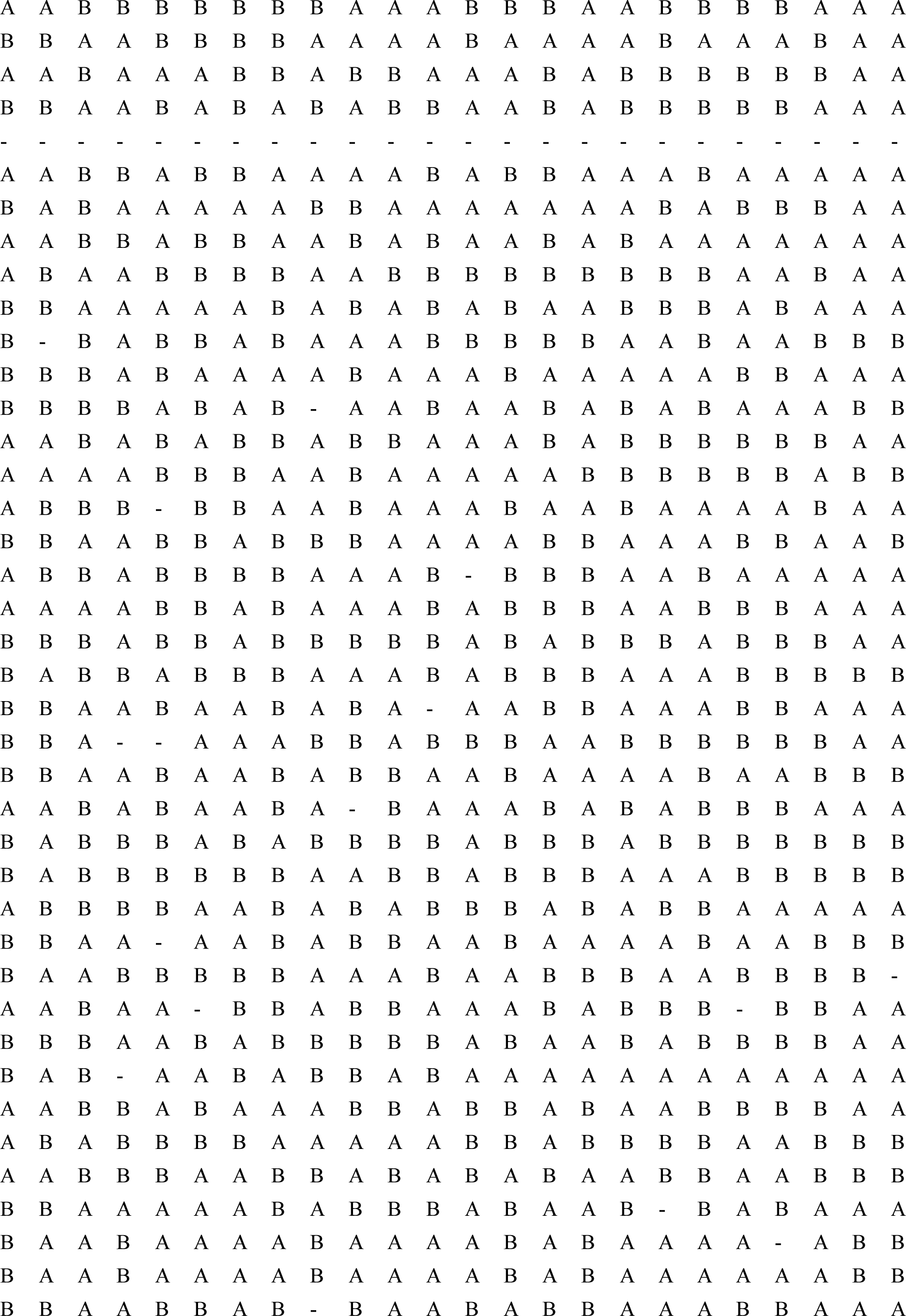
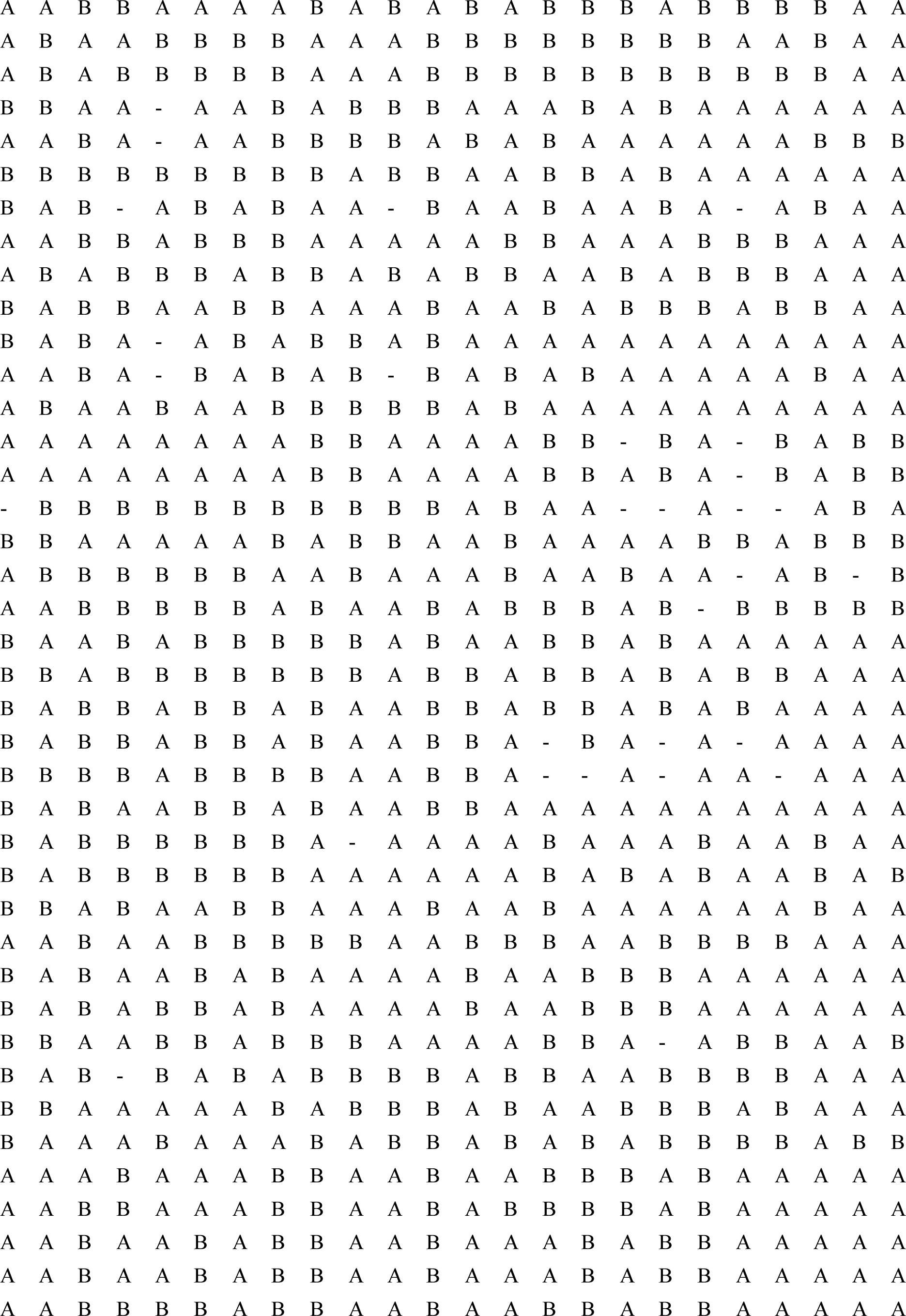
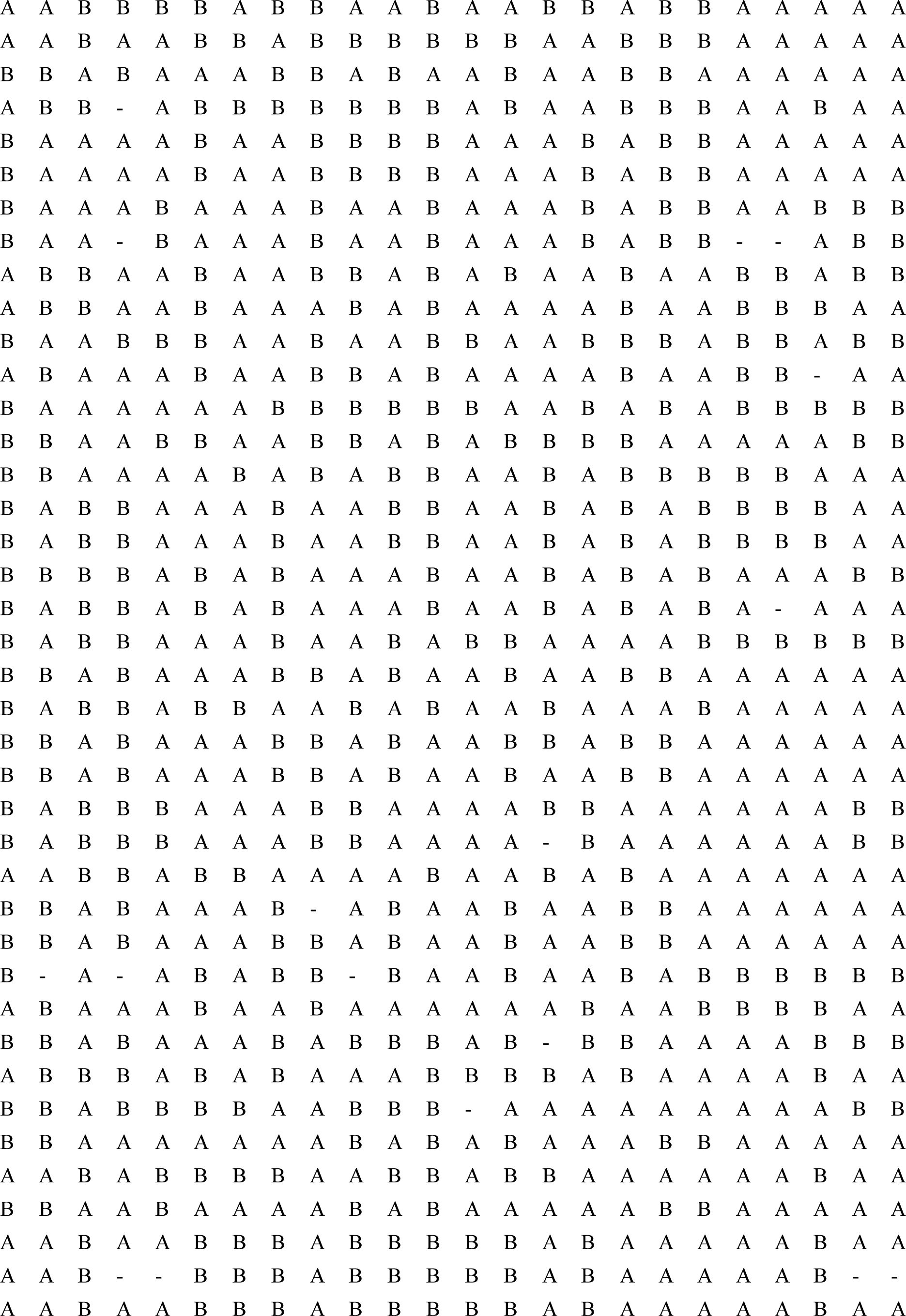
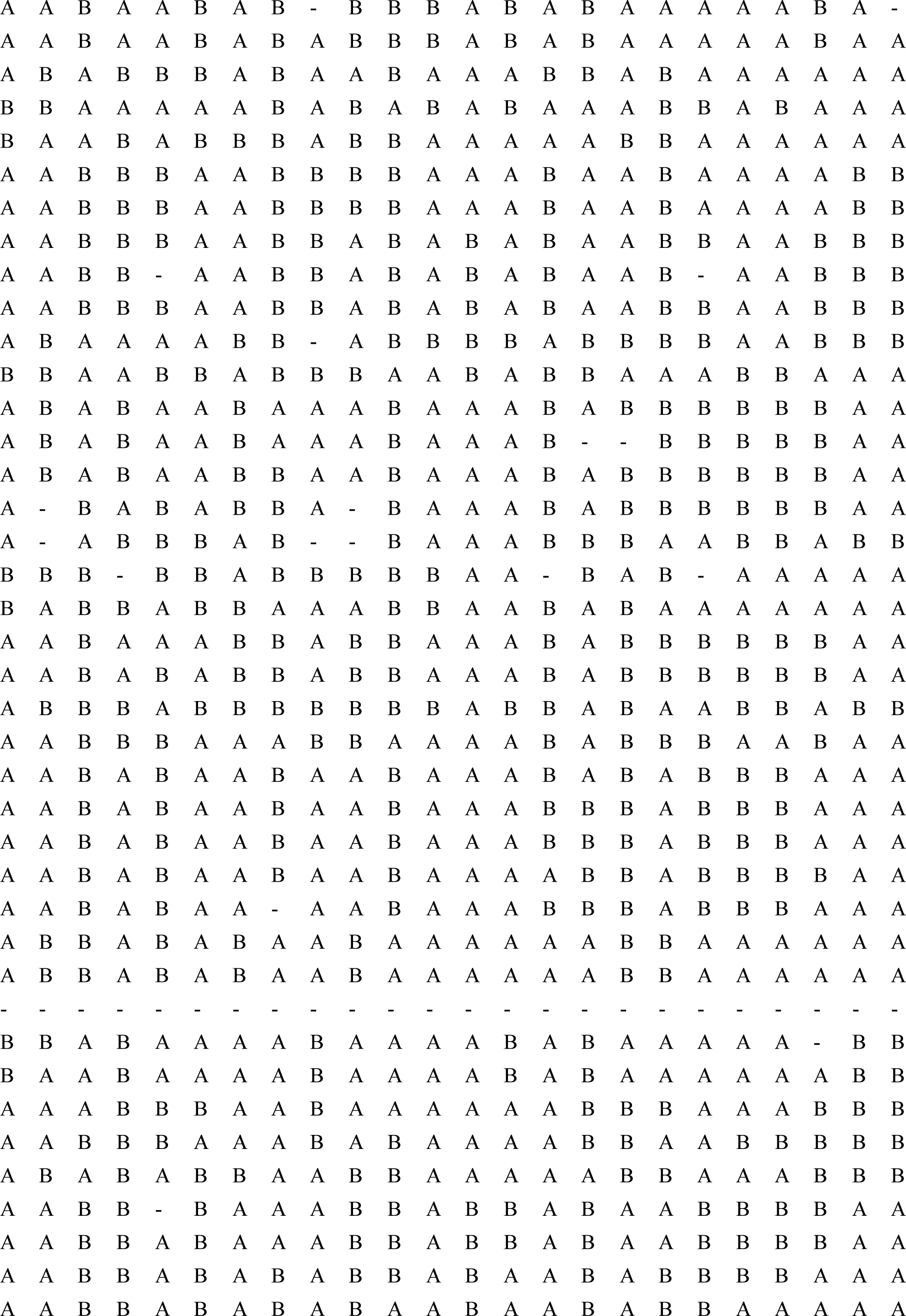
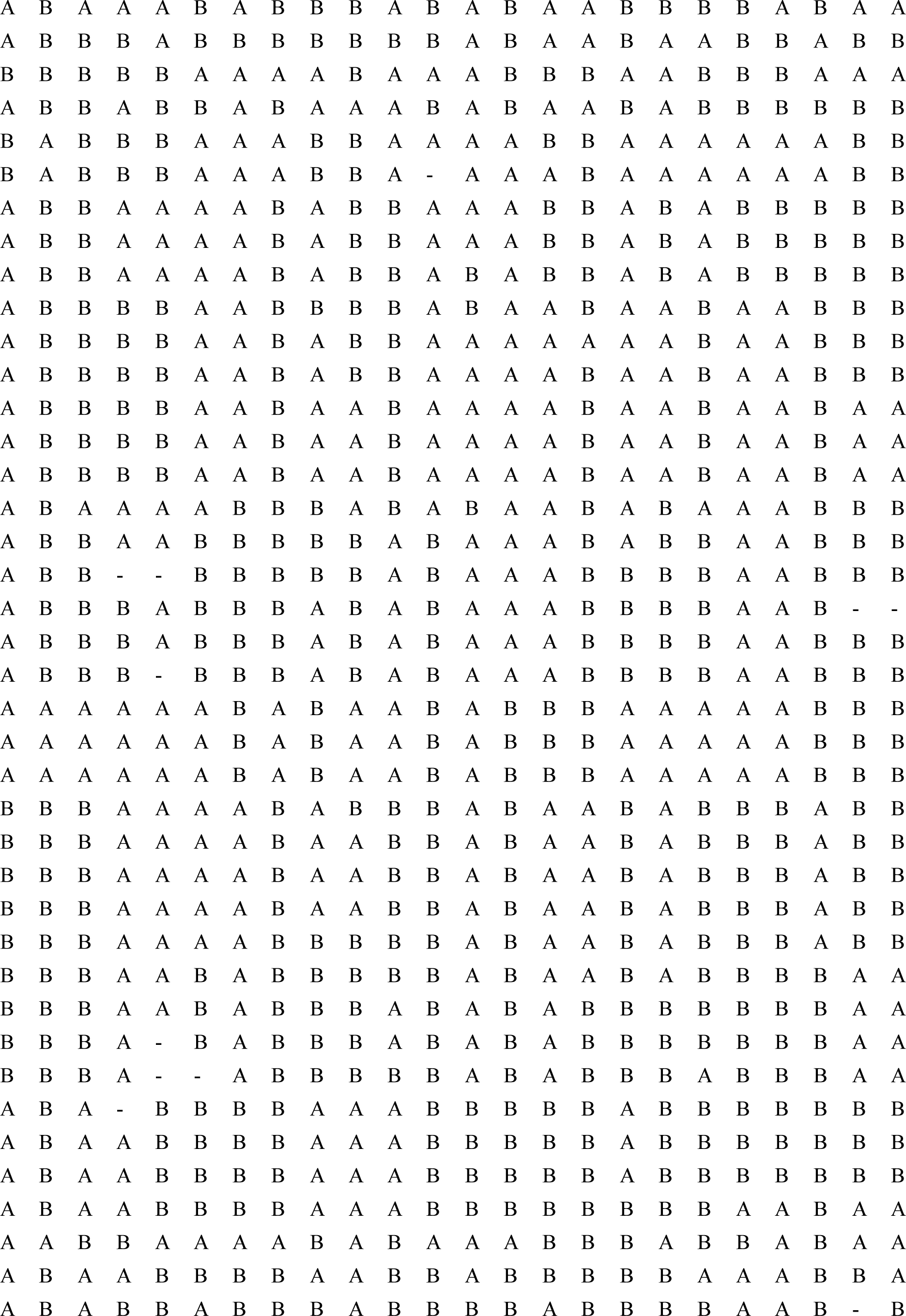
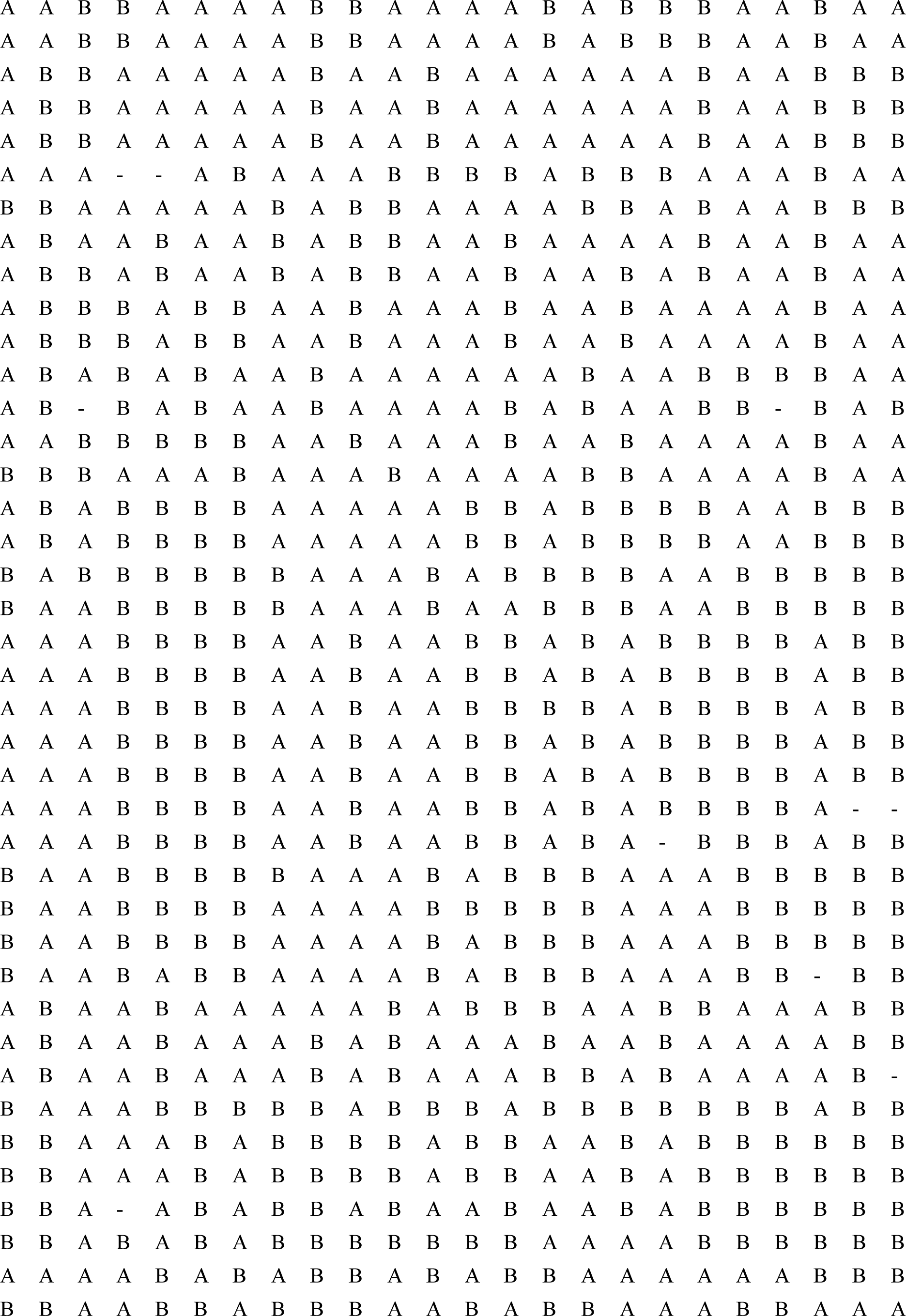
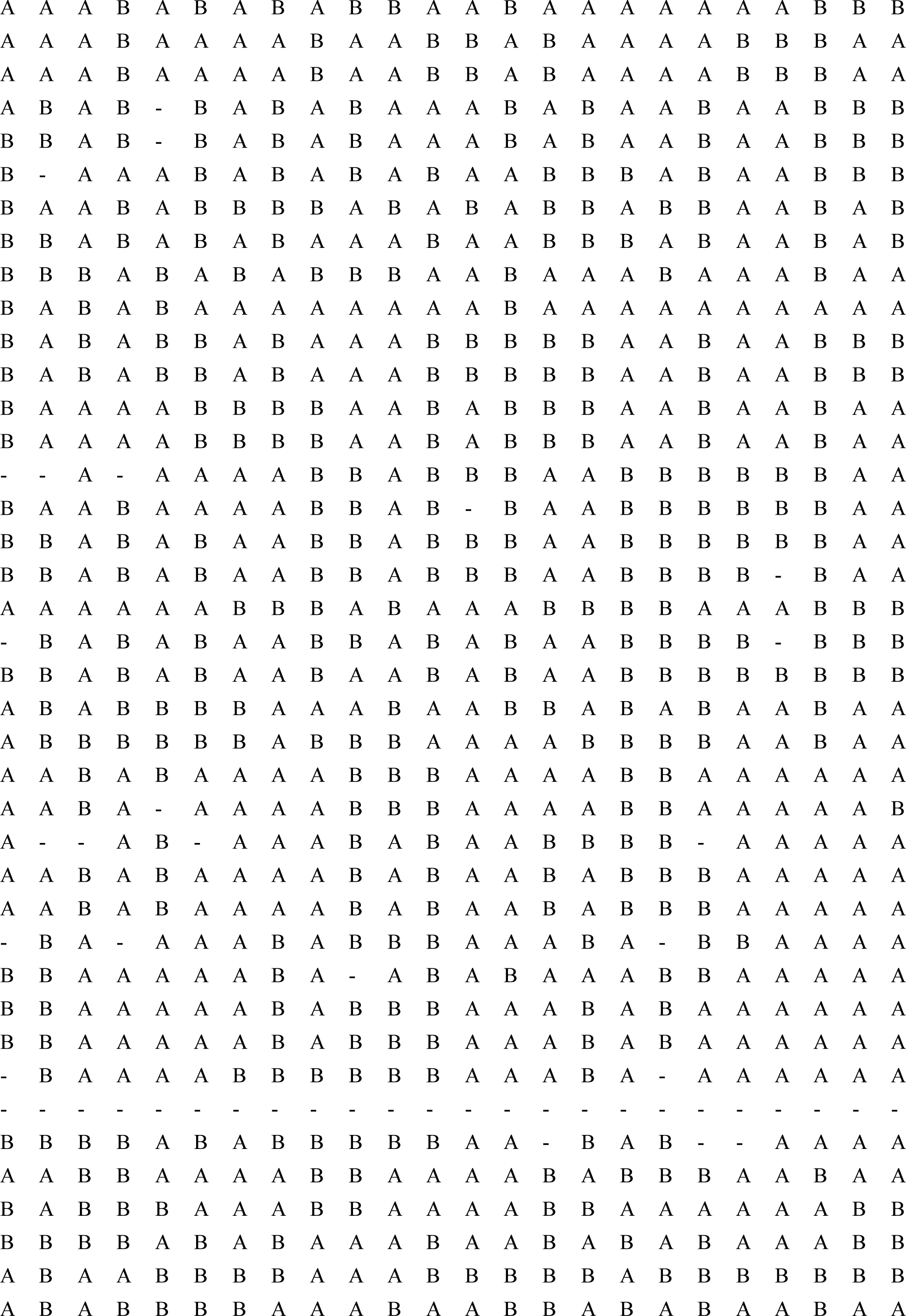
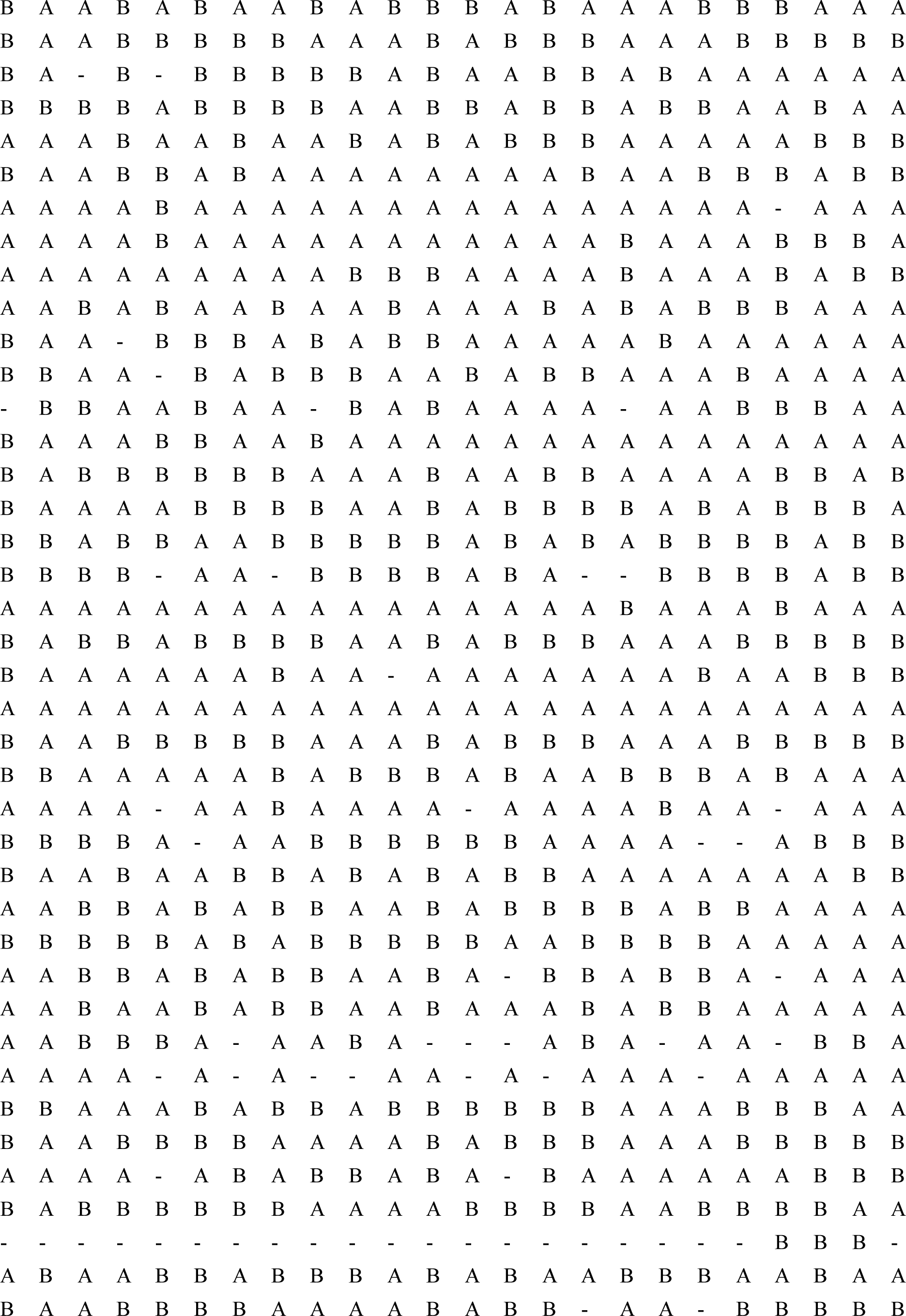
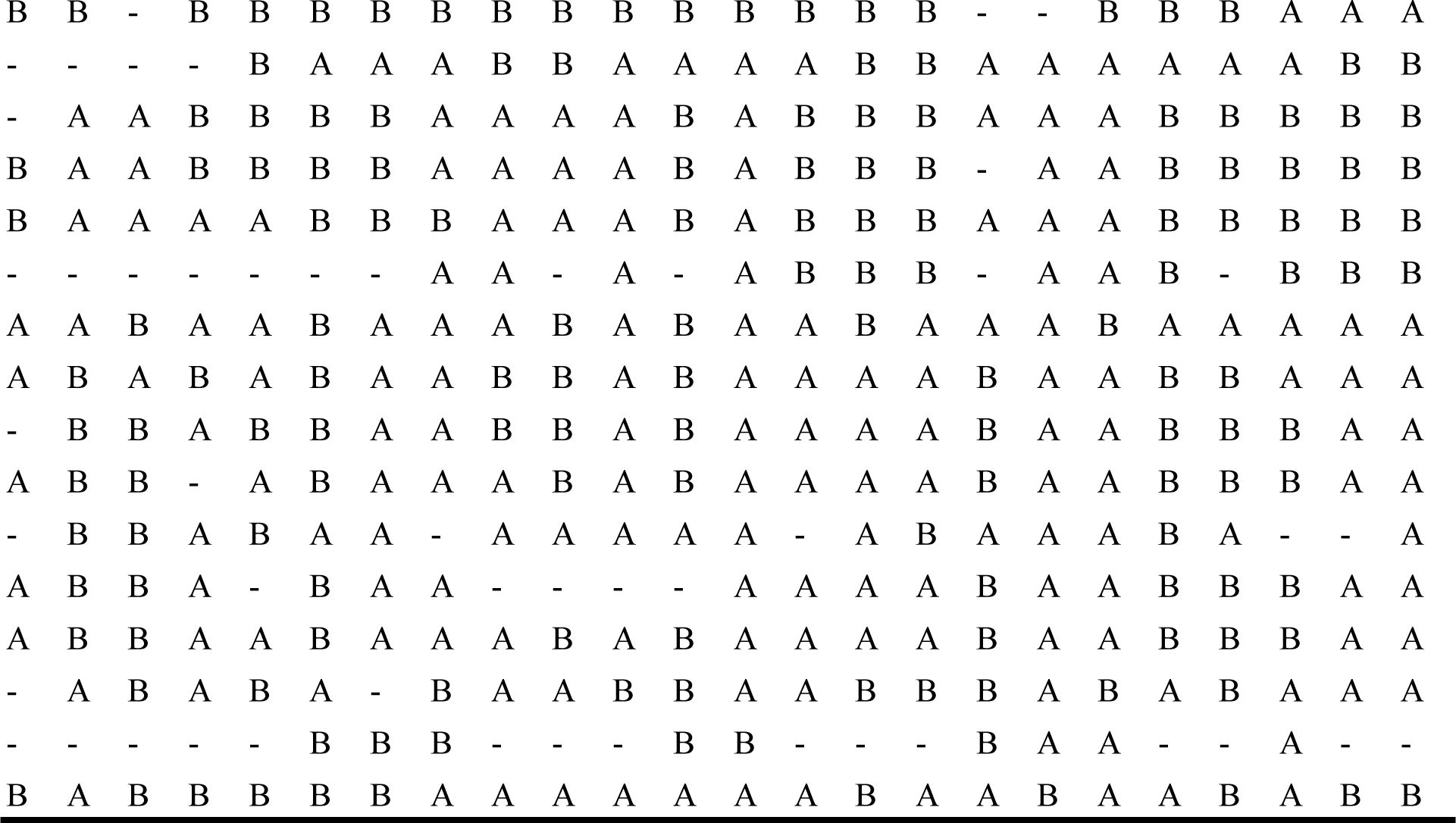
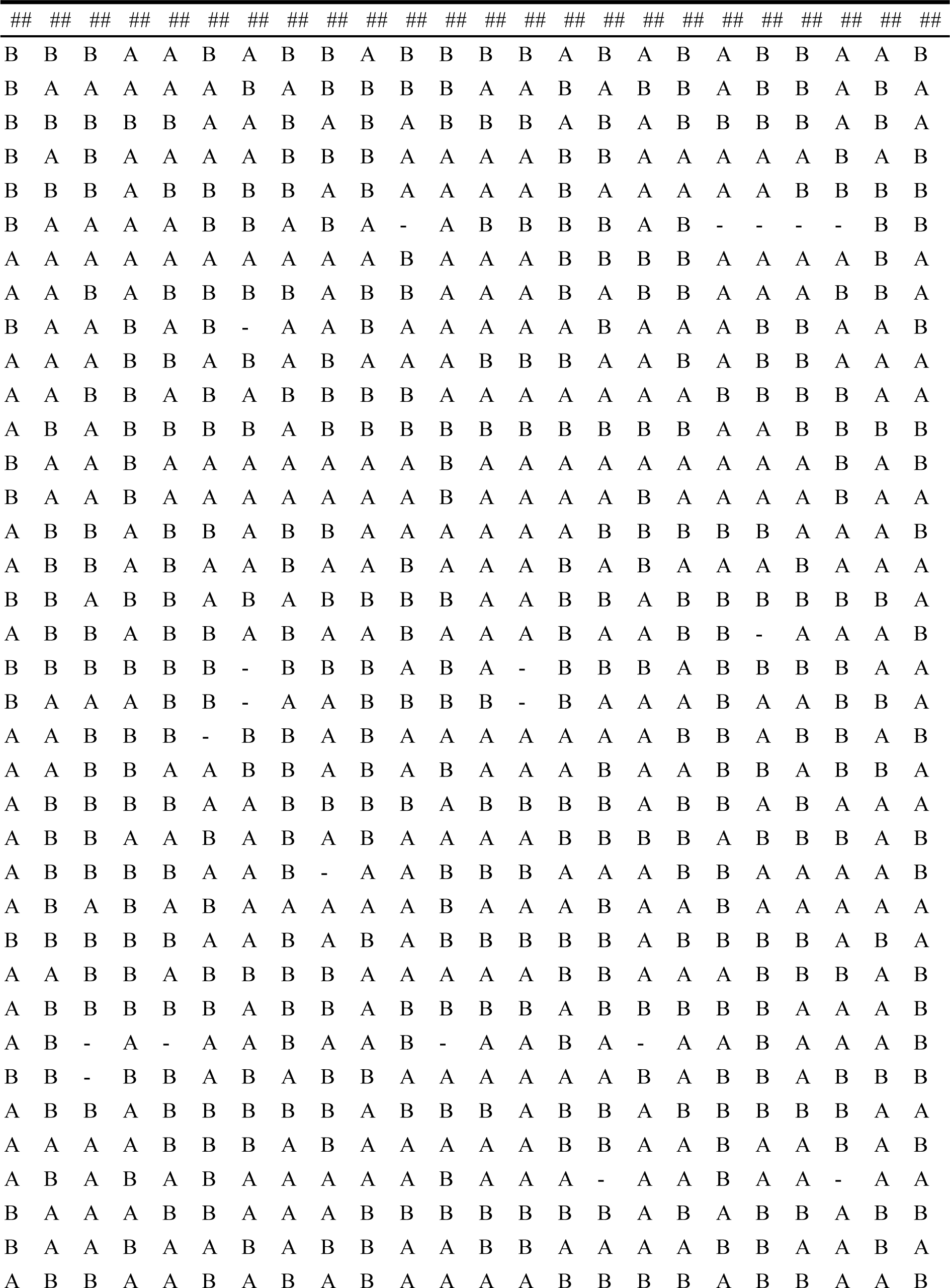
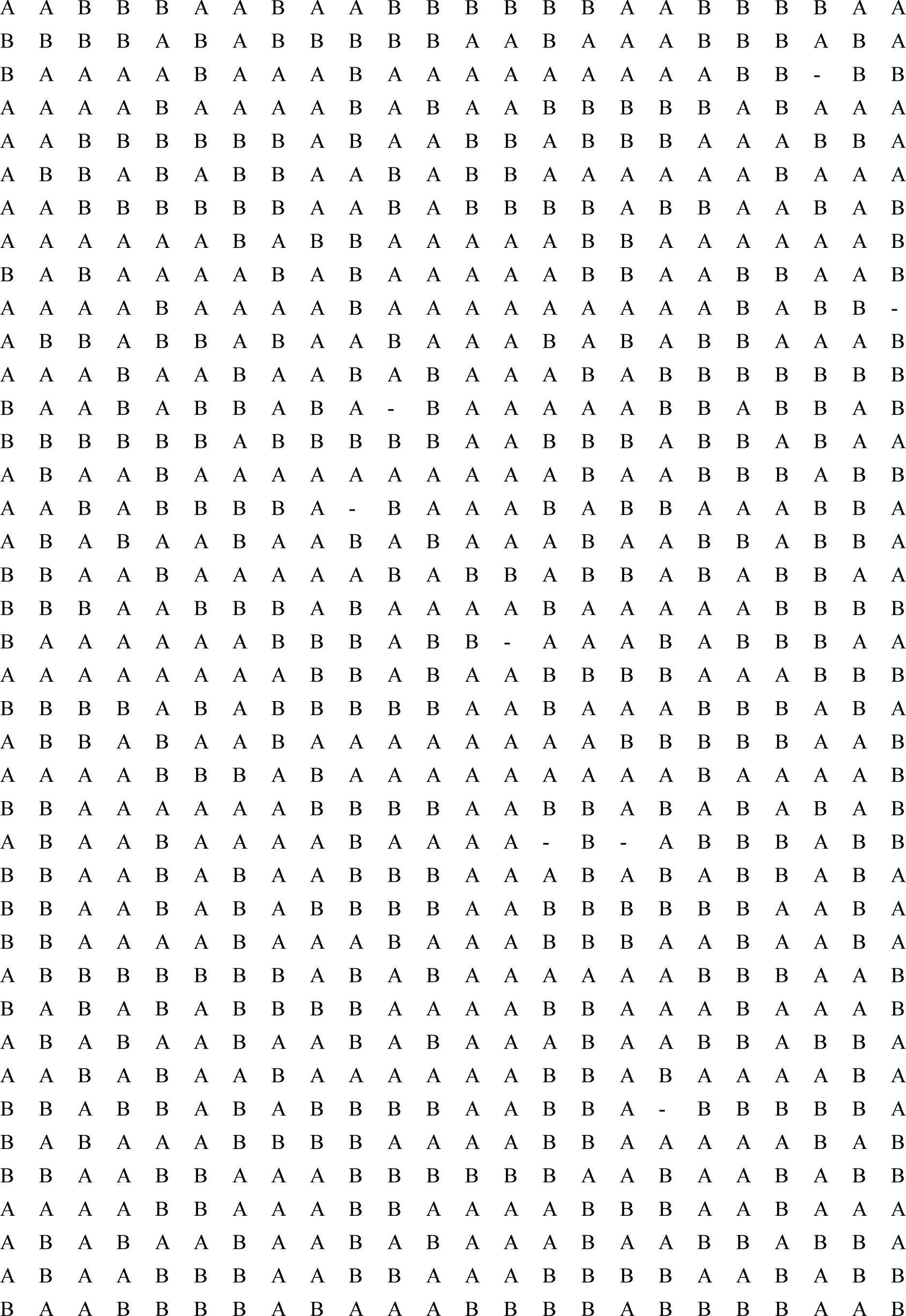
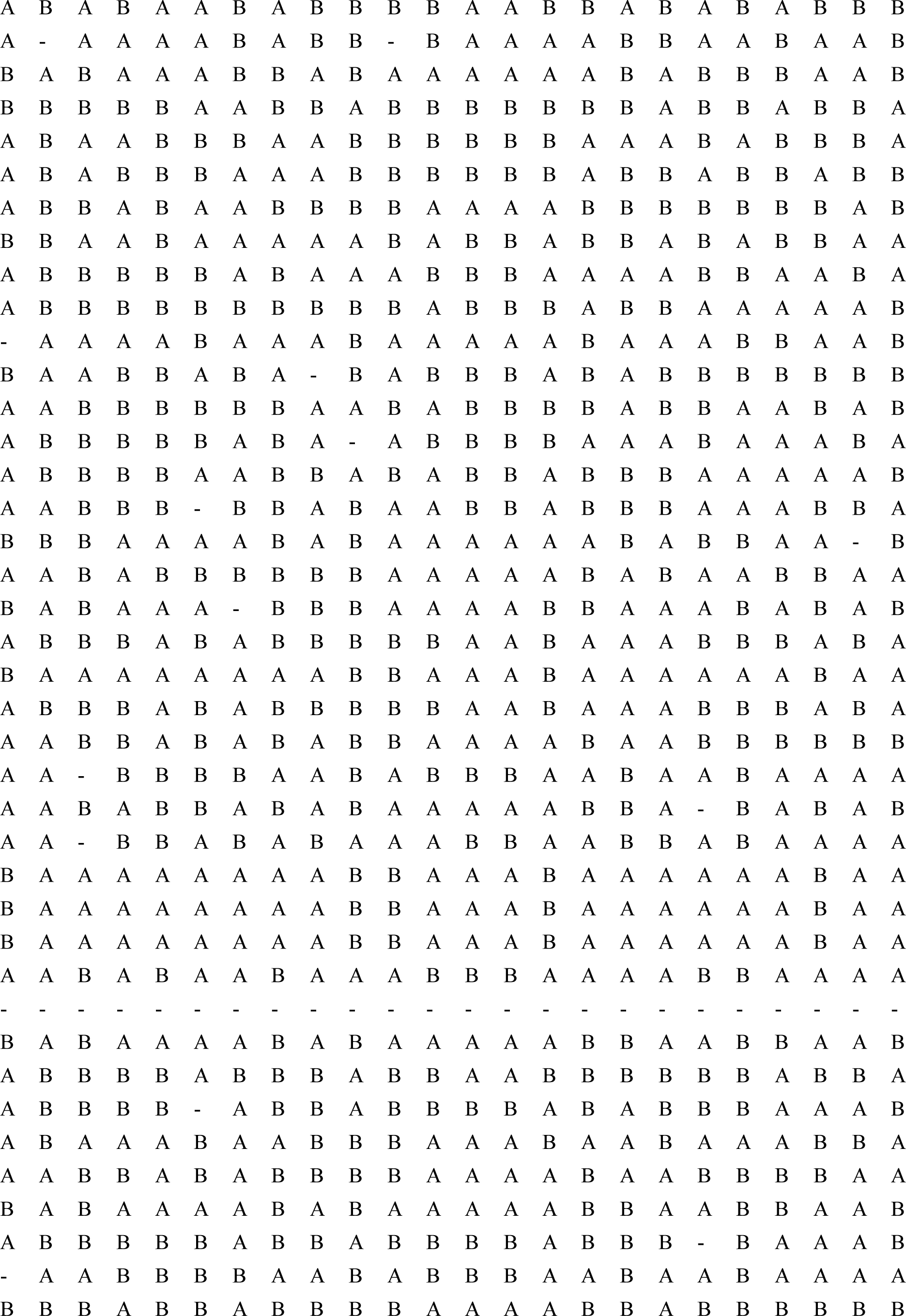
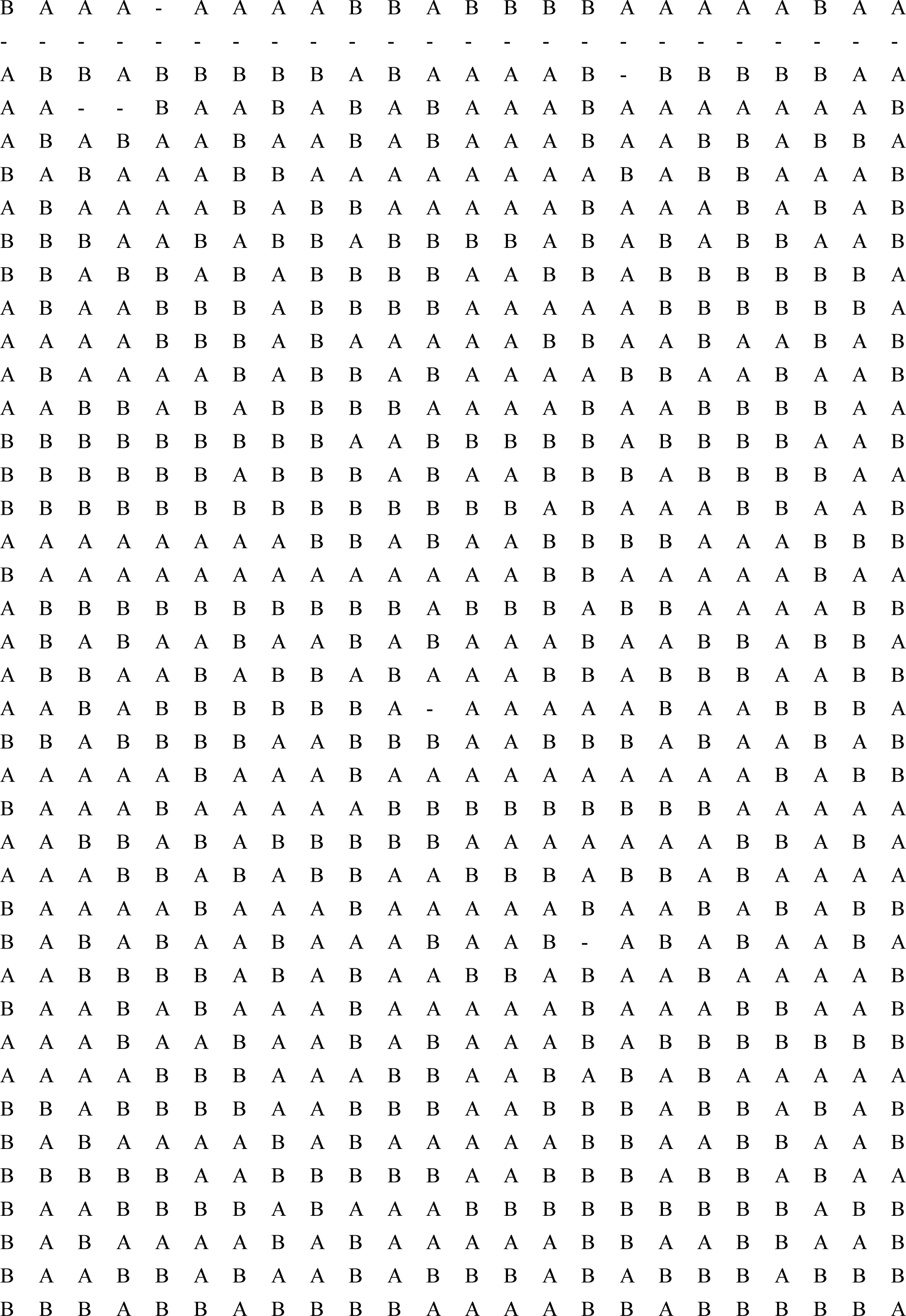
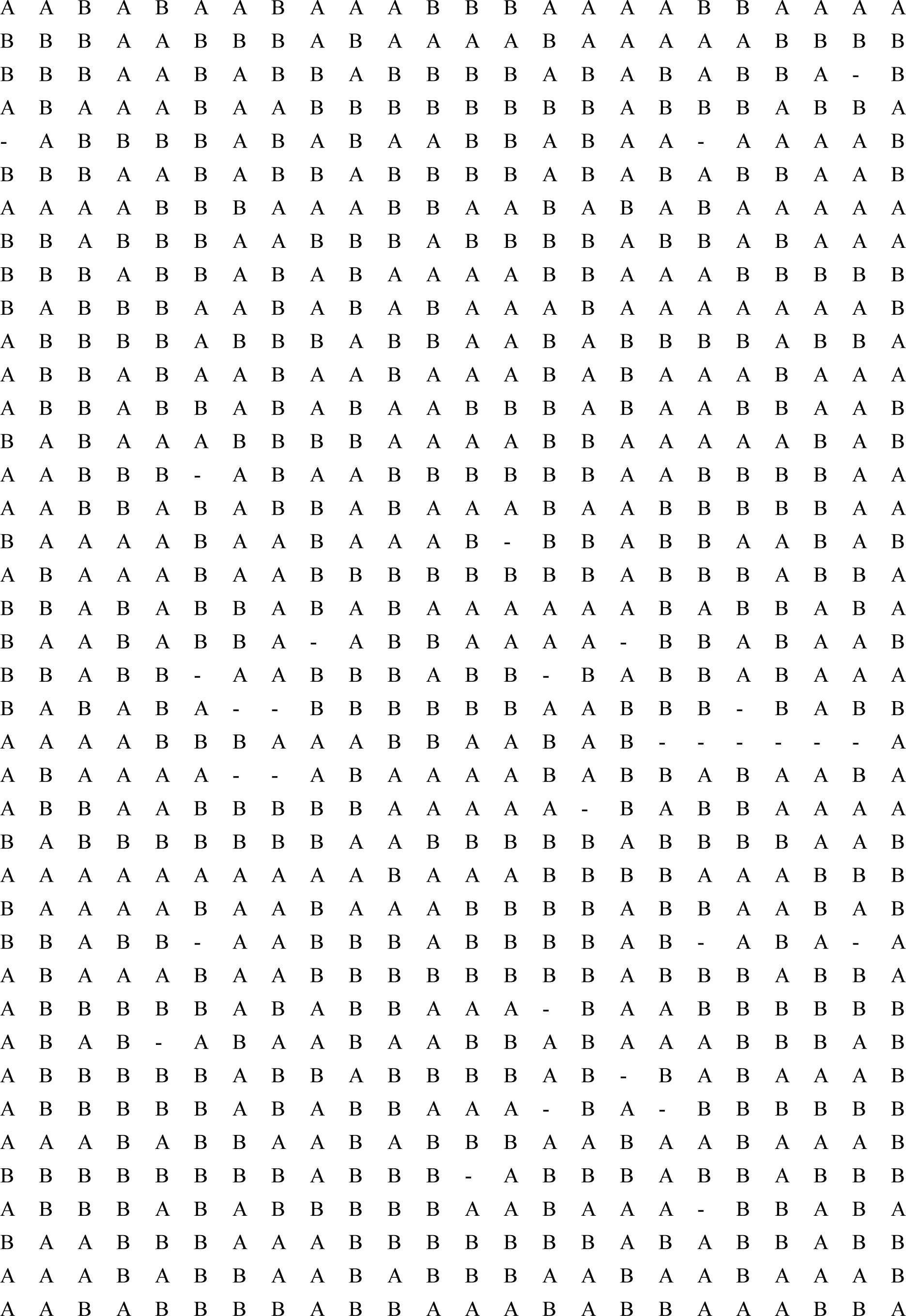
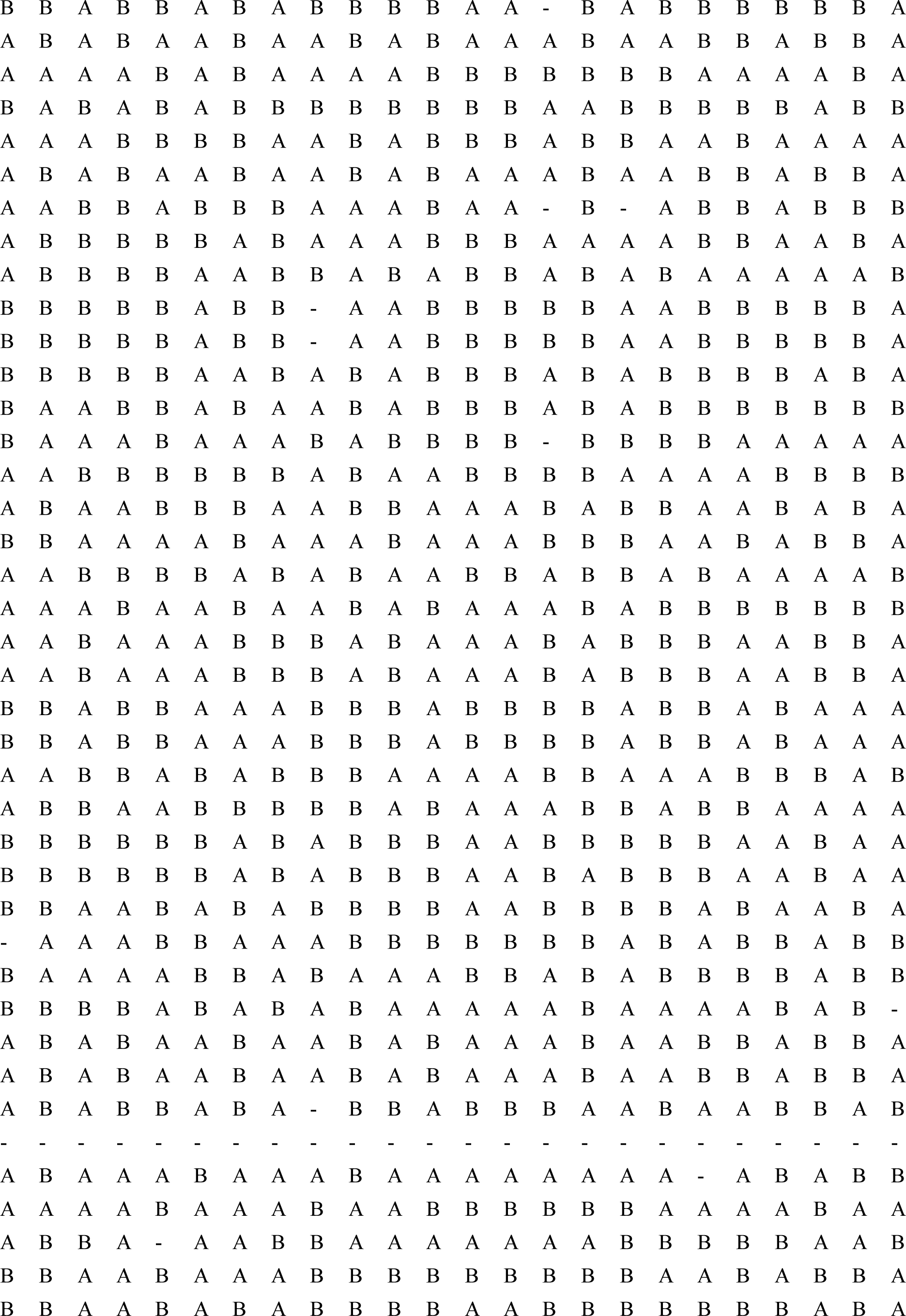
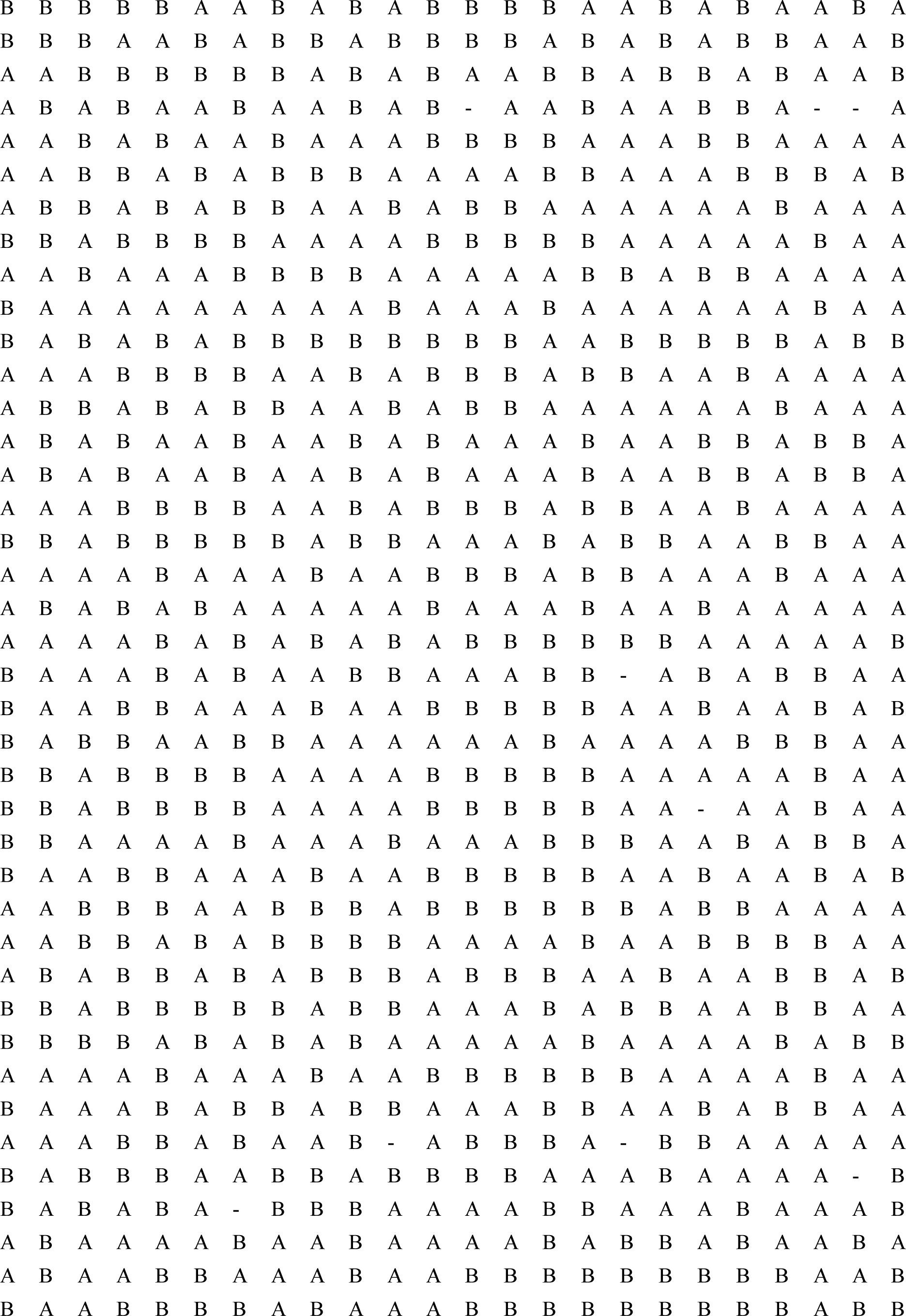
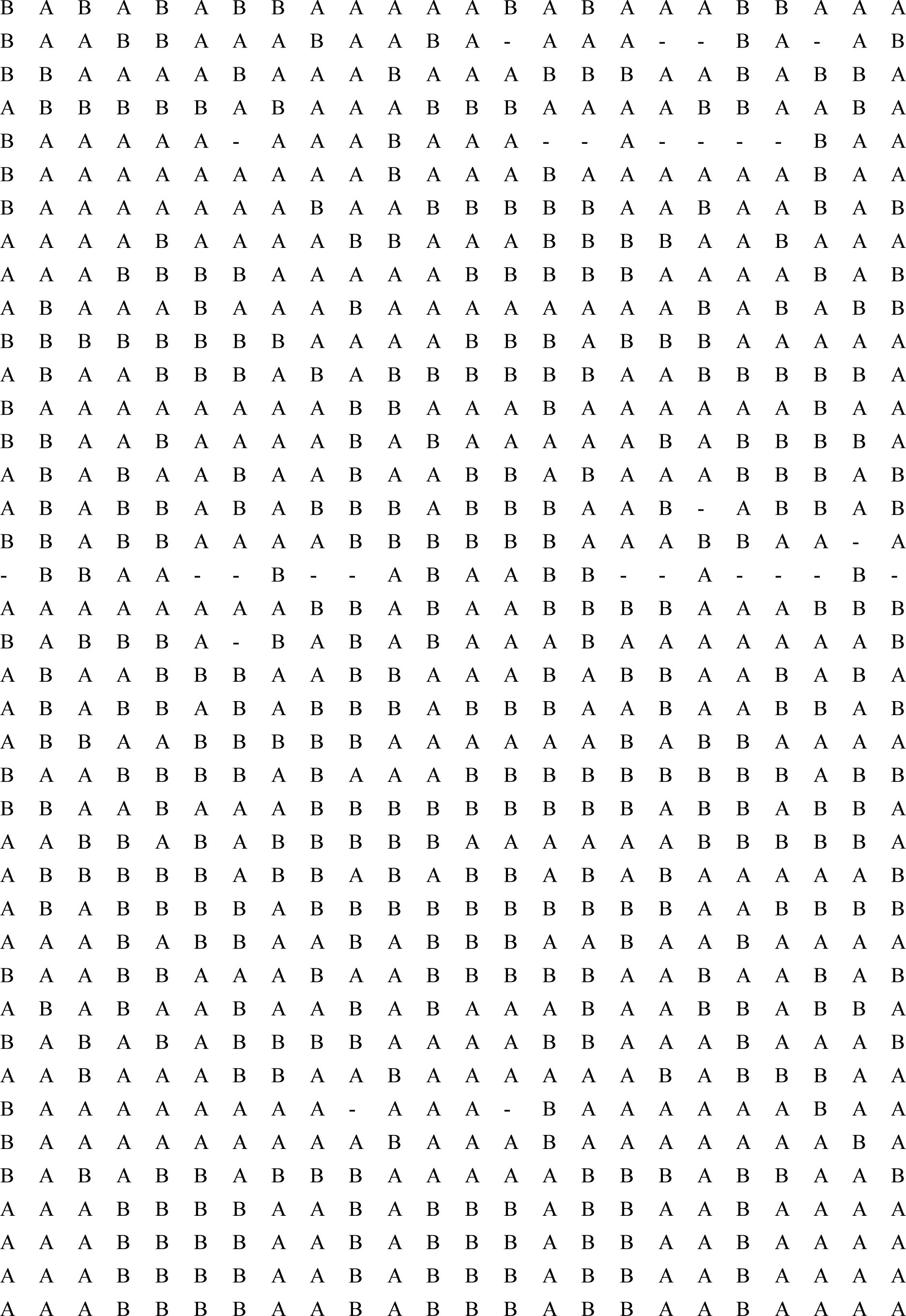
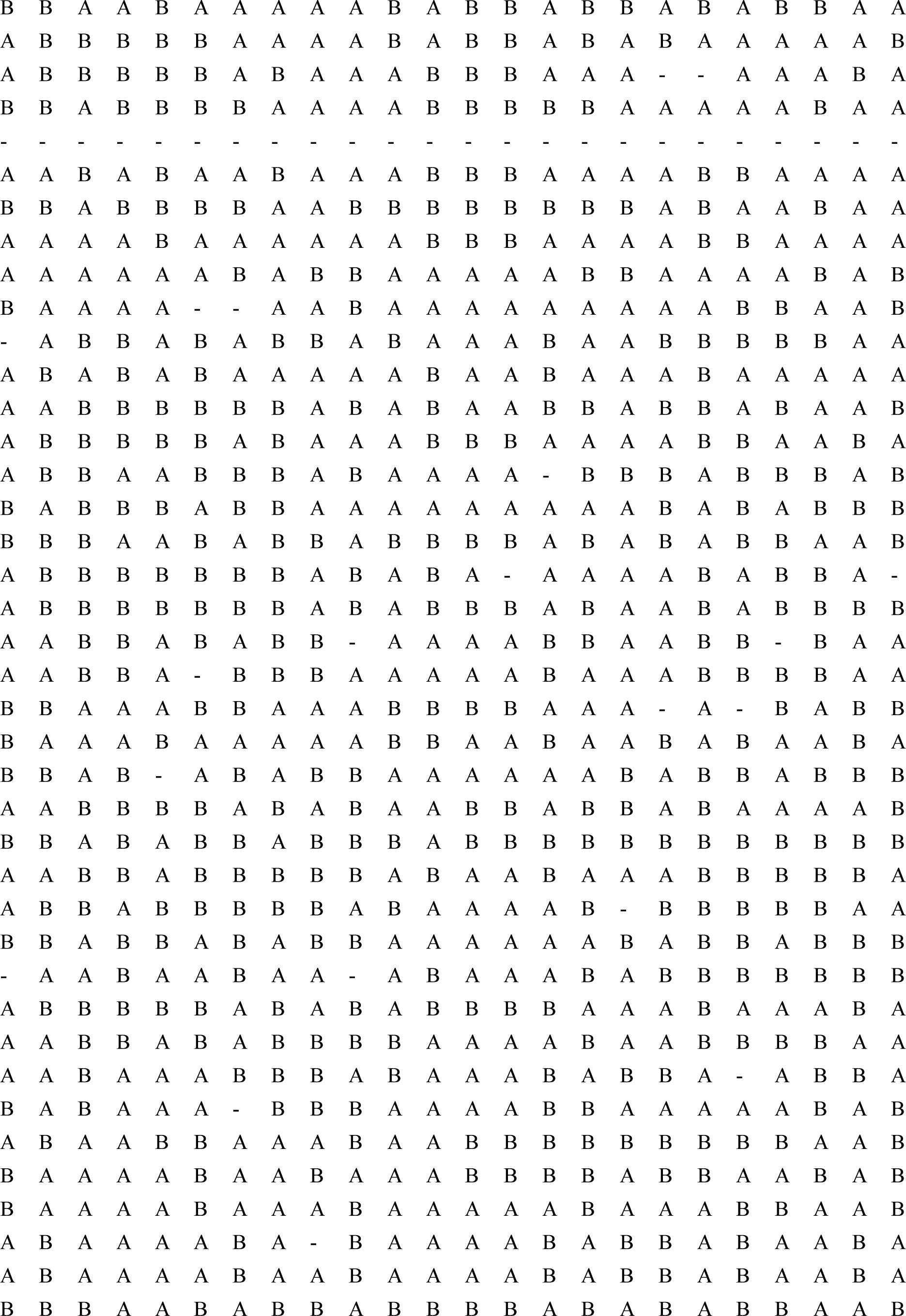
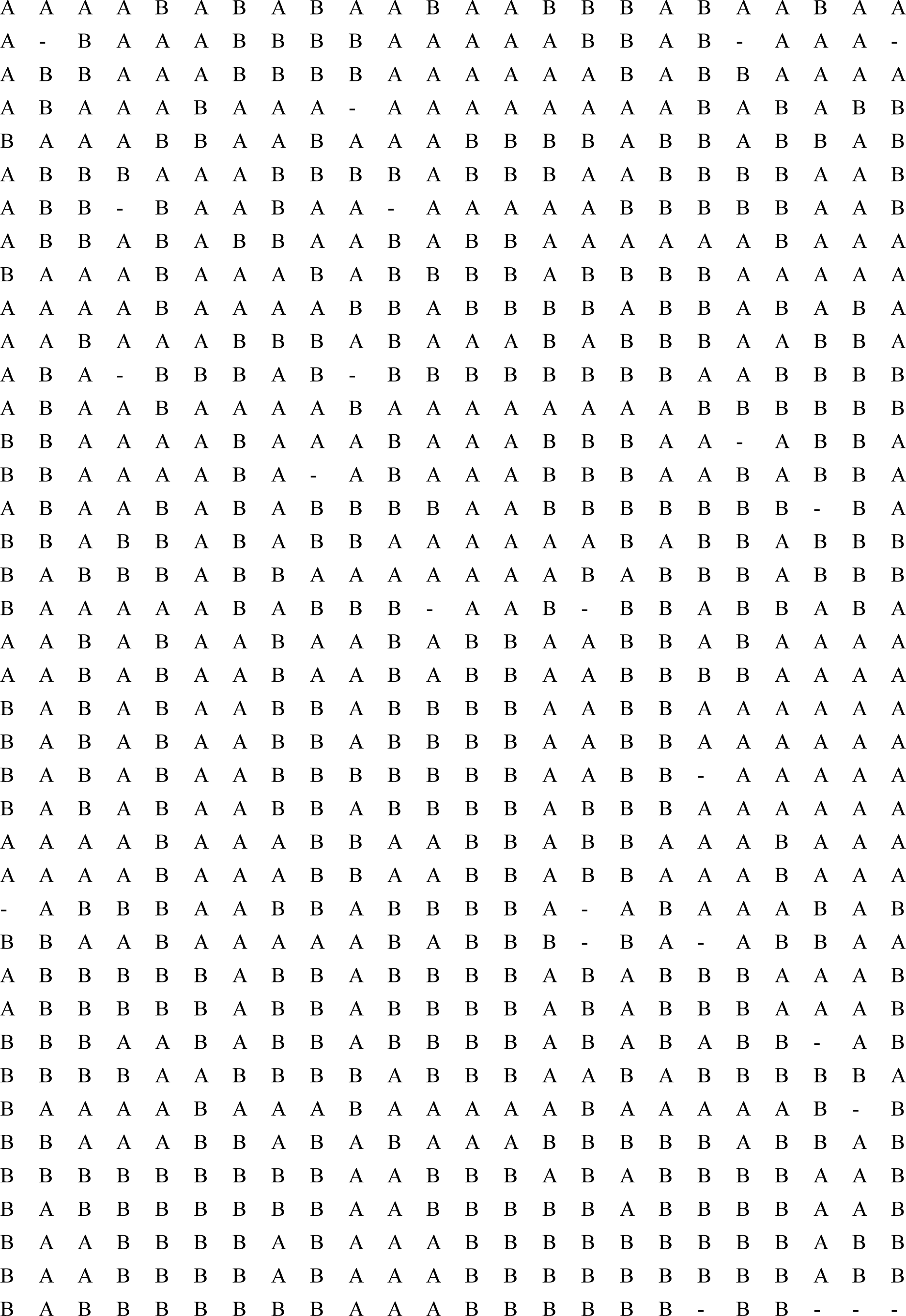
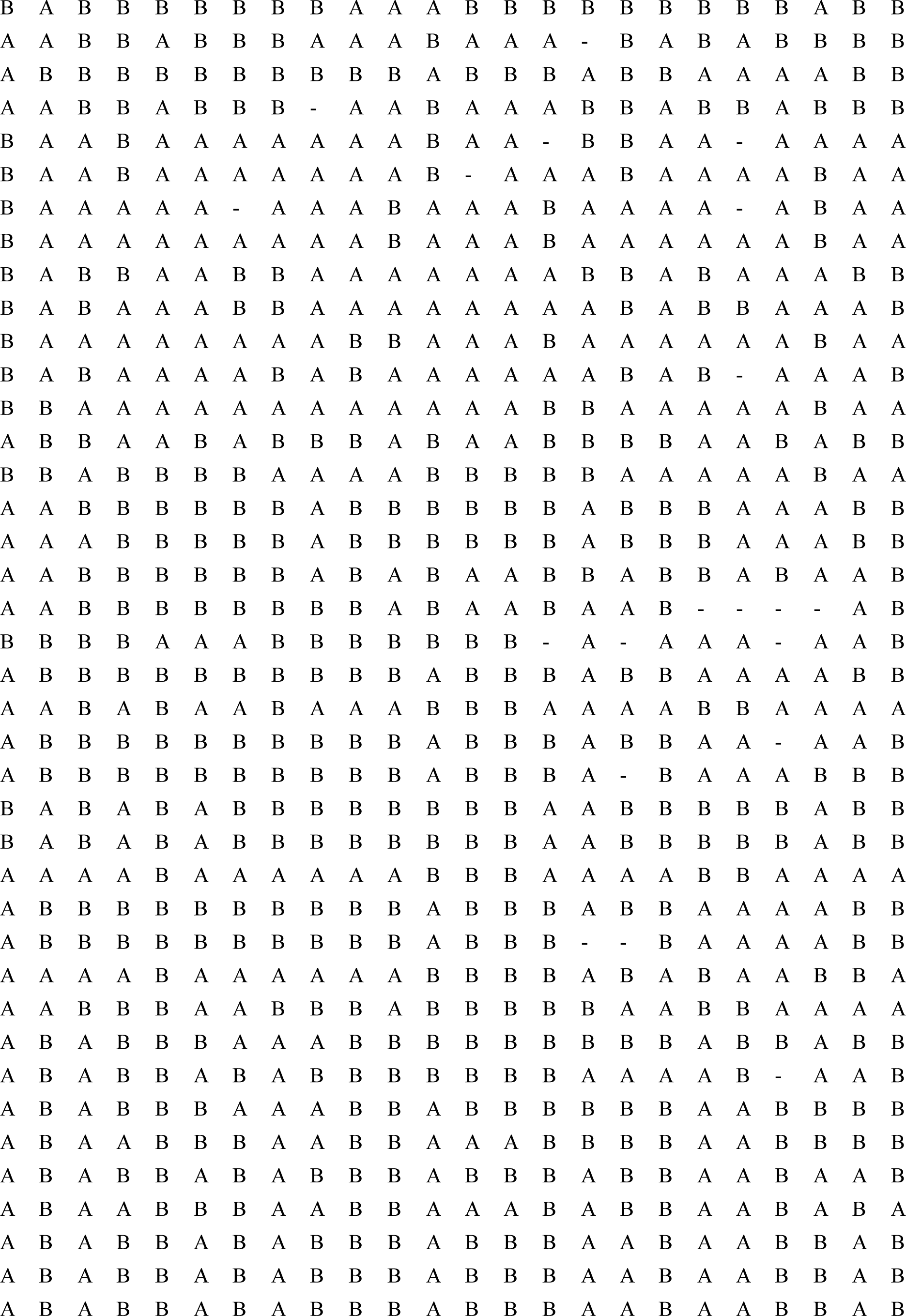
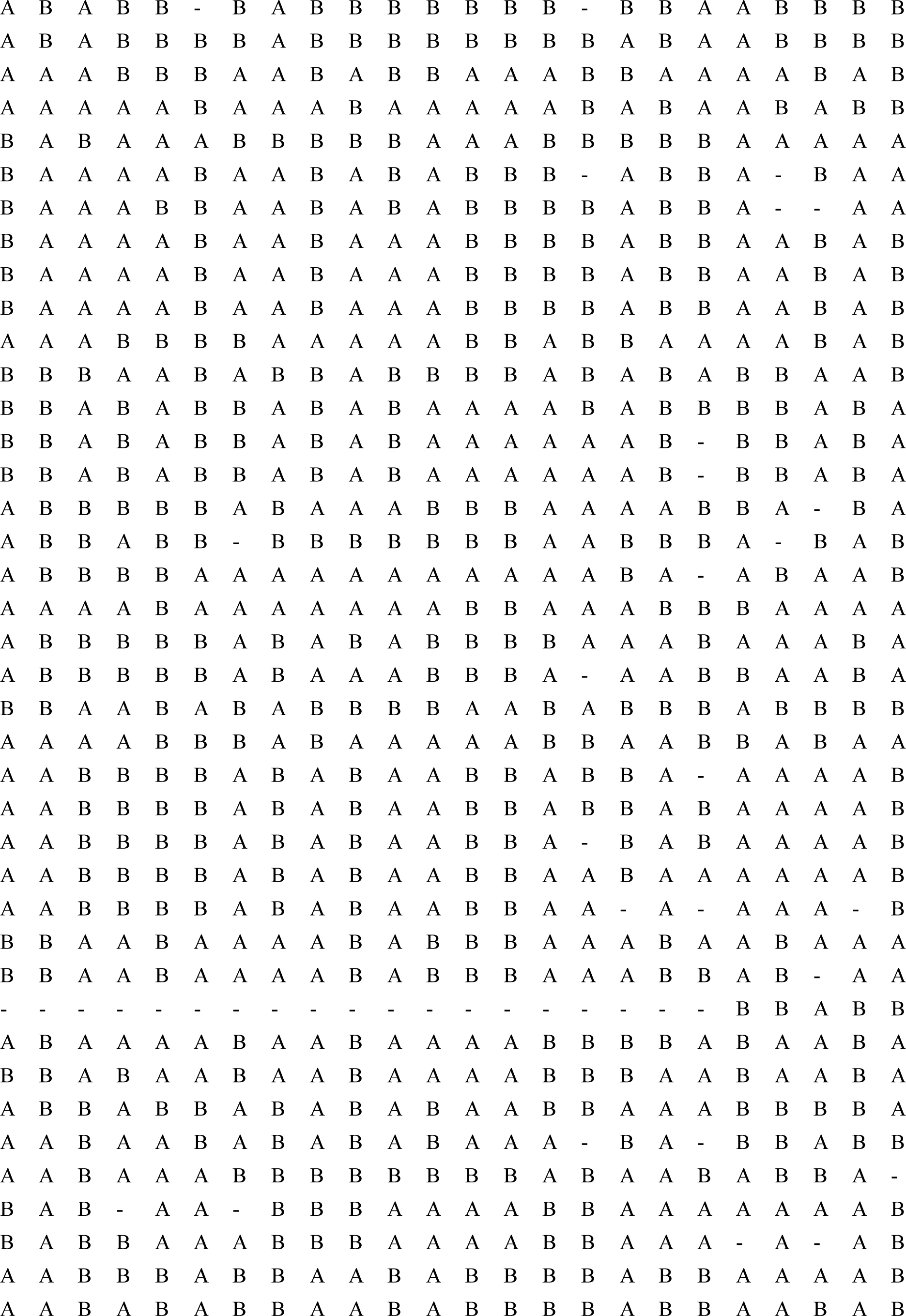
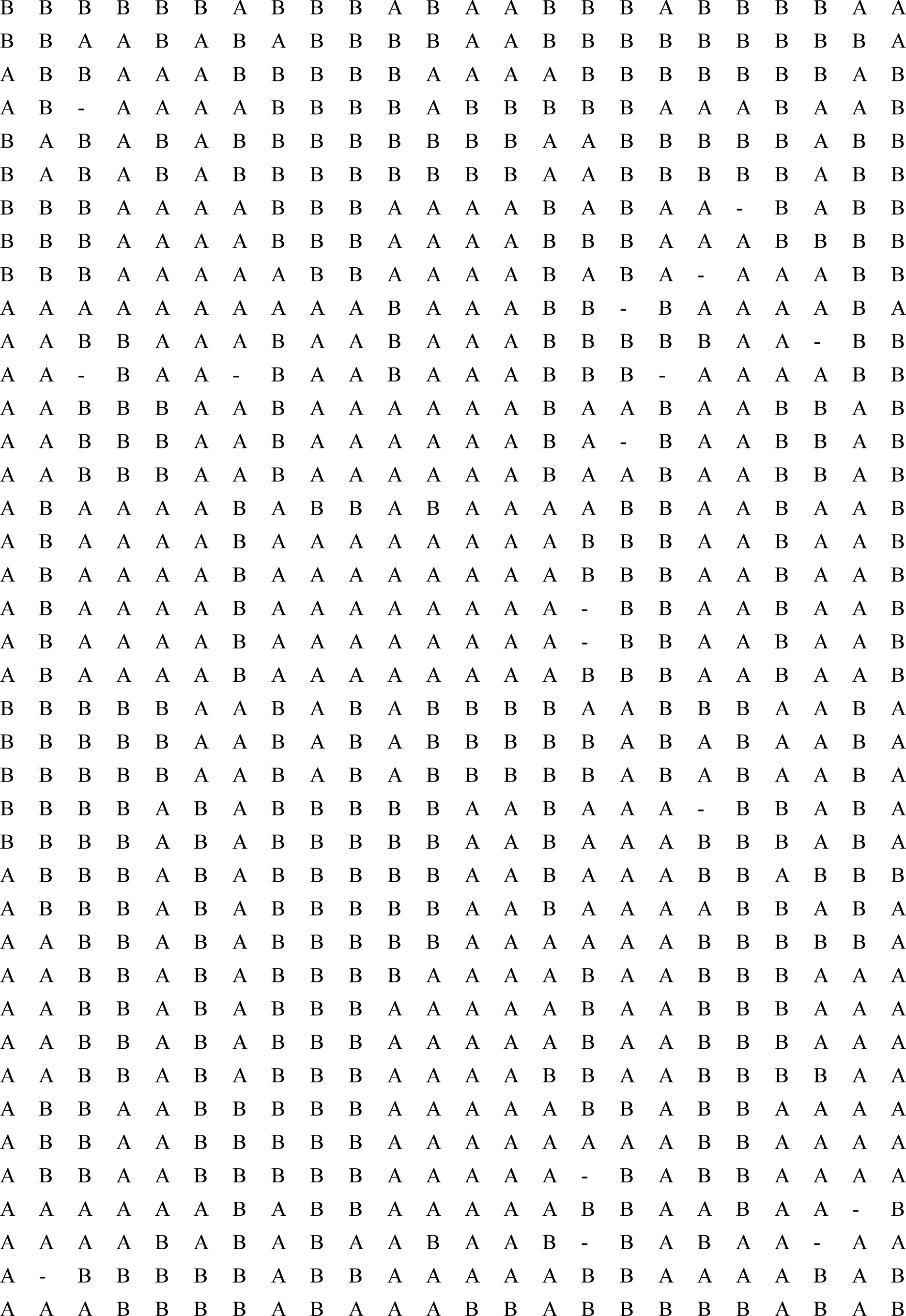
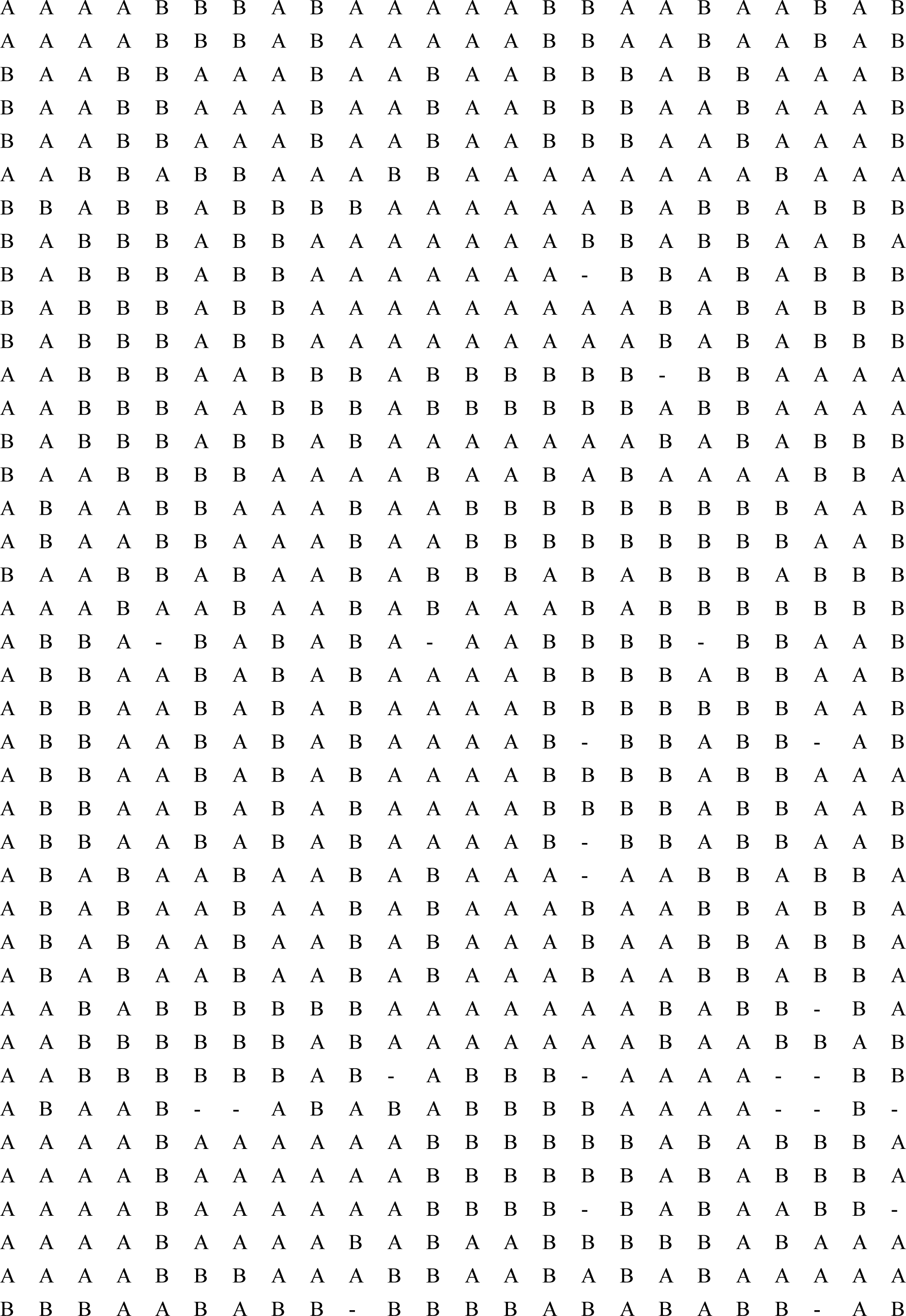
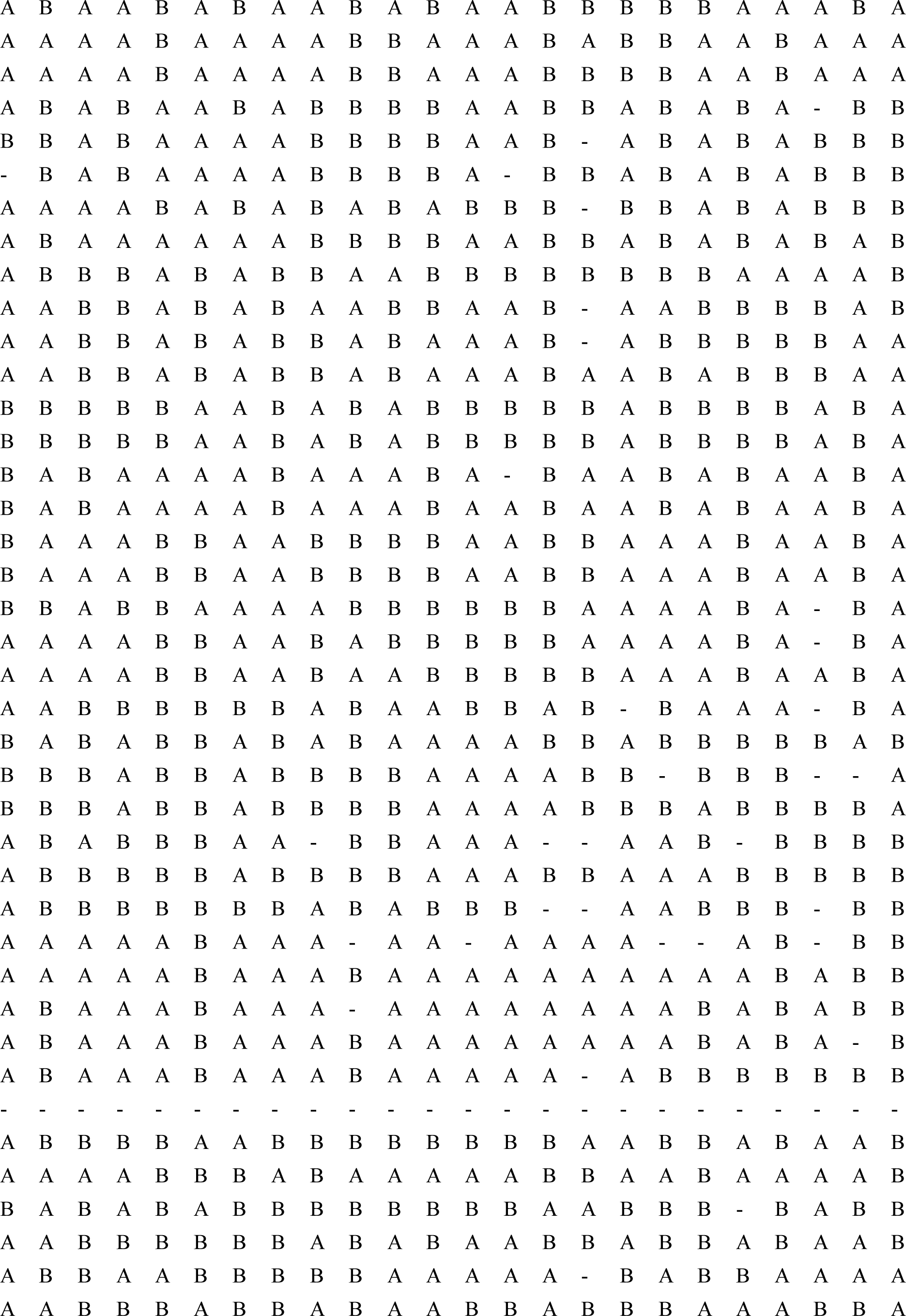
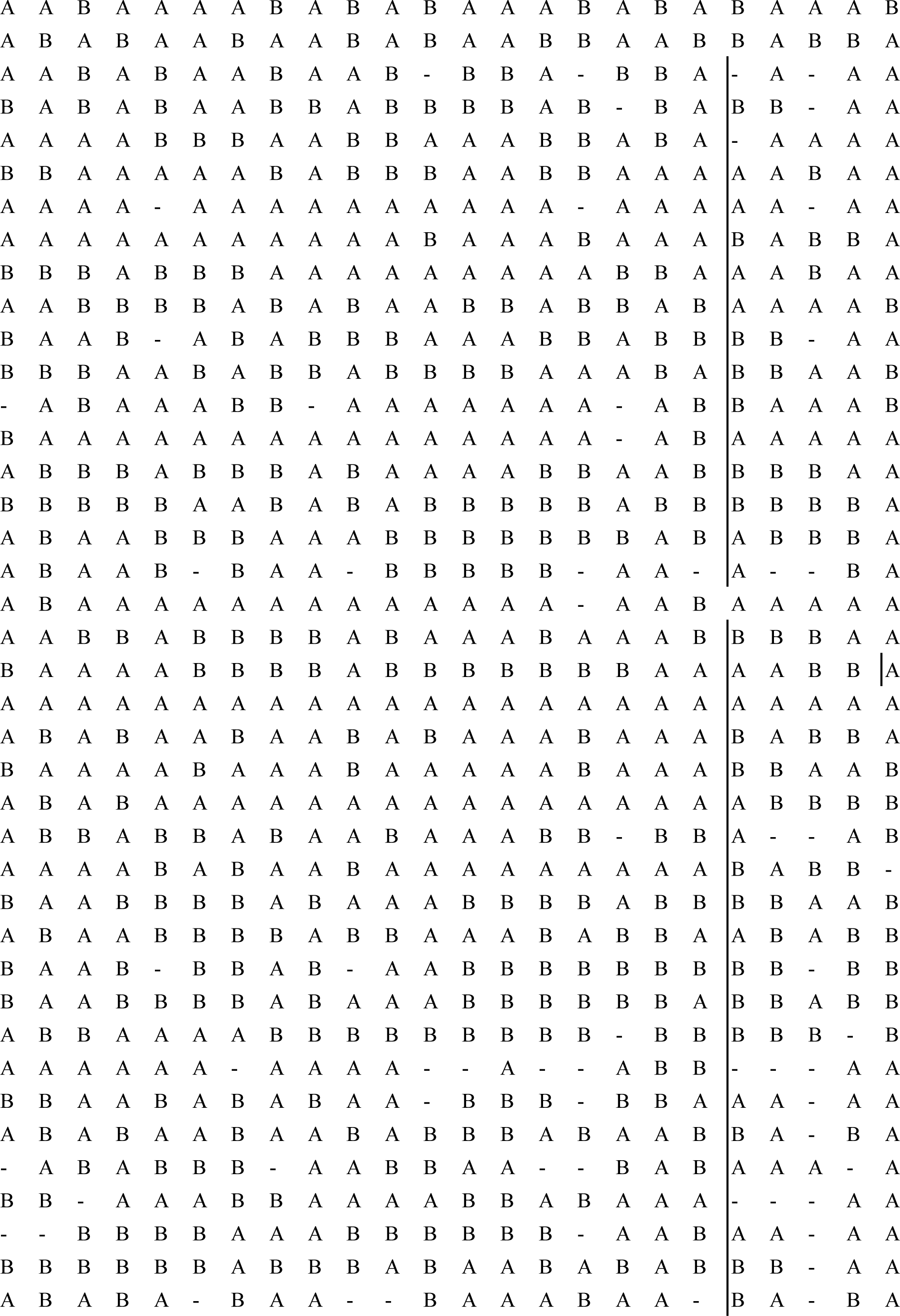
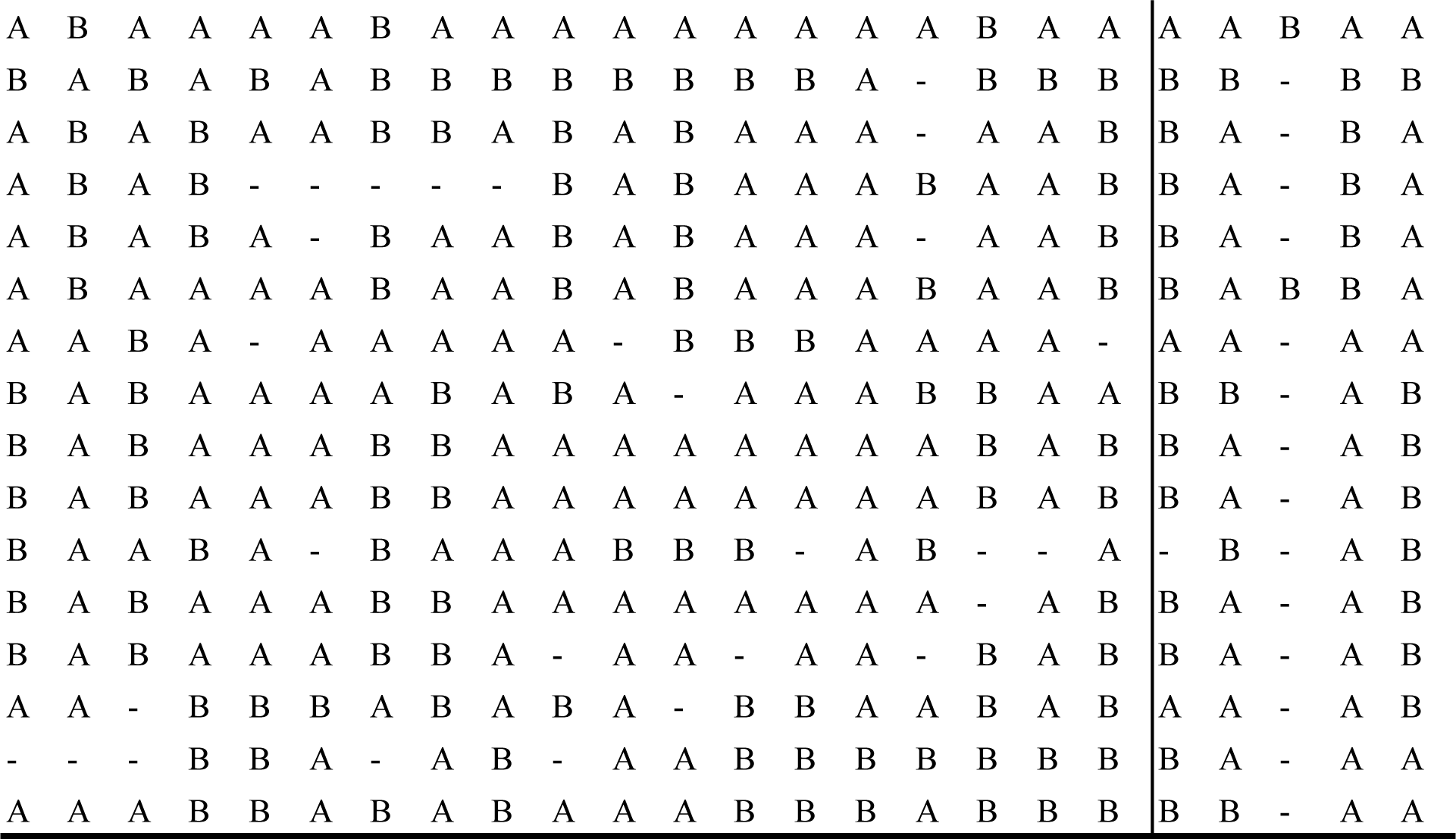
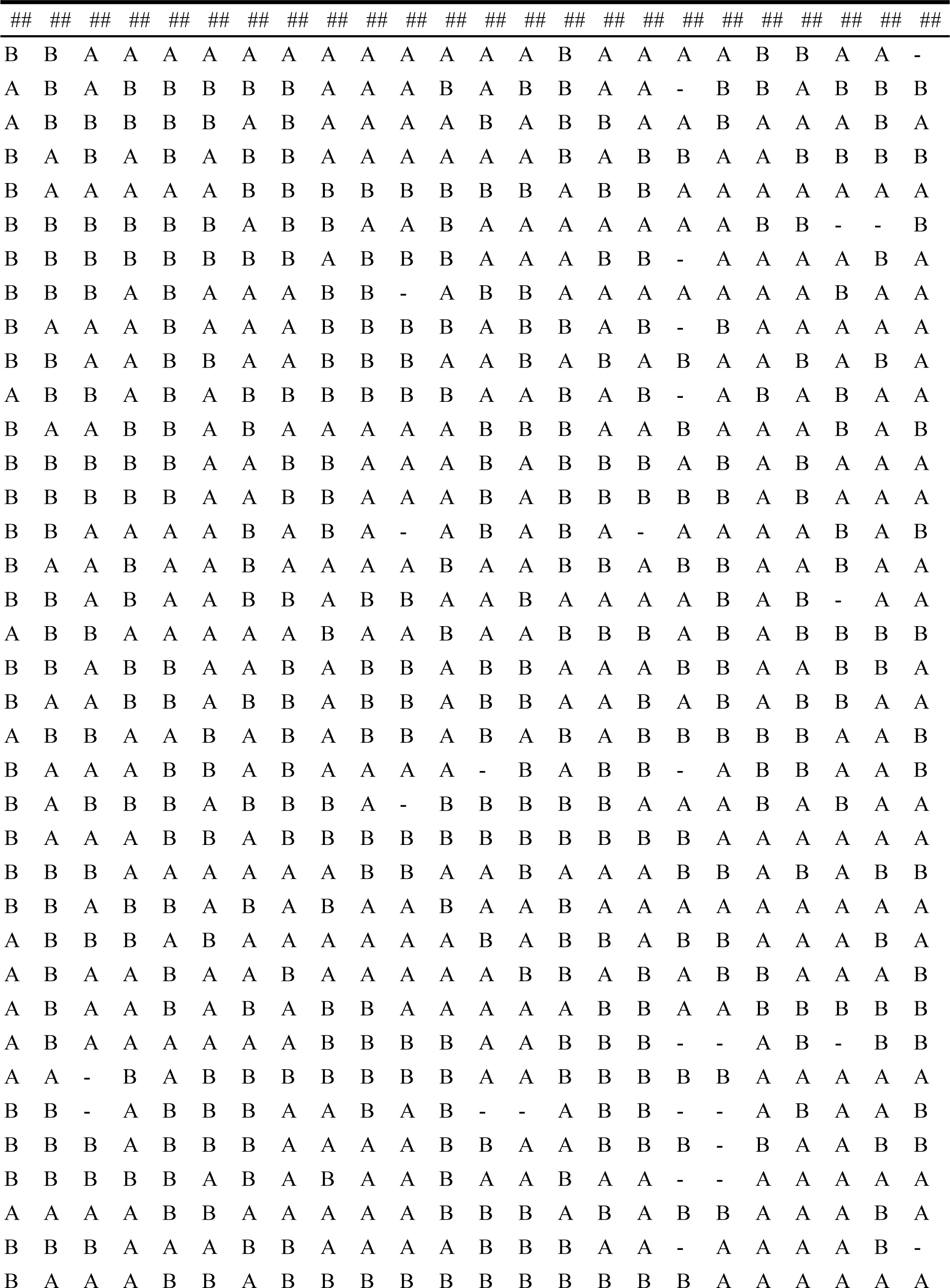
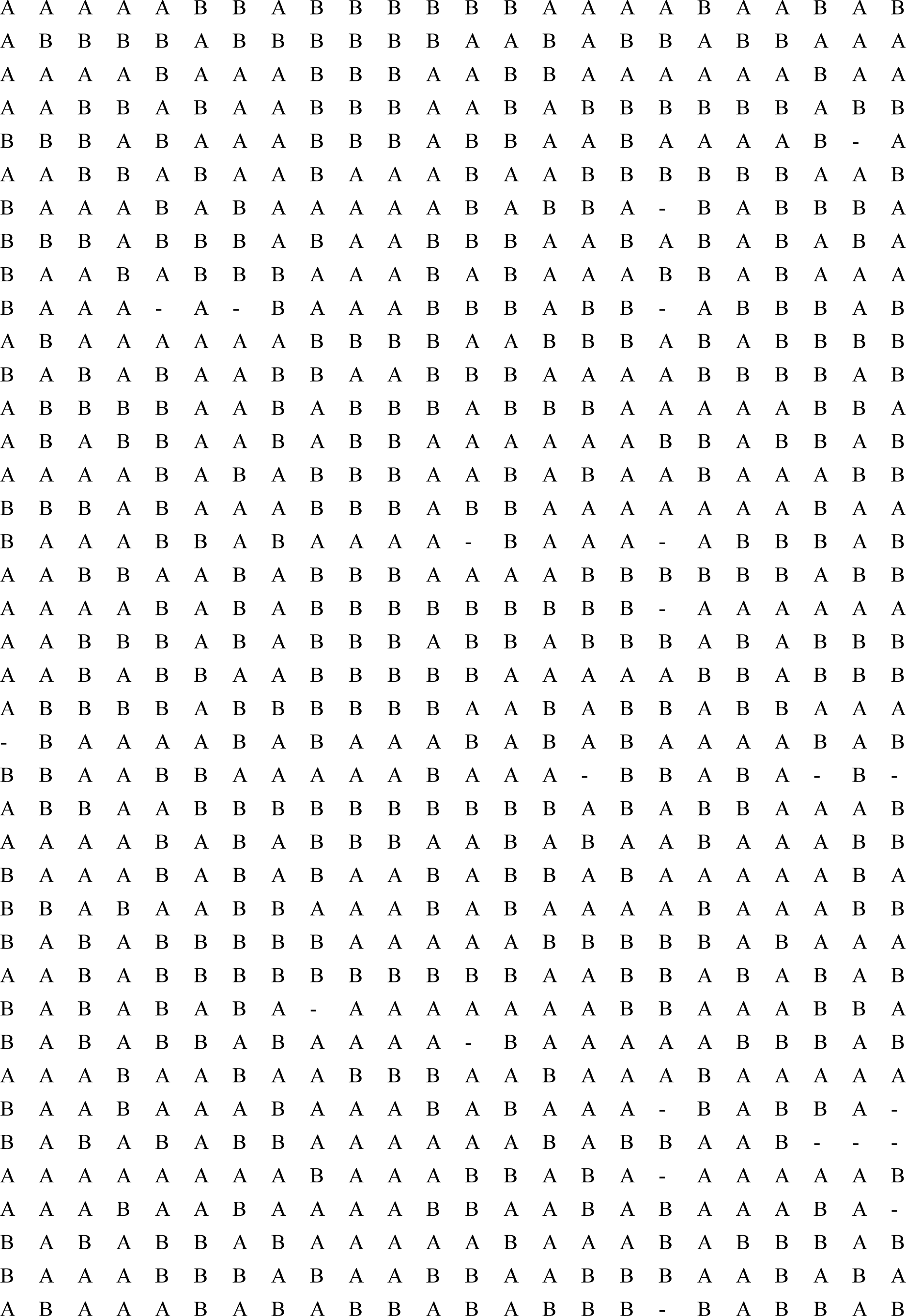
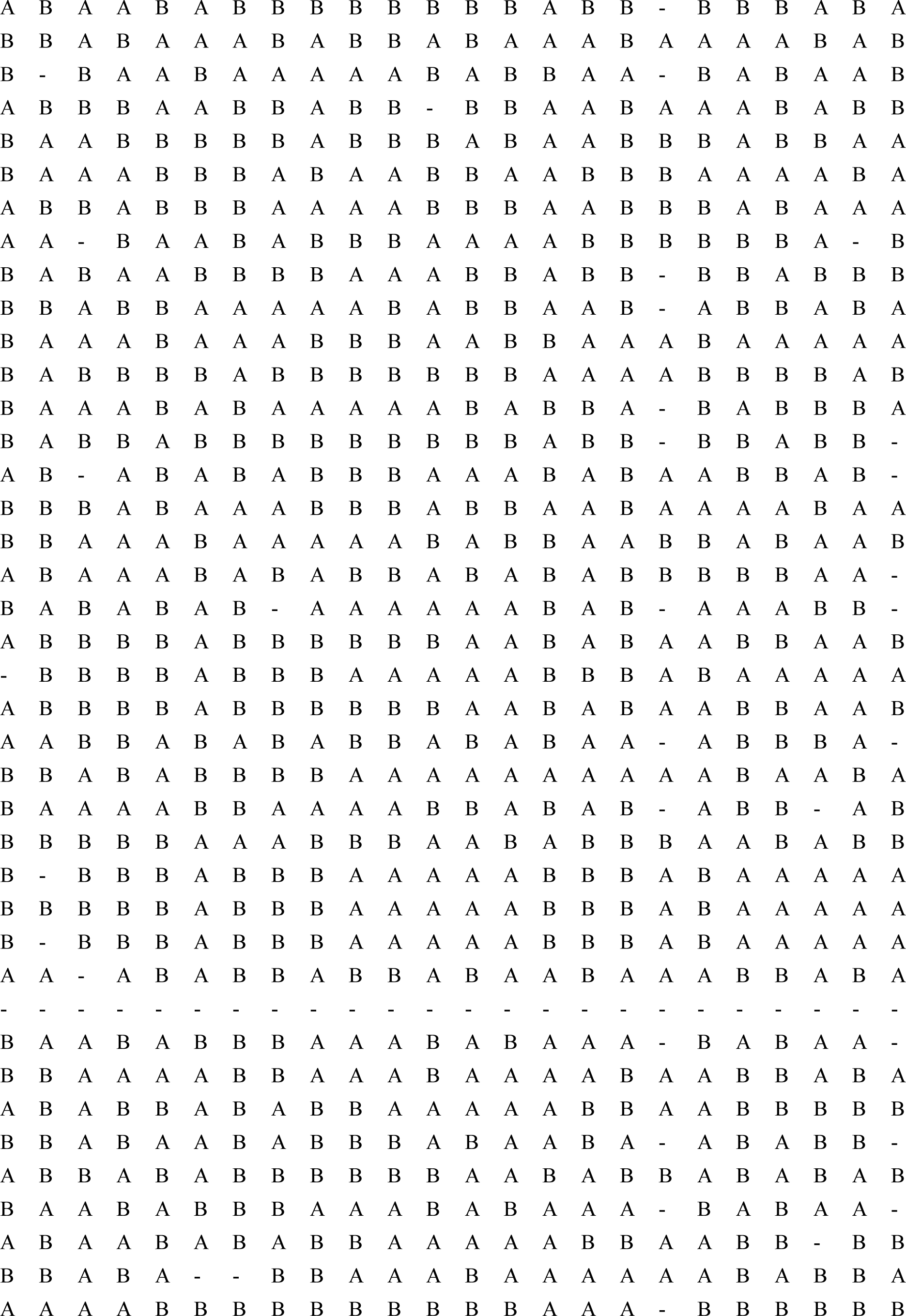
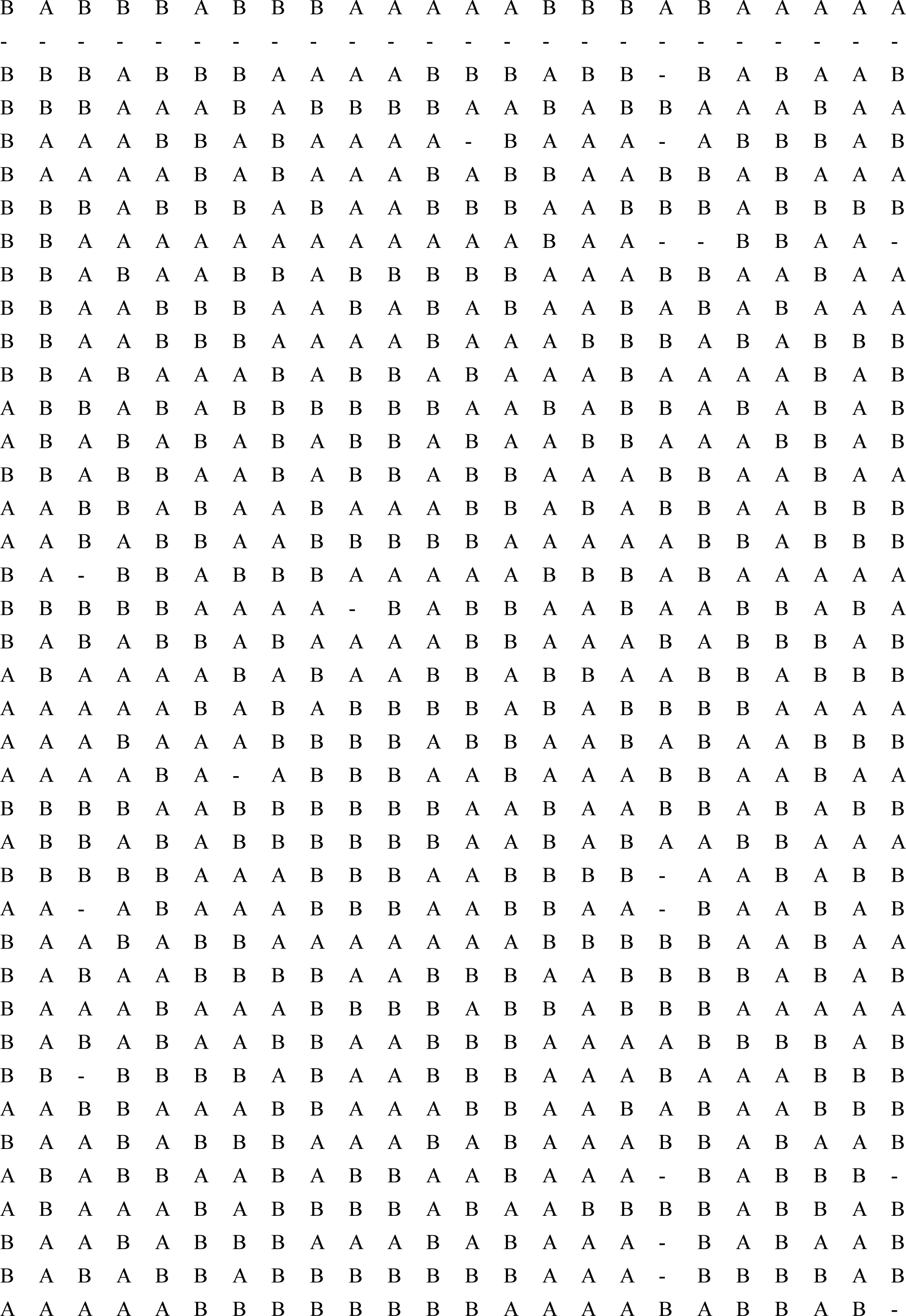
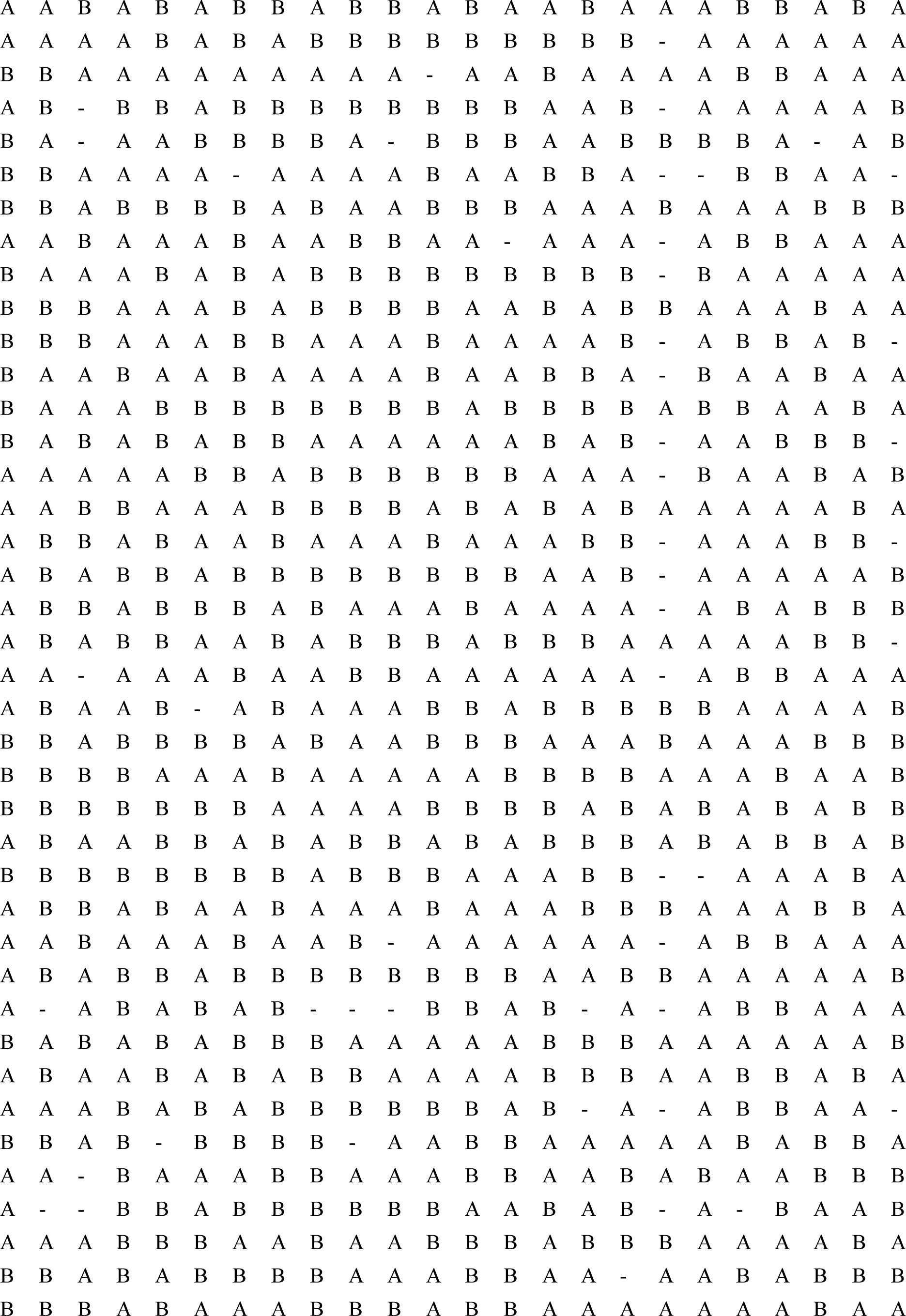
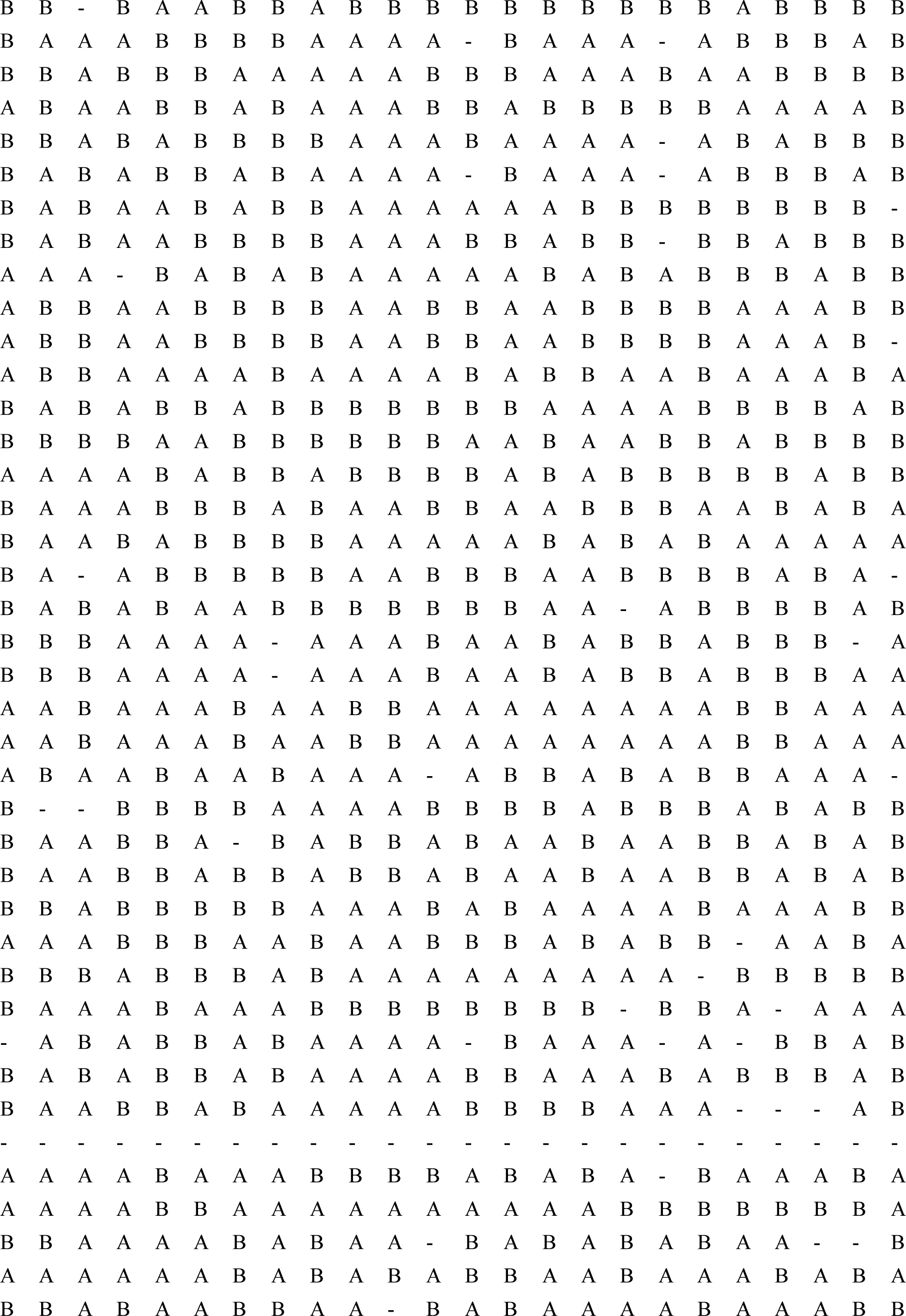
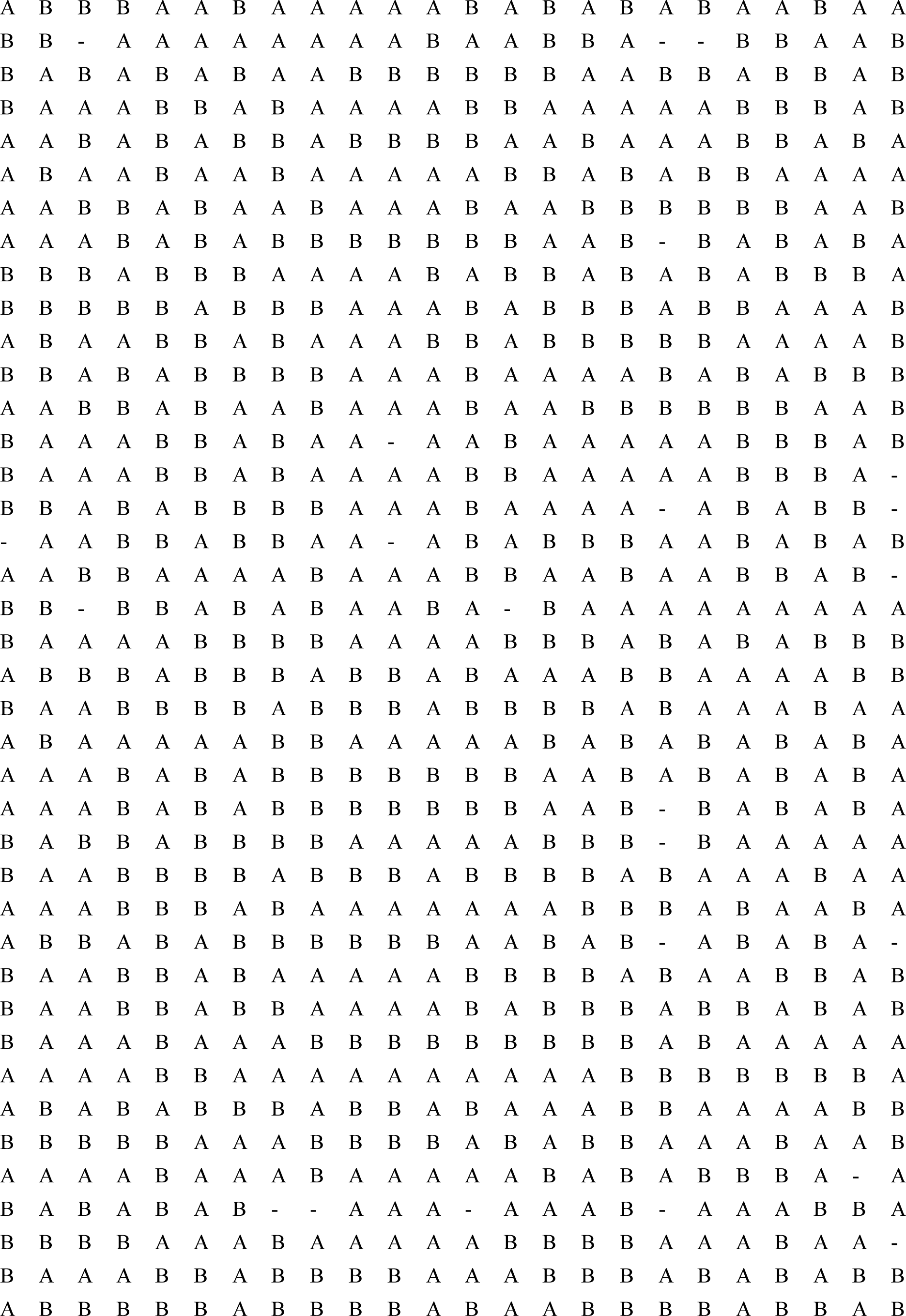
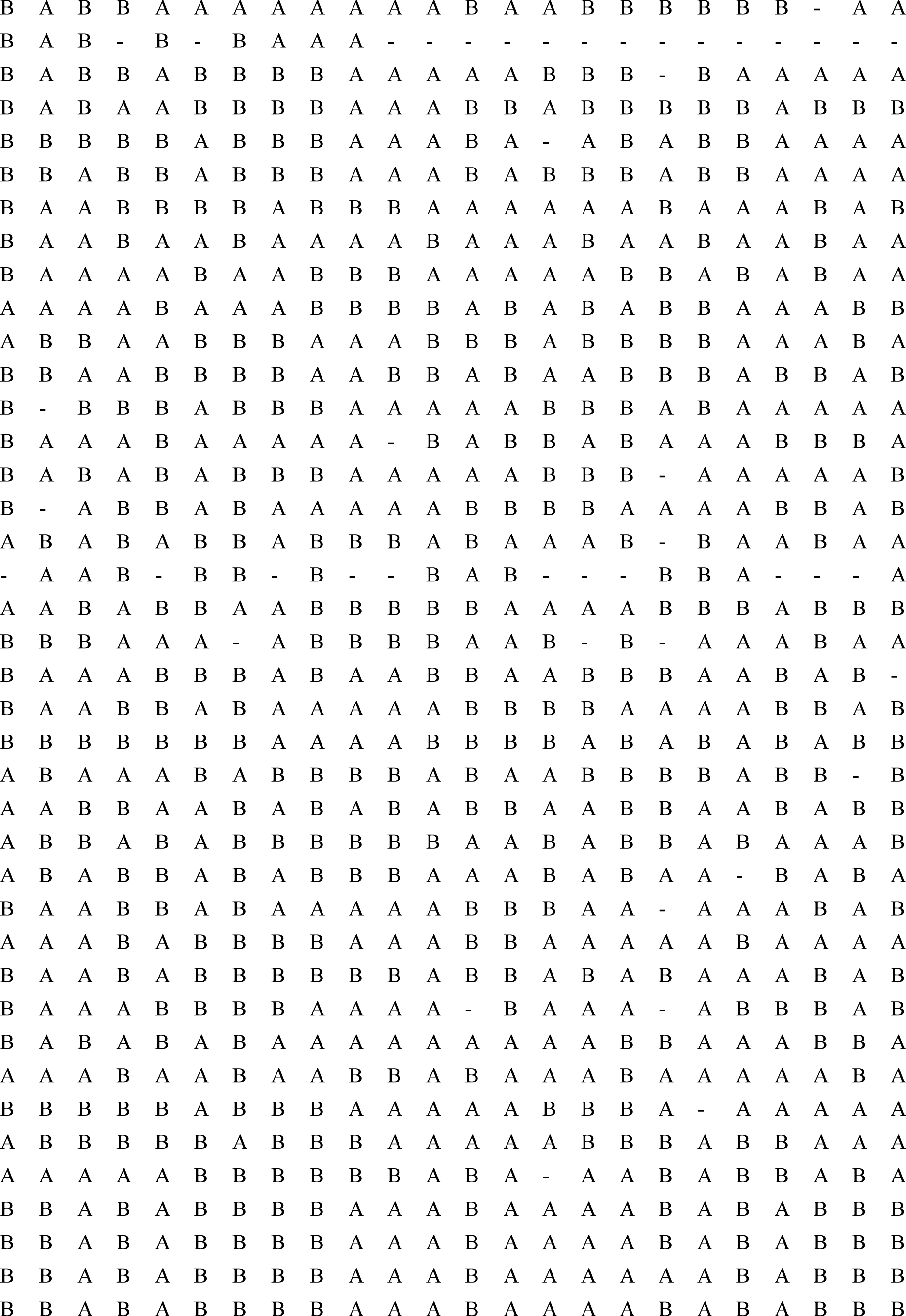
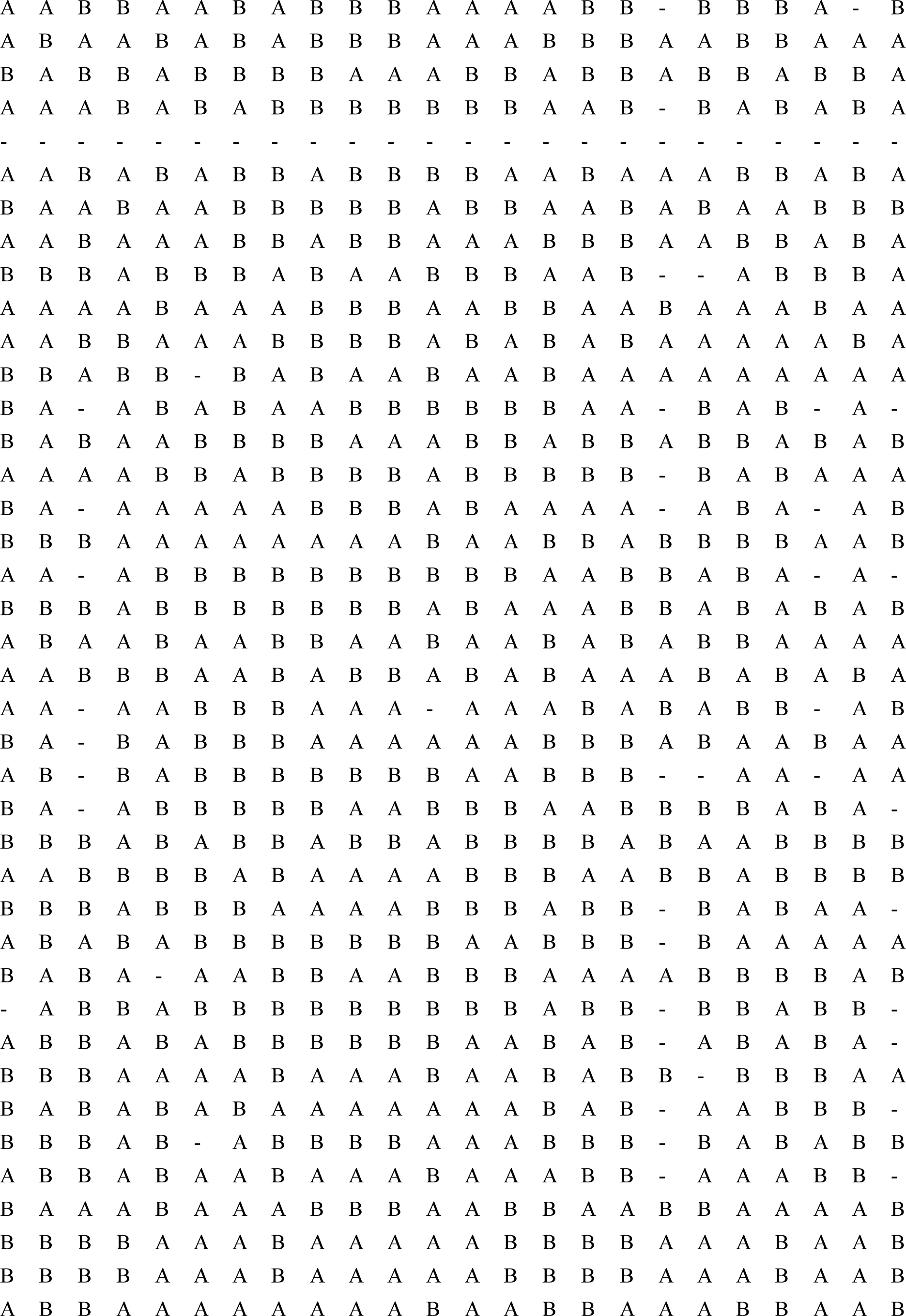
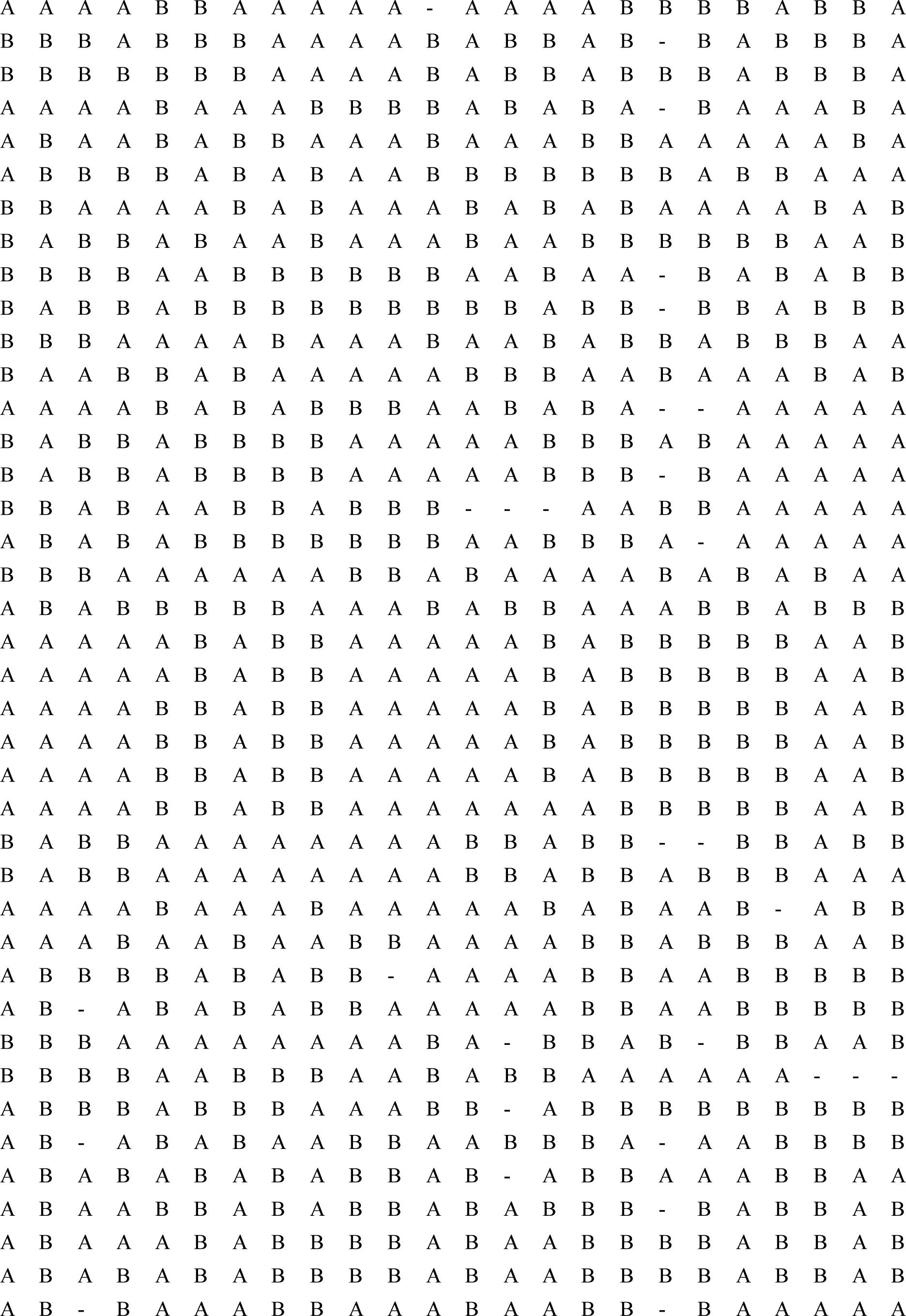
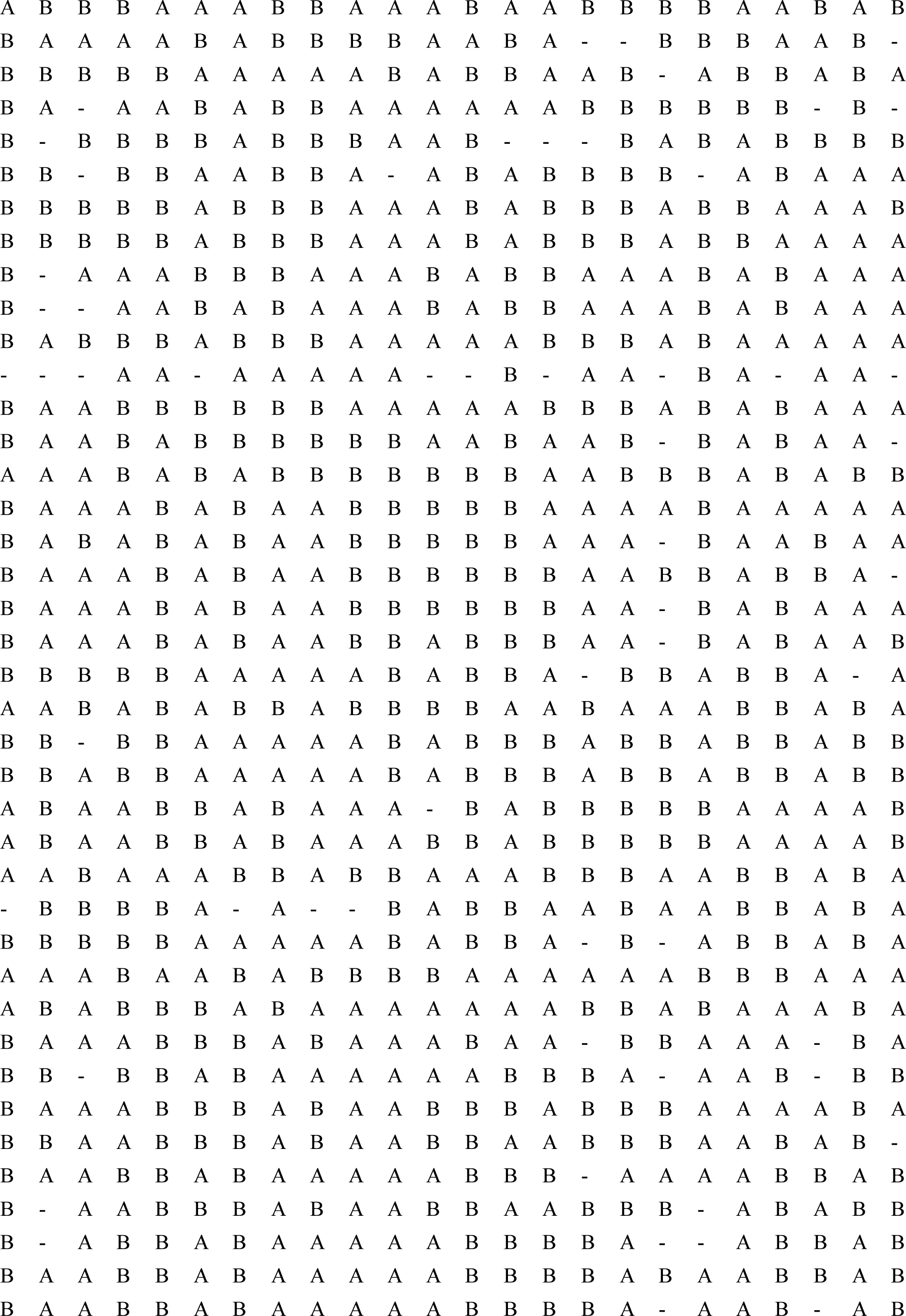
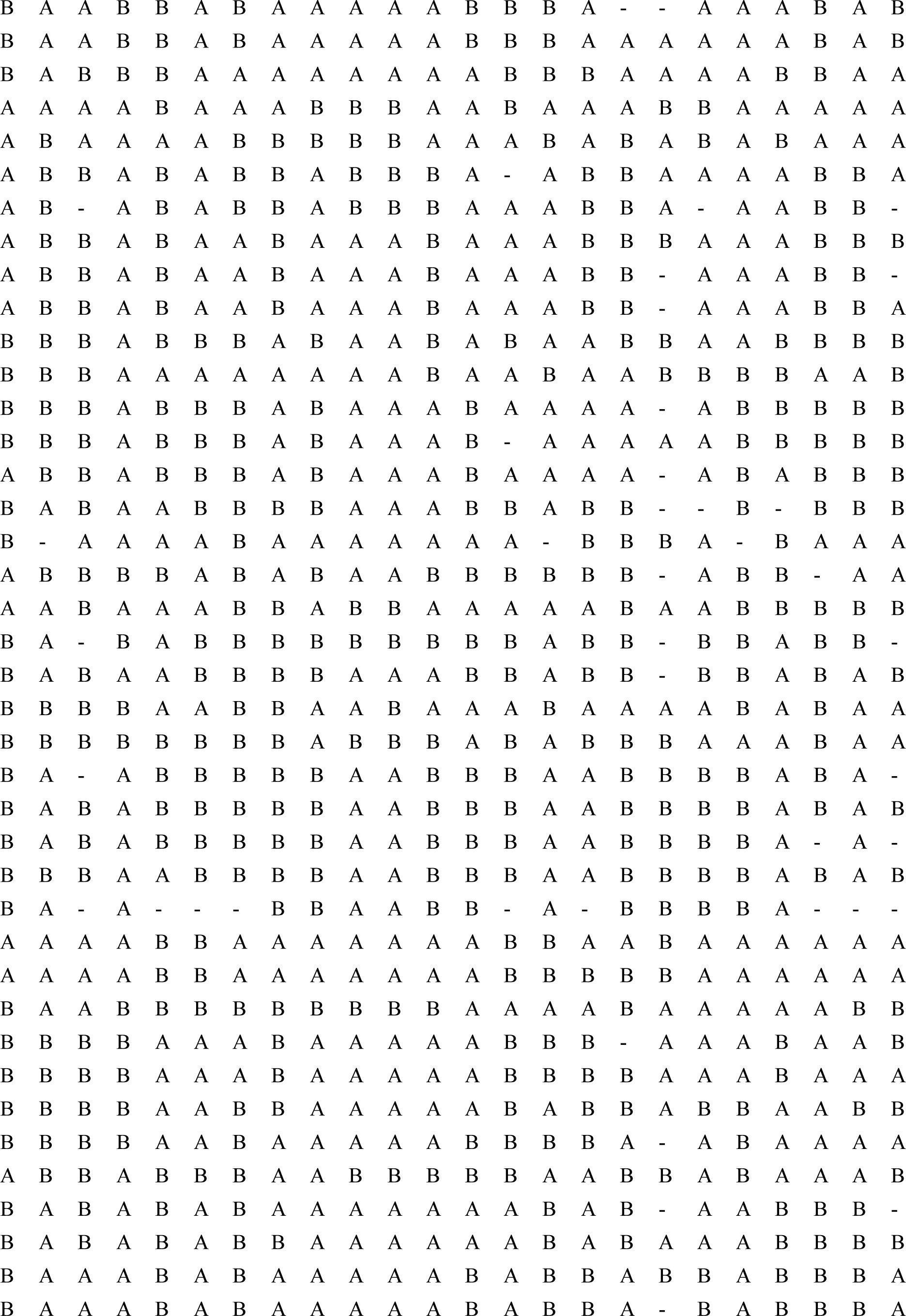
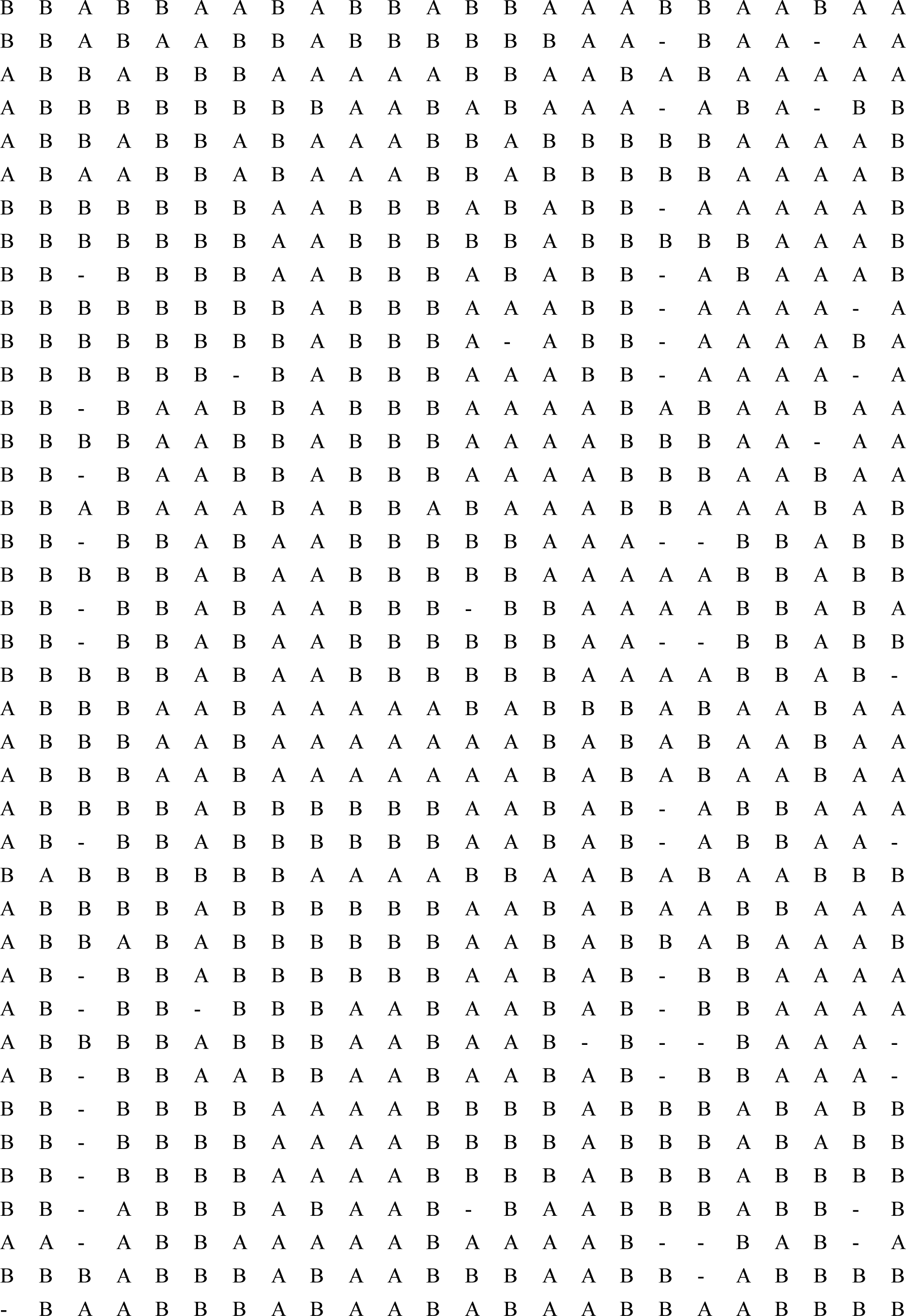
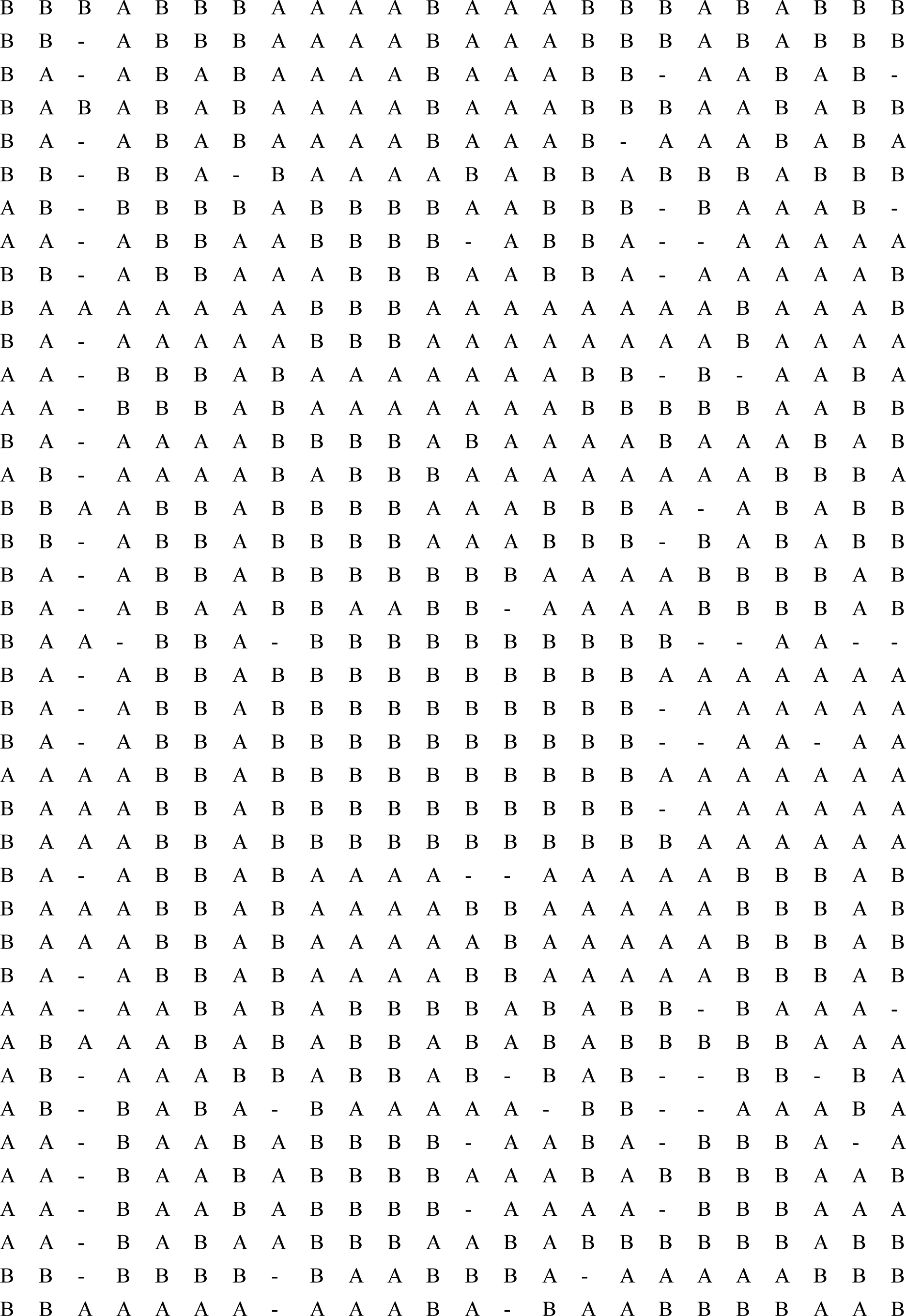
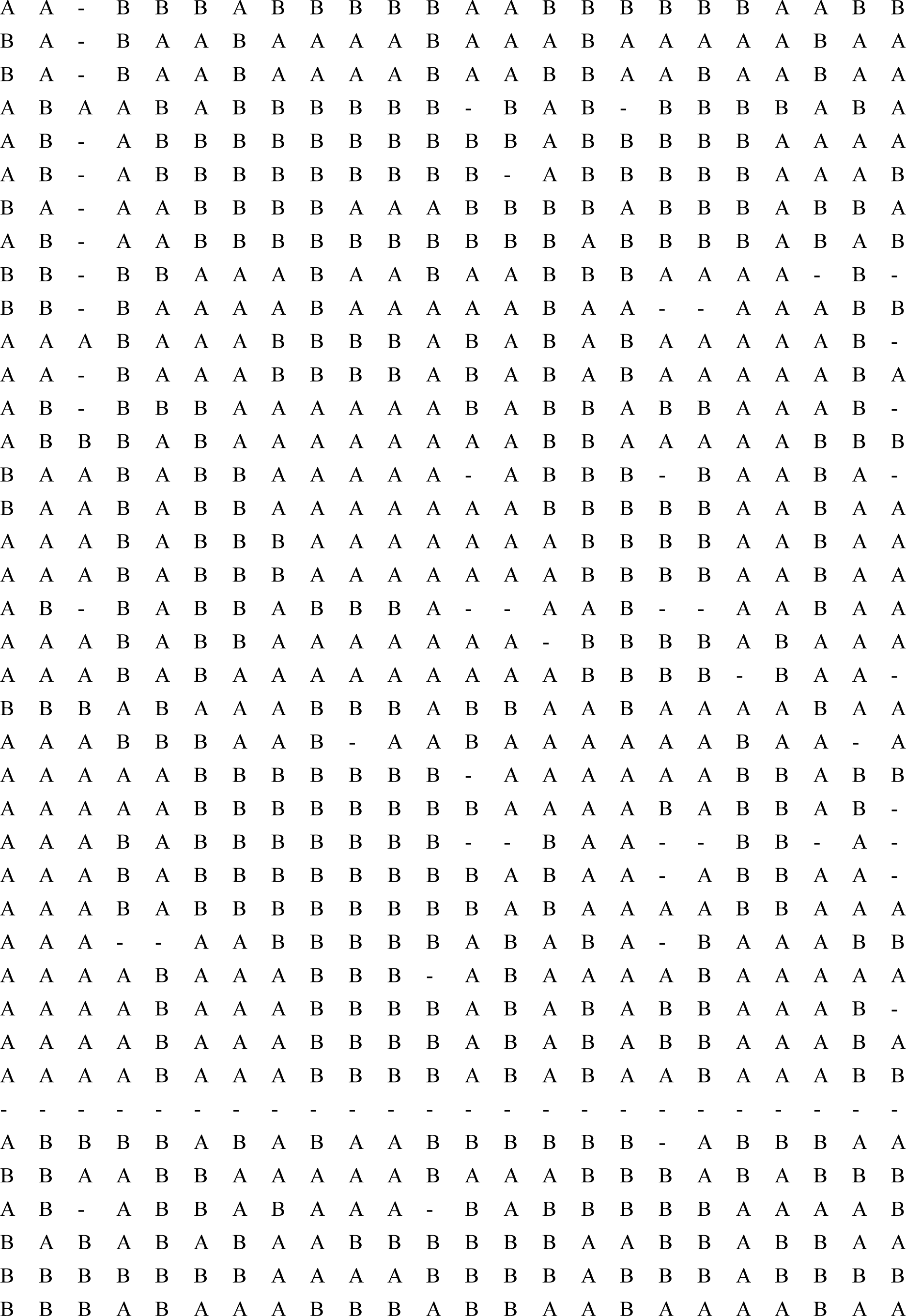
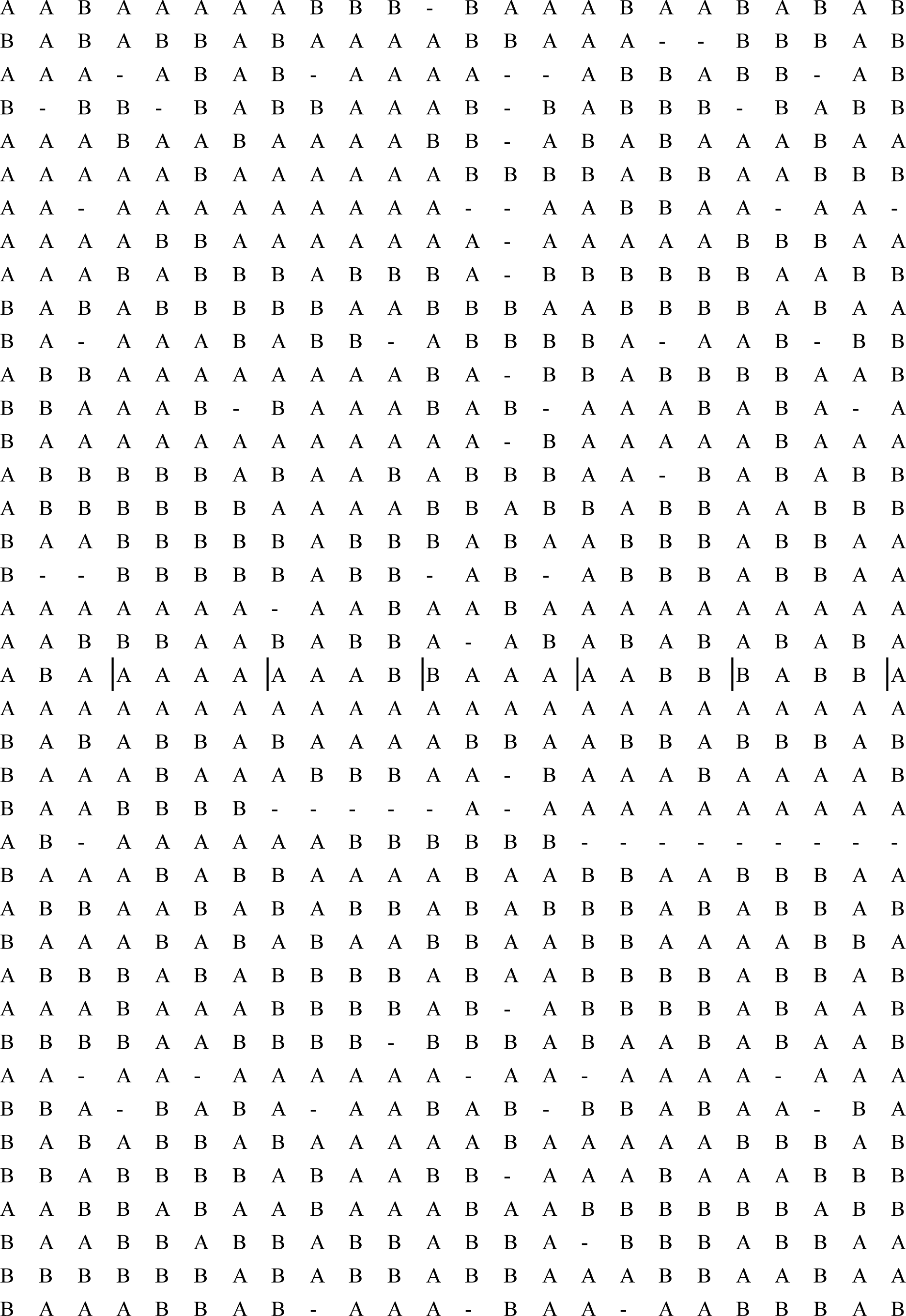
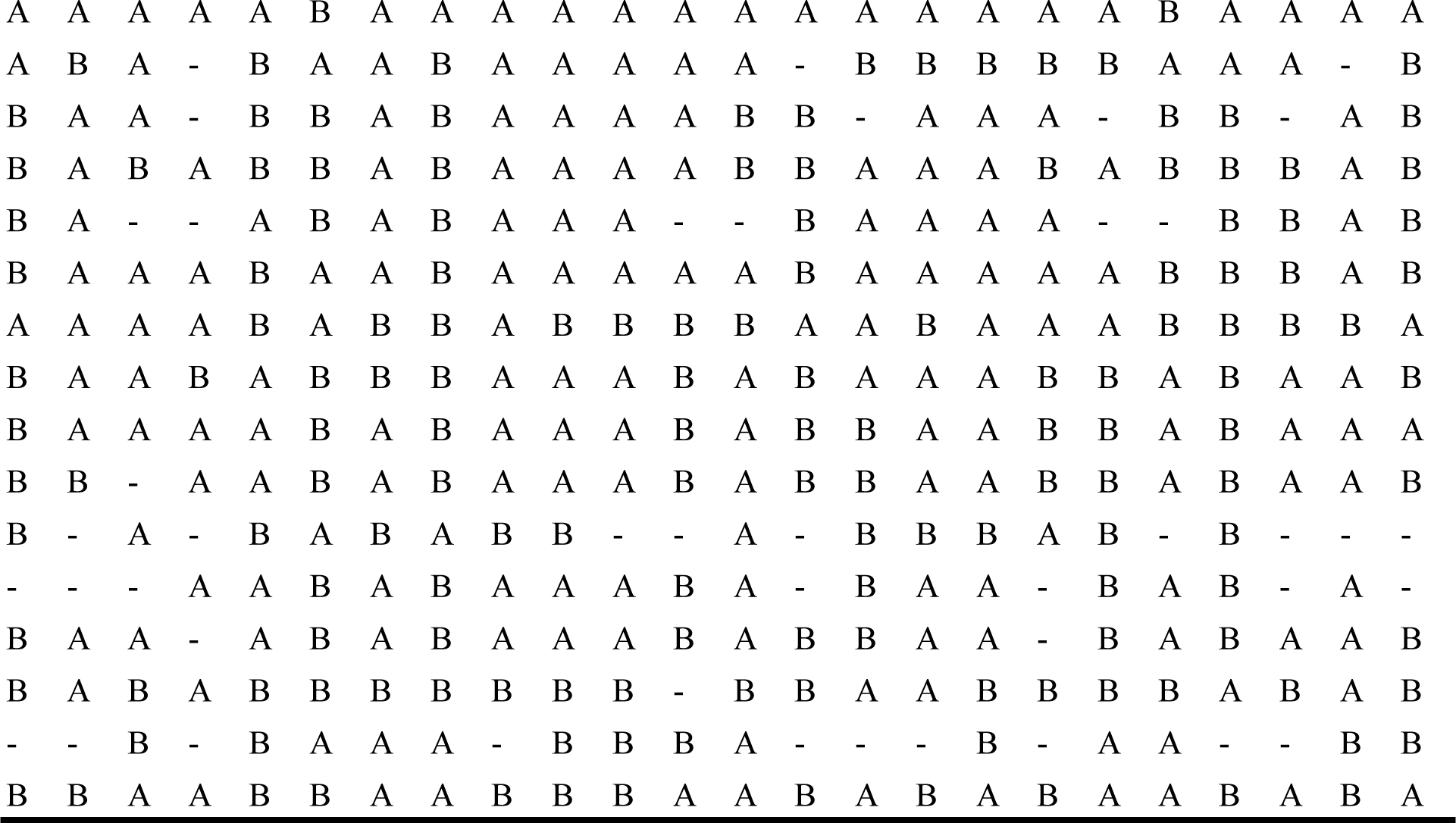
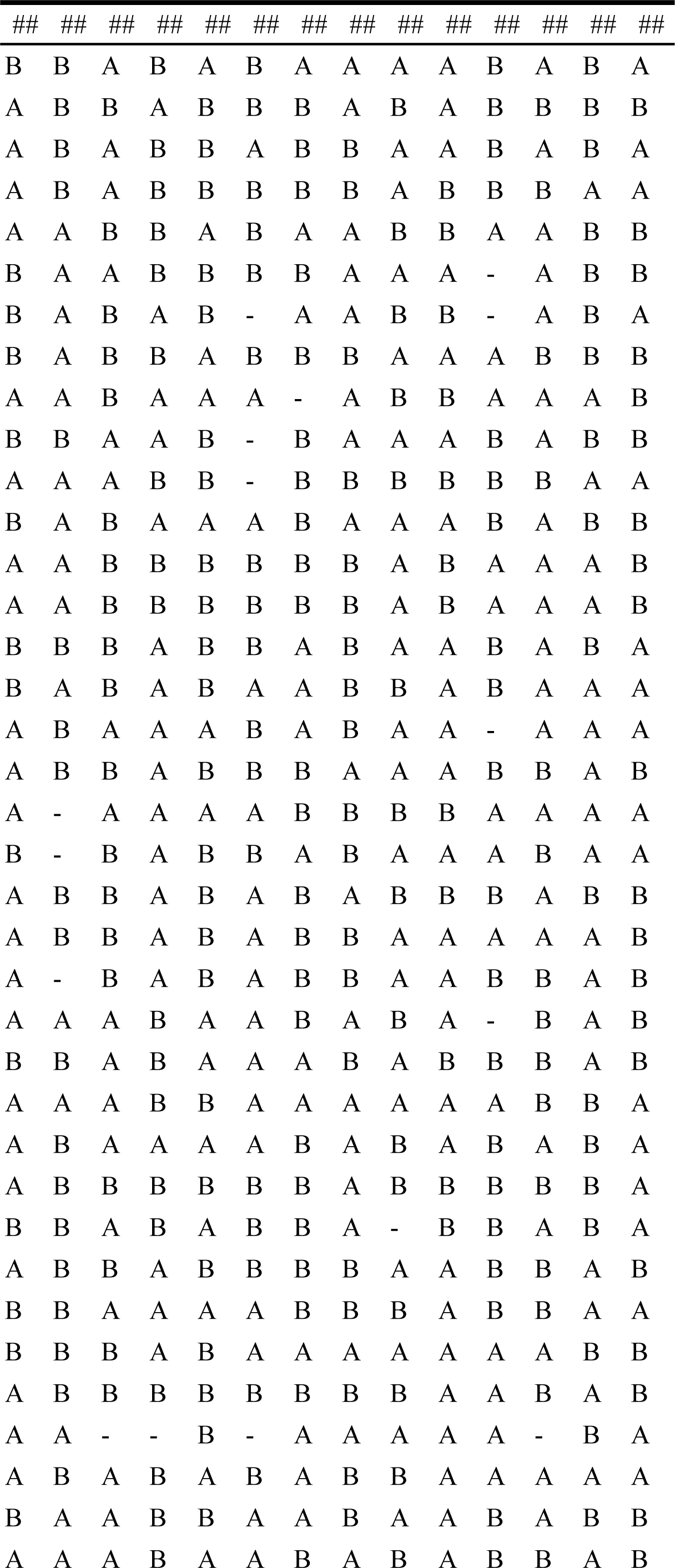
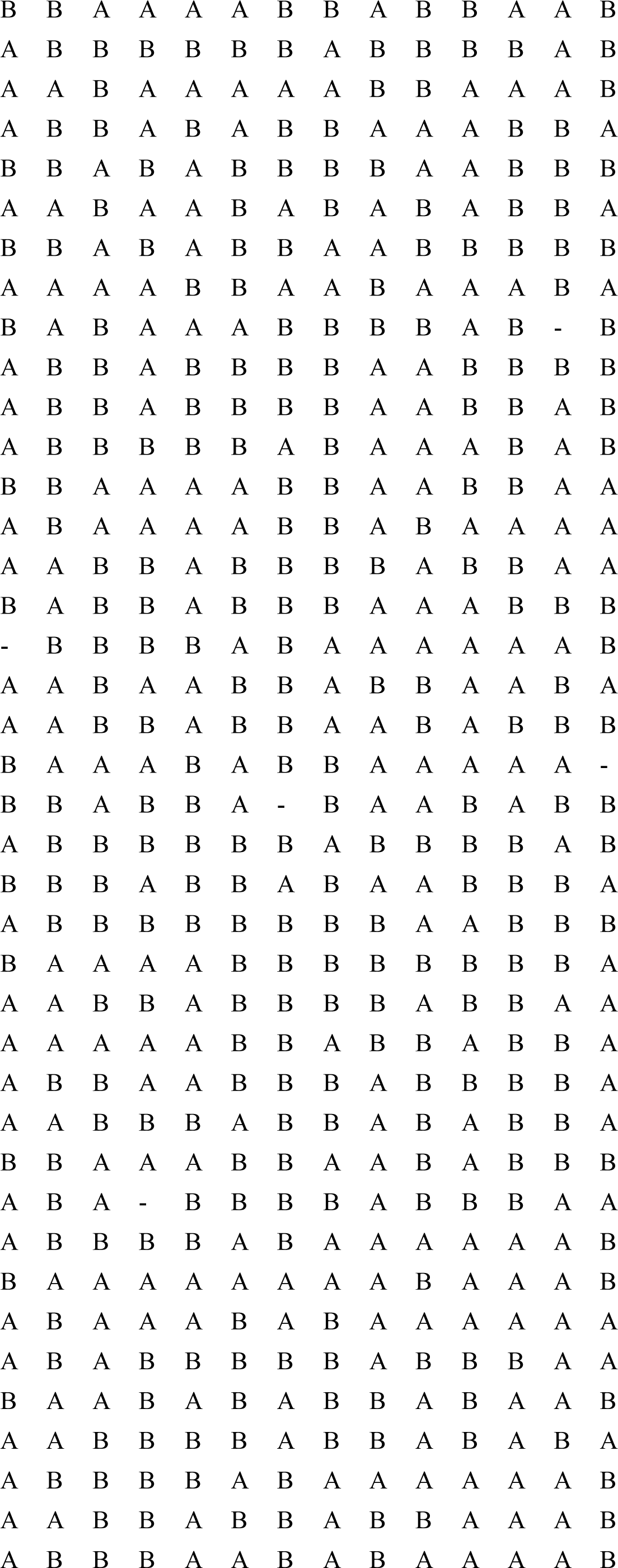
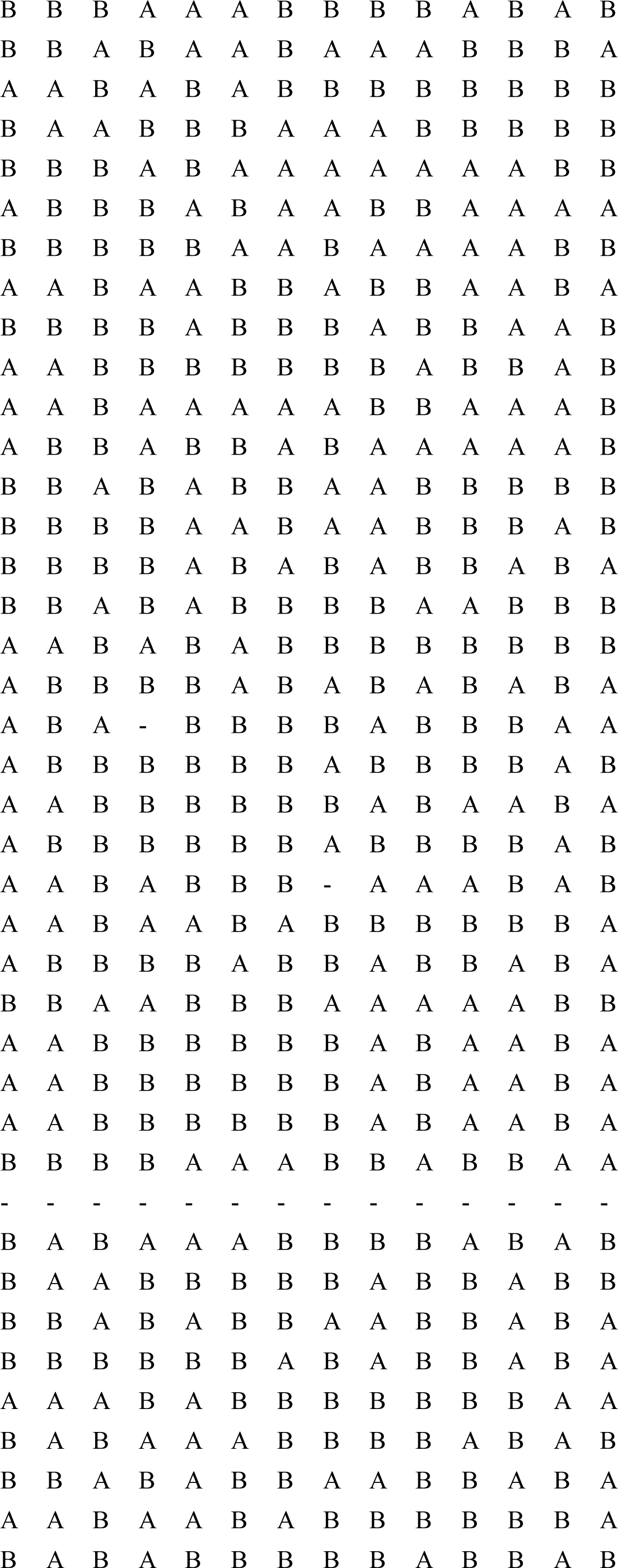
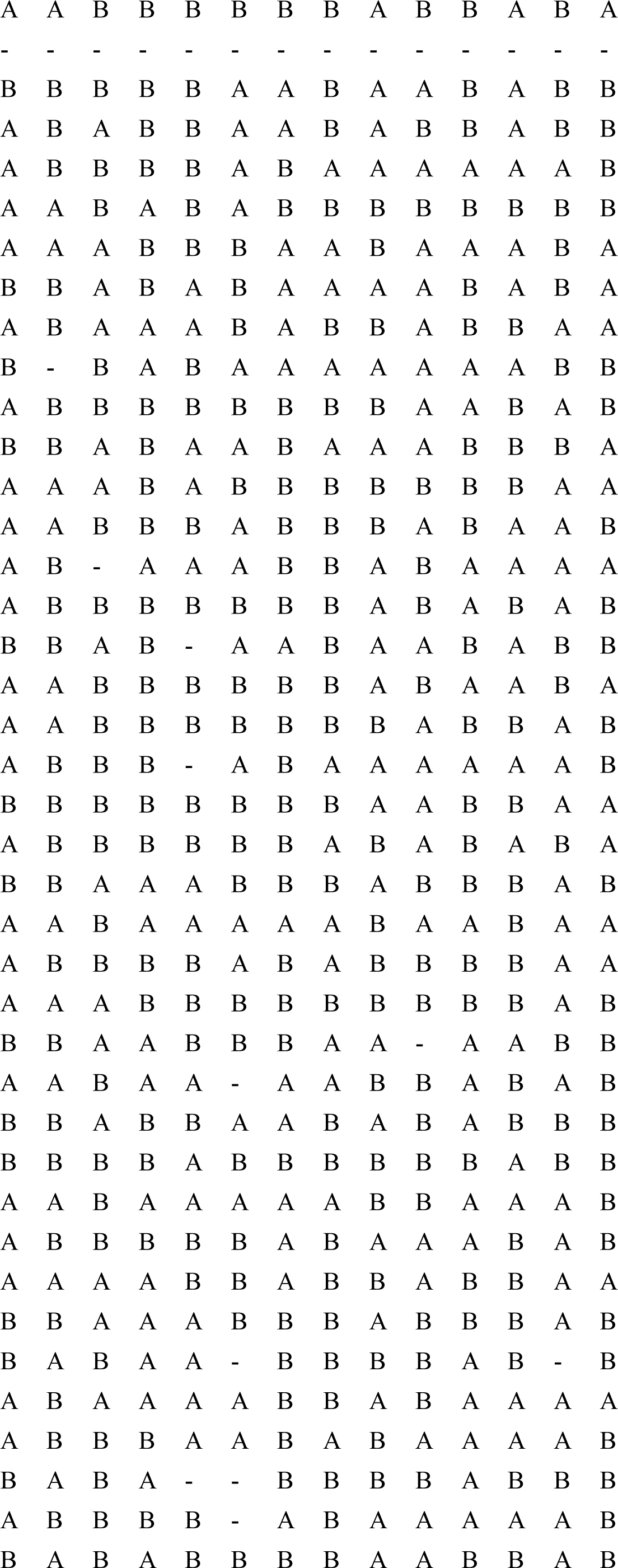
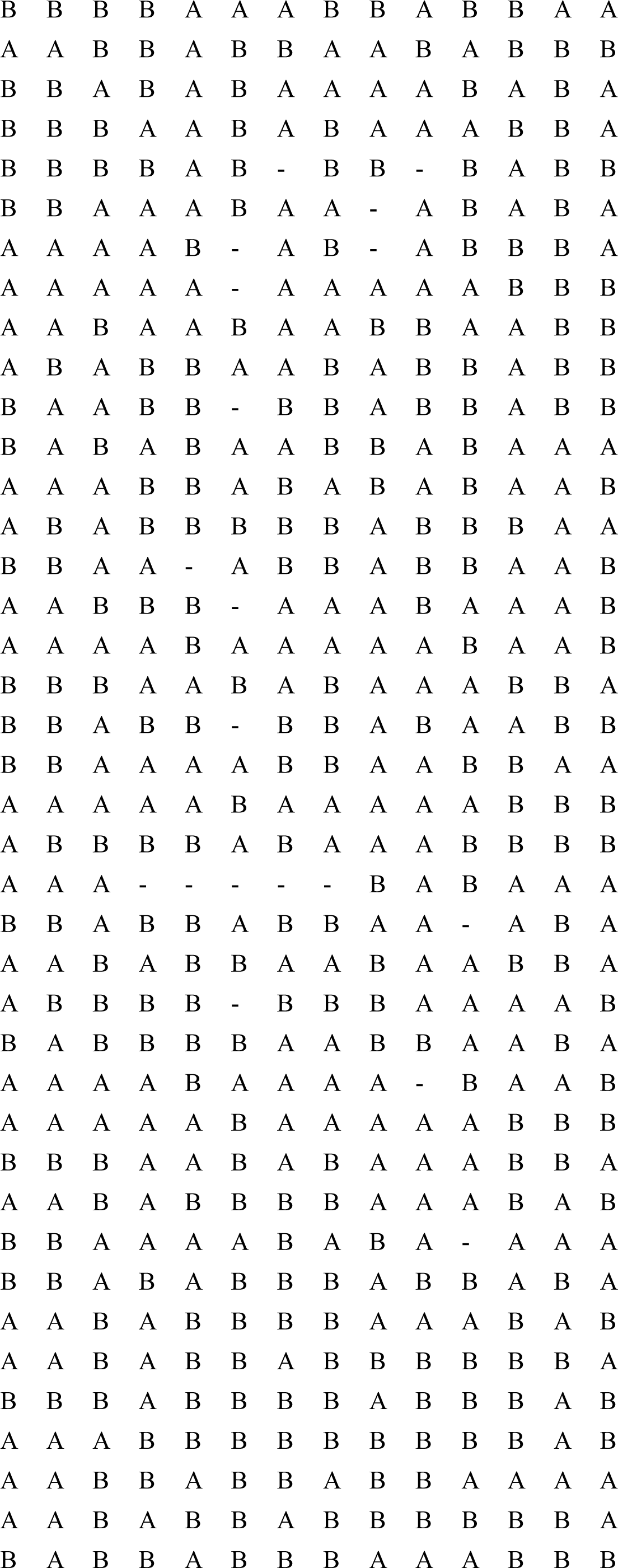
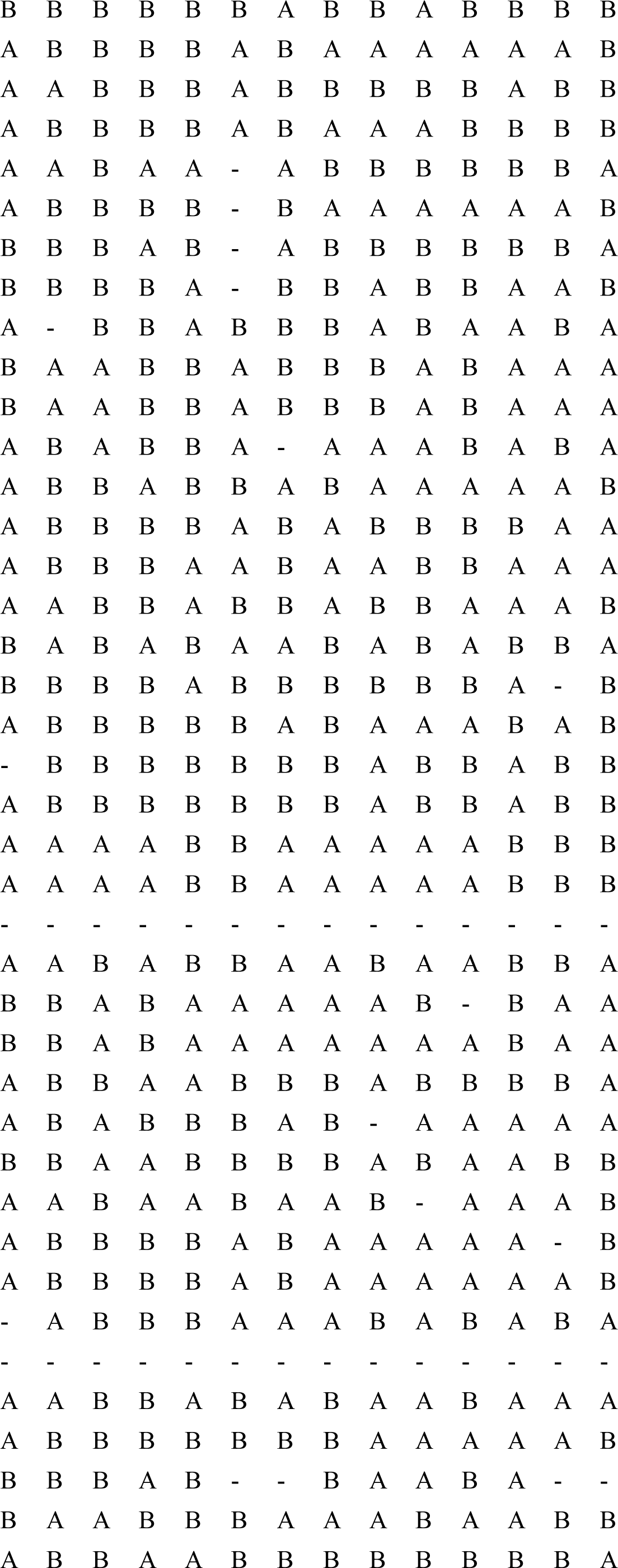
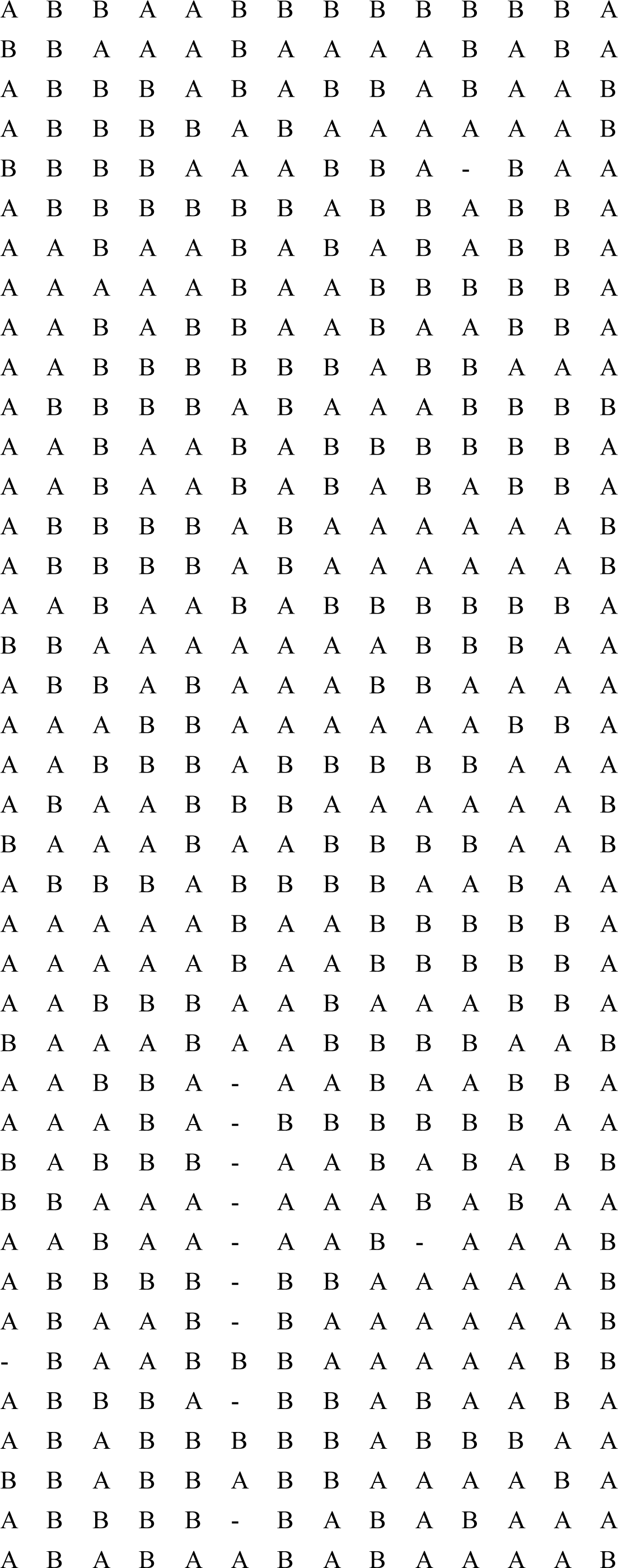
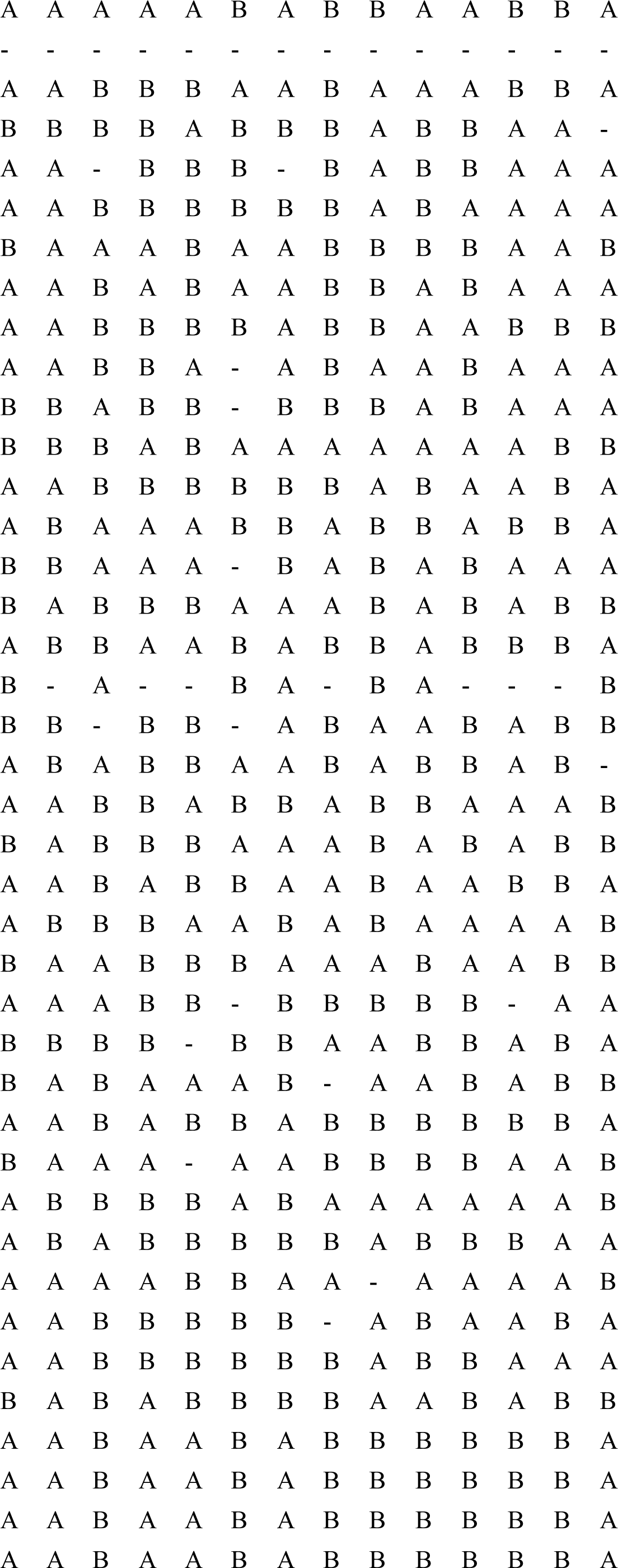
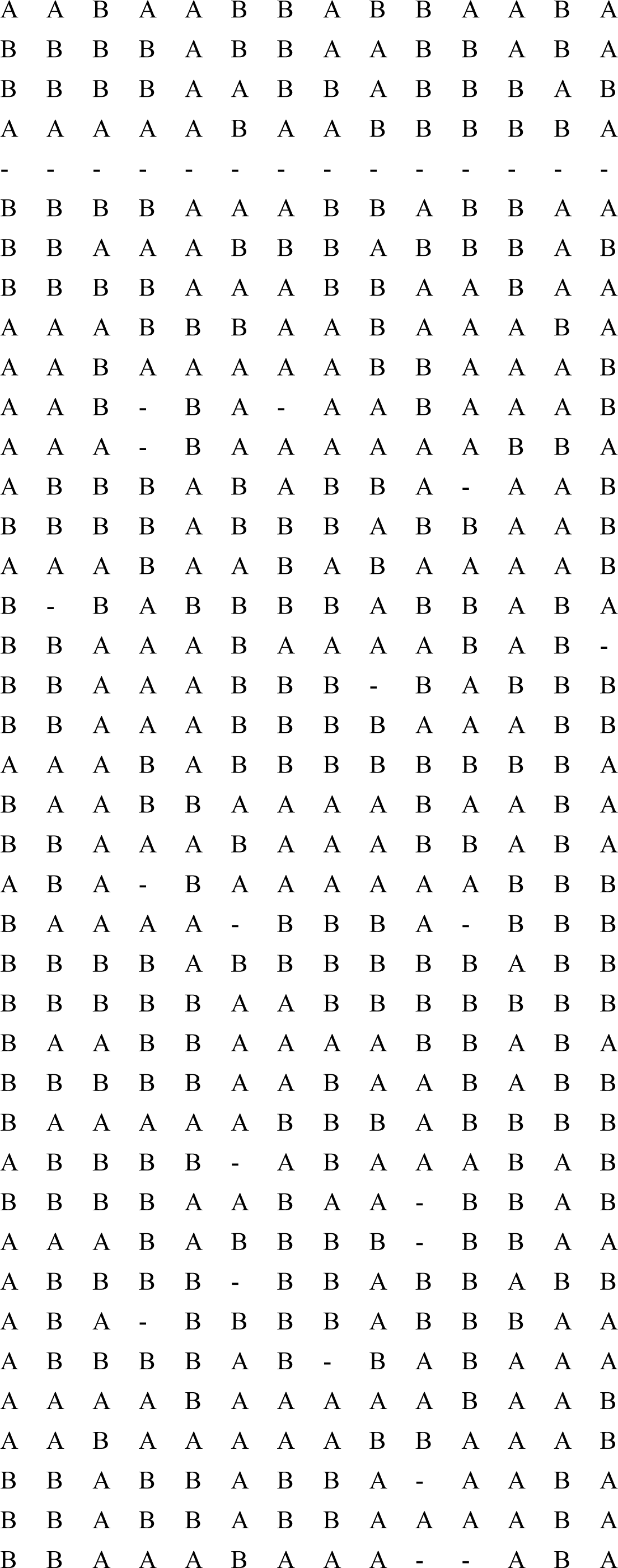
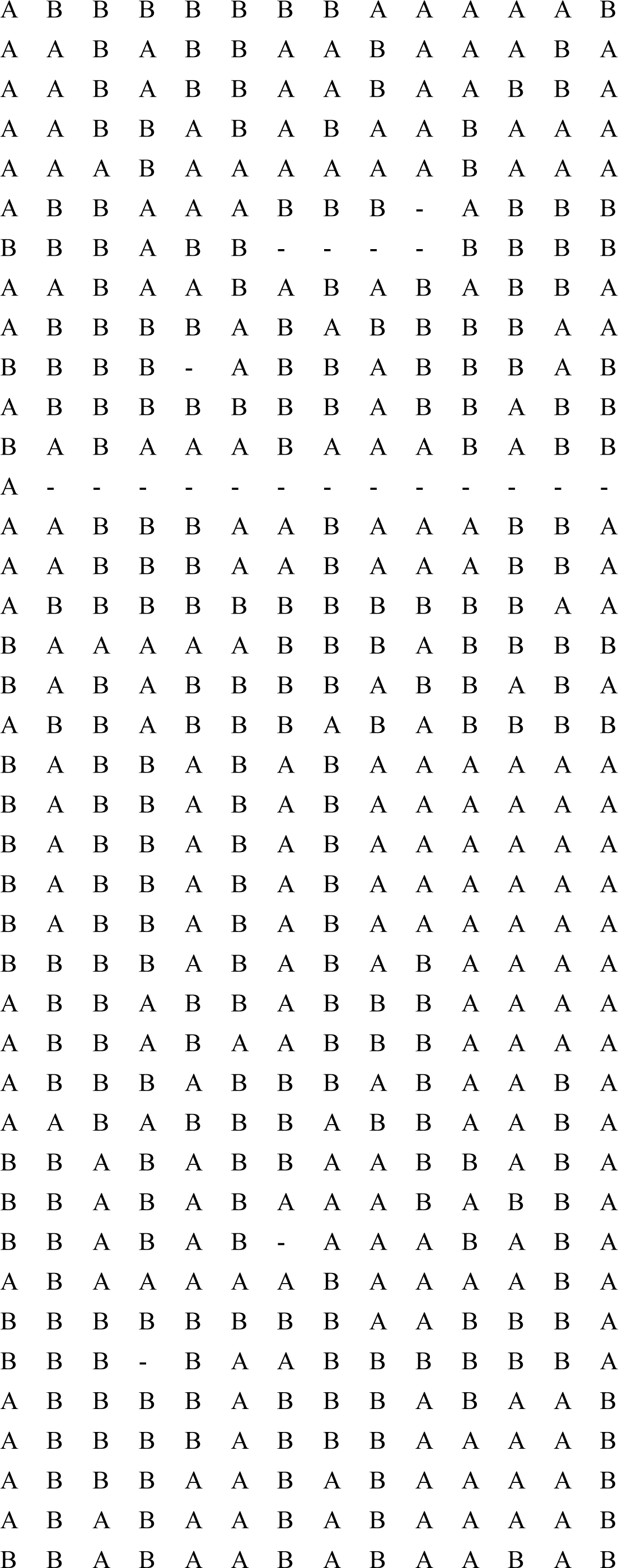
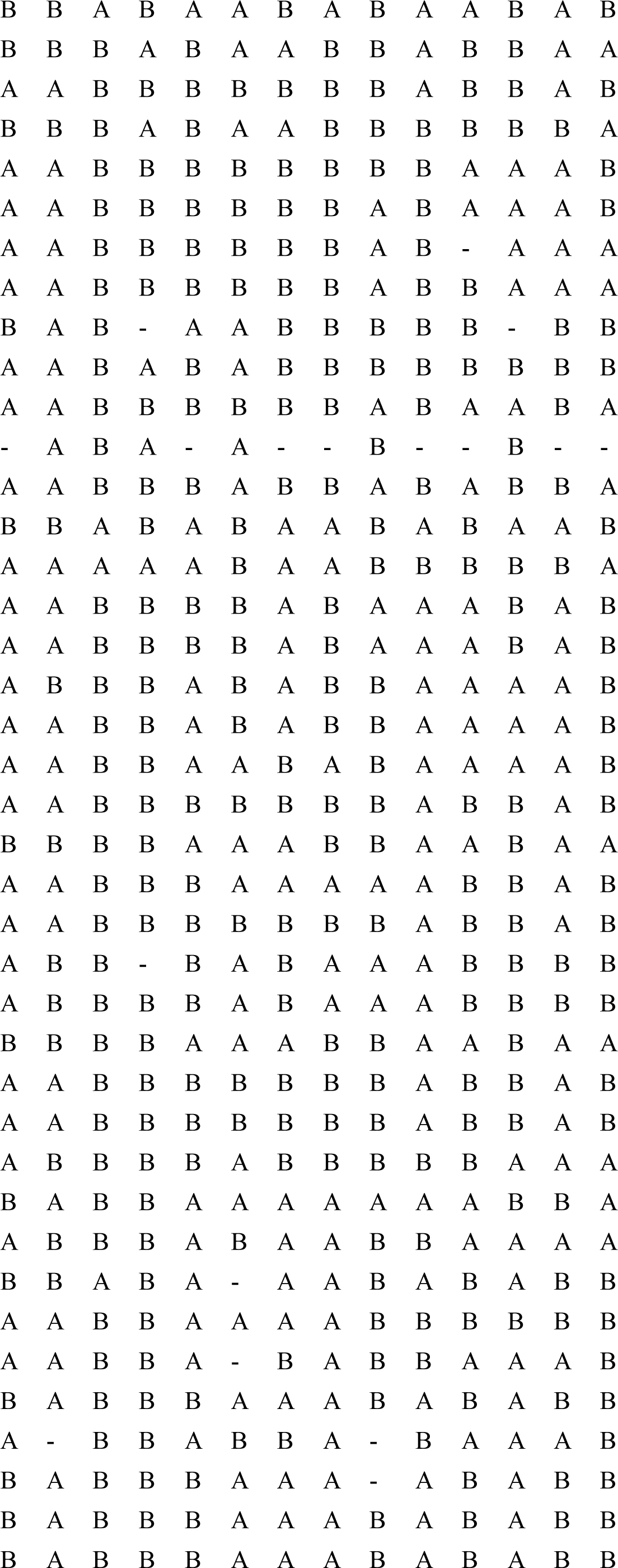
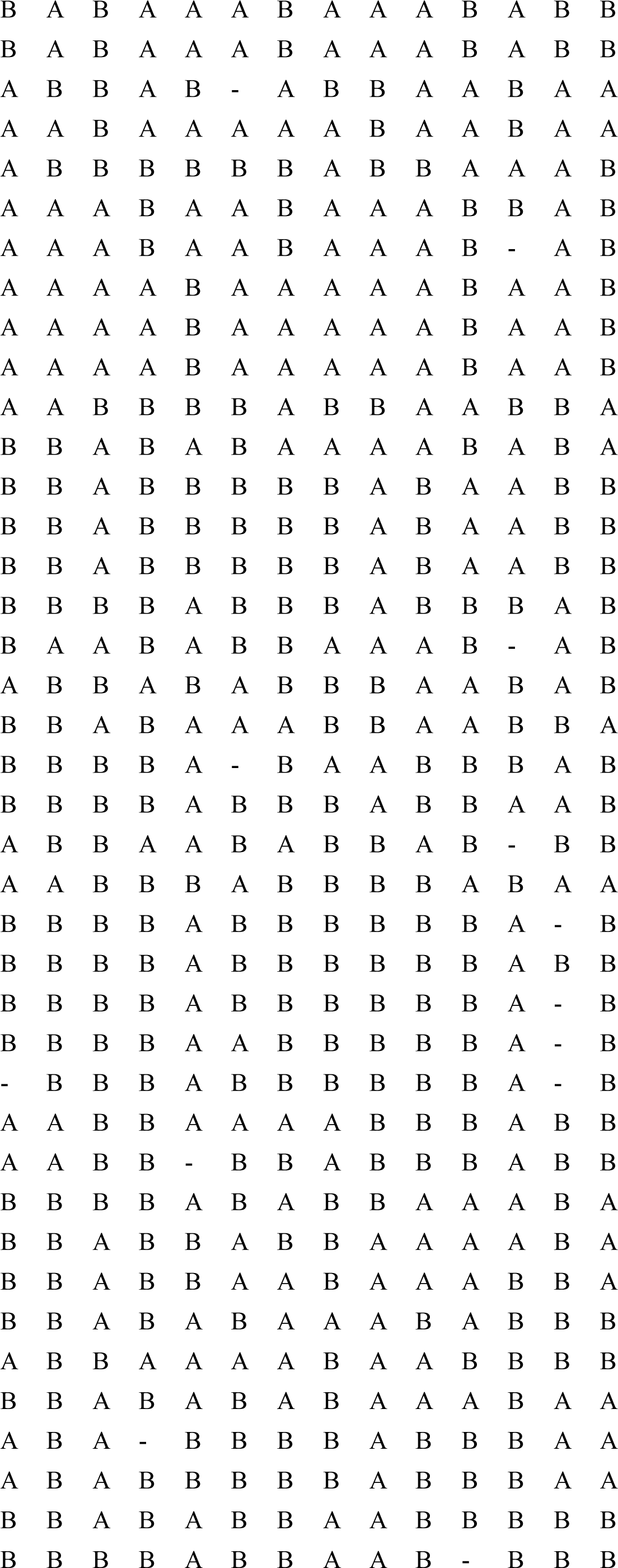
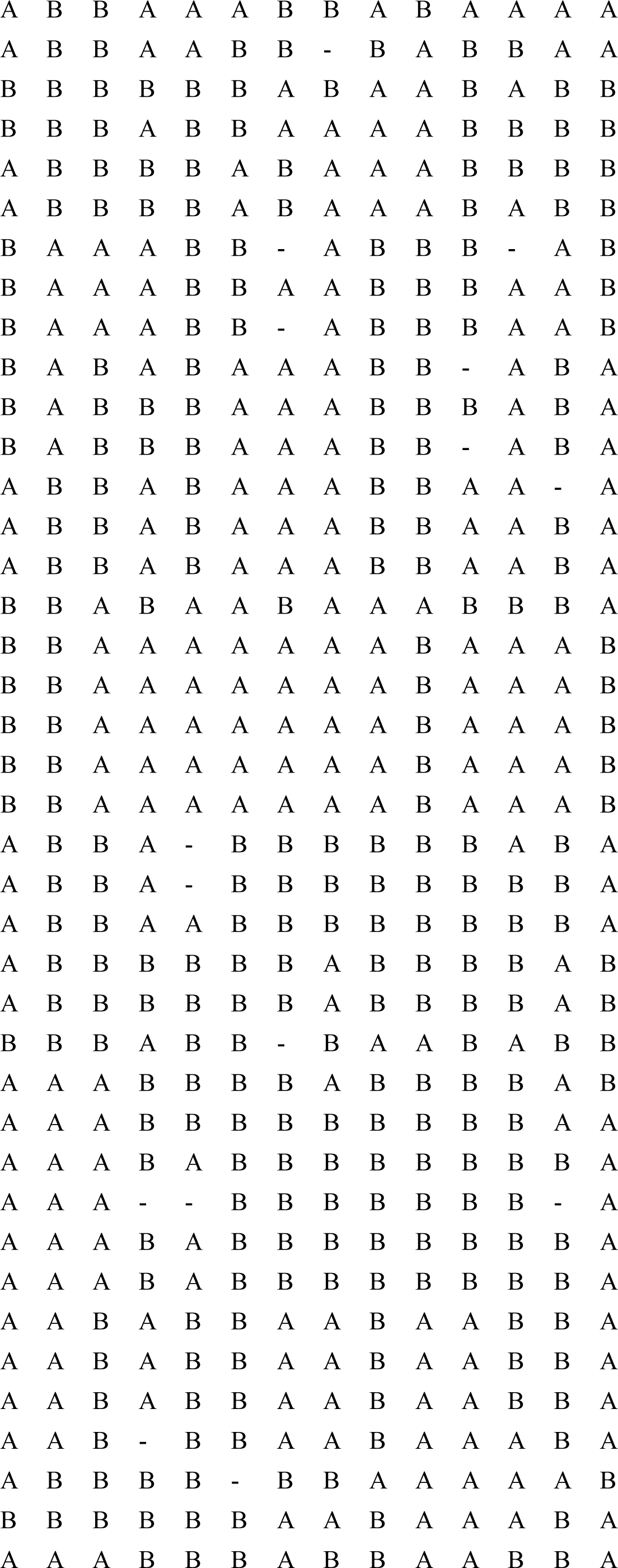
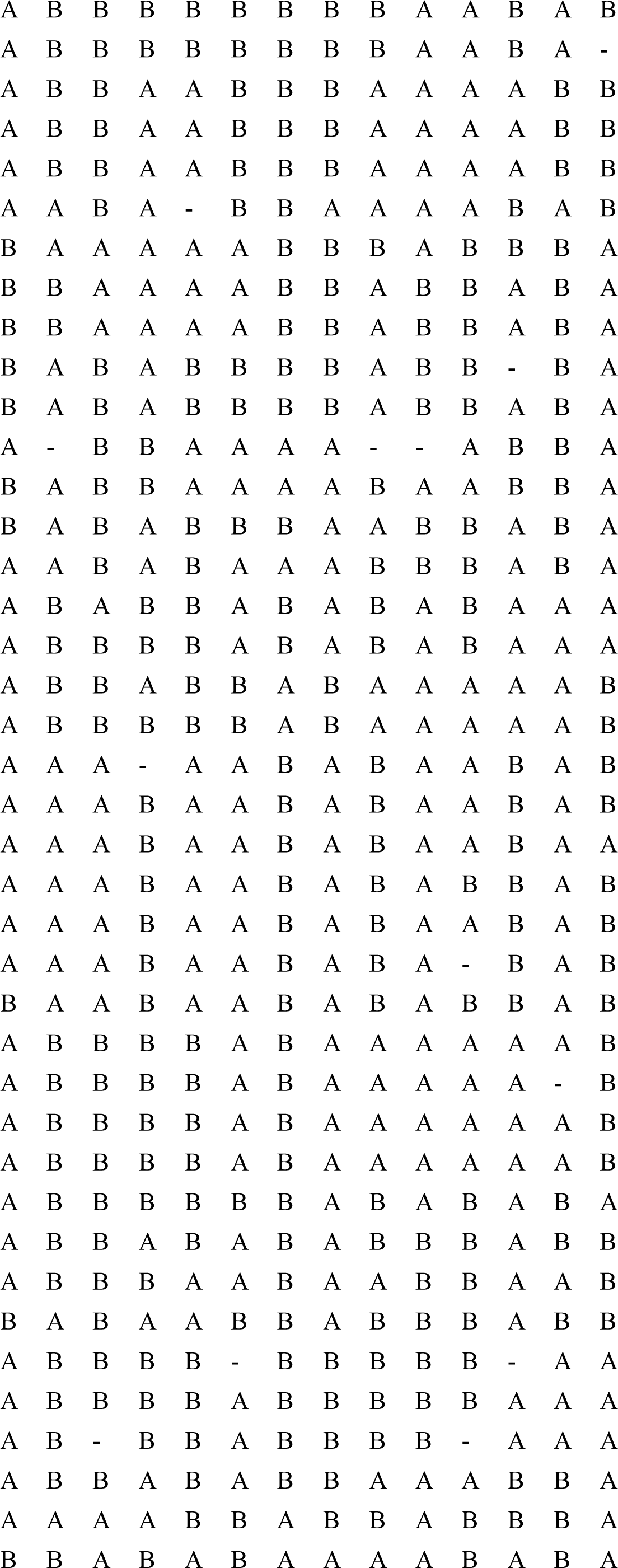
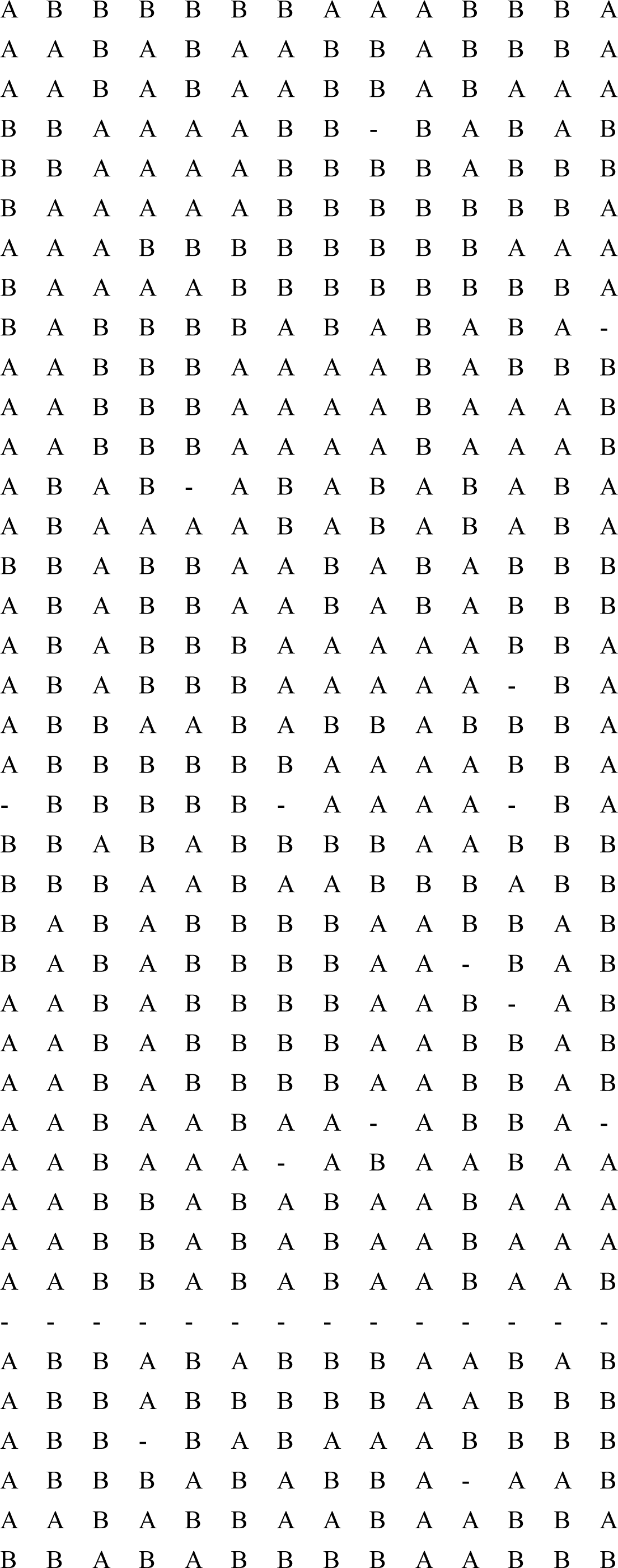
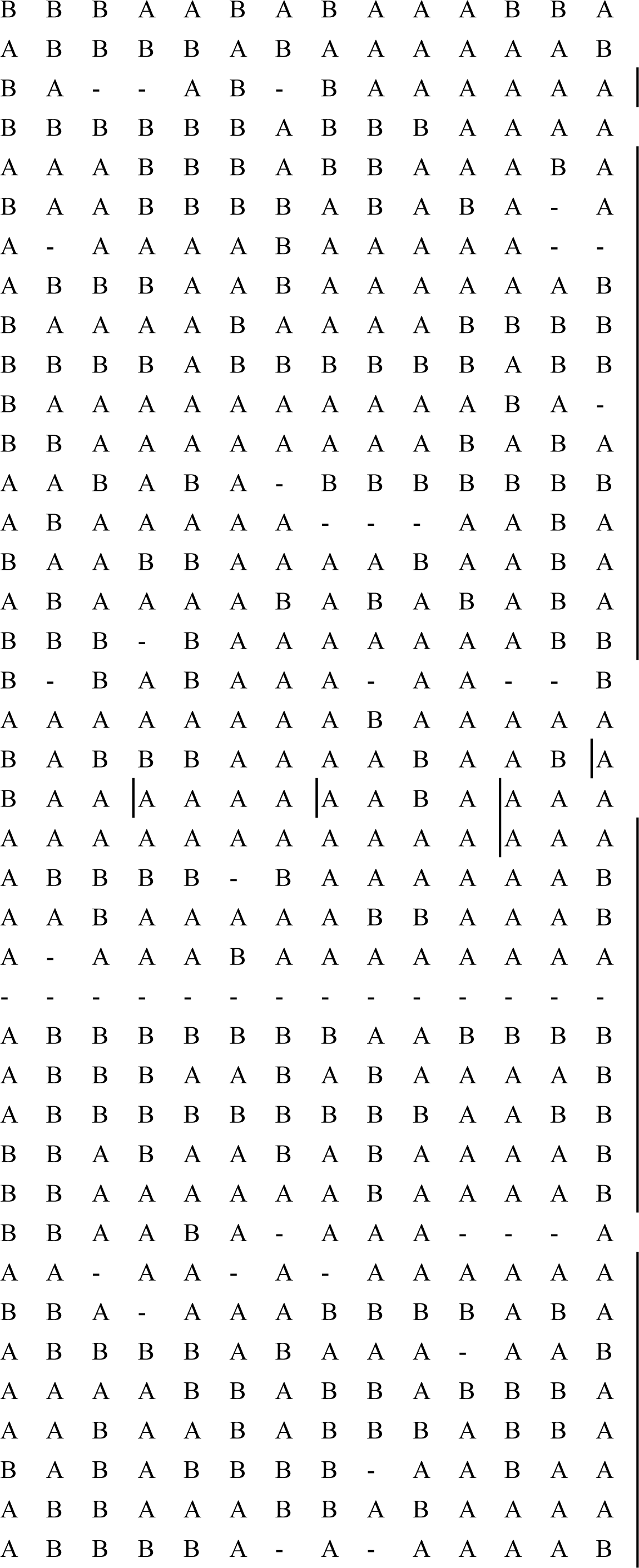
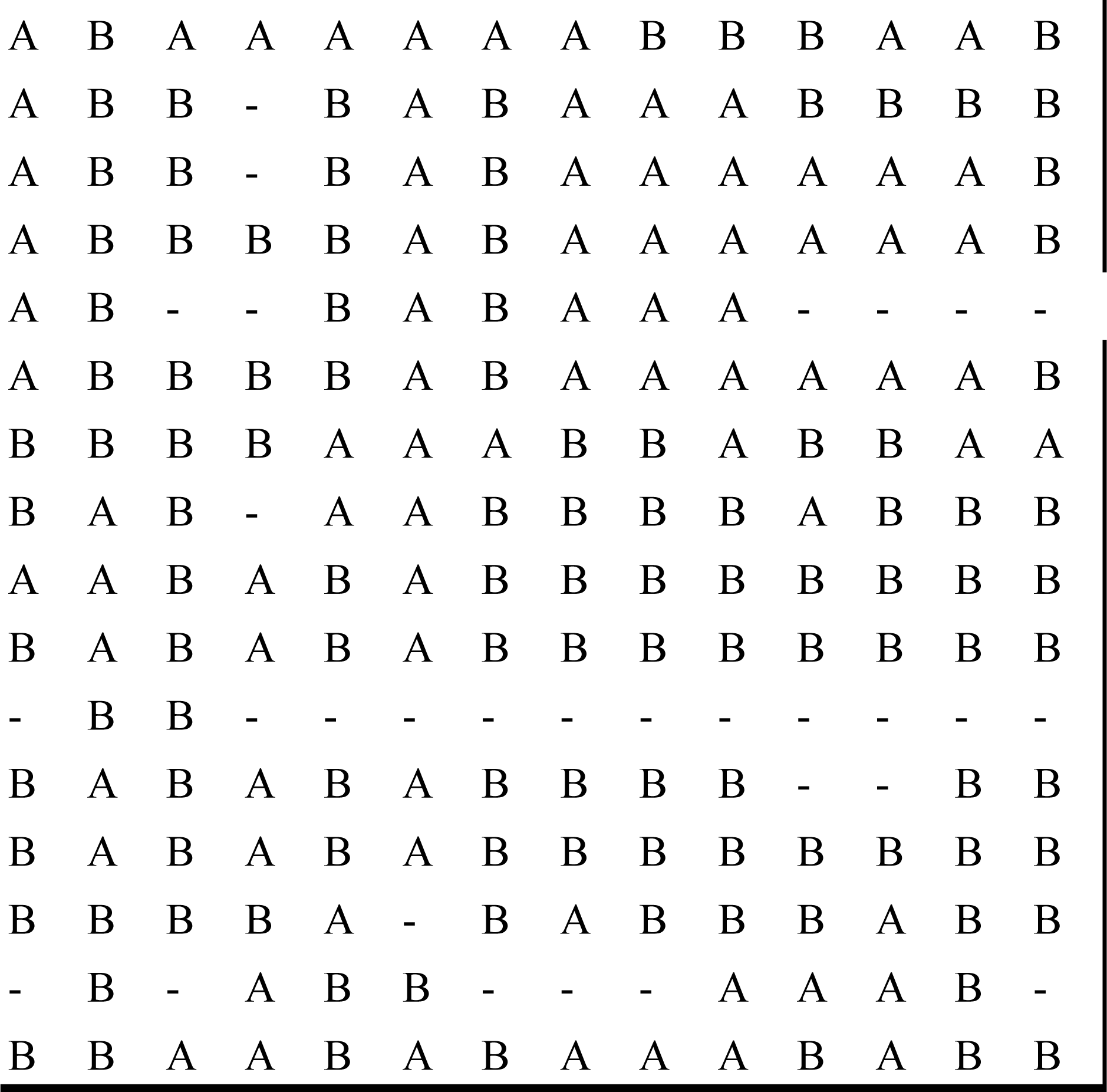
Genotypes of RIL (A), BC/M (B) and BC/P (C) populations. A. Genotypes of RIL population.

